# Gut microbial ecology of Xenopus tadpoles across life stages

**DOI:** 10.1101/2020.05.25.110734

**Authors:** Thibault Scalvenzi, Isabelle Clavereau, Mickaël Bourge, Nicolas Pollet

## Abstract

**Background:** The microorganism world living in amphibians is still largely under-represented and under-studied in the literature. Among anuran amphibians, African clawed frogs of the *Xenopus* genus stand as well-characterized models with an in-depth knowledge of their developmental biological processes including their metamorphosis. In this study, we analyzed the succession of microbial communities and their activities across diverse body habitats of *Xenopus tropicalis* using different approaches including flow cytometry and 16s rDNA gene metabarcoding. We also evaluated the metabolic capacity of the premetamorphic tadpole’s gut microbiome using metagenomic and metatranscriptomic sequencing.

**Results:** We analyzed the bacterial components of the *Xenopus* gut microbiota, the adult gut biogeography, the succession of communities during ontogeny, the impact of the alimentation in shaping the tadpole’s gut bacterial communities and the transmission of skin and fecal bacteria to the eggs. We also identified the most active gut bacteria and their metabolic contribution to tadpole physiology including carbohydrate breakdown, nitrogen recycling, essential amino-acids and vitamin biosynthesis.

**Conclusions:** We present a comprehensive new microbiome dataset of a laboratory amphibian model. Our data provide evidences that studies on the *Xenopus* tadpole model can shed light on the interactions between a vertebrate host and its microbiome. We interpret our findings in light of bile acids being key molecular components regulating the gut microbiome composition during amphibian development and metamorphosis. Further studies into the metabolic interactions between amphibian tadpoles and their microbiota during early development and metamorphosis should provide useful information on the evolution of host-microbiota interactions in vertebrates.

This article has been peer-reviewed and recommended by *Peer Community in Genomics* https://doi.org/10.24072/pci.genomics.100005

## Introduction

Metazoans are vehicles for microbial communities, also named microbiota. The microbiota and its metazoan host have mutualistic interactions and are thought to adapt and evolve as an holobiont (Wilson and Sober, 1989; Gill *et al*., 2006; Zilber-Rosenberg and Rosenberg, 2008; Bordenstein and Theis, 2015). Several studies have highlighted the importance of the microbiota in the function and development of numerous organs such as the alimentary canal, the nervous system and the tegument (Sekirov *et al*., 2010; Sommer and Bäckhed, 2013; Douglas, 2018). The dynamic interaction between a microbiota and its host is under intense scrutiny especially for mammalian species and a handful of model organisms (Colston and Jackson, 2016; Douglas, 2019). However, very little is currently known on the biotic and abiotic interactions between the microbiota of even well-known and classic vertebrate model organisms such as amphibians (Colston and Jackson, 2016; Douglas, 2019; Rebollar and Harris, 2019).

The current paucity of knowledge on amphibian’s microbiome is an historical thumb of nose. Indeed in 1901, Olga Metchnikoff published a note on the influence of microbes on the development of tadpoles (Metchnikoff, 1901). She concluded that microbes were vital to tadpoles because they could not complete their development in sterile conditions, and her results were confirmed by Moro (Moro, 1905). A few years later, these results were challenged by Eugène and Elisabeth Wollmann, who finally observed that *Rana temporaria* tadpoles could complete their metamorphosis in sterile conditions (Wollman, 1913). They also reported that tadpoles could be fed solely with bacteria, pinpointing their nutritive roles (Wollman and Wollman, 1915). Since that time, only few investigations have addressed the interactions between bacteria and tadpoles during development.

Currently, the best-known amphibian model in biology is the African clawed frog. Several features of its life cycle can facilitate investigations on the interactions between an animal host and communities or individual microbial taxa. *Xenopus* have a long lifespan of more than ten years and several tools are available to study their biology including genomic engineering (Vouillot *et al*., 2014). However, there are a few data on *Xenopus* microbiota. Mashoof et al. used 16S rRNA metabarcoding to describe the bacterial diversity in the gut microbiota of adult *Xenopus* and found that *Firmicutes, Bacteroidetes* and *Proteobacteria* were the dominant bacterial phyla like in other vertebrates (Mashoof *et al*., 2013; Colombo *et al*., 2015).

Like many amphibians, *Xenopus* life history is characterized by an aquatic external development. The embryo hatches after one or two days of embryonic development under the protection of the chorion and of a double-layer jelly coat (Bles, E.J., 1906; Faber and Nieuwkoop, 1994). The newly hatched embryo is in direct contact with its environment and usually sticks to the remnants of its jelly coat thanks to secretions from its adhesive gland. The mouth opens one day later, and after about two more days, the small tadpole starts feeding by water filtration. Further growth and development continues during four weeks, a period called premetamorphosis. During the next phase called prometamorphosis, limb growth is significant. Finally, the end of morphogenesis is marked by tail resorption during the metamorphic climax and the adult-shaped froglet is completely formed (Faber and Nieuwkoop, 1994; Brown and Cai, 2007). Most organs, including the gut compartments, are found in both tadpoles and adults but they experience a complete remodeling that enables the continuation of their physiological functions in a different ecological niche (Hourdry *et al*., 1996; Heimeier *et al*., 2010).

Immunologically, post-embryonic development is hallmarked by significant changes especially regarding mucosal immunity (Robert and Ohta, 2009; Pasquier, 2014; Colombo *et al*., 2015). In particular, the climax of metamorphosis is hallmarked by high-levels of glucocorticoids that cause a systemic immuno-depression (Rollins-Smith *et al*., 1997; Sachs and Buchholz, 2019). Thus, amphibian metamorphosis is a post-embryonic developmental step during which host-microbial interactions can play critical roles.

Amphibian tadpoles are almost invariably water-dwelling organisms, and most are microphagous with microbial fermentation contributing up to 20 % of the energy uptake in bullfrog (Clayton, 2005; Pryor and Bjorndal, 2005; Altig *et al*., 2007). After metamorphosis, froglets are still microphagous but they start preying on larger animals as they grow, with a diet composed predominantly of chitin-rich insects and other arthropods. The precise timing of organ and behavioral modifications depends on ecological traits and is known in only a handful of species, including *Xenopus*. As a *Pipidae, Xenopus* tadpoles, metamorphs, froglets and adult frogs live predominantly in water (Bles, E.J., 1906).

Changes in gut bacterial communities during anuran metamorphosis was first studied using culturing methods and more recently using 16S rDNA gene metabarcoding (Fedewa, 2006; Kohl *et al*., 2013; Vences *et al*., 2016; Chai *et al*., 2018; Warne *et al*., 2019; Long *et al*., 2020; Zhang *et al*., 2020). These very interesting studies reported changes in the composition of microbial communities upon metamorphosis, with differences between species. A single study expanded its analysis using metagenomic and metatranscriptomic (Zhang *et al*., 2020). Some limitations of these studies are that they worked on natural populations of non-model organisms without genomic resources. In addition, they typically sampled a single developmental stage and they relied exclusively on DNA extracted from gut samples, yet we are unsure whether this represents viable bacteria (Emerson *et al*., 2017).

Our goal in this work was to survey the phylogenetic and metabolic profiles of the gut microbiota from *Xenopus* tadpoles during development and metamorphosis. We describe the gut microbiome communities found at different life stages and their activities, the communities found in different adult organs, including the skin, stomach, intestine and rectum and the effect of the diet on tadpole’s gut bacterial communities. We also investigated the transmission of bacterial communities between parents and their eggs. Finally, we explored host-gut microbiota metabolic interactions using functional annotations of the tadpole metagenome with the goal of expanding our understanding of its functional potential.

## Methods

### Animals and animal husbandry

We used *Xenopus tropicalis* frogs from the TGA and Sierra Leone strains and we let them reproduce Vusing induced natural mating as described in the supplementary Extended_materials_and_methods. Parents were transferred back to their housing tank while the embryos were let in the mating tanks. Tadpoles were reared in static water throughout embryogenesis and metamorphosis. Tadpoles and adults were euthanized with tricaine methane sulfonate pH 7.5 at 5 g.L^-1^. We used the Nieuwkoop and Faber table of development to identify tadpole’s developmental stages (Faber and Nieuwkoop, 1994).

### Tissue sampling

We dissected euthanized tadpoles or adults in sterile amphibian phosphate buffered saline (aPBS) to collect the gastro-intestinal from the stomach to the rectum. For the sake of simplicity, we used the term “gut” to refer to the whole gastro-intestinal tract of tadpoles or adults throughout the manuscript. Whole embryos and early tadpoles were abundantly rinsed using aPBS before further processing. The skin of adult frogs was sampled after having abundantly rinsed the animals using aPBS. Sampling was performed by swabbing the dorsal and ventral skin areas using a sterile cotton swab. Feces were collected using a sterile Pasteur pipette from frogs that were individually housed in aPBS after reproduction. Samples of two-hundred eggs including their outer and inner jelly coat were collected using a sterile Pasteur pipette from the reproduction aquarium, and washed three times in sterile aPBS. All tissue samples used for DNA extraction and metabarcoding were then immersed in at least ten volumes of ethanol and stored at -20°C until DNA extraction. All tissue samples used for RNA extraction and metabarcoding were kept on ice and then processed for tissue lysis as explained in the Supplementary Extended_materials_and_methods.

### Bacterial purification from intestinal tracts

We homogenized each freshly dissected intestinal tract in 500 μL of ice-cold PBS using a Potter-Elvehjem homogenizer. We then concentrated the bacteria extract by successive filtrations on 40, 20 and 5 μm filters (nylon 40 μm cell strainer from BIOLOGIX; 20 μm net ring from Pharmacia Fine chemicals; Whatman® Puradisc 13 syringe filters 5 μm from SIGMA-ALDRICH®). Next, we harvested the cells by centrifugation at 13 000g for 15 min at room temperature. These pellets were then used for nucleic acid extraction (details in Supplementary Extended_materials_and_methods) or fixation and microscopy (for quality control) or flow cytometry analysis.

### Flow cytometry analysis

We prepared triplicate samples from tadpoles at stages NF45-48, NF51-54, NF56, NF60-61, 15 days froglets and adults. Each sample was made from five tadpole’s guts per replicate for the NF45-48 and NF51-54 stages, single individuals were used for later developmental stages. We fed the tadpoles two days before the first time point (stage NF45-48), and ten days before sampling the froglets. We prepared the filtered bacterial cell samples as described previously and we fixed the resuspended bacterial pellet in 1.0% paraformaldehyde at 4°C during 2 h. We analyzed samples prepared by mixing bacterial samples with fluorescent beads (Flow-Count™ fluorospheres, 10 µm diameter, 1010 beads/µl) to obtain a final concentration of 20 beads/µl. Bacterial samples were stained using propidium iodide (PI) with a MoFlo® Astrio^™^ cytometer (BECKMAN COULTER). Excitation was made at 488 nm and 561 nm for scattering (Forward Scatter, FSC; Side Scatter, SSC) and PI respectively. Emission of PI was collected through a 614/20-nm band pass filter. Each measure was done in triplicate to compute a mean value and the standard deviation for each biological replicate sample. Plots were normalized on 200,000 gated events for the analysis of cell populations.

### Metagenome and metatranscriptome sequencing

We prepared genomic DNA (gDNA) and RNA from freshly prepared 5 μm filtered gut samples prepared as previously described and obtained from five *X. tropicalis* tadpoles at stage NF56 raised in the same aquarium. We obtained 10-15 µg of total RNA and ∼1 µg of DNA per filtered tadpole gut. Metagenomic and metatranscriptomic library construction and sequencing were performed by the Beijing Genomics Institute (BGI) using an Illumina® HighSeq^™^ 2500. The insert sizes of the metagenomic and metatranscriptomic libraries were 170 bp and 180 bp, respectively. We obtained 47,279,786 and 40 000 000 high quality 100bp paired end sequences for metagenomic (ERS716504) and metatranscriptomic (ERS716505), respectively.

### 16S rRNA and 16S rDNA library construction and sequencing

We prepared genomic DNA (gDNA) from dissected tissue samples that were fixed in ethanol and stored at -20°C before processing. RNA was prepared from fresh tissue. gDNA or RNA were used as template for 16S Ribosomal RNA (16S rRNA) or DNA (16S rDNA) amplification. A detailed list of samples used for the 16S rDNA and rRNA library construction is given in Supplementary_table_1. A first set of three replicate PCR reactions was performed to amplify the V3-V4 variable region of the 16S rDNA and rRNA; the PCR condition details and oligonucleotide sequences are given in Supplementary Extended_materials_and_methods. The PCR products were sent to the GeT sequencing facility (France Génomique, INRAE, Toulouse) for library preparation and sequencing. PCR products were purified and quantified by spectrophotometry and a second PCR was performed to integrate a barcode index. The barcoded PCR products were pooled and their quality was checked using capillarity electrophoresis and quantitative PCR. Finally, the PCR products were sequenced using a Miseq (supplied by Illumina®). We obtained a total of 3,647,548 clean read counts from 132 samples, with a minimum of 23,872, a maximum of 179,732 and a median of 43,174 reads. The metadata and number of sequences obtained by sample is given in Supplementary_table_1.

### 16S rRNA and 16s rDNA sequence analysis

We used the FROG pipeline for OTUs identification and taxonomic affiliation (Escudié *et al*., 2018). Overlapping reads were assembled and sequences were dereplicated before clustering using the SWARM algorithm. Chimeric sequences were then removed and OTUs were filtered on abundance for at least 0.005%. Taxonomic affiliation of the OTUs was performed based on the 16S SILVA database (V.123) using blast and RDP classifier. We used script implemented in R to perform different microbiome community analysis (provided on GitHub at https://npollet.github.io/metatetard/) (R Core Team, 2019). Source-sink analysis was performed using Feast (Shenhav *et al*., 2019).

### Taxonomic assignment for metagenome and metatranscriptome sequences

We used MATAM and phyloFlash to extract and assemble full-length or near full-length 16S and 18S rRNA gene sequences from the same set of filtered reads as the one used for the assembly (Pericard *et al*., 2018; Gruber-Vodicka *et al*., 2020). The assembled read and the details of the command lines are provided on GitHub at https://npollet.github.io/metatetard/Metagenome.html.

### Metagenome assembly and gene prediction

We filtered and trimmed reads according to their quality and excluded those derived from *Xenopus* genome using metaWRAP. We selected one assembly of these reads obtained with IDBA-UD using k-mers from 72 to 124 (Peng *et al*., 2012; Uritskiy *et al*., 2018). More details are given in the Supplementary Extended_materials_and_methods. We performed binning on contigs longer than 1000 bp using maxbin, concoct, metabat and DAStool (Alneberg *et al*., 2014; Wu *et al*., 2016; Sieber *et al*., 2018; Kang *et al*., 2019). We visualized binning results using vizbin (Laczny *et al*., 2015). We mapped OTUs, metagenome and metatranscriptome reads to the assembly using BBTools to derive coverage and tpm values at the scaffold and gene levels (BBMap).

We predicted CDS on the assembled metagenome scaffolds using Prokka (Seemann, 2014). We used Minpath for metabolic pathway prediction following the strategy and the tools provided in the IMP pipeline scripts as described in https://metagenomics-workshop.readthedocs.io (Ye and Doak, 2009; Narayanasamy *et al*., 2016). We mapped KEGG and EC identifiers on the Interactive Pathway Explorer V3 (iPATH3) to obtain a map of the tadpole’s gut microbiota metabolic pathway (Darzi *et al*., 2018).

## Results

### Enumeration of bacterial cells in *Xenopus* tadpole’s gut during development

We started to assess changes in tadpole’s gut communities throughout development by enumerating bacteria using flow cytometry (Figure 1). We selected six developmental stages from young tadpoles that just started to feed up to adulthood, through grown-up tadpoles undergoing metamorphosis (Figure 1A and Figure_S1). When we analyzed the distribution of bacterial populations by relative cell size and DNA content, we found that cytometric profiles differed according to life stages (Figure 1A). We observed distinct bacterial populations in mature tadpoles compared to young ones (e.g. populations 50-3 and 56-4 absent or extremely reduced in NF45 tadpoles, Figure 1A and Figure_S1). We also observed distinct cell populations in froglet and adult guts (population Juv-4 and Adu-3 for example, Figure 1A and Figure_S1). These first observations showed qualitative differences in the gut microbiomes across life stages.

**Figure 1:**
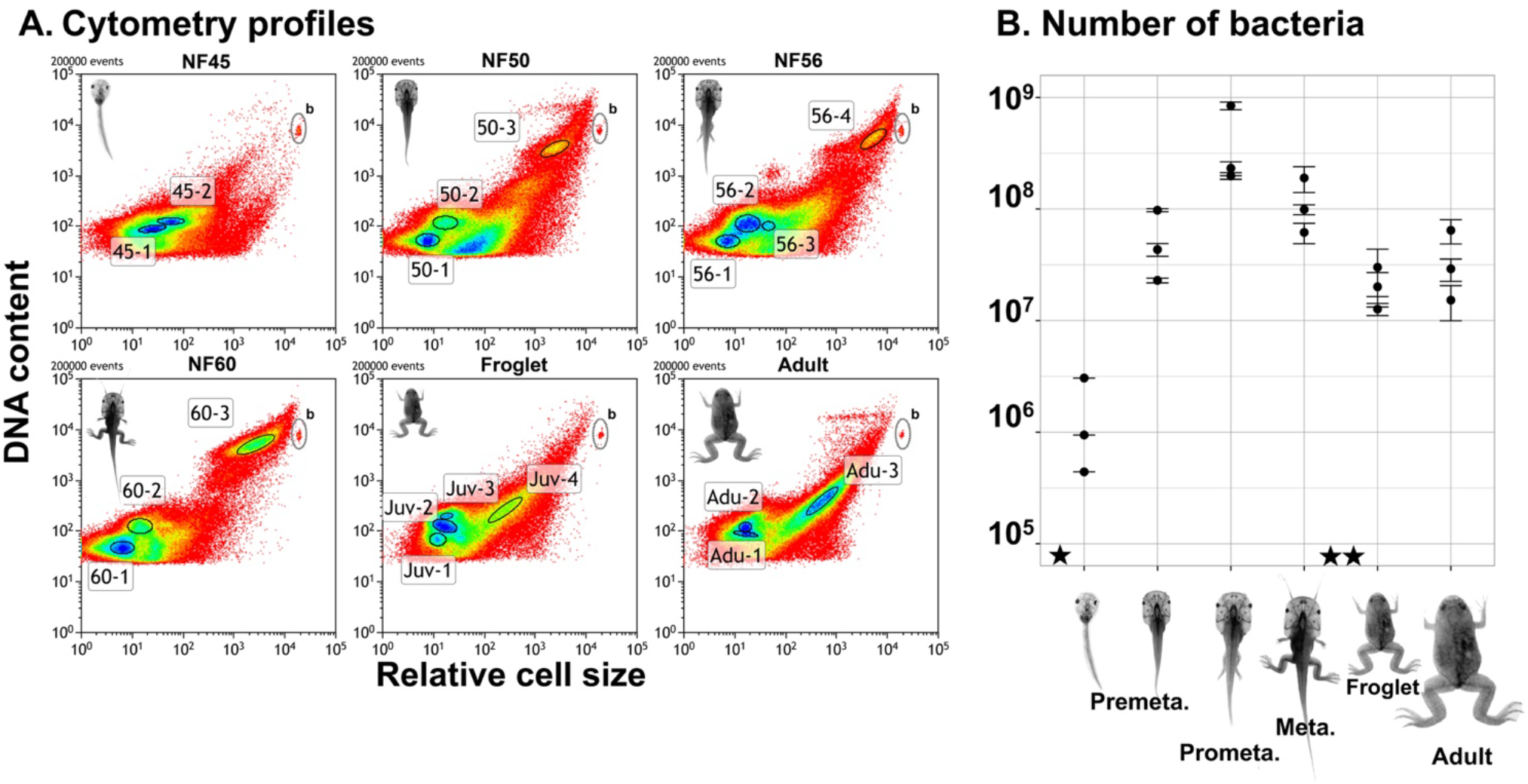
Cytometric profiles of gut bacteria during Xenopus tropicalis development. A. Cytometric profiles. Each dot in the plot represents one event (i.e. a cell), the color represents the density of overlapping point from red (low) to blue (high). The vertical axis represents DNA content by the measurement of propidium iodide (PI) fluorescence (614/20nm) and the horizontal axis represents relative cell size by the measurement of forward scatter. Different bacterial populations are highlighted by an ellipse and identified by a text label. Top-right events labelled with a “b” correspond to the beads used for normalization. Background signals have been removed. B. Enumeration of bacterial load during development. The vertical axis represents the number of bacteria and the horizontal axis corresponds to different stages of development. Each dot corresponds to a biological replicate, the error bar represents the standard deviation obtained by technical replicates. A ★ marks the beginning of feeding and ★★ marks the change from the tadpole to the adult diet.

We quantified the total bacterial populations and determined an average of 1×10^6^ bacteria per individual at the young tadpole stage, 2×10^8^ at the prometamorphic stage and 3×10^7^ in the froglets (Figure 1B), indicating that the gut bacterial mass increased just after the beginning of feeding and decreased simultaneously with the gut size reduction occurring during metamorphosis (Schreiber *et al*., 2005). We found that the bacterial load changed significantly during development (ANOVA P-value = 0.041). The increase between pre-feeding and prometamorphic tadpoles was significant (Tukey adjusted P-value = 0.046). We also observed a tendency for a decrease between prometamorphic tadpoles and adults (Tukey adjusted P-value = 0.073); and between prometamorphic tadpoles and froglets (Tukey adjusted P-value = 0.060).

We conclude from these cytometric data that there are qualitative and quantitative differences in the gut microbiota across *Xenopus* life stages. The tadpole’s gut is colonized by a large population of bacteria at the beginning of feeding, and this bacterial population increases during tadpole’s development, then reduces upon metamorphosis before increasing again upon froglet growth.

### *Xenopus* gut microbial diversity during development and metamorphosis

Since our cytometry results pointed to the presence of an important bacterial population in the tadpole’s guts, we went on to quantify the taxonomic diversity of this microbiota during development. We set out experiments in which we sampled whole tadpole’s guts at different life stages up to adulthood and quantified bacterial taxonomic diversity using 16S rRNA gene profiling. We compared 16s rRNA gene profiles amplified from tadpoles at the premetamorphic stages of development (tadpoles, NF54 to NF56, N=8), at the prometamorphic stages (prometamorphs, NF58 to NF61, N=6), at metamorphic stages (metamorphs, NF 62, N=5), at the end of metamorphosis (froglet, NF66, N=6) and from sexually mature adults (N=8). After filtering low abundance and singleton sequence clusters, we obtained an abundance table for a set of 666 OTUs. According to rarefaction and species accumulation curves, we obtained a depth of sequencing sufficient to obtain a fair evaluation of the richness for the most abundant bacterial species (Figure_S2).

We examined which bacteria constituted these most abundant gut communities (Figure 2 and Figure S3). We observed both global variations in bacterial phyla composition between samples from different life stages (Figure 2A) and inter-individual variations between samples from the same life stage (Figure 2C). In most cases, inter-individual variations were coherent with the global changes observed with the previous or the next life stage, and for this reason we pooled the observations from different samples belonging to the same life stage category for further analysis.

**Figure 2:**
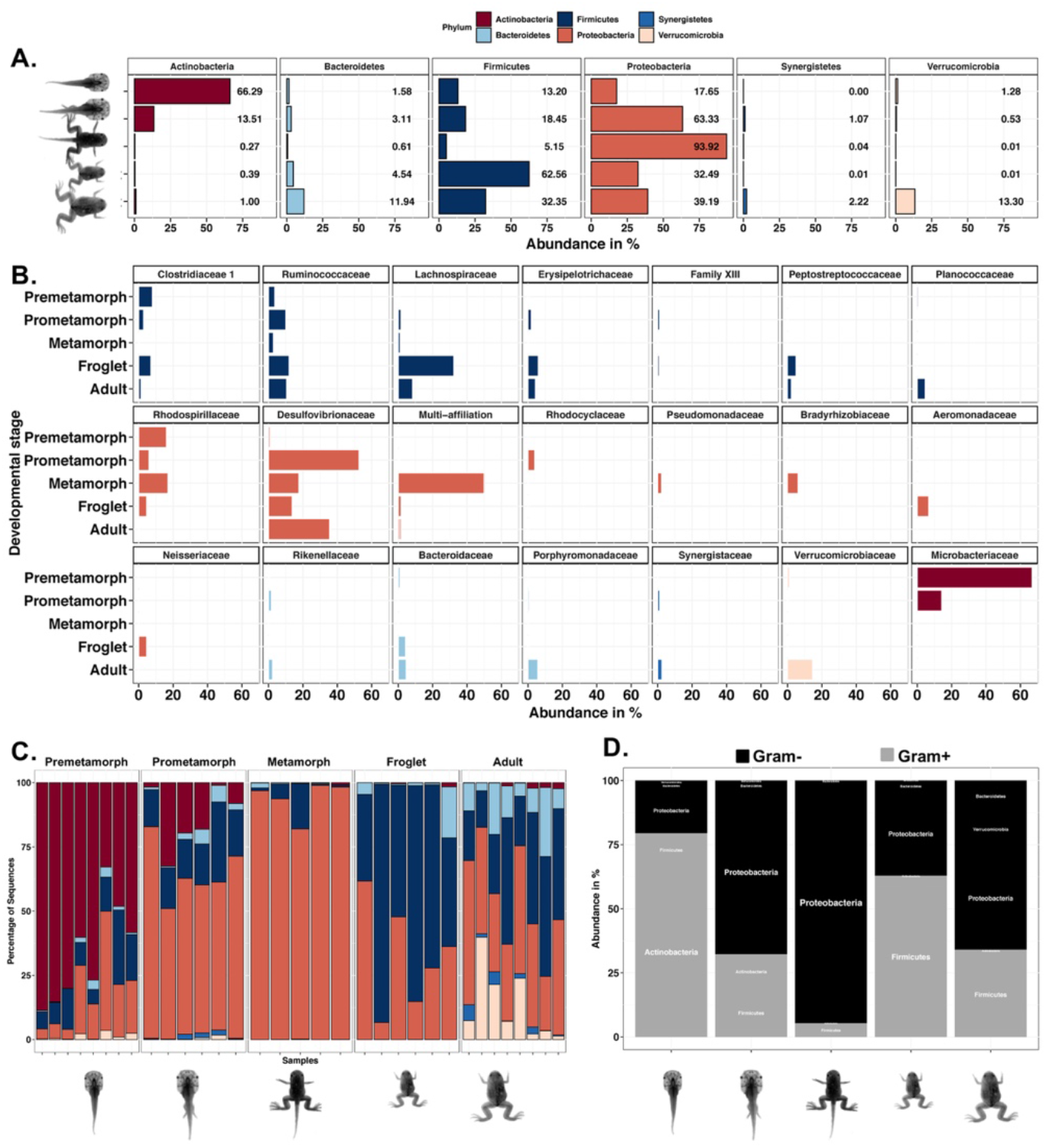
Phylum and Family-level compositional changes in the Xenopus gut microbiome across development. A. Bar plot representation of bacterial diversity at the phylum level. Results for the different life stages studied are shown in the following order, from top to bottom: Premetamorphic tadpoles (NF54 to NF56); Prometamorphic tadpoles (NF58 to NF61); Metamorphic tadpole (NF 62); Froglet (NF66); Adult: sexually mature adults (males and females). B. Bar plot representation of bacterial diversity at the family level, in the same order as A. C. Same as A but viewed at the sample level to highlight interindividual differences. D. Bar plot representation of Gram-positive and Gram-negative bacterial phyla.

At the premetamorphic stage of development, we observed that *Actinobacteria* dominated the tadpole’s gut microbiota (Figure 2A). At this stage, *Proteobacteria* (mostly *Alphaproteobacteria*) and *Firmicutes* were also abundant while *Bacteroidetes* and *Verrucomicrobia* contributions were quantitatively modest. The tadpole’s gut microbiota composition changed with *Proteobacteria* becoming the major phylum (mostly *Deltaproteobacteria*) in prometamorphic tadpoles, followed by *Firmicutes, Actinobacteria, Bacteroidetes, Synergistetes* and *Verrucomicrobia*. The proportion of *Proteobacteria* increased in metamorphic tadpoles (mostly *Alphaproteobacteria*), *Firmicutes* abundance decreased and *Bacteroidetes, Actinobacteria, Synergistetes* and *Verrucomicrobiae* were minor contributors. Once metamorphosis was completed, the gut microbiome of the froglets changed and *Firmicutes* became dominant, followed by *Proteobacteria* (mostly *Deltaproteobacteria*), *Bacteroidetes, Actinobacteria, Synergistetes* and *Verrucomicrobia*. In comparison, the composition of the adult gut microbiome was marked by more balanced abundances of *Proteobacteria* (mostly *Deltaproteobacteria*) and *Firmicutes*, a remarkable abundance of *Verrucomicrobia* (slightly higher than that of *Bacteroidetes)*, and a minor contribution of *Synergistetes* and *Actinobacteria*. The abundance patterns for *Proteobacteria* were marked by different contributions of *Alpha* and *Deltaproteobacteria*.

These patterns of abundances at the phylum level were attributable to a restricted number of families (Figure 2B). At the family level, three genera from three families of *Bacteroidetes* were predominant: *Bacteroides* (*Bacteroidaceae*) with a marked abundance increase at froglet and adult stage, and *dgA-11* (*Rikenellaceae*) with an increase during tadpole growth and then in adult stages, and *Parabacteroides* (*Porphyromonadaceae*) that increased only at the adult stage. *Verrucomicrobiaceae* was the dominant family of *Verrucomicrobia* in adult guts, and it was represented by a single genus, *Akkermansia*. Similarly, a single unknown genus of the *Synergistaceae* family was found at low abundance in prometamorphic tadpoles and in adults. *Synergistetes Microbacteriaceae* was the dominant family of *Actinobacteria* in pre and prometamorphosis tadpole’s guts, and it was represented by a single genus, *Leifsonia*. Seven families of *Firmicutes* were predominant: *Lachnospiraceae* (several genera) and *Peptostreptococcaceae* (one unknown genus) with a marked abundance increase at froglet and adult stage; *Planococcaceae* (one unknown genus) that increased only at the adult stage; *Clostridiaceae 1* (genus “*Clostridium sensu stricto 1*”), *Erysipelotrichaceae* (two genera *Coprobacillus* and *Erysipelatoclostridium*) and *Ruminococcaceae* (genera *Hydrogenoanaerobacterium, Anaerotruncus, Ruminococcaceae*) were found with fluctuating abundances across the life stages, and finally *Family XIII* was found only at low abundance. Eight families of *Proteobacteria* were predominant: *Aeromonadaceae* (genus *Aeromonas*), *Neisseriaceae* (genus *Vogesella*) and a family of *Rhizobiales* (one unknown genus) dominated the froglet gut microbiome; *Rhodospirillaceae* (one unknown genus) exhibited a tendency to decrease across development and were minor in adults; *Desulfovibrionaceae* (genus *Desulfovibrio*) were a minority at premetamorphosis and were abundant in the other stages, *Pseudomonadaceae* (genus *Pseudomonas*) and *Bradyrhizobiaceae* (unknown genus) were found in metamorph’s guts, and *Rhodocyclaceae* (genus *Azovibrio*) in prometamorphic tadpoles.

Altogether, we observed changes in the composition of the five most abundant bacterial phyla making the gut microbiome as development proceeds, both during tadpole growth and metamorphosis. The major shift during tadpole growth was a reduction of *Actinobacteria* and an increase of *Proteobacteria*, and the transition to the adult lifestyle was marked by the increase of *Synergistetes, Verrucomicrobia* and *Firmicutes*. This corresponds to a graded increase of Gram-negative bacteria during tadpole growth and development culminating at metamorphosis before reducing thereafter (Figure 2D).

We also analyzed changes of the estimations of OTU richness and phylogenetic diversity during development (Figure S3, Supplementary_table_2). We identified two important and significant changes: the first was an increase of bacterial species number and diversity that occurred during tadpole growth between pre and prometamorphosis (p=0.000 using betta and the breakaway estimation of richness); and the second was the opposite, namely a significant drop in species richness and diversity that occurred at the climax of metamorphosis (Figure S3 and Supplementary_table_2, (p=0.000 using betta and the breakaway estimation of richness). When focusing on a tadpole-adult comparison, we found that the adult *Xenopus* gut microbiome contained about twice more OTUs than the tadpole one, and its phylogenetic diversity was 1.8 times higher. These differences in microbiome diversity between the premetamorphic tadpole’s guts and the adult ones were significant using various alpha diversity metrics (Figure S3 and Supplementary_table_2).

In parallel to these changes of communities across development, numerous OTUs were shared between life stages. We found 111 OTUs (17%) present at all stages and representing 22% of the adult gut microbiome and 29 to 38% of the other life stage microbiomes (Figure_S4). Another set of 128 OTUs was shared between four life stages, 122 OTUs were shared between three, 151 between two and only 154 OTUs (23%) were found in only one of the developmental stages (Figure_S4). The most prevalent OTUs were detected with a threshold of very low relative abundances (< 5e-05 %, Figure_S4B). Nevertheless, we found a set of ten OTUs that were both common and abundant including an *alpha proteobacterium*, three *Desulfovibrio*, a *Microbacterioaceae*, a *Clostridium stricto sensu*, two *Hydrogenoanaerobacteria*, a *Rhizobiales* and a *Lachnospiraceae* (Figure_S4E). The abundances of the *Alphaproteobacteria*, one *Desulfovibrio*, the *Clostridium sensu stricto* and the *Lachnospiraceae* were the most homogeneous (Figure_S4E).

We searched more specifically which OTUs were characterized by a significant change of abundance across developmental stages using DESeq and identified a set of 22 OTUs corresponding to five phyla and 13 known genera (Figure_S5). The five OTUs most abundant in premetamorph tadpole’s gut but not in adult’s gut were a *Microbacteriaceae* (*Actinobacteria*), a *Rummeliibacillus* and a *Clostridium sensu stricto* 13 (*Firmicutes*), a *Bacteroides* (*Bacteroidetes*) and an *Alpha-proteobacteria* (*Proteobacteria*). The opposite pattern, i.e. more abundant in adult’s guts, was hallmarked by a *Lachnospiracae* (*Firmicutes*), a *Synergistaceae* (*Synergistetes*), a *Ruminococcaceae UCG-014* (*Firmicutes*), a *Rhizobiales* (*Proteobacteria*) and an *Anaerorhabdus furcosa* group (*Firmicutes*).

Significant differences in community composition (beta diversity) were evidenced by microbial community phylogenetic structure and ordination analysis (Figure S3C, Supplementary_table_2). The mean phylogenetic community structure and its dispersion differed according to life stages (Figure_S6). The phylogenetic relatedness of communities changed significantly during development, with negative and lower mean phylogenetic diversity values found in metamorph and froglet samples indicating more phylogenetically clustered microbial communities during metamorphosis. Overall, we conclude that the *Xenopus* gut bacterial community succession during development and metamorphosis underwent significant dynamic changes of composition and structure.

### *Xenopus* microbiota across several gut compartments

To further compare the bacterial diversity along the gut of adult frogs, we dissected the gut in three parts: stomach, intestine and rectum (N=3 for each tissue sample). In parallel, we analyzed the bacterial composition of the feces and the skin of these animals using 16S rRNA gene profiling (N=18 and N=20, respectively). Whereas *Bacteroidetes, Firmicutes* and *Proteobacteria* were the most abundant bacteria in the intestines, the rectum, the feces and the skin microbiota, we observed that the *Tenericutes* (mostly *Mollicutes*) were the most abundant in the stomach (Figure 3A, Supplementary_table_2). Surprisingly we observed a large quantity of *Acidobacteria* (28.8%) in only one feces sample. The stomach, intestine and rectum microbiomes were well differentiated from each other in composition and in structure analysis (PERMANOVA on bray-curtis dissimilarity F=6.93; R2=0.43; p<0.001, Figure 3B and Supplementary_table_2). Feces and skin microbiomes were characterized by their largest variability and were well separated along the x-axis in the ordination analysis.

**Figure 3:**
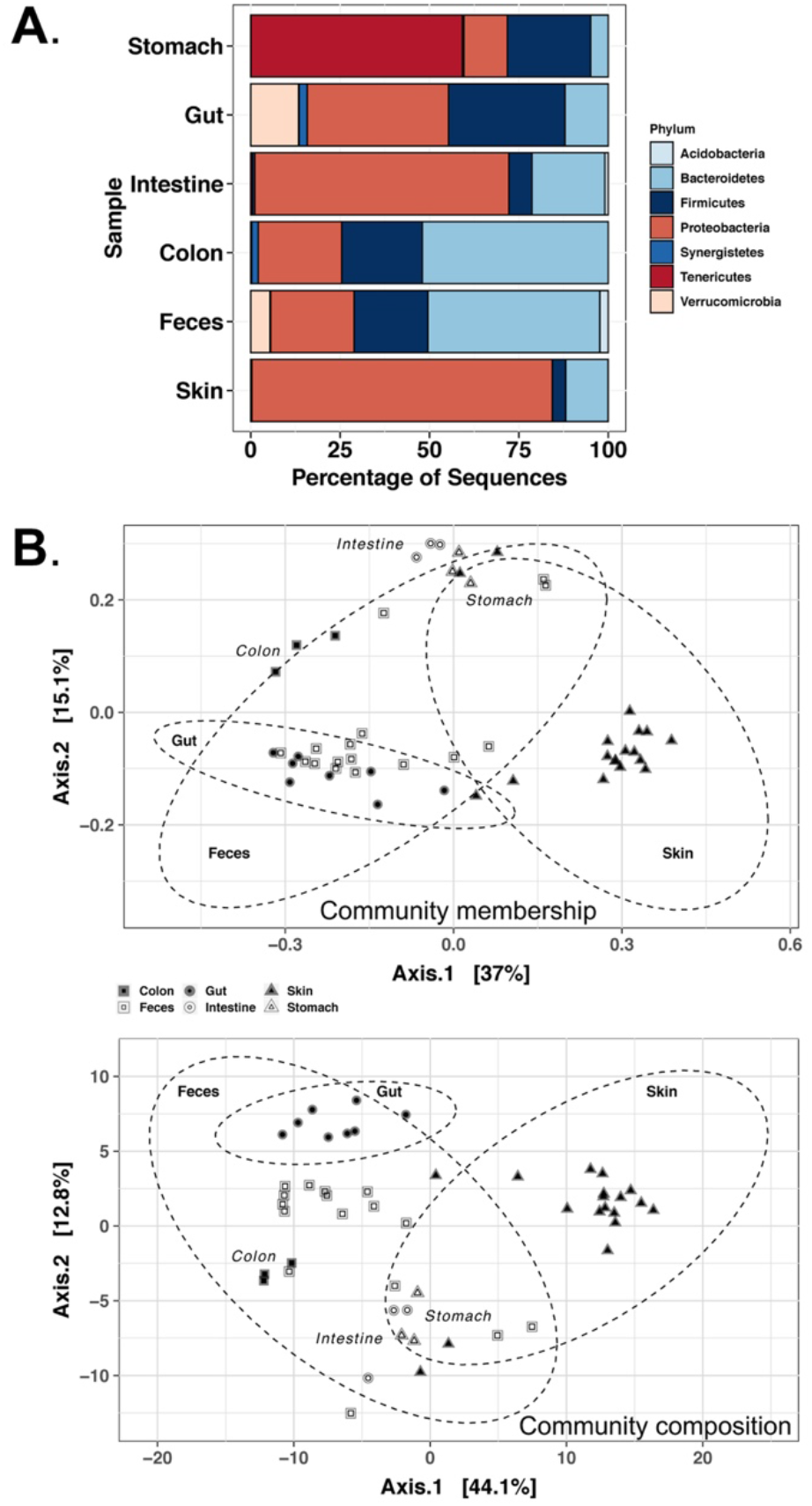
Xenopus microbiota across several adult organs and samples. A. Bar plot representation of bacterial diversity at the phylum level. B. Principal coordinate analysis of microbial communities across several adult organs and samples. Community membership was analyzed using unweighted UniFrac distance, and community structure was analyzed using weighted UniFrac distance. Percentages of the explained variations are indicated on both axes.

Despite these differences, we identified 31 OTUs shared between the three gut sections: 16 were identified as *Bacteroidetes* (including 15 *Bacteroidia*), seven as *Firmicutes* (including six *Clostridia*), six as Proteobacteria (including five *Desulfovibrio*), one *Parcubacteria* and one *Synergistetes*. In conclusion, we showed that the bacterial diversity increased along the anteroposterior axis of the adult *Xenopus* gut, from the stomach to the rectum, and the bacterial community compositions and structures associated with those gut sections were also significantly different (Supplementary_table_2).

### Activity of the *Xenopus* tadpole gut microbiome at the onset of feeding

We then asked when bacterial communities started to be active during development and what bacteria were found at the earliest stages of larval development. When *Xenopus* embryos hatch, their mouth is not open, the archenteron is well formed and the post-anal gut is open. Feeding behavior will only start once the mouth is formed and the gut organogenesis is more advanced, a few days later (Chalmers and Slack, 1998). We took advantage of this life history trait to perform a first investigation of the succession of active bacterial communities at early stages of larval development just after hatching when the mouth is not open (NF 30), before (NF41) and after feeding (NF48); and in growing premetamorphic tadpoles (NF50 and NF52). We set out experiments in which we followed-up embryos and tadpoles from two egg clutches up to the completion of metamorphosis. We targeted active bacteria by collecting RNA from pools of whole embryos (N=2 using pools of 25 embryos) or tadpoles (N=2 using pools of 25 NF48, 20 NF50 or 5 NF52 tadpoles) starting at 24h after fertilization and performing 16S rRNA gene sequencing.

We observed an increase of bacterial species number and diversity up to feeding stage, followed by a transient reduction at stage NF50 (Figure_S7A). We found 103 OTUs present across all stages, among which 11 were abundant OTUs (> 1%). At the earliest stage, we observed that most bacterial activity was attributed to *Proteobacteria* (five OTUs: *Rhodospirillaceae, Aeromonas, Rheinheimera, Pseudomonas, Desulfovibrio*) and *Bacteroidetes* (two *Bacteroides* OTUs, one *Flectobacillus*, one *Rikenella*), that *Firmicutes* (*Clostridium sensu stricto*) and *Synergistetes* (*Synergistaceae*) were already present while *Fusobacteria* (*Fusobacteriales*) and *Verrucomicrobia* (*Akkermansia*) were minor contributors (Figure_S7). We observed a different dynamic of bacterial community successions between the two clutches. In one experiment, there was a gradual decrease of *Proteobacteria* between NF30 and NF50, and an increase thereafter at stage NF52. This was the opposite for *Bacteroidetes*. In the other experiment, *Proteobacteriae* and *Synergistetes* abundances oscillated with a decrease before and after feeding while the abundance of *Bacteroidetes* varied in an opposite manner. This fluctuation was concomitant with a highest abundance of *Synergistetes* that increased after feeding. In both clutches, *Firmicutes* proportion increased up to feeding stage and decreased afterward. Altogether our results highlighted the presence of a rapidly developing microbiota along the first stages of tadpole development, characterized by a shift of the *Proteobacteria*/*Bacteroidetes* ratio.

We used the same strategy to monitor active bacterial communities in tadpoles at different stages of metamorphosis using 16S rRNA sequencing on whole digestive tract RNA extracts (N=2 using pools of five tadpole’s guts). We analyzed the prevalence and the abundance of bacteria as development proceeded. We found a common set of 33 OTUs present in at least 50 % of the samples with a relative abundance of 0.001% or more (Figure 4). All these OTUs were also found at early stages before feeding, and in the gut of tadpoles up to the completion of metamorphosis (Figure 4B). The five commonest OTUs were also the most abundant (> 0.01% abundance), and included unknown or ill-defined species: a *Rikenella*, an *Alphaproteobacteria*, a *Bacteroides*, a *Desulfovibrio* and a *Synergistaceae*. We found a few prevalent bacteria detected with low abundances such as an *Alphaproteobacteria*, two *Ruminococcaceae*, a *Rikenella* and an *Haliscomenobacter*.

**Figure 4:**
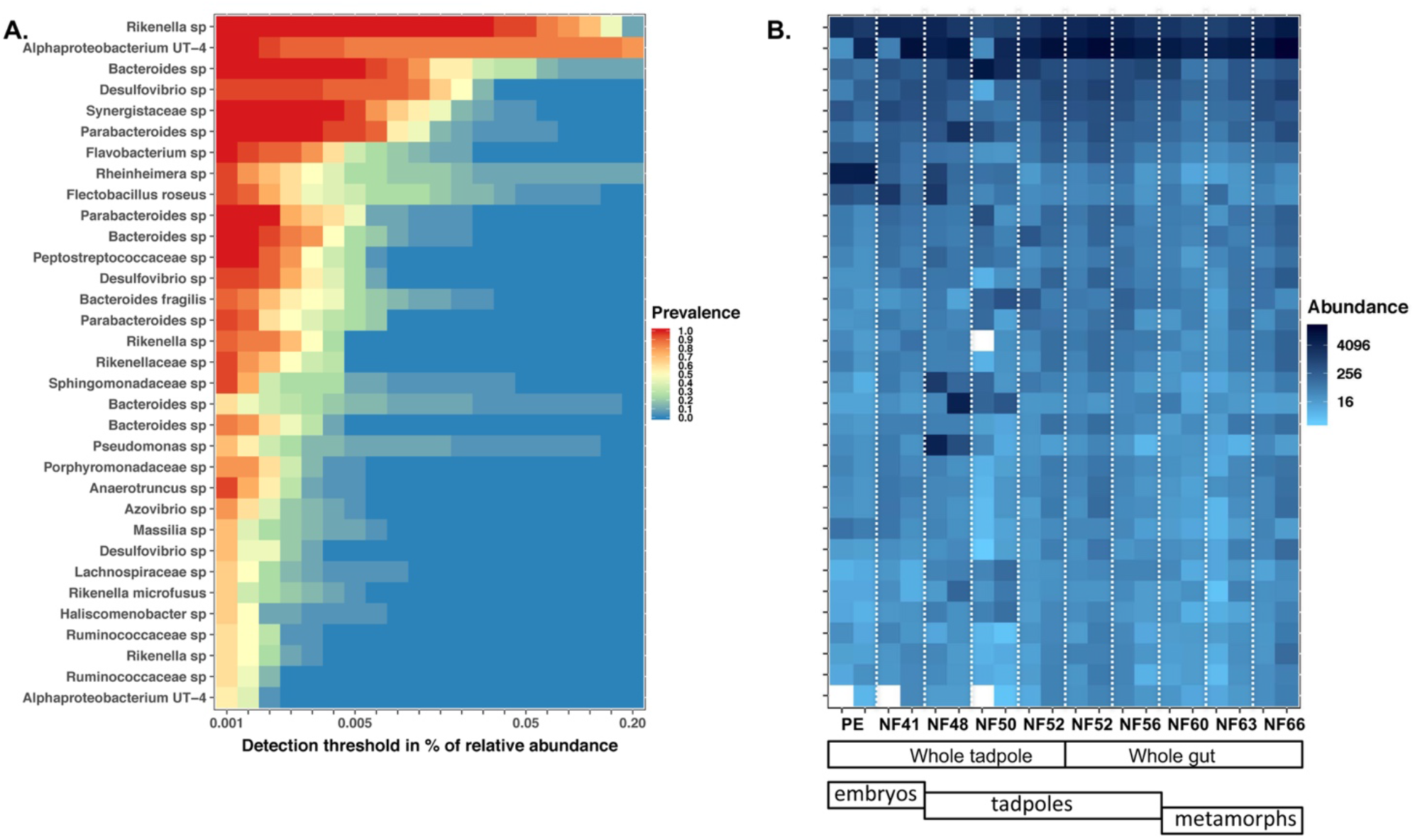
Prevalence and abundances of active bacteria across Xenopus development from post-hatching to metamorphosis. A. Heatmap of prevalence and abundances for 33 OTUs. The detection threshold scale is logarithmic and the metric used was the relative abundance. OTUs were identified using their taxonomic affiliation at the family genus or species level according to the resolution available for a given OTU. This analysis was performed on OTUs characterized by a minimal prevalence of 50 % and a minimal abundance of 0.001%. B. Abundance heatmap for the 33 most prevalent OTUs across the developmental stages sampled. Two clutches were followed up across development, and the white dotted lines separate the two biological replicate samples. The legend for stages is as follows: PE is a post-hatching NF30 tailbud embryonic stage; NF41 is a premetamorphic tadpole non-feeding stage; NF48 is a premetamorphic tadpole feeding stage; NF50; NF 52 and NF56 are premetamorphic tadpole feeding stages; NF60 is a prometamorphic tadpole stage; NF63 are metamorphic tadpole stages; NF66 is a froglet stage.

In a complementary analysis, we used a Bayesian algorithm for source tracking to estimate the proportion of OTUs populating the adult gut that originated from previous stages of development (Figure_S7C) (Shenhav *et al*., 2019). In both experiments, the source of most OTUs was identified as the feeding tadpole’s gut (52 and 95%), a minority was identified as stemming from the embryos (3 and 19%). Altogether, we identified that only few active bacteria found in the embryos before the feeding stage will be later populating the adult gut, and that a majority of tadpole’s active bacteria will pass through metamorphosis and be also present in the adult gut.

### Diet shapes *Xenopus* tadpole’ gut active communities

Since *Xenopus* tadpoles are suspension feeders, we hypothesized that tadpole’s diets could directly contribute to their active gut bacterial communities (Bles, E.J., 1906). We made an experiment in which we fed tadpoles with a commonly used micro planktonic-based diet (food 1, N=9) or a custom meal (food 2, N=9), and monitored their gut microbiome composition using 16S rRNA sequencing on RNAs. The micro planktonic food 1 contained a higher level of fibers (6.4%) than food 2 (2.6%). The custom meal food 2 was sterile, devoid of chitin and richer in available glucids (23.9%). The food regime did not change significantly tadpole growth (Kruskal-Wallis chi-squared = 0.99, df = 1, p = 0.32) or development (Kruskal-Wallis chi-squared = 0.34, df = 1, p = 0.56, Figure_S8). We found a total of 485 OTUs shared between the tadpoles fed with one or another diet, and this was making up the majority of the OTUs found in each condition: 485 out of 581, i.e. 83% for food 1 and 485 out of 531 i.e. 91% for food 2. We did not observe significant changes of OTU richness (log_e_(W)=3.14, p=0.13) and phylodiversity using Faith’s index, (log_e_(W)=3.33, p=0.29). The differences of alpha-diversity measured using the Shannon or the Simpson indices were significant (log_e_(W)=4.34, p=0.001 and log_e_(W)=4.30, p=0.004, respectively) pointing toward a higher specific richness in tadpoles fed with food 1 (micro planktonic-based diet). This difference was confirmed by comparing community structures: the mean standardized effect size of mean phylogenetic distances in communities (SES MPD) was significantly different (log_e_(W)=0.69, p=0.001), as did beta-diversity spread (global PERMDISP F=13.42, df=1, N.perm=999, p=0.001). However, the mean standardized effect size of mean nearest taxon distance (SES MNTD) was not significantly different (log_e_(W)=3.93, p=0.377). The gut bacterial community from tadpoles fed on food 2 (custom meal) was globally more phylogenetically even (Figure_S8). The custom meal was associated with more *Synergistetes* at the expense of *Firmicutes* and *Bacteroidetes* (Figure 5). The taxonomic and phylogenetic composition of gut bacterial communities were also significantly impacted by the diet (PERMANOVA on bray-curtis dissimilarity F=12.64, R2=0.44, p=0.001 and on MNTD dissimilarity F=29.13, R2=0.64, p=0.001), as evidenced on an ordination plot (Figure 5). The relative abundances of all five most abundant Phylum differed according to the diet, the most notable being the *Synergistetes* that accounted for 0.05 to 0.76% in the planktonic diet condition and for 0.80 to 17.72% in the sterile diet condition (Figure 5). We found 82 OTUs whose abundance differed significantly according to diet (Figure_S8). For example, *Chitinophagacae* were abundant in the tadpoles fed with the micro planktonic-based diet, while a *Sphingobacteriales* was most abundant in tadpoles fed with the artificial diet. Altogether, we observed that *Xenopus* tadpoles gut microbial communities could be shaped by their diet without a noticeable impact on growth and development.

**Figure 5:**
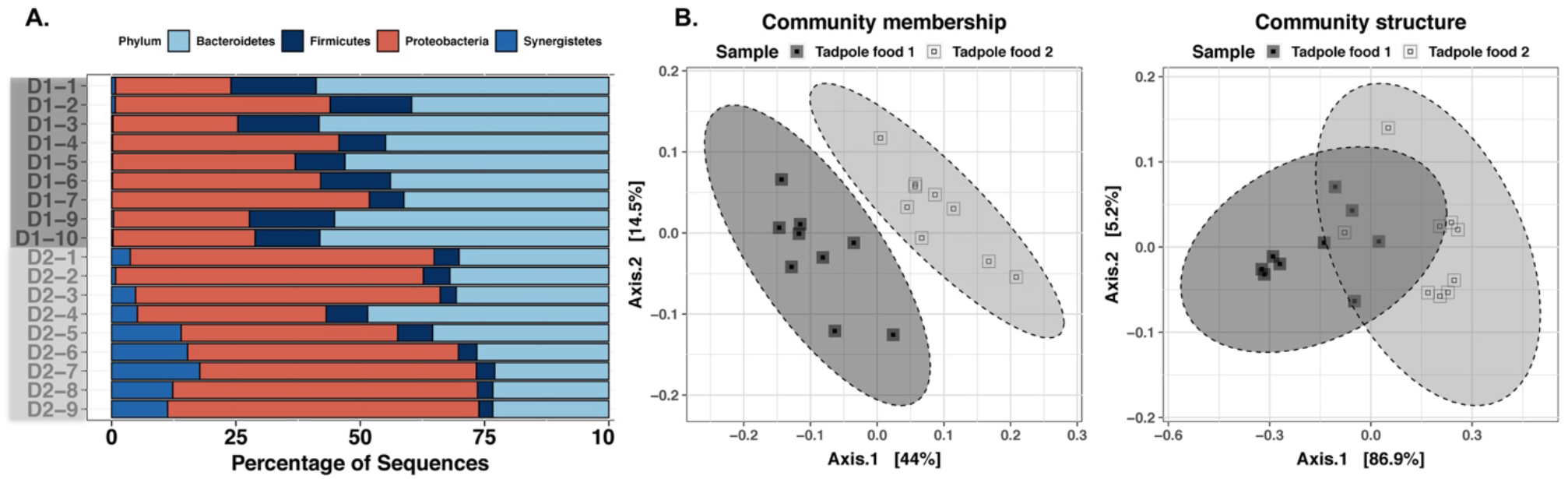
Influence of tadpole’s diet on their gut microbiome. A. Bar plot representation of bacterial diversity at the phylum level. B. Principal coordinate analysis of microbial communities across several adult organs and samples. Community membership was analyzed using unweighted UniFrac distance, and community structure was analyzed using weighted UniFrac distance. Percentages of the explained variations are indicated on both axes. Tadpole Food 1 (D1 samples) was a micro planktonic-based diet and tadpole food 2 (D2 samples) (food 1) was a sterile flour.

### *Xenopus* microbiome transmission

The way bacterial communities are transmitted across individuals and generations is a recurrent question in the study of animal microbiomes. We set out an experiment to evaluate the inheritance of bacterial communities between *Xenopus* parents and their clutch of eggs. We performed eight independent crosses and sampled the skin and the feces of both parents along with their egg clutch (N=16, N=13 and N=8, respectively). When we looked at bacterial diversity, we found that feces contained more *Verrucomicrobia* and *Firmicutes*, while both skin and eggs contained more *Proteobacteria* (Figure 6). In one case, we observed a resemblance between the communities of eggs and feces of one parent (Figure 6B). Globally, the bacterial communities of the skin and of the eggs were significantly more similar to each other than those from feces in terms of community structure and composition as evidenced using different metrics (Figure 6BD, Supplementary_table_2). We also analyzed the proportion of the egg bacteria derived from adult skin and feces using FEAST, a Bayesian algorithm for source tracking (Shenhav *et al*., 2019). In five out of the eight crosses, more than 70% of the bacteria found in the eggs did not come from feces or the skin of the parents. In the three other crosses, we observed that male or female skin made up to 75% of the egg bacteria. In conclusion, we found that the main driver of egg bacterial communities was the environment; the skin and the feces of parents were only minor contributors.

**Figure 6:**
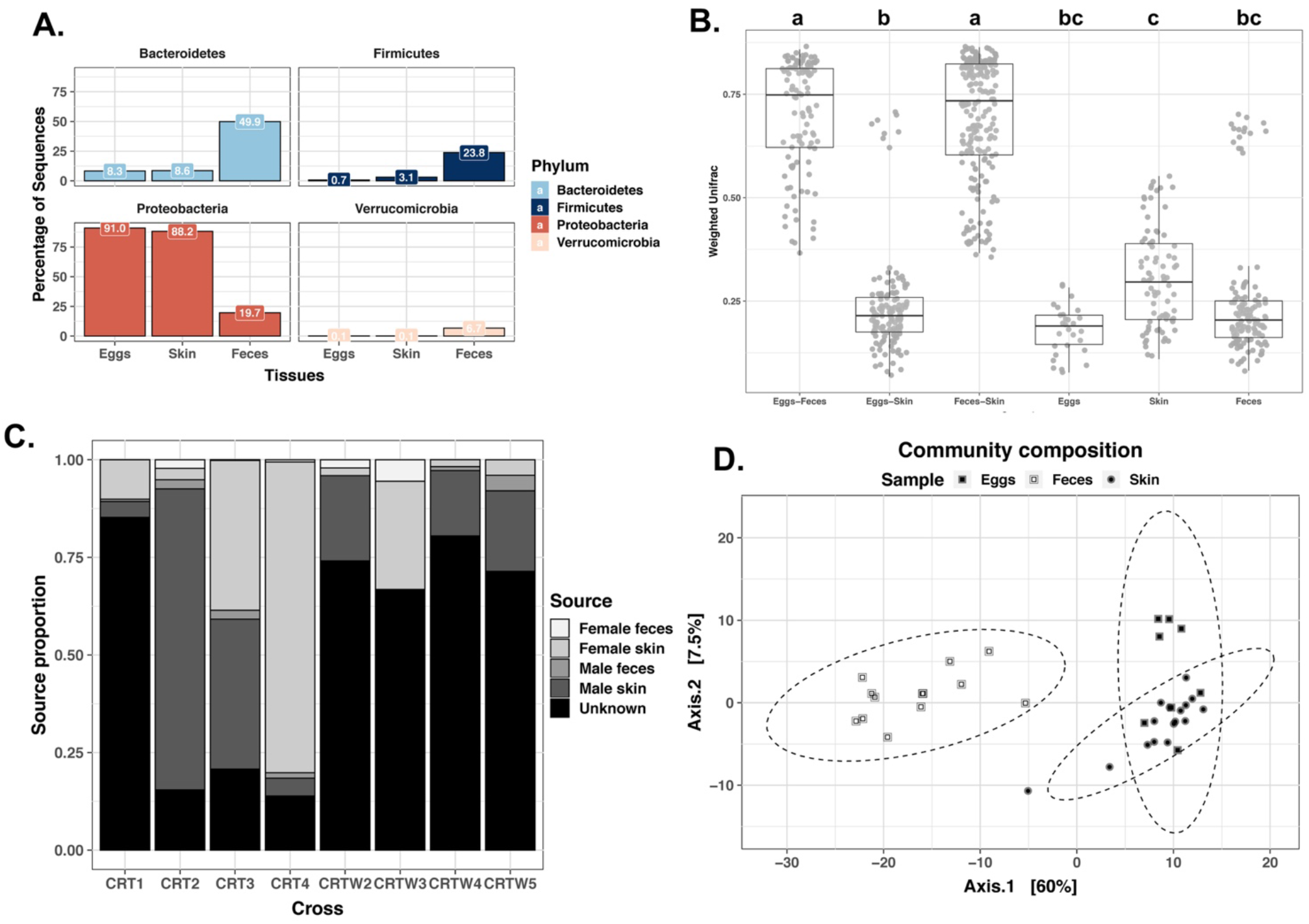
X. tropicalis microbiome transmission. A. Bar plot representation of bacterial diversity at the phylum level. B. Box-plot representation of weighted UniFrac distances between eggs, skin and feces. Letters a, b and c indicate significant differences at the 0.05 level between or within samples according to pairwise Dunn’s rank sum post-hoc test (see Methods section for details). Note the outliers in the feces sample and in the eggs-skin comparison. C. Bar plot representation of source proportions in egg microbiomes. D. Principal coordinate analysis of microbial communities across eggs, feces and skin samples. Community composition was analyzed using PhilR distance. Percentages of explained variations are indicated on both axes.

### Whole shotgun metagenomic and metatranscriptomic analysis of *X. tropicalis* gut microbiota

In the final part of our analysis and since few studies have investigated the genomic functions of the amphibian microbiome, we used a whole shotgun metagenomic and metatranscriptomic approach to better characterize the *Xenopus* gut metagenome and its activity in prometamorphic tadpoles (Figure 7). We obtained a dataset made of 23.6 and 19.8 million paired reads using the metagenomic and metatranscriptomic approach, respectively after filtering out reads mapping to the nuclear or the mitochondrial genome of *X. tropicalis*. We used MATAM, phyloFlash, kraken2 plus bracken and kaiju to perform taxonomic assignation of these reads against nucleotidic and proteic databases (Menzel *et al*., 2016; Pericard *et al*., 2018; Wood *et al*., 2019; Gruber-Vodicka *et al*., 2020). As expected, bacterial reads were predominant, but we identified also a few microeukaryotes, including various parasites (Figure S9, Figure S10, Figure S11, Figure S12). We found low levels of reads assigned to Archaea using kraken2/bracken and kaiju but these results turned out to be false positives upon closer inspection. Indeed, more discriminant sequence comparison tools such as MATAM, phyloFlash or megablast against SILVA or the NCBI non-redundant nucleotide databases evidenced that these reads were of bacterial origin. We also found a few thousands bacteriophage viral sequences using kraken2/bracken or kaiju but we did not investigate them in more details.

**Figure 7:**
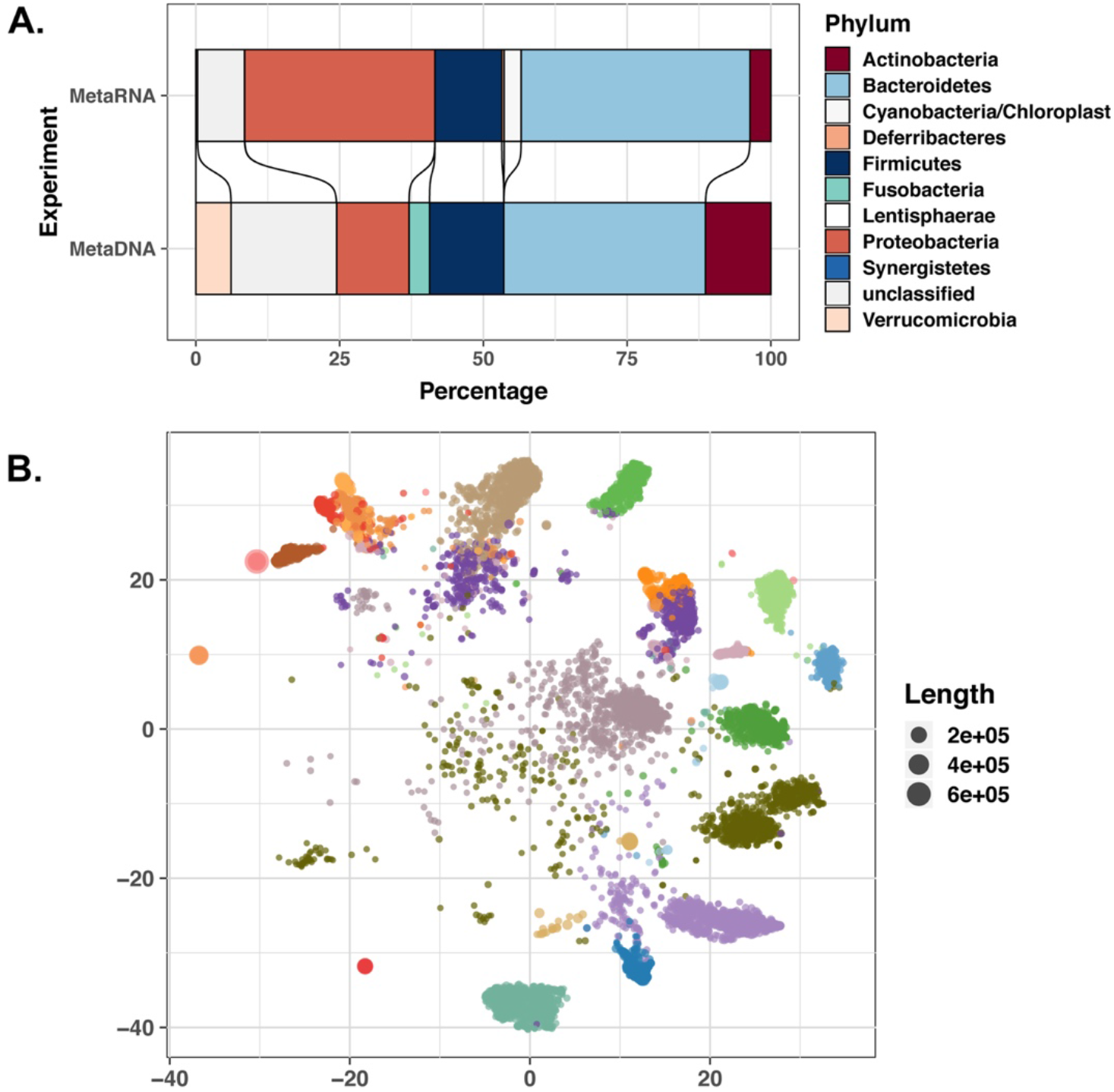
Bacterial taxonomic profiles from X. tropicalis prometamorphic tadpole gut. A. Histogram representation of bacterial diversity at the phylum level observed in a pool of five prometamorphic tadpole’s guts at stage NF56. metaDNA: metagenomic analysis; metaRNA: metatranscriptomic analysis. Each color in the histogram refers to a phylum as described in the legend. B. Vizualisation of the metagenomic assembly. Each bin identified using DASTool is represented in a different colour, each circle is a contig. The circles diameter is proportional to contig length.

Metagenomic and metatranscriptomic sequencing gave similar taxonomic profiles for bacteria according to the number of reads matching 16s rRNA gene sequences (Figure 7A). Using phyloFlash on metagenomic reads, we assembled 16 near full-length SSU sequences with phyloFlash-SPADES; 31 with phyloFlash-emirge and 33 using MATAM (Figure S9, Figure S10). Using metatranscriptomic reads, we assembled 32 near full-length SSU sequences with phyloFlash-spades; 1498 using phyloFlash-emirge and 440 using MATAM (Figure S11, Figure S12). We observed four main dominant phyla: *Bacteroidetes* (mostly *Bacteroides*), *Proteobacteria* (mostly *Desulfovibrio* and *Polynucleobacter*), *Firmicutes* (mostly *Ruminococcus*) and *Actinobacteria* (mostly *Microbacterium*) (Figure 7A).

In conclusion, we confirmed that the *X. tropicalis* tadpole’s gut microbiota harbors a large amount and diversity of bacterial cells, mainly constituted of anaerobic bacteria and common bacteria at the species level with other vertebrate’s gut microbiota.

### *Xenopus tropicalis* gut microbiota genes catalog

To characterize the *Xenopus* microbiome, we compiled a gene catalog from our metagenome and metatranscriptome data sets. We assembled the metagenomic sequence reads using different assemblers and compared the resultant assemblies with metaQUAST (Mikheenko *et al*., 2016). In addition, we mapped back all metagenomic and metatranscriptomic reads and also our previously found OTU sequences (Supplementary table 3). We finally selected one assembly made using idba_ud, based on the rate of properly mapped reads: 93.36% and 77.58% for DNA and RNA reads, respectively and on the number of OTUs identified: 23 OTU sequences matching perfectly and 78 OUT sequences matching with 97% identity to at least one contig. This assembly consisted of 72,183 contigs larger than 500 bp, with a total assembly length of 161.8 Mbp. The contigs N50 size was 6,100 bp with the largest contig being 640,454 bp long (Supplementary table 3). We then used DAStool to identify 22 bins covering 61.5 Mbp, including 14 bins with high completeness (> 90%) and low redundancy (ie contamination, < 8%) (Figure 7B, Supplementary table 4). We found a single perfect alignment of seven OTU sequences in seven different bins, and two bins containing two unrelated OTUs each.

We predicted a total of 141,692 CDS using prokka on the tadpole metagenome assembly. A subset of 87,012 CDS (61.4%) corresponded to hypothetical proteins. At least one metatranscriptomic read could be mapped on 37,926 CDS (26.8%). For more than 95.0% of these “expressed” CDS, the relative expression level was less than 100 reads per kilobase per million (RPKM) while some were highly expressed with a relative expression higher than 10 000 RPKM.

In conclusion, we identified 141,692 CDS from the *X. tropicalis* tadpole gut metagenome. Using metatranscriptomic sequencing, we revealed that about 27% of these protein encoding genes are transcribed and therefore involved in a physiological activity.

### Metabolic profile of the *Xenopus* gut microbiota

To better characterize the metabolic capacity of the *Xenopus* tadpole’s gut metagenome, we used a functional annotation strategy starting from the 141,692 predicted CDS. Out of these, 34,103 were annotated by prokka with a COG identifier and 32,801 by an EC enzyme number. We used Minpath to identify 106 KEGG and 1,281 Metacyc metabolic pathways associated with these known proteins (Ye and Doak, 2009; Figure 8A). Enzymatic activities linked to degradation functions represented 48% of all predicted Metacyc pathways and those linked to biosynthesis represented 45% (Figure 8A). Pathways involved in fermentation represented 0.8% and those involved in detoxification 0.5%. The detoxification potential provided by the gut metagenome included chloroaromatic compounds and terpenoid degradation. A more detailed view of the pathways predicted from the tadpole gut metagenome can be seen on the Krona plots available in Figure_S13 and Figure_S14. Next, we compared the metabolic pathways encoded by the *X. tropicalis* gut metagenome with those encoded by the *X. tropicalis* genome (Figure_S15). To do so, we mapped the EC and KEGG identifiers associated to metagenome-predicted CDS or to annotated *Xenopus* genes on the Interactive Pathway Explorer (Darzi *et al*., 2018). We queried the resulting pathway map using terms corresponding to some of the numerous metabolites known to be derived from microbiome activity in vertebrates (Figure 8B).

**Figure 8:**
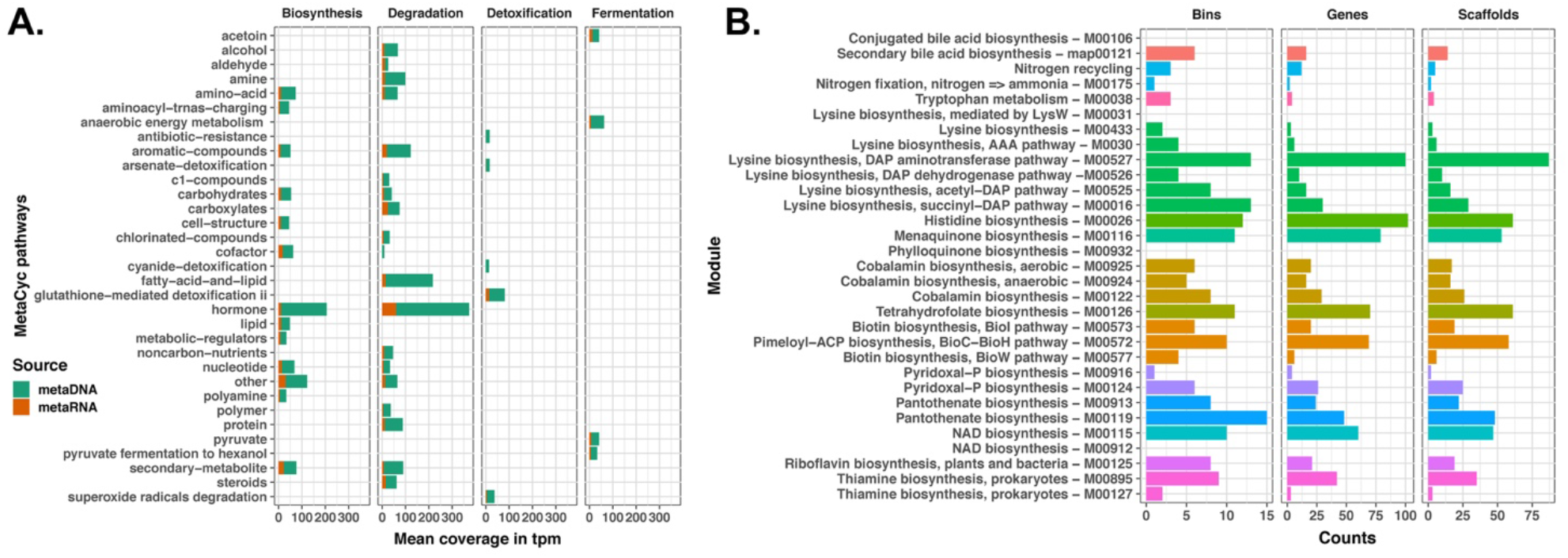
Overview of X. tropicalis tadpole’s microbiota metabolic functions. **A**. This barplot represents the read coverage (x-axis, in transcripts per kilobase million) on genes encoding enzymes participating in various Metacyc pathways (y-axis). **B**. Counts of metagenome bins, genes and scaffolds in which enzymes involved in selected biosynthetic pathways. Expressed genes encoding enzymes involved in selected KEGG modules mentioned on the y-axis were counted.

In herbivorous and omnivorous organisms, gut microbial communities are known to degrade complex carbohydrate fibers and provide short chain fatty acids. We identified 6,823 genes encoding 205 different families of carbohydrate-active enzymes (89 glycoside hydrolase, 50 Glycosyltransferase, 16 Polysaccharide lyase, 16 Carbohydrate esterase and 32 Carbohydrate-Binding Modules as defined in www.cazy.org) and evidence that acetate, propanoate and butyrate biosynthetic pathways were active in the *Xenopus* tadpole gut metagenome (Figure_S16). We also looked for enzymes involved in nitrogen recycling and found three CDS involved in N-fixation (*nifH);* 19 genes encoding urease that degrade urea into ammonia and carbon dioxide (*ureABC)* and two genes encoding uricase homologs (*puuD)*. Thus, tadpole’s gut microbes are involved in nitrogen waste recycling to synthesize amino acids and polypeptides that can act as nutrient sources. Indeed, we found that the known essential amino-acids histidine, lysine and tryptophan could be produced by tadpole gut microbes (Figure 8B, Figure_S16). Another category of molecular exchange between the microbiota and its vertebrate host involves the provision of vitamins (Magnúsdóttir *et al*., 2015; Dearing and Kohl, 2017). We found evidences that various B-vitamins (folate, cobalamin, pantothenate, riboflavin, biotin, thiamin and pyridoxin) and K-vitamin (menaquinone) could be synthesized by the tadpole gut microbiota (Figure 8B, Figure_S14). Similarly, we identified expressed bacterial genes encoding choloylglycine hydrolase involved in the biosynthesis of secondary bile acids (Figure 8B). In conclusion, we identified a set of metabolic pathways from the tadpole gut metagenome that are likely to influence tadpole’s physiology through the action of different metabolites.

## Discussion

We report that a large community of bacteria develops in the tadpole alimentary canal as soon as feeding starts and peaks before metamorphosis. While the gut microbiome biomass diminishes significantly at metamorphosis, it remains important. This probably reflects the feeding behavior of *Xenopus* at metamorphosis: Naitoh and colleagues reported that filter feeding can be observed in metamorphs up to stage NF59-NF61, and adult-like feeding starts at stage NF63 (Naitoh *et al*., 1989). Thus, there is only a short period of time, two to three days, during which the *Xenopus* metamorph stops feeding. Also, the gut itself starts to shorten at the metamorphic stage NF62 and will reduce four-fold to its adult-like length by stage NF66 (Schreiber *et al*., 2005; Heimeier *et al*., 2010). This means that adult feeding behavior occurs before the completion of gut remodeling, and that tadpole and adult digestive physiology overlap at the end of metamorphosis. In an animal-house setting, we deem it likely that *Xenopus* froglets continue to eat the available plankton in the rearing aquarium as long as a new diet is not artificially imposed to them, because they rely upon suction feeding as adults (Bles, E.J., 1906). More generally, the change of eaten food composition is likely to be gradual depending on the ecology of the amphibian species considered. For example, Fedewa reported insect material in the guts of *Bufo terrestris* and *Pseudacris crucifer* at Gosner stage 45-46 (NF64-NF65) i.e. in metamorph before the completion of metamorphosis, and the interval of fasting was the beginning of metamorphosis (Gosner stages 42-44, NF60-NF63) (Fedewa, 2006).

Our metabarcoding data using DNA and RNA-based 16S rRNA gene sequencing further demonstrated that the colonization of the tadpole by microorganisms occurred early during *Xenopus* tadpole development. Moreover, our results proved that *Xenopus* gut bacterial community composition changes during tadpole development and metamorphosis. At first, i.e. before and right after feeding, the change in composition of the microbiota is characterized by an increase of the *Bacteroidetes* / *Proteobacteria* ratio. The overall diversity of the microbiota seems to be reduced just before feeding, possibly because of the development and the activity of innate immune responses of the newly hatched larvae (Pasquier, 2014). After an increase of the microbiota diversity once feeding started, there is another step of reduction that we interpret as the result of an adaptation of the environmental bacterial population to the tadpole’s gut environment. In this period, when the tadpole starts feeding and grows rapidly (NF45 to NF56), bacterial populations increase 1,000-fold but the microbiota composition remains quite the same. During metamorphosis, we observed an important diminution of the bacterial abundance and a change of diversity. Indeed metamorphosis is associated with several modifications of the gut organs and we expected that gut reduction and neuroendocrine changes would impact the gut microbiota, as shown previously for other species (Fedewa, 2006; Kohl *et al*., 2013; Vences *et al*., 2016; Chai *et al*., 2018; Warne *et al*., 2019; Long *et al*., 2020; Zhang *et al*., 2020).

The bacterial diversity observed in the *X. tropicalis* gut was consistent with the results of Mashoof et al. on the diversity observed in *X. laevis* gut (Mashoof *et al*., 2013). Indeed, the same three phyla dominating the gut microbiota were observed in both analyses. The dominance of *Firmicutes, Bacteroidetes* and *Proteobacteria* is also a characteristic of the vertebrate gut in general (Colston and Jackson, 2016; Youngblut *et al*., 2019). Moreover the *Xenopus* gut microbiota diversity increased following the anteroposterior axis of the gut, as observed in human gut microbiota (Sekirov *et al*., 2010). We did not detect Archaea using our metabarcoding methodology, neither did we identify Archaea in our metagenomic and metatranscriptomic data. While we expected to miss Archaea in metabarcoding sequences because we relied on mainly bacteria-targeting primers, the lack of Archaea sequences in the shotgun sequences was more surprising. This constitutes a limitation of our survey and an interesting subject for further investigation in light of the account of methanogenesis in adult *Xenopus* (Saengkerdsub and Ricke, 2014).

We found that the abundance of several *Bacteroides, Bilophila* and *Lachnospiraceae* bacteria increase during tadpole development and metamorphosis. These bacteria are well-known to be abundant in the gastro-intestinal tracts of numerous vertebrates, including humans, and characterized by their resistance to high concentrations of bile salts or their bile acid hydrolase activities (Song *et al*., 2019). In addition, we identified seven CDS encoding an enzyme involved in secondary bile acid biosynthesis. We therefore propose an interpretation of our findings in light of bile acids being key molecular components regulating the gut microbiome composition during amphibian development and metamorphosis. Bile acids are small molecules produced mainly in the liver from cholesterol and further metabolized in the gastro-intestinal tract. They are known to act as molecular messengers between the host and their gut microbiota: on the host side, they play a role in numerous metabolic pathways via their cellular receptors and on the microbiome side they can modulate bacterial growth directly and indirectly (Ridlon *et al*., 2014; Wahlström *et al*., 2016). Bile acids can act directly on the gut microbiome because they have intrinsic antimicrobial activity and indirectly because their host cellular receptors regulate antimicrobial peptides gene expression. In turn, some bacteria encode bile hydrolases that enable the production of secondary bile acids (Joyce *et al*., 2014; Song *et al*., 2019). Therefore, we can consider bile acid composition as a trait of strong evolutionary significance since it links nutrition, gut, liver and gill physiology, host genome evolution and the gut microbiome composition (Reschly *et al*., 2008; Hofmann *et al*., 2010; Ridlon *et al*., 2014). In amphibians, bile acids are present in tadpoles but they increase in quantity and complexity after metamorphosis (Anderson *et al*., 1979; Cole and Little, 1983; Reschly *et al*., 2008). It is known that bile acids composition changes and complexity increases due to changes in host gene expression. We hypothesize that ontogenetic and endocrine regulation of amphibian host genes such as the bile acid nuclear receptor (Farnesoid X receptor, FXR; NR1H4), or its surface G-protein coupled receptor (TGR5) could be initial regulators of gut microbiome composition. FXR, an intra-cellular bile acid-sensing transcription factor of the nuclear factor superfamily, regulates bile acid and lipid homeostasis, and other genes such as FOR in *Xenopus* may play similar roles using different ligands (Seo *et al*., 2002; Reschly *et al*., 2008; Frisch and Alstrup, 2018). It is noteworthy that in mice TGR5 has been shown to link bile acid and thyroid hormone signaling by promoting intracellular thyroid hormone activity via the induction of deiodinase 2 in brown adipose tissue (Watanabe *et al*., 2006). In future studies, manipulation of gene activities, of bile acid composition and concentration and of the microbiome could be used to unravel the diversity and the evolution of these regulations in amphibians and other vertebrates.

We provide evidence that the diet of *Xenopus* tadpoles affected the diversity of their gut microbiome and thus that the mucosal immunity of tadpoles can interact with a variety of bacterial communities (Figure 5). Tadpoles fed a micro planctonic-based diet had a gut microbiome with a higher specific richness compared with tadpoles fed a sterile flour-diet. The latter diet was associated with a lack of *Chitinophagacae* involved in chitin degradation. These results contribute to the growing body of evidence that diet as an environmental factor can affect the composition of the gut microbiota.

Maternal transmission of microbes in animals is a well-known phenomenon but its documentation in amphibians is restricted to only few examples (Funkhouser and Bordenstein, 2013; Walke *et al*., 2014; Warne *et al*., 2019). In amphibians like in other animals, the transmission of microbial communities can be horizontal, vertical or environmental. The vertical transmission mode may depend critically on the reproductive behaviour of the animals. In *Xenopus*, the natural mating behaviour of the couple in amplexus involves looping dances from the bottom to the surface. Eggs are mostly laid by the female when she is near the surface, and immediately fertilized by the male. These movements may lead eggs to stick to the skin of either parent, and since the skin of amphibians harbors a rich microbiota, it can constitute a vector of vertical transmission from either parent. In addition, the eggs are laid through the female cloaca, and the sperm is also emitted by the male cloaca. Since the cloaca is a common anatomical structure shared by the reproductive, excretory and digestive systems, the microbial communities of these three systems may be transmitted to the eggs. Here we sampled the skin and feces of both parents to quantify the transmission of bacteria to the eggs. Quantitatively and using the Bayesian algorithm of source tracker, we found that the bacteria found on the eggs were mostly coming from the environment, and the second source was the skin of either parent. It was notable that a *Chitinimonas* species, a bacterium known to synthesize the antifungal Violacein, was one of the OTU transmitted from both parents in all crosses. This is relevant in the context of *Saprolegnia* egg infections and chytridiomycosis. Recently, Warne and collaborators reported that changes of the woodfrog egg microbiota influenced the bacterial communities at later larval stages (Warne *et al*., 2017, 2019).

*Bacteroidetes, Firmicutes* and *Proteobacteria* were slightly more represented in the metatranscriptome than in metagenome highlighting that those bacteria were present and active in the tadpole gut. *Actinobacteria* were less abundant in metatranscriptome than metagenome, showing a reduced activity. On the contrary, we found that *Verrucomicrobia* were very active.

Our data highlight the important capacity of the microbiota to complement the metabolic pathways of the *Xenopus* genome. We found that the gut microbiota of tadpoles synthesizes short chain fatty acids, essential amino acids and vitamins and recycle nitrogen waste products.

The need to perform investigations on animals from natural populations has rightly been underscored, for example by an interesting review of Sarah Hird (Hird, 2017). Yet we face several methodological difficulties when it comes to sample natural populations. In some studies, animals have been captured in the wild and raised thereafter in a laboratory setting. The current global threats on amphibians makes such studies difficult to realize and restricts severely the breadth of species that can be studied. In addition, the capture of vertebrate animals induces a variable amount of stress, leading to different responses including an immunosuppression (Rollins-Smith *et al*., 1997; Narayan, 2013). While the effect of captivity on amphibian microbiomes has been documented, there is a ample room for possible explanations (Becker *et al*., 2014; Loudon *et al*., 2014; Bataille *et al*., 2016). Also, parasitism is common in natural populations, and it can represent a hidden variable of considerable importance in microbiome studies. For example, the mutualism between the nematode *Gyrinicola batrachiensis* and *Rana catesbeiana* leads to a quicker tadpole development and has been linked to the hindgut fermentation capacity (Pryor and Bjorndal, 2005). Studies on laboratory animals for which rearing conditions are more uniform, with the potential for a better control of parameters and for reproducing experiments in different laboratories are definitively needed to make the link with investigations on natural populations. In this regard, *Xenopus* represents an interesting model since it is a widespread anuran with more than 20 species described in sub-Saharan Africa and classified of least-concern according to IUCN, and there is a vast body of biological knowledge relating to its development, its physiology and its genetic. We hope that our study will constitute a background knowledge enabling future work in the study of amphibian-microbial symbioses.

There were several important limitations to this study. First, marker gene and metagenomic studies are replete of confounding aspects and more studies will be needed to reproduce our observations using alternative technologies and more samples to explore inter-individual variability (Pollock *et al*., 2018; Bharti and Grimm, 2019; McLaren *et al*., 2019). This is especially needed regarding the construction of a metagenome gene catalog. Second, we analyzed the microbiome of *Xenopus* tadpoles in a laboratory setting and this is likely to differ from animals sampled *in natura*. Third, we did not survey Archaea nor microeucaryotes that are important players of the gut microbiome in other vertebrates and this will need dedicated efforts.

## Conclusions

In summary, we described the bacterial components of the *Xenopus* gut microbiota, the adult gut biogeography, the succession of communities during ontogeny, the impact of the alimentation in shaping the tadpole’s gut bacterial communities, the transmission of skin and fecal bacteria to the eggs. We also identified the most active gut bacteria and their metabolic contribution to the tadpole physiology. Keeping in mind the advantages of the *Xenopus* model, our data provided evidences that *X. tropicalis* raised in animal facilities is highly suitable for host-microbiota studies. Our study contributes to the growing body of research on the study of bacterial symbiosis in amphibians and on the evolutionary associations between amphibians and their microbiota. Further studies on the dynamics of the *X. tropicalis* gut microbiota during early development and metamorphosis should provide useful information on the evolution of host-microbiota interactions in vertebrates.

## List of abbreviations

NF: Nieuwkoop and Faber developmental stage

## Data accessibility

The sequence datasets generated and/or analyzed during the current study are available in the EMBL Nucleotide Sequence Database (ENA), metagenomic (ERS716504) and metatranscriptomic (ERS716505) https://www.ebi.ac.uk/ena/data/view/PRJEB9311. 16S rRNA gene sequence data are available at https://www.ebi.ac.uk/ena/data/view/PRJEB38248. OTU sequences are available at http://www.ebi.ac.uk/ena/data/view/LR992138-LR992913. Other data type supporting the findings are available within the article and its supplementary files.

## Supplementary material

R scripts for data analysis are provided on GitHub at https://npollet.github.io/metatetard/.

## Declarations

### Ethics approval and consent to participate

We used standard rearing conditions for Xenopus tadpoles and adults and all animal experiments were conducted in accordance with the regulation on Animal Experimentation (registration number 91–401 provided by Direction Départementale des Services Vétérinaires de l’Essonne).

## Author’s contributions

All authors read and approved the final manuscript. TS and IC performed experimental work under the guidance of NP. MB designed and supervised TC for cytometry, and analyzed cytometry results. TC performed whole genome shotgun sequence analysis. NP performed 16S rRNA gene sequence analysis. TS and NP wrote the manuscript.

## Acknowledgements

We are grateful to the Genotoul bioinformatics platform Toulouse Midi-Pyrenees (Bioinfo Genotoul) for providing computing and storage resources at the start of this project. The present work has benefitted from the facilities and expertise of the cytometry facilities of Imagerie-Gif, (http://www.i2bc.paris-saclay.fr/spip.php?article279). We thank Catherine Zanchetta and the team of the GeT facility for 16S metabarcoding sequencing. We thank Odile Bronchain, Albert Chesneau and Muriel Perron for providing access and use of their *Xenopus* animal facility in Orsay. We thank Yves Carton for pointing us the work of Olga Metchnikoff. We thank Franck Bourrat and Jean-Michel Rossignol for critically reading our manuscript. We are thankful to Wirulda Pootakham, Vanessa Marcelino and one anonymous reviewer for taking the time to review this manuscript. Their comments and suggests have greatly improved the manuscript. Version 4 of this preprint has been peer-reviewed and recommended by Peer Community In Genomics (https://doi.org/10.24072/pci.genomics.100005).

## Appendix

### Additional file 1: Supplementary_table_1.csv

<u>File format</u>: csv

<u>Title of data</u>: Metadata.

<u>Description of data</u>: This file contains the metadata corresponding to samples analysed by 16S rRNA gene metabarcoding.

### Additional file 2: Supplementary_table_2

<u>File format</u>: xlsx

<u>Title of data</u>: Compilation of statistical results in tables.

<u>Description of data</u>: This file contains seven tables compiling the results of significance tests for alpha and beta diversity metrics.

### Additional file 3: Figure_S1

<u>File format</u>: pdf

<u>Title of data</u>: Microbial cell populations identified by flow cytometry across developmental stages. <u>Description of data</u>: Cytometric profiles obtained at different stages of tadpole’s development were used to identify visually bacterial populations. Each population was plotted according to its DNA content and relative size (error bars correspond to the standard error deviation). Vertical axis represents the measurement of fluorescence (propidium iodide -614/20nm) and the horizontal axis the measurement of relative cell size (forward scatter, in arbitrary units). The bacterial populations clearly corresponding across life stages are grouped by a dotted ellipse.

### Additional file 4: Figure_S2

<u>File format</u>: pdf

<u>Title of data</u>: Rarefaction and species accumulation curves.

<u>Description of data</u>: These graphs present rarefaction curves (top) and species accumulation curves (bottom) for the different samples used in the listed experiments. * For premetamorph samples, the rarefaction curve is shown only up to 60,000 reads for the sake of consistency between all plots. Species richness corresponds to OTUs number. A detailed analysis is presented at https://npollet.github.io/metatetard/xpall_rarefaction_phyloseq.html

### Additional file 5: Figure_S3

<u>File format</u>: pdf

<u>Title of data</u>: Bacterial communities in the *Xenopus* gut across development. <u>Description of data</u>:

A. Graphical representation of the bacterial species richness. Each point corresponds to the breakaway richness estimate, error bars represent the measurement error according to the breakaway model. Premet: premetamorphic tadpoles NF54 to NF56. Promet: prometamorphic tadpoles NF58 to NF61; Meta: Metamorphic tadpole NF 62; Fro: Froglet NF66; Adult: sexually mature adults. Horizontal lines represent significant differences between the life stages connected; p=0.000 both between Premet. and Promet. and between Premet. and Adult. B. Representation of the bacterial species diversity measured using the Shannon index. Each large point corresponds to the divnet Shannon estimate from the samples of the same life stage, error bars represent the measurement error. Each small point corresponds to the divnet Shannon estimate from a given sample. The error bar corresponding to the measurement error can not be seen at this scale. Horizontal lines represent significant differences between the life stages connected; p=0.000 in all significant cases including the global test (p=0.875 for Premet-Meta). C. Principal coordinate analysis of gut-associated microbial communities during Xenopus development. Community membership was analyzed using unweighted Unifrac distance, and community composition was analyzed using philR distance. Percentages of the explained variations are indicated on both axes.

### Additional file 6: Figure_S4

<u>File format</u>: pdf

<u>Title of data</u>: Shared and contrasting bacterial communities in the Xenopus gut microbiome across development.

<u>Description of data</u>: A: Venn diagram showing the number of shared OTUs between samples of the same developmental stage category. B: Bar plot of the total number of OTUs in each developmental stage category. C: Number of shared and specific OTUs. D: Heatmap plot of the prevalence and abundances for low-abundance OTUs. The detection threshold scale is logarithmic and the metric used was the relative abundance. This analysis was performed on OTUs characterized by a minimal prevalence of 50 % and a minimal abundance of 1e-5 %. E. Prevalence and abundances of the most common and the most abundant OTUs across development. OTUs were identified using their taxonomic affiliation at the family genus or species level according to the resolution available for a given OTU. This analysis was performed on OTUs characterized by a minimal prevalence of 50 % and a minimal abundance of 0.001%. The right panel shows an abundance heatmap for the same OTUs to visualize their abundances in each sample.

### Additional file 7: Figure_S5

<u>File format</u>: pdf

<u>Title of data</u>: Contrasting OTUs across development

<u>Description of data</u>: A: Abundance fold changes across development. Contrasting OTUs were identied by filtering OTUs with an abundance of at least 100 reads across 20% of sample and using DEseq2 with a significance cutoff of 0.01. B: Heatmap representation of contrasting OTUs abundances across development. The heatmap was produced using a principal component analysis based on UniFrac distances.

### Additional file 8: Figure_S6

<u>File format</u>: pdf

<u>Title of data</u>: Variations of the phylogenetic structure of the *Xenopus* gut microbiota during development. <u>Description of data</u>: Standardized effect sizes of the Mean pairwise distance (left panel) and of the mean nearest taxon distance (right panel) measured using Faith’ PD index for *X. tropicalis* gut microbial communities across developmental life stages as indicated on the x-axis. Letters a, b and c indicate significant differences at the 0.05 level between developmental stages according to pairwise Dunn’s rank sum post-hoc test (see Methods section for details).

### Additional file 9: Figure_S7

<u>File format</u>: pdf

<u>Title of data</u>: *Xenopus* tadpole active bacterial communities: alpha-diversity, bacterial phylum diversity and source proportions

<u>Description of data</u>: A: Bar plot representation of OTU richness and phylogenetic diversity in active bacterial communities. Two clutches were followed up across early development, and the light and dark grey bars correspond to the two biological replicate samples. The legend for stages is as follows: NF30 is a post-hatching tailbud embryonic stage; NF41 is a premetamorphic tadpole non-feeding stage; NF48 is a premetamorphic tadpole feeding stage; NF50 and NF 52 are premetamorphic tadpole feeding stages. B: Bar plot representation of the bacterial phylum diversity. C: Bar plot representation of source proportions in froglet gut microbiome. Abbreviations SL and TGA refer to *X. tropicalis* strain’s names: SL for Sierra Leone strain and TGA for a laboratory population of Adiopodoume strain (Ivory Coast) outbred to Uyere strain (Nigeria).

### Additional file 10: Figure_S8

<u>File format</u>: pdf

<u>Title of data</u>: Impact of the food regime on tadpole growth and development.

<u>Description of data</u>: A: Impact of food diet on tadpole growth and development. Tadpoles were fed using either a micro planktonic-based diet (food 1) or a sterile flour (food 2). Their growth was assessed by weighting them, and their development was monitored by assessing their developmental stage according to the Nieuwkoop and Faber developmental table. The food regime did not change significantly tadpole growth (Kruskal-Wallis chi-squared = 0.99, df = 1, p = 0.32) or development (Kruskal-Wallis chi-squared = 0.34, df = 1, p = 0.56). B: Impact of the food regime on tadpole’s guts microbial communities. These bar plot shows values of the Standardized effect size of the Mean Pairwise Distance (SES MPD) and of the Mean Nearest Taxon Distance (SES MNTD) according to the food regime given to groups of tadpoles. The result of a Wilcoxon signed-rank test is shown above the graphs, with W being the Wilcoxon test statistic computed by this test, p the p-value, r the effect size, CI the confidence interval of the effect size and n the number of observations (see https://indrajeetpatil.github.io/statsExpressions/). C: Abundance ratios of the OTUs found significantly different according to a change of diet.

### Additional file 11: Figure_S9

<u>File format</u>: html

<u>Title of data</u>: Krona plot view of taxonomic affiliations inferred from a MATAM assembly of *X. tropicalis*

tadpole’s gut metagenomic reads.

<u>Description of data</u>: *X. tropicalis* tadpole’s gut metagenomic reads were used as input for the identification and assembly of SSU rRNA sequences using MATAM. Taxonomic affiliations were then inferred using SILVA

### Additional file 12: Figure_S10

<u>File format</u>: html

<u>Title of data</u>: Results of a phyloFlash assembly of *X. tropicalis* tadpole’s gut metagenomic reads.

<u>Description of data</u>: *X. tropicalis* tadpole’s gut metagenomic reads were used as input for the identification and assembly of LSU and SSU rRNA sequences using phyloFlash. The results for Spades and Emirge assembly engines are shown. Taxonomic affiliations were then inferred using SILVA

### Additional file 13: Figure_S11

<u>File format</u>: html

<u>Title of data</u>: Krona plot view of taxonomic affiliations inferred from a MATAM assembly of *X. tropicalis*

tadpole’s gut metatranscriptomic reads.

<u>Description of data</u>: *X. tropicalis* tadpole’s gut metatranscriptomic reads were used as input for the identification and assembly of SSU rRNA sequences using MATAM. Taxonomic affiliations were then inferred using SILVA

### Additional file 14: Figure_S12

<u>File format</u>: html

<u>Title of data</u>: Results of a phyloFlash assembly of *X. tropicalis* tadpole’s gut metatranscriptomic reads. <u>Description of data</u>: *X. tropicalis* tadpole’s gut metatranscriptomic reads were used as input for the identification and assembly of LSU and SSU rRNA sequences using phyloFlash. The results for Spades, Emirge and Trinity (trusted contigs) assembly engines are shown. Taxonomic affiliations were then inferred using SILVA

### Additional file 13: Supplementary table 3

<u>File format</u>: xlsx

<u>Title of data</u>: Comparison of metagenomic assemblies

<u>Description of data</u>: This file contains the metrics obtained using metaQUAST on different assemblies of metagenomic reads.

### Additional file 14: Supplementary table 4

<u>File format</u>: xlsx

<u>Title of data</u>: Results of DASTool binning

<u>Description of data</u>: This file contains the summary of DASTool binning results.

### Additional file 15: Figure_S13

<u>File format</u>: html

<u>Title of data</u>: Krona plot view of predicted KEGG metabolic pathways.

<u>Description of data</u>: KEGG metabolic pathways were predicted based on Minpath inference from EC enzyme numbers obtained using prokka.

### Additional file 16: Figure_S14

<u>File format</u>: html

<u>Title of data</u>: Krona plot view of predicted Metacyc metabolic pathways.

<u>Description of data</u>: Metacyc metabolic pathways were predicted based on Minpath inference from EC enzyme numbers obtained using prokka.

### Additional file 17: Figure_S15

<u>File format</u>: pdf

<u>Title of data</u>: Metabolic map of *X. tropicalis* genome and its gut metagenome.

<u>Description of data</u>: This metabolic pathway highlights in green the metabolic pathways predicted from the *Xenopus* genome and in blue those predicted from the tadpole gut metagenome. This map can be interactively accessed at https://pathways.embl.de/selection/pWbci4bo871W8Qm8XKF

### Additional file 18: Figure_S16

<u>File format</u>: pdf

<u>Title of data</u>: Selected cases of metabolic potential of *X. tropicalis* genome and its gut metagenome. <u>Description of data</u>: Fragments of biosynthetic pathways for common short-chain fatty acids, nitrogen recycling and B-vitamins is depicted.

This metabolic pathway highlights in green the metabolic pathways predicted from the *Xenopus* genome and in blue those predicted from the tadpole gut metagenome. This map can be interactively accessed at https://pathways.embl.de/selection/pWbci4bo871W8Qm8XKF

### Additional file 19: Extended_materials_and_methods

<u>File format</u>: docx

<u>Title of data</u>: Extended materials and methods.

<u>Description of data</u>: This file contains an extended version of the materials and methods

