## Extended_materials_and_methods for "Gut microbial ecology of Xenopus tadpoles across life stages"

### Animals and animal husbandry

We used standard rearing conditions for *Xenopus* tadpoles and adults and all animal experiments were conducted in accordance with the regulation on Animal Experimentation (registration number 91–401). *Xenopus tropicalis* frogs were from our colony of the TGA and Sierra Leone strains and we let them reproduce using induced natural mating. We picked males and females 48 to 72h before mating and injected each of them with 100 $\mu$l of amphibian PBS (0.67X PBS) containing 10-15 IU of human chorionic gonadotropin (Sigma-Aldrich, St. Louis MO). We injected them again with 100µl of amphibian PBS containing 100-150 IU of hormone in the morning of the mating day and placed one male and one female together in the same 16 L polycarbonate mating tank at 25 °C for a maximum of 10 h. Parents were transferred back to their housing tank while the embryos were let in the 16 L mating tanks. Tadpoles were reared in static water throughout embryogenesis and metamorphosis. Water was oxygenated and a third of the volume was manually exchanged on a daily basis. Once a week after the onset of feeding, we cleaned the tanks using a net to remove uneaten sedimented food. After metamorphosis, froglets were transferred to tanks in a flow through system. Tadpoles from stages NF45 to NF66 were fed with Sera Micron (Sera) while froglets and adults were fed with frozen worms (SARL Grebil Père et Fils) and Nasco pellets (Nasco). Tadpoles and adults were euthanized with tricaine methane sulfonate pH 7.5 at 5 $g.L^{-1}$. We used the Nieuwkoop and Faber table of development to identify tadpole's developmental stages [1].

The two batches of tadpoles used for the experiment on the impact of food were issued from the same clutch (named MT, Sierra Leone strain). Three sets of 30 embryos were sorted into three separate 16 L tanks for each diet condition. The set of nine tadpoles used for RNA metabarcoding were raised in the same 16 L tank and collected at the same time point. The two batches of tadpoles used for the RNA metabarcoding time series were each issued from the same clutch and the pools of tadpoles were made from individuals raised in the same rearing tank. The adult frogs sampled were raised in different tanks.

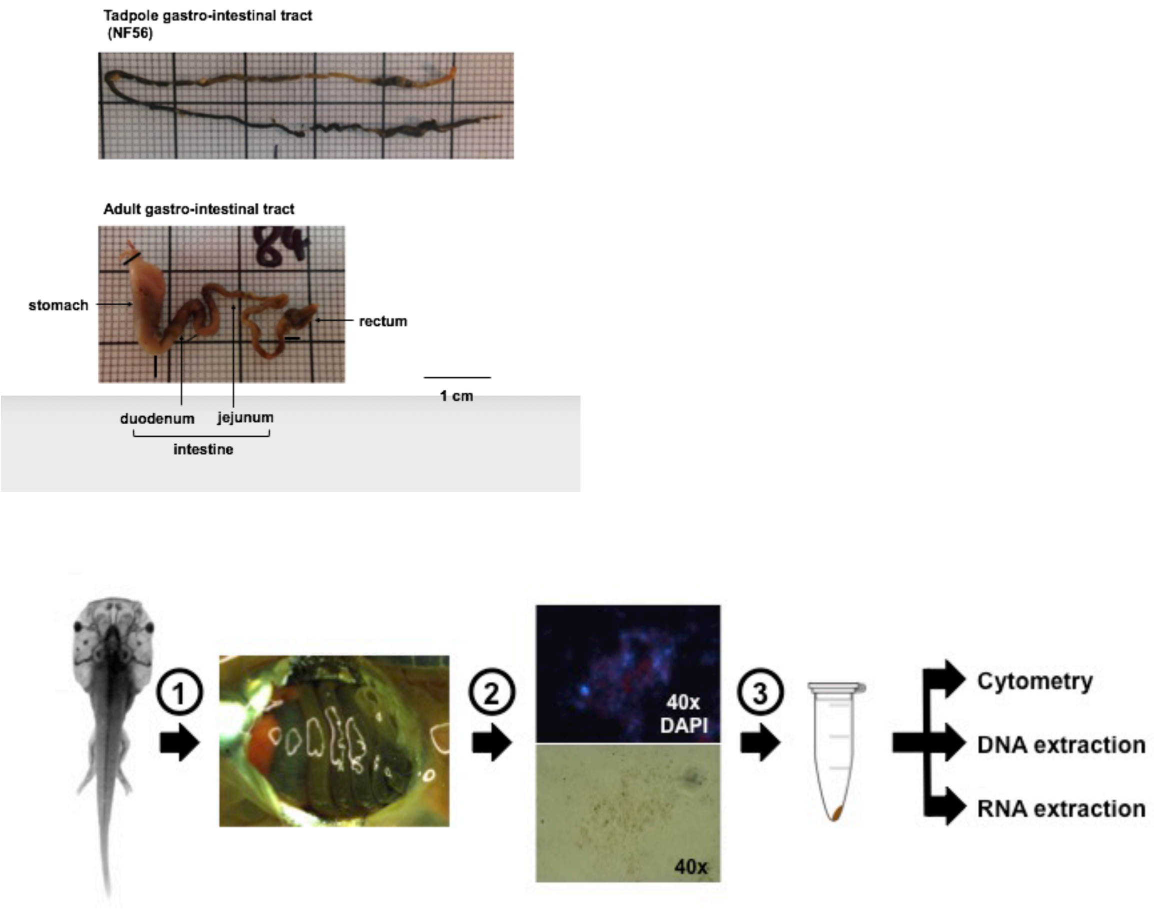


**Figure M1: Bacterial purification from *Xenopus* gut.**
(1) Tadpoles were euthanized and their guts dissected out (see gross anatomy of tadpole and adult gastro-intestinal tract on the top). (2) Guts were homogenized and filtered through 5µm filters. (3) Filtered cells were centrifuged and the pellets used for cytometry, DNA extraction or RNA extraction. The pictures in the second part of the protocol are examples of microscopic visualizations of the filtered cells with DAPI (top) or without DAPI (bottom).

### Tadpole food 1, diet 1 : Sera micron

(https://www.sera.de/fr/produit/aquarium-deau-douce/sera-micron/)

This finely powdered food contains both zooplankton (18% krill) and phytoplankton (51% spirulina).

**Ingredients :** Algae : spiruline (50 %), krill (16 %), fishmeal, shrimp, wheat flour, herbs, alfalfa, nettle, yeast, Ca-caseinate, marine algae, parsley, paprika, spinach, cod liver oil (including 34 % of omega fatty acids), gammares, carrots, whole egg powder, mannane-oligosaccharides (0,4 %), algae *Haematococcus*, *Perna canaliculus*, garlic.

**Analytical constituants :** Proteins 55.0 %, Fat 7.0 %, Fibers 6.4 %, Humidity 5.5 %, Ashes 10.5 %.
**Additives :** Vitamines and provitamines : Vit. A 6.200 UI/kg, Vit. $D_3$ 300 UI/kg, Vit. E (tocopherol D-L-acetate) 20 mg/kg, Vit. $B_1$ 6 mg/kg, Vit. $B_2$ 15 mg/kg, Vit. C stab. (L-ascorbyle monophosphate) 95 mg/kg.

### Tadpole food 2, diet 2 : MOD TESTDIET 5B8V W/ADDED VIT A, MEAL (5C6N catalog 1812140)

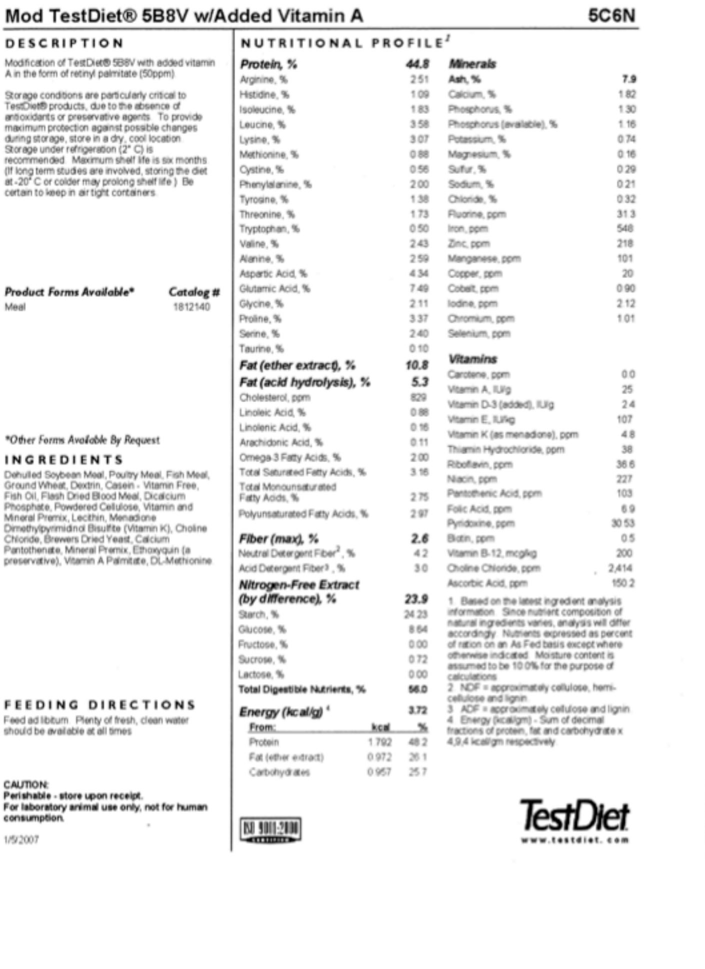

Mod TestDiet® 5B8V w/Added Vitamin A — 5C6N

DESCRIPTION

Modification of TestDiet® 5B8V with added vitamin A in the form of retinyl palmitate (50ppm)

Storage conditions are particularly critical to TestDiet® products, due to the absence of antioxidants or preservative agents. To provide maximum protection against possible changes during storage, store in a dry, cool location. Storage under refrigeration (2° C) is recommended. Maximum shelf life is six months (If long term studies are involved, storing the diet at -20° C or colder may prolong shelf life.) Be certain to keep in air tight containers.

| Product Forms Available* | Catalog # |
|---|---|
| Meal | 1812140 |

*Other Forms Available By Request

INGREDIENTS

Dehulled Soybean Meal, Poultry Meal, Fish Meal, Ground Wheat, Dextrin, Casein - Vitamin Free, Fish Oil, Flash Dried Blood Meal, Dicalcium Phosphate, Powdered Cellulose, Vitamin and Mineral Premix, Lecithin, Menadione Dimethylpyrimidinol Bisulfite (Vitamin K), Choline Chloride, Brewers Dried Yeast, Calcium Pantothenate, Mineral Premix, Ethoxyquin (a preservative), Vitamin A Palmitate, DL-Methionine

FEEDING DIRECTIONS

Feed ad libitum. Plenty of fresh, clean water should be available at all times.

CAUTION:
**Perishable - store upon receipt.**
**For laboratory animal use only, not for human consumption.**

1/5/2007

NUTRITIONAL PROFILE[1]

| | |
|---|---|
| ***Protein, %*** | ***44.8*** |
| Arginine, % | 2.51 |
| Histidine, % | 1.09 |
| Isoleucine, % | 1.83 |
| Leucine, % | 3.58 |
| Lysine, % | 3.07 |
| Methionine, % | 0.88 |
| Cystine, % | 0.56 |
| Phenylalanine, % | 2.00 |
| Tyrosine, % | 1.38 |
| Threonine, % | 1.73 |
| Tryptophan, % | 0.50 |
| Valine, % | 2.43 |
| Alanine, % | 2.59 |
| Aspartic Acid, % | 4.34 |
| Glutamic Acid, % | 7.49 |
| Glycine, % | 2.11 |
| Proline, % | 3.37 |
| Serine, % | 2.40 |
| Taurine, % | 0.10 |
| ***Fat (ether extract), %*** | ***10.8*** |
| ***Fat (acid hydrolysis), %*** | ***5.3*** |
| Cholesterol, ppm | 829 |
| Linoleic Acid, % | 0.88 |
| Linolenic Acid, % | 0.16 |
| Arachidonic Acid, % | 0.11 |
| Omega-3 Fatty Acids, % | 2.00 |
| Total Saturated Fatty Acids, % | 3.16 |
| Total Monounsaturated Fatty Acids, % | 2.75 |
| Polyunsaturated Fatty Acids, % | 2.97 |
| ***Fiber (max), %*** | ***2.6*** |
| Neutral Detergent Fiber[2], % | 4.2 |
| Acid Detergent Fiber[3], % | 3.0 |
| ***Nitrogen-Free Extract (by difference), %*** | ***23.9*** |
| Starch, % | 24.23 |
| Glucose, % | 8.64 |
| Fructose, % | 0.00 |
| Sucrose, % | 0.72 |
| Lactose, % | 0.00 |
| **Total Digestible Nutrients, %** | **56.0** |
| ***Energy (kcal/g)*** [4] | **3.72** |

| From: | kcal | % |
|---|---|---|
| Protein | 1.792 | 48.2 |
| Fat (ether extract) | 0.972 | 26.1 |
| Carbohydrates | 0.957 | 25.7 |

| ***Minerals*** | |
|---|---|
| **Ash, %** | **7.9** |
| Calcium, % | 1.82 |
| Phosphorus, % | 1.30 |
| Phosphorus (available), % | 1.16 |
| Potassium, % | 0.74 |
| Magnesium, % | 0.16 |
| Sulfur, % | 0.29 |
| Sodium, % | 0.21 |
| Chloride, % | 0.32 |
| Fluorine, ppm | 31.3 |
| Iron, ppm | 548 |
| Zinc, ppm | 218 |
| Manganese, ppm | 101 |
| Copper, ppm | 20 |
| Cobalt, ppm | 0.90 |
| Iodine, ppm | 2.12 |
| Chromium, ppm | 1.01 |
| Selenium, ppm | |
| ***Vitamins*** | |
| Carotene, ppm | 0.0 |
| Vitamin A, IU/g | 25 |
| Vitamin D-3 (added), IU/g | 2.4 |
| Vitamin E, IU/kg | 107 |
| Vitamin K (as menadione), ppm | 4.8 |
| Thiamin Hydrochloride, ppm | 38 |
| Riboflavin, ppm | 36.6 |
| Niacin, ppm | 227 |
| Pantothenic Acid, ppm | 103 |
| Folic Acid, ppm | 6.9 |
| Pyridoxine, ppm | 30.53 |
| Biotin, ppm | 0.5 |
| Vitamin B-12, mcg/kg | 200 |
| Choline Chloride, ppm | 2,414 |
| Ascorbic Acid, ppm | 150.2 |

1. Based on the latest ingredient analysis information. Since nutrient composition of natural ingredients varies, analysis will differ accordingly. Nutrients expressed as percent of ration on an As Fed basis except where otherwise indicated. Moisture content is assumed to be 10.0% for the purpose of calculations.
2. NDF = approximately cellulose, hemi-cellulose and lignin.
3. ADF = approximately cellulose and lignin.
4. Energy (kcal/gm) - Sum of decimal fractions of protein, fat and carbohydrate x 4,9,4 kcal/gm respectively.

ISO 9001:2000 certified

TestDiet www.testdiet.com

**Ingredients :** Dehulled Soybean meal, Poultry meal, Fish meal, Ground Wheat, Dextrin, Casein-vitamin free, Fish oil, Flash dried blood meal, Dicalcium phosphate, Powdered cellulose, vitamin and mineral premix, lecithin, menadione, dimethylpyriminidol bisulfite (vitamin K), chiline chloride, brewers dired yeast, calcium pantothenate, mineral premix, ethoxyquin (a preservative), DL-methionine, Vitamin A palmitate.
**Analytical constituants :** Proteins 44.8 %, Fat 10.8 %, Fibers 2.6 %, Nitrogen-Free extract 23.9%, Ashes 7.9 %.

### Cytometry : quantification of bacterial populations across life stages

We counted the number of bacteria per gram of gut weight as follows:

Knowing the initial beads concentrations (1010 beads/µL) and the dilution made with the extract (5 µL plus 500 µL), we computed the concentration (c) of beads in the sample: c=(5*1010)/505=10 beads/µL. Knowing the number of beads counted (N), we computed the volume (v) analyzed during the flow cytometry: v=N/c. Knowing the volume analyzed and the number of bacteria counted, we compute the concentration of bacteria: cb=Nb/v. Knowing the dilution factor for the sample considered (d=20 or d=10 in our experiments) and the total volume of that sample (Vs), we compute the absolute number of bacteria for that sample (B) : B=cb*d*Vs. Finally, we normalized by the mass of the sample to obtain the bacterial load: Q= B/m.

### Bacterial nucleic acids extraction

We used the NucleoSpin® Soil kit (Macherey-Nagel™) to extract genomic DNA (gDNA) from the cell pellet. This cell pellet was initially resuspended in 700 µL of the lysis buffer SL1 and we added 150 µL of the enhancer solution before completing the extraction according to the manufacturer instructions. gDNA was eluted with 50 µL of elution buffer. gDNA concentration was estimated by spectrophotometry with a NanoDrop device and gDNA quality was evaluated with gel electrophoresis on 1% agarose (UltraPure™ Agarose, Life technologies™) gel in 1x TAE buffer stained with SYBR® Safe (Life technologies™).

We extracted RNA by resuspending the cell pellets in 750 µL of TRIzol® (Life technologies™) and transferred the lysates to 2 mL bead beating tubes containing 1g of garnet beads (0.7 mm Garnet Beads, MO BIO Laboratories, Inc.™). We homogenized the cells by bead beating with a Precellys® 24 Dual device (Ozyme) at 5 500 rpm during 30 s. The bead beating step was performed three times and the samples were incubated on ice during 5 min between each step to avoid overheating. Next, we briefly centrifuged the tubes and transferred the supernatants in a new tube to eliminate the beads. We subjected the samples to three cycles of freezing in liquid nitrogen and heating at +70°C for 5 min in a heating block. Next, we centrifuged the tubes at 12 0000g for 10 min at +4°C to remove insoluble materials. We transferred the supernatant to 2 ml Heavy Phase Lock Gel (5 PRIME™) and we added 150 µL of chloroform and mixed by inversion. The Phase Lock Gel were centrifuged at 2 500g for 20 min at 4°C. We then recovered RNA from the aqueous phase by precipitation using 375 µL of isopropanol (SIGMA-ALDRICH) and overnight incubation at -20°C. The RNA pellet was recovered by centrifugation and resuspended in 30 µL of RNase/DNase free water. We estimated RNA quantity by spectrophotometry using a NanoDrop device. RNA quality was evaluated with capillarity gel electrophoresis with RNA Nano chip on an Agilent 2100 Bioanalyzer system (Agilent Technologies™). The capillary gel electrophoresis allowed the observation of bacterial ribosomal RNA from eukaryote tissue by visualizing 16S and 23S prokaryotic ribosomal RNA (Figure M2). We obtained 10-15 µg of total RNA and ~1 µg of DNA per filtered tadpole gut at stage NF 56.

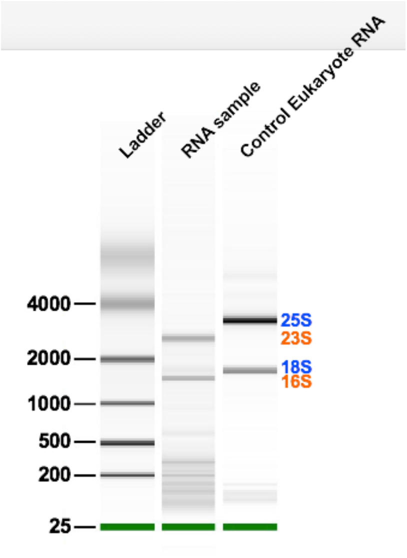


**Figure M2: Microbiome RNA purification from *Xenopus* gut.**
A representative microbiome *Xenopus* tadpole (NF56) gut RNA sample (100 ng) electrophoregram is shown next to a control total Xenopus RNA. This RNA sample was the one used for shotgun metatranscriptomic sequencing.

### DNA extraction from samples fixed in ethanol

We extracted intestinal tracts from *X. tropicalis* fixed in 100% EtOH. Intestines were washed three times in RNase/DNase free water and homogenized in PBS with a potter. We harvested cells by centrifugation at 13 000g for 15 min at room temperature. The pellets were used for gDNA extraction with NucleoSpin® Soil kit (Macherey-Nagel™). gDNA concentration was estimated by spectrophotometry with a NanoDrop device.

### cDNA synthesis

For cDNA synthesis, 1 µg of RNA were mixed with random pentadecamers (2 µL at 150 ng/µL) and RNase/DNase free water (qsp at 19 µL). This mixture was incubated for 5 min at 70°C. Next we added 5x First Buffer from Invitrogen™ (6 µL), 0,1M DTT (1.5 µL), dNTP mix (1.5 µL at 10 mM each), RNase out from Invitrogen™ (1 µL at 40U/µL) and Superscript® III from Invitrogen™ (1 µL at 400U/µL). This mixture was incubated for two to three hours at 46°C. RNA were degraded with RNase H (2 µL at 2U/µL) during 30 min at 37°C and then incubated for 10 min at 70°C to inactivated the RNase H.

### 16S rRNA and 16S rDNA library construction and sequencing

We used universal primers 341F (5'- CCTACGGGNGGCWGCAG -3') and 805R (5'-GACTACHVGGGTATCTAATCC -3') to amplify a ~429bp fragment including the V3-V4 variable regions of the 16S rDNA and 16S rRNA [2]. Illumina adaptors were added in 5' side of both primer, 5'- CTTTCCCTACACGACGCTCTTCCGATCT was added to the 341F2 primer and 5'- GAGTTCAGACGTGTGCTCTTCCGATC to the 805R primer. We used 100ng of gDNA or cDNA diluted at 1/10 (500-1000pg/$\mu$L) in PCR experiments. We assembled PCR reactions in 100 $\mu$l final volume containing 1.7 U of HotMaster™ *Taq* DNA Polymerase (5 PRIME™); 1X HotMaster™ *Taq* buffer (5 PRIME™); 200 nM of each primer; 1 mM dNTP mix; 100 ng of gDNA or 5 $\mu$l of cDNA diluted 1:10X dilution. The following program was used: initial denaturation for 2 min at 95°C, 35 cycles at 95°C for 15s, 47°C for 30s and 68°C for 60s, final extension for 5 min at 68°C. PCR were performed in triplicate and pooled before concentration using a SpeedVac to obtain approximately 55 µL. The PCR products were analyzed by electrophoresis on a 1.5% agarose gel (UltraPure™ Agarose, Life technologies™) in 1x TAE buffer stained with SYBER® Safe (Life technologies™). We observed a single ~ 429 bp amplification product for each sample.

### Metagenome assembly

We used metawrap [3] to control read quality and to filter out reads derived from the nuclear and the mitochondrial *Xenopus tropicalis* genome (assembly version 10 downloaded from www.xenbase.org and mitochondrial genome accession number NC_006839.1) [4]. The initial raw DNA reads data set was obtained from one library of 170 bp insert size using a PE101 strategy and consisted of 47,279,786 high quality paired reads of 100 nt. After metawrap QC we had 47,275,642 reads (99.99%) spanning 4.7 Gbp. The initial raw RNA reads data set consisted of 40,000,000 reads of 100 nt spanning 3.9 Gbp. After QC, the RNA dataset was made of 34,978,866 reads spanning 3.5 Gbp.

We used kraken2 [5] /bracken[6] for taxonomic assignments based on sequence reads with the nucleotidic database PlusPFP (archaea, bacteria, viral, plasmid, human*, UniVec_Core, protozoa, fungi & plant) release 9/19/2020 downloaded from https://benlangmead.github.io/aws-indexes/k2). We used kaiju [7] for taxonomic assignments based on sequence reads with the protein database NCBI BLAST *nr* +euk (103M protein sequences from *nr*: Bacteria, Archaea, Viruses, Fungi and microbial eukaryotes, release 2020-05-25) downloaded from http://kaiju.binf.ku.dk/server. Kraken2 classified 460550 (1.95%) metagenomic reads from the input of 23637821 sequences (4705.09 Mbp).

We used phyloFlash and MATAM to reconstruct near full-length rRNA genes [8].

### Metagenome assembly

We used megahit, metaspades, ray and idbda for metagenome assembly [9–12]. We used the metaQUAST pipeline to compare these assemblies and we mapped back our initial metagenomic and metatranscriptomic reads and also our previously found OTU sequences [13]. We performed binning on contigs longer than 1000 bp using maxbin, concoct, metabat and DAStool [14–17]. We visualized binning results using vizbin [18]. We mapped back the reads to the assembly using BBTools [19].

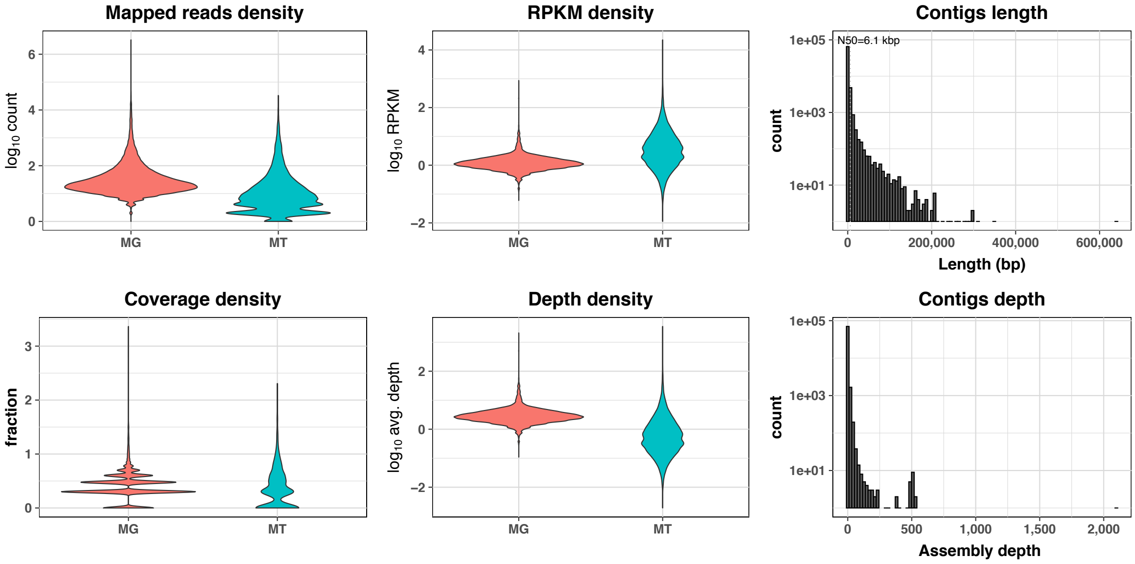


**Figure M3**: Graphical representation of mapping density, coverage and length of metagenomic and metatranscriptomic reads on the metagenomic assembly. Distribution of contigs length for the metagenomic assembly is shown on the top right, and for a trinity assembly on non rRNA reads on the bottom right.

### Metagenome functional annotation

We predicted CDS on the assembled metagenome scaffolds using prokka v.1.14.5 [20]. We used Minpath for metabolic pathway prediction following the strategy and the tools provided in the IMP pipeline scripts as described in https://metagenomics-workshop.readthedocs.io [21, 22]. We mapped KEGG and EC identifiers on the Interactive Pathway Explorer V3 (iPATH3) to obtain a map of the tadpole's gut microbiota metabolic pathway [23, 24][25].
