## Supplementary material for "Gut microbial ecology of Xenopus tadpoles across life stages": Figure_S1

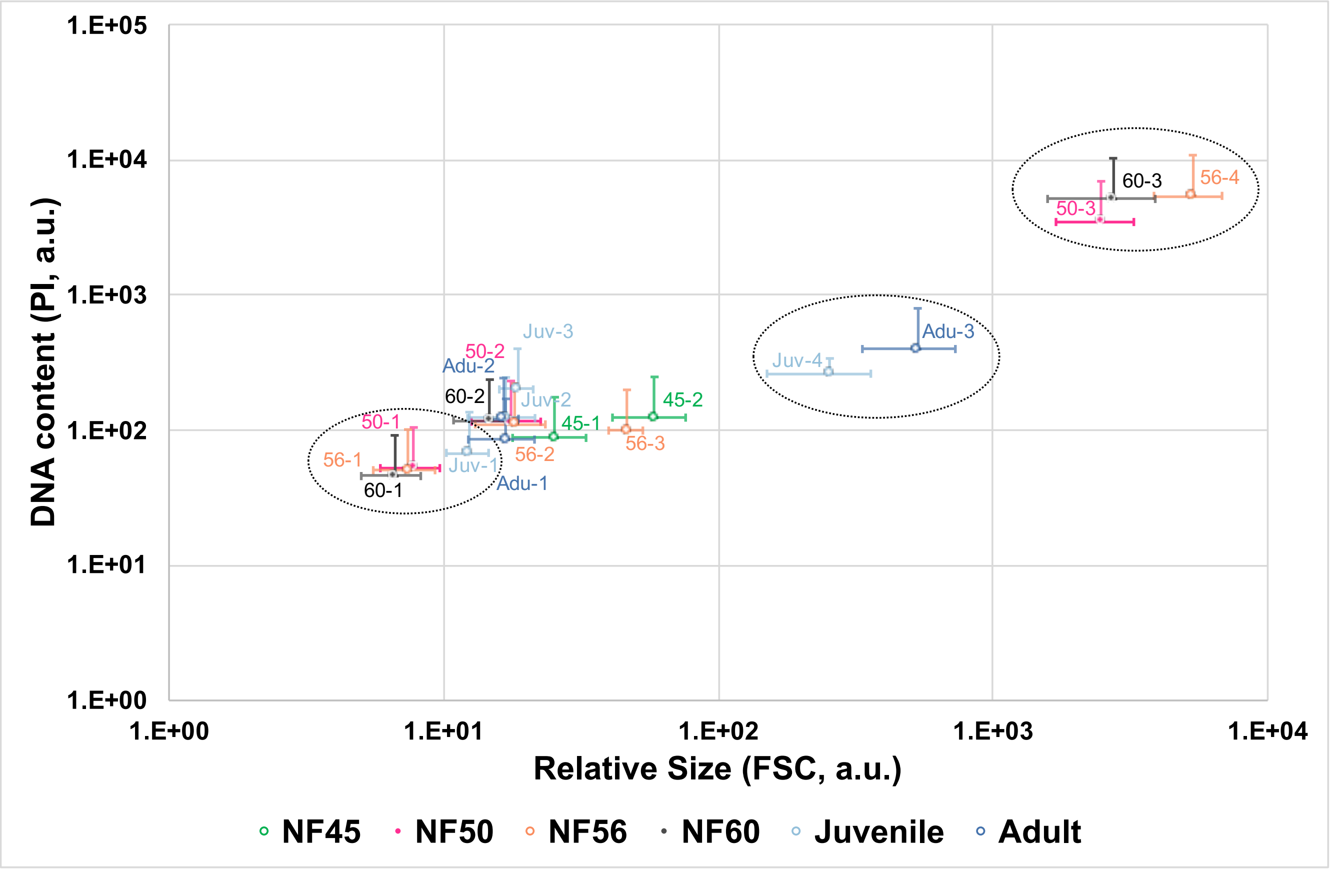


**Figure S1: Microbial cell populations identified by flow cytometry across developmental stages.**
Cytometric profiles obtained at different stages of tadpole's development were used to identify visually bacterial populations. Each population was plotted according to its DNA content and relative size (error bars correspond to the standard error deviation). Vertical axis represents the measurement of fluorescence (propidium iodide - 614/20nm) and the horizontal axis the measurement of relative cell size (forward scatter, in arbitrary units). The bacterial populations clearly corresponding across life stages are grouped by a dotted ellipse.
