## Supplementary material for "Gut microbial ecology of Xenopus tadpoles across life stages": Figure_S2

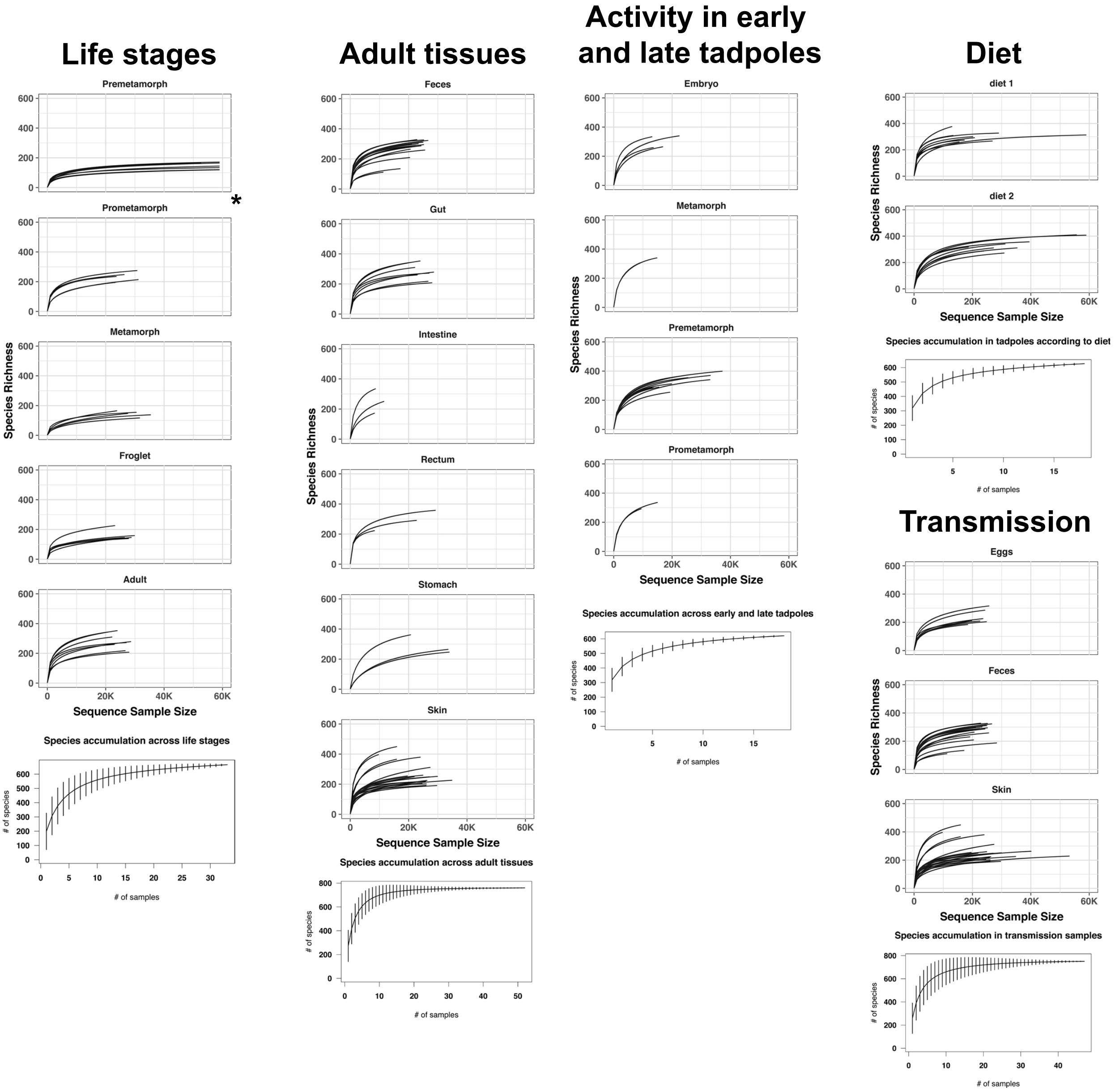


**Figure S2 : Rarefaction and species accumulation curves.**

These graphs present rarefaction curves (top) and species accumulation curves (bottom) for the different samples used in the indicated experiments. * For premetamorph samples, the rarefaction curve is shown only up to 60,000 reads for the sake of consistency between all plots. Species richness corresponds to OTUs number. A detailed analysis is presented at https://npollet.github.io/metatetard/xpall_rarefaction_phyloseq.html
