## Supplementary material for "Gut microbial ecology of Xenopus tadpoles across life stages": Figure_S3

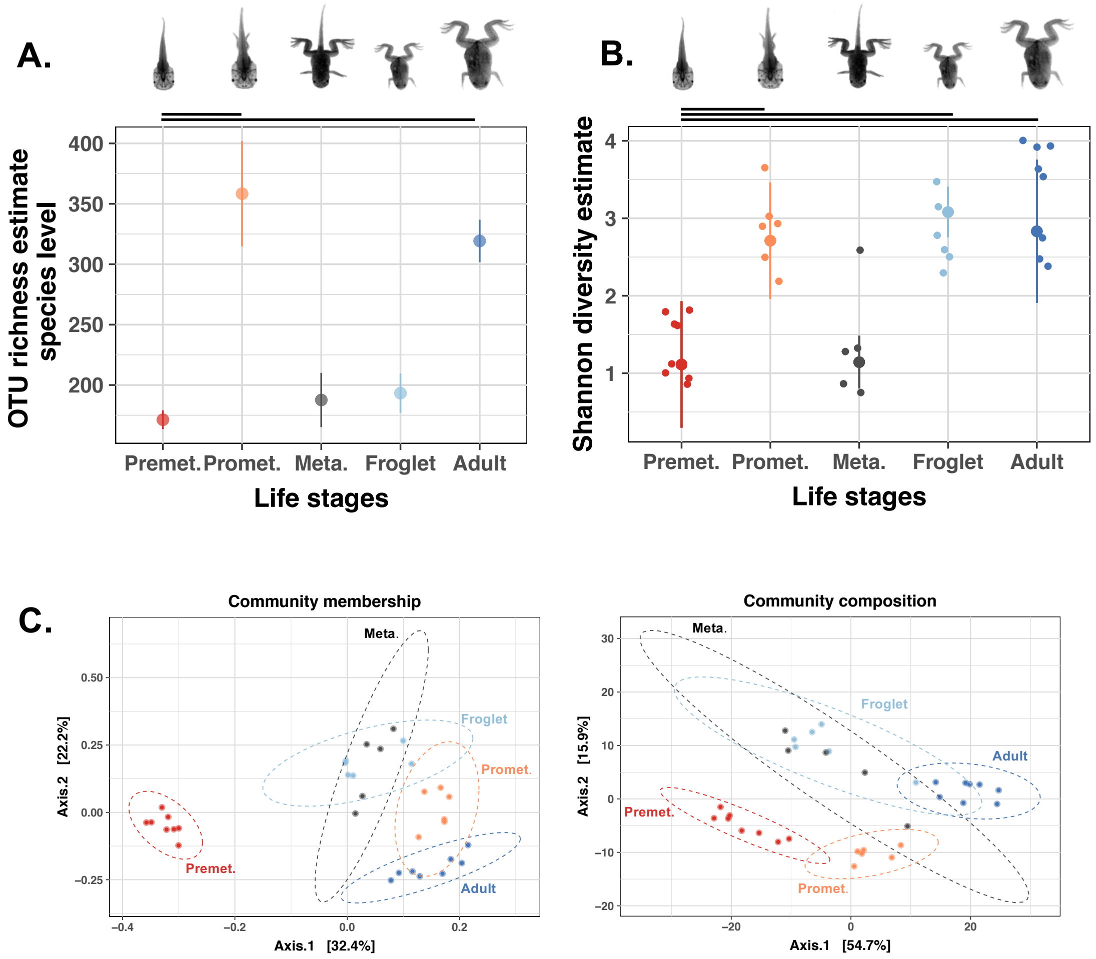


**Figure S3: Bacterial communities in the Xenopus gut across development.**

A. Graphical representation of the bacterial species richness. Each point corresponds to the breakaway richness estimate, error bars represent the measurement error according to the breakaway model. Premet: premetamorphic tadpoles NF54 to NF56 . Promet: prometamorphic tadpoles NF58 to NF61; Meta: Metamorphic tadpole NF 62; Fro: Froglet NF66; Adult: sexually mature adults. Horizontal lines represent significant differences between the life stages connected; p=0.000 both between Premet. and Promet. and between Premet. and Adult. B. Representation of the bacterial species diversity measured using the Shannon index. Each large point corresponds to the divnet Shannon estimate from the samples of the same life stage, error bars represent the measurement error. Each small point corresponds to the divnet Shannon estimate from a given sample. The error bar corresponding to the measurement error can not be seen at this scale. C. Principal coordinate analysis of gut-associated microbial communities during Xenopus development. Community membership was analyzed using unweighted Unifrac distance, and community composition was analyzed using philR distance. Percentages of the explained variations are indicated on both axes.
