## Supplementary material for "Gut microbial ecology of Xenopus tadpoles across life stages": Figure_S4

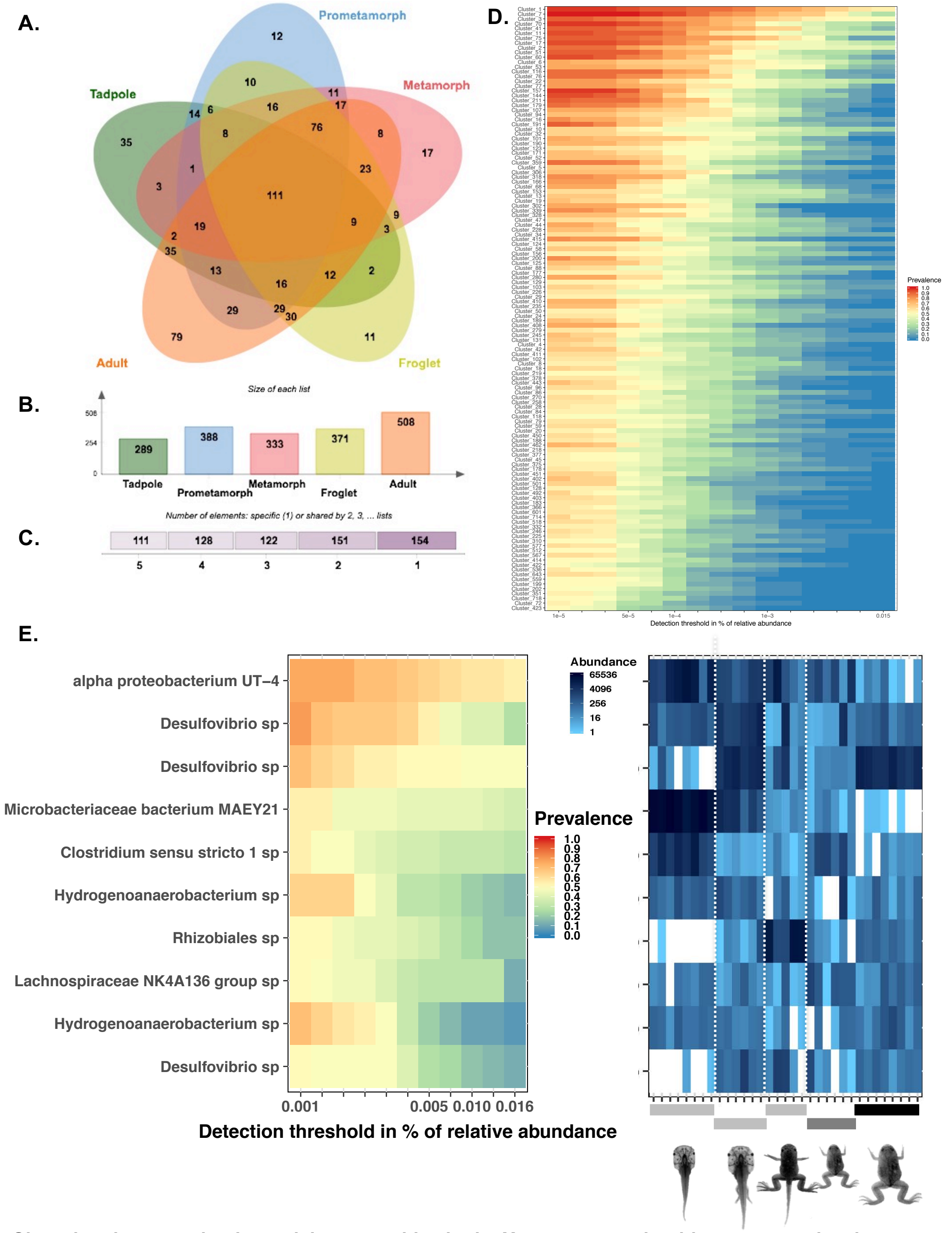


**Figure S4 : Shared and contrasting bacterial communities in the Xenopus gut microbiome across development.**
A. Venn diagram showing the number of shared OTUs between samples of the same developmental stage category. B. Bar plot of the total number of OTUs in each developmental stage category. C. Number of shared and specific OTUs. D. Heatmap plot of the prevalence and abundances for low-abundance OTUs. The detection threshold scale is logarithmic and the metric used was the relative abundance. This analysis was performed on OTUs characterized by a minimal prevalence of 50 % and a minimal abundance of 1e-5 %. E. Prevalence and abundances of the most common and the most abundant OTUs across development. OTUs are identified using their taxonomic affiliation at the family genus or species level according to the resolution available for a given OTU. This analysis was performed on OTUs characterized by a minimal prevalence of 50 % and a minimal abundance of 0.001%. The right panel shows an abundance heatmap for the same OTUs to visualize their abundances in each sample.
