## Supplementary material for "Gut microbial ecology of Xenopus tadpoles across life stages": Figure_S5

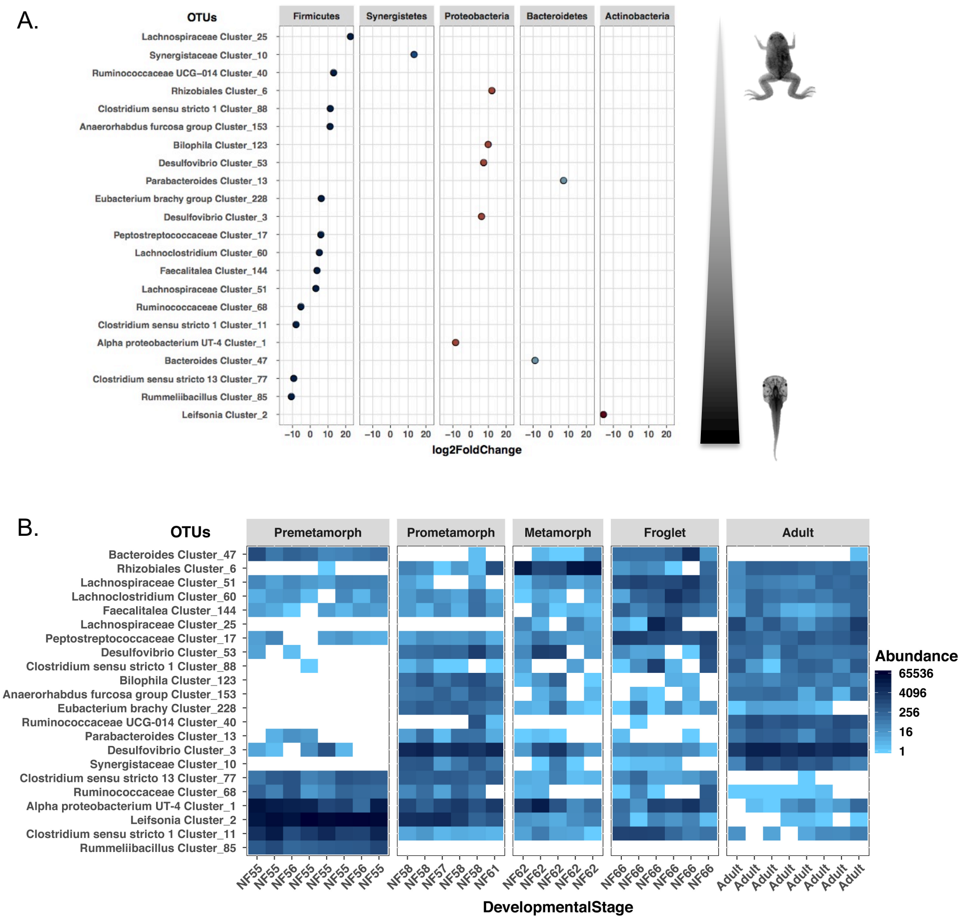


**Figure S5 : Contrasting OTUs across development**

A. Abundance fold changes across development. Contrasting OTUs were identied by filtering OTUs with an abundnce of at least 100 reads across 20% of sample and using DEseq2 with a significance cutoff of 0.01. B. Heatmap representation of contrasting OTUs abundances across development. The heatmap was produced using a principal component analysis based on unifrac distances.
