## Supplementary material for "Gut microbial ecology of Xenopus tadpoles across life stages": Figure_S6

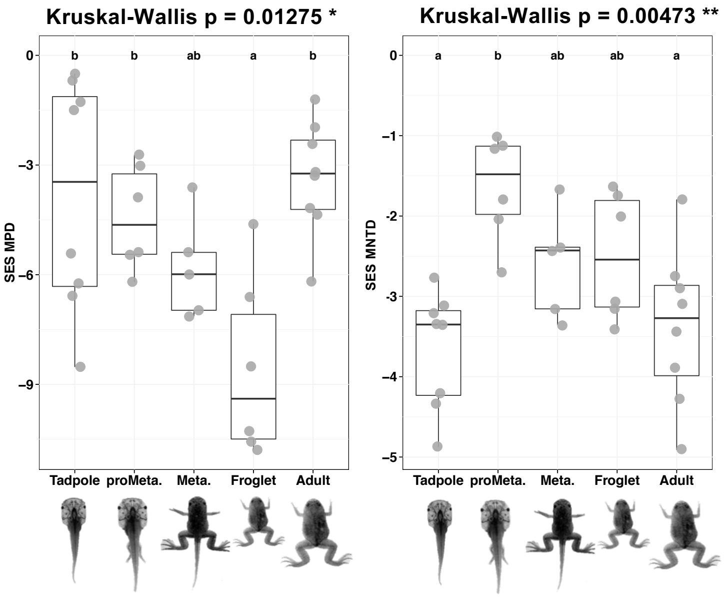


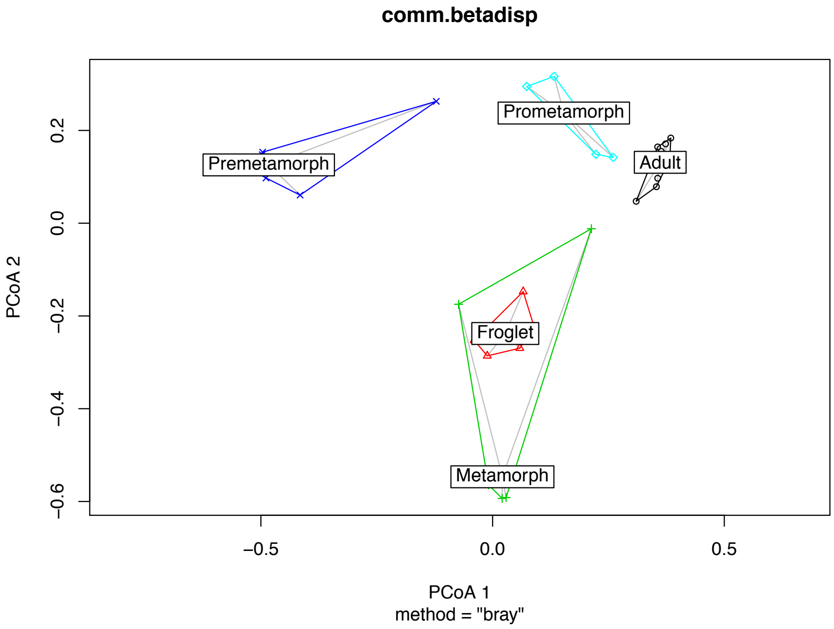


**Figure S6. Variations of the phylogenetic structure of the Xenopus gut microbiota during development.** Standardized effect sizes of the Mean pairwise distance (left panel) and of the mean nearest taxon distance (right panel) measured using Faith' PD index for *X. tropicalis* gut microbial communities across developmental life stages as indicated on the x-axis. Letters a, b and c indicate significant differences at the 0.05 level between developmental stages according to pairwise Dunn's rank sum post-hoc test (see Methods section for details).
