## Supplementary material for "Gut microbial ecology of Xenopus tadpoles across life stages": Figure_S7

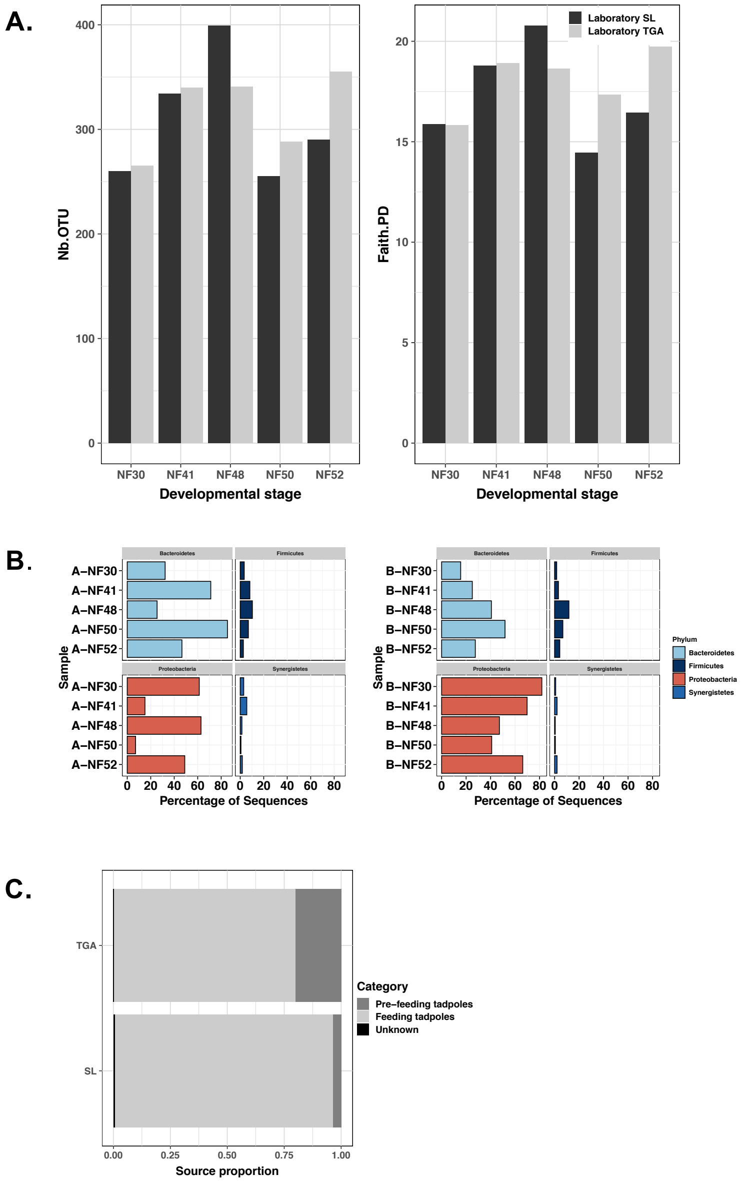


**Figure S7**: **Xenopus tadpole active bacterial communities: alpha-diversity, bacterial phylum diversity and source proportions.**

A. Bar plot representation of OTU richness and phylogenetic diversity in active bacterial communities. Two clutches were followed up across early development, and the light and dark grey bars correspond to the two biological replicate samples. The legend for stages is as follows: NF30 is a post-eclosion tailbud embryonic stage; NF41 is a premetamorphic tadpole non-feeding stage; NF48 is a premetamorphic tadpole feeding stage; NF50 and NF 52 are premetamorphic tadpole feeding stages. B. Bar plot representation of the bacterial phylum diversity. C. Bar plot representation of source proportions in froglet gut microbiome. SL and TGA refer to *X. tropicalis* strain's names : SL for Sierra Leone strain and TGA for a laboratory population of Adiopodoume strain (Ivory Coast) outbred to Uyere strain (Nigeria).
