## Supplementary material for "Gut microbial ecology of Xenopus tadpoles across life stages": Figure_S8

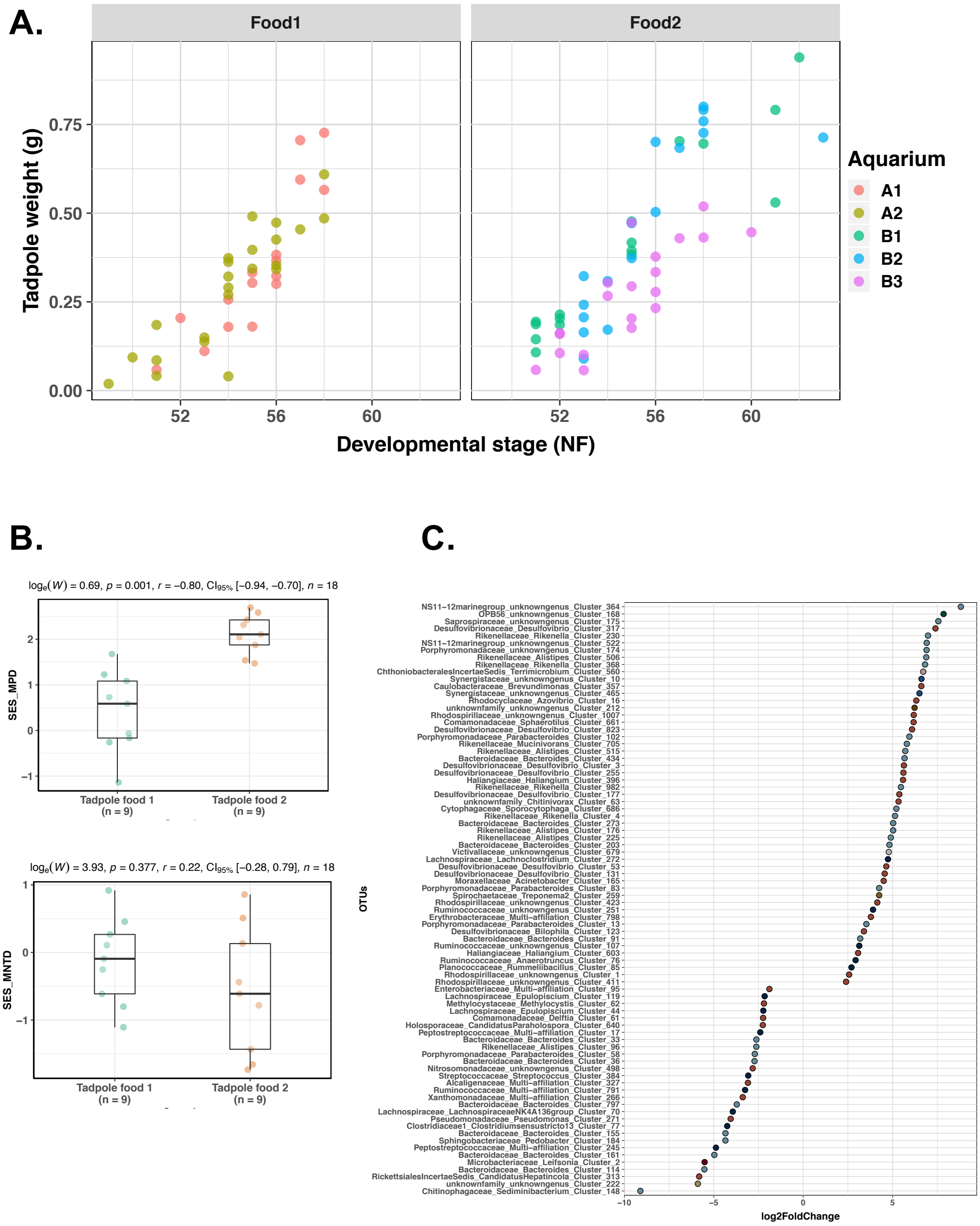


**Figure S8: Impact of the food regime on tadpole growth and development.**

A. Impact of food diet on tadpole growth and development. Tadpoles were fed using either a microplanktonic-based diet (food 1) or a sterile flour (food 2). Their growth was assessed by weighting them, and their developement was monitored by assessing their developmental stage according to the Nieuwkoop and Faber developmental table. The food regime did not change significantly tadpole growth (Kruskal-Wallis chi-squared = 0.99, df = 1, p = 0.32) or development (Kruskal-Wallis chi-squared = 0.34, df = 1, p = 0.56).

B. Impact of the food regime on tadpole's guts microbial communities. These bar plot shows values of the Standardized effect size of the Mean Pairwise Distance (SES MPD) and of the Mean Nearest Taxon Distance (SES MNTD) according to the food regime given to groups of tadpoles.

C. Abundance ratios of the OTUs found significantly different according to a change of diet.
