## Supplementary material for "Gut microbial ecology of Xenopus tadpoles across life stages": Figure_S9: Figure_S9_metaDNA_MATAM_krona.html

Javascript must be enabled to view this page.

magnitude
magnitudeUnassigned

scaffolds.NR.min\_500bp.abd.rdp.fltr.krona

72989.02

13398

13398

13398

13398

13398

13398

59591.02

25602.18

25602.18

25602.18

931.87

931.87

3466.37

3466.37

21203.94

21203.94

2611.83

2611.83

2611.83

2611.83

2611.83

8300.48

8300.48

8300.48

8300.48

8300.48

9179.43

610.83

610.83

610.83

610.83

4785.57

4785.57

4785.57

987

3798.57

3234.7

3234.7

3234.7

3234.7

548.33

548.33

548.33

548.33

4478

4478

4478

4478

4478

9419.1

1750.18

1750.18

1750.18

1750.18

4903.76

4903.76

1034.12

1034.12

1579.33

1579.33

2290.31

2290.31

2765.16

2765.16

2765.16

2765.16
