## Supplementary material for "Gut microbial ecology of Xenopus tadpoles across life stages": Figure_S10: Figure_S10_metaDNA.phyloFlash.html

phyloFlash results summary for library metaDNA


##### image/svg+xml phyloFlash v3.3b3 by Harald Gruber-Vodicka, Elmar Pruesse, Brandon Seah

##### High throughput phylogenetic screening using SSU rRNA gene(s) abundance(s)

Click on report section headers to expand, mouse-over underlined text to see explanations.

### Library name: metaDNA

#### Graphical Summary

Mouseover on panels to expand details.

Mapping identity (%)


30

40

50

60

70

80

90

100


10000

20000

30000

41803
Read-mapping %identity of reads vs. reference database. Lower %identity hits may indicate presence of divergent taxa not represented in the database.

Insert size (bp)


100

200

300

428


1000

2000

3000

4000

5000

6000

7000

8000

9000

10143
Insert sizes for read pairs. Distribution should generally be unimodal; more than one peak may indicate contamination from other libraries.

0.032 % pairs mapped


Mapped pair
Mapped bad pair
Mapped single
Proportion of reads mapped to SSU rRNA database. Typically < 1% for metagenomes, ca. 20% for metatranscriptomes without rRNA depletion or poly-A selection.

Reads assembled


Assembled
Unassembled
Proportion of reads assembled to full-length sequences. High proportion unassembled suggest either assembly failure or high diversity of organisms with low coverage.

Taxonomic summary from reads mapped

Bacteria;Bacteroidota;Bacteroidia;Bacteroidales
11967

Bacteria;Actinobacteriota;Actinobacteria;Micrococcales
5385

Bacteria;Desulfobacterota;Desulfovibrionia;Desulfovibrionales
3781

Bacteria;Firmicutes;Clostridia;Oscillospirales
1937

Bacteria;Verrucomicrobiota;Verrucomicrobiae;Verrucomicrobiales
1809

Bacteria;Firmicutes;Clostridia;Lachnospirales
1456

Bacteria;(Bacteria);(Bacteria);(Bacteria)
1221

Bacteria;Fusobacteriota;Fusobacteriia;Fusobacteriales
891

Bacteria;Firmicutes;Bacilli;Erysipelotrichales
510

Bacteria;Firmicutes;Bacilli;Bacillales
326

Bacteria;Verrucomicrobiota;Verrucomicrobiae;(Verrucomicrobiae)
311

Bacteria;Bacteroidota;Bacteroidia;(Bacteroidia)
304

Bacteria;Firmicutes;(Firmicutes);(Firmicutes)
265

Bacteria;Firmicutes;Clostridia;(Clostridia)
250

Bacteria;Proteobacteria;Gammaproteobacteria;Enterobacterales
246

Bacteria;Proteobacteria;Gammaproteobacteria;Burkholderiales
212

Bacteria;Actinobacteriota;Actinobacteria;(Actinobacteria)
199

Bacteria;Firmicutes;Negativicutes;Acidaminococcales
186

Bacteria;Proteobacteria;Gammaproteobacteria;Legionellales
127

Bacteria;Cyanobacteria;Cyanobacteriia;Chloroplast
122

Bacteria;Firmicutes;Negativicutes;Veillonellales-Selenomonadales
100

Bacteria;Firmicutes;Bacilli;(Bacilli)
93

Bacteria;Firmicutes;Clostridia;Clostridia UCG-014
91

Bacteria;Firmicutes;Clostridia;Peptostreptococcales-Tissierellales
89

Bacteria;Firmicutes;Bacilli;RF39
88

Bacteria;Proteobacteria;Alphaproteobacteria;Rhodobacterales
84

Bacteria;Firmicutes;Bacilli;Lactobacillales
83

Bacteria;Proteobacteria;Alphaproteobacteria;Rhodospirillales
80

Bacteria;Proteobacteria;Alphaproteobacteria;(Alphaproteobacteria)
74

Bacteria;Actinobacteriota;Actinobacteria;Streptomycetales
57

Bacteria;Desulfobacterota;(Desulfobacterota);(Desulfobacterota)
57

Eukaryota;Amorphea;Obazoa;Opisthokonta
53

Bacteria;Proteobacteria;Alphaproteobacteria;Rhizobiales
48

Bacteria;Proteobacteria;(Proteobacteria);(Proteobacteria)
47

Eukaryota;Archaeplastida;Chloroplastida;Charophyta
46

Bacteria;Bacteroidota;Bacteroidia;Flavobacteriales
44

Bacteria;Firmicutes;Bacilli;Staphylococcales
35

Bacteria;Firmicutes;Clostridia;Clostridiales
34

Bacteria;Verrucomicrobiota;Verrucomicrobiae;Pedosphaerales
33

Bacteria;Firmicutes;Negativicutes;(Negativicutes)
29

Bacteria;Actinobacteriota;Actinobacteria;Propionibacteriales
25

Bacteria;Firmicutes;Clostridia;Christensenellales
21

Bacteria;Actinobacteriota;Actinobacteria;Pseudonocardiales
19

Bacteria;Bacteroidota;Bacteroidia;Cytophagales
17

Bacteria;Caldatribacteriota;JS1;terrestrial metagenome
17

Bacteria;Proteobacteria;Alphaproteobacteria;Rickettsiales
16

Bacteria;Firmicutes;Desulfitobacteriia;Desulfitobacteriales
15

Bacteria;Firmicutes;Bacilli;Paenibacillales
12

Bacteria;Actinobacteriota;Actinobacteria;Actinomycetales
12

Bacteria;Verrucomicrobiota;Chlamydiae;Chlamydiales
12

Bacteria;Verrucomicrobiota;Verrucomicrobiae;Chthoniobacterales
11

Bacteria;Firmicutes;Clostridia;Monoglobales
11

Bacteria;Acidobacteriota;Blastocatellia;Blastocatellales
10

Bacteria;Cyanobacteria;Vampirivibrionia;Gastranaerophilales
10

Bacteria;Chloroflexi;Chloroflexia;Chloroflexales
9

Eukaryota;Amorphea;Amoebozoa;Discosea
9

Bacteria;Bacteroidota;(Bacteroidota);(Bacteroidota)
8

Bacteria;Firmicutes;Desulfotomaculia;Desulfotomaculales
8

Bacteria;Proteobacteria;Gammaproteobacteria;Pseudomonadales
8

Bacteria;Verrucomicrobiota;(Verrucomicrobiota);(Verrucomicrobiota)
8

Bacteria;Actinobacteriota;Actinobacteria;Corynebacteriales
7

Bacteria;Proteobacteria;Gammaproteobacteria;(Gammaproteobacteria)
7

Bacteria;Bacteroidota;Bacteroidia;Sphingobacteriales
7

Bacteria;Proteobacteria;Gammaproteobacteria;Xanthomonadales
6

Bacteria;SAR324 clade(Marine group B);uncultured bacterium;(uncultured bacterium)
6

Bacteria;Proteobacteria;Alphaproteobacteria;Sphingomonadales
6

Bacteria;Actinobacteriota;Actinobacteria;Glycomycetales
6

Bacteria;Proteobacteria;Alphaproteobacteria;Defluviicoccales
6

Eukaryota;SAR;Alveolata;Ciliophora
6

Eukaryota;(Eukaryota);(Eukaryota);(Eukaryota)
5

Bacteria;Proteobacteria;Alphaproteobacteria;Paracaedibacterales
5

Bacteria;Proteobacteria;Alphaproteobacteria;uncultured
5

Bacteria;Halanaerobiaeota;Halanaerobiia;Halanaerobiales
5

Bacteria;Firmicutes;Bacilli;RsaHf231
5

Bacteria;Proteobacteria;Alphaproteobacteria;Azospirillales
5

Bacteria;Actinobacteriota;Actinobacteria;Streptosporangiales
4

Bacteria;Proteobacteria;Alphaproteobacteria;Holosporales
4

Bacteria;Verrucomicrobiota;Verrucomicrobiae;Arctic97B-4 marine group
4

Bacteria;Desulfobacterota;Desulfuromonadia;Geobacterales
4

Bacteria;Proteobacteria;Alphaproteobacteria;Zavarziniales
4

Bacteria;Proteobacteria;Alphaproteobacteria;Sneathiellales
3

Bacteria;Proteobacteria;Alphaproteobacteria;Elsterales
3

Bacteria;Desulfobacterota;Desulfobacteria;Desulfobacterales
3

Bacteria;Caldatribacteriota;JS1;uncultured Candidatus Atribacteria bacterium
3

Bacteria;MBNT15;Actinobacteria bacterium RBG\_13\_63\_9;(Actinobacteria bacterium RBG\_13\_63\_9)
3

Bacteria;Actinobacteriota;Thermoleophilia;Solirubrobacterales
3

Bacteria;Desulfobacterota;Desulfuromonadia;PB19
3

Bacteria;Firmicutes;Clostridia;Caldicoprobacterales
3

Bacteria;Firmicutes;Thermoanaerobacteria;Thermoanaerobacterales
3

Bacteria;Firmicutes;Clostridia;Eubacteriales
3

Bacteria;Firmicutes;Syntrophomonadia;Syntrophomonadales
3

Bacteria;Proteobacteria;Alphaproteobacteria;Parvibaculales
3

Bacteria;Firmicutes;Bacilli;Entomoplasmatales
3

Bacteria;Firmicutes;BRH-c20a;uncultured bacterium
2

Bacteria;Firmicutes;Clostridia;uncultured
2

Bacteria;Proteobacteria;Gammaproteobacteria;Methylococcales
2

Bacteria;Proteobacteria;Alphaproteobacteria;Kiloniellales
2

Eukaryota;Discoba;Discicristata;Heterolobosea
2

Bacteria;Firmicutes;Bacilli;Izemoplasmatales
2

Bacteria;Proteobacteria;Alphaproteobacteria;Caulobacterales
2

Bacteria;Verrucomicrobiota;Verrucomicrobiae;Opitutales
2

Bacteria;Firmicutes;Clostridia;Gracilibacteraceae
2

Bacteria;Bacteroidota;Bacteroidia;Chitinophagales
2

Bacteria;Acidobacteriota;Vicinamibacteria;Vicinamibacterales
2

Bacteria;Actinobacteriota;(Actinobacteriota);(Actinobacteriota)
2

Bacteria;Nitrospirota;4-29-1;uncultured bacterium
2

Bacteria;Proteobacteria;Alphaproteobacteria;Kordiimonadales
2

Bacteria;Nitrospirota;Thermodesulfovibrionia;uncultured
1

Bacteria;Acidobacteriota;(Acidobacteriota);(Acidobacteriota)
1

Bacteria;Bacteroidota;Bacteroidia;Bacteroidetes VC2.1 Bac22
1

Bacteria;Proteobacteria;Gammaproteobacteria;Acidithiobacillales
1

Bacteria;Firmicutes;Bacilli;Acholeplasmatales
1

Bacteria;Patescibacteria;Microgenomatia;Candidatus Pacebacteria
1

Bacteria;Proteobacteria;Alphaproteobacteria;AT-s3-44
1

Bacteria;Verrucomicrobiota;Lentisphaeria;Victivallales
1

Eukaryota;SAR;(SAR);(SAR)
1

Bacteria;Caldatribacteriota;JS1;(JS1)
1

Eukaryota;Amorphea;Amoebozoa;Gracilipodida
1

Bacteria;Gemmatimonadota;BD2-11 terrestrial group;uncultured sediment bacterium
1

Bacteria;Proteobacteria;Alphaproteobacteria;Thalassobaculales
1

Bacteria;Chloroflexi;Anaerolineae;Caldilineales
1

Bacteria;Firmicutes;Bacilli;Mycoplasmatales
1

Eukaryota;SAR;Rhizaria;Cercozoa
1

Eukaryota;SAR;Alveolata;(Alveolata)
1

Bacteria;Desulfobacterota;Syntrophia;Syntrophales
1

Bacteria;Dadabacteria;Dadabacteriia;Dadabacteriales
1

Eukaryota;SAR;Alveolata;Protalveolata
1

Bacteria;Deinococcota;Deinococci;Deinococcales
1

Bacteria;Cyanobacteria;Cyanobacteriia;Cyanobacteriales
1

Bacteria;Caldatribacteriota;JS1;uncultured bacterium
1

Eukaryota;Amorphea;Amoebozoa;SAR
1

Bacteria;Proteobacteria;Zetaproteobacteria;Mariprofundales
1

Bacteria;MBNT15;uncultured bacterium;(uncultured bacterium)
1

Bacteria;Synergistota;Synergistia;Synergistales
1

Bacteria;Proteobacteria;Gammaproteobacteria;uncultured
1

Bacteria;Actinobacteriota;Acidimicrobiia;Microtrichales
1

Bacteria;Myxococcota;Polyangia;Haliangiales
1

Bacteria;Actinobacteriota;Actinobacteria;Kineosporiales
1

Bacteria;Firmicutes;Bacilli;Brevibacillales
1

Bacteria;Firmicutes;Incertae Sedis;Gelria
1

Bacteria;Proteobacteria;Alphaproteobacteria;Dstr-E11
1

Bacteria;Proteobacteria;Gammaproteobacteria;PLTA13
1

Bacteria;Firmicutes;Bacilli;Haloplasmatales
1

Bacteria;Nitrospirota;Leptospirillia;Leptospirillales
1

Archaea;Crenarchaeota;Bathyarchaeia;uncultured archaeon
1

Eukaryota;Excavata;Metamonada;Parabasalia
1

Coverage on 18S model


200

300

400

500

600

700

800

900

1000

1100

1200

1300

1400

1500

1600

1700

1822


4

5

6

7

8

9

10

11

12

13

14
Coverage evenness across eukaryotic 18S rRNA gene model from Barrnap, using Nhmmer from random subsample of mapped reads. This helps to detect contamination from tag sequencing libraries (sharp coverage peaks). For the eukaryotic model it is normal to see one or two regions with low coverage because of variable regions in the 18S rRNA gene that are not present in all organisms.

Coverage on 16S model


100

200

300

400

500

600

700

800

900

1000

1100

1200

1300

1400

1507


900

1000

1100

1200

1300

1400

1500

1600

1700

1800

1885
Coverage evenness across prokaryotic 16S rRNA gene model from Barrnap, using Nhmmer from random subsample of mapped reads. This helps to detect contamination from tag sequencing libraries (sharp coverage peaks).


##### Tree of full-length assembled sequences (Click to hide)

Full-length assembled SSU rRNA sequences along with closest hits from SILVA database, in an alignment guide tree produced by MAFFT. This tree helps to visualize the relatedness of sequences in library to known relatives. Colored circles have areas proportional to the number of SSU rRNA reads that map to each respective sequence (re-mapping is done separately for SPAdes, EMIRGE, and trusted-contig full-length sequence sets). Click on the toggle switches to turn them on and off. Additional circle representing proportion of SSU rRNA reads that were not assembled is in lower right corner. Long taxonomy strings of reference sequences are truncated, but the full strings are given in the tables below.

**Color key:** Assembled by SPAdes, Reconstructed by EMIRGE, Extracted from trusted contigs,  Closest-matching reference sequence from SILVA database

Toggle reads mapped:
SPAdes

I

O
EMIRGE

I

O
Reads: 3122
Reads: 1046
Reads: 306
Reads: 2410
Reads: 120
Reads: 1535
Reads: 3020
Reads: 650
Reads: 3427
Reads: 5233
Reads: 116
Reads: 1749
Reads: 1690
Reads: 486
Reads: 133
Reads: 10557
Reads: 388
Reads: 355
Reads: 838
Reads: 5379
Reads: 394
Reads: 2004
Reads: 450
Reads: 4108
Reads: 1658
Reads: 906
Reads: 389
Reads: 540
Reads: 684
Reads: 821
Reads: 133
Reads: 731
Reads: 191
Reads: 1019
Reads: 4274
Reads: 1225
Reads: 1903
Reads: 703
Reads: 962
Reads: 2136
Reads: 8346
Reads: 733
Reads: 565
Reads: 2032
Reads: 1212
Reads: 573
Reads: 490
Unassembled SSU reads: 13162


metaDNA.PFemirge\_66\_0.059048


DQ808599.1.1393 Bacteria;Bacteroidota;Bacteroidia;Bacteroidales;Bacteroidaceae;Bacteroides;uncultured bacterium


DQ799557.1.1402 Bacteria;Bacteroidota;Bacteroidia;Bacteroidales;Bacteroidaceae;Bacteroides;uncultured bacterium


metaDNA.PFemirge\_25\_0.026633


CP012938.2352114.2353639 Bacteria;Bacteroidota;Bacteroidia;Bacteroidales;Bacteroidaceae;Bacteroides;Bacteroides ovatus


metaDNA.PFemirge\_212\_0.007609


LS483487.3478705.3480200 Bacteria;Fusobacteriota;Fusoba ... s;Fusobacteriaceae;Fusobacterium;Fusobacterium ulcerans


metaDNA.PFspades\_9\_9.355891


metaDNA.PFemirge\_83\_0.002874


AJ879783.1.1485 Bacteria;Proteobacteria;Gammaproteobact ... nucleobacter;Polynucleobacter asymbioticus QLW-P1DMWA-1


metaDNA.PFspades\_3\_4.994869


metaDNA.PFemirge\_28\_0.051552


metaDNA.PFemirge\_9\_0.020421


AGTN01029961.1048.2511 Bacteria;Proteobacteria;Alphapro ... netospirillaceae;Magnetospirillum;bioreactor metagenome


FN436123.1.1478 Bacteria;Firmicutes;Clostridia;Oscillospirales;Ruminococcaceae;Angelakisella;uncultured bacterium


metaDNA.PFemirge\_24\_0.075313


metaDNA.PFspades\_1\_21.358624


AY983363.1.1360 Bacteria;Bacteroidota;Bacteroidia;Bacteroidales;Bacteroidaceae;Bacteroides;uncultured bacterium


metaDNA.PFemirge\_633\_0.002690


AJ576387.1.1452 Bacteria;Firmicutes;Bacilli;Erysipelotr ... stridiaceae;Erysipelatoclostridium;uncultured bacterium


metaDNA.PFemirge\_133\_0.040161


metaDNA.PFspades\_2\_6.084291


metaDNA.PFemirge\_193\_0.010396


AP017457.761400.762920 Bacteria;Actinobacteriota;Actino ... obacteriaceae;Aurantimicrobium;Aurantimicrobium minutum


metaDNA.PFemirge\_144\_0.003861


AGZO01000045.56609.58123 Bacteria;Bacteroidota;Bacteroi ... ;Parabacteroides;Parabacteroides goldsteinii CL02T12C30


metaDNA.PFspades\_4\_41.575102


LC094617.1.1418 Bacteria;Actinobacteriota;Actinobacteri ... cteriaceae;Leifsonia;Microbacteriaceae bacterium MAEY21


metaDNA.PFemirge\_255\_0.011580


metaDNA.PFspades\_11\_1.343155


EU776269.1.1279 Bacteria;Firmicutes;Clostridia;Oscillospirales;Hydrogenoanaerobacterium;uncultured bacterium


metaDNA.PFemirge\_13\_0.014234


metaDNA.PFspades\_5\_19.595732


metaDNA.PFemirge\_93\_0.013083


JX105679.1.1392 Bacteria;Bacteroidota;Bacteroidia;Bacteroidales;Bacteroidaceae;Bacteroides;uncultured bacterium


HG934468.2032323.2033833 Bacteria;Bacteroidota;Bacteroi ... dales;Rikenellaceae;Mucinivorans;Mucinivorans hirudinis


metaDNA.PFemirge\_32\_0.047211


metaDNA.PFspades\_12\_1.767901


HM124083.1.1490 Bacteria;Bacteroidota;Bacteroidia;Bacte ... llaceae;Prevotellaceae Ga6A1 group;uncultured bacterium


metaDNA.PFspades\_6\_16.252231


AB298756.1.1471 Bacteria;Firmicutes;Clostridia;Lachnospirales;Lachnospiraceae;Anaerostignum;Anaerotignum aminivorans


LT629973.912339.913840 Bacteria;Verrucomicrobiota;Verru ... es;Akkermansiaceae;Akkermansia;Akkermansia glycaniphila


metaDNA.PFemirge\_6\_0.035316


metaDNA.PFemirge\_127\_0.032601


metaDNA.PFemirge\_124\_0.007919


JQ083789.1.1511 Bacteria;Desulfobacterota;Desulfovibrio ... ales;Desulfovibrionaceae;Bilophila;uncultured bacterium


metaDNA.PFspades\_10\_1.836155


HM630238.1.1413 Bacteria;Bacteroidota;Bacteroidia;Bacteroidales;Rikenellaceae;Rikenella;uncultured bacterium


metaDNA.PFemirge\_169\_0.017424


metaDNA.PFemirge\_8\_0.019814


CBVB010000006.254456.255954 Bacteria;Bacteroidota;Bacte ... Bacteroidaceae;Bacteroides;Bacteroidaceae bacterium MS4


metaDNA.PFspades\_16\_0.786316


KF318250.1.1396 Bacteria;Verrucomicrobiota;Verrucomicro ... iaceae;Akkermansia;uncultured Verrucomicrobia bacterium


CXSE01001446.1371.2879 Bacteria;Bacteroidota;Bacteroidia;Bacteroidales;Bacteroidaceae;Bacteroides;human gut metagenome


metaDNA.PFemirge\_515\_0.013248


AJ867049.1.1500 Bacteria;Desulfobacterota;Desulfovibrio ... les;Desulfovibrionaceae;Bilophila;Bilophila wadsworthia


metaDNA.PFspades\_15\_1.254000


metaDNA.PFemirge\_204\_0.019549


EU158190.1.1510 Bacteria;Firmicutes;Clostridia;Oscillos ... anaerobacterium;Hydrogenoanaerobacterium saccharovorans


metaDNA.PFemirge\_7\_0.099176


AQZZ01000002.9217.10746 Bacteria;Firmicutes;Negativicut ... inococcaceae;Succinispira;Succinispira mobilis DSM 6222


metaDNA.PFemirge\_170\_0.026757


metaDNA.PFemirge\_638\_0.043916


metaDNA.PFspades\_8\_2.715403


metaDNA.PFemirge\_844\_0.021818


DQ793316.1.1393 Bacteria;Bacteroidota;Bacteroidia;Bacteroidales;Tannerellaceae;Parabacteroides;uncultured bacterium


metaDNA.PFspades\_7\_8.330931


AGES01000014.73256.74770 Bacteria;Bacteroidota;Bacteroi ... nerellaceae;Parabacteroides;Parabacteroides sp. HGS0025


LN870298.1.1495 Bacteria;Firmicutes;Clostridia;Oscillospirales;Oscillospiraceae;Intestinimonas;Intestinimonas timonensis


metaDNA.PFemirge\_5\_0.169899


KP231747.1.1515 Bacteria;Desulfobacterota;Desulfovibrio ... vibrionaceae;Desulfovibrio;uncultured Desulfovibrio sp.


metaDNA.PFspades\_13\_3.096195


metaDNA.PFemirge\_239\_0.015009


HG970990.1.1557 Bacteria;Firmicutes;Bacilli;Erysipelotrichales;Erysipelotrichaceae;Breznakia;uncultured bacterium


metaDNA.PFemirge\_424\_0.044579


JN559646.1.1500 Bacteria;Bacteroidota;Bacteroidia;Bacte ... llaceae;Prevotellaceae Ga6A1 group;uncultured bacterium


metaDNA.PFemirge\_36\_0.032254


metaDNA.PFemirge\_423\_0.014054


metaDNA.PFspades\_14\_2.520038


FMDX01000032.25335.26845 Bacteria;Firmicutes;Clostridia ... achnospiraceae;GCA-900066575;uncultured Clostridium sp.


FQ312004.4494006.4495538 Bacteria;Bacteroidota;Bacteroi ... es;Bacteroidaceae;Bacteroides;Bacteroides fragilis 638R


##### Interactive treemap of mapping-based taxonomic read classification (Click to hide)

**Navigation**: Left-click to go down, right-click to go up in taxonomic hierarchy, hover to see counts.

Based on read-mapping hits to reference database, provides an approximate overview of taxonomic composition.

Drawn with Google Visualization API (terms of service)

#### Input parameters

|  |  |
| --- | --- |
| Input command | phyloFlash.pl -lib metaDNA -read1 READ\_QC/metatetard\_DNA/final\_pure\_reads\_1.fastq -read2 READ\_QC/metatetard\_DNA/final\_pure\_reads\_2.fastq -CPUs 24 -everything |
| Forward read file | READ\_QC/metatetard\_DNA/final\_pure\_reads\_1.fastq |
| Reverse read file | READ\_QC/metatetard\_DNA/final\_pure\_reads\_2.fastq |
| Minimum mapping identity | 70% |
| Working folder | /home/pollet/metatetard |
| Database used | /home/anaconda/138.1 |

#### Results

##### Mapping statistics

|  |  |
| --- | --- |
| Input PE-reads | 23637821 |
| Mapped SSU read pairs | 7559 |
| Mapping ratio | 0.032% |
| Fraction assembled | 79.254% |
| Detected median insert size | 154 |
| Used insert size | 220 |
| Insert size standard deviation | 22 |

##### Output files

|  |  |
| --- | --- |
| FASTA file of alignment of all full-length sequences | metaDNA.SSU.collection.alignment.fasta |
| FASTA file of all full-length sequences and their closest database hits | metaDNA.SSU.collection.fasta |
| Newick guide tree from MAFFT alignment of all full-length sequences and closest database hits | metaDNA.SSU.collection.fasta.tree |
| SVG graphic of guide tree from MAFFT alignment of all full-length sequences and closest database hits | metaDNA.SSU.collection.fasta.tree.svg |
| FASTA file of full-length SSU sequences | metaDNA.all.final.fasta |
| Raw BBmap SAM file of initial read mapping to SSU rRNA database | metaDNA.bbmap.sam |
| Log file from EMIRGE sequence reconstruction | metaDNA.emirge.out |
| Reads (fwd) mapping to SSU rRNA database | metaDNA.final\_pure\_reads\_1.fastq.SSU.1.fq |
| Reads (rev) mapping to SSU rRNA database | metaDNA.final\_pure\_reads\_1.fastq.SSU.2.fq |
| SAM file of initial read mapping to SSU rRNA database | metaDNA.final\_pure\_reads\_1.fastq.SSU.sam |
| Mapping identity histogram from BBmap | metaDNA.idhistogram |
| Insert size histogram from BBmap | metaDNA.inserthistogram |
| phyloFlash report in plain text | metaDNA.phyloFlash |
| NTU abundances (truncated to requested taxonomic level) from initial mapping, in CSV format | metaDNA.phyloFlash.NTUabundance.csv |
| SVG graphic of taxonomic composition from initial read mapping | metaDNA.phyloFlash.NTUabundance.csv.svg |
| NTU abundances (untruncated) from initial mapping, in CSV format | metaDNA.phyloFlash.NTUfull\_abundance.csv |
| Taxonomic classification of full-length sequences, in CSV format | metaDNA.phyloFlash.extractedSSUclassifications.csv |
| phyloFlash report in CSV format | metaDNA.phyloFlash.report.csv |
| Taxonomic composition of unassembled SSU reads in CSV format | metaDNA.phyloFlash.unassembled.NTUabundance.csv |
| Log file from SPAdes assembler | metaDNA.spades.out |

##### Taxonomic affiliation of SSU rRNA reads in library

Approximate overview of taxonomic composition of ALL reads, based on mapping hits to SILVA SSU rRNA database using BBmap.

|  |  |
| --- | --- |
| NTUs observed once | 39 |
| NTUs observed twice | 14 |
| NTUs observed three or more times | 93 |
| NTU Chao1 richness estimate | 147.321 |

Taxonomy summarized at level 4. Only displaying taxa with > 3 reads mapped.

| Taxon | Reads |
| --- | --- |
| Bacteria;Bacteroidota;Bacteroidia;Bacteroidales | 11967 |
| Bacteria;Actinobacteriota;Actinobacteria;Micrococcales | 5385 |
| Bacteria;Desulfobacterota;Desulfovibrionia;Desulfovibrionales | 3781 |
| Bacteria;Firmicutes;Clostridia;Oscillospirales | 1937 |
| Bacteria;Verrucomicrobiota;Verrucomicrobiae;Verrucomicrobiales | 1809 |
| Bacteria;Firmicutes;Clostridia;Lachnospirales | 1456 |
| Bacteria;(Bacteria);(Bacteria);(Bacteria) | 1221 |
| Bacteria;Fusobacteriota;Fusobacteriia;Fusobacteriales | 891 |
| Bacteria;Firmicutes;Bacilli;Erysipelotrichales | 510 |
| Bacteria;Firmicutes;Bacilli;Bacillales | 326 |
| Bacteria;Verrucomicrobiota;Verrucomicrobiae;(Verrucomicrobiae) | 311 |
| Bacteria;Bacteroidota;Bacteroidia;(Bacteroidia) | 304 |
| Bacteria;Firmicutes;(Firmicutes);(Firmicutes) | 265 |
| Bacteria;Firmicutes;Clostridia;(Clostridia) | 250 |
| Bacteria;Proteobacteria;Gammaproteobacteria;Enterobacterales | 246 |
| Bacteria;Proteobacteria;Gammaproteobacteria;Burkholderiales | 212 |
| Bacteria;Actinobacteriota;Actinobacteria;(Actinobacteria) | 199 |
| Bacteria;Firmicutes;Negativicutes;Acidaminococcales | 186 |
| Bacteria;Proteobacteria;Gammaproteobacteria;Legionellales | 127 |
| Bacteria;Cyanobacteria;Cyanobacteriia;Chloroplast | 122 |
| Bacteria;Firmicutes;Negativicutes;Veillonellales-Selenomonadales | 100 |
| Bacteria;Firmicutes;Bacilli;(Bacilli) | 93 |
| Bacteria;Firmicutes;Clostridia;Clostridia UCG-014 | 91 |
| Bacteria;Firmicutes;Clostridia;Peptostreptococcales-Tissierellales | 89 |
| Bacteria;Firmicutes;Bacilli;RF39 | 88 |
| Bacteria;Proteobacteria;Alphaproteobacteria;Rhodobacterales | 84 |
| Bacteria;Firmicutes;Bacilli;Lactobacillales | 83 |
| Bacteria;Proteobacteria;Alphaproteobacteria;Rhodospirillales | 80 |
| Bacteria;Proteobacteria;Alphaproteobacteria;(Alphaproteobacteria) | 74 |
| Bacteria;Actinobacteriota;Actinobacteria;Streptomycetales | 57 |
| Bacteria;Desulfobacterota;(Desulfobacterota);(Desulfobacterota) | 57 |
| Eukaryota;Amorphea;Obazoa;Opisthokonta | 53 |
| Bacteria;Proteobacteria;Alphaproteobacteria;Rhizobiales | 48 |
| Bacteria;Proteobacteria;(Proteobacteria);(Proteobacteria) | 47 |
| Eukaryota;Archaeplastida;Chloroplastida;Charophyta | 46 |
| Bacteria;Bacteroidota;Bacteroidia;Flavobacteriales | 44 |
| Bacteria;Firmicutes;Bacilli;Staphylococcales | 35 |
| Bacteria;Firmicutes;Clostridia;Clostridiales | 34 |
| Bacteria;Verrucomicrobiota;Verrucomicrobiae;Pedosphaerales | 33 |
| Bacteria;Firmicutes;Negativicutes;(Negativicutes) | 29 |
| Bacteria;Actinobacteriota;Actinobacteria;Propionibacteriales | 25 |
| Bacteria;Firmicutes;Clostridia;Christensenellales | 21 |
| Bacteria;Actinobacteriota;Actinobacteria;Pseudonocardiales | 19 |
| Bacteria;Caldatribacteriota;JS1;terrestrial metagenome | 17 |
| Bacteria;Bacteroidota;Bacteroidia;Cytophagales | 17 |
| Bacteria;Proteobacteria;Alphaproteobacteria;Rickettsiales | 16 |
| Bacteria;Firmicutes;Desulfitobacteriia;Desulfitobacteriales | 15 |
| Bacteria;Actinobacteriota;Actinobacteria;Actinomycetales | 12 |
| Bacteria;Verrucomicrobiota;Chlamydiae;Chlamydiales | 12 |
| Bacteria;Firmicutes;Bacilli;Paenibacillales | 12 |
| Bacteria;Firmicutes;Clostridia;Monoglobales | 11 |
| Bacteria;Verrucomicrobiota;Verrucomicrobiae;Chthoniobacterales | 11 |
| Bacteria;Acidobacteriota;Blastocatellia;Blastocatellales | 10 |
| Bacteria;Cyanobacteria;Vampirivibrionia;Gastranaerophilales | 10 |
| Bacteria;Chloroflexi;Chloroflexia;Chloroflexales | 9 |
| Eukaryota;Amorphea;Amoebozoa;Discosea | 9 |
| Bacteria;Firmicutes;Desulfotomaculia;Desulfotomaculales | 8 |
| Bacteria;Verrucomicrobiota;(Verrucomicrobiota);(Verrucomicrobiota) | 8 |
| Bacteria;Proteobacteria;Gammaproteobacteria;Pseudomonadales | 8 |
| Bacteria;Bacteroidota;(Bacteroidota);(Bacteroidota) | 8 |
| Bacteria;Proteobacteria;Gammaproteobacteria;(Gammaproteobacteria) | 7 |
| Bacteria;Actinobacteriota;Actinobacteria;Corynebacteriales | 7 |
| Bacteria;Bacteroidota;Bacteroidia;Sphingobacteriales | 7 |
| Bacteria;SAR324 clade(Marine group B);uncultured bacterium;(uncultured bacterium) | 6 |
| Eukaryota;SAR;Alveolata;Ciliophora | 6 |
| Bacteria;Proteobacteria;Alphaproteobacteria;Sphingomonadales | 6 |
| Bacteria;Proteobacteria;Alphaproteobacteria;Defluviicoccales | 6 |
| Bacteria;Actinobacteriota;Actinobacteria;Glycomycetales | 6 |
| Bacteria;Proteobacteria;Gammaproteobacteria;Xanthomonadales | 6 |
| Bacteria;Firmicutes;Bacilli;RsaHf231 | 5 |
| Bacteria;Halanaerobiaeota;Halanaerobiia;Halanaerobiales | 5 |
| Bacteria;Proteobacteria;Alphaproteobacteria;Paracaedibacterales | 5 |
| Bacteria;Proteobacteria;Alphaproteobacteria;uncultured | 5 |
| Bacteria;Proteobacteria;Alphaproteobacteria;Azospirillales | 5 |
| Eukaryota;(Eukaryota);(Eukaryota);(Eukaryota) | 5 |
| Bacteria;Verrucomicrobiota;Verrucomicrobiae;Arctic97B-4 marine group | 4 |
| Bacteria;Actinobacteriota;Actinobacteria;Streptosporangiales | 4 |
| Bacteria;Proteobacteria;Alphaproteobacteria;Holosporales | 4 |
| Bacteria;Proteobacteria;Alphaproteobacteria;Zavarziniales | 4 |
| Bacteria;Desulfobacterota;Desulfuromonadia;Geobacterales | 4 |

##### SSU rRNA assembly-based taxa

Full-length SSU rRNA sequences assembled by SPAdes, matched to SILVA database with Vsearch.

| OTU | Mapped | Cov | DB hit | Taxonomy | % ID | Alnlen | Evalue |
| --- | --- | --- | --- | --- | --- | --- | --- |
| metaDNA.PFspades\_4 | 10557 | 41.575102 | LC094617.1.1418 | Bacteria;Actinobacteriota;Actinobacteria;Micrococcales;Microbacteriaceae;Leifsonia;Microbacteriaceae bacterium MAEY21 | 99.3 | 1418 | -1 |
| metaDNA.PFspades\_5 | 5379 | 19.595732 | HM124083.1.1490 | Bacteria;Bacteroidota;Bacteroidia;Bacteroidales;Prevotellaceae;Prevotellaceae Ga6A1 group;uncultured bacterium | 88.5 | 1493 | -1 |
| metaDNA.PFspades\_1 | 5233 | 21.358624 | KP231747.1.1515 | Bacteria;Desulfobacterota;Desulfovibrionia;Desulfovibrionales;Desulfovibrionaceae;Desulfovibrio;uncultured Desulfovibrio sp. | 96.7 | 1518 | -1 |
| metaDNA.PFspades\_6 | 4108 | 16.252231 | KF318250.1.1396 | Bacteria;Verrucomicrobiota;Verrucomicrobiae;Verrucomicrobiales;Akkermansiaceae;Akkermansia;uncultured Verrucomicrobia bacterium | 91.8 | 1398 | -1 |
| metaDNA.PFspades\_9 | 2410 | 9.355891 | AGES01000014.73256.74770 | Bacteria;Bacteroidota;Bacteroidia;Bacteroidales;Tannerellaceae;Parabacteroides;Parabacteroides sp. HGS0025 | 97.8 | 1508 | -1 |
| metaDNA.PFspades\_7 | 2136 | 8.330931 | HG934468.2032323.2033833 | Bacteria;Bacteroidota;Bacteroidia;Bacteroidales;Rikenellaceae;Mucinivorans;Mucinivorans hirudinis | 91.1 | 1509 | -1 |
| metaDNA.PFspades\_2 | 1690 | 6.084291 | LS483487.3478705.3480200 | Bacteria;Fusobacteriota;Fusobacteriia;Fusobacteriales;Fusobacteriaceae;Fusobacterium;Fusobacterium ulcerans | 100.0 | 1488 | -1 |
| metaDNA.PFspades\_3 | 1535 | 4.994869 | EU158190.1.1510 | Bacteria;Firmicutes;Clostridia;Oscillospirales;Hydrogenoanaerobacterium;Hydrogenoanaerobacterium saccharovorans | 93.5 | 1407 | -1 |
| metaDNA.PFspades\_13 | 733 | 3.096195 | AB298756.1.1471 | Bacteria;Firmicutes;Clostridia;Lachnospirales;Lachnospiraceae;Anaerostignum;Anaerotignum aminivorans | 95.1 | 1354 | -1 |
| metaDNA.PFspades\_8 | 703 | 2.715403 | HG970990.1.1557 | Bacteria;Firmicutes;Bacilli;Erysipelotrichales;Erysipelotrichaceae;Breznakia;uncultured bacterium | 92.6 | 1522 | -1 |
| metaDNA.PFspades\_10 | 540 | 1.836155 | LN870298.1.1495 | Bacteria;Firmicutes;Clostridia;Oscillospirales;Oscillospiraceae;Intestinimonas;Intestinimonas timonensis | 94.5 | 1496 | -1 |
| metaDNA.PFspades\_14 | 490 | 2.520038 | FMDX01000032.25335.26845 | Bacteria;Firmicutes;Clostridia;Lachnospirales;Lachnospiraceae;GCA-900066575;uncultured Clostridium sp. | 93.6 | 1027 | -1 |
| metaDNA.PFspades\_12 | 450 | 1.767901 | AGTN01029961.1048.2511 | Bacteria;Proteobacteria;Alphaproteobacteria;Rhodospirillales;Magnetospirillaceae;Magnetospirillum;bioreactor metagenome | 93.2 | 1466 | -1 |
| metaDNA.PFspades\_11 | 355 | 1.343155 | AQZZ01000002.9217.10746 | Bacteria;Firmicutes;Negativicutes;Acidaminococcales;Acidaminococcaceae;Succinispira;Succinispira mobilis DSM 6222 | 93.6 | 1530 | -1 |
| metaDNA.PFspades\_15 | 191 | 1.254000 | AJ879783.1.1485 | Bacteria;Proteobacteria;Gammaproteobacteria;Burkholderiales;Burkholderiaceae;Polynucleobacter;Polynucleobacter asymbioticus QLW-P1DMWA-1 | 100.0 | 1004 | -1 |
| metaDNA.PFspades\_16 | 133 | 0.786316 | AJ576387.1.1452 | Bacteria;Firmicutes;Bacilli;Erysipelotrichales;Erysipelatoclostridiaceae;Erysipelatoclostridium;uncultured bacterium | 94.4 | 943 | -1 |


##### SSU rRNA reconstruction-based taxa

Full-length SSU rRNA seqeunces reconstructed by EMIRGE, matched to SILVA database by Vsearch.

| OTU | Mapped | Ratio | DB hit | Taxonomy | % ID | Alnlen | Evalue |
| --- | --- | --- | --- | --- | --- | --- | --- |
| metaDNA.PFemirge\_5 | 8346 | 0.169899 | LC094617.1.1418 | Bacteria;Actinobacteriota;Actinobacteria;Micrococcales;Microbacteriaceae;Leifsonia;Microbacteriaceae bacterium MAEY21 | 99.2 | 1419 | -1 |
| metaDNA.PFemirge\_7 | 4274 | 0.099176 | KP231747.1.1515 | Bacteria;Desulfobacterota;Desulfovibrionia;Desulfovibrionales;Desulfovibrionaceae;Desulfovibrio;uncultured Desulfovibrio sp. | 96.4 | 1485 | -1 |
| metaDNA.PFemirge\_24 | 3427 | 0.075313 | KF318250.1.1396 | Bacteria;Verrucomicrobiota;Verrucomicrobiae;Verrucomicrobiales;Akkermansiaceae;Akkermansia;uncultured Verrucomicrobia bacterium | 92.5 | 1355 | -1 |
| metaDNA.PFemirge\_66 | 3122 | 0.059048 | CBVB010000006.254456.255954 | Bacteria;Bacteroidota;Bacteroidia;Bacteroidales;Bacteroidaceae;Bacteroides;Bacteroidaceae bacterium MS4 | 96.9 | 1497 | -1 |
| metaDNA.PFemirge\_28 | 3020 | 0.051552 | JN559646.1.1500 | Bacteria;Bacteroidota;Bacteroidia;Bacteroidales;Prevotellaceae;Prevotellaceae Ga6A1 group;uncultured bacterium | 89.2 | 1494 | -1 |
| metaDNA.PFemirge\_424 | 2032 | 0.044579 | JN559646.1.1500 | Bacteria;Bacteroidota;Bacteroidia;Bacteroidales;Prevotellaceae;Prevotellaceae Ga6A1 group;uncultured bacterium | 89.1 | 1426 | -1 |
| metaDNA.PFemirge\_32 | 2004 | 0.047211 | AP017457.761400.762920 | Bacteria;Actinobacteriota;Actinobacteria;Micrococcales;Microbacteriaceae;Aurantimicrobium;Aurantimicrobium minutum | 94.3 | 1482 | -1 |
| metaDNA.PFemirge\_638 | 1903 | 0.043916 | HM630238.1.1413 | Bacteria;Bacteroidota;Bacteroidia;Bacteroidales;Rikenellaceae;Rikenella;uncultured bacterium | 92.2 | 1377 | -1 |
| metaDNA.PFemirge\_133 | 1749 | 0.040161 | AGES01000014.73256.74770 | Bacteria;Bacteroidota;Bacteroidia;Bacteroidales;Tannerellaceae;Parabacteroides;Parabacteroides sp. HGS0025 | 98.3 | 1498 | -1 |
| metaDNA.PFemirge\_6 | 1658 | 0.035316 | LS483487.3478705.3480200 | Bacteria;Fusobacteriota;Fusobacteriia;Fusobacteriales;Fusobacteriaceae;Fusobacterium;Fusobacterium ulcerans | 100.0 | 1482 | -1 |
| metaDNA.PFemirge\_170 | 1225 | 0.026757 | KP231747.1.1515 | Bacteria;Desulfobacterota;Desulfovibrionia;Desulfovibrionales;Desulfovibrionaceae;Desulfovibrio;uncultured Desulfovibrio sp. | 94.9 | 1366 | -1 |
| metaDNA.PFemirge\_36 | 1212 | 0.032254 | AY983363.1.1360 | Bacteria;Bacteroidota;Bacteroidia;Bacteroidales;Bacteroidaceae;Bacteroides;uncultured bacterium | 96.5 | 1357 | -1 |
| metaDNA.PFemirge\_25 | 1046 | 0.026633 | HQ681866.1.1451 | Bacteria;Bacteroidota;Bacteroidia;Bacteroidales;Bacteroidaceae;Bacteroides;uncultured bacterium | 95.5 | 1379 | -1 |
| metaDNA.PFemirge\_204 | 1019 | 0.019549 | CXSE01001446.1371.2879 | Bacteria;Bacteroidota;Bacteroidia;Bacteroidales;Bacteroidaceae;Bacteroides;human gut metagenome | 94.0 | 1263 | -1 |
| metaDNA.PFemirge\_844 | 962 | 0.021818 | CP012938.3352663.3354184 | Bacteria;Bacteroidota;Bacteroidia;Bacteroidales;Bacteroidaceae;Bacteroides;Bacteroides ovatus | 93.0 | 1404 | -1 |
| metaDNA.PFemirge\_127 | 906 | 0.032601 | FN436123.1.1478 | Bacteria;Firmicutes;Clostridia;Oscillospirales;Ruminococcaceae;Angelakisella;uncultured bacterium | 94.5 | 1253 | -1 |
| metaDNA.PFemirge\_13 | 838 | 0.014234 | FQ312004.4494006.4495538 | Bacteria;Bacteroidota;Bacteroidia;Bacteroidales;Bacteroidaceae;Bacteroides;Bacteroides fragilis 638R | 91.1 | 1389 | -1 |
| metaDNA.PFemirge\_8 | 821 | 0.019814 | GQ449088.1.1384 | Bacteria;Bacteroidota;Bacteroidia;Bacteroidales;Bacteroidaceae;Bacteroides;uncultured bacterium | 94.3 | 1378 | -1 |
| metaDNA.PFemirge\_515 | 731 | 0.013248 | KP231747.1.1515 | Bacteria;Desulfobacterota;Desulfovibrionia;Desulfovibrionales;Desulfovibrionaceae;Desulfovibrio;uncultured Desulfovibrio sp. | 92.9 | 1475 | -1 |
| metaDNA.PFemirge\_169 | 684 | 0.017424 | DQ799557.1.1402 | Bacteria;Bacteroidota;Bacteroidia;Bacteroidales;Bacteroidaceae;Bacteroides;uncultured bacterium | 95.6 | 1388 | -1 |
| metaDNA.PFemirge\_9 | 650 | 0.020421 | EU776269.1.1279 | Bacteria;Firmicutes;Clostridia;Oscillospirales;Hydrogenoanaerobacterium;uncultured bacterium | 92.7 | 1170 | -1 |
| metaDNA.PFemirge\_423 | 573 | 0.014054 | HQ803014.1.1447 | Bacteria;Bacteroidota;Bacteroidia;Bacteroidales;Bacteroidaceae;Bacteroides;uncultured organism | 94.5 | 1320 | -1 |
| metaDNA.PFemirge\_239 | 565 | 0.015009 | HG970990.1.1557 | Bacteria;Firmicutes;Bacilli;Erysipelotrichales;Erysipelotrichaceae;Breznakia;uncultured bacterium | 94.8 | 1280 | -1 |
| metaDNA.PFemirge\_193 | 486 | 0.010396 | AGZO01000045.56609.58123 | Bacteria;Bacteroidota;Bacteroidia;Bacteroidales;Tannerellaceae;Parabacteroides;Parabacteroides goldsteinii CL02T12C30 | 95.0 | 1420 | -1 |
| metaDNA.PFemirge\_93 | 394 | 0.013083 | JX105679.1.1392 | Bacteria;Bacteroidota;Bacteroidia;Bacteroidales;Bacteroidaceae;Bacteroides;uncultured bacterium | 95.6 | 1223 | -1 |
| metaDNA.PFemirge\_124 | 389 | 0.007919 | AJ867049.1.1500 | Bacteria;Desulfobacterota;Desulfovibrionia;Desulfovibrionales;Desulfovibrionaceae;Bilophila;Bilophila wadsworthia | 94.1 | 1501 | -1 |
| metaDNA.PFemirge\_255 | 388 | 0.011580 | DQ808599.1.1393 | Bacteria;Bacteroidota;Bacteroidia;Bacteroidales;Bacteroidaceae;Bacteroides;uncultured bacterium | 96.0 | 1393 | -1 |
| metaDNA.PFemirge\_212 | 306 | 0.007609 | JQ083789.1.1511 | Bacteria;Desulfobacterota;Desulfovibrionia;Desulfovibrionales;Desulfovibrionaceae;Bilophila;uncultured bacterium | 93.7 | 1512 | -1 |
| metaDNA.PFemirge\_144 | 133 | 0.003861 | DQ793316.1.1393 | Bacteria;Bacteroidota;Bacteroidia;Bacteroidales;Tannerellaceae;Parabacteroides;uncultured bacterium | 91.7 | 1175 | -1 |
| metaDNA.PFemirge\_83 | 120 | 0.002874 | LT629973.912339.913840 | Bacteria;Verrucomicrobiota;Verrucomicrobiae;Verrucomicrobiales;Akkermansiaceae;Akkermansia;Akkermansia glycaniphila | 98.9 | 1412 | -1 |
| metaDNA.PFemirge\_633 | 116 | 0.002690 | CP012938.2352114.2353639 | Bacteria;Bacteroidota;Bacteroidia;Bacteroidales;Bacteroidaceae;Bacteroides;Bacteroides ovatus | 93.8 | 1295 | -1 |


##### SSU rRNA from trusted contigs

Full-length SSU rRNA seqeunces extracted from trusted contigs file, matched to SILVA database by Vsearch.

| OTU | Mapped | DB hit | Taxonomy | % ID | Alnlen | Evalue |
| --- | --- | --- | --- | --- | --- | --- |


##### Taxonomic affiliation of unassembled SSU rRNA reads

Approximate overview of taxonomic composition for reads that did NOT assemble into full-length sequences, based on mapping hits to SILVA SSU rRNA database with BBmap.

Taxonomy summarized at level 4. Only displaying taxa with > 3 reads mapped.

| Taxon | Reads |
| --- | --- |
| Bacteria;Bacteroidota;Bacteroidia;Bacteroidales | 2995 |
| Bacteria;Firmicutes;Clostridia;Lachnospirales | 1769 |
| Bacteria;Firmicutes;Clostridia;Oscillospirales | 1719 |
| Bacteria;(Bacteria);(Bacteria);(Bacteria) | 980 |
| Bacteria;Desulfobacterota;Desulfovibrionia;Desulfovibrionales | 647 |
| Bacteria;Actinobacteriota;Actinobacteria;Micrococcales | 471 |
| Bacteria;Proteobacteria;Gammaproteobacteria;Enterobacterales | 375 |
| Bacteria;Firmicutes;Clostridia;(Clostridia) | 373 |
| Bacteria;Firmicutes;Bacilli;Bacillales | 368 |
| Bacteria;Firmicutes;Bacilli;Erysipelotrichales | 359 |
| Bacteria;Firmicutes;(Firmicutes);(Firmicutes) | 248 |
| Bacteria;Bacteroidota;Bacteroidia;(Bacteroidia) | 199 |
| Bacteria;Proteobacteria;Gammaproteobacteria;Burkholderiales | 192 |
| Bacteria;Firmicutes;Clostridia;Clostridia UCG-014 | 172 |
| Bacteria;Firmicutes;Bacilli;RF39 | 171 |
| Bacteria;Firmicutes;Clostridia;Peptostreptococcales-Tissierellales | 166 |
| Bacteria;Proteobacteria;Alphaproteobacteria;Rhodobacterales | 162 |
| Bacteria;Cyanobacteria;Cyanobacteriia;Chloroplast | 145 |
| Bacteria;Proteobacteria;Gammaproteobacteria;Legionellales | 142 |
| Bacteria;Fusobacteriota;Fusobacteriia;Fusobacteriales | 110 |
| Bacteria;Verrucomicrobiota;Verrucomicrobiae;Verrucomicrobiales | 109 |
| Bacteria;Firmicutes;Bacilli;Lactobacillales | 102 |
| Bacteria;Firmicutes;Negativicutes;Acidaminococcales | 99 |
| Bacteria;Proteobacteria;Alphaproteobacteria;Rhodospirillales | 94 |
| Eukaryota;Amorphea;Obazoa;Opisthokonta | 93 |
| Bacteria;Actinobacteriota;Actinobacteria;(Actinobacteria) | 80 |
| Bacteria;Firmicutes;Bacilli;(Bacilli) | 80 |
| Bacteria;Verrucomicrobiota;Verrucomicrobiae;(Verrucomicrobiae) | 71 |
| Bacteria;Actinobacteriota;Actinobacteria;Streptomycetales | 57 |
| Eukaryota;Archaeplastida;Chloroplastida;Charophyta | 50 |
| Bacteria;Firmicutes;Clostridia;Clostridiales | 50 |
| Bacteria;Proteobacteria;Alphaproteobacteria;Rhizobiales | 42 |
| Bacteria;Firmicutes;Bacilli;Staphylococcales | 41 |
| Bacteria;Proteobacteria;(Proteobacteria);(Proteobacteria) | 41 |
| Bacteria;Firmicutes;Clostridia;Christensenellales | 37 |
| Bacteria;Firmicutes;Negativicutes;Veillonellales-Selenomonadales | 36 |
| Bacteria;Bacteroidota;Bacteroidia;Flavobacteriales | 36 |
| Bacteria;Verrucomicrobiota;Verrucomicrobiae;Pedosphaerales | 35 |
| Bacteria;Desulfobacterota;(Desulfobacterota);(Desulfobacterota) | 29 |
| Bacteria;Proteobacteria;Alphaproteobacteria;(Alphaproteobacteria) | 27 |
| Bacteria;Proteobacteria;Alphaproteobacteria;Rickettsiales | 25 |
| Bacteria;Verrucomicrobiota;Chlamydiae;Chlamydiales | 24 |
| Bacteria;Firmicutes;Bacilli;Paenibacillales | 22 |
| Bacteria;Cyanobacteria;Vampirivibrionia;Gastranaerophilales | 19 |
| Bacteria;Caldatribacteriota;JS1;terrestrial metagenome | 18 |
| Bacteria;Bacteroidota;Bacteroidia;Cytophagales | 18 |
| Bacteria;Chloroflexi;Chloroflexia;Chloroflexales | 17 |
| Eukaryota;Amorphea;Amoebozoa;Discosea | 16 |
| Bacteria;Firmicutes;Clostridia;Monoglobales | 16 |
| Bacteria;Firmicutes;Negativicutes;(Negativicutes) | 15 |
| Bacteria;Proteobacteria;Gammaproteobacteria;Pseudomonadales | 15 |
| Bacteria;Firmicutes;Desulfitobacteriia;Desulfitobacteriales | 14 |
| Bacteria;Acidobacteriota;Blastocatellia;Blastocatellales | 13 |
| Bacteria;Bacteroidota;(Bacteroidota);(Bacteroidota) | 12 |
| Bacteria;Actinobacteriota;Actinobacteria;Propionibacteriales | 11 |
| Bacteria;Actinobacteriota;Actinobacteria;Corynebacteriales | 10 |
| Bacteria;Proteobacteria;Alphaproteobacteria;Holosporales | 8 |
| Bacteria;Firmicutes;Desulfotomaculia;Desulfotomaculales | 7 |
| Eukaryota;(Eukaryota);(Eukaryota);(Eukaryota) | 7 |
| Bacteria;Halanaerobiaeota;Halanaerobiia;Halanaerobiales | 6 |
| Eukaryota;SAR;Alveolata;Ciliophora | 6 |
| Bacteria;Desulfobacterota;Desulfuromonadia;PB19 | 6 |
| Bacteria;Firmicutes;Syntrophomonadia;Syntrophomonadales | 6 |
| Bacteria;Caldatribacteriota;JS1;uncultured Candidatus Atribacteria bacterium | 6 |
| Bacteria;Proteobacteria;Gammaproteobacteria;(Gammaproteobacteria) | 6 |
| Bacteria;Actinobacteriota;Thermoleophilia;Solirubrobacterales | 5 |
| Bacteria;Firmicutes;Clostridia;Eubacteriales | 5 |
| Bacteria;MBNT15;Actinobacteria bacterium RBG\_13\_63\_9;(Actinobacteria bacterium RBG\_13\_63\_9) | 5 |
| Bacteria;Firmicutes;Clostridia;Caldicoprobacterales | 4 |
| Bacteria;Firmicutes;Bacilli;RsaHf231 | 4 |
| Bacteria;Firmicutes;Bacilli;Izemoplasmatales | 4 |
| Bacteria;Firmicutes;BRH-c20a;uncultured bacterium | 4 |
| Bacteria;Proteobacteria;Gammaproteobacteria;Xanthomonadales | 4 |
| Bacteria;Firmicutes;Clostridia;uncultured | 4 |
| Bacteria;Firmicutes;Bacilli;Entomoplasmatales | 4 |
| Bacteria;Acidobacteriota;Vicinamibacteria;Vicinamibacterales | 4 |

##### Please cite...

Harald R Gruber-Vodicka, Brandon KB Seah, Elmar Pruesse. 2019 (preprint). phyloFlash - Rapid SSU rRNA profiling and targeted assembly from metagenomes. bioRxiv 521922; doi: https://doi.org/10.1101/521922

###### Cite dependencies when used

- Quast C. et al. 2013. The SILVA ribosomal RNA gene database project: improved data processing and web-based tools. *Nucl. Acids Res.* 41: D590-D596. doi:10.1093/nar/gks1219. Homepage
- Bushnell B. BBMap. Online: https://sourceforge.net/projects/bbmap/
- Bankevich A., et al. 2012. SPAdes: A New Genome Assembly Algorithm and Its Applications to Single-Cell Sequencing. *J. Comput. Biol.* 19 (5): 455-477. doi:10.1089/cmb.2012.0021. Homepage.
- Katoh K., Standley D.M.. 2013. MAFFT Multiple Sequence Alignment Software Version 7: Improvements in Performance and Usability. *Mol. Biol. Evol.* 30 (4): 772-780. doi:10.1093/molbev/mst010. Homepage
- Rognes T. et al. 2016. VSEARCH: a versatile open source tool for metagenomics. *PeerJ* 4: e2584. doi:10.7717/peerj.2584. Homepage
- Miller C.S. et al. 2011. EMIRGE: reconstruction of full-length ribosomal genes from microbial community short read sequencing data. *Genome Biol.* 12: R44. doi:10.1186/gb-2011-12-5-r44. Homepage
- Kopylova E., Noé L., Touzet H. 2012. SortMeRNA: fast and accurate filtering of ribosomal RNAs in metatranscriptomic data. *Bioinformatics* 28 (24): 3211-3217. doi: 10.1093/bioinformatics/bts611. Homepage
- Bedtools. Online: http://bedtools.readthedocs.io/en/latest/
- Seemann T. Barrnap. Online: https://github.com/tseemann/barrnap
- Wheeler T.J., Eddy S.R. 2013. nhmmer: DNA homology search with profile HMMs. *Bioinformatics* 29 (19): 2487-2489. doi:10.1093/bioinformatics/btt403 Homepage
