## Supplementary material for "Gut microbial ecology of Xenopus tadpoles across life stages": Figure_S11: Figure_S11_metaRNA_MATAM_krona.html

Javascript must be enabled to view this page.

magnitude
magnitudeUnassigned

scaffolds.NR.min\_500bp.abd.rdp.fltr.krona

15598937.91

15339457.91

462757.7

462757.7

462757.7

462757.7

462757.7

1011122.51

1011122.51

1011122.51

1011122.51

1011122.51

567220.57

567220.57

60599.5

60599.5

60599.5

506621.07

381870.29

3648.18

378222.11

84420.9

84420.9

40329.88

40329.88

6205990.91

6183861.88

6183861.88

95214.89

95214.89

2925561.28

2873772.08

51789.2

2596136.73

2589135.28

7001.45

566948.98

320550.37

32964.47

102402.46

93872.76

17158.92

22129.03

22129.03

22129.03

22129.03

5161278.26

531321.35

101302.29

101302.29

101302.29

153029.9

153029.9

14285.26

268.42

6060.42

9208.72

7510.19

39123.31

76573.58

229021.1

229021.1

229021.1

1371.1

1371.1

1371.1

29789.9

29789.9

29789.9

16807.06

16807.06

646.72

16160.34

3525830.51

3046720.18

3046720.18

2606940.92

439779.26

479110.33

479110.33

479110.33

711169.2

484595.92

484595.92

3250.03

238847.76

3895.41

238602.72

4094.31

3381.22

3381.22

713.09

713.09

26838.3

26838.3

26838.3

195640.67

195640.67

61174.71

44620.53

89845.43

392957.2

5781.61

5781.61

5781.61

67448.39

67448.39

67448.39

112880.43

63797.28

63797.28

40428.88

40428.88

8654.27

8654.27

206846.77

206846.77

206846.77

1811891.96

1620510.87

1620510.87

2723.53

2723.53

23848.19

23848.19

945988.47

94345.3

62533.07

516049.92

228546.78

44513.4

3883.93

3883.93

42787.76

42787.76

7408.42

7408.42

159221.82

16560.74

79282.87

63378.21

434648.75

96324.46

13389.53

275735.32

49199.44

61121.33

61121.33

61121.33

61121.33

95979.69

95979.69

95979.69

66395.72

29583.97

34280.07

34280.07

34280.07

34280.07

2968.42

2968.42

2968.42

2968.42

2968.42

63182.24

63182.24

63182.24

63182.24

63182.24

47274

47274

47274

47274

47274

5771.34

5771.34

5771.34

5771.34

5771.34

259480

259480

259480

259480

259480

259480
