## Supplementary material for "Gut microbial ecology of Xenopus tadpoles across life stages": Figure_S12: Figure_S12_metaRNA.phyloFlash.html

phyloFlash results summary for library metaRNA


##### image/svg+xml phyloFlash v3.3b3 by Harald Gruber-Vodicka, Elmar Pruesse, Brandon Seah

##### High throughput phylogenetic screening using SSU rRNA gene(s) abundance(s)

Click on report section headers to expand, mouse-over underlined text to see explanations.

### Library name: metaRNA

#### Graphical Summary

Mouseover on panels to expand details.

Mapping identity (%)


20

30

40

50

60

70

80

90

100


1000000

2000000

3000000

4000000

5000000

6000000

7000000

8000000

8947232
Read-mapping %identity of reads vs. reference database. Lower %identity hits may indicate presence of divergent taxa not represented in the database.

Insert size (bp)


100

200

300

400

500

600

700

800

900

1000

1101


1000000

2000000

3052946
Insert sizes for read pairs. Distribution should generally be unimodal; more than one peak may indicate contamination from other libraries.

5.559 % pairs mapped


Mapped pair
Mapped bad pair
Mapped single
Proportion of reads mapped to SSU rRNA database. Typically < 1% for metagenomes, ca. 20% for metatranscriptomes without rRNA depletion or poly-A selection.

Reads assembled


Assembled
Unassembled
Proportion of reads assembled to full-length sequences. High proportion unassembled suggest either assembly failure or high diversity of organisms with low coverage.

Taxonomic summary from reads mapped

Bacteria;Bacteroidota;Bacteroidia;Bacteroidales
3006774

Bacteria;Desulfobacterota;Desulfovibrionia;Desulfovibrionales
1552119

Bacteria;Firmicutes;Negativicutes;Acidaminococcales
876192

Bacteria;Firmicutes;Clostridia;Lachnospirales
292599

Bacteria;Firmicutes;Clostridia;Oscillospirales
262203

Bacteria;Actinobacteriota;Actinobacteria;Micrococcales
242424

Bacteria;(Bacteria);(Bacteria);(Bacteria)
188518

Bacteria;Proteobacteria;Gammaproteobacteria;Enterobacterales
182557

Bacteria;Proteobacteria;Gammaproteobacteria;Burkholderiales
63968

Eukaryota;Amorphea;Obazoa;Opisthokonta
60533

Bacteria;Proteobacteria;Alphaproteobacteria;(Alphaproteobacteria)
55977

Bacteria;Proteobacteria;Alphaproteobacteria;Rhodospirillales
54633

Bacteria;Firmicutes;Bacilli;Erysipelotrichales
54346

Bacteria;Firmicutes;Clostridia;Peptostreptococcales-Tissierellales
34015

Bacteria;Firmicutes;(Firmicutes);(Firmicutes)
30864

Eukaryota;SAR;Rhizaria;Retaria
27001

Bacteria;Bacteroidota;Bacteroidia;(Bacteroidia)
24582

Bacteria;Proteobacteria;Alphaproteobacteria;Paracaedibacterales
23219

Bacteria;Verrucomicrobiota;Verrucomicrobiae;Verrucomicrobiales
22665

Bacteria;Proteobacteria;Alphaproteobacteria;Rhizobiales
20235

Bacteria;Cyanobacteria;Vampirivibrionia;Gastranaerophilales
19394

Bacteria;Firmicutes;Clostridia;(Clostridia)
18290

Bacteria;Deferribacterota;Deferribacteres;Deferribacterales
17546

Eukaryota;Amorphea;Amoebozoa;Discosea
17460

Bacteria;Proteobacteria;(Proteobacteria);(Proteobacteria)
17446

Bacteria;Firmicutes;Bacilli;(Bacilli)
16159

Bacteria;Proteobacteria;Alphaproteobacteria;Zavarziniales
14809

Bacteria;Firmicutes;Negativicutes;(Negativicutes)
14786

Eukaryota;SAR;Stramenopiles;Opalinata
14562

Eukaryota;SAR;Stramenopiles;(Stramenopiles)
13717

Eukaryota;SAR;(SAR);(SAR)
13028

Bacteria;Firmicutes;Negativicutes;Veillonellales-Selenomonadales
10775

Eukaryota;(Eukaryota);(Eukaryota);(Eukaryota)
10686

Bacteria;Firmicutes;Bacilli;Bacillales
10027

Eukaryota;Amorphea;Amoebozoa;Archamoebae
9759

Bacteria;Firmicutes;Bacilli;Paenibacillales
9599

Bacteria;Firmicutes;Clostridia;Clostridia UCG-014
9150

Bacteria;Verrucomicrobiota;Verrucomicrobiae;(Verrucomicrobiae)
8181

Bacteria;Synergistota;Synergistia;Synergistales
7400

Bacteria;Firmicutes;Clostridia;Christensenellales
7225

Bacteria;Actinobacteriota;Actinobacteria;(Actinobacteria)
7078

Bacteria;Desulfobacterota;(Desulfobacterota);(Desulfobacterota)
6867

Bacteria;Proteobacteria;Alphaproteobacteria;Holosporales
5540

Bacteria;Bacteroidota;Bacteroidia;Flavobacteriales
5389

Bacteria;Proteobacteria;Alphaproteobacteria;Caedibacterales
4977

Bacteria;Firmicutes;Bacilli;RF39
4931

Bacteria;Firmicutes;Bacilli;Lactobacillales
4861

Eukaryota;Amorphea;Amoebozoa;(Amoebozoa)
4790

Bacteria;Proteobacteria;Gammaproteobacteria;(Gammaproteobacteria)
4007

Bacteria;Firmicutes;Clostridia;Clostridiales
3922

Bacteria;Verrucomicrobiota;Lentisphaeria;Victivallales
3770

Bacteria;Cyanobacteria;Cyanobacteriia;Chloroplast
3118

Eukaryota;SAR;Alveolata;Ciliophora
3049

Bacteria;Verrucomicrobiota;Chlamydiae;Chlamydiales
3042

Bacteria;Proteobacteria;Alphaproteobacteria;Azospirillales
3029

Bacteria;Proteobacteria;Gammaproteobacteria;Pseudomonadales
2537

Bacteria;Bacteroidota;Bacteroidia;Cytophagales
2383

Bacteria;Bacteroidota;Bacteroidia;Sphingobacteriales
2050

Bacteria;Proteobacteria;Gammaproteobacteria;Xanthomonadales
1991

Bacteria;Bacteroidota;Bacteroidia;Chitinophagales
1826

Eukaryota;SAR;Stramenopiles;Incertae Sedis
1820

Bacteria;Acidobacteriota;Subgroup 22;uncultured bacterium
1780

Bacteria;Proteobacteria;Alphaproteobacteria;Dongiales
1728

Bacteria;Firmicutes;Clostridia;uncultured
1549

Eukaryota;Amorphea;Amoebozoa;uncultured eukaryote
1475

Bacteria;Proteobacteria;Alphaproteobacteria;uncultured
1430

Bacteria;Proteobacteria;Gammaproteobacteria;Legionellales
1414

Bacteria;Proteobacteria;Alphaproteobacteria;Rickettsiales
1336

Bacteria;Actinobacteriota;Actinobacteria;Corynebacteriales
1292

Bacteria;Cyanobacteria;Cyanobacteriia;Cyanobacteriales
1250

Bacteria;Proteobacteria;Alphaproteobacteria;Sphingomonadales
1148

Bacteria;Fusobacteriota;Fusobacteriia;Fusobacteriales
967

Bacteria;Firmicutes;Bacilli;RsaHf231
957

Bacteria;Firmicutes;Bacilli;Staphylococcales
918

Eukaryota;Archaeplastida;Chloroplastida;Charophyta
916

Bacteria;Proteobacteria;Alphaproteobacteria;Reyranellales
886

Bacteria;Desulfobacterota;Desulfuromonadia;Geobacterales
842

Eukaryota;Amorphea;Amoebozoa;Incertae Sedis
813

Bacteria;Proteobacteria;Alphaproteobacteria;Elsterales
786

Bacteria;Proteobacteria;Gammaproteobacteria;Gammaproteobacteria Incertae Sedis
733

Bacteria;Firmicutes;Bacilli;Brevibacillales
715

Bacteria;Proteobacteria;Alphaproteobacteria;Parvibaculales
680

Bacteria;Proteobacteria;Alphaproteobacteria;AT-s3-44
559

Bacteria;Firmicutes;Desulfotomaculia;Desulfotomaculales
518

Bacteria;Firmicutes;Bacilli;Izemoplasmatales
500

Bacteria;Proteobacteria;Alphaproteobacteria;Kiloniellales
497

Bacteria;Actinobacteriota;Actinobacteria;Propionibacteriales
479

Bacteria;Firmicutes;Clostridia;Eubacteriales
474

Bacteria;Proteobacteria;Alphaproteobacteria;Rhodobacterales
450

Bacteria;Proteobacteria;Gammaproteobacteria;Diplorickettsiales
448

Bacteria;Verrucomicrobiota;Kiritimatiellae;Kiritimatiellales
433

Bacteria;Firmicutes;Clostridia;Monoglobales
426

Eukaryota;Amorphea;Amoebozoa;SAR
422

Bacteria;Actinobacteriota;Thermoleophilia;Gaiellales
378

Bacteria;Actinobacteriota;Acidimicrobiia;IMCC26256
377

Bacteria;Firmicutes;Bacilli;Haloplasmatales
364

Bacteria;Desulfobacterota;Desulfobulbia;Desulfobulbales
361

Bacteria;Verrucomicrobiota;Verrucomicrobiae;Chthoniobacterales
357

Bacteria;Actinobacteriota;Coriobacteriia;Coriobacteriales
356

Bacteria;Firmicutes;Clostridia;Clostridia vadinBB60 group
352

Bacteria;Calditrichota;Calditrichia;Calditrichales
348

Bacteria;MBNT15;Actinobacteria bacterium RBG\_13\_63\_9;(Actinobacteria bacterium RBG\_13\_63\_9)
341

Eukaryota;Discoba;Discicristata;Euglenozoa
334

Eukaryota;Amorphea;(Amorphea);(Amorphea)
333

Bacteria;Firmicutes;Clostridia;Peptococcales
318

Eukaryota;SAR;Stramenopiles;MAST-3
315

Bacteria;Cyanobacteria;Vampirivibrionia;Obscuribacterales
297

Bacteria;Proteobacteria;Alphaproteobacteria;Puniceispirillales
281

Bacteria;Firmicutes;Moorellia;Desulfitibacterales
281

Eukaryota;SAR;Stramenopiles;Labyrinthulomycetes
273

Bacteria;Proteobacteria;Gammaproteobacteria;Methylococcales
272

Bacteria;Proteobacteria;Gammaproteobacteria;Acidithiobacillales
257

Bacteria;Firmicutes;Desulfitobacteriia;Desulfitobacteriales
251

Bacteria;Actinobacteriota;Actinobacteria;Pseudonocardiales
248

Eukaryota;SAR;Rhizaria;Cercozoa
232

Eukaryota;SAR;Stramenopiles;Ochrophyta
225

Bacteria;Proteobacteria;Gammaproteobacteria;uncultured
220

Bacteria;Proteobacteria;Alphaproteobacteria;Thalassobaculales
211

Bacteria;Proteobacteria;Gammaproteobacteria;Ectothiorhodospirales
206

Bacteria;Actinobacteriota;Actinobacteria;Actinomycetales
201

Bacteria;Firmicutes;Bacilli;Thermoactinomycetales
188

Bacteria;Acidobacteriota;Holophagae;Subgroup 7
183

Bacteria;Acidobacteriota;Vicinamibacteria;Vicinamibacterales
176

Bacteria;Sva0485;uncultured bacterium;(uncultured bacterium)
170

Eukaryota;SAR;Stramenopiles;Peronosporomycetes
170

Bacteria;Firmicutes;Dethiobacteria;Dethiobacterales
157

Bacteria;Bacteroidota;(Bacteroidota);(Bacteroidota)
154

Bacteria;Firmicutes;Thermoanaerobacteria;Thermoanaerobacterales
137

Bacteria;Firmicutes;Clostridia;Gracilibacteraceae
136

Bacteria;Verrucomicrobiota;Verrucomicrobiae;Pedosphaerales
136

Bacteria;Halanaerobiaeota;Halanaerobiia;Halanaerobiales
135

Eukaryota;Archaeplastida;Chloroplastida;Chlorophyta
133

Bacteria;Spirochaetota;MVP-15;uncultured Firmicutes bacterium
133

Bacteria;Verrucomicrobiota;Omnitrophia;Omnitrophales
116

Bacteria;Actinobacteriota;Actinobacteria;Streptosporangiales
113

Bacteria;Actinobacteriota;Actinobacteria;PeM15
100

Bacteria;Proteobacteria;Alphaproteobacteria;Defluviicoccales
100

Bacteria;Myxococcota;Myxococcia;Myxococcales
98

Bacteria;Firmicutes;Desulfotomaculia;(Desulfotomaculia)
93

Bacteria;Verrucomicrobiota;Verrucomicrobiae;Arctic97B-4 marine group
91

Bacteria;Proteobacteria;Alphaproteobacteria;Kordiimonadales
91

Bacteria;Planctomycetota;Planctomycetes;Pirellulales
90

Bacteria;Verrucomicrobiota;(Verrucomicrobiota);(Verrucomicrobiota)
89

Bacteria;Actinobacteriota;(Actinobacteriota);(Actinobacteriota)
77

Bacteria;Firmicutes;Limnochordia;Limnochordales
76

Bacteria;Spirochaetota;Brachyspirae;Brachyspirales
76

Bacteria;Cyanobacteria;Cyanobacteriia;(Cyanobacteriia)
74

Bacteria;Nitrospirota;Thermodesulfovibrionia;uncultured
74

Bacteria;Verrucomicrobiota;Verrucomicrobiae;Opitutales
73

Bacteria;Firmicutes;Incertae Sedis;Gelria
72

Bacteria;Desulfobacterota;Desulfobacteria;Desulfobacterales
71

Bacteria;Verrucomicrobiota;Lentisphaeria;(Lentisphaeria)
67

Bacteria;Desulfobacterota;Syntrophia;Syntrophales
65

Eukaryota;SAR;Alveolata;Protalveolata
65

Bacteria;Spirochaetota;Spirochaetia;Spirochaetales
62

Bacteria;Myxococcota;Polyangia;mle1-27
61

Bacteria;Chloroflexi;Chloroflexia;Chloroflexales
60

Bacteria;Proteobacteria;Alphaproteobacteria;Ferrovibrionales
60

Bacteria;Myxococcota;Polyangia;Polyangiales
55

Bacteria;Acidobacteriota;Acidobacteriae;Subgroup 13
54

Bacteria;Actinobacteriota;Actinobacteria;Streptomycetales
53

Bacteria;Firmicutes;Bacilli;Exiguobacterales
49

Bacteria;Caldisericota;Caldisericia;Caldisericales
48

Bacteria;Proteobacteria;Alphaproteobacteria;Caulobacterales
47

Bacteria;Planctomycetota;Planctomycetes;Planctomycetales
47

Bacteria;Methylomirabilota;Methylomirabilia;Methylomirabilales
44

Bacteria;Planctomycetota;Phycisphaerae;S-70
42

Eukaryota;SAR;Alveolata;(Alveolata)
42

Bacteria;Desulfobacterota;Desulfuromonadia;Desulfuromonadales
40

Bacteria;Acidobacteriota;(Acidobacteriota);(Acidobacteriota)
40

Bacteria;Planctomycetota;Phycisphaerae;Pla1 lineage
39

Bacteria;Deinococcota;Deinococci;Deinococcales
38

Bacteria;Cloacimonadota;Cloacimonadia;Cloacimonadales
38

Bacteria;Planctomycetota;Planctomycetes;Gemmatales
37

Bacteria;Myxococcota;Polyangia;Haliangiales
37

Bacteria;Firmicutes;Syntrophomonadia;Syntrophomonadales
36

Bacteria;Acidobacteriota;Holophagae;(Holophagae)
32

Bacteria;Desulfobacterota;Desulfuromonadia;(Desulfuromonadia)
32

Bacteria;Dadabacteria;Dadabacteriia;Dadabacteriales
31

Bacteria;Proteobacteria;Gammaproteobacteria;Coxiellales
31

Bacteria;Gemmatimonadota;Gemmatimonadetes;Gemmatimonadales
30

Bacteria;Elusimicrobiota;Elusimicrobia;Elusimicrobiales
30

Bacteria;Firmicutes;Bacilli;Alicyclobacillales
28

Bacteria;Desulfobacterota;Desulfuromonadia;PB19
28

Eukaryota;Amorphea;Amoebozoa;Tubulinea
26

Bacteria;Patescibacteria;Saccharimonadia;Saccharimonadales
26

Bacteria;Campylobacterota;Campylobacteria;Campylobacterales
26

Bacteria;Proteobacteria;Gammaproteobacteria;JTB23
25

Bacteria;Bdellovibrionota;Bdellovibrionia;Bdellovibrionales
25

Eukaryota;Discoba;Discicristata;Heterolobosea
24

Bacteria;Bdellovibrionota;Oligoflexia;0319-6G20
24

Bacteria;Firmicutes;Clostridia;Caldicoprobacterales
24

Bacteria;Firmicutes;uncultured;uncultured bacterium
22

Bacteria;Proteobacteria;Alphaproteobacteria;Sneathiellales
21

Bacteria;Verrucomicrobiota;Lentisphaeria;P.palmC41
21

Bacteria;Actinobacteriota;Acidimicrobiia;Microtrichales
21

Eukaryota;Amorphea;Amoebozoa;metagenome
21

Bacteria;Dependentiae;Babeliae;Babeliales
20

Bacteria;Patescibacteria;Microgenomatia;uncultured sediment bacterium
18

Eukaryota;Archaeplastida;Chloroplastida;(Chloroplastida)
18

Bacteria;Planctomycetota;vadinHA49;uncultured bacterium
17

Eukaryota;uncultured Entamoeba;(uncultured Entamoeba);(uncultured Entamoeba)
16

Eukaryota;Archaeplastida;Rhodophyceae;Bangiales
16

Bacteria;Proteobacteria;Alphaproteobacteria;Acetobacterales
15

Bacteria;Actinobacteriota;Thermoleophilia;Solirubrobacterales
15

Bacteria;MBNT15;Nitrospirae bacterium CG2\_30\_70\_394;(Nitrospirae bacterium CG2\_30\_70\_394)
14

Bacteria;Actinobacteriota;Actinobacteria;Frankiales
14

Bacteria;Myxococcota;Polyangia;(Polyangia)
14

Bacteria;Planctomycetota;Planctomycetes;Isosphaerales
13

Bacteria;Bacteroidota;Kapabacteria;Kapabacteriales
13

Bacteria;Cyanobacteria;Vampirivibrionia;(Vampirivibrionia)
13

Bacteria;Firmicutes;Bacilli;Aneurinibacillales
12

Bacteria;RCP2-54;metagenome;(metagenome)
12

Bacteria;Caldatribacteriota;JS1;uncultured bacterium
12

Bacteria;Verrucomicrobiota;Kiritimatiellae;WCHB1-41
12

Bacteria;Proteobacteria;Gammaproteobacteria;Thiotrichales
11

Eukaryota;SAR;Alveolata;Apicomplexa
11

Bacteria;Desulfobacterota;Desulfobacteria;Desulfatiglandales
11

Bacteria;NB1-j;uncultured bacterium;(uncultured bacterium)
11

Bacteria;Proteobacteria;Gammaproteobacteria;Steroidobacterales
11

Bacteria;Bacteroidota;Bacteroidia;Bacteroidetes VC2.1 Bac22
11

Bacteria;Firmicutes;Desulfotomaculia;Ammonifexales
11

Bacteria;Nitrospirota;4-29-1;uncultured bacterium
10

Bacteria;Bacteroidota;Ignavibacteria;Ignavibacteriales
10

Bacteria;Planctomycetota;Phycisphaerae;Phycisphaerales
10

Bacteria;Marinimicrobia (SAR406 clade);uncultured bacterium;(uncultured bacterium)
9

Bacteria;Firmicutes;D8A-2;uncultured bacterium
9

Bacteria;Firmicutes;Sulfobacillia;Sulfobacillales
9

Eukaryota;SAR;Alveolata;uncultured
9

Bacteria;Patescibacteria;Gracilibacteria;metagenome
9

Bacteria;Hydrogenedentes;Hydrogenedentia;Hydrogenedentiales
9

Bacteria;Proteobacteria;Gammaproteobacteria;HOC36
9

Bacteria;Desulfobacterota;uncultured;uncultured bacterium
8

Bacteria;MBNT15;uncultured delta proteobacterium;(uncultured delta proteobacterium)
8

Eukaryota;Amorphea;Amoebozoa;Schizoplasmodiida
8

Bacteria;Patescibacteria;Gracilibacteria;uncultured bacterium
8

Bacteria;Actinobacteriota;Actinobacteria;Glycomycetales
7

Bacteria;Firmicutes;Limnochordia;MBA03
7

Bacteria;Sva0485;(Sva0485);(Sva0485)
7

Bacteria;Verrucomicrobiota;Verrucomicrobiae;Methylacidiphilales
7

Bacteria;Patescibacteria;Gracilibacteria;(Gracilibacteria)
7

Bacteria;Proteobacteria;Gammaproteobacteria;Cardiobacteriales
7

Bacteria;Proteobacteria;Gammaproteobacteria;Piscirickettsiales
7

Bacteria;Fermentibacterota;Fermentibacteria;Fermentibacterales
7

Bacteria;Proteobacteria;Gammaproteobacteria;Granulosicoccales
7

Eukaryota;Excavata;Metamonada;Preaxostyla
6

Bacteria;Bdellovibrionota;Oligoflexia;Oligoflexales
6

Bacteria;Proteobacteria;Magnetococcia;Magnetococcales
6

Eukaryota;SAR;Stramenopiles;MAST-12
6

Bacteria;Firmicutes;BRH-c20a;uncultured bacterium
6

Bacteria;Myxococcota;bacteriap25;uncultured bacterium
6

Bacteria;Firmicutes;Clostridia;Proteinivoracales
6

Bacteria;Patescibacteria;Microgenomatia;Candidatus Roizmanbacteria
6

Eukaryota;Cryptophyceae;Cryptomonadales;Cryptomonas
6

Bacteria;Acidobacteriota;Blastocatellia;Blastocatellales
6

Eukaryota;SAR;Rhizaria;(Rhizaria)
6

Bacteria;Firmicutes;Symbiobacteriia;Symbiobacteriales
6

Bacteria;Chloroflexi;Anaerolineae;SBR1031
5

Bacteria;Proteobacteria;Alphaproteobacteria;Micavibrionales
5

Bacteria;Firmicutes;Bacilli;Acholeplasmatales
5

Bacteria;RCP2-54;uncultured bacterium;(uncultured bacterium)
5

Bacteria;Acidobacteriota;Thermoanaerobaculia;Thermoanaerobaculales
5

Bacteria;Desulfobacterota;uncultured;uncultured delta proteobacterium
5

Bacteria;Planctomycetota;Phycisphaerae;MSBL9
5

Bacteria;bacterium GBS-1;(bacterium GBS-1);(bacterium GBS-1)
5

Bacteria;Nitrospirota;Nitrospiria;Nitrospirales
5

Bacteria;Firmicutes;Negativicutes;uncultured
4

Eukaryota;Amorphea;Amoebozoa;Gracilipodida
4

Bacteria;Schekmanbacteria;uncultured bacterium;(uncultured bacterium)
4

Bacteria;Elusimicrobiota;Elusimicrobia;(Elusimicrobia)
4

Bacteria;Abditibacteriota;Abditibacteria;Abditibacteriales
4

Bacteria;Nitrospirota;Thermodesulfovibrionia;Magnetobacteriales
4

Bacteria;Bdellovibrionota;Oligoflexia;uncultured
4

Bacteria;Myxococcota;Polyangia;Blfdi19
4

Bacteria;Latescibacterota;Latescibacteria;Latescibacterales
4

Bacteria;Proteobacteria;Gammaproteobacteria;Nitrosococcales
4

Eukaryota;Haptophyta;T58;uncultured marine haptophyte
4

Eukaryota;Amorphea;Amoebozoa;Mycamoeba
4

Bacteria;TA06;uncultured bacterium;(uncultured bacterium)
4

Bacteria;Methylomirabilota;Methylomirabilia;Rokubacteriales
3

Bacteria;Firmicutes;Bacilli;Entomoplasmatales
3

Bacteria;Acidobacteriota;Blastocatellia;Pyrinomonadales
3

Bacteria;Proteobacteria;Gammaproteobacteria;Francisellales
3

Eukaryota;Haptophyta;Prymnesiophyceae;Isochrysidales
3

Bacteria;Campylobacterota;Desulfurellia;Desulfurellales
3

Bacteria;Acidobacteriota;Aminicenantia;Aminicenantales
3

Bacteria;Nitrospirota;(Nitrospirota);(Nitrospirota)
3

Bacteria;Acidobacteriota;Holophagae;Holophagales
3

Bacteria;Bacteroidota;Rhodothermia;Balneolales
3

Eukaryota;Cryptophyceae;Cryptomonadales;(Cryptomonadales)
3

Bacteria;Myxococcota;bacteriap25;metagenome
3

Bacteria;Desulfobacterota;uncultured;Desulfarculaceae bacterium JdFR-97
3

Bacteria;Proteobacteria;Gammaproteobacteria;EV818SWSAP88
3

Bacteria;Actinobacteriota;Actinobacteria;Bifidobacteriales
3

Eukaryota;Amorphea;Amoebozoa;Myxogastria
3

Bacteria;MBNT15;uncultured bacterium;(uncultured bacterium)
3

Bacteria;Fibrobacterota;Fibrobacteria;Fibrobacterales
3

Bacteria;Actinobacteriota;Actinobacteria;Euzebyales
3

Bacteria;Acidobacteriota;Subgroup 22;uncultured Acidobacteriaceae bacterium
3

Bacteria;SAR324 clade(Marine group B);uncultured organism;(uncultured organism)
3

Bacteria;Firmicutes;Moorellia;Moorellales
3

Bacteria;Proteobacteria;Gammaproteobacteria;AT-s2-59
3

Bacteria;Myxococcota;Polyangia;Nannocystales
3

Bacteria;Chloroflexi;Chloroflexia;Thermomicrobiales
3

Bacteria;Spirochaetota;Leptospirae;Leptospirales
2

Bacteria;Chloroflexi;Chloroflexia;Kallotenuales
2

Bacteria;Actinobacteriota;Actinobacteria;uncultured
2

Bacteria;Firmicutes;uncultured;uncultured Clostridiales bacterium
2

Bacteria;Desulfobacterota;Desulfobaccia;Desulfobaccales
2

Bacteria;Acidobacteriota;Subgroup 25;uncultured Acidobacteria bacterium
2

Bacteria;Cyanobacteria;(Cyanobacteria);(Cyanobacteria)
2

Bacteria;Patescibacteria;Parcubacteria;CG1-02-40-25
2

Bacteria;Proteobacteria;Gammaproteobacteria;Thioprofundaceae
2

Bacteria;Nitrospinota;P9X2b3D02;uncultured proteobacterium
2

Bacteria;Acidobacteriota;Blastocatellia;11-24
2

Bacteria;Myxococcota;(Myxococcota);(Myxococcota)
2

Bacteria;Firmicutes;Clostridia;Thermincolales
2

Bacteria;Desulfobacterota;uncultured;uncultured Desulfuromonadales bacterium
2

Bacteria;Planctomycetota;Phycisphaerae;Tepidisphaerales
2

Bacteria;Firmicutes;Natranaerobiia;Natranaerobiales
2

Bacteria;Spirochaetota;MVP-15;uncultured rumen bacterium
2

Bacteria;Actinobacteriota;Coriobacteriia;(Coriobacteriia)
2

Bacteria;Patescibacteria;CPR2;uncultured eubacterium WCHB1-60
2

Bacteria;Desulfobacterota;Syntrophobacteria;Syntrophobacterales
2

Bacteria;Planctomycetota;BD7-11;uncultured bacterium
2

Bacteria;Proteobacteria;Gammaproteobacteria;Arenicellales
2

Bacteria;Chloroflexi;Ktedonobacteria;Ktedonobacterales
2

Bacteria;Proteobacteria;Gammaproteobacteria;Thiomicrospirales
2

Eukaryota;Amorphea;Amoebozoa;Dictyostelia
2

Bacteria;Proteobacteria;Gammaproteobacteria;Salinisphaerales
2

Bacteria;Desulfobacterota;uncultured;delta proteobacterium WX81
2

Bacteria;Chloroflexi;TK10;metagenome
2

Eukaryota;Cryptophyceae;Cryptomonadales;Rhodomonas
2

Bacteria;Poribacteria;(Poribacteria);(Poribacteria)
1

Eukaryota;Archaeplastida;(Archaeplastida);(Archaeplastida)
1

Bacteria;Sva0485;bioreactor metagenome;(bioreactor metagenome)
1

Bacteria;MBNT15;(MBNT15);(MBNT15)
1

Bacteria;Verrucomicrobiota;Lentisphaeria;Oligosphaerales
1

Bacteria;Gemmatimonadota;Longimicrobia;Longimicrobiales
1

Bacteria;Planctomycetota;Phycisphaerae;(Phycisphaerae)
1

Bacteria;Nitrospinota;MT5B39;Nitrospinae bacterium RIFCSPLOWO2\_12\_FULL\_45\_22
1

Bacteria;Firmicutes;Moorellia;Calderihabitantales
1

Bacteria;Desulfobacterota;uncultured;uncultured sludge bacterium H16
1

Bacteria;Chloroflexi;Anaerolineae;Anaerolineales
1

Bacteria;Proteobacteria;Gammaproteobacteria;B2M28
1

Bacteria;MBNT15;uncultured Desulfuromonadales bacterium;(uncultured Desulfuromonadales bacterium)
1

Bacteria;Proteobacteria;Zetaproteobacteria;Mariprofundales
1

Bacteria;Acidobacteriota;Acidobacteriae;Acidobacteriales
1

Eukaryota;Incertae Sedis;Ancyromonadida;(Ancyromonadida)
1

Bacteria;NB1-j;(NB1-j);(NB1-j)
1

Bacteria;Desulfobacterota;Desulfomonilia;Desulfomonilales
1

Bacteria;Cyanobacteria;Cyanobacteriia;Limnotrichales
1

Bacteria;Proteobacteria;Gammaproteobacteria;Ga0077536
1

Bacteria;Proteobacteria;Gammaproteobacteria;UBA10353 marine group
1

Bacteria;Chloroflexi;TK10;uncultured Chloroflexi bacterium
1

Bacteria;Desulfobacterota;Desulfobacteria;(Desulfobacteria)
1

Bacteria;Acidobacteriota;Subgroup 5;uncultured bacterium
1

Bacteria;Firmicutes;uncultured;Clostridia bacterium TQ1
1

Bacteria;Verrucomicrobiota;Kiritimatiellae;(Kiritimatiellae)
1

Bacteria;Elusimicrobiota;Lineage IIa;(Lineage IIa)
1

Bacteria;Acidobacteriota;Subgroup 25;uncultured bacterium
1

Bacteria;Firmicutes;Bacilli;Mycoplasmatales
1

Bacteria;Armatimonadota;Fimbriimonadia;Fimbriimonadales
1

Bacteria;Verrucomicrobiota;Verrucomicrobiae;UA11
1

Bacteria;Proteobacteria;Alphaproteobacteria;Micropepsales
1

Bacteria;Firmicutes;Thermoanaerobacteria;Caldicellulosiruptorales
1

Bacteria;Myxococcota;bacteriap25;Deltaproteobacteria bacterium RIFCSPLOWO2\_02\_56\_12
1

Bacteria;Planctomycetota;Planctomycetes;(Planctomycetes)
1

Bacteria;NB1-j;uncultured organism;(uncultured organism)
1

Bacteria;Latescibacterota;(Latescibacterota);(Latescibacterota)
1

Bacteria;Gemmatimonadota;BD2-11 terrestrial group;uncultured bacterium
1

Bacteria;Chloroflexi;(Chloroflexi);(Chloroflexi)
1

Bacteria;Elusimicrobiota;4-29;uncultured bacterium
1

Bacteria;Firmicutes;Thermaerobacteria;Thermaerobacterales
1

Bacteria;Caldatribacteriota;JS1;terrestrial metagenome
1

Bacteria;Cyanobacteria;Cyanobacteriia;Leptolyngbyales
1

Bacteria;SAR324 clade(Marine group B);uncultured bacterium;(uncultured bacterium)
1

Bacteria;Desulfobacterota;Desulfarculia;Desulfarculales
1

Bacteria;Bdellovibrionota;Bdellovibrionia;Bacteriovoracales
1

Bacteria;Gemmatimonadota;PAUC43f marine benthic group;uncultured bacterium
1

Bacteria;Proteobacteria;Alphaproteobacteria;Tistrellales
1

Archaea;Crenarchaeota;Nitrososphaeria;Nitrososphaerales
1

Bacteria;NB1-j;uncultured delta proteobacterium;(uncultured delta proteobacterium)
1

Bacteria;Firmicutes;Desulfotomaculia;Carboxydothermales
1

Bacteria;Proteobacteria;Alphaproteobacteria;NRL2
1

Bacteria;Elusimicrobiota;Endomicrobia;Endomicrobiales
1

Bacteria;TX1A-33;uncultured bacterium;(uncultured bacterium)
1

Bacteria;Planctomycetota;Pla4 lineage;uncultured bacterium
1

Coverage on 18S model


100

200

300

400

500

600

700

800

900

1000

1100

1200

1300

1400

1500

1634


10

20

30

40

50

60

70

80

90

100

110
Coverage evenness across eukaryotic 18S rRNA gene model from Barrnap, using Nhmmer from random subsample of mapped reads. This helps to detect contamination from tag sequencing libraries (sharp coverage peaks). For the eukaryotic model it is normal to see one or two regions with low coverage because of variable regions in the 18S rRNA gene that are not present in all organisms.

Coverage on 16S model


100

200

300

400

500

600

700

800

900

1000

1100

1200

1300

1379


1000

2000

3000

4000

5000

6420
Coverage evenness across prokaryotic 16S rRNA gene model from Barrnap, using Nhmmer from random subsample of mapped reads. This helps to detect contamination from tag sequencing libraries (sharp coverage peaks).


##### Tree of full-length assembled sequences (Click to hide)

Full-length assembled SSU rRNA sequences along with closest hits from SILVA database, in an alignment guide tree produced by MAFFT. This tree helps to visualize the relatedness of sequences in library to known relatives. Colored circles have areas proportional to the number of SSU rRNA reads that map to each respective sequence (re-mapping is done separately for SPAdes, EMIRGE, and trusted-contig full-length sequence sets). Click on the toggle switches to turn them on and off. Additional circle representing proportion of SSU rRNA reads that were not assembled is in lower right corner. Long taxonomy strings of reference sequences are truncated, but the full strings are given in the tables below.

**Color key:** Assembled by SPAdes, Reconstructed by EMIRGE, Extracted from trusted contigs,  Closest-matching reference sequence from SILVA database

Toggle reads mapped:
SPAdes

I

O
EMIRGE

I

O
Trusted

I

O
Reads: 1524
Reads: 657
Reads: 2141
Reads: 139
Reads: 24
Reads: 217
Reads: 164
Reads: 7519
Reads: 42
Reads: 124
Reads: 401
Reads: 247
Reads: 5058
Reads: 4
Reads: 111
Reads: 761
Reads: 93
Reads: 10
Reads: 70
Reads: 359
Reads: 124
Reads: 941
Reads: 120
Reads: 89965
Reads: 387
Reads: 173
Reads: 25
Reads: 765
Reads: 207
Reads: 2780
Reads: 25
Reads: 3011
Reads: 1896
Reads: 21
Reads: 6905
Reads: 37
Reads: 527
Reads: 19
Reads: 108
Reads: 533
Reads: 93727
Reads: 70
Reads: 2504
Reads: 10
Reads: 12086
Reads: 271
Reads: 1108
Reads: 30259
Reads: 178
Reads: 87227
Reads: 226
Reads: 118
Reads: 135
Reads: 38
Reads: 2425
Reads: 56
Reads: 114
Reads: 77
Reads: 77
Reads: 387
Reads: 11211
Reads: 1962
Reads: 1683
Reads: 819
Reads: 37024
Reads: 7018
Reads: 30801
Reads: 30
Reads: 49
Reads: 38
Reads: 64
Reads: 279
Reads: 67
Reads: 22
Reads: 18
Reads: 79
Reads: 66
Reads: 2414
Reads: 141
Reads: 61
Reads: 21
Reads: 123
Reads: 122
Reads: 2479
Reads: 85
Reads: 672
Reads: 42
Reads: 25
Reads: 143
Reads: 2155
Reads: 49
Reads: 159
Reads: 133
Reads: 20222
Reads: 78
Reads: 13
Reads: 93
Reads: 5796
Reads: 263
Reads: 244
Reads: 47190
Reads: 448
Reads: 389027
Reads: 95
Reads: 29759
Reads: 26367
Reads: 73
Reads: 20
Reads: 38
Reads: 76
Reads: 204
Reads: 651
Reads: 310
Reads: 30
Reads: 357
Reads: 3
Reads: 152
Reads: 1042
Reads: 104
Reads: 487
Reads: 105
Reads: 800
Reads: 17
Reads: 56
Reads: 52
Reads: 627
Reads: 39
Reads: 34
Reads: 34
Reads: 14737
Reads: 14529
Reads: 31
Reads: 39
Reads: 26513
Reads: 74
Reads: 781
Reads: 48
Reads: 18
Reads: 7709
Reads: 2307
Reads: 7503
Reads: 142
Reads: 243
Reads: 4032
Reads: 119
Reads: 2224
Reads: 2282
Reads: 4651
Reads: 31507
Reads: 282
Reads: 10770
Reads: 5
Reads: 113
Reads: 80
Reads: 15
Reads: 81
Reads: 46
Reads: 26103
Reads: 51
Reads: 174
Reads: 30
Reads: 44035
Reads: 97
Reads: 1262
Reads: 42
Reads: 80
Reads: 11651
Reads: 15
Reads: 106
Reads: 57
Reads: 69
Reads: 261
Reads: 4485
Reads: 461834
Reads: 209
Reads: 549
Reads: 34
Reads: 59
Reads: 33
Reads: 25
Reads: 5818
Reads: 1348
Reads: 64
Reads: 169824
Reads: 607
Reads: 2031
Reads: 18911
Reads: 95
Reads: 7352
Reads: 62
Reads: 178
Reads: 4618
Reads: 32
Reads: 164
Reads: 49
Reads: 202
Reads: 38
Reads: 409
Reads: 478
Reads: 7100
Reads: 24
Reads: 128
Reads: 49
Reads: 104
Reads: 1294
Reads: 2782
Reads: 45
Reads: 1006
Reads: 32
Reads: 986
Reads: 2716
Reads: 83
Reads: 162
Reads: 94
Reads: 24
Reads: 30
Reads: 24
Reads: 379
Reads: 786
Reads: 268
Reads: 91
Reads: 25
Reads: 27
Reads: 250
Reads: 35
Reads: 74577
Reads: 53
Reads: 38
Reads: 695
Reads: 16
Reads: 34
Reads: 10475
Reads: 36
Reads: 271
Reads: 62
Reads: 64
Reads: 1518
Reads: 74
Reads: 13
Reads: 50
Reads: 1428
Reads: 457
Reads: 178
Reads: 156
Reads: 262
Reads: 70
Reads: 44
Reads: 36
Reads: 807
Reads: 61
Reads: 291
Reads: 93
Reads: 151
Reads: 259
Reads: 248
Reads: 40417
Reads: 8
Reads: 8472
Reads: 1205
Reads: 124
Reads: 32
Reads: 27513
Reads: 428
Reads: 219
Reads: 38
Reads: 90762
Reads: 388
Reads: 102
Reads: 37
Reads: 398
Reads: 65
Reads: 3
Reads: 95
Reads: 80
Reads: 8
Reads: 92
Reads: 366
Reads: 64
Reads: 219
Reads: 9559
Reads: 284
Reads: 3025
Reads: 14
Reads: 54
Reads: 69
Reads: 17
Reads: 40074
Reads: 1925
Reads: 175
Reads: 7861
Reads: 36
Reads: 98
Reads: 70
Reads: 2672
Reads: 25
Reads: 136635
Reads: 292
Reads: 3897
Reads: 131
Reads: 39
Reads: 41
Reads: 2342
Reads: 87
Reads: 92
Reads: 112
Reads: 75
Reads: 34
Reads: 97
Reads: 309
Reads: 3846
Reads: 827
Reads: 444
Reads: 117
Reads: 5209
Reads: 44
Reads: 142
Reads: 49
Reads: 322
Reads: 90
Reads: 24
Reads: 47
Reads: 50
Reads: 188
Reads: 2077
Reads: 79
Reads: 547
Reads: 117
Reads: 18
Reads: 28
Reads: 38
Reads: 1615
Reads: 46
Reads: 21
Reads: 209
Reads: 423
Reads: 2128
Reads: 27
Reads: 16
Reads: 556
Reads: 22
Reads: 55
Reads: 142
Reads: 407
Reads: 575
Reads: 3317
Reads: 98
Reads: 10
Reads: 58
Reads: 357
Reads: 2624
Reads: 61995
Reads: 340
Reads: 48
Reads: 4090
Reads: 194
Reads: 104
Reads: 806
Reads: 53
Reads: 292
Reads: 40
Reads: 925
Reads: 8
Reads: 1071
Reads: 698
Reads: 54
Reads: 1016
Reads: 32
Reads: 284
Reads: 86
Reads: 1063
Reads: 69
Reads: 61610
Reads: 43978
Reads: 48057
Reads: 16
Reads: 86
Reads: 14
Reads: 284
Reads: 161
Reads: 337
Reads: 243
Reads: 107
Reads: 46
Reads: 609
Reads: 199
Reads: 75
Reads: 77
Reads: 2188
Reads: 43
Reads: 137
Reads: 19
Reads: 9595
Reads: 34155
Reads: 76
Reads: 23
Reads: 163
Reads: 7890
Reads: 1823
Reads: 144
Reads: 61
Reads: 76
Reads: 33
Reads: 22533
Reads: 2311
Reads: 501
Reads: 1340
Reads: 473
Reads: 37
Reads: 73
Reads: 341
Reads: 2151
Reads: 13
Reads: 44
Reads: 32239
Reads: 146
Reads: 41
Reads: 172
Reads: 45
Reads: 63
Reads: 56
Reads: 11
Reads: 311
Reads: 155
Reads: 550
Reads: 43
Reads: 470
Reads: 5097
Reads: 201
Reads: 70
Reads: 300
Reads: 25
Reads: 133978
Reads: 4451
Reads: 421
Reads: 17
Reads: 5
Reads: 46129
Reads: 6646
Reads: 30
Reads: 53
Reads: 133
Reads: 123
Reads: 1273
Reads: 678
Reads: 304
Reads: 42
Reads: 688
Reads: 212
Reads: 36
Reads: 202
Reads: 49
Reads: 1729
Reads: 54
Reads: 56
Reads: NA
Reads: 20
Reads: 2402
Reads: 34553
Reads: 57
Reads: 53
Reads: 77
Reads: 128
Reads: 90
Reads: 102
Reads: 1785595
Reads: 2334
Reads: 41
Reads: 563
Reads: 25
Reads: 77
Reads: 32893
Reads: 45
Reads: 579
Reads: 8296
Reads: 102
Reads: 4553
Reads: 1507
Reads: 48
Reads: 1258
Reads: 1219
Reads: 40
Reads: 155
Reads: 102
Reads: 85064
Reads: 100
Reads: 64
Reads: 32
Reads: 174
Reads: 135
Reads: 52
Reads: 831
Reads: 291
Reads: 19
Reads: 3263
Reads: 4307
Reads: 1846
Reads: 9704
Reads: 2304
Reads: 644
Reads: 389
Reads: 76786
Reads: 115
Reads: 325
Reads: 221
Reads: 117
Reads: 8
Reads: 6
Reads: 45
Reads: 267
Reads: 33
Reads: 400
Reads: 85
Reads: 44
Reads: 638
Reads: 611
Reads: 17
Reads: 152
Reads: 50
Reads: 111
Reads: 88
Reads: 62
Reads: 173119
Reads: 2417
Reads: 91
Reads: 7825
Reads: 96
Reads: 159
Reads: 28
Reads: 44
Reads: 33628
Reads: 78
Reads: 50
Reads: 18
Reads: 44
Reads: 120
Reads: 210
Reads: 37
Reads: 746
Reads: 24
Reads: 64876
Reads: 18731
Reads: 48
Reads: 58
Reads: 32001
Reads: 20651
Reads: 18899
Reads: 199
Reads: 45
Reads: 341
Reads: 217
Reads: 2379
Reads: 953
Reads: 74
Reads: 814
Reads: 489412
Reads: 143
Reads: 5494
Reads: 7602
Reads: 160
Reads: 24
Reads: 32
Reads: 24848
Reads: 2453
Reads: 87
Reads: 230
Reads: 312
Reads: 28
Reads: 21
Reads: 884
Reads: 18
Reads: 15836
Reads: 40
Reads: 105
Reads: 48
Reads: 93
Reads: 151
Reads: 76
Reads: 21
Reads: 191
Reads: 291
Reads: 2206
Reads: 290
Reads: 7567
Reads: 4205
Reads: 71
Reads: 267
Reads: 15
Reads: 548
Reads: 36
Reads: 67
Reads: 98
Reads: 726
Reads: 5051
Reads: 374
Reads: 69
Reads: 29684
Reads: 140
Reads: 123
Reads: 7029
Reads: 44
Reads: 1152939
Reads: 18
Reads: 68
Reads: 1951
Reads: 1304
Reads: 17358
Reads: 266
Reads: 69
Reads: 499
Reads: 2643
Reads: 223
Reads: 1713
Reads: 13003
Reads: 395
Reads: 219
Reads: 1199
Reads: 14
Reads: 3619
Reads: 25942
Reads: 31
Reads: 304
Reads: 279
Reads: 270
Reads: 177423
Reads: 4058
Reads: 32
Reads: 993
Reads: 448044
Reads: 63
Reads: 123
Reads: 80
Reads: 228
Reads: 36
Reads: 116
Reads: 514
Reads: 1142
Reads: 9
Reads: 22
Reads: 30
Reads: 80
Reads: 28
Reads: 214
Reads: 487
Reads: 422201
Reads: 2840
Reads: 1255
Reads: 462
Reads: 35
Reads: 648
Reads: 114
Reads: 29336
Reads: 812
Reads: 1373
Reads: 18
Reads: 4357
Reads: 264
Reads: 41
Reads: 67067
Reads: 228
Reads: 41
Reads: 463
Reads: 354
Reads: 86
Reads: 38
Reads: 1168
Reads: 97
Reads: 456
Reads: 46
Reads: 34
Reads: 64
Reads: 265
Reads: 29
Reads: 32
Reads: 299
Reads: 68
Reads: 296
Reads: 121
Reads: 31
Reads: 441
Reads: 2768
Reads: 794
Reads: 118
Reads: 201
Reads: 443
Reads: 10418
Reads: 66
Reads: 98
Reads: 1634
Reads: 140
Reads: 329
Reads: 102
Reads: 71
Reads: 272
Reads: 64
Reads: 149
Reads: 29
Reads: 17
Reads: 1386
Reads: 28
Reads: 16
Reads: 53
Reads: 16372
Reads: 1801
Reads: 2315
Reads: 69
Reads: 78
Reads: 462
Reads: 218
Reads: 43481
Reads: 76997
Reads: 214
Reads: 49
Reads: 28
Reads: 176
Reads: 76492
Reads: 44
Reads: 279
Reads: 97
Reads: 377
Reads: 3352
Reads: 274
Reads: 1139
Reads: 95
Reads: 156
Reads: 5979
Reads: 280
Reads: 56
Reads: 90
Reads: 33
Reads: 2276
Reads: 24858
Reads: 11342
Reads: 26
Reads: 61
Reads: 342
Reads: 2137
Reads: 27
Reads: 830
Reads: 33
Reads: 338
Reads: 107658
Reads: 24
Reads: 2012
Reads: 2743
Reads: 19569
Reads: 93
Reads: 3689
Reads: 489
Reads: 54
Reads: 355
Reads: 2961
Reads: 81
Reads: 643
Reads: 46
Reads: 21679
Reads: 1359
Reads: 450
Reads: 116
Reads: 213
Reads: 128
Reads: 70
Reads: 84
Reads: 547
Reads: 202
Reads: 754
Reads: 535
Reads: 90
Reads: 92
Reads: 968
Reads: 1189
Reads: 58
Reads: 80
Reads: 54
Reads: 321
Reads: 23
Reads: 53
Reads: 155
Reads: 39306
Reads: 92
Reads: 256
Reads: 87
Reads: 72
Reads: 4634
Reads: 24
Reads: 1947
Reads: 5421
Reads: 1606
Reads: 474
Reads: 25
Reads: 283
Reads: 48
Reads: 31
Reads: 41
Reads: 2366
Reads: 225
Reads: 458
Reads: 273
Reads: 19859
Reads: 253
Reads: 132
Reads: 1735
Reads: 6
Reads: 213
Reads: 48
Reads: 22906
Reads: 105
Reads: 63
Reads: 83
Reads: 60
Reads: 349
Reads: 33
Reads: 113
Reads: 115
Reads: 688
Reads: 51
Reads: 104
Reads: 1855
Reads: 772
Reads: 285
Reads: 656
Reads: 294
Reads: 231
Reads: 63
Reads: 581
Reads: 66
Reads: 923
Reads: 757
Reads: 141
Reads: 32
Reads: 30
Reads: 297
Reads: 37
Reads: 4436
Reads: 227
Reads: 2171
Reads: 12325
Reads: 47
Reads: 256
Reads: 773
Reads: 1282
Reads: 40528
Reads: 29
Reads: 82
Reads: 109
Reads: 11682
Reads: 136
Reads: 161
Reads: 177292
Reads: 72
Reads: 2660
Reads: 547
Reads: 32
Reads: 4
Reads: 16
Reads: 1430
Reads: 28
Reads: 17
Reads: 2153
Reads: 132
Reads: 42
Reads: 108
Reads: 12476
Reads: 28
Reads: 133
Reads: 49
Reads: 439
Reads: 102
Reads: 4069
Reads: 57
Reads: 8453
Reads: 2532
Reads: 5
Reads: 699
Reads: 554970
Reads: 7391
Reads: 212
Reads: 112
Reads: 60409
Reads: 74
Reads: 100
Reads: 117
Reads: 217
Reads: 36
Reads: 37
Reads: 1921
Reads: 967
Reads: 39
Reads: 74
Reads: 628
Reads: 81
Reads: 149
Reads: 1012
Reads: 95153
Reads: 142
Reads: 195
Reads: 486
Reads: 133
Reads: 601
Reads: 52063
Reads: 18
Reads: 76
Reads: 55173
Reads: 795
Reads: 20
Reads: 87977
Reads: 10
Reads: 195
Reads: 12
Reads: 262
Reads: 337
Reads: 16
Reads: 8
Reads: 215
Reads: 10
Reads: 9
Reads: 1789171
Reads: 8
Reads: 32
Reads: 5790
Reads: 14
Reads: 3068
Reads: 949
Reads: 1589
Reads: 1546
Reads: 342
Reads: 121
Reads: 92
Reads: 79
Reads: 204
Reads: 1644
Reads: 30
Reads: 14
Reads: 48
Reads: 283
Reads: 82
Reads: 62
Reads: 4535
Reads: 211
Reads: 51
Reads: 344
Reads: 249
Reads: 1577
Reads: 152
Reads: 93
Reads: 116
Reads: 494
Reads: 973
Reads: 59
Reads: 468
Reads: 303
Reads: 1367
Reads: 19
Reads: 589
Reads: 1
Reads: 372
Reads: 359
Reads: 4738
Reads: 333
Reads: 32
Reads: 184
Reads: 77061
Reads: 13380
Reads: 3064
Reads: 1375
Reads: 16891
Reads: 3033
Reads: 111
Reads: 2058
Reads: 28
Reads: 1014
Reads: 1294
Reads: 212
Reads: 258
Reads: 733
Reads: 5903
Reads: 238
Reads: 694
Reads: 1234
Reads: 63
Reads: 43
Reads: 1867
Reads: 112
Reads: 28
Reads: 49219
Reads: 213
Reads: 521
Reads: 18161
Reads: 667
Reads: 11396
Reads: 3765
Reads: 126
Reads: 62
Reads: 9432
Reads: 245
Reads: 131
Reads: 148
Reads: 516
Reads: 27
Reads: 48
Reads: 26
Reads: 178
Reads: 1147
Reads: 905
Reads: 80
Reads: 216
Reads: 659
Reads: 5807
Reads: 252
Reads: 243
Reads: 654
Reads: 24
Reads: 131
Reads: 747
Reads: 188
Reads: 550
Reads: 813
Reads: 9228
Reads: 23
Reads: 344
Reads: 2626
Reads: 77
Reads: 20949
Reads: 30
Reads: 163
Reads: 73
Reads: 40
Reads: 4338
Reads: 252
Reads: 161
Reads: 48
Reads: 104
Reads: 97
Reads: 24
Reads: 25
Reads: 60
Reads: 15
Reads: 121
Reads: 5590
Reads: 112
Reads: 107
Reads: 106209
Reads: 10835
Reads: 58
Reads: 41455
Reads: 100
Reads: 584
Reads: 74561
Reads: 721
Reads: 106
Reads: 280
Reads: 25
Reads: 292
Reads: 16
Reads: 202
Reads: 23
Reads: 75
Reads: 2418
Reads: 33
Reads: 16613
Reads: 36
Reads: 100
Reads: 14
Reads: 240
Reads: 28453
Reads: 70
Reads: 72
Reads: 73
Reads: 13
Reads: 6
Reads: 727
Reads: 59384
Reads: 16
Reads: 410
Reads: 16
Reads: 80
Reads: 13
Reads: 31
Reads: 3956
Reads: 480
Reads: 100
Reads: 10806
Reads: 97
Reads: 295
Reads: 280
Reads: 873
Reads: 40
Reads: 335
Reads: 48
Reads: 84
Reads: 794
Reads: 726246
Reads: 68
Reads: 28
Reads: 95
Reads: 1088
Reads: 44
Reads: 1441
Reads: 74
Reads: 43
Reads: 280
Reads: 91903
Reads: 124
Reads: 57
Reads: 116
Reads: 113
Reads: 2149
Reads: 27
Reads: 161
Reads: 91
Reads: 3713
Reads: 418
Reads: 38
Reads: 371
Reads: 109
Reads: 329726
Reads: 96
Reads: 92765
Reads: 107
Reads: 282
Reads: 947
Reads: 101
Reads: 268
Reads: 132
Reads: 313
Reads: 53838
Reads: 101
Reads: 22
Reads: 474
Reads: 387
Reads: 5774
Reads: 1207
Reads: 74
Reads: 87
Reads: 2161
Reads: 36
Reads: 166
Reads: 2627
Reads: 30
Reads: 34
Reads: 17
Reads: 220
Reads: 260
Reads: 844
Reads: 39
Reads: 8
Reads: 8
Reads: 475
Reads: 7986
Reads: 47
Reads: 21
Reads: 279194
Reads: 25
Reads: 15
Reads: 44
Reads: 442
Reads: 1194
Reads: 213
Reads: 1857
Reads: 198
Reads: 418
Reads: 56
Reads: 182
Reads: 67
Reads: 53
Reads: 1056
Reads: 39
Reads: 80
Reads: 637
Reads: 215
Reads: 16
Reads: 58
Reads: 676
Reads: 50
Reads: 37
Reads: 200
Reads: 5385
Reads: 67
Reads: 2124
Reads: 113
Reads: 1541
Reads: 725
Reads: 914
Reads: 12978
Reads: 70334
Reads: 58
Reads: 171
Reads: 2938
Reads: 213
Reads: 166
Reads: 79
Reads: 352
Reads: 108
Reads: 43490
Reads: 81
Reads: 34
Reads: 281
Reads: 52
Reads: 11
Reads: 581
Reads: 197
Reads: 13434
Reads: 67
Reads: 315
Reads: 204
Reads: 401
Reads: 380
Reads: 361
Reads: 202
Reads: 18
Reads: 26
Reads: 38
Reads: 174
Reads: 56
Reads: 91
Reads: 96
Reads: 37864
Reads: 52
Reads: 217
Reads: 120
Reads: 4588
Reads: 56
Reads: 131
Reads: 94
Reads: 9069
Reads: 74
Reads: 207
Reads: 83119
Reads: 2103
Reads: 192
Reads: 27
Reads: 61735
Reads: 344
Reads: 68
Reads: 242
Reads: 393909
Reads: 2138
Reads: 57
Reads: 81
Reads: 404
Reads: 67
Reads: 2080
Reads: 24069
Reads: 53
Reads: 32
Reads: 49
Reads: 28215
Reads: 22622
Reads: 577
Reads: 383
Reads: 17821
Reads: 38
Reads: 18
Reads: 8102
Reads: 10711
Reads: 39
Reads: 51
Reads: 196
Reads: 222
Reads: 111
Reads: 583
Reads: 358
Reads: 96
Reads: 94
Reads: 94
Reads: 1179
Reads: 22
Reads: 119
Reads: 156
Reads: 46
Reads: 276
Reads: 125
Reads: 455
Reads: 128
Reads: 78
Reads: 10
Reads: 182
Reads: 10779
Reads: 1628
Reads: 92
Reads: 112
Reads: 41
Reads: 238
Reads: 508
Reads: 117
Reads: 108
Reads: 82
Reads: 78
Reads: 551
Reads: 106
Reads: 378
Reads: 106
Reads: 13
Reads: 635
Reads: 205
Reads: 48
Reads: 21
Reads: 14
Reads: 46
Reads: 84
Reads: 139
Reads: 8
Reads: 40
Reads: 775
Reads: 1084
Reads: 38
Reads: 301942
Reads: 3646
Reads: 6076
Reads: 1334
Reads: 70
Reads: 229
Reads: 40
Reads: 284
Reads: 12332
Reads: 475
Reads: 8
Reads: 2822
Reads: 7983
Reads: 38
Reads: 174
Reads: 146
Reads: 273
Reads: 7163
Reads: 18748
Reads: 265
Reads: 9
Reads: 3858
Reads: 148
Reads: 90773
Reads: 56964
Reads: 47590
Reads: 144
Reads: 54
Reads: 1109
Reads: 100
Reads: 465
Reads: 119239
Reads: 242
Reads: 739
Reads: 55
Reads: 72015
Reads: 19388
Reads: 38
Reads: 36070
Reads: 82
Reads: 120
Reads: 287
Reads: 151
Reads: 31339
Reads: 3496
Reads: 74
Reads: 97
Reads: 154
Reads: 57
Reads: 293
Reads: 32
Reads: 26
Reads: 648
Reads: 5174
Reads: 37
Reads: 1437
Reads: 14882
Reads: 606
Reads: 12863
Reads: 58
Reads: 14104
Reads: 50
Reads: 491
Reads: 186
Reads: 10
Reads: 341
Reads: 44
Reads: 20251
Reads: 5994
Reads: 14969
Reads: 109
Reads: 2865
Reads: 946
Reads: 2195
Reads: 354
Reads: 145
Reads: 36
Reads: 1646
Reads: 4186
Reads: 70
Reads: 643
Reads: 41464
Reads: 37875
Reads: 39
Reads: 73
Reads: 1905
Reads: 213
Reads: 3745
Reads: 159
Reads: 1006
Reads: 149
Reads: 314
Reads: 818
Reads: 33
Reads: 36
Reads: 34
Reads: 2889
Reads: 602
Reads: 1561
Reads: 44
Reads: 28
Reads: 53
Reads: 840
Reads: 133
Reads: 920
Reads: 86
Reads: 30
Reads: 8787
Reads: 236687
Reads: 93120
Reads: 303707
Reads: 878
Reads: 166
Reads: 189
Reads: 50
Reads: 74
Reads: 50
Reads: 34
Reads: 23
Reads: 34520
Reads: 71
Reads: 179
Reads: 80
Reads: 916
Reads: 1300
Reads: 62
Reads: 15
Reads: 10
Reads: 460
Reads: 437
Reads: 22
Reads: 115
Reads: 26
Reads: 465
Reads: 23
Reads: 1508
Reads: 499
Reads: 2986
Reads: 10896
Reads: 25590
Reads: 17
Reads: 140
Reads: 987716
Reads: 96
Reads: 26
Reads: 493
Reads: 2452
Reads: 72
Reads: 14
Reads: 11
Reads: 138
Reads: 39
Reads: 542
Reads: 30
Reads: 16
Reads: 101
Reads: 13
Reads: 182
Reads: 107
Reads: 6201
Reads: 471
Reads: 260
Reads: 245
Reads: 703
Reads: 11468
Reads: 1216
Reads: 130
Reads: 154
Reads: 30
Reads: 32
Reads: 21
Reads: 480
Reads: 179
Reads: 12617
Reads: 28369
Reads: 54
Reads: 16725
Reads: 47
Reads: 40
Reads: 64
Reads: 31
Reads: 38
Reads: 288
Reads: 31
Reads: 70
Reads: 5674
Reads: 47
Reads: 69
Reads: 8
Reads: 2774
Reads: 84
Reads: 2812
Reads: 11
Reads: 8262
Reads: 986
Reads: 155
Reads: 38691
Reads: 54
Reads: 1983
Reads: 140
Reads: 12
Reads: 25641
Reads: 87
Reads: 9039
Reads: 83142
Reads: 3478
Reads: 2070
Reads: 2488
Reads: 251
Reads: 248
Reads: 193
Reads: 30
Reads: 39
Reads: 349358
Reads: 7377
Reads: 461
Reads: 68
Reads: 33
Reads: 1371
Reads: 1767
Reads: 32
Reads: 38
Reads: 99
Reads: 443
Reads: 238
Reads: 139
Reads: 201
Reads: 12944
Reads: 94069
Reads: 99
Reads: 76
Reads: 145
Unassembled SSU reads: 446095


DQ800679.1.1366 Bacteria;Firmicutes;Clostridia;Lachnosp ... ae;[Ruminococcus] gauvreauii group;uncultured bacterium


HM630179.1.1443 Bacteria;Cyanobacteria;Vampirivibrionia;Gastranaerophilales;uncultured bacterium


metaRNA.PFemirge\_27699\_0.000031


GQ492245.1.1349 Bacteria;Firmicutes;Bacilli;Erysipelotr ... tridiaceae;Candidatus Stoquefichus;uncultured bacterium


metaRNA.PFemirge\_20664\_0.000055


metaRNA.PFemirge\_5324\_0.000152


HQ815291.1.1415 Bacteria;Firmicutes;Clostridia;Lachnosp ... eae;[Ruminococcus] gauvreauii group;uncultured organism


metaRNA.PFemirge\_23284\_0.000031


metaRNA.PFemirge\_25672\_0.000001


metaRNA.PFemirge\_28668\_0.000017


metaRNA.PFemirge\_16300\_0.000027


KY122019.1.1428 Bacteria;Proteobacteria;Alphaproteobacteria;Rhizobiales;Rhizobiaceae;Mesorhizobium;Mesorhizobium sp.


metaRNA.PFemirge\_2278\_0.000785


metaRNA.PFemirge\_8488\_0.000006


HM038002.1.1488 Bacteria;Firmicutes;Bacilli;Erysipelotr ... eae;[Anaerorhabdus] furcosa group;Anaerorhabdus furcosa


FJ881199.1.1509 Bacteria;Desulfobacterota;Desulfovibrio ... les;Desulfovibrionaceae;uncultured;uncultured bacterium


metaRNA.PFemirge\_7109\_0.000012


AY721628.1.1491 Bacteria;Desulfobacterota;Desulfovibrio ... lfovibrionaceae;Desulfovibrio;bacterium 'New Zealand D'


metaRNA.PFemirge\_3372\_0.000322


metaRNA.PFemirge\_24301\_0.000003


AB088925.1.1354 Bacteria;Bacteroidota;Bacteroidia;Bacte ... s-D38 termite group;uncultured Bacteroidaceae bacterium


metaRNA.PFemirge\_4545\_0.000219


DQ796609.1.1391 Bacteria;Firmicutes;Clostridia;Oscillospirales;Ruminococcaceae;Faecalibacterium;uncultured bacterium


metaRNA.PFemirge\_10915\_0.000003


metaRNA.PFemirge\_9673\_0.000013


metaRNA.PFemirge\_4924\_0.000568


EU461102.1.1401 Bacteria;Firmicutes;Clostridia;Lachnospirales;Lachnospiraceae;Dorea;uncultured bacterium


JN792358.1.1395 Bacteria;Firmicutes;Clostridia;Peptostr ... ptococcaceae;Paraclostridium;uncultured Clostridium sp.


metaRNA.PFemirge\_1733\_0.000003


metaRNA.PFemirge\_13123\_0.000001


FJ901820.1.1378 Bacteria;MBNT15;uncultured delta proteobacterium


metaRNA.PFemirge\_25888\_0.000015


metaRNA.PFemirge\_27879\_0.000016


DQ797819.1.1407 Bacteria;Desulfobacterota;Desulfovibrio ... ;Desulfovibrionaceae;Desulfovibrio;uncultured bacterium


metaRNA.PFemirge\_11222\_0.000001


EU468659.1.1353 Bacteria;Firmicutes;Clostridia;Lachnospirales;Lachnospiraceae;uncultured;uncultured bacterium


metaRNA.PFemirge\_17143\_0.000013


metaRNA.PFemirge\_19772\_0.000019


metaRNA.PFemirge\_17011\_0.005371


metaRNA.PFemirge\_27920\_0.000064


metaRNA.PFemirge\_13109\_0.000020


KT952726.1.1486 Bacteria;Firmicutes;Clostridia;Oscillospirales;Ruminococcaceae;uncultured;uncultured bacterium


DQ817375.1.1403 Bacteria;Firmicutes;Clostridia;Clostrid ... iaceae;Clostridium sensu stricto 1;uncultured bacterium


EU158190.1.1510 Bacteria;Firmicutes;Clostridia;Oscillos ... anaerobacterium;Hydrogenoanaerobacterium saccharovorans


metaRNA.PFemirge\_3621\_0.000002


metaRNA.PFemirge\_7265\_0.000032


metaRNA.PFemirge\_7939\_0.000028


BAAU01002038.417.1926 Bacteria;Firmicutes;Clostridia;Lachnospirales;Lachnospiraceae;uncultured;human gut metagenome


metaRNA.PFemirge\_15599\_0.001175


HQ588840.1.1529 Bacteria;Proteobacteria;Gammaproteobact ... dales;Pseudomonadaceae;Pseudomonas;Pseudomonas sp. BZ77


metaRNA.PFemirge\_10106\_0.000002


JN619526.1.1407 Bacteria;Synergistota;Synergistia;Synergistales;Synergistaceae;uncultured;uncultured bacterium


metaRNA.PFemirge\_12946\_0.000158


metaRNA.PFemirge\_16283\_0.000167


metaRNA.PFemirge\_11478\_0.000002


LC028669.1.1525 Bacteria;Firmicutes;Clostridia;Oscillospirales;Butyricicoccaceae;UCG-009;uncultured bacterium


metaRNA.PFemirge\_28094\_0.001063


metaRNA.PFemirge\_13158\_0.000003


metaRNA.PFemirge\_28125\_0.000035


metaRNA.PFemirge\_23470\_0.000003


EU775673.1.1399 Bacteria;Proteobacteria;Gammaproteobact ... obacteriaceae;Escherichia-Shigella;uncultured bacterium


metaRNA.PFemirge\_10180\_0.000008


GQ898302.1.1489 Bacteria;Bacteroidota;Bacteroidia;Bacteroidales;Bacteroidaceae;Bacteroides;uncultured bacterium


metaRNA.PFemirge\_16968\_0.000152


GBID01005713.1.1428 Bacteria;Bacteroidota;Bacteroidia;B ... ceae;Bacteroides;Blattella germanica (German cockroach)


metaRNA.PFemirge\_15251\_0.005416


FPLS01017333.1.1416 Bacteria;Proteobacteria;Gammaproteo ... proteobacteria Incertae Sedis;Unknown Family;metagenome


metaRNA.PFemirge\_7353\_0.000002


metaRNA.PFemirge\_12447\_0.000311


JF830134.1.1428 Bacteria;Proteobacteria;Alphaproteobact ... Rhodobacteraceae;Pseudorhodobacter;uncultured bacterium


JQ084791.1.1476 Bacteria;Firmicutes;Clostridia;Oscillospirales;Ruminococcaceae;Anaerotruncus;uncultured bacterium


metaRNA.PFemirge\_12794\_0.000013


metaRNA.PFemirge\_15716\_0.001541


JN896612.1.1485 Bacteria;Actinobacteriota;Actinobacteria;Corynebacteriales;Nocardiaceae;Nocardia;Nocardia sp. FSN271


HQ741370.1.1454 Bacteria;Firmicutes;Clostridia;Lachnospirales;Lachnospiraceae;Epulopiscium;uncultured bacterium


metaRNA.PFemirge\_1634\_0.000024


metaRNA.PFemirge\_27660\_0.000081


metaRNA.PFspades\_24\_180.812745


EU466098.1.1227 Bacteria;Firmicutes;Bacilli;Erysipelotrichales;Erysipelotrichaceae;Turicibacter;uncultured bacterium


metaRNA.PFemirge\_22786\_0.000038


metaRNA.PFemirge\_2866\_0.007591


metaRNA.PFemirge\_27669\_0.000009


metaRNA.PFemirge\_23421\_0.000006


metaRNA.PFemirge\_2056\_0.000009


metaRNA.PFtrusted\_14\_0


metaRNA.PFemirge\_9024\_0.000780


KC836026.1.1497 Bacteria;Proteobacteria;Gammaproteobact ... ales;Rhodocyclaceae;Dechlorobacter;uncultured bacterium


metaRNA.PFemirge\_23142\_0.000006


metaRNA.PFemirge\_13120\_0.000006


EU771244.1.1399 Bacteria;Firmicutes;Clostridia;Lachnosp ... aceae;Lachnospiraceae NC2004 group;uncultured bacterium


metaRNA.PFemirge\_16169\_0.000005


metaRNA.PFemirge\_24172\_0.000017


metaRNA.PFemirge\_12795\_0.000251


metaRNA.PFspades\_22\_54.022814


metaRNA.PFemirge\_14379\_0.000010


metaRNA.PFemirge\_10074\_0.000131


DQ794177.1.1393 Bacteria;Firmicutes;Clostridia;Oscillos ... bacterium] coprostanoligenes group;uncultured bacterium


metaRNA.PFemirge\_12905\_0.000056


metaRNA.PFemirge\_2360\_0.002347


AGTS01000010.187.1721 Bacteria;Proteobacteria;Gammaprot ... lderiales;Comamonadaceae;Acidovorax;Acidovorax sp. NO-1


metaRNA.PFemirge\_11849\_0.001477


metaRNA.PFemirge\_7737\_0.002818


DQ809030.1.1400 Bacteria;Bacteroidota;Bacteroidia;Bacteroidales;Rikenellaceae;Alistipes;uncultured bacterium


metaRNA.PFemirge\_18796\_0.000002


EU777338.1.1348 Bacteria;Bacteroidota;Bacteroidia;Bacteroidales;Rikenellaceae;dgA-11 gut group;uncultured bacterium


metaRNA.PFemirge\_22906\_0.000002


metaRNA.PFemirge\_3312\_0.000004


FJ951888.1.1489 Bacteria;Bacteroidota;Bacteroidia;Bacteroidales;Rikenellaceae;Alistipes;uncultured bacterium


metaRNA.PFemirge\_22176\_0.000008


metaRNA.PFemirge\_13617\_0.000003


metaRNA.PFemirge\_12656\_0.000021


metaRNA.PFemirge\_12520\_0.000004


metaRNA.PFemirge\_7187\_0.000001


metaRNA.PFemirge\_7058\_0.000007


metaRNA.PFemirge\_4565\_0.000003


metaRNA.PFspades\_21\_13.924386


metaRNA.PFemirge\_25902\_0.000006


HM017143.1.1714 Eukaryota;Amorphea;Amoebozoa;Discosea;Flabellinia;Dactylopodida;Angulamoeba;Angulamoeba microcystivorans


metaRNA.PFemirge\_18876\_0.000005


metaRNA.PFemirge\_16322\_0.000001


KC358008.1.1260 Bacteria;Bacteroidota;Bacteroidia;Bacteroidales;Barnesiellaceae;uncultured;uncultured bacterium


metaRNA.PFemirge\_3590\_0.000010


metaRNA.PFemirge\_4086\_0.000010


metaRNA.PFemirge\_10437\_0.000692


metaRNA.PFemirge\_19870\_0.000021


metaRNA.PFemirge\_22496\_0.000324


metaRNA.PFemirge\_13282\_0.000002


metaRNA.PFemirge\_16359\_0.000008


metaRNA.PFemirge\_30246\_0.000023


HQ766029.1.1454 Bacteria;Firmicutes;Clostridia;Oscillospirales;Ruminococcaceae;Subdoligranulum;uncultured organism


HQ774646.1.1457 Bacteria;Proteobacteria;Gammaproteobact ... iales;Sutterellaceae;Parasutterella;uncultured organism


metaRNA.PFemirge\_7042\_0.000760


metaRNA.PFemirge\_16410\_0.000002


EF511528.1.1486 Bacteria;Proteobacteria;Gammaproteobact ... ;Xanthomonadaceae;Stenotrophomonas;uncultured bacterium


metaRNA.PFemirge\_19358\_0.000346


JN093128.1.1497 Bacteria;Firmicutes;Clostridia;Clostrid ... iaceae;Clostridium sensu stricto 13;Clostridium punense


FWEL01000003.4055.5565 Bacteria;Firmicutes;Clostridia;O ... eae;uncultured;Pseudoflavonifractor sp. Marseille-P3106


metaRNA.PFemirge\_23650\_0.000000


metaRNA.PFemirge\_11898\_0.001264


EU775240.1.1308 Bacteria;Bacteroidota;Bacteroidia;Bacte ... votellaceae;Prevotellaceae UCG-003;uncultured bacterium


metaRNA.PFemirge\_25998\_0.000014


metaRNA.PFemirge\_11870\_0.000005


metaRNA.PFemirge\_23125\_0.000014


metaRNA.PFemirge\_11565\_0.001286


EU136680.1.2019 Bacteria;Bacteroidota;Bacteroidia;Bacteroidales;Bacteroidaceae;Bacteroides;Bacteroides uniformis


AY957925.1.1536 Bacteria;Proteobacteria;Gammaproteobacteria;Legionellales;Legionellaceae;Legionella;uncultured bacterium


metaRNA.PFemirge\_12566\_0.000019


metaRNA.PFemirge\_24040\_0.000018


AB234530.1.1368 Bacteria;Desulfobacterota;Desulfovibrio ... ibrionaceae;uncultured;uncultured delta proteobacterium


metaRNA.PFemirge\_7153\_0.002586


metaRNA.PFemirge\_28178\_0.000003


metaRNA.PFemirge\_7295\_0.019734


metaRNA.PFemirge\_6145\_0.000003


metaRNA.PFemirge\_28104\_0.006325


DQ799921.1.1390 Bacteria;Firmicutes;Clostridia;Oscillospirales;Ruminococcaceae;Faecalibacterium;uncultured bacterium


EU772885.1.1375 Bacteria;Firmicutes;Clostridia;Lachnospirales;Lachnospiraceae;Agathobacter;uncultured bacterium


JF830174.1.1513 Bacteria;Proteobacteria;Gammaproteobact ... les;Enterobacteriaceae;Plesiomonas;uncultured bacterium


metaRNA.PFspades\_8\_105.065262


GQ448053.1.1387 Bacteria;Firmicutes;Clostridia;Peptostr ... caceae;[Eubacterium] nodatum group;uncultured bacterium


metaRNA.PFemirge\_26253\_0.000018


FR749936.1.1524 Bacteria;Firmicutes;Clostridia;Lachnosp ... ubacterium] fissicatena group;Faecalicatena fissicatena


JN680616.1.1459 Bacteria;Firmicutes;Clostridia;Oscillos ... icoccaceae;UCG-009;uncultured Ruminococcaceae bacterium


metaRNA.PFemirge\_26540\_0.000005


metaRNA.PFemirge\_19789\_0.000002


metaRNA.PFemirge\_6959\_0.000005


metaRNA.PFemirge\_17428\_0.000005


metaRNA.PFemirge\_1573\_0.000053


metaRNA.PFemirge\_11834\_0.000029


EU777108.1.1328 Bacteria;Firmicutes;Clostridia;Oscillospirales;Ruminococcaceae;Incertae Sedis;uncultured bacterium


metaRNA.PFemirge\_26488\_0.000004


metaRNA.PFemirge\_16808\_0.000043


EU465560.1.1391 Bacteria;Firmicutes;Clostridia;Lachnosp ... aceae;Lachnospiraceae AC2044 group;uncultured bacterium


metaRNA.PFemirge\_30244\_0.000034


metaRNA.PFemirge\_6131\_0.000014


KT952808.1.1480 Bacteria;Bacteroidota;Bacteroidia;Bacteroidales;Rs-E47 termite group;uncultured bacterium


metaRNA.PFemirge\_21111\_0.000149


metaRNA.PFemirge\_22898\_0.000015


metaRNA.PFemirge\_17357\_0.000059


JX457179.1.1492 Bacteria;Firmicutes;Bacilli;Erysipelotr ... stridiaceae;Erysipelatoclostridium;uncultured bacterium


HQ757204.1.1439 Bacteria;Bacteroidota;Bacteroidia;Bacteroidales;Bacteroidaceae;Bacteroides;uncultured organism


EU472567.1.1360 Bacteria;Firmicutes;Clostridia;Oscillospirales;Ruminococcaceae;Ruminococcus;uncultured bacterium


metaRNA.PFemirge\_11298\_0.000012


FQXY01000005.366.1875 Bacteria;Bacteroidota;Bacteroidia ... teroidaceae;Bacteroides;Bacteroides sp. Marseille-P3132


EF559167.1.1492 Bacteria;Firmicutes;Clostridia;Lachnospirales;Lachnospiraceae;Anaerostignum;uncultured bacterium


AF083009.1.1734 Eukaryota;Amorphea;Obazoa;Opisthokonta; ... Nematoda;Chromadorea;Rhabditida;Diploscapter sp. PS1897


metaRNA.PFemirge\_9685\_0.000101


metaRNA.PFemirge\_2588\_0.000002


metaRNA.PFemirge\_18330\_0.000006


DQ808767.1.1394 Bacteria;Firmicutes;Clostridia;Lachnosp ... ospiraceae;Lachnospiraceae UCG-008;uncultured bacterium


DQ810054.1.1385 Bacteria;Firmicutes;Clostridia;Lachnospirales;Lachnospiraceae;Lachnoclostridium;uncultured bacterium


metaRNA.PFemirge\_6115\_0.000010


DQ806700.1.1391 Bacteria;Firmicutes;Clostridia;Oscillospirales;Oscillospiraceae;Colidextribacter;uncultured bacterium


metaRNA.PFemirge\_30232\_0.000056


metaRNA.PFemirge\_11130\_0.000005


metaRNA.PFemirge\_22134\_0.000006


metaRNA.PFemirge\_20218\_0.000002


metaRNA.PFemirge\_14964\_0.001020


metaRNA.PFemirge\_1686\_0.000861


U81750.1.1482 Bacteria;Firmicutes;Negativicutes;Acidaminococcales;Acidaminococcaceae;uncultured;uncultured bacterium


metaRNA.PFemirge\_5680\_0.000003


EF626688.1.1481 Bacteria;Proteobacteria;Gammaproteobact ... nadales;Xanthomonadaceae;Luteimonas;Luteimonas aquatica


AGES01000014.73256.74770 Bacteria;Bacteroidota;Bacteroi ... nerellaceae;Parabacteroides;Parabacteroides sp. HGS0025


HG970990.1.1557 Bacteria;Firmicutes;Bacilli;Erysipelotrichales;Erysipelotrichaceae;Breznakia;uncultured bacterium


metaRNA.PFemirge\_1784\_0.000005


metaRNA.PFemirge\_27756\_0.002917


CP002345.1438824.1440339 Bacteria;Bacteroidota;Bacteroi ... cteraceae;Paludibacter;Paludibacter propionicigenes WB4


metaRNA.PFemirge\_24949\_0.000006


metaRNA.PFemirge\_22812\_0.000073


metaRNA.PFemirge\_15303\_0.000004


CP000875.5447186.5448660 Bacteria;Chloroflexi;Chlorofle ... onaceae;Herpetosiphon;Herpetosiphon aurantiacus DSM 785


metaRNA.PFemirge\_7277\_0.000003


metaRNA.PFemirge\_28141\_0.001026


metaRNA.PFemirge\_6811\_0.000674


metaRNA.PFemirge\_16116\_0.001083


metaRNA.PFemirge\_25952\_0.000006


FJ269067.1.1493 Bacteria;Bacteroidota;Bacteroidia;Bacte ... s;iron-reducing bacterium enrichment culture clone HN86


AM930363.1.1475 Bacteria;Firmicutes;Clostridia;Clostrid ... iaceae;Clostridium sensu stricto 1;uncultured bacterium


metaRNA.PFemirge\_20554\_0.000011


EU775615.1.1257 Bacteria;Firmicutes;Clostridia;Peptostr ... s;Peptostreptococcaceae;Romboutsia;uncultured bacterium


metaRNA.PFemirge\_944\_0.000203


metaRNA.PFemirge\_13173\_0.000011


KF464766.1.1221 Bacteria;Proteobacteria;Gammaproteobacteria;Enterobacterales;Yersiniaceae;Yersinia;uncultured bacterium


metaRNA.PFspades\_19\_11.010999


EF096599.1.1399 Bacteria;Firmicutes;Clostridia;Lachnosp ... ceae;Lachnospiraceae NK4A136 group;uncultured bacterium


metaRNA.PFemirge\_19929\_0.000020


metaRNA.PFemirge\_1474\_0.000427


metaRNA.PFemirge\_27668\_0.005292


metaRNA.PFemirge\_3476\_0.000057


metaRNA.PFemirge\_4327\_0.001436


metaRNA.PFemirge\_20985\_0.000000


metaRNA.PFemirge\_27885\_0.000004


metaRNA.PFemirge\_25338\_0.000012


metaRNA.PFemirge\_28095\_0.000008


metaRNA.PFemirge\_6260\_0.000009


metaRNA.PFemirge\_17569\_0.000004


DQ793765.1.1405 Bacteria;Firmicutes;Clostridia;Oscillos ... bacterium] coprostanoligenes group;uncultured bacterium


metaRNA.PFemirge\_10716\_0.001165


metaRNA.PFemirge\_22739\_0.000005


metaRNA.PFemirge\_9053\_0.000013


metaRNA.PFemirge\_25175\_0.000001


metaRNA.PFemirge\_11813\_0.004412


GQ448098.1.1374 Bacteria;Firmicutes;Clostridia;Christen ... ceae;Christensenellaceae R-7 group;uncultured bacterium


metaRNA.PFemirge\_20745\_0.000017


metaRNA.PFemirge\_3684\_0.000086


metaRNA.PFemirge\_21473\_0.000005


DQ673521.1.1485 Bacteria;Firmicutes;Clostridia;Lachnosp ... achnospiraceae NK4A136 group;uncultured rumen bacterium


DQ799425.1.1396 Bacteria;Bacteroidota;Bacteroidia;Bacteroidales;Tannerellaceae;Parabacteroides;uncultured bacterium


metaRNA.PFemirge\_24475\_0.000031


metaRNA.PFspades\_20\_61.734259


metaRNA.PFemirge\_18649\_0.000001


metaRNA.PFemirge\_2385\_0.000143


metaRNA.PFemirge\_25787\_0.000008


FJ881357.1.1492 Bacteria;Firmicutes;Clostridia;Oscillospirales;Butyricicoccaceae;UCG-009;uncultured bacterium


EU779425.1.1388 Bacteria;Firmicutes;Clostridia;Oscillos ... bacterium] coprostanoligenes group;uncultured bacterium


metaRNA.PFemirge\_14416\_0.000005


metaRNA.PFemirge\_7082\_0.000018


metaRNA.PFemirge\_2652\_0.000820


AM930333.1.1446 Bacteria;Halanaerobiaeota;Halanaerobiia;Halanaerobiales;Halanaerobiaceae;Halocella;uncultured bacterium


metaRNA.PFemirge\_14997\_0.030526


HQ772787.1.1440 Bacteria;Firmicutes;Clostridia;Lachnospirales;Lachnospiraceae;Agathobacter;uncultured organism


metaRNA.PFemirge\_10811\_0.000021


JN391840.1.1499 Bacteria;Proteobacteria;Gammaproteobact ... deriales;Comamonadaceae;uncultured;uncultured bacterium


EU773260.1.1381 Bacteria;Firmicutes;Clostridia;Lachnospirales;Lachnospiraceae;Tyzzerella;uncultured bacterium


metaRNA.PFemirge\_28268\_0.000173


metaRNA.PFemirge\_10705\_0.000004


metaRNA.PFemirge\_12065\_0.000003


metaRNA.PFemirge\_17303\_0.000008


metaRNA.PFemirge\_26124\_0.000002


metaRNA.PFemirge\_16020\_0.000293


FPLP01013193.1.1403 Bacteria;Cyanobacteria;Vampirivibri ... ;Obscuribacteraceae;Candidatus Obscuribacter;metagenome


metaRNA.PFemirge\_5603\_0.000219


metaRNA.PFemirge\_28124\_0.000008


metaRNA.PFemirge\_4725\_0.027537


KM098109.1.1519 Bacteria;Firmicutes;Clostridia;Oscillospirales;Ruminococcaceae;UBA1819;Ruthenibacterium lactatiformans


metaRNA.PFspades\_27\_3.570273


metaRNA.PFemirge\_13295\_0.000130


AB534319.1.1609 Eukaryota;Amorphea;Obazoa;Opisthokonta; ... mycotina;Dothideomycetes;Pleosporales;uncultured fungus


metaRNA.PFemirge\_17782\_0.001614


metaRNA.PFemirge\_5190\_0.000008


EU510692.1.1406 Bacteria;Desulfobacterota;Desulfovibrio ... les;Desulfovibrionaceae;uncultured;uncultured bacterium


AY554146.1.1441 Bacteria;Desulfobacterota;Desulfovibrio ... esulfovibrionaceae;Desulfovibrio;Desulfovibrio sp. LNB2


metaRNA.PFemirge\_4693\_0.000270


metaRNA.PFemirge\_11772\_0.000013


AB517712.1.1448 Bacteria;Bacteroidota;Bacteroidia;Bacte ... inilabiliaceae;Saccharicrinis;Saccharicrinis fermentans


metaRNA.PFemirge\_18220\_0.000008


metaRNA.PFemirge\_27675\_0.001813


AB243814.1.1495 Bacteria;Bacteroidota;Bacteroidia;Bacteroidales;Rikenellaceae;uncultured;uncultured bacterium


LN810638.1.1401 Bacteria;Bacteroidota;Bacteroidia;Cytophagales;Spirosomaceae;Flectobacillus;Flectobacillus rhizosphaerae


metaRNA.PFemirge\_25314\_0.000004


metaRNA.PFemirge\_11100\_0.000006


HQ385613.1.1405 Bacteria;Proteobacteria;Gammaproteobact ... eria Incertae Sedis;Unknown Family;uncultured bacterium


metaRNA.PFemirge\_26303\_0.000003


metaRNA.PFemirge\_3431\_0.000028


metaRNA.PFemirge\_8952\_0.000006


GFIG01012561.7.1417 Bacteria;Proteobacteria;Gammaproteo ... erichia-Shigella;Coregonus lavaretus (common whitefish)


GU304321.1.1480 Bacteria;Firmicutes;Clostridia;Lachnosp ... ceae;Lachnospiraceae UCG-008;uncultured rumen bacterium


GALZ01007484.194.1638 Bacteria;Cyanobacteria;Cyanobacteriia;Chloroplast;Actinidia chinensis


metaRNA.PFemirge\_25721\_0.000441


metaRNA.PFemirge\_23238\_0.000034


metaRNA.PFemirge\_12796\_0.000920


metaRNA.PFemirge\_22189\_0.000024


metaRNA.PFemirge\_23725\_0.000005


metaRNA.PFemirge\_15683\_0.000008


metaRNA.PFemirge\_11327\_0.000006


AY957923.1.1520 Bacteria;Proteobacteria;Gammaproteobact ... eriales;Comamonadaceae;Curvibacter;uncultured bacterium


metaRNA.PFemirge\_7857\_0.000091


metaRNA.PFemirge\_12893\_0.000289


metaRNA.PFemirge\_19200\_0.000005


metaRNA.PFemirge\_15229\_0.000039


metaRNA.PFemirge\_17065\_0.000009


metaRNA.PFemirge\_27721\_0.000096


metaRNA.PFemirge\_27708\_0.002097


metaRNA.PFemirge\_30255\_0.000046


metaRNA.PFemirge\_12490\_0.000030


EU777648.1.1410 Bacteria;Firmicutes;Bacilli;Lactobacillales;Streptococcaceae;Lactococcus;uncultured bacterium


metaRNA.PFemirge\_27849\_0.000032


LN650648.388047.389538 Bacteria;Firmicutes;Clostridia;P ... les;Peptostreptococcaceae;Romboutsia;Romboutsia hominis


DQ794860.1.1394 Bacteria;Firmicutes;Clostridia;Lachnosp ... ;[Eubacterium] oxidoreducens group;uncultured bacterium


metaRNA.PFemirge\_27752\_0.000024


DQ805650.1.1385 Bacteria;Firmicutes;Clostridia;Lachnosp ... ospiraceae;Lachnospiraceae UCG-008;uncultured bacterium


metaRNA.PFemirge\_20056\_0.000005


metaRNA.PFemirge\_9797\_0.000003


metaRNA.PFemirge\_3072\_0.000018


AGZO01000045.56609.58123 Bacteria;Bacteroidota;Bacteroi ... ;Parabacteroides;Parabacteroides goldsteinii CL02T12C30


metaRNA.PFemirge\_511\_0.000319


metaRNA.PFemirge\_6044\_0.000013


metaRNA.PFemirge\_5928\_0.000011


HQ741466.1.1426 Bacteria;Firmicutes;Clostridia;Clostrid ... iaceae;Clostridium sensu stricto 1;uncultured bacterium


metaRNA.PFemirge\_6116\_0.000006


DQ808503.1.1399 Bacteria;Bacteroidota;Bacteroidia;Bacteroidales;Tannerellaceae;Parabacteroides;uncultured bacterium


metaRNA.PFemirge\_22979\_0.000007


metaRNA.PFemirge\_24647\_0.000018


metaRNA.PFemirge\_7189\_0.000004


FJ477666.1.1463 Bacteria;Proteobacteria;Gammaproteobact ... acteriaceae;Escherichia-Shigella;uncultured Pantoea sp.


metaRNA.PFspades\_18\_408.152000


metaRNA.PFemirge\_8438\_0.000003


metaRNA.PFemirge\_13894\_0.000003


metaRNA.PFemirge\_3393\_0.000032


metaRNA.PFemirge\_26451\_0.000003


AY457069.1.1483 Bacteria;Actinobacteriota;Actinobacteri ... Mycobacteriaceae;Mycobacterium;Mycobacterium peregrinum


DQ393283.1.1453 Bacteria;Cyanobacteria;Cyanobacteriia;C ... ae;Arthrospira PCC-7345;Arthrospira platensis SAG 21.99


metaRNA.PFemirge\_21054\_0.000002


metaRNA.PFemirge\_690\_0.000751


JN032891.1.1470 Bacteria;Desulfobacterota;Desulfovibrio ... ;Desulfovibrionaceae;Desulfovibrio;uncultured bacterium


metaRNA.PFemirge\_1026\_0.000005


metaRNA.PFemirge\_27473\_0.000022


DQ798990.1.1390 Bacteria;Firmicutes;Clostridia;Lachnosp ... aceae;[Ruminococcus] torques group;uncultured bacterium


metaRNA.PFemirge\_3834\_0.000009


HQ317401.1.1485 Bacteria;Cyanobacteria;Cyanobacteriia;Chloroplast;Haematococcus lacustris


JX198613.1.1484 Bacteria;Firmicutes;Clostridia;Lachnosp ... ae;[Ruminococcus] gauvreauii group;uncultured bacterium


HQ143300.1.1507 Bacteria;Proteobacteria;Gammaproteobact ... obacteriaceae;Escherichia-Shigella;uncultured bacterium


metaRNA.PFemirge\_14238\_0.000020


EU777328.1.1296 Bacteria;Firmicutes;Bacilli;RF39;uncultured bacterium


metaRNA.PFemirge\_5205\_0.000169


metaRNA.PFemirge\_2119\_0.000005


metaRNA.PFemirge\_4081\_0.000002


EU508752.1.1287 Bacteria;Firmicutes;Clostridia;Oscillospirales;Ruminococcaceae;uncultured;uncultured bacterium


metaRNA.PFemirge\_24603\_0.000004


EU885024.1.1445 Bacteria;Firmicutes;Clostridia;Oscillospirales;Ruminococcaceae;uncultured;uncultured bacterium


metaRNA.PFemirge\_4739\_0.000130


metaRNA.PFemirge\_15194\_0.000074


EF098038.1.1349 Bacteria;Firmicutes;Clostridia;Lachnosp ... aceae;Lachnospiraceae NC2004 group;uncultured bacterium


metaRNA.PFemirge\_6032\_0.000096


metaRNA.PFemirge\_22745\_0.000012


metaRNA.PFemirge\_20578\_0.000005


metaRNA.PFemirge\_27568\_0.000007


metaRNA.PFemirge\_16721\_0.000004


metaRNA.PFemirge\_10325\_0.000006


metaRNA.PFemirge\_14552\_0.000054


metaRNA.PFemirge\_23902\_0.000002


AY126453.1.1511 Bacteria;Proteobacteria;Gammaproteobacteria;Burkholderiales;Rhodocyclaceae;Azospira;Azospira sp. PCC


metaRNA.PFemirge\_7425\_0.000007


AB447718.1.1450 Bacteria;Firmicutes;Clostridia;Oscillospirales;Oscillospiraceae;Colidextribacter;uncultured bacterium


metaRNA.PFemirge\_26246\_0.000009


metaRNA.PFemirge\_24245\_0.000002


metaRNA.PFemirge\_27653\_0.000009


metaRNA.PFemirge\_5519\_0.000005


metaRNA.PFemirge\_9475\_0.004681


AB612398.1.1470 Bacteria;Verrucomicrobiota;Lentisphaeria;Victivallales;Victivallaceae;uncultured rumen bacterium


metaRNA.PFemirge\_6206\_0.000006


KC189686.1.1429 Bacteria;Proteobacteria;Alphaproteobact ... rales;Rhodobacteraceae;Rhodobacter;uncultured bacterium


HQ777507.1.1443 Bacteria;Firmicutes;Clostridia;Lachnosp ... nospiraceae;Lachnospiraceae UCG-006;uncultured organism


JF830177.1.1510 Bacteria;Proteobacteria;Gammaproteobact ... adales;Moraxellaceae;Acinetobacter;uncultured bacterium


AB176219.1.1449 Bacteria;Proteobacteria;Alphaproteobact ... ;Sphingomonadaceae;Novosphingobium;uncultured bacterium


metaRNA.PFemirge\_17272\_0.000885


metaRNA.PFemirge\_12096\_0.000024


metaRNA.PFemirge\_21594\_0.000009


EU778828.1.1398 Bacteria;Bacteroidota;Bacteroidia;Bacteroidales;Tannerellaceae;Parabacteroides;uncultured bacterium


EU475681.1.1364 Bacteria;Firmicutes;Clostridia;Peptostr ... s;Peptostreptococcaceae;uncultured;uncultured bacterium


metaRNA.PFemirge\_20887\_0.000008


KT952743.1.1492 Bacteria;Bacteroidota;Bacteroidia;Bacteroidales;Rikenellaceae;Rikenella;uncultured bacterium


metaRNA.PFemirge\_12854\_0.001735


EF400740.1.1495 Bacteria;Firmicutes;Clostridia;Oscillospirales;Butyricicoccaceae;UCG-009;uncultured bacterium


EF688147.1.1469 Bacteria;Firmicutes;Clostridia;Oscillospirales;UCG-010;uncultured bacterium


metaRNA.PFemirge\_6783\_0.000035


metaRNA.PFemirge\_8965\_0.000036


metaRNA.PFemirge\_27913\_0.000002


HAFU01069247.55.1411 Bacteria;Proteobacteria;Gammaprote ... domonadaceae;Pseudomonas;Spumella-like flagellate JBL14


metaRNA.PFemirge\_9005\_0.009032


metaRNA.PFemirge\_27748\_0.000033


DQ307722.1.1396 Bacteria;Bacteroidota;Bacteroidia;Bacte ... enellaceae;Alistipes;uncultured Bacteroidetes bacterium


metaRNA.PFemirge\_30245\_0.000213


KR822473.1.1454 Bacteria;Firmicutes;Clostridia;Peptostr ... ovoracaceae;[Eubacterium] nodatum group;bacterium PM5-7


metaRNA.PFemirge\_20055\_0.000008


GU179922.1.1487 Bacteria;Firmicutes;Clostridia;Clostridia UCG-014;uncultured Firmicutes bacterium


LC028726.1.1525 Bacteria;Firmicutes;Clostridia;Lachnospirales;Lachnospiraceae;uncultured;uncultured bacterium


metaRNA.PFemirge\_11159\_0.000109


GQ448213.1.1432 Bacteria;Firmicutes;Clostridia;Lachnosp ... ceae;Lachnospiraceae NK4A136 group;uncultured bacterium


metaRNA.PFemirge\_21996\_0.000030


LC028724.1.1522 Bacteria;Firmicutes;Clostridia;Lachnosp ... aceae;Lachnospiraceae FCS020 group;uncultured bacterium


EU794122.1.1443 Bacteria;Firmicutes;Clostridia;Oscillos ... ccaceae;uncultured;uncultured Ruminococcaceae bacterium


metaRNA.PFemirge\_25519\_0.000007


metaRNA.PFemirge\_9100\_0.000003


metaRNA.PFemirge\_15445\_0.000011


metaRNA.PFemirge\_21215\_0.000007


AB307664.1.1358 Bacteria;Proteobacteria;Alphaproteobact ... les;Magnetospirillaceae;uncultured;uncultured bacterium


GQ871730.1.1506 Bacteria;Proteobacteria;Gammaproteobact ... dales;Pseudomonadaceae;Pseudomonas;uncultured bacterium


metaRNA.PFemirge\_23838\_0.000015


FPLS01027059.10.1473 Bacteria;Proteobacteria;Alphaproteobacteria;Rhizobiales;Devosiaceae;Devosia;metagenome


metaRNA.PFemirge\_13192\_0.000130


metaRNA.PFemirge\_16199\_0.000007


metaRNA.PFemirge\_16170\_0.000009


metaRNA.PFemirge\_21587\_0.001205


HQ761492.1.1450 Bacteria;Firmicutes;Clostridia;Lachnosp ... raceae;[Ruminococcus] torques group;uncultured organism


metaRNA.PFemirge\_6835\_0.000022


metaRNA.PFemirge\_16889\_0.000942


metaRNA.PFemirge\_12400\_0.000001


metaRNA.PFemirge\_2115\_0.000008


metaRNA.PFemirge\_7723\_0.000003


metaRNA.PFemirge\_26073\_0.000001


metaRNA.PFemirge\_27873\_0.002329


metaRNA.PFemirge\_2011\_0.000127


metaRNA.PFemirge\_24922\_0.000023


metaRNA.PFemirge\_7289\_0.000472


EU762199.1.1370 Bacteria;Bacteroidota;Bacteroidia;Bacteroidales;Bacteroidaceae;Bacteroides;uncultured bacterium


metaRNA.PFemirge\_837\_0.000003


EU937930.1.1450 Bacteria;Proteobacteria;Alphaproteobact ... ;Sphingomonadaceae;Novosphingobium;uncultured bacterium


metaRNA.PFemirge\_8230\_0.000014


DQ800244.1.1377 Bacteria;Firmicutes;Clostridia;Oscillospirales;Ruminococcaceae;Faecalibacterium;uncultured bacterium


DQ806157.1.1383 Bacteria;Firmicutes;Clostridia;Oscillospirales;Ruminococcaceae;Faecalibacterium;uncultured bacterium


metaRNA.PFemirge\_18144\_0.000007


KC961269.1.1495 Bacteria;Proteobacteria;Gammaproteobact ... ales;Comamonadaceae;Hydrogenophaga;uncultured bacterium


CDZJ01005172.1679.3161 Bacteria;Proteobacteria;Alphaproteobacteria;Rhodospirillales;uncultured;gut metagenome


metaRNA.PFemirge\_27167\_0.000019


metaRNA.PFemirge\_24538\_0.000003


metaRNA.PFemirge\_16223\_0.004950


metaRNA.PFemirge\_26216\_0.000003


DQ799380.1.1399 Bacteria;Bacteroidota;Bacteroidia;Bacteroidales;Rikenellaceae;Alistipes;uncultured bacterium


GQ359970.1.1475 Bacteria;Firmicutes;Clostridia;Clostrid ... iaceae;Clostridium sensu stricto 1;uncultured bacterium


metaRNA.PFemirge\_17236\_0.000028


metaRNA.PFemirge\_5136\_0.000008


metaRNA.PFemirge\_23516\_0.000004


FJ366372.1.1363 Bacteria;Bacteroidota;Bacteroidia;Bacteroidales;Rikenellaceae;Alistipes;uncultured bacterium


metaRNA.PFemirge\_16743\_0.000008


metaRNA.PFemirge\_26221\_0.000146


metaRNA.PFemirge\_4079\_0.000011


GQ867454.1.1492 Bacteria;Firmicutes;Clostridia;Oscillospirales;Ruminococcaceae;Fournierella;uncultured bacterium


metaRNA.PFemirge\_1297\_0.000008


DQ797856.1.1399 Bacteria;Bacteroidota;Bacteroidia;Bacteroidales;Rikenellaceae;Alistipes;uncultured bacterium


AB626942.1.1521 Bacteria;Firmicutes;Clostridia;Lachnosp ... ospiraceae;Lachnospiraceae UCG-008;uncultured bacterium


metaRNA.PFemirge\_11184\_0.000013


metaRNA.PFemirge\_22660\_0.000007


metaRNA.PFemirge\_12328\_0.000003


metaRNA.PFemirge\_11728\_0.000004


metaRNA.PFemirge\_24111\_0.000032


metaRNA.PFemirge\_15803\_0.000040


DQ816804.1.1464 Bacteria;Firmicutes;Clostridia;Peptostr ... lales;Anaerovoracaceae;Anaerovorax;uncultured bacterium


HQ767134.1.1448 Bacteria;Bacteroidota;Bacteroidia;Bacteroidales;Tannerellaceae;Parabacteroides;uncultured organism


AB554230.1.1487 Bacteria;Bacteroidota;Bacteroidia;Bacteroidales;Rikenellaceae;Alistipes;Alistipes finegoldii


EU772341.1.1387 Bacteria;Firmicutes;Clostridia;Oscillos ... bacterium] coprostanoligenes group;uncultured bacterium


EU454363.1.1392 Bacteria;Firmicutes;Clostridia;Lachnosp ... ceae;Lachnospiraceae NK4A136 group;uncultured bacterium


metaRNA.PFemirge\_25883\_0.000009


KX858536.1.1393 Bacteria;Proteobacteria;Gammaproteobacteria;Pseudomonadales;Pseudomonadaceae;Pseudomonas;Pseudomonas sp.


metaRNA.PFemirge\_8407\_0.000134


metaRNA.PFemirge\_23392\_0.000003


metaRNA.PFemirge\_16166\_0.000023


JQ083789.1.1511 Bacteria;Desulfobacterota;Desulfovibrio ... ales;Desulfovibrionaceae;Bilophila;uncultured bacterium


metaRNA.PFemirge\_27292\_0.000002


MG367100.1.1491 Bacteria;Firmicutes;Clostridia;Oscillos ... xtribacter;uncultured Clostridiales Family IV bacterium


metaRNA.PFemirge\_27279\_0.000031


metaRNA.PFemirge\_21756\_0.000003


metaRNA.PFemirge\_20908\_0.000012


EU475534.1.1401 Bacteria;Firmicutes;Clostridia;Lachnosp ... aceae;Lachnospiraceae AC2044 group;uncultured bacterium


HQ751117.1.1433 Bacteria;Firmicutes;Clostridia;Lachnosp ... nospiraceae;Lachnospiraceae UCG-004;uncultured organism


DQ794702.1.1380 Bacteria;Firmicutes;Clostridia;Lachnosp ... ospiraceae;Lachnospiraceae UCG-004;uncultured bacterium


metaRNA.PFemirge\_25115\_0.000008


AB916502.1.1486 Bacteria;Bacteroidota;Bacteroidia;Bacteroidales;Marinifilaceae;Butyricimonas;Butyricimonas paravirosa


metaRNA.PFemirge\_27994\_0.000003


metaRNA.PFemirge\_10860\_0.000017


metaRNA.PFemirge\_530\_0.000008


EU461103.1.1392 Bacteria;Firmicutes;Clostridia;Lachnosp ... ospiraceae;Lachnospiraceae UCG-008;uncultured bacterium


metaRNA.PFemirge\_21700\_0.000030


KP866212.1.1451 Bacteria;Bacteroidota;Bacteroidia;Flavo ... obacteriaceae;Flavobacterium;Flavobacterium sp. HMF3121


metaRNA.PFemirge\_4539\_0.000156


metaRNA.PFemirge\_16332\_0.000026


EU464332.1.1399 Bacteria;Bacteroidota;Bacteroidia;Bacteroidales;Rikenellaceae;dgA-11 gut group;uncultured bacterium


metaRNA.PFemirge\_23347\_0.000124


metaRNA.PFemirge\_13214\_0.000008


MH699149.1.1471 Bacteria;Actinobacteriota;Actinobacteri ... ales;Microbacteriaceae;Microbacterium;Mycobacterium sp.


metaRNA.PFemirge\_26618\_0.000000


metaRNA.PFemirge\_20401\_0.000002


EU616637.1.1399 Bacteria;Bacteroidota;Bacteroidia;Bacte ... enellaceae;Rikenella;uncultured Rikenellaceae bacterium


metaRNA.PFemirge\_18121\_0.000002


JN038827.1.1489 Bacteria;Proteobacteria;Gammaproteobact ... tterellaceae;uncultured;uncultured beta proteobacterium


metaRNA.PFemirge\_6963\_0.000087


MF085166.1.1529 Bacteria;Proteobacteria;Gammaproteobact ... les;Acidithiobacillaceae;KCM-B-112;uncultured bacterium


AY328682.1.1446 Bacteria;Proteobacteria;Alphaproteobact ... terales;Hyphomonadaceae;uncultured;uncultured bacterium


metaRNA.PFemirge\_25146\_0.000005


metaRNA.PFemirge\_19506\_0.000005


metaRNA.PFemirge\_1478\_0.000009


metaRNA.PFemirge\_14075\_0.000005


metaRNA.PFtrusted\_12\_0


metaRNA.PFemirge\_8590\_0.000001


metaRNA.PFemirge\_16440\_0.000003


metaRNA.PFemirge\_11245\_0.000110


EU939391.1.1539 Bacteria;Firmicutes;Clostridia;Eubacteriales;Eubacteriaceae;Eubacterium;uncultured bacterium


metaRNA.PFemirge\_7284\_0.000002


metaRNA.PFemirge\_13924\_0.000022


metaRNA.PFemirge\_27638\_0.000027


metaRNA.PFemirge\_16686\_0.000010


metaRNA.PFemirge\_8147\_0.000037


metaRNA.PFemirge\_7544\_0.000820


KX465194.1.1643 Eukaryota;SAR;Alveolata;Ciliophora;Intr ... itostomatea;Haptoria;Arcuospathidium;uncultured ciliate


metaRNA.PFemirge\_18350\_0.000004


metaRNA.PFemirge\_13831\_0.000002


JX222921.1.1476 Bacteria;Firmicutes;Clostridia;Peptostr ... lales;Anaerovoracaceae;Anaerovorax;uncultured bacterium


metaRNA.PFemirge\_3064\_0.000093


metaRNA.PFemirge\_3901\_0.000008


metaRNA.PFemirge\_9657\_0.000599


metaRNA.PFemirge\_27687\_0.004843


FJ716048.1.1407 Bacteria;Proteobacteria;Gammaproteobact ... es;Enterobacteriaceae;Buttiauxella;uncultured bacterium


metaRNA.PFemirge\_17111\_0.000035


metaRNA.PFemirge\_3096\_0.000003


KC994754.1.1487 Bacteria;Actinobacteriota;Actinobacteri ... eriaceae;Agromyces;uncultured Actinomycetales bacterium


metaRNA.PFemirge\_27715\_0.001416


metaRNA.PFemirge\_15548\_0.000132


metaRNA.PFemirge\_23625\_0.000016


metaRNA.PFemirge\_10528\_0.000144


metaRNA.PFemirge\_4063\_0.000001


metaRNA.PFemirge\_8912\_0.000005


metaRNA.PFemirge\_9376\_0.000013


JX457157.1.1486 Bacteria;Bacteroidota;Bacteroidia;Bacteroidales;Tannerellaceae;Parabacteroides;uncultured bacterium


KF441638.1.1495 Bacteria;Bacteroidota;Bacteroidia;Flavo ... vobacteriaceae;Flavobacterium;Flavobacterium sp. 7A-207


metaRNA.PFemirge\_28093\_0.000089


metaRNA.PFemirge\_20621\_0.000002


metaRNA.PFemirge\_28223\_0.009837


metaRNA.PFemirge\_28188\_0.000023


metaRNA.PFemirge\_27927\_0.000007


metaRNA.PFemirge\_16552\_0.000664


metaRNA.PFemirge\_23431\_0.000015


metaRNA.PFemirge\_6921\_0.000052


EU775320.1.1281 Bacteria;Firmicutes;Clostridia;Oscillospirales;Ruminococcaceae;Ruminococcus;uncultured bacterium


KF104611.1.1373 Bacteria;Proteobacteria;Gammaproteobact ... ;Xanthomonadaceae;Stenotrophomonas;uncultured bacterium


metaRNA.PFemirge\_5586\_0.000016


metaRNA.PFemirge\_13159\_0.000108


metaRNA.PFemirge\_12069\_0.000009


metaRNA.PFemirge\_17372\_0.004551


metaRNA.PFemirge\_15835\_0.002820


metaRNA.PFemirge\_588\_0.005742


metaRNA.PFemirge\_6359\_0.000004


metaRNA.PFemirge\_26515\_0.000007


metaRNA.PFemirge\_5919\_0.000002


GDHD01010040.119.1623 Bacteria;Cyanobacteria;Cyanobacteriia;Chloroplast;Cymbidium faberi


metaRNA.PFemirge\_22900\_0.000058


metaRNA.PFemirge\_23861\_0.000009


metaRNA.PFemirge\_16925\_0.000100


metaRNA.PFemirge\_6413\_0.000035


metaRNA.PFemirge\_3258\_0.000008


metaRNA.PFemirge\_22967\_0.000002


DQ793254.1.1377 Bacteria;Firmicutes;Clostridia;Oscillospirales;Ruminococcaceae;Faecalibacterium;uncultured bacterium


metaRNA.PFemirge\_5654\_0.000047


metaRNA.PFemirge\_12765\_0.000006


metaRNA.PFemirge\_23557\_0.000007


metaRNA.PFemirge\_14760\_0.000012


metaRNA.PFemirge\_12583\_0.000152


metaRNA.PFemirge\_16878\_0.000013


metaRNA.PFemirge\_18878\_0.000005


metaRNA.PFemirge\_10289\_0.000002


metaRNA.PFemirge\_27775\_0.000338


metaRNA.PFemirge\_24591\_0.002806


metaRNA.PFemirge\_23719\_0.000008


metaRNA.PFemirge\_25380\_0.000002


metaRNA.PFemirge\_1864\_0.000004


EU773965.1.1389 Bacteria;Firmicutes;Clostridia;Lachnosp ... aceae;[Ruminococcus] torques group;uncultured bacterium


GQ867427.1.1505 Bacteria;Proteobacteria;Gammaproteobact ... obacteriaceae;Escherichia-Shigella;uncultured bacterium


metaRNA.PFemirge\_12250\_0.000656


KM650565.1.1462 Bacteria;Desulfobacterota;Desulfovibrio ... fovibrionaceae;Desulfovibrio;uncultured proteobacterium


metaRNA.PFemirge\_12524\_0.000118


metaRNA.PFemirge\_16774\_0.000075


KX589545.1.1498 Bacteria;Firmicutes;Bacilli;Alicyclobacillales;Alicyclobacillaceae;Tumebacillus;uncultured bacterium


HQ804310.1.1461 Bacteria;Firmicutes;Clostridia;Lachnosp ... nospiraceae;Lachnospiraceae UCG-004;uncultured organism


DQ807369.1.1362 Bacteria;Firmicutes;Clostridia;Peptostr ... s;Peptostreptococcaceae;uncultured;uncultured bacterium


CBVB010000006.254456.255954 Bacteria;Bacteroidota;Bacte ... Bacteroidaceae;Bacteroides;Bacteroidaceae bacterium MS4


LARG01000025.1.1402 Bacteria;Bacteroidota;Bacteroidia;Bacteroidales;Rikenellaceae;Alistipes;Alistipes sp. N15.MGS-157


DQ800642.1.1301 Bacteria;Firmicutes;Clostridia;Christen ... ceae;Christensenellaceae R-7 group;uncultured bacterium


metaRNA.PFemirge\_27385\_0.000009


HQ813979.1.1426 Bacteria;Firmicutes;Clostridia;Lachnospirales;Lachnospiraceae;uncultured;uncultured organism


metaRNA.PFemirge\_18613\_0.000046


JQ820153.1.1498 Bacteria;Firmicutes;Clostridia;Oscillospirales;Oscillospiraceae;uncultured;uncultured bacterium


GQ389185.1.1501 Bacteria;Proteobacteria;Gammaproteobact ... dales;Pseudomonadaceae;Pseudomonas;uncultured bacterium


metaRNA.PFemirge\_15917\_0.000003


metaRNA.PFemirge\_3434\_0.000815


metaRNA.PFemirge\_26086\_0.000005


metaRNA.PFemirge\_25536\_0.000062


metaRNA.PFemirge\_9252\_0.000096


metaRNA.PFemirge\_28411\_0.000010


metaRNA.PFemirge\_21967\_0.000007


HQ740432.1.1447 Bacteria;Bacteroidota;Bacteroidia;Bacteroidales;Marinifilaceae;Odoribacter;uncultured organism


metaRNA.PFemirge\_2567\_0.000039


metaRNA.PFemirge\_27990\_0.000007


metaRNA.PFemirge\_16722\_0.000315


metaRNA.PFemirge\_20640\_0.000002


metaRNA.PFemirge\_10801\_0.000001


metaRNA.PFemirge\_6069\_0.003151


metaRNA.PFemirge\_17352\_0.000022


metaRNA.PFemirge\_3979\_0.000007


DQ825005.1.1403 Bacteria;Bacteroidota;Bacteroidia;Bacteroidales;Bacteroidaceae;Bacteroides;uncultured bacterium


metaRNA.PFemirge\_24271\_0.000007


metaRNA.PFemirge\_21922\_0.000006


KC331222.1.1485 Bacteria;Actinobacteriota;Actinobacteri ... ales;Microbacteriaceae;Leucobacter;uncultured bacterium


metaRNA.PFemirge\_4467\_0.000005


metaRNA.PFemirge\_8156\_0.000003


GQ448730.1.1400 Bacteria;Bacteroidota;Bacteroidia;Bacteroidales;Bacteroidaceae;Bacteroides;uncultured bacterium


metaRNA.PFemirge\_16100\_0.000002


metaRNA.PFemirge\_3525\_0.000015


metaRNA.PFemirge\_21077\_0.000002


metaRNA.PFemirge\_13494\_0.000061


metaRNA.PFemirge\_23135\_0.000009


KY190699.1.1411 Bacteria;Proteobacteria;Alphaproteobact ... les;Sphingomonadaceae;Sphingomonas;uncultured bacterium


metaRNA.PFemirge\_16518\_0.000033


metaRNA.PFemirge\_21547\_0.000410


HQ781428.1.1455 Bacteria;Firmicutes;Clostridia;Lachnospirales;Lachnospiraceae;uncultured;uncultured organism


metaRNA.PFemirge\_4671\_0.000009


HM159970.1.1478 Bacteria;Proteobacteria;Gammaproteobact ... rales;Aeromonadaceae;Aeromonas;uncultured Aeromonas sp.


EU772665.1.1358 Bacteria;Firmicutes;Clostridia;Lachnosp ... ospiraceae;Lachnospiraceae UCG-008;uncultured bacterium


metaRNA.PFemirge\_21052\_0.000003


metaRNA.PFemirge\_3510\_0.000010


metaRNA.PFemirge\_21456\_0.000004


EU778073.1.1322 Bacteria;Bacteroidota;Bacteroidia;Bacteroidales;Rikenellaceae;Alistipes;uncultured bacterium


DQ800771.1.1398 Bacteria;Proteobacteria;Gammaproteobact ... obacteriaceae;Escherichia-Shigella;uncultured bacterium


metaRNA.PFemirge\_16294\_0.010316


GCKC01172265.99.1551 Bacteria;Proteobacteria;Alphaproteobacteria;Rickettsiales;Mitochondria;Gleditsia sinensis


metaRNA.PFemirge\_13062\_0.000607


MG321612.1.1372 Bacteria;Bacteroidota;Bacteroidia;Bacteroidales;Tannerellaceae;Parabacteroides;Parabacteroides sp.


HQ806032.1.1447 Bacteria;Bacteroidota;Bacteroidia;Bacteroidales;Tannerellaceae;Parabacteroides;uncultured organism


metaRNA.PFemirge\_12730\_0.000104


DQ456478.1.1451 Bacteria;Firmicutes;Clostridia;Oscillospirales;Oscillospiraceae;Colidextribacter;uncultured bacterium


metaRNA.PFemirge\_5673\_0.000025


metaRNA.PFemirge\_16700\_0.000000


CR933278.1.1273 Bacteria;Firmicutes;Clostridia;Lachnosp ... ospiraceae;Lachnospiraceae UCG-004;uncultured bacterium


metaRNA.PFspades\_30\_283.402893


DQ799601.1.1400 Bacteria;Bacteroidota;Bacteroidia;Bacteroidales;Bacteroidaceae;Bacteroides;uncultured bacterium


AB610559.1.1481 Bacteria;Firmicutes;Clostridia;Lachnospirales;Lachnospiraceae;Lachnoclostridium;Clostridium sp. K7


metaRNA.PFemirge\_6452\_0.000408


DQ797886.1.1381 Bacteria;Firmicutes;Clostridia;Lachnosp ... ospiraceae;Lachnospiraceae UCG-004;uncultured bacterium


metaRNA.PFemirge\_3265\_0.000001


metaRNA.PFemirge\_20181\_0.000003


metaRNA.PFemirge\_12036\_0.000018


U14932.1.1433 Bacteria;Spirochaetota;Brachyspirae;Brachyspirales;Brachyspiraceae;Brachyspira;Brachyspira hyodysenteriae


metaRNA.PFemirge\_24781\_0.000004


DQ681227.1.1525 Bacteria;Firmicutes;Clostridia;Lachnospirales;Lachnospiraceae;Robinsoniella;Robinsoniella peoriensis


metaRNA.PFemirge\_9609\_0.000017


metaRNA.PFemirge\_22505\_0.000009


metaRNA.PFemirge\_11872\_0.000003


metaRNA.PFemirge\_2664\_0.000003


metaRNA.PFemirge\_22396\_0.000054


metaRNA.PFemirge\_12601\_0.000023


EU773113.1.1391 Bacteria;Bacteroidota;Bacteroidia;Bacte ... laceae;Rikenellaceae RC9 gut group;uncultured bacterium


metaRNA.PFemirge\_25223\_0.000006


metaRNA.PFemirge\_15940\_0.000017


metaRNA.PFemirge\_15432\_0.000003


metaRNA.PFemirge\_16208\_0.000135


metaRNA.PFemirge\_2017\_0.000013


metaRNA.PFemirge\_23882\_0.000006


metaRNA.PFemirge\_20171\_0.000000


metaRNA.PFemirge\_19766\_0.000003


metaRNA.PFemirge\_3029\_0.000135


DQ339713.1.1435 Bacteria;Firmicutes;Clostridia;Clostrid ... aceae;Clostridium sensu stricto 13;uncultured bacterium


DQ793376.1.1284 Bacteria;Firmicutes;Clostridia;Oscillospirales;Butyricicoccaceae;Butyricicoccus;uncultured bacterium


metaRNA.PFemirge\_6702\_0.001703


KM650266.1.1454 Bacteria;Firmicutes;Clostridia;Oscillospirales;Oscillospiraceae;UCG-002;uncultured proteobacterium


metaRNA.PFemirge\_5371\_0.000002


metaRNA.PFemirge\_26250\_0.000011


metaRNA.PFemirge\_2228\_0.000010


metaRNA.PFemirge\_25608\_0.000012


metaRNA.PFemirge\_15469\_0.000008


EU850512.1.1455 Bacteria;Firmicutes;Clostridia;Oscillospirales;Ruminococcaceae;Ruminococcus;uncultured rumen bacterium


DQ799526.1.1402 Bacteria;Bacteroidota;Bacteroidia;Bacteroidales;Bacteroidaceae;Bacteroides;uncultured bacterium


metaRNA.PFemirge\_18465\_0.000004


metaRNA.PFemirge\_2917\_0.057600


metaRNA.PFemirge\_7151\_0.000042


HQ741595.1.1456 Bacteria;Firmicutes;Clostridia;Lachnospirales;Lachnospiraceae;Epulopiscium;uncultured bacterium


EU828415.1.1454 Bacteria;Proteobacteria;Gammaproteobact ... Oxalobacteraceae;Janthinobacterium;uncultured bacterium


metaRNA.PFemirge\_16362\_0.000010


metaRNA.PFemirge\_25820\_0.000001


metaRNA.PFemirge\_27926\_0.000006


metaRNA.PFemirge\_26214\_0.000007


metaRNA.PFspades\_35\_119.998907


metaRNA.PFemirge\_24794\_0.000006


metaRNA.PFemirge\_5628\_0.000045


metaRNA.PFemirge\_6295\_0.000755


metaRNA.PFemirge\_26187\_0.000008


metaRNA.PFemirge\_1849\_0.000866


metaRNA.PFemirge\_13064\_0.000029


JN680575.1.1450 Bacteria;Bacteroidota;Bacteroidia;Bacte ... ae;Dysgonomonas;uncultured Porphyromonadaceae bacterium


metaRNA.PFemirge\_26046\_0.000009


EU505366.1.1398 Bacteria;Bacteroidota;Bacteroidia;Bacteroidales;Tannerellaceae;Parabacteroides;uncultured bacterium


metaRNA.PFemirge\_22995\_0.000180


AF479766.1.2092 Bacteria;Proteobacteria;Gammaproteobact ... iales;Rhodocyclaceae;Dechloromonas;Dechloromonas sp. HZ


metaRNA.PFemirge\_14707\_0.000621


metaRNA.PFemirge\_18865\_0.000003


AB294338.1.1433 Bacteria;Bacteroidota;Bacteroidia;Cytophagales;Cytophagaceae;Cytophaga;uncultured bacterium


metaRNA.PFemirge\_28079\_0.000005


metaRNA.PFemirge\_23138\_0.000015


metaRNA.PFemirge\_6249\_0.005745


KU667224.1.1485 Bacteria;Bacteroidota;Bacteroidia;Bacteroidales;Rikenellaceae;uncultured;uncultured bacterium


metaRNA.PFemirge\_25956\_0.000062


metaRNA.PFemirge\_2702\_0.000002


metaRNA.PFemirge\_5798\_0.000004


GQ448690.1.1389 Bacteria;Firmicutes;Clostridia;Lachnosp ... ;[Eubacterium] oxidoreducens group;uncultured bacterium


metaRNA.PFemirge\_19219\_0.000015


metaRNA.PFemirge\_23226\_0.000034


metaRNA.PFemirge\_21042\_0.000014


metaRNA.PFtrusted\_11\_0


JX457238.1.1481 Bacteria;Bacteroidota;Bacteroidia;Bacteroidales;Rikenellaceae;Alistipes;uncultured bacterium


metaRNA.PFemirge\_12067\_0.000096


metaRNA.PFemirge\_26597\_0.000001


metaRNA.PFemirge\_7262\_0.000361


KC748467.1.1433 Bacteria;Firmicutes;Bacilli;Lactobacillales;Streptococcaceae;Streptococcus;Streptococcus iniae


metaRNA.PFemirge\_27811\_0.000524


KP182152.1.1434 Bacteria;Proteobacteria;Alphaproteobact ... Incertae Sedis;Phreatobacter;Phreatobacter sp. 61DPR27


JQ248079.1.1513 Bacteria;Firmicutes;Clostridia;Oscillospirales;Oscillospiraceae;Flavonifractor;uncultured bacterium


metaRNA.PFemirge\_6740\_0.000168


metaRNA.PFemirge\_171\_0.000565


metaRNA.PFemirge\_62\_0.000158


metaRNA.PFemirge\_9241\_0.000060


DQ810172.1.1365 Bacteria;Firmicutes;Bacilli;Erysipelotrichales;Erysipelotrichaceae;Dielma;uncultured bacterium


metaRNA.PFemirge\_3735\_0.000836


metaRNA.PFemirge\_11448\_0.003731


DQ807222.1.1401 Bacteria;Bacteroidota;Bacteroidia;Bacteroidales;Bacteroidaceae;Bacteroides;uncultured bacterium


metaRNA.PFemirge\_21127\_0.000013


EU775304.1.1384 Bacteria;Firmicutes;Clostridia;Lachnosp ... ospiraceae;Lachnospiraceae UCG-008;uncultured bacterium


metaRNA.PFemirge\_16188\_0.000021


metaRNA.PFemirge\_26979\_0.000065


metaRNA.PFemirge\_16724\_0.000006


DQ799557.1.1402 Bacteria;Bacteroidota;Bacteroidia;Bacteroidales;Bacteroidaceae;Bacteroides;uncultured bacterium


metaRNA.PFemirge\_19434\_0.000001


metaRNA.PFemirge\_30265\_0.000015


metaRNA.PFemirge\_13003\_0.000004


CABV01003337.48680.50227 Bacteria;Verrucomicrobiota;Ver ... robiales;Rubritaleaceae;Luteolibacter;Oikopleura dioica


metaRNA.PFemirge\_20446\_0.000003


GQ480160.1.1510 Bacteria;Firmicutes;Negativicutes;Acidaminococcales;Acidaminococcaceae;Succinispira;uncultured bacterium


metaRNA.PFemirge\_5097\_0.000004


metaRNA.PFemirge\_14766\_0.000059


metaRNA.PFemirge\_14514\_0.000032


metaRNA.PFemirge\_8284\_0.000003


LC028597.1.1518 Bacteria;Bacteroidota;Bacteroidia;Bacteroidales;Rikenellaceae;dgA-11 gut group;uncultured bacterium


metaRNA.PFemirge\_28106\_0.000113


metaRNA.PFemirge\_30259\_0.000380


metaRNA.PFemirge\_17678\_0.000001


GQ448680.1.1388 Bacteria;Firmicutes;Clostridia;Lachnospirales;Lachnospiraceae;uncultured bacterium


metaRNA.PFemirge\_15909\_0.000018


metaRNA.PFemirge\_5247\_0.000004


metaRNA.PFemirge\_27764\_0.000008


metaRNA.PFemirge\_16960\_0.000007


metaRNA.PFemirge\_23063\_0.000006


metaRNA.PFemirge\_27691\_0.017422


metaRNA.PFemirge\_20555\_0.000182


metaRNA.PFemirge\_3004\_0.000002


JX105679.1.1392 Bacteria;Bacteroidota;Bacteroidia;Bacteroidales;Bacteroidaceae;Bacteroides;uncultured bacterium


metaRNA.PFemirge\_6802\_0.000938


metaRNA.PFemirge\_23597\_0.000021


metaRNA.PFemirge\_18543\_0.000023


AB934800.1.1322 Bacteria;Proteobacteria;Alphaproteobacteria;Rhodospirillales;uncultured;uncultured bacterium


JN547325.1.1576 Eukaryota;Amorphea;Amoebozoa;Discosea;Flabellinia;Dactylopodida;Angulamoeba;uncultured eukaryote


metaRNA.PFemirge\_23234\_0.000001


EU772882.1.1401 Bacteria;Firmicutes;Clostridia;Oscillospirales;Butyricicoccaceae;uncultured;uncultured bacterium


CP011612.1889203.1890758 Bacteria;Proteobacteria;Gammap ... les;Enterobacteriaceae;Citrobacter;Citrobacter freundii


FM177879.1.1437 Bacteria;Proteobacteria;Alphaproteobacteria;Rhizobiales;Rhizobiaceae;Shinella;Shinella fusca


metaRNA.PFemirge\_5576\_0.000002


metaRNA.PFemirge\_4769\_0.002101


metaRNA.PFemirge\_8057\_0.000005


metaRNA.PFemirge\_7726\_0.000003


metaRNA.PFemirge\_23962\_0.000004


metaRNA.PFemirge\_5707\_0.000006


KF773008.1.1446 Bacteria;Firmicutes;Clostridia;Lachnospirales;Lachnospiraceae;Hungatella;uncultured bacterium


EU775261.1.1286 Bacteria;Firmicutes;Clostridia;Lachnospirales;Lachnospiraceae;Agathobacter;uncultured bacterium


ABYH01000014.137.1655 Bacteria;Bacteroidota;Bacteroidia ... eae;Parabacteroides;Parabacteroides johnsonii DSM 18315


HAEN01058188.41.1551 Bacteria;Proteobacteria;Gammaprote ... eriales;Comamonadaceae;Rhizobacter;Bragasellus peltatus


metaRNA.PFemirge\_7175\_0.000004


metaRNA.PFemirge\_22117\_0.000063


metaRNA.PFemirge\_23903\_0.000006


EU511065.1.1410 Bacteria;Firmicutes;Clostridia;Oscillospirales;Oscillospiraceae;uncultured;uncultured bacterium


metaRNA.PFemirge\_9441\_0.000053


metaRNA.PFemirge\_16589\_0.000005


JN391619.1.1506 Bacteria;Proteobacteria;Gammaproteobacteria;Burkholderiales;Rhodocyclaceae;Thauera;uncultured bacterium


AM490748.1.1504 Bacteria;Myxococcota;Polyangia;mle1-27;uncultured bacterium


metaRNA.PFemirge\_15660\_0.004618


DQ809267.1.1395 Bacteria;Bacteroidota;Bacteroidia;Bacteroidales;Marinifilaceae;Butyricimonas;uncultured bacterium


metaRNA.PFspades\_38\_79.249441


metaRNA.PFemirge\_25233\_0.000006


metaRNA.PFemirge\_23233\_0.000006


metaRNA.PFemirge\_6071\_0.001656


metaRNA.PFspades\_5\_75.033820


metaRNA.PFemirge\_812\_0.001165


metaRNA.PFemirge\_27169\_0.000020


metaRNA.PFemirge\_26690\_0.000003


EF644509.1.1469 Bacteria;Firmicutes;Clostridia;Oscillospirales;Oscillospiraceae;Colidextribacter;uncultured bacterium


metaRNA.PFemirge\_23419\_0.000002


HQ761452.1.1440 Bacteria;Bacteroidota;Bacteroidia;Bacteroidales;Tannerellaceae;Parabacteroides;uncultured organism


metaRNA.PFemirge\_26413\_0.000001


metaRNA.PFemirge\_9202\_0.000233


metaRNA.PFemirge\_26268\_0.000097


metaRNA.PFemirge\_21475\_0.000087


metaRNA.PFemirge\_7296\_0.000138


DQ805627.1.1385 Bacteria;Firmicutes;Clostridia;Lachnosp ... ospiraceae;Lachnospiraceae UCG-004;uncultured bacterium


metaRNA.PFspades\_2\_1753.138489


metaRNA.PFemirge\_7308\_0.000010


HM124083.1.1490 Bacteria;Bacteroidota;Bacteroidia;Bacte ... llaceae;Prevotellaceae Ga6A1 group;uncultured bacterium


metaRNA.PFemirge\_17542\_0.001328


metaRNA.PFemirge\_12464\_0.000497


EU775041.1.1395 Bacteria;Bacteroidota;Bacteroidia;Bacteroidales;Bacteroidaceae;Bacteroides;uncultured bacterium


CP016102.16052.17542 Bacteria;Firmicutes;Clostridia;Pep ... treptococcaceae;Clostridioides;Clostridioides difficile


metaRNA.PFemirge\_6842\_0.000023


metaRNA.PFemirge\_17871\_0.000036


DQ810138.1.1391 Bacteria;Firmicutes;Clostridia;Lachnospirales;Lachnospiraceae;Agathobacter;uncultured bacterium


metaRNA.PFemirge\_14306\_0.000004


metaRNA.PFspades\_41\_161.204363


metaRNA.PFemirge\_2779\_0.000359


AY693826.1.1487 Bacteria;Bacteroidota;Bacteroidia;Bacteroidales;Rikenellaceae;uncultured;uncultured bacterium


metaRNA.PFemirge\_28083\_0.000016


metaRNA.PFemirge\_19894\_0.000009


metaRNA.PFemirge\_13650\_0.000076


EU509593.1.1402 Bacteria;Desulfobacterota;Desulfovibrio ... les;Desulfovibrionaceae;uncultured;uncultured bacterium


metaRNA.PFemirge\_26764\_0.000004


metaRNA.PFemirge\_22720\_0.000001


CEAB01046477.78830.80315 Bacteria;Proteobacteria;Alphaproteobacteria;Rhodospirillales;uncultured;gut metagenome


EU776025.1.1313 Bacteria;Firmicutes;Bacilli;Erysipelotrichales;Erysipelotrichaceae;Holdemanella;uncultured bacterium


DQ796132.1.1379 Bacteria;Firmicutes;Clostridia;Oscillospirales;Ruminococcaceae;Faecalibacterium;uncultured bacterium


DI433844.4.1378 Bacteria;Firmicutes;Bacilli;Bacillales;Bacillaceae;Bacillus;Bacillus sp. EBW4


DQ905437.1.1471 Bacteria;Firmicutes;Clostridia;Oscillospirales;Ruminococcaceae;uncultured;uncultured bacterium


metaRNA.PFemirge\_26639\_0.000029


metaRNA.PFemirge\_20677\_0.000001


DQ456069.1.1449 Bacteria;Bacteroidota;Bacteroidia;Bacteroidales;Bacteroidaceae;Bacteroides;uncultured bacterium


metaRNA.PFemirge\_12879\_0.002607


FJ269095.1.1502 Bacteria;Bacteroidota;Bacteroidia;Bacte ... on-reducing bacterium enrichment culture clone HN-HFO47


metaRNA.PFemirge\_22551\_0.000002


DQ804757.1.1391 Bacteria;Firmicutes;Clostridia;Lachnosp ... aceae;[Ruminococcus] torques group;uncultured bacterium


metaRNA.PFemirge\_4967\_0.000011


metaRNA.PFemirge\_23053\_0.000013


EU772907.1.1401 Bacteria;Bacteroidota;Bacteroidia;Bacteroidales;Bacteroidaceae;Bacteroides;uncultured bacterium


metaRNA.PFemirge\_19102\_0.000009


metaRNA.PFemirge\_14600\_0.000012


metaRNA.PFemirge\_14304\_0.000005


metaRNA.PFemirge\_16163\_0.000017


metaRNA.PFemirge\_99\_0.000013


metaRNA.PFemirge\_22806\_0.000050


metaRNA.PFemirge\_6039\_0.000506


JX223620.1.1368 Bacteria;Firmicutes;Clostridia;Clostrid ... aceae;Clostridium sensu stricto 12;uncultured bacterium


EF404130.1.1478 Bacteria;Firmicutes;Clostridia;Oscillospirales;Ruminococcaceae;Ruminococcus;uncultured bacterium


AY321958.1.1429 Bacteria;Actinobacteriota;Coriobacterii ... riales;Eggerthellaceae;Eggerthella;Eggerthella sinensis


metaRNA.PFemirge\_17925\_0.003077


LN879502.2473726.2475252 Bacteria;Verrucomicrobiota;Chl ... Protochlamydia;Candidatus Protochlamydia naegleriophila


metaRNA.PFemirge\_27964\_0.000571


KC669122.1.1486 Bacteria;Firmicutes;Clostridia;Lachnospirales;Lachnospiraceae;Epulopiscium;uncultured bacterium


HQ793579.1.1446 Bacteria;Bacteroidota;Bacteroidia;Bacteroidales;Marinifilaceae;Butyricimonas;uncultured organism


KP687196.1.1423 Bacteria;Proteobacteria;Alphaproteobacteria;Holosporales;Holosporaceae;uncultured;uncultured bacterium


metaRNA.PFemirge\_2883\_0.000311


metaRNA.PFemirge\_18550\_0.000031


metaRNA.PFemirge\_16483\_0.000024


metaRNA.PFemirge\_4839\_0.000003


metaRNA.PFemirge\_18207\_0.000010


metaRNA.PFemirge\_2466\_0.000003


metaRNA.PFemirge\_27729\_0.000013


metaRNA.PFemirge\_6401\_0.000005


metaRNA.PFemirge\_22890\_0.000077


metaRNA.PFemirge\_12515\_0.001993


HQ815341.1.1435 Bacteria;Firmicutes;Clostridia;Peptostr ... ptostreptococcaceae;Intestinibacter;uncultured organism


EU775986.1.1401 Bacteria;Bacteroidota;Bacteroidia;Bacteroidales;Bacteroidaceae;Bacteroides;uncultured bacterium


EU509830.1.1376 Bacteria;Firmicutes;Clostridia;Oscillospirales;Ruminococcaceae;Ruminococcus;uncultured bacterium


CP008712.314129.315667 Bacteria;Firmicutes;Bacilli;Bacillales;Bacillaceae;Bacillus;Bacillus cereus


EU765827.1.1360 Bacteria;Bacteroidota;Bacteroidia;Bacteroidales;Bacteroidaceae;Bacteroides;uncultured bacterium


metaRNA.PFemirge\_17917\_0.000030


HQ775105.1.1456 Bacteria;Firmicutes;Clostridia;Lachnospirales;Lachnospiraceae;Hungatella;uncultured organism


FJ366291.1.1363 Bacteria;Firmicutes;Clostridia;Oscillospirales;Oscillospiraceae;UCG-002;uncultured bacterium


EU777807.1.1400 Bacteria;Proteobacteria;Gammaproteobact ... obacteriaceae;Escherichia-Shigella;uncultured bacterium


KT952840.1.1494 Bacteria;Firmicutes;Clostridia;Oscillospirales;Oscillospiraceae;Flavonifractor;uncultured bacterium


metaRNA.PFemirge\_6437\_0.000006


KU978243.1.1517 Bacteria;Proteobacteria;Gammaproteobact ... dales;Pseudomonadaceae;Pseudomonas;uncultured bacterium


metaRNA.PFemirge\_26379\_0.003009


metaRNA.PFemirge\_23074\_0.000021


AY945907.1.1506 Bacteria;Proteobacteria;Gammaproteobact ... ales;Rhodocyclaceae;Propionivibrio;uncultured bacterium


metaRNA.PFemirge\_11902\_0.000016


KC311367.1.1475 Bacteria;Firmicutes;Clostridia;Oscillos ... iraceae;Intestinimonas;Intestinimonas butyriciproducens


metaRNA.PFemirge\_1920\_0.000512


metaRNA.PFemirge\_20156\_0.000004


HE575399.1.1900 Eukaryota;Amorphea;Amoebozoa;Incertae Sedis;Ischnamoeba;uncultured Lobosea


metaRNA.PFemirge\_6635\_0.084218


EU765284.1.1347 Bacteria;Firmicutes;Clostridia;Lachnosp ... ospiraceae;Lachnospiraceae UCG-008;uncultured bacterium


metaRNA.PFemirge\_19175\_0.000002


metaRNA.PFemirge\_22972\_0.000009


metaRNA.PFemirge\_17335\_0.001368


HQ748426.1.1449 Bacteria;Firmicutes;Clostridia;Lachnospirales;Lachnospiraceae;Lachnoclostridium;uncultured organism


metaRNA.PFemirge\_9349\_0.000138


metaRNA.PFemirge\_3793\_0.001333


HQ802642.1.1374 Bacteria;Firmicutes;Clostridia;Lachnospirales;Lachnospiraceae;Epulopiscium;uncultured organism


metaRNA.PFemirge\_4037\_0.000034


metaRNA.PFemirge\_9931\_0.000007


metaRNA.PFemirge\_6164\_0.000004


metaRNA.PFspades\_13\_11.888227


metaRNA.PFemirge\_18586\_0.000031


metaRNA.PFemirge\_23328\_0.000250


metaRNA.PFemirge\_1105\_0.000501


metaRNA.PFemirge\_30235\_0.000216


metaRNA.PFemirge\_12489\_0.000063


metaRNA.PFemirge\_27730\_0.000033


metaRNA.PFemirge\_20037\_0.000002


metaRNA.PFemirge\_21658\_0.000263


metaRNA.PFemirge\_15318\_0.003179


JN391753.1.1499 Bacteria;Proteobacteria;Gammaproteobact ... deriales;Comamonadaceae;uncultured;uncultured bacterium


AB574482.1.1489 Bacteria;Bacteroidota;Bacteroidia;Bacte ... nerellaceae;Parabacteroides;Parabacteroides chinchillae


metaRNA.PFemirge\_21900\_0.000011


metaRNA.PFemirge\_5076\_0.000024


GU390143.1.1290 Bacteria;Bacteroidota;Bacteroidia;Bacteroidales;Prolixibacteraceae;BSV13;uncultured bacterium


JN559628.1.1499 Bacteria;Bacteroidota;Bacteroidia;Bacteroidales;Bacteroidaceae;Bacteroides;uncultured bacterium


metaRNA.PFemirge\_13713\_0.000009


HW066305.1.1532 Bacteria;Firmicutes;Clostridia;Clostridia UCG-014;unidentified


EU777145.1.1347 Bacteria;Firmicutes;Clostridia;Christen ... ceae;Christensenellaceae R-7 group;uncultured bacterium


EU767529.1.1356 Bacteria;Bacteroidota;Bacteroidia;Bacteroidales;Bacteroidaceae;Bacteroides;uncultured bacterium


metaRNA.PFemirge\_14522\_0.000011


metaRNA.PFemirge\_7089\_0.010108


metaRNA.PFemirge\_17754\_0.000258


metaRNA.PFemirge\_9837\_0.000002


GU293185.1.1498 Bacteria;Proteobacteria;Gammaproteobact ... ales;Alteromonadaceae;Rheinheimera;uncultured bacterium


metaRNA.PFemirge\_16699\_0.000110


GQ898343.1.1489 Bacteria;Bacteroidota;Bacteroidia;Bacteroidales;Bacteroidaceae;Bacteroides;uncultured bacterium


metaRNA.PFspades\_17\_1555.440517


JX222815.1.1491 Bacteria;Proteobacteria;Gammaproteobact ... iales;Comamonadaceae;Aquabacterium;uncultured bacterium


LT629973.912339.913840 Bacteria;Verrucomicrobiota;Verru ... es;Akkermansiaceae;Akkermansia;Akkermansia glycaniphila


metaRNA.PFemirge\_17423\_0.000007


metaRNA.PFemirge\_16959\_0.000007


metaRNA.PFemirge\_4780\_0.000004


metaRNA.PFemirge\_11419\_0.000039


metaRNA.PFemirge\_30248\_0.000013


metaRNA.PFemirge\_20523\_0.000009


metaRNA.PFemirge\_13624\_0.000038


metaRNA.PFemirge\_6726\_0.000920


metaRNA.PFemirge\_3270\_0.000002


metaRNA.PFemirge\_26408\_0.000001


metaRNA.PFemirge\_17705\_0.000003


metaRNA.PFemirge\_4892\_0.000019


AB599904.1.1384 Bacteria;Proteobacteria;Alphaproteobact ... gomonadaceae;Novosphingobium;alpha proteobacterium INB3


metaRNA.PFemirge\_21457\_0.000003


metaRNA.PFemirge\_4015\_0.000023


DQ129271.1.1384 Bacteria;Actinobacteriota;Actinobacteri ... iales;Nocardioidaceae;Nocardioides;uncultured bacterium


metaRNA.PFemirge\_8537\_0.000004


metaRNA.PFspades\_4\_1638.275299


CP011073.4368909.4370448 Bacteria;Bacteroidota;Bacteroidia;Bacteroidales;Bacteroidaceae;Bacteroides;Bacteroides fragilis


metaRNA.PFemirge\_23133\_0.000160


metaRNA.PFemirge\_17584\_0.000104


metaRNA.PFemirge\_21888\_0.000086


metaRNA.PFemirge\_16860\_0.000009


JF266314.1.1315 Bacteria;Cyanobacteria;Vampirivibrionia;Obscuribacterales;Obscuribacteraceae;uncultured bacterium


metaRNA.PFemirge\_27670\_0.000071


CP014223.544406.545927 Bacteria;Firmicutes;Clostridia;L ... iraceae;Anaerostignum;Anaerotignum propionicum DSM 1682


HQ756942.1.1450 Bacteria;Firmicutes;Clostridia;Lachnosp ... iraceae;[Ruminococcus] gnavus group;uncultured organism


metaRNA.PFemirge\_24461\_0.000011


metaRNA.PFemirge\_18570\_0.001437


DQ809859.1.1362 Bacteria;Firmicutes;Clostridia;Oscillospirales;Oscillospiraceae;uncultured;uncultured bacterium


EU834774.1.1419 Bacteria;Proteobacteria;Alphaproteobact ... iales Incertae Sedis;Phreatobacter;uncultured bacterium


FN563303.1.1455 Bacteria;Firmicutes;Clostridia;Lachnospirales;Lachnospiraceae;Eisenbergiella;uncultured bacterium


metaRNA.PFemirge\_23715\_0.000282


GQ448528.1.1397 Bacteria;Firmicutes;Clostridia;Clostridia UCG-014;uncultured bacterium


metaRNA.PFemirge\_23141\_0.000006


metaRNA.PFemirge\_2446\_0.000014


metaRNA.PFemirge\_185\_0.000244


metaRNA.PFemirge\_23236\_0.000271


AF192152.1.1542 Bacteria;Desulfobacterota;Desulfovibrio ... s;Desulfovibrionaceae;Desulfovibrio;Desulfovibrio piger


metaRNA.PFemirge\_13476\_0.000005


metaRNA.PFspades\_3\_246.403614


HM778909.1.1458 Bacteria;Proteobacteria;Gammaproteobact ... les;Enterobacteriaceae;Plesiomonas;uncultured bacterium


EU766155.1.1353 Bacteria;Bacteroidota;Bacteroidia;Bacteroidales;Bacteroidaceae;Bacteroides;uncultured bacterium


metaRNA.PFemirge\_15658\_0.000008


metaRNA.PFemirge\_11292\_0.000008


metaRNA.PFemirge\_22798\_0.000024


metaRNA.PFemirge\_16117\_0.000028


DQ799866.1.1410 Bacteria;Bacteroidota;Bacteroidia;Bacteroidales;Rikenellaceae;Alistipes;uncultured bacterium


DQ795713.1.1384 Bacteria;Firmicutes;Clostridia;Lachnosp ... ospiraceae;Lachnospiraceae UCG-008;uncultured bacterium


metaRNA.PFemirge\_25050\_0.000004


metaRNA.PFemirge\_24669\_0.000001


GQ448684.1.1384 Bacteria;Bacteroidota;Bacteroidia;Bacteroidales;Rikenellaceae;Alistipes;uncultured bacterium


metaRNA.PFemirge\_617\_0.000065


metaRNA.PFemirge\_13867\_0.000006


metaRNA.PFemirge\_24351\_0.000045


DQ794241.1.1398 Bacteria;Bacteroidota;Bacteroidia;Bacteroidales;Rikenellaceae;Alistipes;uncultured bacterium


metaRNA.PFemirge\_21949\_0.000007


metaRNA.PFemirge\_20441\_0.000029


GQ360030.1.1497 Bacteria;Proteobacteria;Gammaproteobact ... acterales;Aeromonadaceae;Aeromonas;uncultured bacterium


metaRNA.PFemirge\_26323\_0.000009


metaRNA.PFemirge\_26002\_0.000049


metaRNA.PFemirge\_26536\_0.000003


EU475758.1.1213 Bacteria;Firmicutes;Clostridia;Lachnosp ... ospiraceae;Lachnospiraceae UCG-008;uncultured bacterium


metaRNA.PFemirge\_28066\_0.000004


metaRNA.PFemirge\_17635\_0.000012


metaRNA.PFemirge\_20855\_0.000009


metaRNA.PFemirge\_30262\_0.000015


metaRNA.PFemirge\_17488\_0.000014


metaRNA.PFemirge\_25985\_0.000003


metaRNA.PFemirge\_16639\_0.000058


metaRNA.PFemirge\_22655\_0.000210


MF145698.1.1460 Bacteria;Firmicutes;Clostridia;Peptostr ... ococcaceae;Terrisporobacter;Terrisporobacter glycolicus


EU508041.1.1387 Bacteria;Firmicutes;Clostridia;Lachnosp ... ospiraceae;Lachnospiraceae UCG-008;uncultured bacterium


metaRNA.PFemirge\_14141\_0.000050


KM650567.1.1481 Bacteria;Firmicutes;Negativicutes;Acida ... inococcaceae;uncultured;uncultured Firmicutes bacterium


metaRNA.PFemirge\_6300\_0.000007


metaRNA.PFemirge\_21668\_0.000001


metaRNA.PFemirge\_23414\_0.000014


metaRNA.PFspades\_15\_49.033463


metaRNA.PFemirge\_21727\_0.000007


HQ759402.1.1446 Bacteria;Firmicutes;Clostridia;Clostridia UCG-014;uncultured organism


EU560783.1.1502 Bacteria;Proteobacteria;Gammaproteobact ... obacteriaceae;Escherichia-Shigella;uncultured bacterium


metaRNA.PFemirge\_16525\_0.000007


FJ393126.1.1488 Bacteria;Bacteroidota;Bacteroidia;Bacteroidales;Bacteroidaceae;Bacteroides;uncultured Bacteroides sp.


metaRNA.PFemirge\_15099\_0.000018


metaRNA.PFemirge\_26028\_0.000012


metaRNA.PFemirge\_23621\_0.000030


GQ449092.1.1375 Bacteria;Firmicutes;Clostridia;Lachnospirales;Lachnospiraceae;Tyzzerella;uncultured bacterium


metaRNA.PFemirge\_5138\_0.000654


metaRNA.PFemirge\_20318\_0.000011


MBVN01000043.446670.448353 Eukaryota;Amorphea;Obazoa;Op ... -Candida clade;Saccharomyces cerevisiae (baker's yeast)


metaRNA.PFemirge\_2507\_0.000048


metaRNA.PFemirge\_27580\_0.000012


metaRNA.PFemirge\_24983\_0.000020


metaRNA.PFemirge\_25227\_0.000002


EF401958.1.1488 Bacteria;Bacteroidota;Bacteroidia;Bacteroidales;Tannerellaceae;Parabacteroides;uncultured bacterium


metaRNA.PFemirge\_27950\_0.000002


DQ808071.1.1266 Bacteria;Firmicutes;Negativicutes;Acida ... inococcaceae;Phascolarctobacterium;uncultured bacterium


metaRNA.PFemirge\_1410\_0.000139


GQ898126.1.1466 Bacteria;Proteobacteria;Alphaproteobacteria;Rhodospirillales;uncultured;uncultured bacterium


FJ269051.1.1487 Bacteria;Bacteroidota;Bacteroidia;Bacte ... s;iron-reducing bacterium enrichment culture clone HN11


metaRNA.PFemirge\_17240\_0.000004


metaRNA.PFemirge\_20801\_0.000002


EU460205.1.1360 Bacteria;Proteobacteria;Alphaproteobacteria;Rhodospirillales;uncultured;uncultured bacterium


KT952737.1.1475 Bacteria;Firmicutes;Clostridia;Oscillospirales;Ruminococcaceae;Pygmaiobacter;uncultured bacterium


metaRNA.PFemirge\_16372\_0.000003


metaRNA.PFspades\_14\_56.928036


metaRNA.PFemirge\_4684\_0.000107


EU766033.1.1357 Bacteria;Bacteroidota;Bacteroidia;Bacteroidales;Bacteroidaceae;Bacteroides;uncultured bacterium


AB298756.1.1471 Bacteria;Firmicutes;Clostridia;Lachnospirales;Lachnospiraceae;Anaerostignum;Anaerotignum aminivorans


metaRNA.PFemirge\_11640\_0.000331


DQ797586.1.1394 Bacteria;Firmicutes;Clostridia;Lachnosp ... raceae;[Eubacterium] eligens group;uncultured bacterium


metaRNA.PFemirge\_3348\_0.000007


HQ741361.1.1434 Bacteria;Firmicutes;Clostridia;Lachnospirales;Lachnospiraceae;Epulopiscium;uncultured bacterium


FJ868767.1.1498 Bacteria;Firmicutes;Clostridia;Lachnosp ... ospiraceae;Lachnospiraceae UCG-008;uncultured bacterium


KF945050.1.1264 Bacteria;Myxococcota;Polyangia;Haliangiales;Haliangiaceae;Haliangium;uncultured bacterium


metaRNA.PFemirge\_22892\_0.000014


JF160900.1.1352 Bacteria;Firmicutes;Clostridia;Oscillospirales;Oscillospiraceae;uncultured;uncultured bacterium


HM445966.1.1478 Bacteria;Firmicutes;Bacilli;Erysipelotrichales;Erysipelotrichaceae;Erysipelothrix;uncultured bacterium


metaRNA.PFemirge\_28799\_0.000088


metaRNA.PFemirge\_28444\_0.000111


metaRNA.PFspades\_16\_159.465154


metaRNA.PFemirge\_18143\_0.002475


metaRNA.PFemirge\_18749\_0.000101


AJ867049.1.1500 Bacteria;Desulfobacterota;Desulfovibrio ... les;Desulfovibrionaceae;Bilophila;Bilophila wadsworthia


metaRNA.PFemirge\_6250\_0.000010


metaRNA.PFemirge\_28453\_0.000003


metaRNA.PFemirge\_11867\_0.000139


metaRNA.PFtrusted\_8\_0


metaRNA.PFemirge\_3087\_0.000005


DQ115999.1.1504 Bacteria;Firmicutes;Clostridia;Lachnospirales;Lachnospiraceae;uncultured;uncultured bacterium


metaRNA.PFemirge\_20835\_0.000027


metaRNA.PFemirge\_16730\_0.000011


CBEW010000180.8693.10203 Bacteria;Cyanobacteria;Vampirivibrionia;Gastranaerophilales;Clostridium sp. CAG:306


metaRNA.PFemirge\_20894\_0.000018


FXDJ01000009.1407067.1408576 Bacteria;Bacteroidota;Bact ... erellaceae;uncultured;Porphyromonadaceae bacterium MV12


DQ799406.1.1391 Bacteria;Bacteroidota;Bacteroidia;Bacteroidales;Bacteroidaceae;Bacteroides;uncultured bacterium


metaRNA.PFemirge\_13393\_0.000478


KF697581.1.1227 Bacteria;Nitrospirota;Nitrospiria;Nitrospirales;Nitrospiraceae;Nitrospira;uncultured bacterium


EU663602.1.1492 Bacteria;Bacteroidota;Bacteroidia;Bacteroidales;Marinifilaceae;uncultured;uncultured bacterium


metaRNA.PFemirge\_30229\_0.000284


metaRNA.PFemirge\_14293\_0.000033


EU774419.1.1295 Bacteria;Firmicutes;Clostridia;Christen ... ceae;Christensenellaceae R-7 group;uncultured bacterium


metaRNA.PFemirge\_11750\_0.000002


AB486052.1.1370 Bacteria;Firmicutes;Clostridia;Oscillospirales;Oscillospiraceae;Colidextribacter;uncultured bacterium


KM651117.1.1438 Bacteria;Firmicutes;Clostridia;Oscillos ... prostanoligenes group;uncultured Spirochaetes bacterium


metaRNA.PFemirge\_15856\_0.000016


metaRNA.PFemirge\_7053\_0.000377


metaRNA.PFemirge\_15280\_0.000028


GQ417416.1.1463 Bacteria;Proteobacteria;Gammaproteobact ... s;Morganellaceae;Providencia;uncultured Providencia sp.


metaRNA.PFemirge\_17978\_0.000014


metaRNA.PFemirge\_7984\_0.000019


metaRNA.PFemirge\_5397\_0.000004


EU776216.1.1354 Bacteria;Firmicutes;Clostridia;Lachnospirales;Lachnospiraceae;uncultured;uncultured bacterium


metaRNA.PFemirge\_569\_0.000450


metaRNA.PFemirge\_26390\_0.001415


metaRNA.PFemirge\_23168\_0.000598


metaRNA.PFemirge\_12525\_0.000002


metaRNA.PFemirge\_20799\_0.000008


metaRNA.PFemirge\_15152\_0.000041


metaRNA.PFemirge\_27933\_0.000300


EU774348.1.1286 Bacteria;Firmicutes;Clostridia;Clostrid ... iaceae;Clostridium sensu stricto 1;uncultured bacterium


HQ796347.1.1441 Bacteria;Bacteroidota;Bacteroidia;Bacteroidales;Bacteroidaceae;Bacteroides;uncultured organism


EU778905.1.1289 Bacteria;Firmicutes;Clostridia;Oscillospirales;Ruminococcaceae;Faecalibacterium;uncultured bacterium


metaRNA.PFemirge\_7721\_0.000014


metaRNA.PFemirge\_17373\_0.000091


metaRNA.PFemirge\_18128\_0.000009


metaRNA.PFemirge\_28067\_0.000010


DQ799369.1.1397 Bacteria;Bacteroidota;Bacteroidia;Bacteroidales;Bacteroidaceae;Bacteroides;uncultured bacterium


metaRNA.PFemirge\_2531\_0.005487


metaRNA.PFemirge\_26399\_0.000003


DQ807591.1.1392 Bacteria;Bacteroidota;Bacteroidia;Bacteroidales;Bacteroidaceae;Bacteroides;uncultured bacterium


AY957942.1.1505 Bacteria;Actinobacteriota;Actinobacteri ... les;Mycobacteriaceae;Mycobacterium;uncultured bacterium


AF371590.1.1437 Bacteria;Firmicutes;Clostridia;Lachnospirales;Lachnospiraceae;uncultured;uncultured bacterium


metaRNA.PFemirge\_5563\_0.000393


KP231747.1.1515 Bacteria;Desulfobacterota;Desulfovibrio ... vibrionaceae;Desulfovibrio;uncultured Desulfovibrio sp.


metaRNA.PFemirge\_23282\_0.000303


FP929047.78761.80257 Bacteria;Actinobacteriota;Coriobac ... ellaceae;Gordonibacter;Gordonibacter pamelaeae 7-10-1-b


metaRNA.PFemirge\_15680\_0.005455


metaRNA.PFemirge\_28012\_0.000012


metaRNA.PFemirge\_16455\_0.000348


metaRNA.PFemirge\_15773\_0.000014


metaRNA.PFemirge\_25432\_0.000006


metaRNA.PFemirge\_19940\_0.000013


metaRNA.PFemirge\_10859\_0.000367


metaRNA.PFemirge\_27917\_0.000004


metaRNA.PFemirge\_6691\_0.000004


metaRNA.PFemirge\_19061\_0.000014


metaRNA.PFemirge\_27674\_0.002611


EU772357.1.1394 Bacteria;Bacteroidota;Bacteroidia;Bacteroidales;p-251-o5;uncultured bacterium


metaRNA.PFemirge\_2964\_0.000728


JN217164.1.1510 Bacteria;Proteobacteria;Gammaproteobact ... les;Acidithiobacillaceae;KCM-B-112;uncultured bacterium


metaRNA.PFemirge\_11899\_0.000015


DQ800054.1.1388 Bacteria;Firmicutes;Clostridia;Oscillospirales;Oscillospiraceae;uncultured;uncultured bacterium


metaRNA.PFemirge\_26334\_0.000019


metaRNA.PFemirge\_2566\_0.000026


metaRNA.PFemirge\_8548\_0.000006


CCFF01000015.520926.522414 Bacteria;Firmicutes;Clostrid ... aceae;Clostridium sensu stricto 13;Clostridium sp. CL-6


EF679189.1.1505 Bacteria;Proteobacteria;Gammaproteobact ... rales;Aeromonadaceae;Aeromonas;uncultured Aeromonas sp.


FM204998.1.1455 Bacteria;Firmicutes;Clostridia;Lachnosp ... Lachnospiraceae NC2004 group;uncultured Clostridium sp.


EU504186.1.1398 Bacteria;Firmicutes;Clostridia;Lachnosp ... ceae;Lachnospiraceae NK4A136 group;uncultured bacterium


metaRNA.PFemirge\_7563\_0.000014


EF097622.1.1400 Bacteria;Firmicutes;Clostridia;Oscillospirales;Oscillospiraceae;Colidextribacter;uncultured bacterium


metaRNA.PFemirge\_25014\_0.000142


JQ191011.1.1340 Bacteria;Firmicutes;Bacilli;RF39;uncultured bacterium


DQ809572.1.1407 Bacteria;Firmicutes;Bacilli;Erysipelotrichales;Erysipelotrichaceae;Holdemanella;uncultured bacterium


HQ807877.1.1477 Bacteria;Firmicutes;Negativicutes;Veill ... nadales;Selenomonadaceae;uncultured;uncultured organism


metaRNA.PFemirge\_23415\_0.000071


metaRNA.PFemirge\_25689\_0.000012


metaRNA.PFemirge\_23832\_0.000475


metaRNA.PFemirge\_16302\_0.000164


GQ871713.1.1490 Bacteria;Bacteroidota;Bacteroidia;Bacteroidales;Rikenellaceae;Alistipes;uncultured bacterium


metaRNA.PFemirge\_21766\_0.000003


CP012937.1474579.1476104 Bacteria;Bacteroidota;Bacteroi ... Bacteroidaceae;Bacteroides;Bacteroides thetaiotaomicron


metaRNA.PFemirge\_20522\_0.000007


metaRNA.PFemirge\_8631\_0.000216


KX431291.1.1467 Bacteria;Firmicutes;Clostridia;Peptostr ... s;Peptostreptococcaceae;Romboutsia;uncultured bacterium


metaRNA.PFemirge\_7248\_0.000159


KT952740.1.1467 Bacteria;Verrucomicrobiota;Kiritimatiel ... ales;Kiritimatiellaceae;uncultured;uncultured bacterium


metaRNA.PFemirge\_3615\_0.000018


AGXU01000010.481004.482520 Bacteria;Bacteroidota;Bacter ... acteroidaceae;Bacteroides;Bacteroides ovatus CL03T12C18


metaRNA.PFemirge\_26583\_0.000005


metaRNA.PFemirge\_6529\_0.000007


metaRNA.PFemirge\_8296\_0.000066


HQ761491.1.1432 Bacteria;Firmicutes;Clostridia;Lachnosp ... raceae;Lachnospiraceae AC2044 group;uncultured organism


AM275521.1.1477 Bacteria;Bacteroidota;Bacteroidia;Bacteroidales;Bacteroidaceae;Bacteroides;uncultured bacterium


metaRNA.PFemirge\_27051\_0.000002


metaRNA.PFemirge\_8104\_0.000009


metaRNA.PFemirge\_5226\_0.000012


metaRNA.PFemirge\_14034\_0.003854


FN428761.1.1496 Bacteria;Bacteroidota;Bacteroidia;Chitinophagales;Chitinophagaceae;Edaphobaculum;uncultured bacterium


HM445960.1.1486 Bacteria;Bacteroidota;Bacteroidia;Bacteroidales;Rikenellaceae;uncultured;uncultured bacterium


metaRNA.PFemirge\_27128\_0.000013


metaRNA.PFemirge\_17674\_0.000028


metaRNA.PFemirge\_10256\_0.000017


FJ825522.1.1343 Bacteria;Firmicutes;Clostridia;Peptostr ... s;Peptostreptococcaceae;Romboutsia;uncultured bacterium


KF704746.1.1383 Bacteria;Proteobacteria;Alphaproteobacteria;Rhizobiales;Rhizobiaceae;Shinella;Shinella zoogloeoides


metaRNA.PFemirge\_16667\_0.000017


CU926683.1.1355 Bacteria;Bacteroidota;Bacteroidia;Bacteroidales;Tannerellaceae;Macellibacteroides;uncultured bacterium


metaRNA.PFemirge\_16109\_0.000006


metaRNA.PFemirge\_17783\_0.000002


metaRNA.PFemirge\_27842\_0.000220


metaRNA.PFemirge\_10311\_0.000290


BAAZ01000703.1371.2875 Bacteria;Bacteroidota;Bacteroidi ... les;Tannerellaceae;Parabacteroides;human gut metagenome


AB509219.1.1467 Bacteria;Firmicutes;Negativicutes;Acida ... inococcaceae;Phascolarctobacterium;uncultured bacterium


metaRNA.PFemirge\_7011\_0.000156


metaRNA.PFemirge\_12627\_0.000040


LN995859.1.1478 Bacteria;Verrucomicrobiota;Chlamydiae;C ... tured;Neochlamydia sp. Trut23-12-2015\_Venoge-Embouchure


HQ783986.1.1437 Bacteria;Bacteroidota;Bacteroidia;Bacteroidales;Tannerellaceae;Parabacteroides;uncultured organism


EU777237.1.1343 Bacteria;Firmicutes;Clostridia;Lachnosp ... ospiraceae;Lachnospiraceae UCG-008;uncultured bacterium


metaRNA.PFemirge\_19201\_0.000002


metaRNA.PFemirge\_12072\_0.000009


EU776022.1.1355 Bacteria;Firmicutes;Clostridia;Peptostr ... s;Peptostreptococcaceae;uncultured;uncultured bacterium


HM630210.1.1461 Bacteria;Firmicutes;Bacilli;RF39;uncultured bacterium


metaRNA.PFemirge\_9234\_0.000003


EU773321.1.1396 Bacteria;Firmicutes;Clostridia;Oscillospirales;Oscillospiraceae;uncultured;uncultured bacterium


metaRNA.PFemirge\_15328\_0.000002


metaRNA.PFemirge\_24494\_0.000005


metaRNA.PFemirge\_1425\_0.000140


HQ778039.1.1464 Bacteria;Bacteroidota;Bacteroidia;Bacteroidales;Barnesiellaceae;Coprobacter;uncultured organism


metaRNA.PFemirge\_30258\_0.000571


AY957892.1.1500 Bacteria;Cyanobacteria;Vampirivibrionia;Obscuribacterales;Obscuribacteraceae;uncultured bacterium


EU459185.1.1385 Bacteria;Firmicutes;Clostridia;Lachnospirales;Lachnospiraceae;Epulopiscium;uncultured bacterium


metaRNA.PFemirge\_15717\_0.000002


metaRNA.PFemirge\_1169\_0.000023


MF383477.1.1442 Bacteria;Proteobacteria;Gammaproteobacteria;Pseudomonadales;Pseudomonadaceae;Pseudomonas;Pseudomonas sp.


metaRNA.PFemirge\_23423\_0.000049


GQ448331.1.1393 Bacteria;Bacteroidota;Bacteroidia;Bacteroidales;Bacteroidaceae;Bacteroides;uncultured bacterium


EU776035.1.1400 Bacteria;Proteobacteria;Gammaproteobact ... obacteriaceae;Escherichia-Shigella;uncultured bacterium


FJ393118.1.1494 Bacteria;Firmicutes;Clostridia;Oscillos ... spiraceae;uncultured;uncultured Clostridiales bacterium


metaRNA.PFemirge\_20308\_0.000025


KT002331.1.1465 Bacteria;Firmicutes;Clostridia;Peptostr ... tococcaceae;Paeniclostridium;uncultured Clostridium sp.


metaRNA.PFemirge\_13374\_0.000003


metaRNA.PFemirge\_6749\_0.000183


DQ793316.1.1393 Bacteria;Bacteroidota;Bacteroidia;Bacteroidales;Tannerellaceae;Parabacteroides;uncultured bacterium


metaRNA.PFemirge\_4551\_0.000003


metaRNA.PFemirge\_5940\_0.000090


JX221903.1.1461 Bacteria;Proteobacteria;Alphaproteobacteria;Holosporales;Holosporaceae;uncultured;uncultured bacterium


metaRNA.PFemirge\_19080\_0.000003


metaRNA.PFemirge\_13838\_0.005820


metaRNA.PFemirge\_13651\_0.000003


metaRNA.PFemirge\_12393\_0.000003


KM651159.1.1495 Bacteria;Firmicutes;Clostridia;Lachnospirales;Lachnospiraceae;Hungatella;uncultured Firmicutes bacterium


metaRNA.PFemirge\_12596\_0.000015


metaRNA.PFemirge\_18627\_0.000006


AF502202.1.1376 Bacteria;Proteobacteria;Alphaproteobacteria;Rickettsiales;Mitochondria;uncultured bacterium


DQ824259.1.1398 Bacteria;Bacteroidota;Bacteroidia;Bacteroidales;Bacteroidaceae;Bacteroides;uncultured bacterium


metaRNA.PFemirge\_25382\_0.000016


metaRNA.PFemirge\_24309\_0.000005


FPLP01011680.8.1529 Bacteria;Proteobacteria;Gammaproteobacteria;Enterobacterales;Aeromonadaceae;Aeromonas;metagenome


EU776080.1.1388 Bacteria;Firmicutes;Clostridia;Lachnosp ... aceae;Lachnospiraceae NC2004 group;uncultured bacterium


AB552872.1.1469 Bacteria;Verrucomicrobiota;Verrucomicro ... rucomicrobiaceae;Brevifollis;Brevifollis gellanilyticus


metaRNA.PFemirge\_27844\_0.000018


metaRNA.PFemirge\_23065\_0.000024


metaRNA.PFemirge\_30234\_0.000078


metaRNA.PFemirge\_12029\_0.000002


metaRNA.PFemirge\_18308\_0.000006


metaRNA.PFemirge\_2697\_0.000130


metaRNA.PFemirge\_14145\_0.000012


metaRNA.PFemirge\_5047\_0.000017


metaRNA.PFemirge\_17293\_0.000099


metaRNA.PFemirge\_23152\_0.000010


metaRNA.PFemirge\_18472\_0.000010


metaRNA.PFemirge\_22518\_0.000005


metaRNA.PFemirge\_1728\_0.000036


HQ761714.1.1441 Bacteria;Bacteroidota;Bacteroidia;Bacteroidales;Tannerellaceae;Parabacteroides;uncultured organism


metaRNA.PFemirge\_12531\_0.000003


metaRNA.PFemirge\_23054\_0.000108


metaRNA.PFemirge\_27719\_0.000142


FM213077.1.1512 Bacteria;Proteobacteria;Gammaproteobact ... ales;Rhodanobacteraceae;Tahibacter;uncultured bacterium


HQ697726.1.1505 Bacteria;Proteobacteria;Gammaproteobact ... les;Acidithiobacillaceae;KCM-B-112;uncultured bacterium


metaRNA.PFemirge\_634\_0.000009


DQ817541.1.1479 Bacteria;Proteobacteria;Gammaproteobact ... les;Chromobacteriaceae;Crenobacter;uncultured bacterium


metaRNA.PFemirge\_22129\_0.000002


JN703378.1.1490 Bacteria;Bacteroidota;Bacteroidia;Bacteroidales;Barnesiellaceae;Coprobacter;Coprobacter fastidiosus


metaRNA.PFemirge\_21002\_0.000004


metaRNA.PFemirge\_5756\_0.000092


metaRNA.PFemirge\_10739\_0.000004


metaRNA.PFemirge\_754\_0.000780


GQ388996.1.1510 Bacteria;Proteobacteria;Gammaproteobact ... dales;Pseudomonadaceae;Pseudomonas;uncultured bacterium


metaRNA.PFemirge\_14656\_0.000017


metaRNA.PFemirge\_25652\_0.000166


metaRNA.PFemirge\_7297\_0.001339


metaRNA.PFemirge\_23222\_0.000005


metaRNA.PFemirge\_28036\_0.000003


metaRNA.PFemirge\_5633\_0.000088


metaRNA.PFemirge\_16457\_0.000413


metaRNA.PFemirge\_5532\_0.003110


HM556834.1.1374 Bacteria;Cyanobacteria;Cyanobacteriia;Chloroplast;uncultured bacterium


EF096560.1.1389 Bacteria;Firmicutes;Clostridia;Lachnosp ... ospiraceae;Lachnospiraceae UCG-008;uncultured bacterium


metaRNA.PFemirge\_6254\_0.000005


metaRNA.PFemirge\_18679\_0.000010


metaRNA.PFemirge\_26243\_0.000042


EU772076.1.1400 Bacteria;Firmicutes;Clostridia;Lachnosp ... ospiraceae;Lachnospiraceae UCG-008;uncultured bacterium


metaRNA.PFemirge\_22949\_0.000009


metaRNA.PFemirge\_30231\_0.000042


metaRNA.PFemirge\_9530\_0.000005


LN573922.1.1381 Bacteria;Planctomycetota;Planctomycetes;Gemmatales;Gemmataceae;uncultured;uncultured bacterium


metaRNA.PFemirge\_2499\_0.043392


metaRNA.PFemirge\_19427\_0.000020


EU508987.1.1386 Bacteria;Firmicutes;Clostridia;Oscillospirales;Ruminococcaceae;Ruminococcus;uncultured bacterium


DQ801443.1.1401 Bacteria;Bacteroidota;Bacteroidia;Bacteroidales;Bacteroidaceae;Bacteroides;uncultured bacterium


metaRNA.PFemirge\_7881\_0.000314


metaRNA.PFemirge\_26565\_0.000107


metaRNA.PFemirge\_27319\_0.000003


KC994683.1.1482 Bacteria;Bacteroidota;Bacteroidia;Cytophagales;Spirosomaceae;Runella;uncultured Runella sp.


metaRNA.PFemirge\_30233\_0.000082


metaRNA.PFemirge\_10872\_0.000001


metaRNA.PFemirge\_12701\_0.000008


metaRNA.PFemirge\_16270\_0.000003


metaRNA.PFemirge\_24892\_0.000001


LIUU01000102.1484.3268 Eukaryota;Amorphea;Obazoa;Opisth ... etaceae;Zygosaccharomyces;Saccharomyces cerevisiae W303


metaRNA.PFemirge\_1247\_0.000466


DQ807523.1.1284 Bacteria;Firmicutes;Clostridia;Lachnosp ... aceae;[Ruminococcus] torques group;uncultured bacterium


JN653056.1.1473 Bacteria;Desulfobacterota;Desulfovibrio ... ;Desulfovibrio;uncultured Desulfovibrionaceae bacterium


metaRNA.PFemirge\_14889\_0.000011


HQ697737.1.1504 Bacteria;Proteobacteria;Gammaproteobact ... les;Diplorickettsiaceae;uncultured;uncultured bacterium


HQ779103.1.1440 Bacteria;Firmicutes;Clostridia;Lachnosp ... raceae;Lachnospiraceae NC2004 group;uncultured organism


metaRNA.PFemirge\_27006\_0.000032


metaRNA.PFemirge\_6989\_0.000076


AM265457.1.1438 Bacteria;Firmicutes;Clostridia;Oscillospirales;Oscillospiraceae;Colidextribacter;uncultured bacterium


DQ806118.1.1388 Bacteria;Firmicutes;Clostridia;Lachnosp ... ospiraceae;Lachnospiraceae UCG-008;uncultured bacterium


EU454507.1.1407 Bacteria;Firmicutes;Clostridia;Oscillospirales;Butyricicoccaceae;UCG-009;uncultured bacterium


AB746664.1.1536 Bacteria;Firmicutes;Clostridia;Lachnosp ... ceae;Lachnospiraceae NK4A136 group;uncultured bacterium


metaRNA.PFemirge\_13870\_0.000633


metaRNA.PFemirge\_9528\_0.000003


AB034016.1.1515 Bacteria;Cyanobacteria;Vampirivibrionia;Gastranaerophilales;uncultured rumen bacterium 4C0d-2


metaRNA.PFemirge\_25582\_0.000010


metaRNA.PFemirge\_21161\_0.000003


GU302791.1.1495 Bacteria;Firmicutes;Clostridia;Christen ... hristensenellaceae R-7 group;uncultured rumen bacterium


metaRNA.PFemirge\_2602\_0.000036


metaRNA.PFemirge\_19936\_0.000026


HM630205.1.1449 Bacteria;Bacteroidota;Bacteroidia;Bacteroidales;Rikenellaceae;Rikenella;uncultured bacterium


HQ778029.1.1454 Bacteria;Firmicutes;Clostridia;Lachnospirales;Lachnospiraceae;Lachnoclostridium;uncultured organism


DQ795734.1.1397 Bacteria;Firmicutes;Clostridia;Lachnosp ... ae;[Ruminococcus] gauvreauii group;uncultured bacterium


metaRNA.PFemirge\_2462\_0.000030


metaRNA.PFemirge\_24886\_0.000009


metaRNA.PFemirge\_8940\_0.001871


EU775962.1.1393 Bacteria;Bacteroidota;Bacteroidia;Bacteroidales;Bacteroidaceae;Bacteroides;uncultured bacterium


metaRNA.PFemirge\_4143\_0.000112


metaRNA.PFemirge\_19950\_0.000001


metaRNA.PFemirge\_16690\_0.000006


metaRNA.PFemirge\_28311\_0.021522


EU777679.1.1373 Bacteria;Firmicutes;Clostridia;Clostrid ... iaceae;Clostridium sensu stricto 1;uncultured bacterium


metaRNA.PFemirge\_27694\_0.000596


metaRNA.PFemirge\_27313\_0.000197


metaRNA.PFemirge\_12098\_0.000009


metaRNA.PFtrusted\_9\_0


metaRNA.PFemirge\_7236\_0.000005


metaRNA.PFemirge\_350\_0.000007


metaRNA.PFemirge\_12790\_0.000229


metaRNA.PFemirge\_17214\_0.000011


GQ448246.1.1389 Bacteria;Bacteroidota;Bacteroidia;Bacteroidales;Bacteroidaceae;Bacteroides;uncultured bacterium


metaRNA.PFemirge\_12523\_0.000091


metaRNA.PFemirge\_23318\_0.000006


FPLN01009549.9.1530 Bacteria;Proteobacteria;Gammaproteo ... ales;Enterobacteriaceae;Escherichia-Shigella;metagenome


EU773857.1.1364 Bacteria;Firmicutes;Clostridia;Peptostr ... s;Peptostreptococcaceae;uncultured;uncultured bacterium


metaRNA.PFemirge\_8467\_0.000207


metaRNA.PFemirge\_8530\_0.000070


metaRNA.PFemirge\_3320\_0.000004


metaRNA.PFemirge\_22310\_0.000028


metaRNA.PFemirge\_28266\_0.000003


metaRNA.PFemirge\_1572\_0.000005


metaRNA.PFemirge\_16875\_0.000021


metaRNA.PFemirge\_22896\_0.000249


metaRNA.PFemirge\_1887\_0.011605


metaRNA.PFemirge\_26340\_0.000017


metaRNA.PFemirge\_27718\_0.000020


FPLS01020683.16.1482 Bacteria;Proteobacteria;Alphaprote ... omonadales;Sphingomonadaceae;Novosphingobium;metagenome


metaRNA.PFemirge\_5717\_0.000041


DQ797995.1.1380 Bacteria;Firmicutes;Clostridia;Oscillospirales;Oscillospiraceae;uncultured;uncultured bacterium


FJ823944.1.1497 Bacteria;Proteobacteria;Gammaproteobact ... ales;Comamonadaceae;uncultured;uncultured Comamonas sp.


CU920243.2.1323 Bacteria;Planctomycetota;Planctomycetes;Planctomycetales;Rubinisphaeraceae;SH-PL14;uncultured bacterium


EU778864.1.1377 Bacteria;Firmicutes;Clostridia;Lachnosp ... ospiraceae;Lachnospiraceae UCG-008;uncultured bacterium


metaRNA.PFemirge\_16323\_0.000005


metaRNA.PFemirge\_14692\_0.000045


metaRNA.PFemirge\_28131\_0.013761


AB649277.1.1490 Bacteria;Firmicutes;Bacilli;Erysipelotrichales;Erysipelotrichaceae;Faecalitalea;Absiella dolichum


MAIQ01000023.175.1690 Bacteria;Firmicutes;Clostridia;Ch ... les;Christensenellaceae;Christensenella sp. AF73-05CM02


metaRNA.PFemirge\_12919\_0.000002


ADLN01000129.202.1639 Bacteria;Firmicutes;Clostridia;La ... spiraceae;Incertae Sedis;Hungatella hathewayi WAL-18680


metaRNA.PFemirge\_28135\_0.000007


AJ576387.1.1452 Bacteria;Firmicutes;Bacilli;Erysipelotr ... stridiaceae;Erysipelatoclostridium;uncultured bacterium


metaRNA.PFemirge\_21965\_0.004375


metaRNA.PFemirge\_4206\_0.000185


metaRNA.PFemirge\_23960\_0.000001


EF454925.1.1440 Bacteria;Synergistota;Synergistia;Syner ... gistaceae;uncultured;uncultured Synergistetes bacterium


metaRNA.PFemirge\_9501\_0.010235


EU773851.1.1360 Bacteria;Firmicutes;Clostridia;Peptostr ... s;Peptostreptococcaceae;uncultured;uncultured bacterium


metaRNA.PFemirge\_5105\_0.000003


metaRNA.PFemirge\_16462\_0.000050


metaRNA.PFemirge\_24081\_0.000001


metaRNA.PFemirge\_16647\_0.000019


metaRNA.PFemirge\_7416\_0.000030


metaRNA.PFemirge\_22783\_0.000004


CDTW01027470.3361.4864 Bacteria;Bacteroidota;Bacteroidia;Bacteroidales;Rikenellaceae;Alistipes;gut metagenome


metaRNA.PFemirge\_26828\_0.000001


metaRNA.PFemirge\_19845\_0.000019


metaRNA.PFemirge\_10925\_0.000001


metaRNA.PFemirge\_21528\_0.000002


metaRNA.PFspades\_1\_6007.471877


FJ374211.1.1495 Bacteria;Firmicutes;Clostridia;Christensenellales;Christensenellaceae;uncultured;uncultured bacterium


DQ807393.1.1301 Bacteria;Firmicutes;Clostridia;Lachnospirales;Lachnospiraceae;Anaerostipes;uncultured bacterium


DQ793916.1.1399 Bacteria;Bacteroidota;Bacteroidia;Bacteroidales;Bacteroidaceae;Bacteroides;uncultured bacterium


metaRNA.PFemirge\_23926\_0.000005


metaRNA.PFemirge\_16460\_0.000006


metaRNA.PFemirge\_6509\_0.000467


metaRNA.PFemirge\_18633\_0.000002


EU469747.1.1392 Bacteria;Firmicutes;Clostridia;Lachnospirales;Lachnospiraceae;28-4;uncultured bacterium


metaRNA.PFemirge\_28459\_0.000059


EU622704.1.1488 Bacteria;Firmicutes;Clostridia;Oscillospirales;Oscillospiraceae;Colidextribacter;uncultured bacterium


metaRNA.PFemirge\_26903\_0.000064


LARG01000024.765.2171 Bacteria;Bacteroidota;Bacteroidia;Bacteroidales;Rikenellaceae;Alistipes;Alistipes sp. N15.MGS-157


metaRNA.PFemirge\_21155\_0.000022


metaRNA.PFemirge\_9916\_0.000359


metaRNA.PFemirge\_30238\_0.000025


metaRNA.PFemirge\_7079\_0.000003


metaRNA.PFemirge\_16271\_0.000008


GU559813.1.1427 Bacteria;Firmicutes;Negativicutes;Veill ... oromusaceae;Anaeromusa-Anaeroarcus;uncultured bacterium


LGUD01000026.3673.5196 Bacteria;Proteobacteria;Gammapro ... cyclaceae;Dechlorosoma;Rhodocyclaceae bacterium Paddy-1


metaRNA.PFemirge\_19539\_0.000004


EF454852.1.1475 Bacteria;Desulfobacterota;Desulfovibrio ... eae;uncultured;uncultured Desulfovibrionaceae bacterium


metaRNA.PFemirge\_11241\_0.000058


AY916388.1.1341 Bacteria;Proteobacteria;Alphaproteobacteria;Rhodospirillales;uncultured;uncultured bacterium


metaRNA.PFemirge\_13422\_0.000143


metaRNA.PFemirge\_22204\_0.000004


metaRNA.PFemirge\_10002\_0.000002


JN012610.1.1443 Bacteria;Firmicutes;Clostridia;Oscillospirales;Ruminococcaceae;Anaerotruncus;uncultured bacterium


metaRNA.PFemirge\_21667\_0.000015


metaRNA.PFemirge\_27758\_0.000017


EU771929.1.1386 Bacteria;Firmicutes;Clostridia;Lachnosp ... ospiraceae;Lachnospiraceae UCG-008;uncultured bacterium


AM934838.1.1381 Bacteria;Dadabacteria;Dadabacteriia;Dadabacteriales;uncultured candidate division SBR1093 bacterium


EU776960.1.1399 Bacteria;Proteobacteria;Gammaproteobact ... obacteriaceae;Escherichia-Shigella;uncultured bacterium


metaRNA.PFemirge\_25565\_0.000008


metaRNA.PFemirge\_8356\_0.000005


metaRNA.PFemirge\_16161\_0.000079


metaRNA.PFemirge\_6590\_0.000028


metaRNA.PFemirge\_12661\_0.000008


metaRNA.PFemirge\_14112\_0.000025


metaRNA.PFemirge\_5910\_0.000239


metaRNA.PFemirge\_16099\_0.000208


X71855.1.1501 Bacteria;Firmicutes;Clostridia;Lachnospir ... spiraceae;Lachnoclostridium;[Clostridium] xylanolyticum


KC189649.1.1474 Bacteria;Bacteroidota;Bacteroidia;Chitinophagales;Saprospiraceae;uncultured;uncultured bacterium


EU465589.1.1360 Bacteria;Firmicutes;Clostridia;Lachnosp ... ceae;Lachnospiraceae NK4A136 group;uncultured bacterium


FPLL01003728.5.1485 Bacteria;Proteobacteria;Gammaproteobacteria;Xanthomonadales;Rhodanobacteraceae;Tahibacter;metagenome


metaRNA.PFemirge\_21898\_0.000007


metaRNA.PFemirge\_10927\_0.000021


metaRNA.PFemirge\_500\_0.000003


metaRNA.PFemirge\_24307\_0.000051


FLKF01000010.1355.2888 Bacteria;Firmicutes;Negativicute ... eae;Anaerosinus;Clostridiales bacterium Marseille-P2411


metaRNA.PFemirge\_27935\_0.000081


KF318232.1.1402 Bacteria;Bacteroidota;Bacteroidia;Bacte ... oidaceae;Bacteroides;uncultured Bacteroidetes bacterium


metaRNA.PFemirge\_11285\_0.000004


AB595134.1.1484 Bacteria;Firmicutes;Clostridia;Eubacteriales;Eubacteriaceae;Eubacterium;Eubacterium limosum


ABVO01000045.3575.5089 Bacteria;Bacteroidota;Bacteroidi ... teroidaceae;Bacteroides;Bacteroides eggerthii DSM 20697


metaRNA.PFemirge\_598\_0.000032


HM630238.1.1413 Bacteria;Bacteroidota;Bacteroidia;Bacteroidales;Rikenellaceae;Rikenella;uncultured bacterium


JX457277.1.1488 Bacteria;Bacteroidota;Bacteroidia;Bacteroidales;Bacteroidaceae;Bacteroides;uncultured bacterium


KM251124.1.1542 Bacteria;Firmicutes;Clostridia;Oscillospirales;Hydrogenoanaerobacterium;uncultured bacterium


DQ659895.1.1400 Bacteria;Actinobacteriota;Actinobacteria;Corynebacteriales;Nocardiaceae;Nocardia;Nocardia abscessus


metaRNA.PFemirge\_11161\_0.000016


metaRNA.PFemirge\_12928\_0.000016


metaRNA.PFemirge\_27707\_0.000010


metaRNA.PFemirge\_10478\_0.000063


metaRNA.PFemirge\_8549\_0.000000


JF490034.1.1450 Bacteria;Proteobacteria;Alphaproteobact ... gnetospirillaceae;uncultured;alpha proteobacterium UT-4


LC194927.1.1488 Bacteria;Firmicutes;Clostridia;Lachnosp ... ceae;Lachnospiraceae UCG-008;uncultured Clostridium sp.


metaRNA.PFemirge\_27747\_0.000017


metaRNA.PFemirge\_16933\_0.000070


BBDT01000057.3432.4945 Bacteria;Firmicutes;Bacilli;Erys ... ceae;Coprobacillus;Coprobacillus cateniformis JCM 10604


metaRNA.PFemirge\_17626\_0.000010


metaRNA.PFemirge\_30251\_0.000064


metaRNA.PFemirge\_18253\_0.000499


metaRNA.PFemirge\_9589\_0.000015


metaRNA.PFemirge\_16705\_0.005309


KF318250.1.1396 Bacteria;Verrucomicrobiota;Verrucomicro ... iaceae;Akkermansia;uncultured Verrucomicrobia bacterium


metaRNA.PFemirge\_13425\_0.001237


metaRNA.PFemirge\_8289\_0.000288


HG917472.1.1448 Bacteria;Proteobacteria;Gammaproteobact ... les;Chitinimonadaceae;Chitinivorax;uncultured bacterium


metaRNA.PFemirge\_27722\_0.000401


KM650913.1.1442 Bacteria;Bacteroidota;Bacteroidia;Bacte ... oidaceae;Bacteroides;uncultured Bacteroidetes bacterium


FQ658622.1.1363 Bacteria;Proteobacteria;Gammaproteobact ... drogenophilaceae;Thiobacillus;uncultured soil bacterium


metaRNA.PFemirge\_7161\_0.003040


metaRNA.PFemirge\_23519\_0.000340


DQ824480.1.1373 Bacteria;Actinobacteriota;Coriobacterii ... riales;Eggerthellaceae;Eggerthella;uncultured bacterium


EU511665.1.1382 Bacteria;Firmicutes;Clostridia;Lachnosp ... ospiraceae;Lachnospiraceae UCG-008;uncultured bacterium


metaRNA.PFemirge\_16556\_0.000006


JX457309.1.1485 Bacteria;Bacteroidota;Bacteroidia;Bacteroidales;Rs-E47 termite group;uncultured bacterium


metaRNA.PFemirge\_14604\_0.000176


metaRNA.PFemirge\_25825\_0.000004


metaRNA.PFemirge\_19515\_0.000091


GQ449181.1.1364 Bacteria;Bacteroidota;Bacteroidia;Bacte ... laceae;Rikenellaceae RC9 gut group;uncultured bacterium


metaRNA.PFemirge\_16369\_0.000269


EU234274.1.1501 Bacteria;Proteobacteria;Gammaproteobacteria;Burkholderiales;Rhodocyclaceae;C39;uncultured bacterium


metaRNA.PFemirge\_25886\_0.000008


metaRNA.PFemirge\_3711\_0.000017


GQ480135.1.1486 Bacteria;Bacteroidota;Bacteroidia;Chitinophagales;Saprospiraceae;Haliscomenobacter;uncultured bacterium


DQ800695.1.1386 Bacteria;Firmicutes;Clostridia;Lachnospirales;Lachnospiraceae;Lachnoclostridium;uncultured bacterium


metaRNA.PFemirge\_23310\_0.000097


metaRNA.PFemirge\_6600\_0.000395


metaRNA.PFemirge\_28304\_0.000021


metaRNA.PFemirge\_11903\_0.000099


metaRNA.PFemirge\_3699\_0.000078


HQ739338.1.1431 Bacteria;Bacteroidota;Bacteroidia;Bacteroidales;Bacteroidaceae;Bacteroides;uncultured bacterium


metaRNA.PFemirge\_23145\_0.000025


metaRNA.PFemirge\_26665\_0.000003


DQ905881.1.1578 Bacteria;Bacteroidota;Bacteroidia;Bacteroidales;Bacteroidaceae;Bacteroides;uncultured bacterium


KP975338.1.1480 Bacteria;Proteobacteria;Gammaproteobact ... acterales;Aeromonadaceae;Aeromonas;uncultured bacterium


metaRNA.PFemirge\_22634\_0.000016


metaRNA.PFemirge\_26078\_0.000014


LARD01000021.11.1513 Bacteria;Cyanobacteria;Vampirivibrionia;Gastranaerophilales;Clostridium sp. K4410.MGS-306


metaRNA.PFemirge\_26869\_0.000003


metaRNA.PFemirge\_18360\_0.002973


metaRNA.PFemirge\_6283\_0.000007


metaRNA.PFemirge\_5019\_0.000087


metaRNA.PFspades\_25\_97.307312


metaRNA.PFemirge\_4201\_0.000063


metaRNA.PFemirge\_10824\_0.001220


metaRNA.PFemirge\_27787\_0.000299


metaRNA.PFemirge\_8249\_0.000024


metaRNA.PFemirge\_21028\_0.000008


metaRNA.PFspades\_10\_39.599860


metaRNA.PFemirge\_22315\_0.000005


metaRNA.PFemirge\_10712\_0.000005


AB821059.1.1460 Bacteria;Proteobacteria;Alphaproteobact ... s;Xanthobacteraceae;Bradyrhizobium;uncultured bacterium


metaRNA.PFemirge\_16949\_0.000007


metaRNA.PFemirge\_5554\_0.000160


FR799325.1.1443 Bacteria;Proteobacteria;Alphaproteobact ... nthobacteraceae;Azorhizobium;Azorhizobium oxalatiphilum


KJ725364.1.1442 Bacteria;Proteobacteria;Gammaproteobacteria;Burkholderiales;Chromobacteriaceae;Vogesella;Vibrio cholerae


metaRNA.PFemirge\_10921\_0.000003


FP929032.881290.882801 Bacteria;Bacteroidota;Bacteroidia;Bacteroidales;Rikenellaceae;Alistipes;Alistipes shahii WAL 8301


metaRNA.PFemirge\_21241\_0.000018


metaRNA.PFemirge\_5789\_0.000006


metaRNA.PFemirge\_20191\_0.000004


metaRNA.PFemirge\_16058\_0.000088


JN607058.1.1380 Bacteria;Verrucomicrobiota;Chlamydiae;Chlamydiales;Parachlamydiaceae;Neochlamydia;uncultured bacterium


metaRNA.PFemirge\_13989\_0.000254


metaRNA.PFemirge\_20527\_0.000002


metaRNA.PFemirge\_26324\_0.000002


EU663601.1.1489 Bacteria;Bacteroidota;Bacteroidia;Bacteroidales;Marinifilaceae;Butyricimonas;uncultured bacterium


metaRNA.PFemirge\_16812\_0.000006


metaRNA.PFemirge\_637\_0.000307


metaRNA.PFemirge\_7034\_0.000006


EU772114.1.1390 Bacteria;Firmicutes;Clostridia;Peptostr ... caceae;[Eubacterium] nodatum group;uncultured bacterium


metaRNA.PFemirge\_19321\_0.000007


metaRNA.PFemirge\_18963\_0.000130


metaRNA.PFemirge\_26180\_0.000003


metaRNA.PFemirge\_18380\_0.000005


metaRNA.PFemirge\_27887\_0.000076


metaRNA.PFemirge\_13618\_0.000005


GU174132.1.1481 Bacteria;Firmicutes;Clostridia;Lachnosp ... ospiraceae;Lachnospiraceae UCG-009;uncultured bacterium


metaRNA.PFemirge\_21019\_0.000048


metaRNA.PFemirge\_19176\_0.000253


metaRNA.PFemirge\_6166\_0.000389


metaRNA.PFemirge\_19034\_0.000004


metaRNA.PFemirge\_14922\_0.000022


metaRNA.PFemirge\_6977\_0.000305


DQ799101.1.1373 Bacteria;Firmicutes;Clostridia;Oscillospirales;Ruminococcaceae;Faecalibacterium;uncultured bacterium


metaRNA.PFemirge\_26055\_0.000012


JN559646.1.1500 Bacteria;Bacteroidota;Bacteroidia;Bacte ... llaceae;Prevotellaceae Ga6A1 group;uncultured bacterium


metaRNA.PFspades\_6\_72.965356


AB198564.1.1344 Bacteria;Elusimicrobiota;Elusimicrobia; ... eae;Elusimicrobium;uncultured Termite group 1 bacterium


metaRNA.PFemirge\_19798\_0.000002


metaRNA.PFemirge\_13330\_0.000023


EU772602.1.1240 Bacteria;Proteobacteria;Gammaproteobact ... obacteriaceae;Escherichia-Shigella;uncultured bacterium


metaRNA.PFemirge\_23708\_0.000015


metaRNA.PFemirge\_22960\_0.000002


DQ805525.1.1380 Bacteria;Firmicutes;Clostridia;Lachnosp ... ospiraceae;Lachnospiraceae UCG-008;uncultured bacterium


metaRNA.PFemirge\_30260\_0.000080


EU472520.1.1399 Bacteria;Firmicutes;Clostridia;Lachnosp ... aceae;Lachnospiraceae AC2044 group;uncultured bacterium


metaRNA.PFemirge\_20627\_0.000004


metaRNA.PFemirge\_7126\_0.000043


metaRNA.PFemirge\_18993\_0.000006


HM123962.1.1510 Bacteria;Desulfobacterota;Desulfovibrio ... les;Desulfovibrionaceae;uncultured;uncultured bacterium


metaRNA.PFemirge\_13244\_0.000017


CU926592.1.1355 Bacteria;Bacteroidota;Bacteroidia;Bacteroidales;Porphyromonadaceae;Porphyromonas;uncultured bacterium


metaRNA.PFemirge\_9592\_0.000007


metaRNA.PFemirge\_8137\_0.000002


metaRNA.PFemirge\_11197\_0.000002


AY494079.33242.34761 Bacteria;Proteobacteria;Alphaproteobacteria;Rickettsiales;Mitochondria;Fucus vesiculosus


metaRNA.PFemirge\_26884\_0.000016


EU457153.1.1393 Bacteria;Firmicutes;Clostridia;Oscillospirales;Ruminococcaceae;Anaerotruncus;uncultured bacterium


metaRNA.PFemirge\_5887\_0.000002


metaRNA.PFemirge\_11104\_0.000018


FJ848388.1.1476 Bacteria;Proteobacteria;Alphaproteobacteria;Rhodospirillales;uncultured;uncultured bacterium


metaRNA.PFemirge\_10978\_0.000740


metaRNA.PFemirge\_23699\_0.000008


metaRNA.PFemirge\_23113\_0.000004


metaRNA.PFemirge\_10226\_0.014067


metaRNA.PFemirge\_27732\_0.002573


EU763324.1.1351 Bacteria;Firmicutes;Clostridia;Lachnosp ... ceae;Lachnospiraceae NK4A136 group;uncultured bacterium


EU778614.1.1403 Bacteria;Bacteroidota;Bacteroidia;Bacteroidales;Rikenellaceae;Alistipes;uncultured bacterium


metaRNA.PFemirge\_20655\_0.000007


EU104080.1.1428 Bacteria;Bacteroidota;Bacteroidia;Chitinophagales;Saprospiraceae;uncultured;uncultured bacterium


HQ813540.1.1430 Bacteria;Bacteroidota;Bacteroidia;Bacteroidales;Tannerellaceae;Parabacteroides;uncultured organism


metaRNA.PFtrusted\_10\_0


metaRNA.PFemirge\_4408\_0.000004


metaRNA.PFemirge\_3472\_0.000036


DQ355176.1.1464 Bacteria;Desulfobacterota;Desulfovibrio ... vibrionaceae;Desulfovibrio;uncultured Desulfovibrio sp.


metaRNA.PFspades\_9\_311.027739


metaRNA.PFemirge\_11224\_0.000318


metaRNA.PFemirge\_6605\_0.000009


metaRNA.PFemirge\_27320\_0.000082


metaRNA.PFemirge\_15283\_0.000001


HQ385555.1.1318 Bacteria;Proteobacteria;Alphaproteobact ... rales;Rhodobacteraceae;Rhodobacter;uncultured bacterium


metaRNA.PFemirge\_2932\_0.000015


metaRNA.PFemirge\_19967\_0.000019


metaRNA.PFemirge\_15497\_0.000007


metaRNA.PFemirge\_19051\_0.000004


metaRNA.PFemirge\_19085\_0.000007


metaRNA.PFemirge\_13111\_0.000429


metaRNA.PFemirge\_5611\_0.000002


metaRNA.PFemirge\_27736\_0.001373


metaRNA.PFemirge\_24895\_0.000008


metaRNA.PFemirge\_11165\_0.000012


metaRNA.PFemirge\_7319\_0.000002


metaRNA.PFemirge\_8975\_0.000015


metaRNA.PFemirge\_2498\_0.010021


LC036226.1.1496 Bacteria;Bacteroidota;Bacteroidia;Bacteroidales;Paludibacteraceae;H1;uncultured Paludibacter sp.


metaRNA.PFemirge\_22167\_0.000009


AF157057.1.1502 Bacteria;Deferribacterota;Deferribacter ... rribacteraceae;Mucispirillum;bacterium 'Lincoln Park 3'


metaRNA.PFemirge\_5128\_0.000006


metaRNA.PFemirge\_10848\_0.000006


metaRNA.PFemirge\_18132\_0.000001


metaRNA.PFemirge\_26292\_0.000001


metaRNA.PFemirge\_22441\_0.000050


FJ868757.1.1502 Bacteria;Bacteroidota;Bacteroidia;Bacteroidales;Porphyromonadaceae;Porphyromonas;uncultured bacterium


metaRNA.PFemirge\_14512\_0.003498


HM779545.1.1509 Bacteria;Proteobacteria;Gammaproteobact ... dales;Pseudomonadaceae;Pseudomonas;uncultured bacterium


metaRNA.PFemirge\_11376\_0.000004


metaRNA.PFemirge\_15735\_0.000012


metaRNA.PFemirge\_2521\_0.000002


ASTA01000042.17380.18894 Bacteria;Firmicutes;Clostridia ... uminococcaceae;Anaerotruncus;Anaerotruncus sp. G3(2012)


metaRNA.PFemirge\_27966\_0.000003


metaRNA.PFemirge\_15465\_0.000001


EU777150.1.1350 Bacteria;Firmicutes;Clostridia;Lachnosp ... ospiraceae;Lachnospiraceae UCG-004;uncultured bacterium


metaRNA.PFemirge\_5958\_0.000005


LN833517.1.1512 Bacteria;Proteobacteria;Gammaproteobact ... riaceae;Escherichia-Shigella;uncultured Escherichia sp.


CP029554.2592470.2594015 Bacteria;Proteobacteria;Gammap ... riaceae;Chromobacterium;Chromobacterium sp. IIBBL 274-1


metaRNA.PFemirge\_27667\_0.000530


metaRNA.PFemirge\_30228\_0.000122


FPLP01007766.16.1549 Bacteria;Myxococcota;Polyangia;Haliangiales;Haliangiaceae;Haliangium;metagenome


metaRNA.PFemirge\_23481\_0.000008


metaRNA.PFemirge\_27684\_0.001116


metaRNA.PFemirge\_12807\_0.000003


EU775302.1.1379 Bacteria;Firmicutes;Clostridia;Lachnosp ... aceae;Lachnospiraceae AC2044 group;uncultured bacterium


metaRNA.PFemirge\_28197\_0.000021


metaRNA.PFemirge\_24439\_0.000029


FPLP01012861.16.1542 Bacteria;Verrucomicrobiota;Verruco ... comicrobiales;Verrucomicrobiaceae;uncultured;metagenome


GQ133618.1.1320 Bacteria;Firmicutes;Bacilli;Erysipelotrichales;Erysipelotrichaceae;Erysipelothrix;uncultured bacterium


KF500761.1.1329 Bacteria;Proteobacteria;Gammaproteobacteria;Burkholderiales;uncultured;uncultured bacterium


metaRNA.PFemirge\_4338\_0.000074


metaRNA.PFemirge\_21142\_0.000004


EU778872.1.1287 Bacteria;Firmicutes;Clostridia;Lachnosp ... ospiraceae;Lachnospiraceae UCG-004;uncultured bacterium


metaRNA.PFemirge\_2689\_0.000027


JN391820.1.1494 Bacteria;Proteobacteria;Gammaproteobact ... riales;Comamonadaceae;Sphaerotilus;uncultured bacterium


metaRNA.PFemirge\_7796\_0.000017


EU772683.1.1391 Bacteria;Firmicutes;Clostridia;Lachnospirales;Lachnospiraceae;Blautia;uncultured bacterium


metaRNA.PFemirge\_9231\_0.000012


EU466086.1.1375 Bacteria;Firmicutes;Clostridia;Lachnospirales;Lachnospiraceae;Dorea;uncultured bacterium


metaRNA.PFemirge\_23221\_0.000003


metaRNA.PFemirge\_12071\_0.024427


MG754439.1.1392 Bacteria;Proteobacteria;Alphaproteobacteria;Rhizobiales;Beijerinckiaceae;Bosea;Bosea sp.


metaRNA.PFemirge\_8290\_0.000001


JN680618.1.1455 Bacteria;Firmicutes;Clostridia;Oscillos ... icoccaceae;UCG-008;uncultured Ruminococcaceae bacterium


HQ697694.1.1452 Bacteria;Proteobacteria;Alphaproteobact ... gnetospirillaceae;Magnetospirillum;uncultured bacterium


metaRNA.PFemirge\_24146\_0.000003


EU453128.1.1406 Bacteria;Firmicutes;Clostridia;Lachnosp ... ospiraceae;Lachnospiraceae UCG-004;uncultured bacterium


GQ360012.1.1488 Bacteria;Bacteroidota;Bacteroidia;Bacteroidales;Tannerellaceae;Macellibacteroides;uncultured bacterium


EF632748.1.1502 Bacteria;Firmicutes;Clostridia;Clostrid ... iaceae;Clostridium sensu stricto 8;uncultured bacterium


metaRNA.PFemirge\_27575\_0.000002


metaRNA.PFemirge\_19251\_0.000448


metaRNA.PFemirge\_12307\_0.000001


metaRNA.PFemirge\_8305\_0.000107


metaRNA.PFemirge\_19773\_0.000009


JF709929.1.1398 Bacteria;Firmicutes;Bacilli;Exiguobacterales;Exiguobacteraceae;Exiguobacterium;uncultured bacterium


metaRNA.PFemirge\_20165\_0.000003


metaRNA.PFemirge\_15396\_0.000009


metaRNA.PFtrusted\_13\_0


metaRNA.PFemirge\_28136\_0.000016


LT223609.1.1490 Bacteria;Bacteroidota;Bacteroidia;Bacteroidales;Tannerellaceae;Parabacteroides;Parabacteroides faecis


metaRNA.PFemirge\_7707\_0.000007


JF824799.1.1423 Bacteria;Bacteroidota;Bacteroidia;Bacteroidales;Rikenellaceae;Alistipes;Alistipes timonensis JC136


metaRNA.PFemirge\_2761\_0.000011


KC431919.1.1365 Bacteria;Proteobacteria;Gammaproteobacteria;Legionellales;Legionellaceae;Legionella;uncultured bacterium


metaRNA.PFemirge\_21199\_0.000006


LC222972.1.1573 Eukaryota;Amorphea;Amoebozoa;uncultured eukaryote


metaRNA.PFemirge\_17368\_0.000291


metaRNA.PFemirge\_21495\_0.000001


JN680650.1.1446 Bacteria;Firmicutes;Clostridia;Oscillos ... aceae;Ruminococcus;uncultured Lachnospiraceae bacterium


metaRNA.PFemirge\_2918\_0.000022


metaRNA.PFemirge\_23080\_0.000007


DQ799295.1.1402 Bacteria;Bacteroidota;Bacteroidia;Bacteroidales;Bacteroidaceae;Bacteroides;uncultured bacterium


metaRNA.PFemirge\_22312\_0.001807


HQ777443.1.1457 Bacteria;Bacteroidota;Bacteroidia;Bacteroidales;Bacteroidaceae;Bacteroides;uncultured organism


metaRNA.PFemirge\_8123\_0.000005


metaRNA.PFemirge\_27894\_0.000013


metaRNA.PFemirge\_7454\_0.000029


metaRNA.PFemirge\_30242\_0.000027


metaRNA.PFemirge\_15962\_0.019534


metaRNA.PFemirge\_7092\_0.000017


metaRNA.PFemirge\_3306\_0.002933


FJ184323.1.1507 Bacteria;Proteobacteria;Gammaproteobact ... homonadaceae;Stenotrophomonas;uncultured soil bacterium


metaRNA.PFemirge\_10948\_0.000015


metaRNA.PFemirge\_12364\_0.000025


EU655834.1.1314 Bacteria;Firmicutes;Clostridia;Oscillospirales;Ruminococcaceae;UBA1819;uncultured bacterium


EU467709.1.1380 Bacteria;Bacteroidota;Bacteroidia;Bacteroidales;Bacteroidaceae;Bacteroides;uncultured bacterium


metaRNA.PFemirge\_6899\_0.000172


DQ807465.1.1395 Bacteria;Firmicutes;Clostridia;Oscillospirales;Oscillospiraceae;uncultured;uncultured bacterium


DQ795028.1.1385 Bacteria;Firmicutes;Clostridia;Oscillospirales;Oscillospiraceae;UCG-002;uncultured bacterium


metaRNA.PFemirge\_27610\_0.000006


metaRNA.PFemirge\_19378\_0.000042


metaRNA.PFemirge\_24401\_0.000007


metaRNA.PFemirge\_24201\_0.000316


EU772735.1.1293 Bacteria;Firmicutes;Clostridia;Oscillospirales;Butyricicoccaceae;uncultured;uncultured bacterium


CR933273.1.1287 Bacteria;Firmicutes;Clostridia;Lachnosp ... ospiraceae;Lachnospiraceae UCG-004;uncultured bacterium


metaRNA.PFemirge\_27678\_0.002836


metaRNA.PFemirge\_8971\_0.000011


metaRNA.PFemirge\_26157\_0.000005


metaRNA.PFemirge\_4857\_0.000005


EU454443.1.1393 Bacteria;Firmicutes;Clostridia;Lachnosp ... ospiraceae;Lachnospiraceae UCG-008;uncultured bacterium


GQ451215.1.1487 Bacteria;Firmicutes;Clostridia;Oscillospirales;Ruminococcaceae;Ruminococcus;uncultured bacterium


EF403674.1.1517 Bacteria;Firmicutes;Negativicutes;Acida ... inococcaceae;Phascolarctobacterium;uncultured bacterium


metaRNA.PFemirge\_12037\_0.000068


metaRNA.PFemirge\_8090\_0.003659


metaRNA.PFemirge\_4647\_0.000088


metaRNA.PFemirge\_21184\_0.000012


metaRNA.PFemirge\_27211\_0.000011


metaRNA.PFemirge\_15810\_0.000104


metaRNA.PFemirge\_27209\_0.000001


KJ624065.29957.31422 Bacteria;Cyanobacteria;Cyanobacteriia;Chloroplast;Trachydiscus minutus


CP026122.3396659.3398209 Bacteria;Proteobacteria;Gammap ... acterales;Aeromonadaceae;Aeromonas;Aeromonas sp. ASNIH5


ATXW01000003.334809.336317 Bacteria;Bacteroidota;Bacter ... ;Rikenellaceae;Rikenella;Rikenella microfusus DSM 15922


metaRNA.PFemirge\_19805\_0.000004


metaRNA.PFemirge\_10137\_0.000214


KC682971.1.1497 Bacteria;Planctomycetota;Planctomycetes;Planctomycetales;Rubinisphaeraceae;SH-PL14;uncultured bacterium


metaRNA.PFemirge\_20726\_0.000004


BABG01001469.752.2264 Bacteria;Firmicutes;Clostridia;La ... ;Lachnospiraceae;Lachnoclostridium;human gut metagenome


EU771988.1.1377 Bacteria;Firmicutes;Clostridia;Oscillospirales;Ruminococcaceae;Fournierella;uncultured bacterium


metaRNA.PFemirge\_24650\_0.000006


metaRNA.PFemirge\_6423\_0.000004


EU469473.1.1385 Bacteria;Firmicutes;Clostridia;Lachnosp ... ae;[Ruminococcus] gauvreauii group;uncultured bacterium


metaRNA.PFemirge\_2460\_0.000052


metaRNA.PFemirge\_28014\_0.000010


metaRNA.PFemirge\_4651\_0.000048


JQ923952.1.1496 Bacteria;Proteobacteria;Gammaproteobact ... yclaceae;Candidatus Accumulibacter;uncultured bacterium


metaRNA.PFemirge\_22064\_0.000005


metaRNA.PFemirge\_20408\_0.000001


metaRNA.PFemirge\_14287\_0.000002


metaRNA.PFemirge\_15034\_0.000305


metaRNA.PFemirge\_10080\_0.000615


metaRNA.PFemirge\_13047\_0.000042


KM360180.1.1492 Bacteria;Firmicutes;Clostridia;Lachnospirales;Lachnospiraceae;Lachnoclostridium;Clostridium sp. BPY5


metaRNA.PFemirge\_19857\_0.000002


metaRNA.PFspades\_42\_1972.710648


metaRNA.PFemirge\_26291\_0.000003


KP411849.1.1479 Bacteria;Myxococcota;Myxococcia;Myxococcales;Myxococcaceae;P3OB-42;uncultured bacterium


metaRNA.PFemirge\_24437\_0.000003


metaRNA.PFemirge\_30256\_0.000020


metaRNA.PFemirge\_22451\_0.000005


metaRNA.PFemirge\_27690\_0.000328


metaRNA.PFemirge\_12921\_0.000010


metaRNA.PFemirge\_9718\_0.000112


metaRNA.PFemirge\_27760\_0.000012


metaRNA.PFemirge\_22986\_0.000045


metaRNA.PFemirge\_6196\_0.000004


metaRNA.PFemirge\_23786\_0.000024


metaRNA.PFemirge\_13718\_0.000008


metaRNA.PFemirge\_11626\_0.000007


metaRNA.PFemirge\_4783\_0.000126


metaRNA.PFemirge\_19494\_0.000005


metaRNA.PFemirge\_27532\_0.000021


metaRNA.PFemirge\_12202\_0.000068


AB721051.1.1697 Eukaryota;Amorphea;Amoebozoa;SAR;Strame ... omycetes;Sorodiplophrys;uncultured freshwater eukaryote


EU778697.1.1353 Bacteria;Bacteroidota;Bacteroidia;Bacteroidales;Rikenellaceae;SP3-e08;uncultured bacterium


metaRNA.PFemirge\_16650\_0.000005


metaRNA.PFemirge\_3806\_0.000001


DQ809702.1.1364 Bacteria;Firmicutes;Clostridia;Peptostr ... s;Peptostreptococcaceae;uncultured;uncultured bacterium


metaRNA.PFemirge\_12306\_0.000010


metaRNA.PFemirge\_11662\_0.000003


metaRNA.PFemirge\_5015\_0.000007


metaRNA.PFemirge\_3326\_0.000003


metaRNA.PFemirge\_24079\_0.000050


metaRNA.PFemirge\_6695\_0.000395


HQ741592.1.1434 Bacteria;Firmicutes;Clostridia;Lachnospirales;Lachnospiraceae;Epulopiscium;uncultured bacterium


LC094617.1.1418 Bacteria;Actinobacteriota;Actinobacteri ... cteriaceae;Leifsonia;Microbacteriaceae bacterium MAEY21


KF068124.1.1369 Bacteria;Bacteroidota;Bacteroidia;Bacteroidales;Tannerellaceae;Parabacteroides;uncultured bacterium


KM650312.1.1481 Bacteria;Firmicutes;Negativicutes;Acida ... inococcaceae;uncultured;uncultured Firmicutes bacterium


metaRNA.PFemirge\_15952\_0.000004


metaRNA.PFemirge\_636\_0.000124


metaRNA.PFemirge\_25403\_0.000012


metaRNA.PFemirge\_23804\_0.000232


HQ813653.1.1429 Bacteria;Firmicutes;Clostridia;Oscillospirales;Ruminococcaceae;Subdoligranulum;uncultured organism


metaRNA.PFemirge\_6134\_0.000063


metaRNA.PFemirge\_14297\_0.000030


metaRNA.PFemirge\_11249\_0.000931


DQ676995.1.1498 Bacteria;Actinobacteriota;Actinobacteri ... ae;Propionicicella;iron-reducing enrichment clone Cl-A3


metaRNA.PFemirge\_27695\_0.006100


EU772837.1.1306 Bacteria;Bacteroidota;Bacteroidia;Bacteroidales;Bacteroidaceae;Bacteroides;uncultured bacterium


metaRNA.PFemirge\_15056\_0.000153


metaRNA.PFemirge\_28041\_0.000006


metaRNA.PFemirge\_17190\_0.000412


metaRNA.PFemirge\_10253\_0.000037


metaRNA.PFemirge\_13981\_0.000054


metaRNA.PFemirge\_22977\_0.000003


DQ798058.1.1390 Bacteria;Firmicutes;Clostridia;Lachnosp ... aceae;[Ruminococcus] torques group;uncultured bacterium


metaRNA.PFemirge\_19090\_0.000036


DQ809036.1.1400 Bacteria;Bacteroidota;Bacteroidia;Bacteroidales;Bacteroidaceae;Bacteroides;uncultured bacterium


metaRNA.PFemirge\_15906\_0.000034


CP011914.3962957.3964425 Bacteria;Firmicutes;Clostridia;Eubacteriales;Eubacteriaceae;Eubacterium;Eubacterium limosum


CP009057.2296841.2298384 Bacteria;Bacteroidota;Bacteroidia;Bacteroidales;Bacteroidaceae;Bacteroides;Bacteroides dorei


DQ234243.1.1534 Bacteria;Proteobacteria;Gammaproteobact ... aceae;Sphaerotilus;uncultured Burkholderiales bacterium


metaRNA.PFemirge\_7599\_0.004233


AQHX01000018.1891.3401 Bacteria;Bacteroidota;Bacteroidi ... acteroides salyersiae WAL 10018 = DSM 18765 = JCM 12988


KT072190.1.1634 Eukaryota;SAR;Stramenopiles;Labyrinthulomycetes;Amphitraemidae;uncultured eukaryote


metaRNA.PFemirge\_11344\_0.000006


metaRNA.PFemirge\_12361\_0.000007


FBXW01000008.267.1775 Bacteria;Firmicutes;Clostridia;Oscillospirales;Oscillospiraceae;uncultured;Intestinimonas sp. GM5


metaRNA.PFemirge\_11239\_0.000029


HQ802049.1.1451 Bacteria;Firmicutes;Clostridia;Lachnospirales;Lachnospiraceae;uncultured;uncultured organism


CP022655.4386843.4388399 Bacteria;Firmicutes;Bacilli;Pa ... Paenibacillaceae;Paenibacillus;Paenibacillus sp. RUD330


metaRNA.PFemirge\_3497\_0.000015


metaRNA.PFemirge\_26517\_0.000001


metaRNA.PFemirge\_16370\_0.000246


metaRNA.PFemirge\_22501\_0.000008


KX823723.1.1477 Bacteria;Firmicutes;Clostridia;Oscillospirales;Oscillospiraceae;NK4A214 group;uncultured bacterium


metaRNA.PFemirge\_6591\_0.002127


metaRNA.PFemirge\_10288\_0.000011


EU510219.1.1285 Bacteria;Firmicutes;Clostridia;Lachnosp ... ospiraceae;Lachnospiraceae UCG-008;uncultured bacterium


EU454993.1.1404 Bacteria;Firmicutes;Clostridia;Oscillospirales;Oscillospiraceae;uncultured;uncultured bacterium


metaRNA.PFemirge\_237\_0.000025


AY571491.1.1466 Bacteria;Proteobacteria;Alphaproteobacteria;Rickettsiales;uncultured;uncultured bacterium


JN653038.1.1443 Bacteria;Bacteroidota;Bacteroidia;Bacte ... ceae;uncultured;uncultured Porphyromonadaceae bacterium


metaRNA.PFemirge\_15795\_0.000015


FJ184339.1.1505 Bacteria;Proteobacteria;Gammaproteobact ... homonadaceae;Stenotrophomonas;uncultured soil bacterium


metaRNA.PFemirge\_8831\_0.000040


metaRNA.PFemirge\_4975\_0.000004


metaRNA.PFemirge\_27061\_0.000022


metaRNA.PFemirge\_20331\_0.000022


DQ795751.1.1277 Bacteria;Firmicutes;Clostridia;Oscillospirales;Butyricicoccaceae;Butyricicoccus;uncultured bacterium


metaRNA.PFemirge\_25548\_0.000004


metaRNA.PFemirge\_21816\_0.000002


FJ374184.1.1494 Bacteria;Bacteroidota;Bacteroidia;Bacteroidales;Dysgonomonadaceae;uncultured;uncultured bacterium


FJ365071.1.1328 Bacteria;Proteobacteria;Alphaproteobacteria;Rhodospirillales;uncultured;uncultured bacterium


metaRNA.PFemirge\_17727\_0.000002


metaRNA.PFemirge\_26159\_0.000042


metaRNA.PFemirge\_2928\_0.000009


DQ799377.1.1384 Bacteria;Firmicutes;Clostridia;Oscillospirales;Ruminococcaceae;Ruminococcus;uncultured bacterium


metaRNA.PFemirge\_8730\_0.000006


metaRNA.PFemirge\_12927\_0.000008


DQ815225.1.1263 Bacteria;Desulfobacterota;Desulfovibrio ... les;Desulfovibrionaceae;uncultured;uncultured bacterium


metaRNA.PFemirge\_13253\_0.004234


metaRNA.PFemirge\_23846\_0.000006


metaRNA.PFemirge\_22403\_0.000019


metaRNA.PFemirge\_4771\_0.000016


metaRNA.PFemirge\_16159\_0.001568


metaRNA.PFemirge\_2123\_0.000004


metaRNA.PFemirge\_26364\_0.000031


KY445639.1.1525 Bacteria;Proteobacteria;Gammaproteobacteria;Burkholderiales;Comamonadaceae;uncultured;Ideonella sp.


metaRNA.PFemirge\_19453\_0.000002


metaRNA.PFemirge\_8798\_0.000631


JQ885561.1.1458 Bacteria;Proteobacteria;Gammaproteobact ... robacteriaceae;Enterobacter;uncultured Enterobacter sp.


HQ791782.1.1436 Bacteria;Bacteroidota;Bacteroidia;Bacteroidales;Bacteroidaceae;Bacteroides;uncultured organism


metaRNA.PFemirge\_22723\_0.000013


metaRNA.PFemirge\_24043\_0.000018


metaRNA.PFemirge\_12623\_0.005308


metaRNA.PFemirge\_10687\_0.000639


AF245350.1.1489 Bacteria;Proteobacteria;Gammaproteobact ... les;Rhodocyclaceae;uncultured;uncultured bacterium SA35


metaRNA.PFemirge\_1217\_0.000014


metaRNA.PFemirge\_6540\_0.000006


metaRNA.PFemirge\_13372\_0.003476


EU283367.1.1440 Bacteria;Bacteroidota;Bacteroidia;Chiti ... ae;Haliscomenobacter;uncultured Bacteroidetes bacterium


metaRNA.PFemirge\_16703\_0.000022


metaRNA.PFemirge\_16817\_0.000003


KJ615121.1.1238 Bacteria;Proteobacteria;Alphaproteobacteria;Rhizobiales;Beijerinckiaceae;Bosea;uncultured bacterium


metaRNA.PFemirge\_14690\_0.000022


GQ867583.1.1492 Bacteria;Bacteroidota;Bacteroidia;Bacteroidales;Bacteroidaceae;Bacteroides;uncultured bacterium


metaRNA.PFemirge\_25631\_0.023054


metaRNA.PFemirge\_4071\_0.000143


LT631507.1.1440 Bacteria;Firmicutes;Clostridia;Peptostr ... ovoracaceae;Anaerovorax;Bacilliculturomica massiliensis


LN563559.1.1331 Bacteria;Bacteroidota;Bacteroidia;Cytop ... ophilaceae;Candidatus Amoebophilus;uncultured bacterium


metaRNA.PFemirge\_26455\_0.000004


metaRNA.PFemirge\_4901\_0.000008


HM779020.1.1471 Bacteria;Proteobacteria;Gammaproteobact ... les;Enterobacteriaceae;Plesiomonas;uncultured bacterium


metaRNA.PFemirge\_1548\_0.000008


EU778290.1.1377 Bacteria;Firmicutes;Clostridia;Lachnosp ... aceae;Lachnospiraceae NC2004 group;uncultured bacterium


metaRNA.PFemirge\_14307\_0.000006


GQ493239.1.1362 Bacteria;Firmicutes;Bacilli;Erysipelotr ... stridiaceae;Erysipelatoclostridium;uncultured bacterium


metaRNA.PFemirge\_27763\_0.002974


GQ448681.1.1394 Bacteria;Bacteroidota;Bacteroidia;Bacteroidales;Marinifilaceae;Odoribacter;uncultured bacterium


metaRNA.PFemirge\_7443\_0.002130


metaRNA.PFemirge\_10840\_0.000008


metaRNA.PFemirge\_22148\_0.000002


metaRNA.PFemirge\_27705\_0.000033


metaRNA.PFemirge\_2358\_0.002376


metaRNA.PFemirge\_4741\_0.000796


metaRNA.PFemirge\_6685\_0.000147


DQ809316.1.1289 Bacteria;Firmicutes;Clostridia;Oscillospirales;Ruminococcaceae;Subdoligranulum;uncultured bacterium


metaRNA.PFemirge\_11694\_0.000035


metaRNA.PFemirge\_17975\_0.002583


EU136696.1.1944 Bacteria;Bacteroidota;Bacteroidia;Bacteroidales;Bacteroidaceae;Bacteroides;Bacteroides massiliensis


metaRNA.PFemirge\_20325\_0.000004


metaRNA.PFemirge\_4290\_0.000046


metaRNA.PFemirge\_11874\_0.000453


metaRNA.PFemirge\_21649\_0.000048


metaRNA.PFemirge\_23880\_0.000002


metaRNA.PFemirge\_1497\_0.000007


metaRNA.PFemirge\_16811\_0.000008


HM630239.1.1414 Bacteria;Firmicutes;Bacilli;Erysipelotr ... stridiaceae;Erysipelatoclostridium;uncultured bacterium


HM630236.1.1451 Bacteria;Bacteroidota;Bacteroidia;Bacteroidales;Rikenellaceae;dgA-11 gut group;uncultured bacterium


EU772041.1.1407 Bacteria;Firmicutes;Negativicutes;Acida ... inococcaceae;Phascolarctobacterium;uncultured bacterium


APKA01105891.43.1849 Eukaryota;Amorphea;Obazoa;Opisthok ... zoa;Rotifera;Bdelloidea;Adinetida;Biomphalaria glabrata


metaRNA.PFemirge\_20099\_0.000012


metaRNA.PFemirge\_24632\_0.000006


metaRNA.PFemirge\_24325\_0.000037


metaRNA.PFemirge\_16638\_0.000657


metaRNA.PFemirge\_17976\_0.000032


ABFK02000017.223676.225190 Bacteria;Bacteroidota;Bacter ... ;Rikenellaceae;Alistipes;Alistipes putredinis DSM 17216


metaRNA.PFemirge\_18858\_0.000018


metaRNA.PFemirge\_4069\_0.000010


EU465626.1.1378 Bacteria;Firmicutes;Clostridia;Lachnosp ... ceae;Lachnospiraceae NK4A136 group;uncultured bacterium


EU765828.1.1335 Bacteria;Firmicutes;Clostridia;Oscillospirales;Ruminococcaceae;Faecalibacterium;uncultured bacterium


metaRNA.PFemirge\_23442\_0.000004


metaRNA.PFemirge\_26585\_0.000003


DQ824289.1.1398 Bacteria;Bacteroidota;Bacteroidia;Bacteroidales;Bacteroidaceae;Bacteroides;uncultured bacterium


metaRNA.PFemirge\_18287\_0.000026


GQ135950.1.1352 Bacteria;Firmicutes;Clostridia;Clostrid ... iaceae;Clostridium sensu stricto 1;uncultured bacterium


EF403711.1.1488 Bacteria;Bacteroidota;Bacteroidia;Bacteroidales;Bacteroidaceae;Bacteroides;uncultured bacterium


metaRNA.PFemirge\_16443\_0.000064


AY916210.1.1353 Bacteria;Bacteroidota;Bacteroidia;Bacteroidales;Bacteroidaceae;Bacteroides;uncultured bacterium


metaRNA.PFemirge\_30249\_0.000019


DQ238611.1.1497 Bacteria;Firmicutes;Clostridia;Clostrid ... iaceae;Clostridium sensu stricto 1;uncultured bacterium


AY452004.1.1502 Bacteria;Firmicutes;Clostridia;Lachnosp ... ospiraceae;Lachnospiraceae UCG-004;uncultured bacterium


metaRNA.PFemirge\_20425\_0.000055


metaRNA.PFemirge\_10119\_0.000008


EU774249.1.1294 Bacteria;Firmicutes;Clostridia;Oscillospirales;Butyricicoccaceae;uncultured;uncultured bacterium


AF371540.1.1455 Bacteria;Firmicutes;Clostridia;Lachnospirales;Lachnospiraceae;Blautia;uncultured bacterium


metaRNA.PFemirge\_28024\_0.000027


metaRNA.PFemirge\_7333\_0.000021


DQ798128.1.1365 Bacteria;Firmicutes;Clostridia;Oscillospirales;Ruminococcaceae;Ruminococcus;uncultured bacterium


metaRNA.PFemirge\_28152\_0.000012


EU775698.1.1374 Bacteria;Firmicutes;Clostridia;Clostrid ... iaceae;Clostridium sensu stricto 1;uncultured bacterium


metaRNA.PFemirge\_5891\_0.000001


metaRNA.PFemirge\_26911\_0.000000


metaRNA.PFemirge\_18462\_0.001364


metaRNA.PFemirge\_1171\_0.000094


LK056647.1.1423 Bacteria;Proteobacteria;Alphaproteobact ... ingomonadaceae;Novosphingobium;Novosphingobium piscinae


metaRNA.PFemirge\_27871\_0.000006


EU762749.1.1373 Bacteria;Bacteroidota;Bacteroidia;Bacteroidales;Bacteroidaceae;Bacteroides;uncultured bacterium


DQ824384.1.1400 Bacteria;Bacteroidota;Bacteroidia;Bacteroidales;Bacteroidaceae;Bacteroides;uncultured bacterium


DQ808516.1.1400 Bacteria;Bacteroidota;Bacteroidia;Bacteroidales;Bacteroidaceae;Bacteroides;uncultured bacterium


metaRNA.PFemirge\_15841\_0.000011


metaRNA.PFemirge\_17982\_0.000003


HM778328.1.1446 Bacteria;Proteobacteria;Gammaproteobact ... les;Enterobacteriaceae;Plesiomonas;uncultured bacterium


metaRNA.PFemirge\_12309\_0.000016


metaRNA.PFemirge\_12903\_0.000007


metaRNA.PFemirge\_9028\_0.000019


metaRNA.PFemirge\_17433\_0.000009


metaRNA.PFemirge\_22056\_0.000025


GQ175524.1.1451 Bacteria;Bacteroidota;Bacteroidia;Bacteroidales;Rikenellaceae;Rikenella;uncultured bacterium


EF024902.1.1792 Eukaryota;Archaeplastida;Chloroplastida ... matophyta;Magnoliophyta;Phaseoleae environmental sample


metaRNA.PFemirge\_27774\_0.000013


EU475815.1.1389 Bacteria;Firmicutes;Clostridia;Lachnosp ... ospiraceae;Lachnospiraceae UCG-008;uncultured bacterium


AM743169.416435.417979 Bacteria;Proteobacteria;Gammapro ... eae;Stenotrophomonas;Stenotrophomonas maltophilia K279a


JRGF01000044.3.1537 Bacteria;Bacteroidota;Bacteroidia;Bacteroidales;Rikenellaceae;Alistipes;Alistipes inops


metaRNA.PFemirge\_12955\_0.000010


JN619858.1.1417 Bacteria;Bacteroidota;Bacteroidia;Bacteroidales;Rikenellaceae;Alistipes;uncultured bacterium


metaRNA.PFemirge\_17717\_0.000011


DQ807211.1.1386 Bacteria;Firmicutes;Clostridia;Oscillospirales;Ruminococcaceae;Incertae Sedis;uncultured bacterium


DQ824247.1.1290 Bacteria;Firmicutes;Clostridia;Lachnospirales;Lachnospiraceae;Lachnoclostridium;uncultured bacterium


metaRNA.PFemirge\_6704\_0.000046


metaRNA.PFemirge\_21460\_0.000044


metaRNA.PFemirge\_12416\_0.000004


metaRNA.PFemirge\_26525\_0.000032


metaRNA.PFemirge\_18548\_0.000043


metaRNA.PFemirge\_15025\_0.000015


metaRNA.PFemirge\_13515\_0.000014


metaRNA.PFemirge\_10448\_0.000001


metaRNA.PFemirge\_12401\_0.000009


metaRNA.PFemirge\_12011\_0.000016


metaRNA.PFemirge\_15966\_0.000007


metaRNA.PFemirge\_26218\_0.000001


EU507529.1.1407 Bacteria;Firmicutes;Clostridia;Oscillospirales;Butyricicoccaceae;UCG-009;uncultured bacterium


metaRNA.PFemirge\_22735\_0.000002


HQ827970.1.1471 Bacteria;Bacteroidota;Bacteroidia;Chitinophagales;Saprospiraceae;uncultured;uncultured bacterium


MG428876.1.1497 Bacteria;Proteobacteria;Gammaproteobact ... bacterales;Aeromonadaceae;Aeromonas;Aeromonas encheleia


metaRNA.PFemirge\_21386\_0.000316


EU772234.1.1393 Bacteria;Firmicutes;Clostridia;Christen ... ceae;Christensenellaceae R-7 group;uncultured bacterium


AF011457.1.2048 Eukaryota;Discoba;Discicristata;Heterolobosea;Tetramitia;Naegleria;Naegleria galeacystis


metaRNA.PFemirge\_25365\_0.000174


metaRNA.PFemirge\_25640\_0.000004


CP026007.74890.76443 Bacteria;Firmicutes;Bacilli;Bacillales;Planococcaceae;Lysinibacillus;Lysinibacillus sp. YS11


JN869169.1.1481 Bacteria;Proteobacteria;Alphaproteobacteria;Reyranellales;Reyranellaceae;Reyranella;uncultured bacterium


GU222249.1.1456 Bacteria;Proteobacteria;Gammaproteobact ... iales;Comamonadaceae;Simplicispira;uncultured bacterium


EU774669.1.1387 Bacteria;Firmicutes;Clostridia;Peptostr ... caceae;[Eubacterium] nodatum group;uncultured bacterium


FR669217.1.1452 Bacteria;Firmicutes;Clostridia;Oscillospirales;Oscillospiraceae;Colidextribacter;uncultured bacterium


metaRNA.PFemirge\_9950\_0.013428


metaRNA.PFemirge\_23456\_0.000422


metaRNA.PFemirge\_13026\_0.000916


metaRNA.PFemirge\_2279\_0.000061


JN680597.1.1451 Bacteria;Bacteroidota;Bacteroidia;Bacte ... enellaceae;Alistipes;uncultured Rikenellaceae bacterium


EU510318.1.1396 Bacteria;Firmicutes;Clostridia;Oscillospirales;Oscillospiraceae;Colidextribacter;uncultured bacterium


EF400726.1.1488 Bacteria;Bacteroidota;Bacteroidia;Bacteroidales;Marinifilaceae;Odoribacter;uncultured bacterium


metaRNA.PFemirge\_25367\_0.000009


HQ761439.1.1422 Bacteria;Firmicutes;Clostridia;Clostridia vadinBB60 group;uncultured organism


FM957479.1.1497 Bacteria;Proteobacteria;Gammaproteobacteria;Enterobacterales;Vibrionaceae;Vibrio;Vibrio sp. MY-2008-U67


KC253315.1.1470 Bacteria;Bacteroidota;Bacteroidia;Cytophagales;Spirosomaceae;Runella;uncultured bacterium


BAAY01003437.486.1979 Bacteria;Firmicutes;Clostridia;Lachnospirales;Lachnospiraceae;uncultured;human gut metagenome


metaRNA.PFemirge\_22671\_0.000052


metaRNA.PFemirge\_20021\_0.000069


KM205539.1.1438 Bacteria;Firmicutes;Clostridia;Clostrid ... e;Clostridium sensu stricto 3;uncultured soil bacterium


metaRNA.PFemirge\_22623\_0.000102


metaRNA.PFemirge\_6724\_0.004080


JX198651.1.1513 Bacteria;Firmicutes;Clostridia;Oscillospirales;Butyricicoccaceae;UCG-009;uncultured bacterium


KF318251.1.1396 Bacteria;Firmicutes;Clostridia;Oscillos ... inococcaceae;uncultured;uncultured Firmicutes bacterium


FPLS01033705.10.1524 Bacteria;Planctomycetota;Planctomycetes;Pirellulales;Pirellulaceae;Pir4 lineage;metagenome


metaRNA.PFemirge\_16632\_0.000037


metaRNA.PFemirge\_20030\_0.000000


metaRNA.PFemirge\_16642\_0.000815


metaRNA.PFemirge\_26353\_0.002068


EU457166.1.1396 Bacteria;Bacteroidota;Bacteroidia;Bacteroidales;Bacteroidaceae;Bacteroides;uncultured bacterium


metaRNA.PFemirge\_11749\_0.000005


metaRNA.PFemirge\_21157\_0.000045


metaRNA.PFemirge\_6554\_0.000060


metaRNA.PFemirge\_2582\_0.000049


metaRNA.PFspades\_11\_30.944644


metaRNA.PFemirge\_27697\_0.002996


metaRNA.PFemirge\_7133\_0.000004


metaRNA.PFemirge\_11932\_0.000002


metaRNA.PFemirge\_13684\_0.000421


metaRNA.PFemirge\_23134\_0.000040


metaRNA.PFspades\_7\_377.945982


metaRNA.PFemirge\_6917\_0.003777


metaRNA.PFemirge\_4199\_0.004230


AB264620.1.1463 Bacteria;Firmicutes;Clostridia;Lachnosp ... ospiraceae NC2004 group;Clostridiaceae bacterium bSSV31


metaRNA.PFemirge\_16901\_0.000019


metaRNA.PFemirge\_1876\_0.000003


GQ135998.1.1368 Bacteria;Firmicutes;Clostridia;Peptostr ... llales;Anaerovoracaceae;uncultured;uncultured bacterium


metaRNA.PFemirge\_15725\_0.000455


metaRNA.PFemirge\_18125\_0.000005


metaRNA.PFemirge\_12452\_0.000004


EU776450.1.1387 Bacteria;Firmicutes;Clostridia;Oscillospirales;Ruminococcaceae;Ruminococcus;uncultured bacterium


DQ802739.1.1400 Bacteria;Firmicutes;Clostridia;Oscillospirales;Ruminococcaceae;Incertae Sedis;uncultured bacterium


DQ808514.1.1304 Bacteria;Bacteroidota;Bacteroidia;Bacteroidales;Bacteroidaceae;Bacteroides;uncultured bacterium


metaRNA.PFemirge\_12918\_0.003212


AJ421915.1.1402 Bacteria;Proteobacteria;Gammaproteobact ... monadaceae;Sphaerotilus;uncultured beta proteobacterium


EU104138.1.1438 Bacteria;Bacteroidota;Bacteroidia;Chitinophagales;Saprospiraceae;Haliscomenobacter;uncultured bacterium


JN012636.1.1457 Bacteria;Firmicutes;Clostridia;Lachnospirales;Lachnospiraceae;uncultured;uncultured bacterium


metaRNA.PFemirge\_10492\_0.000020


metaRNA.PFemirge\_16640\_0.000098


HQ741327.1.1434 Bacteria;Firmicutes;Clostridia;Clostrid ... iaceae;Clostridium sensu stricto 1;uncultured bacterium


metaRNA.PFemirge\_23333\_0.000004


metaRNA.PFemirge\_6935\_0.004836


JN680601.1.1450 Bacteria;Bacteroidota;Bacteroidia;Bacte ... enellaceae;Alistipes;uncultured Rikenellaceae bacterium


FJ534971.1.1485 Bacteria;Bacteroidota;Bacteroidia;Bacteroidales;Rikenellaceae;uncultured;uncultured bacterium


metaRNA.PFemirge\_13238\_0.001251


FJ542833.1.1492 Bacteria;Verrucomicrobiota;Verrucomicro ... ;Terrimicrobium;uncultured Verrucomicrobiales bacterium


metaRNA.PFemirge\_23694\_0.000013


metaRNA.PFspades\_33\_227.152688


metaRNA.PFemirge\_5805\_0.000005


EU777165.1.1344 Bacteria;Firmicutes;Clostridia;Oscillospirales;Oscillospiraceae;Oscillibacter;uncultured bacterium


metaRNA.PFemirge\_20173\_0.000015


metaRNA.PFemirge\_26412\_0.000030


metaRNA.PFemirge\_5013\_0.000028


DQ801177.1.1348 Bacteria;Bacteroidota;Bacteroidia;Bacteroidales;Bacteroidaceae;Bacteroides;uncultured bacterium


DQ798901.1.1391 Bacteria;Firmicutes;Clostridia;Lachnospirales;Lachnospiraceae;Dorea;uncultured bacterium


metaRNA.PFemirge\_17188\_0.002435


JN868790.1.1523 Bacteria;Proteobacteria;Gammaproteobact ... deriales;Comamonadaceae;uncultured;uncultured bacterium


metaRNA.PFemirge\_13727\_0.000597


metaRNA.PFemirge\_17446\_0.000009


metaRNA.PFemirge\_13802\_0.000006


metaRNA.PFemirge\_30252\_0.000015


metaRNA.PFemirge\_20200\_0.000004


JN688021.1.1446 Bacteria;Bacteroidota;Bacteroidia;Bacte ... aceae;uncultured;bacterium enrichment culture clone M02


metaRNA.PFemirge\_2576\_0.000068


KR029407.1.1499 Bacteria;Proteobacteria;Gammaproteobact ... ;Yersiniaceae;Serratia;Serratia sp. mixed culture X13-6


metaRNA.PFemirge\_17776\_0.000002


EU776418.1.1406 Bacteria;Firmicutes;Clostridia;Lachnosp ... ospiraceae;Lachnospiraceae UCG-004;uncultured bacterium


metaRNA.PFemirge\_19442\_0.000006


metaRNA.PFemirge\_12198\_0.000070


metaRNA.PFemirge\_4585\_0.000735


metaRNA.PFemirge\_24874\_0.000003


metaRNA.PFemirge\_18832\_0.000298


metaRNA.PFemirge\_9752\_0.000858


metaRNA.PFemirge\_6703\_0.000042


KJ600954.1.1517 Bacteria;Proteobacteria;Gammaproteobact ... ;Xanthomonadaceae;Stenotrophomonas;uncultured bacterium


metaRNA.PFemirge\_7230\_0.000729


metaRNA.PFemirge\_8465\_0.000009


metaRNA.PFemirge\_16674\_0.001307


metaRNA.PFemirge\_18240\_0.000015


KX508033.1.1494 Bacteria;Proteobacteria;Gammaproteobact ... iales;Comamonadaceae;Aquabacterium;uncultured bacterium


HQ816006.1.1448 Bacteria;Bacteroidota;Bacteroidia;Bacteroidales;Bacteroidaceae;Bacteroides;uncultured organism


metaRNA.PFemirge\_28017\_0.000059


metaRNA.PFemirge\_12901\_0.000019


metaRNA.PFemirge\_16522\_0.000074


metaRNA.PFemirge\_5733\_0.000056


metaRNA.PFemirge\_26363\_0.000004


metaRNA.PFemirge\_3462\_0.001424


FJ825520.1.1372 Bacteria;Firmicutes;Clostridia;Christen ... ceae;Christensenellaceae R-7 group;uncultured bacterium


metaRNA.PFemirge\_17001\_0.001040


metaRNA.PFemirge\_2897\_0.000694


metaRNA.PFemirge\_23451\_0.000011


metaRNA.PFemirge\_20782\_0.000343


metaRNA.PFemirge\_24960\_0.000086


metaRNA.PFemirge\_28020\_0.000045


metaRNA.PFemirge\_8463\_0.000025


metaRNA.PFemirge\_18986\_0.000116


metaRNA.PFemirge\_16610\_0.000002


metaRNA.PFemirge\_16160\_0.000419


metaRNA.PFemirge\_8001\_0.000285


metaRNA.PFemirge\_20993\_0.000007


metaRNA.PFemirge\_11838\_0.000075


JF513192.1.1510 Bacteria;Proteobacteria;Gammaproteobact ... les;Moraxellaceae;Acinetobacter;Acinetobacter baumannii


metaRNA.PFemirge\_15943\_0.000843


metaRNA.PFemirge\_5412\_0.003665


metaRNA.PFemirge\_28072\_0.000003


metaRNA.PFemirge\_2903\_0.000018


metaRNA.PFspades\_12\_8.205331


metaRNA.PFemirge\_13667\_0.000145


EU466072.1.1394 Bacteria;Firmicutes;Clostridia;Christen ... ceae;Christensenellaceae R-7 group;uncultured bacterium


metaRNA.PFemirge\_19666\_0.000065


metaRNA.PFemirge\_23640\_0.000013


metaRNA.PFemirge\_13642\_0.000107


metaRNA.PFemirge\_17158\_0.000007


metaRNA.PFemirge\_16574\_0.000014


metaRNA.PFemirge\_11223\_0.000214


LOQJ01002372.643.1942 Eukaryota;Amorphea;Obazoa;Opistho ... charomycetaceae;Saccharomyces;Saccharomyces pastorianus


metaRNA.PFemirge\_27410\_0.000001


AGTN01029961.1048.2511 Bacteria;Proteobacteria;Alphapro ... netospirillaceae;Magnetospirillum;bioreactor metagenome


EU458567.1.1358 Bacteria;Proteobacteria;Alphaproteobacteria;Rhodospirillales;uncultured;uncultured bacterium


metaRNA.PFemirge\_27870\_0.000005


metaRNA.PFemirge\_11718\_0.000005


metaRNA.PFemirge\_24801\_0.000649


metaRNA.PFemirge\_17589\_0.000056


metaRNA.PFemirge\_22821\_0.000029


EF406661.1.1497 Bacteria;Firmicutes;Clostridia;Lachnosp ... aceae;[Ruminococcus] torques group;uncultured bacterium


FJ880485.1.1495 Bacteria;Firmicutes;Clostridia;Lachnospirales;Lachnospiraceae;uncultured;uncultured bacterium


EF098082.1.1412 Bacteria;Bacteroidota;Bacteroidia;Bacteroidales;Muribaculaceae;uncultured bacterium


metaRNA.PFemirge\_25653\_0.000004


metaRNA.PFemirge\_12504\_0.000006


metaRNA.PFemirge\_30263\_0.000013


DQ799802.1.1391 Bacteria;Firmicutes;Clostridia;Lachnosp ... ospiraceae;Lachnospiraceae UCG-008;uncultured bacterium


metaRNA.PFemirge\_25011\_0.000022


MG428882.1.1489 Bacteria;Bacteroidota;Bacteroidia;Bacte ... eroidaceae;Bacteroides;Bacteroides paurosaccharolyticus


metaRNA.PFemirge\_4824\_0.000005


metaRNA.PFemirge\_28001\_0.000400


DQ797781.1.1401 Bacteria;Bacteroidota;Bacteroidia;Bacteroidales;Tannerellaceae;Parabacteroides;uncultured bacterium


metaRNA.PFemirge\_21595\_0.000006


metaRNA.PFemirge\_7880\_0.000005


CP022983.633553.635105 Bacteria;Firmicutes;Bacilli;Bacillales;Bacillaceae;Bacillus;Bacillus kochii


metaRNA.PFemirge\_18629\_0.001209


metaRNA.PFemirge\_4517\_0.016085


metaRNA.PFemirge\_109\_0.004985


metaRNA.PFemirge\_6699\_0.011454


MF040523.1.1509 Bacteria;Verrucomicrobiota;Verrucomicro ... robiaceae;FukuN18 freshwater group;uncultured bacterium


metaRNA.PFemirge\_3501\_0.000070


metaRNA.PFemirge\_23661\_0.000023


FPLS01048670.18.1522 Bacteria;Bacteroidota;Bacteroidia;Bacteroidales;Paludibacteraceae;uncultured;metagenome


HQ681763.1.1459 Bacteria;Firmicutes;Clostridia;Oscillospirales;Ruminococcaceae;Ruminococcus;uncultured bacterium


HQ792908.1.1451 Bacteria;Firmicutes;Clostridia;Oscillospirales;Butyricicoccaceae;UCG-008;uncultured organism


HG792196.1.1487 Bacteria;Firmicutes;Bacilli;Erysipelotr ... pelatoclostridiaceae;Coprobacillus;uncultured bacterium


metaRNA.PFemirge\_12199\_0.000012


metaRNA.PFemirge\_14697\_0.000010


KC633480.1.1496 Bacteria;Proteobacteria;Gammaproteobact ... mamonadaceae;Acidovorax;uncultured beta proteobacterium


KJ563218.1.1771 Eukaryota;Amorphea;Obazoa;Opisthokonta; ... dermataceae;Paraphysoderma;Paraphysoderma sedebokerense


HQ757038.1.1425 Bacteria;Firmicutes;Clostridia;Lachnospirales;Lachnospiraceae;Blautia;uncultured organism


metaRNA.PFemirge\_18757\_0.000009


EU560770.1.1499 Bacteria;Proteobacteria;Gammaproteobact ... dales;Pseudomonadaceae;Pseudomonas;uncultured bacterium


metaRNA.PFemirge\_1882\_0.000004


KJ874069.1.1323 Bacteria;Bacteroidota;Bacteroidia;Bacte ... laceae;Rikenellaceae RC9 gut group;uncultured bacterium


FN667250.1.1445 Bacteria;Firmicutes;Clostridia;Oscillos ... inococcaceae;Anaerotruncus;uncultured compost bacterium


CDZS01002192.2025.3531 Bacteria;Bacteroidota;Bacteroidia;Bacteroidales;Bacteroidaceae;Bacteroides;gut metagenome


ABJL02000006.58.1568 Bacteria;Bacteroidota;Bacteroidia; ... oidaceae;Bacteroides;Bacteroides intestinalis DSM 17393


GQ871719.1.1509 Bacteria;Firmicutes;Bacilli;Erysipelotrichales;Erysipelotrichaceae;Faecalitalea;uncultured bacterium


M97959.3.1817 Eukaryota;SAR;Stramenopiles;Ochrophyta;Phaeophyceae;Fucales;Fucus;Fucus distichus


metaRNA.PFemirge\_20262\_0.000004


EU542517.1.1444 Bacteria;Firmicutes;Clostridia;Oscillospirales;Oscillospiraceae;Intestinimonas;uncultured bacterium


metaRNA.PFemirge\_25880\_0.000002


metaRNA.PFemirge\_7412\_0.002524


EU774505.1.1241 Bacteria;Firmicutes;Clostridia;Clostridia UCG-014;uncultured bacterium


metaRNA.PFemirge\_1589\_0.000007


HQ793252.1.1444 Bacteria;Bacteroidota;Bacteroidia;Bacteroidales;Bacteroidaceae;Bacteroides;uncultured organism


GQ448867.1.1381 Bacteria;Bacteroidota;Bacteroidia;Bacteroidales;Tannerellaceae;Parabacteroides;uncultured bacterium


HG934468.2032323.2033833 Bacteria;Bacteroidota;Bacteroi ... dales;Rikenellaceae;Mucinivorans;Mucinivorans hirudinis


EU771706.1.1407 Bacteria;Verrucomicrobiota;Lentisphaeria;Victivallales;Victivallaceae;uncultured bacterium


metaRNA.PFemirge\_16289\_0.000095


metaRNA.PFemirge\_3395\_0.000008


metaRNA.PFemirge\_23292\_0.000017


AB547644.1.1485 Bacteria;Bacteroidota;Bacteroidia;Bacteroidales;Bacteroidaceae;Bacteroides;Bacteroides oleiciplenus


metaRNA.PFemirge\_8584\_0.000080


EU775026.1.1400 Bacteria;Bacteroidota;Bacteroidia;Bacteroidales;Bacteroidaceae;Bacteroides;uncultured bacterium


metaRNA.PFemirge\_26231\_0.000007


metaRNA.PFemirge\_20738\_0.000001


metaRNA.PFemirge\_25220\_0.000001


metaRNA.PFemirge\_5036\_0.000110


MF175255.1.1500 Bacteria;Firmicutes;Clostridia;Lachnospirales;Lachnospiraceae;Lachnoclostridium;Clostridium sp.


metaRNA.PFemirge\_11629\_0.000050


metaRNA.PFemirge\_3185\_0.000003


EU771874.1.1278 Bacteria;Firmicutes;Bacilli;Erysipelotrichales;Erysipelatoclostridiaceae;uncultured;uncultured bacterium


metaRNA.PFemirge\_23766\_0.000010


metaRNA.PFemirge\_12736\_0.000007


metaRNA.PFemirge\_17308\_0.000014


metaRNA.PFemirge\_11040\_0.000003


metaRNA.PFemirge\_15322\_0.000011


DQ824104.1.1398 Bacteria;Firmicutes;Clostridia;Oscillospirales;Butyricicoccaceae;Butyricicoccus;uncultured bacterium


DQ807770.1.1395 Bacteria;Firmicutes;Clostridia;Lachnospirales;Lachnospiraceae;Roseburia;uncultured bacterium


HQ119254.1.1464 Bacteria;Proteobacteria;Alphaproteobacteria;Dongiales;Dongiaceae;Dongia;uncultured bacterium


metaRNA.PFemirge\_14375\_0.000013


MG367106.1.1496 Bacteria;Firmicutes;Clostridia;Lachnosp ... ostridium;uncultured Clostridiales Family XIV bacterium


JN217178.1.1510 Bacteria;Proteobacteria;Gammaproteobact ... les;Acidithiobacillaceae;KCM-B-112;uncultured bacterium


metaRNA.PFemirge\_1731\_0.000166


HQ761737.1.1467 Bacteria;Bacteroidota;Bacteroidia;Bacteroidales;Tannerellaceae;Parabacteroides;uncultured organism


metaRNA.PFemirge\_9898\_0.000807


metaRNA.PFemirge\_7513\_0.001841


metaRNA.PFemirge\_12177\_0.000004


EU772065.1.1302 Bacteria;Bacteroidota;Bacteroidia;Bacte ... votellaceae;Prevotellaceae UCG-004;uncultured bacterium


EU471853.1.1298 Bacteria;Firmicutes;Clostridia;Christen ... ceae;Christensenellaceae R-7 group;uncultured bacterium


JTDA01000039.175.1712 Bacteria;Bacteroidota;Bacteroidia;Bacteroidales;Barnesiellaceae;Coprobacter;Coprobacter secundus


metaRNA.PFemirge\_14455\_0.000015


FJ534957.1.1501 Bacteria;Firmicutes;Clostridia;Lachnosp ... aceae;Lachnospiraceae FCS020 group;uncultured bacterium


metaRNA.PFemirge\_3509\_0.047992


metaRNA.PFemirge\_25313\_0.000007


GQ871737.1.1494 Bacteria;Firmicutes;Clostridia;Oscillospirales;Ruminococcaceae;Anaerofilum;uncultured bacterium


metaRNA.PFemirge\_27621\_0.000005


metaRNA.PFemirge\_17247\_0.000036


metaRNA.PFemirge\_2636\_0.000336


metaRNA.PFemirge\_19535\_0.000007


metaRNA.PFemirge\_24291\_0.000002


CYSK01000016.265066.266548 Bacteria;Firmicutes;Clostrid ... ostridium sensu stricto 8;Clostridiaceae bacterium mt10


metaRNA.PFemirge\_4544\_0.000000


metaRNA.PFemirge\_16459\_0.000006


AY983363.1.1360 Bacteria;Bacteroidota;Bacteroidia;Bacteroidales;Bacteroidaceae;Bacteroides;uncultured bacterium


DQ803378.1.1392 Bacteria;Firmicutes;Clostridia;Lachnospirales;Lachnospiraceae;uncultured;uncultured bacterium


metaRNA.PFemirge\_1632\_0.000008


CP022115.2965538.2967082 Bacteria;Proteobacteria;Gammap ... es;Aquaspirillaceae;Laribacter;Laribacter hongkongensis


metaRNA.PFemirge\_27888\_0.000148


metaRNA.PFemirge\_24550\_0.000001


AATF01000761.367.1874 Bacteria;Firmicutes;Clostridia;Oscillospirales;Ruminococcaceae;Ruminococcus;mouse gut metagenome


metaRNA.PFemirge\_18904\_0.000005


metaRNA.PFemirge\_27531\_0.000013


FJ658504.1.1402 Bacteria;Proteobacteria;Gammaproteobact ... cterales;Morganellaceae;Morganella;uncultured bacterium


metaRNA.PFemirge\_17932\_0.000003


DQ799104.1.1398 Bacteria;Bacteroidota;Bacteroidia;Bacteroidales;Bacteroidaceae;Bacteroides;uncultured bacterium


AJ289162.1.1486 Bacteria;Proteobacteria;Gammaproteobact ... ales;Cellvibrionaceae;Cellvibrio;Cellvibrio gandavensis


metaRNA.PFemirge\_26443\_0.000017


EF401177.1.1491 Bacteria;Firmicutes;Clostridia;Lachnospirales;Lachnospiraceae;uncultured;uncultured bacterium


metaRNA.PFemirge\_7266\_0.000005


EU464167.1.1397 Bacteria;Bacteroidota;Bacteroidia;Bacteroidales;Barnesiellaceae;uncultured;uncultured bacterium


metaRNA.PFemirge\_19649\_0.000406


AB614872.1.1469 Bacteria;Verrucomicrobiota;Lentisphaeria;Victivallales;Victivallaceae;uncultured rumen bacterium


metaRNA.PFemirge\_1117\_0.000229


metaRNA.PFemirge\_14769\_0.000018


metaRNA.PFemirge\_1829\_0.000025


metaRNA.PFemirge\_12625\_0.000154


GU303678.1.1524 Bacteria;Firmicutes;Bacilli;RF39;uncultured rumen bacterium


metaRNA.PFemirge\_16111\_0.001253


metaRNA.PFemirge\_25780\_0.000272


metaRNA.PFemirge\_11693\_0.000022


HQ761612.1.1444 Bacteria;Bacteroidota;Bacteroidia;Bacteroidales;Tannerellaceae;Parabacteroides;uncultured organism


metaRNA.PFemirge\_16056\_0.000004


EF667904.1.1463 Bacteria;Bacteroidota;Bacteroidia;Cytop ... rosomaceae;Emticicia;uncultured Bacteroidetes bacterium


LT598589.1.1424 Bacteria;Firmicutes;Clostridia;Oscillospirales;Butyricicoccaceae;UCG-008;Intestinibacillus massiliensis


metaRNA.PFemirge\_30254\_0.000016


GU056304.1.1502 Bacteria;Proteobacteria;Gammaproteobact ... e;Azonexus;Azonexus sp. enrichment culture clone ECC4-1


EU503945.1.1397 Bacteria;Firmicutes;Clostridia;Oscillospirales;Butyricicoccaceae;Butyricicoccus;uncultured bacterium


metaRNA.PFemirge\_3014\_0.000005


AB901579.1.1807 Eukaryota;Amorphea;Obazoa;Opisthokonta; ... ozoa;Rotifera;Monogononta;Ploimida;uncultured eukaryote


metaRNA.PFemirge\_27324\_0.000004


metaRNA.PFemirge\_26444\_0.000026


AJ629069.1.1499 Bacteria;Firmicutes;Bacilli;Erysipelotrichales;Erysipelotrichaceae;Breznakia;Breznakia pachnodae


metaRNA.PFemirge\_11095\_0.000044


metaRNA.PFemirge\_22844\_0.000909


LN870298.1.1495 Bacteria;Firmicutes;Clostridia;Oscillospirales;Oscillospiraceae;Intestinimonas;Intestinimonas timonensis


metaRNA.PFspades\_23\_155.328872


metaRNA.PFemirge\_20039\_0.000011


metaRNA.PFemirge\_6863\_0.000788


metaRNA.PFemirge\_4742\_0.000004


JX000037.1.1482 Bacteria;Firmicutes;Clostridia;Oscillospirales;Oscillospiraceae;uncultured;uncultured bacterium


KJ807937.1.1493 Bacteria;Proteobacteria;Gammaproteobact ... deriales;Comamonadaceae;uncultured;uncultured bacterium


metaRNA.PFemirge\_35\_0.000002


metaRNA.PFemirge\_12197\_0.000012


metaRNA.PFemirge\_19580\_0.000003


metaRNA.PFemirge\_775\_0.000004


metaRNA.PFemirge\_2798\_0.000019


metaRNA.PFemirge\_6700\_0.000006


EU772528.1.1373 Bacteria;Firmicutes;Clostridia;Lachnosp ... aceae;Lachnospiraceae NC2004 group;uncultured bacterium


metaRNA.PFemirge\_27744\_0.000011


FN794258.1.1496 Bacteria;Proteobacteria;Gammaproteobacteria;Burkholderiales;Rhodocyclaceae;Azospira;uncultured bacterium


DQ793508.1.1385 Bacteria;Firmicutes;Clostridia;Oscillospirales;UCG-011;uncultured bacterium


metaRNA.PFemirge\_14207\_0.000051


CP012068.1576987.1578505 Bacteria;Firmicutes;Clostridia ... Eubacterium] nodatum group;Eubacterium sulci ATCC 35585


metaRNA.PFemirge\_14643\_0.000003


AB574480.1.1486 Bacteria;Bacteroidota;Bacteroidia;Bacteroidales;Bacteroidaceae;Bacteroides;Bacteroides faecichinchillae


metaRNA.PFemirge\_23106\_0.000008


metaRNA.PFemirge\_3956\_0.000000


DQ456246.1.1451 Bacteria;Bacteroidota;Bacteroidia;Bacteroidales;Bacteroidaceae;Bacteroides;uncultured bacterium


DQ824136.1.1393 Bacteria;Bacteroidota;Bacteroidia;Bacteroidales;Bacteroidaceae;Bacteroides;uncultured bacterium


metaRNA.PFemirge\_7720\_0.000296


metaRNA.PFemirge\_24547\_0.000006


metaRNA.PFemirge\_27765\_0.000112


metaRNA.PFemirge\_15075\_0.000001


metaRNA.PFemirge\_2154\_0.000043


FJ715990.1.1502 Bacteria;Proteobacteria;Gammaproteobact ... acterales;Aeromonadaceae;Aeromonas;uncultured bacterium


metaRNA.PFemirge\_9884\_0.000063


metaRNA.PFemirge\_4007\_0.000023


metaRNA.PFemirge\_2210\_0.001759


metaRNA.PFemirge\_27746\_0.000012


JN559712.1.1498 Bacteria;Bacteroidota;Bacteroidia;Bacteroidales;Bacteroidaceae;Bacteroides;uncultured bacterium


metaRNA.PFemirge\_10734\_0.000127


HM630228.1.1408 Bacteria;Firmicutes;Clostridia;Clostridia vadinBB60 group;uncultured bacterium


EU459250.1.1293 Bacteria;Firmicutes;Clostridia;Peptostr ... ostreptococcaceae;Paeniclostridium;uncultured bacterium


metaRNA.PFemirge\_16157\_0.000021


DQ808618.1.1395 Bacteria;Bacteroidota;Bacteroidia;Bacteroidales;Bacteroidaceae;Bacteroides;uncultured bacterium


metaRNA.PFemirge\_26285\_0.000002


metaRNA.PFemirge\_15784\_0.001731


GQ503874.1.1501 Bacteria;Desulfobacterota;Desulfovibrio ... ulfovibrio;bacterium enrichment culture clone Ecwsrb037


EU775230.1.1391 Bacteria;Firmicutes;Clostridia;Lachnosp ... aceae;Lachnospiraceae AC2044 group;uncultured bacterium


metaRNA.PFemirge\_27396\_0.000005


EU778633.1.1385 Bacteria;Firmicutes;Clostridia;Lachnospirales;Lachnospiraceae;Moryella;uncultured bacterium


CZBH01000001.999036.1000544 Bacteria;Bacteroidota;Bacteroidia;Bacteroidales;Rikenellaceae;Alistipes;Alistipes sp. cv1


metaRNA.PFemirge\_28310\_0.000170


metaRNA.PFemirge\_5051\_0.007470


metaRNA.PFemirge\_8071\_0.000339


metaRNA.PFemirge\_23679\_0.000081


EU778426.1.1385 Bacteria;Firmicutes;Clostridia;Clostrid ... iaceae;Clostridium sensu stricto 1;uncultured bacterium


HQ661204.1.1414 Bacteria;Proteobacteria;Alphaproteobacteria;Holosporales;Holosporaceae;uncultured;uncultured bacterium


GQ480147.1.1537 Bacteria;Proteobacteria;Alphaproteobacteria;Holosporales;Holosporaceae;uncultured;uncultured bacterium


metaRNA.PFemirge\_1575\_0.000184


metaRNA.PFemirge\_8044\_0.000160


JN680664.1.1439 Bacteria;Firmicutes;Clostridia;Peptostr ... erium] brachy group;uncultured Eubacteriaceae bacterium


HQ778371.1.1457 Bacteria;Firmicutes;Clostridia;Lachnospirales;Lachnospiraceae;Lachnoclostridium;uncultured organism


metaRNA.PFemirge\_20784\_0.000026


metaRNA.PFemirge\_6786\_0.000015


JQ608060.1.1417 Bacteria;Firmicutes;Clostridia;Oscillospirales;Butyricicoccaceae;UCG-009;bacterium NLAE-zl-C349


metaRNA.PFemirge\_6511\_0.000007


metaRNA.PFemirge\_19862\_0.000005


metaRNA.PFemirge\_20010\_0.017832


EF096958.1.1403 Bacteria;Bacteroidota;Bacteroidia;Bacteroidales;Bacteroidaceae;Bacteroides;uncultured bacterium


DQ794175.1.1406 Bacteria;Bacteroidota;Bacteroidia;Bacteroidales;Bacteroidaceae;Bacteroides;uncultured bacterium


AY212697.1.1526 Bacteria;Proteobacteria;Gammaproteobacteria;Burkholderiales;Rhodocyclaceae;Azospira;uncultured bacterium


metaRNA.PFemirge\_17760\_0.000272


metaRNA.PFemirge\_11747\_0.000053


metaRNA.PFemirge\_12017\_0.000004


metaRNA.PFemirge\_23794\_0.000008


metaRNA.PFemirge\_25869\_0.000016


metaRNA.PFemirge\_23365\_0.000011


AM183061.1.1440 Bacteria;Firmicutes;Clostridia;Clostrid ... iaceae;Clostridium sensu stricto 1;uncultured bacterium


metaRNA.PFemirge\_1445\_0.000005


metaRNA.PFemirge\_23154\_0.000006


metaRNA.PFemirge\_26295\_0.000002


metaRNA.PFemirge\_5990\_0.000119


metaRNA.PFemirge\_25491\_0.000005


metaRNA.PFemirge\_10947\_0.000006


AM936169.1.1362 Bacteria;Proteobacteria;Gammaproteobact ... eae;Immundisolibacter;uncultured Chromatiales bacterium


FCNU01000016.57.1576 Bacteria;Verrucomicrobiota;Chlamyd ... ydiaceae;Candidatus Protochlamydia;Parachlamydia sp. C2


metaRNA.PFemirge\_15643\_0.000014


metaRNA.PFemirge\_4309\_0.001028


MH031289.1.1458 Bacteria;Bacteroidota;Bacteroidia;Chiti ... spiraceae;uncultured;uncultured Bacteroidetes bacterium


metaRNA.PFemirge\_17427\_0.005542


EU776269.1.1279 Bacteria;Firmicutes;Clostridia;Oscillospirales;Hydrogenoanaerobacterium;uncultured bacterium


metaRNA.PFemirge\_12327\_0.000005


GQ503787.1.1505 Bacteria;Desulfobacterota;Desulfovibrio ... io;Desulfovibrio sp. enrichment culture clone Jdgsrb034


metaRNA.PFemirge\_25860\_0.000002


metaRNA.PFemirge\_26799\_0.000024


##### Interactive treemap of mapping-based taxonomic read classification (Click to hide)

**Navigation**: Left-click to go down, right-click to go up in taxonomic hierarchy, hover to see counts.

Based on read-mapping hits to reference database, provides an approximate overview of taxonomic composition.

Drawn with Google Visualization API (terms of service)

#### Input parameters

|  |  |
| --- | --- |
| Input command | phyloFlash.pl -lib metaRNA -read1 final\_pure\_reads\_1.fastq -read2 final\_pure\_reads\_2.fastq -CPUs 24 -everything -trusted Xt\_metatrin.fasta |
| Forward read file | final\_pure\_reads\_1.fastq |
| Reverse read file | final\_pure\_reads\_2.fastq |
| Minimum mapping identity | 70% |
| Working folder | /home/pollet/metatetard/metatranscriptomic/phyloflash\_output2 |
| Database used | /home/anaconda/138.1 |

#### Results

##### Mapping statistics

|  |  |
| --- | --- |
| Input PE-reads | 17489433 |
| Mapped SSU read pairs | 972323 |
| Mapping ratio | 5.559% |
| Fraction assembled | 96.966% |
| Detected median insert size | 159 |
| Used insert size | 220 |
| Insert size standard deviation | 27 |

##### Output files

|  |  |
| --- | --- |
| FASTA file of alignment of all full-length sequences | metaRNA.SSU.collection.alignment.fasta |
| FASTA file of all full-length sequences and their closest database hits | metaRNA.SSU.collection.fasta |
| Newick guide tree from MAFFT alignment of all full-length sequences and closest database hits | metaRNA.SSU.collection.fasta.tree |
| SVG graphic of guide tree from MAFFT alignment of all full-length sequences and closest database hits | metaRNA.SSU.collection.fasta.tree.svg |
| FASTA file of full-length SSU sequences | metaRNA.all.final.fasta |
| Raw BBmap SAM file of initial read mapping to SSU rRNA database | metaRNA.bbmap.sam |
| Log file from EMIRGE sequence reconstruction | metaRNA.emirge.out |
| Reads (fwd) mapping to SSU rRNA database | metaRNA.final\_pure\_reads\_1.fastq.SSU.1.fq |
| Reads (rev) mapping to SSU rRNA database | metaRNA.final\_pure\_reads\_1.fastq.SSU.2.fq |
| SAM file of initial read mapping to SSU rRNA database | metaRNA.final\_pure\_reads\_1.fastq.SSU.sam |
| Mapping identity histogram from BBmap | metaRNA.idhistogram |
| Insert size histogram from BBmap | metaRNA.inserthistogram |
| phyloFlash report in plain text | metaRNA.phyloFlash |
| NTU abundances (truncated to requested taxonomic level) from initial mapping, in CSV format | metaRNA.phyloFlash.NTUabundance.csv |
| SVG graphic of taxonomic composition from initial read mapping | metaRNA.phyloFlash.NTUabundance.csv.svg |
| NTU abundances (untruncated) from initial mapping, in CSV format | metaRNA.phyloFlash.NTUfull\_abundance.csv |
| Taxonomic classification of full-length sequences, in CSV format | metaRNA.phyloFlash.extractedSSUclassifications.csv |
| phyloFlash report in CSV format | metaRNA.phyloFlash.report.csv |
| Taxonomic composition of unassembled SSU reads in CSV format | metaRNA.phyloFlash.unassembled.NTUabundance.csv |
| Log file from SPAdes assembler | metaRNA.spades.out |
| Concatenated GFF file of SSU rRNA sequences for all three models in trusted contigs | metaRNA.trusted.all.gff |
| SAM file of read mapping vs trusted contigs | metaRNA.trusted.bbmap.sam |

##### Taxonomic affiliation of SSU rRNA reads in library

Approximate overview of taxonomic composition of ALL reads, based on mapping hits to SILVA SSU rRNA database using BBmap.

|  |  |
| --- | --- |
| NTUs observed once | 55 |
| NTUs observed twice | 29 |
| NTUs observed three or more times | 304 |
| NTU Chao1 richness estimate | 356.155 |

Taxonomy summarized at level 4. Only displaying taxa with > 3 reads mapped.

| Taxon | Reads |
| --- | --- |
| Bacteria;Bacteroidota;Bacteroidia;Bacteroidales | 3006774 |
| Bacteria;Desulfobacterota;Desulfovibrionia;Desulfovibrionales | 1552119 |
| Bacteria;Firmicutes;Negativicutes;Acidaminococcales | 876192 |
| Bacteria;Firmicutes;Clostridia;Lachnospirales | 292599 |
| Bacteria;Firmicutes;Clostridia;Oscillospirales | 262203 |
| Bacteria;Actinobacteriota;Actinobacteria;Micrococcales | 242424 |
| Bacteria;(Bacteria);(Bacteria);(Bacteria) | 188518 |
| Bacteria;Proteobacteria;Gammaproteobacteria;Enterobacterales | 182557 |
| Bacteria;Proteobacteria;Gammaproteobacteria;Burkholderiales | 63968 |
| Eukaryota;Amorphea;Obazoa;Opisthokonta | 60533 |
| Bacteria;Proteobacteria;Alphaproteobacteria;(Alphaproteobacteria) | 55977 |
| Bacteria;Proteobacteria;Alphaproteobacteria;Rhodospirillales | 54633 |
| Bacteria;Firmicutes;Bacilli;Erysipelotrichales | 54346 |
| Bacteria;Firmicutes;Clostridia;Peptostreptococcales-Tissierellales | 34015 |
| Bacteria;Firmicutes;(Firmicutes);(Firmicutes) | 30864 |
| Eukaryota;SAR;Rhizaria;Retaria | 27001 |
| Bacteria;Bacteroidota;Bacteroidia;(Bacteroidia) | 24582 |
| Bacteria;Proteobacteria;Alphaproteobacteria;Paracaedibacterales | 23219 |
| Bacteria;Verrucomicrobiota;Verrucomicrobiae;Verrucomicrobiales | 22665 |
| Bacteria;Proteobacteria;Alphaproteobacteria;Rhizobiales | 20235 |
| Bacteria;Cyanobacteria;Vampirivibrionia;Gastranaerophilales | 19394 |
| Bacteria;Firmicutes;Clostridia;(Clostridia) | 18290 |
| Bacteria;Deferribacterota;Deferribacteres;Deferribacterales | 17546 |
| Eukaryota;Amorphea;Amoebozoa;Discosea | 17460 |
| Bacteria;Proteobacteria;(Proteobacteria);(Proteobacteria) | 17446 |
| Bacteria;Firmicutes;Bacilli;(Bacilli) | 16159 |
| Bacteria;Proteobacteria;Alphaproteobacteria;Zavarziniales | 14809 |
| Bacteria;Firmicutes;Negativicutes;(Negativicutes) | 14786 |
| Eukaryota;SAR;Stramenopiles;Opalinata | 14562 |
| Eukaryota;SAR;Stramenopiles;(Stramenopiles) | 13717 |
| Eukaryota;SAR;(SAR);(SAR) | 13028 |
| Bacteria;Firmicutes;Negativicutes;Veillonellales-Selenomonadales | 10775 |
| Eukaryota;(Eukaryota);(Eukaryota);(Eukaryota) | 10686 |
| Bacteria;Firmicutes;Bacilli;Bacillales | 10027 |
| Eukaryota;Amorphea;Amoebozoa;Archamoebae | 9759 |
| Bacteria;Firmicutes;Bacilli;Paenibacillales | 9599 |
| Bacteria;Firmicutes;Clostridia;Clostridia UCG-014 | 9150 |
| Bacteria;Verrucomicrobiota;Verrucomicrobiae;(Verrucomicrobiae) | 8181 |
| Bacteria;Synergistota;Synergistia;Synergistales | 7400 |
| Bacteria;Firmicutes;Clostridia;Christensenellales | 7225 |
| Bacteria;Actinobacteriota;Actinobacteria;(Actinobacteria) | 7078 |
| Bacteria;Desulfobacterota;(Desulfobacterota);(Desulfobacterota) | 6867 |
| Bacteria;Proteobacteria;Alphaproteobacteria;Holosporales | 5540 |
| Bacteria;Bacteroidota;Bacteroidia;Flavobacteriales | 5389 |
| Bacteria;Proteobacteria;Alphaproteobacteria;Caedibacterales | 4977 |
| Bacteria;Firmicutes;Bacilli;RF39 | 4931 |
| Bacteria;Firmicutes;Bacilli;Lactobacillales | 4861 |
| Eukaryota;Amorphea;Amoebozoa;(Amoebozoa) | 4790 |
| Bacteria;Proteobacteria;Gammaproteobacteria;(Gammaproteobacteria) | 4007 |
| Bacteria;Firmicutes;Clostridia;Clostridiales | 3922 |
| Bacteria;Verrucomicrobiota;Lentisphaeria;Victivallales | 3770 |
| Bacteria;Cyanobacteria;Cyanobacteriia;Chloroplast | 3118 |
| Eukaryota;SAR;Alveolata;Ciliophora | 3049 |
| Bacteria;Verrucomicrobiota;Chlamydiae;Chlamydiales | 3042 |
| Bacteria;Proteobacteria;Alphaproteobacteria;Azospirillales | 3029 |
| Bacteria;Proteobacteria;Gammaproteobacteria;Pseudomonadales | 2537 |
| Bacteria;Bacteroidota;Bacteroidia;Cytophagales | 2383 |
| Bacteria;Bacteroidota;Bacteroidia;Sphingobacteriales | 2050 |
| Bacteria;Proteobacteria;Gammaproteobacteria;Xanthomonadales | 1991 |
| Bacteria;Bacteroidota;Bacteroidia;Chitinophagales | 1826 |
| Eukaryota;SAR;Stramenopiles;Incertae Sedis | 1820 |
| Bacteria;Acidobacteriota;Subgroup 22;uncultured bacterium | 1780 |
| Bacteria;Proteobacteria;Alphaproteobacteria;Dongiales | 1728 |
| Bacteria;Firmicutes;Clostridia;uncultured | 1549 |
| Eukaryota;Amorphea;Amoebozoa;uncultured eukaryote | 1475 |
| Bacteria;Proteobacteria;Alphaproteobacteria;uncultured | 1430 |
| Bacteria;Proteobacteria;Gammaproteobacteria;Legionellales | 1414 |
| Bacteria;Proteobacteria;Alphaproteobacteria;Rickettsiales | 1336 |
| Bacteria;Actinobacteriota;Actinobacteria;Corynebacteriales | 1292 |
| Bacteria;Cyanobacteria;Cyanobacteriia;Cyanobacteriales | 1250 |
| Bacteria;Proteobacteria;Alphaproteobacteria;Sphingomonadales | 1148 |
| Bacteria;Fusobacteriota;Fusobacteriia;Fusobacteriales | 967 |
| Bacteria;Firmicutes;Bacilli;RsaHf231 | 957 |
| Bacteria;Firmicutes;Bacilli;Staphylococcales | 918 |
| Eukaryota;Archaeplastida;Chloroplastida;Charophyta | 916 |
| Bacteria;Proteobacteria;Alphaproteobacteria;Reyranellales | 886 |
| Bacteria;Desulfobacterota;Desulfuromonadia;Geobacterales | 842 |
| Eukaryota;Amorphea;Amoebozoa;Incertae Sedis | 813 |
| Bacteria;Proteobacteria;Alphaproteobacteria;Elsterales | 786 |
| Bacteria;Proteobacteria;Gammaproteobacteria;Gammaproteobacteria Incertae Sedis | 733 |
| Bacteria;Firmicutes;Bacilli;Brevibacillales | 715 |
| Bacteria;Proteobacteria;Alphaproteobacteria;Parvibaculales | 680 |
| Bacteria;Proteobacteria;Alphaproteobacteria;AT-s3-44 | 559 |
| Bacteria;Firmicutes;Desulfotomaculia;Desulfotomaculales | 518 |
| Bacteria;Firmicutes;Bacilli;Izemoplasmatales | 500 |
| Bacteria;Proteobacteria;Alphaproteobacteria;Kiloniellales | 497 |
| Bacteria;Actinobacteriota;Actinobacteria;Propionibacteriales | 479 |
| Bacteria;Firmicutes;Clostridia;Eubacteriales | 474 |
| Bacteria;Proteobacteria;Alphaproteobacteria;Rhodobacterales | 450 |
| Bacteria;Proteobacteria;Gammaproteobacteria;Diplorickettsiales | 448 |
| Bacteria;Verrucomicrobiota;Kiritimatiellae;Kiritimatiellales | 433 |
| Bacteria;Firmicutes;Clostridia;Monoglobales | 426 |
| Eukaryota;Amorphea;Amoebozoa;SAR | 422 |
| Bacteria;Actinobacteriota;Thermoleophilia;Gaiellales | 378 |
| Bacteria;Actinobacteriota;Acidimicrobiia;IMCC26256 | 377 |
| Bacteria;Firmicutes;Bacilli;Haloplasmatales | 364 |
| Bacteria;Desulfobacterota;Desulfobulbia;Desulfobulbales | 361 |
| Bacteria;Verrucomicrobiota;Verrucomicrobiae;Chthoniobacterales | 357 |
| Bacteria;Actinobacteriota;Coriobacteriia;Coriobacteriales | 356 |
| Bacteria;Firmicutes;Clostridia;Clostridia vadinBB60 group | 352 |
| Bacteria;Calditrichota;Calditrichia;Calditrichales | 348 |
| Bacteria;MBNT15;Actinobacteria bacterium RBG\_13\_63\_9;(Actinobacteria bacterium RBG\_13\_63\_9) | 341 |
| Eukaryota;Discoba;Discicristata;Euglenozoa | 334 |
| Eukaryota;Amorphea;(Amorphea);(Amorphea) | 333 |
| Bacteria;Firmicutes;Clostridia;Peptococcales | 318 |
| Eukaryota;SAR;Stramenopiles;MAST-3 | 315 |
| Bacteria;Cyanobacteria;Vampirivibrionia;Obscuribacterales | 297 |
| Bacteria;Proteobacteria;Alphaproteobacteria;Puniceispirillales | 281 |
| Bacteria;Firmicutes;Moorellia;Desulfitibacterales | 281 |
| Eukaryota;SAR;Stramenopiles;Labyrinthulomycetes | 273 |
| Bacteria;Proteobacteria;Gammaproteobacteria;Methylococcales | 272 |
| Bacteria;Proteobacteria;Gammaproteobacteria;Acidithiobacillales | 257 |
| Bacteria;Firmicutes;Desulfitobacteriia;Desulfitobacteriales | 251 |
| Bacteria;Actinobacteriota;Actinobacteria;Pseudonocardiales | 248 |
| Eukaryota;SAR;Rhizaria;Cercozoa | 232 |
| Eukaryota;SAR;Stramenopiles;Ochrophyta | 225 |
| Bacteria;Proteobacteria;Gammaproteobacteria;uncultured | 220 |
| Bacteria;Proteobacteria;Alphaproteobacteria;Thalassobaculales | 211 |
| Bacteria;Proteobacteria;Gammaproteobacteria;Ectothiorhodospirales | 206 |
| Bacteria;Actinobacteriota;Actinobacteria;Actinomycetales | 201 |
| Bacteria;Firmicutes;Bacilli;Thermoactinomycetales | 188 |
| Bacteria;Acidobacteriota;Holophagae;Subgroup 7 | 183 |
| Bacteria;Acidobacteriota;Vicinamibacteria;Vicinamibacterales | 176 |
| Bacteria;Sva0485;uncultured bacterium;(uncultured bacterium) | 170 |
| Eukaryota;SAR;Stramenopiles;Peronosporomycetes | 170 |
| Bacteria;Firmicutes;Dethiobacteria;Dethiobacterales | 157 |
| Bacteria;Bacteroidota;(Bacteroidota);(Bacteroidota) | 154 |
| Bacteria;Firmicutes;Thermoanaerobacteria;Thermoanaerobacterales | 137 |
| Bacteria;Verrucomicrobiota;Verrucomicrobiae;Pedosphaerales | 136 |
| Bacteria;Firmicutes;Clostridia;Gracilibacteraceae | 136 |
| Bacteria;Halanaerobiaeota;Halanaerobiia;Halanaerobiales | 135 |
| Eukaryota;Archaeplastida;Chloroplastida;Chlorophyta | 133 |
| Bacteria;Spirochaetota;MVP-15;uncultured Firmicutes bacterium | 133 |
| Bacteria;Verrucomicrobiota;Omnitrophia;Omnitrophales | 116 |
| Bacteria;Actinobacteriota;Actinobacteria;Streptosporangiales | 113 |
| Bacteria;Proteobacteria;Alphaproteobacteria;Defluviicoccales | 100 |
| Bacteria;Actinobacteriota;Actinobacteria;PeM15 | 100 |
| Bacteria;Myxococcota;Myxococcia;Myxococcales | 98 |
| Bacteria;Firmicutes;Desulfotomaculia;(Desulfotomaculia) | 93 |
| Bacteria;Verrucomicrobiota;Verrucomicrobiae;Arctic97B-4 marine group | 91 |
| Bacteria;Proteobacteria;Alphaproteobacteria;Kordiimonadales | 91 |
| Bacteria;Planctomycetota;Planctomycetes;Pirellulales | 90 |
| Bacteria;Verrucomicrobiota;(Verrucomicrobiota);(Verrucomicrobiota) | 89 |
| Bacteria;Actinobacteriota;(Actinobacteriota);(Actinobacteriota) | 77 |
| Bacteria;Firmicutes;Limnochordia;Limnochordales | 76 |
| Bacteria;Spirochaetota;Brachyspirae;Brachyspirales | 76 |
| Bacteria;Nitrospirota;Thermodesulfovibrionia;uncultured | 74 |
| Bacteria;Cyanobacteria;Cyanobacteriia;(Cyanobacteriia) | 74 |
| Bacteria;Verrucomicrobiota;Verrucomicrobiae;Opitutales | 73 |
| Bacteria;Firmicutes;Incertae Sedis;Gelria | 72 |
| Bacteria;Desulfobacterota;Desulfobacteria;Desulfobacterales | 71 |
| Bacteria;Verrucomicrobiota;Lentisphaeria;(Lentisphaeria) | 67 |
| Eukaryota;SAR;Alveolata;Protalveolata | 65 |
| Bacteria;Desulfobacterota;Syntrophia;Syntrophales | 65 |
| Bacteria;Spirochaetota;Spirochaetia;Spirochaetales | 62 |
| Bacteria;Myxococcota;Polyangia;mle1-27 | 61 |
| Bacteria;Chloroflexi;Chloroflexia;Chloroflexales | 60 |
| Bacteria;Proteobacteria;Alphaproteobacteria;Ferrovibrionales | 60 |
| Bacteria;Myxococcota;Polyangia;Polyangiales | 55 |
| Bacteria;Acidobacteriota;Acidobacteriae;Subgroup 13 | 54 |
| Bacteria;Actinobacteriota;Actinobacteria;Streptomycetales | 53 |
| Bacteria;Firmicutes;Bacilli;Exiguobacterales | 49 |
| Bacteria;Caldisericota;Caldisericia;Caldisericales | 48 |
| Bacteria;Proteobacteria;Alphaproteobacteria;Caulobacterales | 47 |
| Bacteria;Planctomycetota;Planctomycetes;Planctomycetales | 47 |
| Bacteria;Methylomirabilota;Methylomirabilia;Methylomirabilales | 44 |
| Bacteria;Planctomycetota;Phycisphaerae;S-70 | 42 |
| Eukaryota;SAR;Alveolata;(Alveolata) | 42 |
| Bacteria;Acidobacteriota;(Acidobacteriota);(Acidobacteriota) | 40 |
| Bacteria;Desulfobacterota;Desulfuromonadia;Desulfuromonadales | 40 |
| Bacteria;Planctomycetota;Phycisphaerae;Pla1 lineage | 39 |
| Bacteria;Deinococcota;Deinococci;Deinococcales | 38 |
| Bacteria;Cloacimonadota;Cloacimonadia;Cloacimonadales | 38 |
| Bacteria;Planctomycetota;Planctomycetes;Gemmatales | 37 |
| Bacteria;Myxococcota;Polyangia;Haliangiales | 37 |
| Bacteria;Firmicutes;Syntrophomonadia;Syntrophomonadales | 36 |
| Bacteria;Desulfobacterota;Desulfuromonadia;(Desulfuromonadia) | 32 |
| Bacteria;Acidobacteriota;Holophagae;(Holophagae) | 32 |
| Bacteria;Proteobacteria;Gammaproteobacteria;Coxiellales | 31 |
| Bacteria;Dadabacteria;Dadabacteriia;Dadabacteriales | 31 |
| Bacteria;Elusimicrobiota;Elusimicrobia;Elusimicrobiales | 30 |
| Bacteria;Gemmatimonadota;Gemmatimonadetes;Gemmatimonadales | 30 |
| Bacteria;Desulfobacterota;Desulfuromonadia;PB19 | 28 |
| Bacteria;Firmicutes;Bacilli;Alicyclobacillales | 28 |
| Bacteria;Patescibacteria;Saccharimonadia;Saccharimonadales | 26 |
| Bacteria;Campylobacterota;Campylobacteria;Campylobacterales | 26 |
| Eukaryota;Amorphea;Amoebozoa;Tubulinea | 26 |
| Bacteria;Bdellovibrionota;Bdellovibrionia;Bdellovibrionales | 25 |
| Bacteria;Proteobacteria;Gammaproteobacteria;JTB23 | 25 |
| Eukaryota;Discoba;Discicristata;Heterolobosea | 24 |
| Bacteria;Firmicutes;Clostridia;Caldicoprobacterales | 24 |
| Bacteria;Bdellovibrionota;Oligoflexia;0319-6G20 | 24 |
| Bacteria;Firmicutes;uncultured;uncultured bacterium | 22 |
| Bacteria;Actinobacteriota;Acidimicrobiia;Microtrichales | 21 |
| Bacteria;Proteobacteria;Alphaproteobacteria;Sneathiellales | 21 |
| Bacteria;Verrucomicrobiota;Lentisphaeria;P.palmC41 | 21 |
| Eukaryota;Amorphea;Amoebozoa;metagenome | 21 |
| Bacteria;Dependentiae;Babeliae;Babeliales | 20 |
| Bacteria;Patescibacteria;Microgenomatia;uncultured sediment bacterium | 18 |
| Eukaryota;Archaeplastida;Chloroplastida;(Chloroplastida) | 18 |
| Bacteria;Planctomycetota;vadinHA49;uncultured bacterium | 17 |
| Eukaryota;uncultured Entamoeba;(uncultured Entamoeba);(uncultured Entamoeba) | 16 |
| Eukaryota;Archaeplastida;Rhodophyceae;Bangiales | 16 |
| Bacteria;Proteobacteria;Alphaproteobacteria;Acetobacterales | 15 |
| Bacteria;Actinobacteriota;Thermoleophilia;Solirubrobacterales | 15 |
| Bacteria;Myxococcota;Polyangia;(Polyangia) | 14 |
| Bacteria;Actinobacteriota;Actinobacteria;Frankiales | 14 |
| Bacteria;MBNT15;Nitrospirae bacterium CG2\_30\_70\_394;(Nitrospirae bacterium CG2\_30\_70\_394) | 14 |
| Bacteria;Cyanobacteria;Vampirivibrionia;(Vampirivibrionia) | 13 |
| Bacteria;Planctomycetota;Planctomycetes;Isosphaerales | 13 |
| Bacteria;Bacteroidota;Kapabacteria;Kapabacteriales | 13 |
| Bacteria;Firmicutes;Bacilli;Aneurinibacillales | 12 |
| Bacteria;Caldatribacteriota;JS1;uncultured bacterium | 12 |
| Bacteria;Verrucomicrobiota;Kiritimatiellae;WCHB1-41 | 12 |
| Bacteria;RCP2-54;metagenome;(metagenome) | 12 |
| Bacteria;Proteobacteria;Gammaproteobacteria;Thiotrichales | 11 |
| Bacteria;Proteobacteria;Gammaproteobacteria;Steroidobacterales | 11 |
| Eukaryota;SAR;Alveolata;Apicomplexa | 11 |
| Bacteria;NB1-j;uncultured bacterium;(uncultured bacterium) | 11 |
| Bacteria;Desulfobacterota;Desulfobacteria;Desulfatiglandales | 11 |
| Bacteria;Firmicutes;Desulfotomaculia;Ammonifexales | 11 |
| Bacteria;Bacteroidota;Bacteroidia;Bacteroidetes VC2.1 Bac22 | 11 |
| Bacteria;Planctomycetota;Phycisphaerae;Phycisphaerales | 10 |
| Bacteria;Bacteroidota;Ignavibacteria;Ignavibacteriales | 10 |
| Bacteria;Nitrospirota;4-29-1;uncultured bacterium | 10 |
| Eukaryota;SAR;Alveolata;uncultured | 9 |
| Bacteria;Patescibacteria;Gracilibacteria;metagenome | 9 |
| Bacteria;Firmicutes;D8A-2;uncultured bacterium | 9 |
| Bacteria;Proteobacteria;Gammaproteobacteria;HOC36 | 9 |
| Bacteria;Marinimicrobia (SAR406 clade);uncultured bacterium;(uncultured bacterium) | 9 |
| Bacteria;Firmicutes;Sulfobacillia;Sulfobacillales | 9 |
| Bacteria;Hydrogenedentes;Hydrogenedentia;Hydrogenedentiales | 9 |
| Bacteria;MBNT15;uncultured delta proteobacterium;(uncultured delta proteobacterium) | 8 |
| Eukaryota;Amorphea;Amoebozoa;Schizoplasmodiida | 8 |
| Bacteria;Patescibacteria;Gracilibacteria;uncultured bacterium | 8 |
| Bacteria;Desulfobacterota;uncultured;uncultured bacterium | 8 |
| Bacteria;Sva0485;(Sva0485);(Sva0485) | 7 |
| Bacteria;Fermentibacterota;Fermentibacteria;Fermentibacterales | 7 |
| Bacteria;Proteobacteria;Gammaproteobacteria;Cardiobacteriales | 7 |
| Bacteria;Proteobacteria;Gammaproteobacteria;Piscirickettsiales | 7 |
| Bacteria;Patescibacteria;Gracilibacteria;(Gracilibacteria) | 7 |
| Bacteria;Actinobacteriota;Actinobacteria;Glycomycetales | 7 |
| Bacteria;Verrucomicrobiota;Verrucomicrobiae;Methylacidiphilales | 7 |
| Bacteria;Firmicutes;Limnochordia;MBA03 | 7 |
| Bacteria;Proteobacteria;Gammaproteobacteria;Granulosicoccales | 7 |
| Eukaryota;Cryptophyceae;Cryptomonadales;Cryptomonas | 6 |
| Eukaryota;SAR;Rhizaria;(Rhizaria) | 6 |
| Bacteria;Patescibacteria;Microgenomatia;Candidatus Roizmanbacteria | 6 |
| Eukaryota;Excavata;Metamonada;Preaxostyla | 6 |
| Bacteria;Firmicutes;BRH-c20a;uncultured bacterium | 6 |
| Bacteria;Acidobacteriota;Blastocatellia;Blastocatellales | 6 |
| Bacteria;Firmicutes;Clostridia;Proteinivoracales | 6 |
| Bacteria;Bdellovibrionota;Oligoflexia;Oligoflexales | 6 |
| Bacteria;Myxococcota;bacteriap25;uncultured bacterium | 6 |
| Bacteria;Firmicutes;Symbiobacteriia;Symbiobacteriales | 6 |
| Bacteria;Proteobacteria;Magnetococcia;Magnetococcales | 6 |
| Eukaryota;SAR;Stramenopiles;MAST-12 | 6 |
| Bacteria;Planctomycetota;Phycisphaerae;MSBL9 | 5 |
| Bacteria;Nitrospirota;Nitrospiria;Nitrospirales | 5 |
| Bacteria;bacterium GBS-1;(bacterium GBS-1);(bacterium GBS-1) | 5 |
| Bacteria;Desulfobacterota;uncultured;uncultured delta proteobacterium | 5 |
| Bacteria;Firmicutes;Bacilli;Acholeplasmatales | 5 |
| Bacteria;Chloroflexi;Anaerolineae;SBR1031 | 5 |
| Bacteria;Proteobacteria;Alphaproteobacteria;Micavibrionales | 5 |
| Bacteria;Acidobacteriota;Thermoanaerobaculia;Thermoanaerobaculales | 5 |
| Bacteria;RCP2-54;uncultured bacterium;(uncultured bacterium) | 5 |
| Eukaryota;Haptophyta;T58;uncultured marine haptophyte | 4 |
| Bacteria;Latescibacterota;Latescibacteria;Latescibacterales | 4 |
| Bacteria;Myxococcota;Polyangia;Blfdi19 | 4 |
| Eukaryota;Amorphea;Amoebozoa;Mycamoeba | 4 |
| Eukaryota;Amorphea;Amoebozoa;Gracilipodida | 4 |
| Bacteria;Bdellovibrionota;Oligoflexia;uncultured | 4 |
| Bacteria;Firmicutes;Negativicutes;uncultured | 4 |
| Bacteria;TA06;uncultured bacterium;(uncultured bacterium) | 4 |
| Bacteria;Nitrospirota;Thermodesulfovibrionia;Magnetobacteriales | 4 |
| Bacteria;Abditibacteriota;Abditibacteria;Abditibacteriales | 4 |
| Bacteria;Elusimicrobiota;Elusimicrobia;(Elusimicrobia) | 4 |
| Bacteria;Schekmanbacteria;uncultured bacterium;(uncultured bacterium) | 4 |
| Bacteria;Proteobacteria;Gammaproteobacteria;Nitrosococcales | 4 |

##### SSU rRNA assembly-based taxa

Full-length SSU rRNA sequences assembled by SPAdes, matched to SILVA database with Vsearch.

| OTU | Mapped | Cov | DB hit | Taxonomy | % ID | Alnlen | Evalue |
| --- | --- | --- | --- | --- | --- | --- | --- |
| metaRNA.PFspades\_1 | 1789171 | 6007.471877 | GQ480160.1.1510 | Bacteria;Firmicutes;Negativicutes;Acidaminococcales;Acidaminococcaceae;Succinispira;uncultured bacterium | 94.7 | 1509 | -1 |
| metaRNA.PFspades\_2 | 489412 | 1753.138489 | LC094617.1.1418 | Bacteria;Actinobacteriota;Actinobacteria;Micrococcales;Microbacteriaceae;Leifsonia;Microbacteriaceae bacterium MAEY21 | 99.3 | 1418 | -1 |
| metaRNA.PFspades\_17 | 448044 | 1555.440517 | DQ393283.1.1453 | Bacteria;Cyanobacteria;Cyanobacteriia;Cyanobacteriales;Phormidiaceae;Arthrospira PCC-7345;Arthrospira platensis SAG 21.99 | 93.2 | 1253 | -1 |
| metaRNA.PFspades\_4 | 422201 | 1638.275299 | AGTN01029961.1048.2511 | Bacteria;Proteobacteria;Alphaproteobacteria;Rhodospirillales;Magnetospirillaceae;Magnetospirillum;bioreactor metagenome | 93.4 | 1466 | -1 |
| metaRNA.PFspades\_42 | 279194 | 1972.710648 | JN653056.1.1473 | Bacteria;Desulfobacterota;Desulfovibrionia;Desulfovibrionales;Desulfovibrionaceae;Desulfovibrio;uncultured Desulfovibrionaceae bacterium | 96.6 | 939 | -1 |
| metaRNA.PFspades\_7 | 90773 | 377.945982 | AF157057.1.1502 | Bacteria;Deferribacterota;Deferribacteres;Deferribacterales;Deferribacteraceae;Mucispirillum;bacterium 'Lincoln Park 3' | 91.9 | 1481 | -1 |
| metaRNA.PFspades\_18 | 74577 | 408.152000 | APKA01105891.43.1849 | Eukaryota;Amorphea;Obazoa;Opisthokonta;Holozoa;Choanozoa;Metazoa;Animalia;BCP clade;Bilateria;Protostomia;Lophotrochozoa;Rotifera;Bdelloidea;Adinetida;Biomphalaria glabrata | 100.0 | 1203 | -1 |
| metaRNA.PFspades\_9 | 74561 | 311.027739 | CP022983.633553.635105 | Bacteria;Firmicutes;Bacilli;Bacillales;Bacillaceae;Bacillus;Bacillus kochii | 87.6 | 1501 | -1 |
| metaRNA.PFspades\_3 | 67067 | 246.403614 | KF318250.1.1396 | Bacteria;Verrucomicrobiota;Verrucomicrobiae;Verrucomicrobiales;Akkermansiaceae;Akkermansia;uncultured Verrucomicrobia bacterium | 91.8 | 1398 | -1 |
| metaRNA.PFspades\_30 | 46129 | 283.402893 | GU585668.1.1471 | Bacteria;Firmicutes;Bacilli;Erysipelotrichales;Erysipelotrichaceae;[Anaerorhabdus] furcosa group;Anaerorhabdus furcosa | 98.4 | 994 | -1 |
| metaRNA.PFspades\_16 | 43481 | 159.465154 | EF400740.1.1495 | Bacteria;Firmicutes;Clostridia;Oscillospirales;Butyricicoccaceae;UCG-009;uncultured bacterium | 93.4 | 1308 | -1 |
| metaRNA.PFspades\_33 | 36070 | 227.152688 | HG970990.1.1557 | Bacteria;Firmicutes;Bacilli;Erysipelotrichales;Erysipelotrichaceae;Breznakia;uncultured bacterium | 91.1 | 975 | -1 |
| metaRNA.PFspades\_35 | 32893 | 119.998907 | AB649276.1.1495 | Bacteria;Firmicutes;Clostridia;Lachnospirales;Lachnospiraceae;Anaerostignum;Anaerotignum propionicum | 97.2 | 1002 | -1 |
| metaRNA.PFspades\_24 | 30259 | 180.812745 | GU293195.1.1464 | Bacteria;Firmicutes;Clostridia;Peptostreptococcales-Tissierellales;Peptostreptococcaceae;Romboutsia;uncultured bacterium | 99.4 | 1100 | -1 |
| metaRNA.PFspades\_23 | 28369 | 155.328872 | DQ817541.1.1479 | Bacteria;Proteobacteria;Gammaproteobacteria;Burkholderiales;Chromobacteriaceae;Crenobacter;uncultured bacterium | 99.6 | 1135 | -1 |
| metaRNA.PFspades\_8 | 26367 | 105.065262 | JX457309.1.1485 | Bacteria;Bacteroidota;Bacteroidia;Bacteroidales;Rs-E47 termite group;uncultured bacterium | 92.9 | 1465 | -1 |
| metaRNA.PFspades\_41 | 24848 | 161.204363 | DQ808998.1.1391 | Bacteria;Firmicutes;Clostridia;Oscillospirales;Butyricicoccaceae;UCG-009;uncultured bacterium | 94.9 | 954 | -1 |
| metaRNA.PFspades\_6 | 20949 | 72.965356 | CBEW010000180.8693.10203 | Bacteria;Cyanobacteria;Vampirivibrionia;Gastranaerophilales;Clostridium sp. CAG:306 | 96.8 | 1471 | -1 |
| metaRNA.PFspades\_5 | 20651 | 75.033820 | EF454925.1.1440 | Bacteria;Synergistota;Synergistia;Synergistales;Synergistaceae;uncultured;uncultured Synergistetes bacterium | 92.9 | 1431 | -1 |
| metaRNA.PFspades\_38 | 18731 | 79.249441 | KM098109.1.1519 | Bacteria;Firmicutes;Clostridia;Oscillospirales;Ruminococcaceae;UBA1819;Ruthenibacterium lactatiformans | 93.6 | 981 | -1 |
| metaRNA.PFspades\_25 | 18161 | 97.307312 | JX576025.1.1509 | Bacteria;Proteobacteria;Gammaproteobacteria;Burkholderiales;Rhodocyclaceae;Azospira;uncultured Azospira sp. | 95.3 | 986 | -1 |
| metaRNA.PFspades\_14 | 16372 | 56.928036 | EU158190.1.1510 | Bacteria;Firmicutes;Clostridia;Oscillospirales;Hydrogenoanaerobacterium;Hydrogenoanaerobacterium saccharovorans | 92.8 | 1411 | -1 |
| metaRNA.PFspades\_20 | 11651 | 61.734259 | KM650567.1.1481 | Bacteria;Firmicutes;Negativicutes;Acidaminococcales;Acidaminococcaceae;uncultured;uncultured Firmicutes bacterium | 94.6 | 1167 | -1 |
| metaRNA.PFspades\_22 | 11211 | 54.022814 | KR822473.1.1454 | Bacteria;Firmicutes;Clostridia;Peptostreptococcales-Tissierellales;Anaerovoracaceae;[Eubacterium] nodatum group;bacterium PM5-7 | 98.9 | 1144 | -1 |
| metaRNA.PFspades\_15 | 10418 | 49.033463 | DQ800642.1.1301 | Bacteria;Firmicutes;Clostridia;Christensenellales;Christensenellaceae;Christensenellaceae R-7 group;uncultured bacterium | 95.5 | 1292 | -1 |
| metaRNA.PFspades\_10 | 9432 | 39.599860 | BBDT01000057.3432.4945 | Bacteria;Firmicutes;Bacilli;Erysipelotrichales;Erysipelatoclostridiaceae;Coprobacillus;Coprobacillus cateniformis JCM 10604 | 95.7 | 1461 | -1 |
| metaRNA.PFspades\_11 | 7163 | 30.944644 | AB614872.1.1469 | Bacteria;Verrucomicrobiota;Lentisphaeria;Victivallales;Victivallaceae;uncultured rumen bacterium | 94.6 | 1376 | -1 |
| metaRNA.PFspades\_13 | 2643 | 11.888227 | AY571491.1.1466 | Bacteria;Proteobacteria;Alphaproteobacteria;Rickettsiales;uncultured;uncultured bacterium | 96.5 | 1358 | -1 |
| metaRNA.PFspades\_21 | 2414 | 13.924386 | X97806.1.1771 | Eukaryota;Amorphea;Obazoa;Opisthokonta;Nucletmycea;Fungi;Dikarya;Ascomycota;Saccharomycotina;Saccharomycetes;Saccharomycetales;Saccharomycetaceae;Saccharomyces;Saccharomyces paradoxus | 100.0 | 1151 | -1 |
| metaRNA.PFspades\_19 | 2224 | 11.010999 | AY457069.1.1483 | Bacteria;Actinobacteriota;Actinobacteria;Corynebacteriales;Mycobacteriaceae;Mycobacterium;Mycobacterium peregrinum | 99.9 | 1183 | -1 |
| metaRNA.PFspades\_12 | 1905 | 8.205331 | KC253315.1.1470 | Bacteria;Bacteroidota;Bacteroidia;Cytophagales;Spirosomaceae;Runella;uncultured bacterium | 98.2 | 1407 | -1 |
| metaRNA.PFspades\_27 | 607 | 3.570273 | AB595134.1.1484 | Bacteria;Firmicutes;Clostridia;Eubacteriales;Eubacteriaceae;Eubacterium;Eubacterium limosum | 97.0 | 1082 | -1 |


##### SSU rRNA reconstruction-based taxa

Full-length SSU rRNA seqeunces reconstructed by EMIRGE, matched to SILVA database by Vsearch.

| OTU | Mapped | Ratio | DB hit | Taxonomy | % ID | Alnlen | Evalue |
| --- | --- | --- | --- | --- | --- | --- | --- |
| metaRNA.PFemirge\_2917 | 1785595 | 0.057600 | GQ480160.1.1510 | Bacteria;Firmicutes;Negativicutes;Acidaminococcales;Acidaminococcaceae;Succinispira;uncultured bacterium | 94.7 | 1502 | -1 |
| metaRNA.PFemirge\_6635 | 1152939 | 0.084218 | AB574482.1.1489 | Bacteria;Bacteroidota;Bacteroidia;Bacteroidales;Tannerellaceae;Parabacteroides;Parabacteroides chinchillae | 96.2 | 1480 | -1 |
| metaRNA.PFemirge\_3509 | 987716 | 0.047992 | GQ503787.1.1505 | Bacteria;Desulfobacterota;Desulfovibrionia;Desulfovibrionales;Desulfovibrionaceae;Desulfovibrio;Desulfovibrio sp. enrichment culture clone Jdgsrb034 | 95.6 | 1456 | -1 |
| metaRNA.PFemirge\_12071 | 726246 | 0.024427 | HQ764819.1.1444 | Bacteria;Bacteroidota;Bacteroidia;Bacteroidales;Bacteroidaceae;Bacteroides;uncultured organism | 94.2 | 1379 | -1 |
| metaRNA.PFemirge\_28311 | 554970 | 0.021522 | KP231747.1.1515 | Bacteria;Desulfobacterota;Desulfovibrionia;Desulfovibrionales;Desulfovibrionaceae;Desulfovibrio;uncultured Desulfovibrio sp. | 95.6 | 1446 | -1 |
| metaRNA.PFemirge\_14997 | 461834 | 0.030526 | LC094617.1.1418 | Bacteria;Actinobacteriota;Actinobacteria;Micrococcales;Microbacteriaceae;Leifsonia;Microbacteriaceae bacterium MAEY21 | 99.2 | 1414 | -1 |
| metaRNA.PFemirge\_25631 | 393909 | 0.023054 | KP231747.1.1515 | Bacteria;Desulfobacterota;Desulfovibrionia;Desulfovibrionales;Desulfovibrionaceae;Desulfovibrio;uncultured Desulfovibrio sp. | 91.0 | 1326 | -1 |
| metaRNA.PFemirge\_7295 | 389027 | 0.019734 | CP022754.4526807.4528346 | Bacteria;Bacteroidota;Bacteroidia;Bacteroidales;Tannerellaceae;Parabacteroides;Parabacteroides sp. CT06 | 94.5 | 1486 | -1 |
| metaRNA.PFemirge\_20010 | 349358 | 0.017832 | EU775041.1.1395 | Bacteria;Bacteroidota;Bacteroidia;Bacteroidales;Bacteroidaceae;Bacteroides;uncultured bacterium | 92.5 | 1391 | -1 |
| metaRNA.PFemirge\_15962 | 329726 | 0.019534 | AGTN01029961.1048.2511 | Bacteria;Proteobacteria;Alphaproteobacteria;Rhodospirillales;Magnetospirillaceae;Magnetospirillum;bioreactor metagenome | 93.0 | 1401 | -1 |
| metaRNA.PFemirge\_6699 | 303707 | 0.011454 | HM630236.1.1451 | Bacteria;Bacteroidota;Bacteroidia;Bacteroidales;Rikenellaceae;dgA-11 gut group;uncultured bacterium | 89.9 | 1303 | -1 |
| metaRNA.PFemirge\_9950 | 301942 | 0.013428 | DQ355176.1.1464 | Bacteria;Desulfobacterota;Desulfovibrionia;Desulfovibrionales;Desulfovibrionaceae;Desulfovibrio;uncultured Desulfovibrio sp. | 93.8 | 1452 | -1 |
| metaRNA.PFemirge\_4517 | 236687 | 0.016085 | CBVB010000006.254456.255954 | Bacteria;Bacteroidota;Bacteroidia;Bacteroidales;Bacteroidaceae;Bacteroides;Bacteroidaceae bacterium MS4 | 96.7 | 1495 | -1 |
| metaRNA.PFemirge\_7089 | 177423 | 0.010108 | LN870298.1.1495 | Bacteria;Firmicutes;Clostridia;Oscillospirales;Oscillospiraceae;Intestinimonas;Intestinimonas timonensis | 94.1 | 1487 | -1 |
| metaRNA.PFemirge\_2499 | 177292 | 0.043392 | KM360180.1.1492 | Bacteria;Firmicutes;Clostridia;Lachnospirales;Lachnospiraceae;Lachnoclostridium;Clostridium sp. BPY5 | 93.0 | 1491 | -1 |
| metaRNA.PFemirge\_27691 | 173119 | 0.017422 | EF454852.1.1475 | Bacteria;Desulfobacterota;Desulfovibrionia;Desulfovibrionales;Desulfovibrionaceae;uncultured;uncultured Desulfovibrionaceae bacterium | 90.8 | 1468 | -1 |
| metaRNA.PFemirge\_4725 | 169824 | 0.027537 | AQHW01000007.13.1527 | Bacteria;Bacteroidota;Bacteroidia;Bacteroidales;Tannerellaceae;Parabacteroides;Parabacteroides gordonii MS-1 | 90.2 | 1450 | -1 |
| metaRNA.PFemirge\_16223 | 136635 | 0.004950 | EU509593.1.1402 | Bacteria;Desulfobacterota;Desulfovibrionia;Desulfovibrionales;Desulfovibrionaceae;uncultured;uncultured bacterium | 90.4 | 1400 | -1 |
| metaRNA.PFemirge\_16294 | 133978 | 0.010316 | DQ797781.1.1401 | Bacteria;Bacteroidota;Bacteroidia;Bacteroidales;Tannerellaceae;Parabacteroides;uncultured bacterium | 92.1 | 1386 | -1 |
| metaRNA.PFemirge\_12918 | 119239 | 0.003212 | DQ115999.1.1504 | Bacteria;Firmicutes;Clostridia;Lachnospirales;Lachnospiraceae;uncultured;uncultured bacterium | 90.7 | 1385 | -1 |
| metaRNA.PFemirge\_2531 | 107658 | 0.005487 | LN833517.1.1512 | Bacteria;Proteobacteria;Gammaproteobacteria;Enterobacterales;Enterobacteriaceae;Escherichia-Shigella;uncultured Escherichia sp. | 92.3 | 1347 | -1 |
| metaRNA.PFemirge\_10226 | 106209 | 0.014067 | KP231747.1.1515 | Bacteria;Desulfobacterota;Desulfovibrionia;Desulfovibrionales;Desulfovibrionaceae;Desulfovibrio;uncultured Desulfovibrio sp. | 92.6 | 1437 | -1 |
| metaRNA.PFemirge\_1887 | 95153 | 0.011605 | JN653056.1.1473 | Bacteria;Desulfobacterota;Desulfovibrionia;Desulfovibrionales;Desulfovibrionaceae;Desulfovibrio;uncultured Desulfovibrionaceae bacterium | 91.5 | 1476 | -1 |
| metaRNA.PFemirge\_17427 | 94069 | 0.005542 | DQ808618.1.1395 | Bacteria;Bacteroidota;Bacteroidia;Bacteroidales;Bacteroidaceae;Bacteroides;uncultured bacterium | 95.8 | 1395 | -1 |
| metaRNA.PFemirge\_15251 | 93727 | 0.005416 | HM630238.1.1413 | Bacteria;Bacteroidota;Bacteroidia;Bacteroidales;Rikenellaceae;Rikenella;uncultured bacterium | 92.0 | 1396 | -1 |
| metaRNA.PFemirge\_109 | 93120 | 0.004985 | APKA01105891.43.1849 | Eukaryota;Amorphea;Obazoa;Opisthokonta;Holozoa;Choanozoa;Metazoa;Animalia;BCP clade;Bilateria;Protostomia;Lophotrochozoa;Rotifera;Bdelloidea;Adinetida;Biomphalaria glabrata | 99.9 | 1772 | -1 |
| metaRNA.PFemirge\_3306 | 92765 | 0.002933 | HM124083.1.1490 | Bacteria;Bacteroidota;Bacteroidia;Bacteroidales;Prevotellaceae;Prevotellaceae Ga6A1 group;uncultured bacterium | 89.7 | 1482 | -1 |
| metaRNA.PFemirge\_9005 | 90762 | 0.009032 | JN653056.1.1473 | Bacteria;Desulfobacterota;Desulfovibrionia;Desulfovibrionales;Desulfovibrionaceae;Desulfovibrio;uncultured Desulfovibrionaceae bacterium | 95.6 | 1461 | -1 |
| metaRNA.PFemirge\_17011 | 89965 | 0.005371 | DQ799369.1.1397 | Bacteria;Bacteroidota;Bacteroidia;Bacteroidales;Bacteroidaceae;Bacteroides;uncultured bacterium | 95.5 | 1397 | -1 |
| metaRNA.PFemirge\_9501 | 87977 | 0.010235 | KP231747.1.1515 | Bacteria;Desulfobacterota;Desulfovibrionia;Desulfovibrionales;Desulfovibrionaceae;Desulfovibrio;uncultured Desulfovibrio sp. | 94.5 | 1428 | -1 |
| metaRNA.PFemirge\_2866 | 87227 | 0.007591 | CP011073.4368909.4370448 | Bacteria;Bacteroidota;Bacteroidia;Bacteroidales;Bacteroidaceae;Bacteroides;Bacteroides fragilis | 94.1 | 1508 | -1 |
| metaRNA.PFemirge\_6249 | 85064 | 0.005745 | AF157057.1.1502 | Bacteria;Deferribacterota;Deferribacteres;Deferribacterales;Deferribacteraceae;Mucispirillum;bacterium 'Lincoln Park 3' | 91.1 | 1481 | -1 |
| metaRNA.PFemirge\_5051 | 83142 | 0.007470 | MG428876.1.1497 | Bacteria;Proteobacteria;Gammaproteobacteria;Enterobacterales;Aeromonadaceae;Aeromonas;Aeromonas encheleia | 97.4 | 1492 | -1 |
| metaRNA.PFemirge\_12623 | 83119 | 0.005308 | AB574480.1.1486 | Bacteria;Bacteroidota;Bacteroidia;Bacteroidales;Bacteroidaceae;Bacteroides;Bacteroides faecichinchillae | 93.2 | 1419 | -1 |
| metaRNA.PFemirge\_16705 | 77061 | 0.005309 | EU616637.1.1399 | Bacteria;Bacteroidota;Bacteroidia;Bacteroidales;Rikenellaceae;Rikenella;uncultured Rikenellaceae bacterium | 94.2 | 1353 | -1 |
| metaRNA.PFemirge\_18143 | 76997 | 0.002475 | EU775962.1.1393 | Bacteria;Bacteroidota;Bacteroidia;Bacteroidales;Bacteroidaceae;Bacteroides;uncultured bacterium | 94.0 | 1391 | -1 |
| metaRNA.PFemirge\_11448 | 76786 | 0.003731 | KM360180.1.1492 | Bacteria;Firmicutes;Clostridia;Lachnospirales;Lachnospiraceae;Lachnoclostridium;Clostridium sp. BPY5 | 94.6 | 1451 | -1 |
| metaRNA.PFemirge\_6935 | 72015 | 0.004836 | CP012938.3352663.3354184 | Bacteria;Bacteroidota;Bacteroidia;Bacteroidales;Bacteroidaceae;Bacteroides;Bacteroides ovatus | 94.1 | 1489 | -1 |
| metaRNA.PFemirge\_27695 | 70334 | 0.006100 | KP231747.1.1515 | Bacteria;Desulfobacterota;Desulfovibrionia;Desulfovibrionales;Desulfovibrionaceae;Desulfovibrio;uncultured Desulfovibrio sp. | 94.5 | 1425 | -1 |
| metaRNA.PFemirge\_15660 | 64876 | 0.004618 | HQ819238.1.1451 | Bacteria;Bacteroidota;Bacteroidia;Bacteroidales;Bacteroidaceae;Bacteroides;uncultured organism | 98.6 | 1076 | -1 |
| metaRNA.PFemirge\_27687 | 61995 | 0.004843 | JX457157.1.1486 | Bacteria;Bacteroidota;Bacteroidia;Bacteroidales;Tannerellaceae;Parabacteroides;uncultured bacterium | 95.4 | 1484 | -1 |
| metaRNA.PFemirge\_13372 | 61735 | 0.003476 | AB470343.1.1485 | Bacteria;Bacteroidota;Bacteroidia;Bacteroidales;Tannerellaceae;Parabacteroides;Parabacteroides gordonii | 90.1 | 1374 | -1 |
| metaRNA.PFemirge\_17372 | 61610 | 0.004551 | KF318250.1.1396 | Bacteria;Verrucomicrobiota;Verrucomicrobiae;Verrucomicrobiales;Akkermansiaceae;Akkermansia;uncultured Verrucomicrobia bacterium | 93.8 | 1304 | -1 |
| metaRNA.PFemirge\_14512 | 59384 | 0.003498 | HQ757038.1.1425 | Bacteria;Firmicutes;Clostridia;Lachnospirales;Lachnospiraceae;Blautia;uncultured organism | 93.7 | 1336 | -1 |
| metaRNA.PFemirge\_6917 | 56964 | 0.003777 | HM038002.1.1488 | Bacteria;Firmicutes;Bacilli;Erysipelotrichales;Erysipelotrichaceae;[Anaerorhabdus] furcosa group;Anaerorhabdus furcosa | 97.7 | 1463 | -1 |
| metaRNA.PFemirge\_21965 | 55173 | 0.004375 | DQ817375.1.1403 | Bacteria;Firmicutes;Clostridia;Clostridiales;Clostridiaceae;Clostridium sensu stricto 1;uncultured bacterium | 97.5 | 1372 | -1 |
| metaRNA.PFemirge\_27678 | 53838 | 0.002836 | CDZC01018442.2079.3583 | Bacteria;Bacteroidota;Bacteroidia;Bacteroidales;Bacteroidaceae;Bacteroides;gut metagenome | 92.3 | 1502 | -1 |
| metaRNA.PFemirge\_28131 | 52063 | 0.013761 | KP231747.1.1515 | Bacteria;Desulfobacterota;Desulfovibrionia;Desulfovibrionales;Desulfovibrionaceae;Desulfovibrio;uncultured Desulfovibrio sp. | 89.8 | 1389 | -1 |
| metaRNA.PFemirge\_18360 | 49219 | 0.002973 | EU454363.1.1392 | Bacteria;Firmicutes;Clostridia;Lachnospirales;Lachnospiraceae;Lachnospiraceae NK4A136 group;uncultured bacterium | 92.7 | 1392 | -1 |
| metaRNA.PFemirge\_588 | 48057 | 0.005742 | X60408.1.1503 | Bacteria;Proteobacteria;Gammaproteobacteria;Enterobacterales;Aeromonadaceae;Aeromonas;Aeromonas caviae | 95.0 | 1576 | -1 |
| metaRNA.PFemirge\_4199 | 47590 | 0.004230 | JX576025.1.1509 | Bacteria;Proteobacteria;Gammaproteobacteria;Burkholderiales;Rhodocyclaceae;Azospira;uncultured Azospira sp. | 90.5 | 1439 | -1 |
| metaRNA.PFemirge\_7153 | 47190 | 0.002586 | HG934468.2032323.2033833 | Bacteria;Bacteroidota;Bacteroidia;Bacteroidales;Rikenellaceae;Mucinivorans;Mucinivorans hirudinis | 91.7 | 1487 | -1 |
| metaRNA.PFemirge\_11813 | 44035 | 0.004412 | AB554230.1.1487 | Bacteria;Bacteroidota;Bacteroidia;Bacteroidales;Rikenellaceae;Alistipes;Alistipes finegoldii | 91.3 | 1449 | -1 |
| metaRNA.PFemirge\_15835 | 43978 | 0.002820 | EU616637.1.1399 | Bacteria;Bacteroidota;Bacteroidia;Bacteroidales;Rikenellaceae;Rikenella;uncultured Rikenellaceae bacterium | 88.2 | 1351 | -1 |
| metaRNA.PFemirge\_7599 | 43490 | 0.004233 | AQHX01000018.1891.3401 | Bacteria;Bacteroidota;Bacteroidia;Bacteroidales;Bacteroidaceae;Bacteroides;Bacteroides salyersiae WAL 10018 = DSM 18765 = JCM 12988 | 91.5 | 1470 | -1 |
| metaRNA.PFemirge\_15943 | 41464 | 0.000843 | KP231747.1.1515 | Bacteria;Desulfobacterota;Desulfovibrionia;Desulfovibrionales;Desulfovibrionaceae;Desulfovibrio;uncultured Desulfovibrio sp. | 90.3 | 1404 | -1 |
| metaRNA.PFemirge\_5532 | 40528 | 0.003110 | EF400740.1.1495 | Bacteria;Firmicutes;Clostridia;Oscillospirales;Butyricicoccaceae;UCG-009;uncultured bacterium | 95.4 | 1382 | -1 |
| metaRNA.PFemirge\_9475 | 40417 | 0.004681 | AB821059.1.1460 | Bacteria;Proteobacteria;Alphaproteobacteria;Rhizobiales;Xanthobacteraceae;Bradyrhizobium;uncultured bacterium | 89.9 | 1345 | -1 |
| metaRNA.PFemirge\_27873 | 40074 | 0.002329 | CP012937.5517779.5519304 | Bacteria;Bacteroidota;Bacteroidia;Bacteroidales;Bacteroidaceae;Bacteroides;Bacteroides thetaiotaomicron | 91.0 | 1484 | -1 |
| metaRNA.PFemirge\_14034 | 39306 | 0.003854 | FJ269067.1.1493 | Bacteria;Bacteroidota;Bacteroidia;Bacteroidales;Bacteroidaceae;Bacteroides;iron-reducing bacterium enrichment culture clone HN86 | 88.8 | 1312 | -1 |
| metaRNA.PFemirge\_2210 | 38691 | 0.001759 | AB649276.1.1495 | Bacteria;Firmicutes;Clostridia;Lachnospirales;Lachnospiraceae;Anaerostignum;Anaerotignum propionicum | 97.5 | 1453 | -1 |
| metaRNA.PFemirge\_5412 | 37875 | 0.003665 | ABJL02000006.58.1568 | Bacteria;Bacteroidota;Bacteroidia;Bacteroidales;Bacteroidaceae;Bacteroides;Bacteroides intestinalis DSM 17393 | 93.1 | 1503 | -1 |
| metaRNA.PFemirge\_13253 | 37864 | 0.004234 | HQ785173.1.1448 | Bacteria;Bacteroidota;Bacteroidia;Bacteroidales;Bacteroidaceae;Bacteroides;uncultured organism | 93.1 | 1419 | -1 |
| metaRNA.PFemirge\_2360 | 37024 | 0.002347 | HG970990.1.1557 | Bacteria;Firmicutes;Bacilli;Erysipelotrichales;Erysipelotrichaceae;Breznakia;uncultured bacterium | 93.0 | 1480 | -1 |
| metaRNA.PFemirge\_6702 | 34553 | 0.001703 | GQ360012.1.1488 | Bacteria;Bacteroidota;Bacteroidia;Bacteroidales;Tannerellaceae;Macellibacteroides;uncultured bacterium | 88.4 | 1493 | -1 |
| metaRNA.PFemirge\_7412 | 34520 | 0.002524 | FJ534957.1.1501 | Bacteria;Firmicutes;Clostridia;Lachnospirales;Lachnospiraceae;Lachnospiraceae FCS020 group;uncultured bacterium | 94.6 | 1405 | -1 |
| metaRNA.PFemirge\_24591 | 34155 | 0.002806 | AY916388.1.1341 | Bacteria;Proteobacteria;Alphaproteobacteria;Rhodospirillales;uncultured;uncultured bacterium | 95.2 | 1341 | -1 |
| metaRNA.PFemirge\_4769 | 33628 | 0.002101 | GQ871713.1.1490 | Bacteria;Bacteroidota;Bacteroidia;Bacteroidales;Rikenellaceae;Alistipes;uncultured bacterium | 87.2 | 1497 | -1 |
| metaRNA.PFemirge\_6069 | 32239 | 0.003151 | ABFK02000017.223676.225190 | Bacteria;Bacteroidota;Bacteroidia;Bacteroidales;Rikenellaceae;Alistipes;Alistipes putredinis DSM 17216 | 95.1 | 1487 | -1 |
| metaRNA.PFemirge\_6071 | 32001 | 0.001656 | EF403697.1.1493 | Bacteria;Bacteroidota;Bacteroidia;Bacteroidales;Bacteroidaceae;Bacteroides;uncultured bacterium | 91.1 | 1498 | -1 |
| metaRNA.PFemirge\_27668 | 31507 | 0.005292 | KP231747.1.1515 | Bacteria;Desulfobacterota;Desulfovibrionia;Desulfovibrionales;Desulfovibrionaceae;Desulfovibrio;uncultured Desulfovibrio sp. | 95.2 | 1504 | -1 |
| metaRNA.PFemirge\_17188 | 31339 | 0.002435 | AB574482.1.1489 | Bacteria;Bacteroidota;Bacteroidia;Bacteroidales;Tannerellaceae;Parabacteroides;Parabacteroides chinchillae | 90.7 | 1396 | -1 |
| metaRNA.PFemirge\_7737 | 30801 | 0.002818 | AQHW01000007.13.1527 | Bacteria;Bacteroidota;Bacteroidia;Bacteroidales;Tannerellaceae;Parabacteroides;Parabacteroides gordonii MS-1 | 97.1 | 1408 | -1 |
| metaRNA.PFemirge\_28104 | 29759 | 0.006325 | AB574482.1.1489 | Bacteria;Bacteroidota;Bacteroidia;Bacteroidales;Tannerellaceae;Parabacteroides;Parabacteroides chinchillae | 94.7 | 1480 | -1 |
| metaRNA.PFemirge\_26379 | 29684 | 0.003009 | MG367100.1.1491 | Bacteria;Firmicutes;Clostridia;Oscillospirales;Oscillospiraceae;Colidextribacter;uncultured Clostridiales Family IV bacterium | 91.4 | 1282 | -1 |
| metaRNA.PFemirge\_18570 | 29336 | 0.001437 | AGES01000014.73256.74770 | Bacteria;Bacteroidota;Bacteroidia;Bacteroidales;Tannerellaceae;Parabacteroides;Parabacteroides sp. HGS0025 | 92.6 | 1397 | -1 |
| metaRNA.PFemirge\_2498 | 28453 | 0.010021 | FJ881199.1.1509 | Bacteria;Desulfobacterota;Desulfovibrionia;Desulfovibrionales;Desulfovibrionaceae;uncultured;uncultured bacterium | 93.2 | 1506 | -1 |
| metaRNA.PFemirge\_2358 | 28215 | 0.002376 | FR749935.1.1522 | Bacteria;Firmicutes;Clostridia;Lachnospirales;Lachnospiraceae;[Eubacterium] fissicatena group;Faecalicatena fissicatena | 98.2 | 1469 | -1 |
| metaRNA.PFemirge\_12854 | 27513 | 0.001735 | DQ817541.1.1479 | Bacteria;Proteobacteria;Gammaproteobacteria;Burkholderiales;Chromobacteriaceae;Crenobacter;uncultured bacterium | 99.4 | 1397 | -1 |
| metaRNA.PFemirge\_27756 | 26513 | 0.002917 | AB238925.1.1486 | Bacteria;Bacteroidota;Bacteroidia;Bacteroidales;Tannerellaceae;Parabacteroides;Parabacteroides distasonis | 93.7 | 1398 | -1 |
| metaRNA.PFemirge\_10716 | 26103 | 0.001165 | EU234274.1.1501 | Bacteria;Proteobacteria;Gammaproteobacteria;Burkholderiales;Rhodocyclaceae;C39;uncultured bacterium | 97.5 | 1339 | -1 |
| metaRNA.PFemirge\_15318 | 25942 | 0.003179 | EF401958.1.1488 | Bacteria;Bacteroidota;Bacteroidia;Bacteroidales;Tannerellaceae;Parabacteroides;uncultured bacterium | 91.4 | 1401 | -1 |
| metaRNA.PFemirge\_15784 | 25641 | 0.001731 | HQ761452.1.1440 | Bacteria;Bacteroidota;Bacteroidia;Bacteroidales;Tannerellaceae;Parabacteroides;uncultured organism | 91.5 | 1304 | -1 |
| metaRNA.PFemirge\_7513 | 25590 | 0.001841 | JX457309.1.1485 | Bacteria;Bacteroidota;Bacteroidia;Bacteroidales;Rs-E47 termite group;uncultured bacterium | 92.8 | 1466 | -1 |
| metaRNA.PFemirge\_26390 | 24858 | 0.001415 | EU510219.1.1285 | Bacteria;Firmicutes;Clostridia;Lachnospirales;Lachnospiraceae;Lachnospiraceae UCG-008;uncultured bacterium | 93.0 | 1280 | -1 |
| metaRNA.PFemirge\_7443 | 24069 | 0.002130 | CP028523.140101.141607 | Bacteria;Firmicutes;Clostridia;Peptostreptococcales-Tissierellales;Peptostreptococcaceae;Clostridioides;Clostridioides difficile | 99.6 | 1393 | -1 |
| metaRNA.PFemirge\_13838 | 22906 | 0.005820 | GQ898343.1.1489 | Bacteria;Bacteroidota;Bacteroidia;Bacteroidales;Bacteroidaceae;Bacteroides;uncultured bacterium | 96.1 | 1423 | -1 |
| metaRNA.PFemirge\_4741 | 22622 | 0.000796 | AB298756.1.1471 | Bacteria;Firmicutes;Clostridia;Lachnospirales;Lachnospiraceae;Anaerostignum;Anaerotignum aminivorans | 94.3 | 1429 | -1 |
| metaRNA.PFemirge\_3434 | 22533 | 0.000815 | AY945907.1.1506 | Bacteria;Proteobacteria;Gammaproteobacteria;Burkholderiales;Rhodocyclaceae;Propionivibrio;uncultured bacterium | 95.4 | 1418 | -1 |
| metaRNA.PFemirge\_27674 | 21679 | 0.002611 | AQHX01000018.1891.3401 | Bacteria;Bacteroidota;Bacteroidia;Bacteroidales;Bacteroidaceae;Bacteroides;Bacteroides salyersiae WAL 10018 = DSM 18765 = JCM 12988 | 94.5 | 1500 | -1 |
| metaRNA.PFemirge\_3462 | 20251 | 0.001424 | CXSE01001446.1371.2879 | Bacteria;Bacteroidota;Bacteroidia;Bacteroidales;Bacteroidaceae;Bacteroides;human gut metagenome | 94.7 | 1466 | -1 |
| metaRNA.PFemirge\_11898 | 20222 | 0.001264 | GQ360012.1.1488 | Bacteria;Bacteroidota;Bacteroidia;Bacteroidales;Tannerellaceae;Macellibacteroides;uncultured bacterium | 87.4 | 1452 | -1 |
| metaRNA.PFemirge\_23423 | 19859 | 0.000049 | GQ360012.1.1488 | Bacteria;Bacteroidota;Bacteroidia;Bacteroidales;Tannerellaceae;Macellibacteroides;uncultured bacterium | 87.5 | 1358 | -1 |
| metaRNA.PFemirge\_15680 | 19569 | 0.005455 | DQ797819.1.1407 | Bacteria;Desulfobacterota;Desulfovibrionia;Desulfovibrionales;Desulfovibrionaceae;Desulfovibrio;uncultured bacterium | 92.2 | 1406 | -1 |
| metaRNA.PFemirge\_13238 | 19388 | 0.001251 | JX198651.1.1513 | Bacteria;Firmicutes;Clostridia;Oscillospirales;Butyricicoccaceae;UCG-009;uncultured bacterium | 92.8 | 1331 | -1 |
| metaRNA.PFemirge\_17782 | 18911 | 0.001614 | DQ806700.1.1391 | Bacteria;Firmicutes;Clostridia;Oscillospirales;Oscillospiraceae;Colidextribacter;uncultured bacterium | 91.2 | 1191 | -1 |
| metaRNA.PFemirge\_812 | 18899 | 0.001165 | HM017143.1.1714 | Eukaryota;Amorphea;Amoebozoa;Discosea;Flabellinia;Dactylopodida;Angulamoeba;Angulamoeba microcystivorans | 95.9 | 1432 | -1 |
| metaRNA.PFemirge\_27697 | 18748 | 0.002996 | AM275521.1.1477 | Bacteria;Bacteroidota;Bacteroidia;Bacteroidales;Bacteroidaceae;Bacteroides;uncultured bacterium | 94.0 | 1465 | -1 |
| metaRNA.PFemirge\_17975 | 17821 | 0.002583 | GQ448246.1.1389 | Bacteria;Bacteroidota;Bacteroidia;Bacteroidales;Bacteroidaceae;Bacteroides;uncultured bacterium | 93.4 | 1389 | -1 |
| metaRNA.PFemirge\_3793 | 17358 | 0.001333 | CBEW010000180.8693.10203 | Bacteria;Cyanobacteria;Vampirivibrionia;Gastranaerophilales;Clostridium sp. CAG:306 | 96.7 | 1473 | -1 |
| metaRNA.PFemirge\_7161 | 16891 | 0.003040 | KC994754.1.1487 | Bacteria;Actinobacteriota;Actinobacteria;Micrococcales;Microbacteriaceae;Agromyces;uncultured Actinomycetales bacterium | 96.1 | 1425 | -1 |
| metaRNA.PFemirge\_6863 | 16725 | 0.000788 | EU622704.1.1488 | Bacteria;Firmicutes;Clostridia;Oscillospirales;Oscillospiraceae;Colidextribacter;uncultured bacterium | 91.7 | 1395 | -1 |
| metaRNA.PFemirge\_27736 | 16613 | 0.001373 | MG367100.1.1491 | Bacteria;Firmicutes;Clostridia;Oscillospirales;Oscillospiraceae;Colidextribacter;uncultured Clostridiales Family IV bacterium | 93.6 | 1328 | -1 |
| metaRNA.PFemirge\_12879 | 15836 | 0.002607 | AF129440.1.1436 | Bacteria;Proteobacteria;Gammaproteobacteria;Enterobacterales;Enterobacteriaceae;Klebsiella;Klebsiella oxytoca | 90.8 | 1420 | -1 |
| metaRNA.PFemirge\_2897 | 14969 | 0.000694 | FBXW01000008.267.1775 | Bacteria;Firmicutes;Clostridia;Oscillospirales;Oscillospiraceae;uncultured;Intestinimonas sp. GM5 | 94.8 | 1483 | -1 |
| metaRNA.PFemirge\_9752 | 14882 | 0.000858 | GQ898126.1.1466 | Bacteria;Proteobacteria;Alphaproteobacteria;Rhodospirillales;uncultured;uncultured bacterium | 95.8 | 1444 | -1 |
| metaRNA.PFemirge\_14964 | 14737 | 0.001020 | JN619526.1.1407 | Bacteria;Synergistota;Synergistia;Synergistales;Synergistaceae;uncultured;uncultured bacterium | 93.1 | 1278 | -1 |
| metaRNA.PFemirge\_1686 | 14529 | 0.000861 | AF479766.1.2092 | Bacteria;Proteobacteria;Gammaproteobacteria;Burkholderiales;Rhodocyclaceae;Dechloromonas;Dechloromonas sp. HZ | 99.1 | 1419 | -1 |
| metaRNA.PFemirge\_16674 | 14104 | 0.001307 | EU464332.1.1399 | Bacteria;Bacteroidota;Bacteroidia;Bacteroidales;Rikenellaceae;dgA-11 gut group;uncultured bacterium | 90.6 | 1397 | -1 |
| metaRNA.PFemirge\_6591 | 13434 | 0.002127 | EF559167.1.1492 | Bacteria;Firmicutes;Clostridia;Lachnospirales;Lachnospiraceae;Anaerostignum;uncultured bacterium | 89.4 | 1395 | -1 |
| metaRNA.PFemirge\_13425 | 13380 | 0.001237 | AM265457.1.1438 | Bacteria;Firmicutes;Clostridia;Oscillospirales;Oscillospiraceae;Colidextribacter;uncultured bacterium | 93.3 | 1383 | -1 |
| metaRNA.PFemirge\_1105 | 13003 | 0.000501 | HW066305.1.1532 | Bacteria;Firmicutes;Clostridia;Clostridia UCG-014;unidentified | 97.1 | 1465 | -1 |
| metaRNA.PFemirge\_11249 | 12978 | 0.000931 | KR822473.1.1454 | Bacteria;Firmicutes;Clostridia;Peptostreptococcales-Tissierellales;Anaerovoracaceae;[Eubacterium] nodatum group;bacterium PM5-7 | 98.7 | 1449 | -1 |
| metaRNA.PFemirge\_4309 | 12944 | 0.001028 | AB034016.1.1515 | Bacteria;Cyanobacteria;Vampirivibrionia;Gastranaerophilales;uncultured rumen bacterium 4C0d-2 | 94.8 | 1469 | -1 |
| metaRNA.PFemirge\_7230 | 12863 | 0.000729 | CEAB01046477.78830.80315 | Bacteria;Proteobacteria;Alphaproteobacteria;Rhodospirillales;uncultured;gut metagenome | 93.3 | 1444 | -1 |
| metaRNA.PFemirge\_22844 | 12617 | 0.000909 | GQ449181.1.1364 | Bacteria;Bacteroidota;Bacteroidia;Bacteroidales;Rikenellaceae;Rikenellaceae RC9 gut group;uncultured bacterium | 89.7 | 1279 | -1 |
| metaRNA.PFemirge\_13870 | 12476 | 0.000633 | KM650567.1.1481 | Bacteria;Firmicutes;Negativicutes;Acidaminococcales;Acidaminococcaceae;uncultured;uncultured Firmicutes bacterium | 92.8 | 1370 | -1 |
| metaRNA.PFemirge\_6724 | 12332 | 0.004080 | CEBY01043209.9650.11156 | Bacteria;Bacteroidota;Bacteroidia;Bacteroidales;Rikenellaceae;Alistipes;gut metagenome | 90.8 | 1486 | -1 |
| metaRNA.PFemirge\_7297 | 12325 | 0.001339 | HM630236.1.1451 | Bacteria;Bacteroidota;Bacteroidia;Bacteroidales;Rikenellaceae;dgA-11 gut group;uncultured bacterium | 90.9 | 1411 | -1 |
| metaRNA.PFemirge\_15716 | 12086 | 0.001541 | EU771706.1.1407 | Bacteria;Verrucomicrobiota;Lentisphaeria;Victivallales;Victivallaceae;uncultured bacterium | 88.8 | 1397 | -1 |
| metaRNA.PFemirge\_22949 | 11682 | 0.000009 | KP231747.1.1515 | Bacteria;Desulfobacterota;Desulfovibrionia;Desulfovibrionales;Desulfovibrionaceae;Desulfovibrio;uncultured Desulfovibrio sp. | 94.0 | 1216 | -1 |
| metaRNA.PFemirge\_16111 | 11468 | 0.001253 | FJ366372.1.1363 | Bacteria;Bacteroidota;Bacteroidia;Bacteroidales;Rikenellaceae;Alistipes;uncultured bacterium | 91.5 | 1223 | -1 |
| metaRNA.PFemirge\_10824 | 11396 | 0.001220 | AB234530.1.1368 | Bacteria;Desulfobacterota;Desulfovibrionia;Desulfovibrionales;Desulfovibrionaceae;uncultured;uncultured delta proteobacterium | 94.9 | 1315 | -1 |
| metaRNA.PFemirge\_23168 | 11342 | 0.000598 | FJ825520.1.1372 | Bacteria;Firmicutes;Clostridia;Christensenellales;Christensenellaceae;Christensenellaceae R-7 group;uncultured bacterium | 91.9 | 1211 | -1 |
| metaRNA.PFemirge\_9898 | 10896 | 0.000807 | LT725663.1.1450 | Bacteria;Bacteroidota;Bacteroidia;Bacteroidales;Tannerellaceae;Parabacteroides;Parabacteroides sp. Marseille-P3668 | 94.3 | 1450 | -1 |
| metaRNA.PFemirge\_27732 | 10835 | 0.002573 | CP012937.2383895.2385420 | Bacteria;Bacteroidota;Bacteroidia;Bacteroidales;Bacteroidaceae;Bacteroides;Bacteroides thetaiotaomicron | 94.4 | 1420 | -1 |
| metaRNA.PFemirge\_27684 | 10806 | 0.001116 | ABVO01000045.3575.5089 | Bacteria;Bacteroidota;Bacteroidia;Bacteroidales;Bacteroidaceae;Bacteroides;Bacteroides eggerthii DSM 20697 | 93.5 | 1489 | -1 |
| metaRNA.PFemirge\_18462 | 10779 | 0.001364 | HG934468.2032323.2033833 | Bacteria;Bacteroidota;Bacteroidia;Bacteroidales;Rikenellaceae;Mucinivorans;Mucinivorans hirudinis | 89.9 | 1393 | -1 |
| metaRNA.PFemirge\_4327 | 10770 | 0.001436 | JQ083789.1.1511 | Bacteria;Desulfobacterota;Desulfovibrionia;Desulfovibrionales;Desulfovibrionaceae;Bilophila;uncultured bacterium | 93.2 | 1500 | -1 |
| metaRNA.PFemirge\_21649 | 10711 | 0.000048 | DI433844.4.1378 | Bacteria;Firmicutes;Bacilli;Bacillales;Bacillaceae;Bacillus;Bacillus sp. EBW4 | 94.5 | 1357 | -1 |
| metaRNA.PFemirge\_690 | 10475 | 0.000751 | LC222972.1.1573 | Eukaryota;Amorphea;Amoebozoa;uncultured eukaryote | 99.5 | 1480 | -1 |
| metaRNA.PFemirge\_171 | 9704 | 0.000565 | AB901579.1.1807 | Eukaryota;Amorphea;Obazoa;Opisthokonta;Holozoa;Choanozoa;Metazoa;Animalia;BCP clade;Bilateria;Protostomia;Lophotrochozoa;Rotifera;Monogononta;Ploimida;uncultured eukaryote | 98.5 | 1668 | -1 |
| metaRNA.PFemirge\_27775 | 9595 | 0.000338 | BAAZ01000703.1371.2875 | Bacteria;Bacteroidota;Bacteroidia;Bacteroidales;Tannerellaceae;Parabacteroides;human gut metagenome | 92.7 | 1227 | -1 |
| metaRNA.PFemirge\_21587 | 9559 | 0.001205 | EU772341.1.1387 | Bacteria;Firmicutes;Clostridia;Oscillospirales;[Eubacterium] coprostanoligenes group;uncultured bacterium | 91.7 | 1212 | -1 |
| metaRNA.PFemirge\_6166 | 9228 | 0.000389 | KT952743.1.1492 | Bacteria;Bacteroidota;Bacteroidia;Bacteroidales;Rikenellaceae;Rikenella;uncultured bacterium | 92.0 | 1474 | -1 |
| metaRNA.PFemirge\_8798 | 9069 | 0.000631 | KC311367.1.1475 | Bacteria;Firmicutes;Clostridia;Oscillospirales;Oscillospiraceae;Intestinimonas;Intestinimonas butyriciproducens | 95.6 | 1471 | -1 |
| metaRNA.PFemirge\_28310 | 9039 | 0.000170 | KP231747.1.1515 | Bacteria;Desulfobacterota;Desulfovibrionia;Desulfovibrionales;Desulfovibrionaceae;Desulfovibrio;uncultured Desulfovibrio sp. | 93.9 | 1504 | -1 |
| metaRNA.PFemirge\_18629 | 8787 | 0.001209 | DQ799406.1.1391 | Bacteria;Bacteroidota;Bacteroidia;Bacteroidales;Bacteroidaceae;Bacteroides;uncultured bacterium | 93.9 | 1385 | -1 |
| metaRNA.PFemirge\_17272 | 8472 | 0.000885 | EU507529.1.1407 | Bacteria;Firmicutes;Clostridia;Oscillospirales;Butyricicoccaceae;UCG-009;uncultured bacterium | 94.1 | 1390 | -1 |
| metaRNA.PFemirge\_8940 | 8453 | 0.001871 | KP231747.1.1515 | Bacteria;Desulfobacterota;Desulfovibrionia;Desulfovibrionales;Desulfovibrionaceae;Desulfovibrio;uncultured Desulfovibrio sp. | 92.2 | 1360 | -1 |
| metaRNA.PFemirge\_6295 | 8296 | 0.000755 | HQ806032.1.1447 | Bacteria;Bacteroidota;Bacteroidia;Bacteroidales;Tannerellaceae;Parabacteroides;uncultured organism | 94.4 | 1454 | -1 |
| metaRNA.PFemirge\_2154 | 8262 | 0.000043 | HM123962.1.1510 | Bacteria;Desulfobacterota;Desulfovibrionia;Desulfovibrionales;Desulfovibrionaceae;uncultured;uncultured bacterium | 89.7 | 1274 | -1 |
| metaRNA.PFemirge\_11874 | 8102 | 0.000453 | JN680601.1.1450 | Bacteria;Bacteroidota;Bacteroidia;Bacteroidales;Rikenellaceae;Alistipes;uncultured Rikenellaceae bacterium | 96.1 | 1379 | -1 |
| metaRNA.PFemirge\_10080 | 7986 | 0.000615 | EF400726.1.1488 | Bacteria;Bacteroidota;Bacteroidia;Bacteroidales;Marinifilaceae;Odoribacter;uncultured bacterium | 89.4 | 1270 | -1 |
| metaRNA.PFemirge\_26353 | 7983 | 0.002068 | ADLN01000129.202.1639 | Bacteria;Firmicutes;Clostridia;Lachnospirales;Lachnospiraceae;Incertae Sedis;Hungatella hathewayi WAL-18680 | 90.2 | 1239 | -1 |
| metaRNA.PFemirge\_12250 | 7890 | 0.000656 | KC669122.1.1486 | Bacteria;Firmicutes;Clostridia;Lachnospirales;Lachnospiraceae;Epulopiscium;uncultured bacterium | 98.6 | 1317 | -1 |
| metaRNA.PFemirge\_7289 | 7861 | 0.000472 | HG792196.1.1487 | Bacteria;Firmicutes;Bacilli;Erysipelotrichales;Erysipelatoclostridiaceae;Coprobacillus;uncultured bacterium | 95.9 | 1449 | -1 |
| metaRNA.PFemirge\_6802 | 7825 | 0.000938 | AGES01000014.73256.74770 | Bacteria;Bacteroidota;Bacteroidia;Bacteroidales;Tannerellaceae;Parabacteroides;Parabacteroides sp. HGS0025 | 89.9 | 1491 | -1 |
| metaRNA.PFemirge\_28141 | 7709 | 0.001026 | KP231747.1.1515 | Bacteria;Desulfobacterota;Desulfovibrionia;Desulfovibrionales;Desulfovibrionaceae;Desulfovibrio;uncultured Desulfovibrio sp. | 92.3 | 1404 | -1 |
| metaRNA.PFemirge\_12464 | 7602 | 0.000497 | HQ793579.1.1446 | Bacteria;Bacteroidota;Bacteroidia;Bacteroidales;Marinifilaceae;Butyricimonas;uncultured organism | 96.8 | 1254 | -1 |
| metaRNA.PFemirge\_27964 | 7567 | 0.000571 | KP944118.1.1420 | Bacteria;Bacteroidota;Bacteroidia;Bacteroidales;Bacteroidaceae;Bacteroides;Bacteroides thetaiotaomicron | 94.2 | 1353 | -1 |
| metaRNA.PFemirge\_2278 | 7519 | 0.000785 | LC028669.1.1525 | Bacteria;Firmicutes;Clostridia;Oscillospirales;Butyricicoccaceae;UCG-009;uncultured bacterium | 94.4 | 1326 | -1 |
| metaRNA.PFemirge\_16116 | 7503 | 0.001083 | EU616637.1.1399 | Bacteria;Bacteroidota;Bacteroidia;Bacteroidales;Rikenellaceae;Rikenella;uncultured Rikenellaceae bacterium | 92.1 | 1321 | -1 |
| metaRNA.PFemirge\_27694 | 7391 | 0.000596 | HQ778533.1.1446 | Bacteria;Bacteroidota;Bacteroidia;Bacteroidales;Tannerellaceae;Parabacteroides;uncultured organism | 95.0 | 1386 | -1 |
| metaRNA.PFemirge\_17760 | 7377 | 0.000272 | EU504186.1.1398 | Bacteria;Firmicutes;Clostridia;Lachnospirales;Lachnospiraceae;Lachnospiraceae NK4A136 group;uncultured bacterium | 92.7 | 1391 | -1 |
| metaRNA.PFemirge\_4693 | 7352 | 0.000270 | AB574482.1.1489 | Bacteria;Bacteroidota;Bacteroidia;Bacteroidales;Tannerellaceae;Parabacteroides;Parabacteroides chinchillae | 92.1 | 1496 | -1 |
| metaRNA.PFemirge\_12796 | 7100 | 0.000920 | HQ761612.1.1444 | Bacteria;Bacteroidota;Bacteroidia;Bacteroidales;Tannerellaceae;Parabacteroides;uncultured organism | 91.2 | 1430 | -1 |
| metaRNA.PFemirge\_1920 | 7029 | 0.000512 | CP022115.2965538.2967082 | Bacteria;Proteobacteria;Gammaproteobacteria;Burkholderiales;Aquaspirillaceae;Laribacter;Laribacter hongkongensis | 99.6 | 1416 | -1 |
| metaRNA.PFemirge\_11849 | 7018 | 0.001477 | CXSE01001446.1371.2879 | Bacteria;Bacteroidota;Bacteroidia;Bacteroidales;Bacteroidaceae;Bacteroides;human gut metagenome | 94.8 | 1387 | -1 |
| metaRNA.PFemirge\_28094 | 6905 | 0.001063 | JQ083789.1.1511 | Bacteria;Desulfobacterota;Desulfovibrionia;Desulfovibrionales;Desulfovibrionaceae;Bilophila;uncultured bacterium | 91.1 | 1506 | -1 |
| metaRNA.PFemirge\_6452 | 6646 | 0.000408 | AB649277.1.1490 | Bacteria;Firmicutes;Bacilli;Erysipelotrichales;Erysipelotrichaceae;Faecalitalea;Absiella dolichum | 93.6 | 1342 | -1 |
| metaRNA.PFemirge\_19649 | 6201 | 0.000406 | MG367100.1.1491 | Bacteria;Firmicutes;Clostridia;Oscillospirales;Oscillospiraceae;Colidextribacter;uncultured Clostridiales Family IV bacterium | 92.6 | 1384 | -1 |
| metaRNA.PFemirge\_13026 | 6076 | 0.000916 | DQ856557.1.1487 | Bacteria;Bacteroidota;Bacteroidia;Bacteroidales;Bacteroidaceae;Bacteroides;uncultured bacterium | 93.7 | 1444 | -1 |
| metaRNA.PFemirge\_17001 | 5994 | 0.001040 | AB554233.1.1486 | Bacteria;Bacteroidota;Bacteroidia;Bacteroidales;Rikenellaceae;Alistipes;Alistipes shahii | 94.9 | 1356 | -1 |
| metaRNA.PFemirge\_7053 | 5979 | 0.000377 | EF400726.1.1488 | Bacteria;Bacteroidota;Bacteroidia;Bacteroidales;Marinifilaceae;Odoribacter;uncultured bacterium | 94.7 | 1485 | -1 |
| metaRNA.PFemirge\_6600 | 5903 | 0.000395 | HQ745788.1.1434 | Bacteria;Firmicutes;Clostridia;Peptostreptococcales-Tissierellales;Peptostreptococcaceae;Romboutsia;uncultured organism | 97.4 | 1253 | -1 |
| metaRNA.PFemirge\_16020 | 5818 | 0.000293 | KP231747.1.1515 | Bacteria;Desulfobacterota;Desulfovibrionia;Desulfovibrionales;Desulfovibrionaceae;Desulfovibrio;uncultured Desulfovibrio sp. | 94.1 | 1233 | -1 |
| metaRNA.PFemirge\_637 | 5807 | 0.000307 | JN547325.1.1576 | Eukaryota;Amorphea;Amoebozoa;Discosea;Flabellinia;Dactylopodida;Angulamoeba;uncultured eukaryote | 93.4 | 1297 | -1 |
| metaRNA.PFemirge\_11565 | 5796 | 0.001286 | HQ760530.1.1440 | Bacteria;Bacteroidota;Bacteroidia;Bacteroidales;Tannerellaceae;Parabacteroides;uncultured organism | 93.3 | 1442 | -1 |
| metaRNA.PFemirge\_6509 | 5790 | 0.000467 | EF401817.1.1484 | Bacteria;Bacteroidota;Bacteroidia;Bacteroidales;Bacteroidaceae;Bacteroides;uncultured bacterium | 90.9 | 1488 | -1 |
| metaRNA.PFemirge\_8090 | 5774 | 0.003659 | AB470343.1.1485 | Bacteria;Bacteroidota;Bacteroidia;Bacteroidales;Tannerellaceae;Parabacteroides;Parabacteroides gordonii | 94.9 | 1478 | -1 |
| metaRNA.PFemirge\_14207 | 5674 | 0.000051 | JX105679.1.1392 | Bacteria;Bacteroidota;Bacteroidia;Bacteroidales;Bacteroidaceae;Bacteroides;uncultured bacterium | 94.9 | 1295 | -1 |
| metaRNA.PFemirge\_10978 | 5590 | 0.000740 | FR669217.1.1452 | Bacteria;Firmicutes;Clostridia;Oscillospirales;Oscillospiraceae;Colidextribacter;uncultured bacterium | 93.6 | 1318 | -1 |
| metaRNA.PFemirge\_17542 | 5494 | 0.001328 | DQ799526.1.1402 | Bacteria;Bacteroidota;Bacteroidia;Bacteroidales;Bacteroidaceae;Bacteroides;uncultured bacterium | 92.2 | 1400 | -1 |
| metaRNA.PFemirge\_10311 | 5421 | 0.000290 | HM630210.1.1461 | Bacteria;Firmicutes;Bacilli;RF39;uncultured bacterium | 97.0 | 1301 | -1 |
| metaRNA.PFemirge\_6695 | 5385 | 0.000395 | DQ817541.1.1479 | Bacteria;Proteobacteria;Gammaproteobacteria;Burkholderiales;Chromobacteriaceae;Crenobacter;uncultured bacterium | 94.8 | 1417 | -1 |
| metaRNA.PFemirge\_16166 | 5209 | 0.000023 | HQ761452.1.1440 | Bacteria;Bacteroidota;Bacteroidia;Bacteroidales;Tannerellaceae;Parabacteroides;uncultured organism | 92.6 | 1306 | -1 |
| metaRNA.PFemirge\_4585 | 5174 | 0.000735 | CP003153.1901384.1902805 | Bacteria;Proteobacteria;Gammaproteobacteria;Burkholderiales;Rhodocyclaceae;Azospira;Azospira oryzae PS | 95.2 | 1410 | -1 |
| metaRNA.PFemirge\_21547 | 5097 | 0.000410 | DQ793254.1.1377 | Bacteria;Firmicutes;Clostridia;Oscillospirales;Ruminococcaceae;Faecalibacterium;uncultured bacterium | 91.1 | 1287 | -1 |
| metaRNA.PFemirge\_4545 | 5058 | 0.000219 | EU158190.1.1510 | Bacteria;Firmicutes;Clostridia;Oscillospirales;Hydrogenoanaerobacterium;Hydrogenoanaerobacterium saccharovorans | 94.1 | 1316 | -1 |
| metaRNA.PFemirge\_12515 | 5051 | 0.001993 | HQ681866.1.1451 | Bacteria;Bacteroidota;Bacteroidia;Bacteroidales;Bacteroidaceae;Bacteroides;uncultured bacterium | 93.5 | 1452 | -1 |
| metaRNA.PFemirge\_17626 | 4738 | 0.000010 | LARG01000025.1.1402 | Bacteria;Bacteroidota;Bacteroidia;Bacteroidales;Rikenellaceae;Alistipes;Alistipes sp. N15.MGS-157 | 89.4 | 1392 | -1 |
| metaRNA.PFemirge\_1474 | 4651 | 0.000427 | CP021963.420966.422519 | Bacteria;Proteobacteria;Gammaproteobacteria;Enterobacterales;Enterobacteriaceae;Citrobacter;Citrobacter youngae | 98.9 | 1495 | -1 |
| metaRNA.PFemirge\_16109 | 4634 | 0.000006 | DQ307722.1.1396 | Bacteria;Bacteroidota;Bacteroidia;Bacteroidales;Rikenellaceae;Alistipes;uncultured Bacteroidetes bacterium | 87.5 | 1403 | -1 |
| metaRNA.PFemirge\_27675 | 4618 | 0.001813 | AB574481.1.1484 | Bacteria;Bacteroidota;Bacteroidia;Bacteroidales;Bacteroidaceae;Bacteroides;Bacteroides stercorirosoris | 95.7 | 1473 | -1 |
| metaRNA.PFemirge\_16159 | 4588 | 0.001568 | CDZC01018442.2079.3583 | Bacteria;Bacteroidota;Bacteroidia;Bacteroidales;Bacteroidaceae;Bacteroides;gut metagenome | 92.1 | 1405 | -1 |
| metaRNA.PFemirge\_1849 | 4553 | 0.000866 | CP011612.1889203.1890758 | Bacteria;Proteobacteria;Gammaproteobacteria;Enterobacterales;Enterobacteriaceae;Citrobacter;Citrobacter freundii | 88.2 | 1541 | -1 |
| metaRNA.PFemirge\_16161 | 4535 | 0.000079 | DQ799295.1.1402 | Bacteria;Bacteroidota;Bacteroidia;Bacteroidales;Bacteroidaceae;Bacteroides;uncultured bacterium | 89.4 | 1408 | -1 |
| metaRNA.PFemirge\_2652 | 4485 | 0.000820 | AF192152.1.1542 | Bacteria;Desulfobacterota;Desulfovibrionia;Desulfovibrionales;Desulfovibrionaceae;Desulfovibrio;Desulfovibrio piger | 90.7 | 1499 | -1 |
| metaRNA.PFemirge\_13062 | 4451 | 0.000607 | CBVB010000006.254456.255954 | Bacteria;Bacteroidota;Bacteroidia;Bacteroidales;Bacteroidaceae;Bacteroides;Bacteroidaceae bacterium MS4 | 94.4 | 1443 | -1 |
| metaRNA.PFemirge\_754 | 4436 | 0.000780 | CP003153.1901384.1902805 | Bacteria;Proteobacteria;Gammaproteobacteria;Burkholderiales;Rhodocyclaceae;Azospira;Azospira oryzae PS | 94.3 | 1435 | -1 |
| metaRNA.PFemirge\_185 | 4357 | 0.000244 | LIUU01000102.1484.3268 | Eukaryota;Amorphea;Obazoa;Opisthokonta;Nucletmycea;Fungi;Dikarya;Ascomycota;Saccharomycotina;Saccharomycetes;Saccharomycetales;Saccharomycetaceae;Zygosaccharomyces;Saccharomyces cerevisiae W303 | 98.5 | 1630 | -1 |
| metaRNA.PFemirge\_30260 | 4338 | 0.000080 | KP231747.1.1515 | Bacteria;Desulfobacterota;Desulfovibrionia;Desulfovibrionales;Desulfovibrionaceae;Desulfovibrio;uncultured Desulfovibrio sp. | 91.4 | 1516 | -1 |
| metaRNA.PFemirge\_27811 | 4307 | 0.000524 | DQ115999.1.1504 | Bacteria;Firmicutes;Clostridia;Lachnospirales;Lachnospiraceae;uncultured;uncultured bacterium | 90.9 | 1504 | -1 |
| metaRNA.PFemirge\_2883 | 4205 | 0.000311 | CZBH01000001.999036.1000544 | Bacteria;Bacteroidota;Bacteroidia;Bacteroidales;Rikenellaceae;Alistipes;Alistipes sp. cv1 | 96.9 | 1503 | -1 |
| metaRNA.PFemirge\_8001 | 4186 | 0.000285 | LARD01000021.11.1513 | Bacteria;Cyanobacteria;Vampirivibrionia;Gastranaerophilales;Clostridium sp. K4410.MGS-306 | 94.9 | 1471 | -1 |
| metaRNA.PFemirge\_27715 | 4090 | 0.001416 | JX105679.1.1392 | Bacteria;Bacteroidota;Bacteroidia;Bacteroidales;Bacteroidaceae;Bacteroides;uncultured bacterium | 95.1 | 1311 | -1 |
| metaRNA.PFemirge\_2462 | 4069 | 0.000030 | DQ355176.1.1464 | Bacteria;Desulfobacterota;Desulfovibrionia;Desulfovibrionales;Desulfovibrionaceae;Desulfovibrio;uncultured Desulfovibrio sp. | 91.0 | 1464 | -1 |
| metaRNA.PFemirge\_17754 | 4058 | 0.000258 | EU503945.1.1397 | Bacteria;Firmicutes;Clostridia;Oscillospirales;Butyricicoccaceae;Butyricicoccus;uncultured bacterium | 91.8 | 1391 | -1 |
| metaRNA.PFemirge\_944 | 4032 | 0.000203 | GQ480147.1.1537 | Bacteria;Proteobacteria;Alphaproteobacteria;Holosporales;Holosporaceae;uncultured;uncultured bacterium | 99.0 | 1463 | -1 |
| metaRNA.PFemirge\_27667 | 3956 | 0.000530 | LT223609.1.1490 | Bacteria;Bacteroidota;Bacteroidia;Bacteroidales;Tannerellaceae;Parabacteroides;Parabacteroides faecis | 93.5 | 1487 | -1 |
| metaRNA.PFemirge\_17236 | 3897 | 0.000028 | EU457166.1.1396 | Bacteria;Bacteroidota;Bacteroidia;Bacteroidales;Bacteroidaceae;Bacteroides;uncultured bacterium | 93.2 | 1389 | -1 |
| metaRNA.PFemirge\_13684 | 3858 | 0.000421 | HQ789658.1.1441 | Bacteria;Bacteroidota;Bacteroidia;Bacteroidales;Bacteroidaceae;Bacteroides;uncultured organism | 91.3 | 1229 | -1 |
| metaRNA.PFemirge\_15803 | 3846 | 0.000040 | EU510692.1.1406 | Bacteria;Desulfobacterota;Desulfovibrionia;Desulfovibrionales;Desulfovibrionaceae;uncultured;uncultured bacterium | 89.0 | 1405 | -1 |
| metaRNA.PFemirge\_27787 | 3765 | 0.000299 | KP231747.1.1515 | Bacteria;Desulfobacterota;Desulfovibrionia;Desulfovibrionales;Desulfovibrionaceae;Desulfovibrio;uncultured Desulfovibrio sp. | 92.9 | 1517 | -1 |
| metaRNA.PFemirge\_19666 | 3745 | 0.000065 | DQ806118.1.1388 | Bacteria;Firmicutes;Clostridia;Lachnospirales;Lachnospiraceae;Lachnospiraceae UCG-008;uncultured bacterium | 91.7 | 1192 | -1 |
| metaRNA.PFemirge\_22312 | 3713 | 0.001807 | KP720604.1.1444 | Bacteria;Proteobacteria;Gammaproteobacteria;Enterobacterales;Enterobacteriaceae;Enterobacter;Enterobacter sp. IPC1-8 | 92.2 | 1314 | -1 |
| metaRNA.PFemirge\_16455 | 3689 | 0.000348 | DQ808516.1.1400 | Bacteria;Bacteroidota;Bacteroidia;Bacteroidales;Bacteroidaceae;Bacteroides;uncultured bacterium | 94.6 | 1396 | -1 |
| metaRNA.PFemirge\_23456 | 3646 | 0.000422 | EU766033.1.1357 | Bacteria;Bacteroidota;Bacteroidia;Bacteroidales;Bacteroidaceae;Bacteroides;uncultured bacterium | 93.2 | 1357 | -1 |
| metaRNA.PFemirge\_21658 | 3619 | 0.000263 | FJ881357.1.1492 | Bacteria;Firmicutes;Clostridia;Oscillospirales;Butyricicoccaceae;UCG-009;uncultured bacterium | 92.0 | 1288 | -1 |
| metaRNA.PFemirge\_13727 | 3496 | 0.000597 | CP012937.1474579.1476104 | Bacteria;Bacteroidota;Bacteroidia;Bacteroidales;Bacteroidaceae;Bacteroides;Bacteroides thetaiotaomicron | 93.2 | 1375 | -1 |
| metaRNA.PFemirge\_8071 | 3478 | 0.000339 | HQ810916.1.1454 | Bacteria;Firmicutes;Clostridia;Oscillospirales;Butyricicoccaceae;UCG-009;uncultured organism | 92.3 | 1368 | -1 |
| metaRNA.PFemirge\_13393 | 3352 | 0.000478 | DQ856503.1.1485 | Bacteria;Bacteroidota;Bacteroidia;Bacteroidales;Bacteroidaceae;Bacteroides;uncultured bacterium | 94.9 | 1362 | -1 |
| metaRNA.PFemirge\_7544 | 3317 | 0.000820 | AGES01000014.73256.74770 | Bacteria;Bacteroidota;Bacteroidia;Bacteroidales;Tannerellaceae;Parabacteroides;Parabacteroides sp. HGS0025 | 94.1 | 1482 | -1 |
| metaRNA.PFemirge\_7262 | 3263 | 0.000361 | GQ871713.1.1490 | Bacteria;Bacteroidota;Bacteroidia;Bacteroidales;Rikenellaceae;Alistipes;uncultured bacterium | 88.5 | 1432 | -1 |
| metaRNA.PFemirge\_28459 | 3068 | 0.000059 | KP231747.1.1515 | Bacteria;Desulfobacterota;Desulfovibrionia;Desulfovibrionales;Desulfovibrionaceae;Desulfovibrio;uncultured Desulfovibrio sp. | 93.6 | 1504 | -1 |
| metaRNA.PFemirge\_8289 | 3064 | 0.000288 | LT598589.1.1424 | Bacteria;Firmicutes;Clostridia;Oscillospirales;Butyricicoccaceae;UCG-008;Intestinibacillus massiliensis | 94.8 | 1357 | -1 |
| metaRNA.PFemirge\_23519 | 3033 | 0.000340 | KP944135.1.1430 | Bacteria;Bacteroidota;Bacteroidia;Bacteroidales;Bacteroidaceae;Bacteroides;Bacteroides fragilis | 88.4 | 1322 | -1 |
| metaRNA.PFemirge\_16889 | 3025 | 0.000942 | JX105679.1.1392 | Bacteria;Bacteroidota;Bacteroidia;Bacteroidales;Bacteroidaceae;Bacteroides;uncultured bacterium | 90.6 | 1402 | -1 |
| metaRNA.PFemirge\_12946 | 3011 | 0.000158 | DQ800642.1.1301 | Bacteria;Firmicutes;Clostridia;Christensenellales;Christensenellaceae;Christensenellaceae R-7 group;uncultured bacterium | 96.1 | 1293 | -1 |
| metaRNA.PFemirge\_1731 | 2986 | 0.000166 | JQ191011.1.1340 | Bacteria;Firmicutes;Bacilli;RF39;uncultured bacterium | 95.3 | 1282 | -1 |
| metaRNA.PFemirge\_10859 | 2961 | 0.000367 | ADLN01000129.202.1639 | Bacteria;Firmicutes;Clostridia;Lachnospirales;Lachnospiraceae;Incertae Sedis;Hungatella hathewayi WAL-18680 | 94.8 | 1394 | -1 |
| metaRNA.PFemirge\_17190 | 2938 | 0.000412 | DQ856503.1.1485 | Bacteria;Bacteroidota;Bacteroidia;Bacteroidales;Bacteroidaceae;Bacteroides;uncultured bacterium | 92.9 | 1396 | -1 |
| metaRNA.PFemirge\_24801 | 2889 | 0.000649 | HM630236.1.1451 | Bacteria;Bacteroidota;Bacteroidia;Bacteroidales;Rikenellaceae;dgA-11 gut group;uncultured bacterium | 90.7 | 1341 | -1 |
| metaRNA.PFemirge\_20782 | 2865 | 0.000343 | EU457153.1.1393 | Bacteria;Firmicutes;Clostridia;Oscillospirales;Ruminococcaceae;Anaerotruncus;uncultured bacterium | 91.7 | 1378 | -1 |
| metaRNA.PFemirge\_23133 | 2840 | 0.000160 | EU460205.1.1360 | Bacteria;Proteobacteria;Alphaproteobacteria;Rhodospirillales;uncultured;uncultured bacterium | 94.3 | 1358 | -1 |
| metaRNA.PFemirge\_16642 | 2822 | 0.000815 | EU136680.1.2019 | Bacteria;Bacteroidota;Bacteroidia;Bacteroidales;Bacteroidaceae;Bacteroides;Bacteroides uniformis | 91.8 | 1396 | -1 |
| metaRNA.PFemirge\_27765 | 2812 | 0.000112 | CP012937.757465.758990 | Bacteria;Bacteroidota;Bacteroidia;Bacteroidales;Bacteroidaceae;Bacteroides;Bacteroides thetaiotaomicron | 93.2 | 1399 | -1 |
| metaRNA.PFemirge\_12893 | 2782 | 0.000289 | AB218288.1.1449 | Bacteria;Bacteroidota;Bacteroidia;Bacteroidales;Tannerellaceae;Parabacteroides;uncultured Bacteroidaceae bacterium | 93.4 | 1331 | -1 |
| metaRNA.PFemirge\_15599 | 2780 | 0.001175 | GQ480147.1.1537 | Bacteria;Proteobacteria;Alphaproteobacteria;Holosporales;Holosporaceae;uncultured;uncultured bacterium | 86.1 | 1169 | -1 |
| metaRNA.PFemirge\_7720 | 2774 | 0.000296 | GQ898126.1.1466 | Bacteria;Proteobacteria;Alphaproteobacteria;Rhodospirillales;uncultured;uncultured bacterium | 91.6 | 1449 | -1 |
| metaRNA.PFemirge\_22655 | 2768 | 0.000210 | KX431291.1.1467 | Bacteria;Firmicutes;Clostridia;Peptostreptococcales-Tissierellales;Peptostreptococcaceae;Romboutsia;uncultured bacterium | 84.0 | 1454 | -1 |
| metaRNA.PFemirge\_23282 | 2743 | 0.000303 | GQ175524.1.1451 | Bacteria;Bacteroidota;Bacteroidia;Bacteroidales;Rikenellaceae;Rikenella;uncultured bacterium | 92.3 | 1359 | -1 |
| metaRNA.PFemirge\_27708 | 2716 | 0.002097 | DQ456246.1.1451 | Bacteria;Bacteroidota;Bacteroidia;Bacteroidales;Bacteroidaceae;Bacteroides;uncultured bacterium | 89.4 | 1294 | -1 |
| metaRNA.PFemirge\_27167 | 2672 | 0.000019 | HM778909.1.1458 | Bacteria;Proteobacteria;Gammaproteobacteria;Enterobacterales;Enterobacteriaceae;Plesiomonas;uncultured bacterium | 98.6 | 1117 | -1 |
| metaRNA.PFemirge\_7881 | 2660 | 0.000314 | FJ848388.1.1476 | Bacteria;Proteobacteria;Alphaproteobacteria;Rhodospirillales;uncultured;uncultured bacterium | 91.9 | 1403 | -1 |
| metaRNA.PFemirge\_10137 | 2627 | 0.000214 | KP231747.1.1515 | Bacteria;Desulfobacterota;Desulfovibrionia;Desulfovibrionales;Desulfovibrionaceae;Desulfovibrio;uncultured Desulfovibrio sp. | 94.2 | 1273 | -1 |
| metaRNA.PFemirge\_6977 | 2626 | 0.000305 | AB574482.1.1489 | Bacteria;Bacteroidota;Bacteroidia;Bacteroidales;Tannerellaceae;Parabacteroides;Parabacteroides chinchillae | 90.2 | 1484 | -1 |
| metaRNA.PFemirge\_9657 | 2624 | 0.000599 | HQ761737.1.1467 | Bacteria;Bacteroidota;Bacteroidia;Bacteroidales;Tannerellaceae;Parabacteroides;uncultured organism | 94.2 | 1461 | -1 |
| metaRNA.PFemirge\_4143 | 2532 | 0.000112 | AY457069.1.1483 | Bacteria;Actinobacteriota;Actinobacteria;Corynebacteriales;Mycobacteriaceae;Mycobacterium;Mycobacterium peregrinum | 99.9 | 1409 | -1 |
| metaRNA.PFemirge\_12447 | 2504 | 0.000311 | HM778328.1.1446 | Bacteria;Proteobacteria;Gammaproteobacteria;Enterobacterales;Enterobacteriaceae;Plesiomonas;uncultured bacterium | 97.8 | 1431 | -1 |
| metaRNA.PFemirge\_1575 | 2488 | 0.000184 | LN879502.2473726.2475252 | Bacteria;Verrucomicrobiota;Chlamydiae;Chlamydiales;Parachlamydiaceae;Candidatus Protochlamydia;Candidatus Protochlamydia naegleriophila | 96.4 | 1235 | -1 |
| metaRNA.PFemirge\_10437 | 2479 | 0.000692 | HQ775084.1.1453 | Bacteria;Bacteroidota;Bacteroidia;Bacteroidales;Barnesiellaceae;Coprobacter;uncultured organism | 90.2 | 1292 | -1 |
| metaRNA.PFemirge\_2779 | 2453 | 0.000359 | GQ175524.1.1451 | Bacteria;Bacteroidota;Bacteroidia;Bacteroidales;Rikenellaceae;Rikenella;uncultured bacterium | 93.2 | 1268 | -1 |
| metaRNA.PFemirge\_2636 | 2452 | 0.000336 | FWEL01000003.4055.5565 | Bacteria;Firmicutes;Clostridia;Oscillospirales;Oscillospiraceae;uncultured;Pseudoflavonifractor sp. Marseille-P3106 | 95.5 | 1495 | -1 |
| metaRNA.PFemirge\_9024 | 2425 | 0.000780 | DQ355176.1.1464 | Bacteria;Desulfobacterota;Desulfovibrionia;Desulfovibrionales;Desulfovibrionaceae;Desulfovibrio;uncultured Desulfovibrio sp. | 90.9 | 1386 | -1 |
| metaRNA.PFemirge\_13111 | 2418 | 0.000429 | FXDJ01000009.1407067.1408576 | Bacteria;Bacteroidota;Bacteroidia;Bacteroidales;Tannerellaceae;uncultured;Porphyromonadaceae bacterium MV12 | 90.2 | 1447 | -1 |
| metaRNA.PFemirge\_20555 | 2417 | 0.000182 | GQ448681.1.1394 | Bacteria;Bacteroidota;Bacteroidia;Bacteroidales;Marinifilaceae;Odoribacter;uncultured bacterium | 92.4 | 1389 | -1 |
| metaRNA.PFemirge\_3029 | 2402 | 0.000135 | AGZO01000045.56609.58123 | Bacteria;Bacteroidota;Bacteroidia;Bacteroidales;Tannerellaceae;Parabacteroides;Parabacteroides goldsteinii CL02T12C30 | 87.1 | 1498 | -1 |
| metaRNA.PFemirge\_9202 | 2379 | 0.000233 | HM779020.1.1471 | Bacteria;Proteobacteria;Gammaproteobacteria;Enterobacterales;Enterobacteriaceae;Plesiomonas;uncultured bacterium | 98.4 | 1469 | -1 |
| metaRNA.PFemirge\_1425 | 2366 | 0.000140 | AY957925.1.1536 | Bacteria;Proteobacteria;Gammaproteobacteria;Legionellales;Legionellaceae;Legionella;uncultured bacterium | 99.6 | 1498 | -1 |
| metaRNA.PFemirge\_26221 | 2342 | 0.000146 | DQ809316.1.1289 | Bacteria;Firmicutes;Clostridia;Oscillospirales;Ruminococcaceae;Subdoligranulum;uncultured bacterium | 93.9 | 1245 | -1 |
| metaRNA.PFemirge\_7151 | 2334 | 0.000042 | CP012937.5517779.5519304 | Bacteria;Bacteroidota;Bacteroidia;Bacteroidales;Bacteroidaceae;Bacteroides;Bacteroides thetaiotaomicron | 92.9 | 1487 | -1 |
| metaRNA.PFemirge\_11640 | 2315 | 0.000331 | JRGF01000044.3.1537 | Bacteria;Bacteroidota;Bacteroidia;Bacteroidales;Rikenellaceae;Alistipes;Alistipes inops | 94.8 | 1447 | -1 |
| metaRNA.PFemirge\_26086 | 2311 | 0.000005 | EU469747.1.1392 | Bacteria;Firmicutes;Clostridia;Lachnospirales;Lachnospiraceae;28-4;uncultured bacterium | 92.1 | 1290 | -1 |
| metaRNA.PFemirge\_6811 | 2307 | 0.000674 | MG428882.1.1489 | Bacteria;Bacteroidota;Bacteroidia;Bacteroidales;Bacteroidaceae;Bacteroides;Bacteroides paurosaccharolyticus | 92.5 | 1321 | -1 |
| metaRNA.PFemirge\_62 | 2304 | 0.000158 | HE575399.1.1900 | Eukaryota;Amorphea;Amoebozoa;Incertae Sedis;Ischnamoeba;uncultured Lobosea | 98.3 | 1531 | -1 |
| metaRNA.PFemirge\_19929 | 2282 | 0.000020 | EU508041.1.1387 | Bacteria;Firmicutes;Clostridia;Lachnospirales;Lachnospiraceae;Lachnospiraceae UCG-008;uncultured bacterium | 90.9 | 1305 | -1 |
| metaRNA.PFemirge\_569 | 2276 | 0.000450 | CP026007.74890.76443 | Bacteria;Firmicutes;Bacilli;Bacillales;Planococcaceae;Lysinibacillus;Lysinibacillus sp. YS11 | 86.4 | 1520 | -1 |
| metaRNA.PFemirge\_6039 | 2206 | 0.000506 | MG367100.1.1491 | Bacteria;Firmicutes;Clostridia;Oscillospirales;Oscillospiraceae;Colidextribacter;uncultured Clostridiales Family IV bacterium | 97.6 | 1482 | -1 |
| metaRNA.PFemirge\_28020 | 2195 | 0.000045 | DQ806118.1.1388 | Bacteria;Firmicutes;Clostridia;Lachnospirales;Lachnospiraceae;Lachnospiraceae UCG-008;uncultured bacterium | 93.0 | 1161 | -1 |
| metaRNA.PFemirge\_12583 | 2188 | 0.000152 | EU885024.1.1445 | Bacteria;Firmicutes;Clostridia;Oscillospirales;Ruminococcaceae;uncultured;uncultured bacterium | 94.9 | 1277 | -1 |
| metaRNA.PFemirge\_25652 | 2171 | 0.000166 | KP231747.1.1515 | Bacteria;Desulfobacterota;Desulfovibrionia;Desulfovibrionales;Desulfovibrionaceae;Desulfovibrio;uncultured Desulfovibrio sp. | 95.2 | 1195 | -1 |
| metaRNA.PFemirge\_15810 | 2161 | 0.000104 | EU776418.1.1406 | Bacteria;Firmicutes;Clostridia;Lachnospirales;Lachnospiraceae;Lachnospiraceae UCG-004;uncultured bacterium | 89.1 | 1405 | -1 |
| metaRNA.PFemirge\_7042 | 2155 | 0.000760 | AY693826.1.1487 | Bacteria;Bacteroidota;Bacteroidia;Bacteroidales;Rikenellaceae;uncultured;uncultured bacterium | 92.0 | 1407 | -1 |
| metaRNA.PFemirge\_1247 | 2153 | 0.000466 | AY554146.1.1441 | Bacteria;Desulfobacterota;Desulfovibrionia;Desulfovibrionales;Desulfovibrionaceae;Desulfovibrio;Desulfovibrio sp. LNB2 | 92.0 | 1439 | -1 |
| metaRNA.PFemirge\_16722 | 2151 | 0.000315 | GQ360012.1.1488 | Bacteria;Bacteroidota;Bacteroidia;Bacteroidales;Tannerellaceae;Macellibacteroides;uncultured bacterium | 89.3 | 1400 | -1 |
| metaRNA.PFemirge\_17368 | 2149 | 0.000291 | AB746664.1.1536 | Bacteria;Firmicutes;Clostridia;Lachnospirales;Lachnospiraceae;Lachnospiraceae NK4A136 group;uncultured bacterium | 93.6 | 1341 | -1 |
| metaRNA.PFemirge\_5324 | 2141 | 0.000152 | FJ374211.1.1495 | Bacteria;Firmicutes;Clostridia;Christensenellales;Christensenellaceae;uncultured;uncultured bacterium | 96.4 | 1485 | -1 |
| metaRNA.PFemirge\_4071 | 2138 | 0.000143 | JN391619.1.1506 | Bacteria;Proteobacteria;Gammaproteobacteria;Burkholderiales;Rhodocyclaceae;Thauera;uncultured bacterium | 94.1 | 1423 | -1 |
| metaRNA.PFemirge\_27933 | 2137 | 0.000300 | HQ760643.1.1454 | Bacteria;Bacteroidota;Bacteroidia;Bacteroidales;Bacteroidaceae;Bacteroides;uncultured organism | 92.4 | 1331 | -1 |
| metaRNA.PFemirge\_636 | 2124 | 0.000124 | CP008712.314129.315667 | Bacteria;Firmicutes;Bacilli;Bacillales;Bacillaceae;Bacillus;Bacillus cereus | 93.3 | 1509 | -1 |
| metaRNA.PFemirge\_10687 | 2103 | 0.000639 | FR749935.1.1522 | Bacteria;Firmicutes;Clostridia;Lachnospirales;Lachnospiraceae;[Eubacterium] fissicatena group;Faecalicatena fissicatena | 95.5 | 1389 | -1 |
| metaRNA.PFemirge\_27763 | 2080 | 0.002974 | DQ794241.1.1398 | Bacteria;Bacteroidota;Bacteroidia;Bacteroidales;Rikenellaceae;Alistipes;uncultured bacterium | 92.3 | 1394 | -1 |
| metaRNA.PFemirge\_4539 | 2077 | 0.000156 | KF840789.1.1401 | Bacteria;Cyanobacteria;Cyanobacteriia;Cyanobacteriales;Phormidiaceae;Arthrospira PCC-7345;Arthrospira platensis 540 | 99.8 | 1101 | -1 |
| metaRNA.PFemirge\_23679 | 2070 | 0.000081 | HQ784719.1.1452 | Bacteria;Bacteroidota;Bacteroidia;Bacteroidales;Bacteroidaceae;Bacteroides;uncultured organism | 93.7 | 1294 | -1 |
| metaRNA.PFemirge\_14604 | 2058 | 0.000176 | KP866212.1.1451 | Bacteria;Bacteroidota;Bacteroidia;Flavobacteriales;Flavobacteriaceae;Flavobacterium;Flavobacterium sp. HMF3121 | 96.9 | 1272 | -1 |
| metaRNA.PFemirge\_13295 | 2031 | 0.000130 | LT631507.1.1440 | Bacteria;Firmicutes;Clostridia;Peptostreptococcales-Tissierellales;Anaerovoracaceae;Anaerovorax;Bacilliculturomica massiliensis | 96.0 | 1367 | -1 |
| metaRNA.PFemirge\_5563 | 2012 | 0.000393 | FJ868767.1.1498 | Bacteria;Firmicutes;Clostridia;Lachnospirales;Lachnospiraceae;Lachnospiraceae UCG-008;uncultured bacterium | 92.5 | 1483 | -1 |
| metaRNA.PFemirge\_10734 | 1983 | 0.000127 | MH031289.1.1458 | Bacteria;Bacteroidota;Bacteroidia;Chitinophagales;Saprospiraceae;uncultured;uncultured Bacteroidetes bacterium | 94.2 | 1195 | -1 |
| metaRNA.PFemirge\_14379 | 1962 | 0.000010 | GU559813.1.1427 | Bacteria;Firmicutes;Negativicutes;Veillonellales-Selenomonadales;Sporomusaceae;Anaeromusa-Anaeroarcus;uncultured bacterium | 89.7 | 1403 | -1 |
| metaRNA.PFemirge\_17335 | 1951 | 0.001368 | FJ366372.1.1363 | Bacteria;Bacteroidota;Bacteroidia;Bacteroidales;Rikenellaceae;Alistipes;uncultured bacterium | 87.0 | 1367 | -1 |
| metaRNA.PFemirge\_27842 | 1947 | 0.000220 | FJ879941.1.1485 | Bacteria;Bacteroidota;Bacteroidia;Bacteroidales;Tannerellaceae;Parabacteroides;uncultured bacterium | 93.5 | 1481 | -1 |
| metaRNA.PFemirge\_2011 | 1925 | 0.000127 | MAIQ01000023.175.1690 | Bacteria;Firmicutes;Clostridia;Christensenellales;Christensenellaceae;Christensenella sp. AF73-05CM02 | 98.5 | 1490 | -1 |
| metaRNA.PFemirge\_8467 | 1921 | 0.000207 | EU158190.1.1510 | Bacteria;Firmicutes;Clostridia;Oscillospirales;Hydrogenoanaerobacterium;Hydrogenoanaerobacterium saccharovorans | 93.1 | 1310 | -1 |
| metaRNA.PFemirge\_16283 | 1896 | 0.000167 | LARG01000025.1.1402 | Bacteria;Bacteroidota;Bacteroidia;Bacteroidales;Rikenellaceae;Alistipes;Alistipes sp. N15.MGS-157 | 94.6 | 1395 | -1 |
| metaRNA.PFemirge\_22634 | 1867 | 0.000016 | LN870298.1.1495 | Bacteria;Firmicutes;Clostridia;Oscillospirales;Oscillospiraceae;Intestinimonas;Intestinimonas timonensis | 94.2 | 1290 | -1 |
| metaRNA.PFemirge\_9718 | 1857 | 0.000112 | AY571491.1.1466 | Bacteria;Proteobacteria;Alphaproteobacteria;Rickettsiales;uncultured;uncultured bacterium | 96.8 | 1317 | -1 |
| metaRNA.PFemirge\_2697 | 1855 | 0.000130 | JN559646.1.1500 | Bacteria;Bacteroidota;Bacteroidia;Bacteroidales;Prevotellaceae;Prevotellaceae Ga6A1 group;uncultured bacterium | 87.8 | 1397 | -1 |
| metaRNA.PFemirge\_6740 | 1846 | 0.000168 | JN038827.1.1489 | Bacteria;Proteobacteria;Gammaproteobacteria;Burkholderiales;Sutterellaceae;uncultured;uncultured beta proteobacterium | 98.1 | 1399 | -1 |
| metaRNA.PFemirge\_12524 | 1823 | 0.000118 | CP009057.2296841.2298384 | Bacteria;Bacteroidota;Bacteroidia;Bacteroidales;Bacteroidaceae;Bacteroides;Bacteroides dorei | 89.2 | 1270 | -1 |
| metaRNA.PFemirge\_4684 | 1801 | 0.000107 | KC253315.1.1470 | Bacteria;Bacteroidota;Bacteroidia;Cytophagales;Spirosomaceae;Runella;uncultured bacterium | 97.2 | 1426 | -1 |
| metaRNA.PFemirge\_23365 | 1767 | 0.000011 | FN665653.10808.12310 | Bacteria;Firmicutes;Clostridia;Peptostreptococcales-Tissierellales;Peptostreptococcaceae;Clostridioides;Clostridioides difficile M120 | 93.8 | 1360 | -1 |
| metaRNA.PFemirge\_6749 | 1735 | 0.000183 | ASTA01000042.17380.18894 | Bacteria;Firmicutes;Clostridia;Oscillospirales;Ruminococcaceae;Anaerotruncus;Anaerotruncus sp. G3(2012) | 94.3 | 1323 | -1 |
| metaRNA.PFemirge\_16208 | 1729 | 0.000135 | LC028597.1.1518 | Bacteria;Bacteroidota;Bacteroidia;Bacteroidales;Rikenellaceae;dgA-11 gut group;uncultured bacterium | 87.6 | 1240 | -1 |
| metaRNA.PFemirge\_23328 | 1713 | 0.000250 | AB470343.1.1485 | Bacteria;Bacteroidota;Bacteroidia;Bacteroidales;Tannerellaceae;Parabacteroides;Parabacteroides gordonii | 91.8 | 1359 | -1 |
| metaRNA.PFemirge\_10074 | 1683 | 0.000131 | EF667904.1.1463 | Bacteria;Bacteroidota;Bacteroidia;Cytophagales;Spirosomaceae;Emticicia;uncultured Bacteroidetes bacterium | 94.7 | 1292 | -1 |
| metaRNA.PFemirge\_16160 | 1646 | 0.000419 | AE015928.3034714.3036195 | Bacteria;Bacteroidota;Bacteroidia;Bacteroidales;Bacteroidaceae;Bacteroides;Bacteroides thetaiotaomicron VPI-5482 | 92.4 | 1398 | -1 |
| metaRNA.PFemirge\_13422 | 1644 | 0.000143 | MBVN01000043.446670.448353 | Eukaryota;Amorphea;Obazoa;Opisthokonta;Nucletmycea;Fungi;Dikarya;Ascomycota;Saccharomycotina;Saccharomycetes;Saccharomycetales;Phaffomycetaceae;Cyberlindnera-Candida clade;Saccharomyces cerevisiae (baker's yeast) | 99.8 | 888 | -1 |
| metaRNA.PFemirge\_15099 | 1634 | 0.000018 | DQ355176.1.1464 | Bacteria;Desulfobacterota;Desulfovibrionia;Desulfovibrionales;Desulfovibrionaceae;Desulfovibrio;uncultured Desulfovibrio sp. | 90.9 | 1391 | -1 |
| metaRNA.PFemirge\_1171 | 1628 | 0.000094 | AM743169.416435.417979 | Bacteria;Proteobacteria;Gammaproteobacteria;Xanthomonadales;Xanthomonadaceae;Stenotrophomonas;Stenotrophomonas maltophilia K279a | 98.7 | 1499 | -1 |
| metaRNA.PFemirge\_6963 | 1615 | 0.000087 | JX457277.1.1488 | Bacteria;Bacteroidota;Bacteroidia;Bacteroidales;Bacteroidaceae;Bacteroides;uncultured bacterium | 91.7 | 1426 | -1 |
| metaRNA.PFemirge\_7011 | 1606 | 0.000156 | LC194927.1.1488 | Bacteria;Firmicutes;Clostridia;Lachnospirales;Lachnospiraceae;Lachnospiraceae UCG-008;uncultured Clostridium sp. | 94.0 | 1416 | -1 |
| metaRNA.PFemirge\_21155 | 1589 | 0.000022 | HQ748426.1.1449 | Bacteria;Firmicutes;Clostridia;Lachnospirales;Lachnospiraceae;Lachnoclostridium;uncultured organism | 93.6 | 1297 | -1 |
| metaRNA.PFemirge\_16099 | 1577 | 0.000208 | EU778614.1.1403 | Bacteria;Bacteroidota;Bacteroidia;Bacteroidales;Rikenellaceae;Alistipes;uncultured bacterium | 95.1 | 1402 | -1 |
| metaRNA.PFemirge\_22821 | 1561 | 0.000029 | EU773857.1.1364 | Bacteria;Firmicutes;Clostridia;Peptostreptococcales-Tissierellales;Peptostreptococcaceae;uncultured;uncultured bacterium | 93.2 | 1348 | -1 |
| metaRNA.PFemirge\_9916 | 1546 | 0.000359 | CP012706.2989623.2991162 | Bacteria;Bacteroidota;Bacteroidia;Bacteroidales;Bacteroidaceae;Bacteroides;Bacteroides fragilis | 91.7 | 1441 | -1 |
| metaRNA.PFemirge\_23804 | 1541 | 0.000232 | AY916210.1.1353 | Bacteria;Bacteroidota;Bacteroidia;Bacteroidales;Bacteroidaceae;Bacteroides;uncultured bacterium | 96.5 | 1152 | -1 |
| metaRNA.PFemirge\_27699 | 1524 | 0.000031 | GQ898302.1.1489 | Bacteria;Bacteroidota;Bacteroidia;Bacteroidales;Bacteroidaceae;Bacteroides;uncultured bacterium | 94.1 | 1467 | -1 |
| metaRNA.PFemirge\_5205 | 1518 | 0.000169 | JQ923952.1.1496 | Bacteria;Proteobacteria;Gammaproteobacteria;Burkholderiales;Rhodocyclaceae;Candidatus Accumulibacter;uncultured bacterium | 96.2 | 1413 | -1 |
| metaRNA.PFemirge\_15322 | 1508 | 0.000011 | AQHW01000007.13.1527 | Bacteria;Bacteroidota;Bacteroidia;Bacteroidales;Tannerellaceae;Parabacteroides;Parabacteroides gordonii MS-1 | 87.3 | 1236 | -1 |
| metaRNA.PFemirge\_13064 | 1507 | 0.000029 | KP231747.1.1515 | Bacteria;Desulfobacterota;Desulfovibrionia;Desulfovibrionales;Desulfovibrionaceae;Desulfovibrio;uncultured Desulfovibrio sp. | 95.5 | 560 | -1 |
| metaRNA.PFemirge\_8305 | 1441 | 0.000107 | KT952808.1.1480 | Bacteria;Bacteroidota;Bacteroidia;Bacteroidales;Rs-E47 termite group;uncultured bacterium | 92.1 | 1283 | -1 |
| metaRNA.PFemirge\_18832 | 1437 | 0.000298 | JF160900.1.1352 | Bacteria;Firmicutes;Clostridia;Oscillospirales;Oscillospiraceae;uncultured;uncultured bacterium | 92.8 | 1354 | -1 |
| metaRNA.PFemirge\_12701 | 1430 | 0.000008 | KM651159.1.1495 | Bacteria;Firmicutes;Clostridia;Lachnospirales;Lachnospiraceae;Hungatella;uncultured Firmicutes bacterium | 92.2 | 1264 | -1 |
| metaRNA.PFemirge\_4739 | 1428 | 0.000130 | AQHX01000018.1891.3401 | Bacteria;Bacteroidota;Bacteroidia;Bacteroidales;Bacteroidaceae;Bacteroides;Bacteroides salyersiae WAL 10018 = DSM 18765 = JCM 12988 | 92.5 | 1499 | -1 |
| metaRNA.PFemirge\_1410 | 1386 | 0.000139 | FCNU01000016.57.1576 | Bacteria;Verrucomicrobiota;Chlamydiae;Chlamydiales;Parachlamydiaceae;Candidatus Protochlamydia;Parachlamydia sp. C2 | 94.7 | 1111 | -1 |
| metaRNA.PFemirge\_27722 | 1375 | 0.000401 | CP012937.5517779.5519304 | Bacteria;Bacteroidota;Bacteroidia;Bacteroidales;Bacteroidaceae;Bacteroides;Bacteroides thetaiotaomicron | 94.2 | 1388 | -1 |
| metaRNA.PFemirge\_23141 | 1373 | 0.000006 | EU472567.1.1360 | Bacteria;Firmicutes;Clostridia;Oscillospirales;Ruminococcaceae;Ruminococcus;uncultured bacterium | 91.2 | 1356 | -1 |
| metaRNA.PFemirge\_25869 | 1371 | 0.000016 | DQ824104.1.1398 | Bacteria;Firmicutes;Clostridia;Oscillospirales;Butyricicoccaceae;Butyricicoccus;uncultured bacterium | 90.9 | 1211 | -1 |
| metaRNA.PFemirge\_12928 | 1367 | 0.000016 | CP012938.2352114.2353639 | Bacteria;Bacteroidota;Bacteroidia;Bacteroidales;Bacteroidaceae;Bacteroides;Bacteroides ovatus | 92.5 | 1432 | -1 |
| metaRNA.PFemirge\_2964 | 1359 | 0.000728 | HG934468.2032323.2033833 | Bacteria;Bacteroidota;Bacteroidia;Bacteroidales;Rikenellaceae;Mucinivorans;Mucinivorans hirudinis | 89.1 | 1452 | -1 |
| metaRNA.PFemirge\_5603 | 1348 | 0.000219 | CBVB010000006.254456.255954 | Bacteria;Bacteroidota;Bacteroidia;Bacteroidales;Bacteroidaceae;Bacteroides;Bacteroidaceae bacterium MS4 | 93.6 | 1319 | -1 |
| metaRNA.PFemirge\_9252 | 1340 | 0.000096 | JN869169.1.1481 | Bacteria;Proteobacteria;Alphaproteobacteria;Reyranellales;Reyranellaceae;Reyranella;uncultured bacterium | 98.7 | 1403 | -1 |
| metaRNA.PFemirge\_2279 | 1334 | 0.000061 | AY554146.1.1441 | Bacteria;Desulfobacterota;Desulfovibrionia;Desulfovibrionales;Desulfovibrionaceae;Desulfovibrio;Desulfovibrio sp. LNB2 | 91.9 | 1253 | -1 |
| metaRNA.PFemirge\_9349 | 1304 | 0.000138 | AB614872.1.1469 | Bacteria;Verrucomicrobiota;Lentisphaeria;Victivallales;Victivallaceae;uncultured rumen bacterium | 94.5 | 1165 | -1 |
| metaRNA.PFemirge\_8584 | 1300 | 0.000080 | HQ807877.1.1477 | Bacteria;Firmicutes;Negativicutes;Veillonellales-Selenomonadales;Selenomonadaceae;uncultured;uncultured organism | 92.8 | 1414 | -1 |
| metaRNA.PFemirge\_16369 | 1294 | 0.000269 | JX457157.1.1486 | Bacteria;Bacteroidota;Bacteroidia;Bacteroidales;Tannerellaceae;Parabacteroides;uncultured bacterium | 95.0 | 1401 | -1 |
| metaRNA.PFemirge\_7857 | 1294 | 0.000091 | JN680664.1.1439 | Bacteria;Firmicutes;Clostridia;Peptostreptococcales-Tissierellales;Anaerovoracaceae;[Eubacterium] brachy group;uncultured Eubacteriaceae bacterium | 95.2 | 1390 | -1 |
| metaRNA.PFemirge\_16457 | 1282 | 0.000413 | DQ809030.1.1400 | Bacteria;Bacteroidota;Bacteroidia;Bacteroidales;Rikenellaceae;Alistipes;uncultured bacterium | 93.6 | 1399 | -1 |
| metaRNA.PFemirge\_9609 | 1273 | 0.000017 | CP022151.4897185.4898740 | Bacteria;Proteobacteria;Gammaproteobacteria;Enterobacterales;Enterobacteriaceae;Citrobacter;Citrobacter freundii | 94.4 | 1399 | -1 |
| metaRNA.PFemirge\_3684 | 1262 | 0.000086 | HQ697737.1.1504 | Bacteria;Proteobacteria;Gammaproteobacteria;Diplorickettsiales;Diplorickettsiaceae;uncultured;uncultured bacterium | 94.9 | 1393 | -1 |
| metaRNA.PFemirge\_22995 | 1258 | 0.000180 | GQ493239.1.1362 | Bacteria;Firmicutes;Bacilli;Erysipelotrichales;Erysipelatoclostridiaceae;Erysipelatoclostridium;uncultured bacterium | 92.5 | 1363 | -1 |
| metaRNA.PFemirge\_17584 | 1255 | 0.000104 | AY452004.1.1502 | Bacteria;Firmicutes;Clostridia;Lachnospirales;Lachnospiraceae;Lachnospiraceae UCG-004;uncultured bacterium | 89.3 | 1329 | -1 |
| metaRNA.PFemirge\_3699 | 1234 | 0.000078 | FPLS01017333.1.1416 | Bacteria;Proteobacteria;Gammaproteobacteria;Gammaproteobacteria Incertae Sedis;Unknown Family;metagenome | 93.7 | 1391 | -1 |
| metaRNA.PFemirge\_14707 | 1219 | 0.000621 | KP944118.1.1420 | Bacteria;Bacteroidota;Bacteroidia;Bacteroidales;Bacteroidaceae;Bacteroides;Bacteroides thetaiotaomicron | 94.2 | 1356 | -1 |
| metaRNA.PFemirge\_25780 | 1216 | 0.000272 | EU772837.1.1306 | Bacteria;Bacteroidota;Bacteroidia;Bacteroidales;Bacteroidaceae;Bacteroides;uncultured bacterium | 93.6 | 1306 | -1 |
| metaRNA.PFemirge\_4647 | 1207 | 0.000088 | KR029407.1.1499 | Bacteria;Proteobacteria;Gammaproteobacteria;Enterobacterales;Yersiniaceae;Serratia;Serratia sp. mixed culture X13-6 | 96.4 | 1407 | -1 |
| metaRNA.PFemirge\_12096 | 1205 | 0.000024 | DQ794241.1.1398 | Bacteria;Bacteroidota;Bacteroidia;Bacteroidales;Rikenellaceae;Alistipes;uncultured bacterium | 88.3 | 1245 | -1 |
| metaRNA.PFemirge\_27730 | 1199 | 0.000033 | DQ799104.1.1398 | Bacteria;Bacteroidota;Bacteroidia;Bacteroidales;Bacteroidaceae;Bacteroides;uncultured bacterium | 90.3 | 1397 | -1 |
| metaRNA.PFemirge\_27690 | 1194 | 0.000328 | FQXY01000005.366.1875 | Bacteria;Bacteroidota;Bacteroidia;Bacteroidales;Bacteroidaceae;Bacteroides;Bacteroides sp. Marseille-P3132 | 94.1 | 1398 | -1 |
| metaRNA.PFemirge\_7248 | 1189 | 0.000159 | HM630236.1.1451 | Bacteria;Bacteroidota;Bacteroidia;Bacteroidales;Rikenellaceae;dgA-11 gut group;uncultured bacterium | 90.1 | 1303 | -1 |
| metaRNA.PFemirge\_23442 | 1179 | 0.000004 | KM360180.1.1492 | Bacteria;Firmicutes;Clostridia;Lachnospirales;Lachnospiraceae;Lachnoclostridium;Clostridium sp. BPY5 | 91.6 | 1351 | -1 |
| metaRNA.PFemirge\_617 | 1168 | 0.000065 | MG754439.1.1392 | Bacteria;Proteobacteria;Alphaproteobacteria;Rhizobiales;Beijerinckiaceae;Bosea;Bosea sp. | 86.5 | 1552 | -1 |
| metaRNA.PFemirge\_16058 | 1147 | 0.000088 | DQ799601.1.1400 | Bacteria;Bacteroidota;Bacteroidia;Bacteroidales;Bacteroidaceae;Bacteroides;uncultured bacterium | 94.1 | 1221 | -1 |
| metaRNA.PFemirge\_6726 | 1142 | 0.000920 | CBVB010000006.254456.255954 | Bacteria;Bacteroidota;Bacteroidia;Bacteroidales;Bacteroidaceae;Bacteroides;Bacteroidaceae bacterium MS4 | 92.8 | 1437 | -1 |
| metaRNA.PFemirge\_14293 | 1139 | 0.000033 | AB214328.1.1484 | Bacteria;Bacteroidota;Bacteroidia;Bacteroidales;Bacteroidaceae;Bacteroides;Bacteroides intestinalis | 93.4 | 1405 | -1 |
| metaRNA.PFemirge\_15725 | 1109 | 0.000455 | FJ716048.1.1407 | Bacteria;Proteobacteria;Gammaproteobacteria;Enterobacterales;Enterobacteriaceae;Buttiauxella;uncultured bacterium | 97.2 | 1376 | -1 |
| metaRNA.PFemirge\_27660 | 1108 | 0.000081 | KP231747.1.1515 | Bacteria;Desulfobacterota;Desulfovibrionia;Desulfovibrionales;Desulfovibrionaceae;Desulfovibrio;uncultured Desulfovibrio sp. | 93.9 | 1444 | -1 |
| metaRNA.PFemirge\_19251 | 1088 | 0.000448 | DQ800695.1.1386 | Bacteria;Firmicutes;Clostridia;Lachnospirales;Lachnospiraceae;Lachnoclostridium;uncultured bacterium | 92.5 | 1389 | -1 |
| metaRNA.PFemirge\_25365 | 1084 | 0.000174 | AB934800.1.1322 | Bacteria;Proteobacteria;Alphaproteobacteria;Rhodospirillales;uncultured;uncultured bacterium | 94.5 | 1322 | -1 |
| metaRNA.PFemirge\_28223 | 1071 | 0.009837 | KP231747.1.1515 | Bacteria;Desulfobacterota;Desulfovibrionia;Desulfovibrionales;Desulfovibrionaceae;Desulfovibrio;uncultured Desulfovibrio sp. | 91.3 | 1406 | -1 |
| metaRNA.PFemirge\_13159 | 1063 | 0.000108 | HQ681866.1.1451 | Bacteria;Bacteroidota;Bacteroidia;Bacteroidales;Bacteroidaceae;Bacteroides;uncultured bacterium | 92.6 | 1451 | -1 |
| metaRNA.PFemirge\_4783 | 1056 | 0.000126 | JQ820153.1.1498 | Bacteria;Firmicutes;Clostridia;Oscillospirales;Oscillospiraceae;uncultured;uncultured bacterium | 90.2 | 1465 | -1 |
| metaRNA.PFemirge\_21111 | 1042 | 0.000149 | EU467709.1.1380 | Bacteria;Bacteroidota;Bacteroidia;Bacteroidales;Bacteroidaceae;Bacteroides;uncultured bacterium | 92.0 | 1383 | -1 |
| metaRNA.PFemirge\_16552 | 1016 | 0.000664 | JX105679.1.1392 | Bacteria;Bacteroidota;Bacteroidia;Bacteroidales;Bacteroidaceae;Bacteroides;uncultured bacterium | 88.7 | 1371 | -1 |
| metaRNA.PFemirge\_19515 | 1014 | 0.000091 | EU776450.1.1387 | Bacteria;Firmicutes;Clostridia;Oscillospirales;Ruminococcaceae;Ruminococcus;uncultured bacterium | 92.3 | 1386 | -1 |
| metaRNA.PFemirge\_22896 | 1012 | 0.000249 | DQ800679.1.1366 | Bacteria;Firmicutes;Clostridia;Lachnospirales;Lachnospiraceae;[Ruminococcus] gauvreauii group;uncultured bacterium | 93.8 | 1363 | -1 |
| metaRNA.PFemirge\_15229 | 1006 | 0.000039 | CP026122.3396659.3398209 | Bacteria;Proteobacteria;Gammaproteobacteria;Enterobacterales;Aeromonadaceae;Aeromonas;Aeromonas sp. ASNIH5 | 92.5 | 1388 | -1 |
| metaRNA.PFemirge\_13642 | 1006 | 0.000107 | AM183080.1.1438 | Bacteria;Firmicutes;Clostridia;Clostridiales;Clostridiaceae;Clostridium sensu stricto 1;uncultured bacterium | 97.3 | 1316 | -1 |
| metaRNA.PFemirge\_16699 | 993 | 0.000110 | DQ824296.1.1398 | Bacteria;Bacteroidota;Bacteroidia;Bacteroidales;Bacteroidaceae;Bacteroides;uncultured bacterium | 87.9 | 1284 | -1 |
| metaRNA.PFemirge\_27721 | 986 | 0.000096 | CP012937.1474579.1476104 | Bacteria;Bacteroidota;Bacteroidia;Bacteroidales;Bacteroidaceae;Bacteroides;Bacteroides thetaiotaomicron | 92.7 | 1391 | -1 |
| metaRNA.PFemirge\_9884 | 986 | 0.000063 | JX006733.1.1426 | Bacteria;Desulfobacterota;Desulfovibrionia;Desulfovibrionales;Desulfovibrionaceae;Bilophila;bacterium NLAE-zl-H528 | 90.8 | 1219 | -1 |
| metaRNA.PFemirge\_27935 | 973 | 0.000081 | KP231747.1.1515 | Bacteria;Desulfobacterota;Desulfovibrionia;Desulfovibrionales;Desulfovibrionaceae;Desulfovibrio;uncultured Desulfovibrio sp. | 91.6 | 1411 | -1 |
| metaRNA.PFemirge\_8631 | 968 | 0.000216 | KX823723.1.1477 | Bacteria;Firmicutes;Clostridia;Oscillospirales;Oscillospiraceae;NK4A214 group;uncultured bacterium | 91.4 | 1302 | -1 |
| metaRNA.PFemirge\_8530 | 967 | 0.000070 | LN995859.1.1478 | Bacteria;Verrucomicrobiota;Chlamydiae;Chlamydiales;Parachlamydiaceae;uncultured;Neochlamydia sp. Trut23-12-2015\_Venoge-Embouchure | 97.5 | 1379 | -1 |
| metaRNA.PFemirge\_26268 | 953 | 0.000097 | HQ815291.1.1415 | Bacteria;Firmicutes;Clostridia;Lachnospirales;Lachnospiraceae;[Ruminococcus] gauvreauii group;uncultured organism | 94.4 | 1288 | -1 |
| metaRNA.PFemirge\_26903 | 949 | 0.000064 | EU777108.1.1328 | Bacteria;Firmicutes;Clostridia;Oscillospirales;Ruminococcaceae;Incertae Sedis;uncultured bacterium | 92.9 | 1261 | -1 |
| metaRNA.PFemirge\_6899 | 947 | 0.000172 | AGZO01000045.56609.58123 | Bacteria;Bacteroidota;Bacteroidia;Bacteroidales;Tannerellaceae;Parabacteroides;Parabacteroides goldsteinii CL02T12C30 | 91.5 | 1490 | -1 |
| metaRNA.PFemirge\_24960 | 946 | 0.000086 | LT725663.1.1450 | Bacteria;Bacteroidota;Bacteroidia;Bacteroidales;Tannerellaceae;Parabacteroides;Parabacteroides sp. Marseille-P3668 | 94.0 | 1235 | -1 |
| metaRNA.PFemirge\_17143 | 941 | 0.000013 | GU179922.1.1487 | Bacteria;Firmicutes;Clostridia;Clostridia UCG-014;uncultured Firmicutes bacterium | 93.3 | 1149 | -1 |
| metaRNA.PFemirge\_28093 | 925 | 0.000089 | AB574482.1.1489 | Bacteria;Bacteroidota;Bacteroidia;Bacteroidales;Tannerellaceae;Parabacteroides;Parabacteroides chinchillae | 93.8 | 1497 | -1 |
| metaRNA.PFemirge\_23054 | 923 | 0.000108 | HQ792908.1.1451 | Bacteria;Firmicutes;Clostridia;Oscillospirales;Butyricicoccaceae;UCG-008;uncultured organism | 91.5 | 1103 | -1 |
| metaRNA.PFemirge\_28001 | 920 | 0.000400 | EU766155.1.1353 | Bacteria;Bacteroidota;Bacteroidia;Bacteroidales;Bacteroidaceae;Bacteroides;uncultured bacterium | 94.8 | 1215 | -1 |
| metaRNA.PFemirge\_23292 | 916 | 0.000017 | CP011307.631177.632643 | Bacteria;Firmicutes;Clostridia;Oscillospirales;Oscillospiraceae;Intestinimonas;Intestinimonas butyriciproducens | 94.9 | 1295 | -1 |
| metaRNA.PFemirge\_14297 | 914 | 0.000030 | GBID01005713.1.1428 | Bacteria;Bacteroidota;Bacteroidia;Bacteroidales;Bacteroidaceae;Bacteroides;Blattella germanica (German cockroach) | 90.6 | 1187 | -1 |
| metaRNA.PFemirge\_13989 | 905 | 0.000254 | HQ761491.1.1432 | Bacteria;Firmicutes;Clostridia;Lachnospirales;Lachnospiraceae;Lachnospiraceae AC2044 group;uncultured organism | 92.1 | 1339 | -1 |
| metaRNA.PFemirge\_26639 | 884 | 0.000029 | HQ697694.1.1452 | Bacteria;Proteobacteria;Alphaproteobacteria;Rhodospirillales;Magnetospirillaceae;Magnetospirillum;uncultured bacterium | 90.0 | 1200 | -1 |
| metaRNA.PFemirge\_3501 | 878 | 0.000070 | GDHD01010040.119.1623 | Bacteria;Cyanobacteria;Cyanobacteriia;Chloroplast;Cymbidium faberi | 97.0 | 1475 | -1 |
| metaRNA.PFemirge\_4338 | 873 | 0.000074 | HQ791782.1.1436 | Bacteria;Bacteroidota;Bacteroidia;Bacteroidales;Bacteroidaceae;Bacteroides;uncultured organism | 88.7 | 1266 | -1 |
| metaRNA.PFemirge\_4651 | 844 | 0.000048 | KU978243.1.1517 | Bacteria;Proteobacteria;Gammaproteobacteria;Pseudomonadales;Pseudomonadaceae;Pseudomonas;uncultured bacterium | 95.9 | 1491 | -1 |
| metaRNA.PFemirge\_25011 | 840 | 0.000022 | GQ448965.1.1384 | Bacteria;Bacteroidota;Bacteroidia;Bacteroidales;Bacteroidaceae;Bacteroides;uncultured bacterium | 92.7 | 1199 | -1 |
| metaRNA.PFemirge\_17373 | 830 | 0.000091 | KF318251.1.1396 | Bacteria;Firmicutes;Clostridia;Oscillospirales;Ruminococcaceae;uncultured;uncultured Firmicutes bacterium | 96.6 | 1314 | -1 |
| metaRNA.PFemirge\_25883 | 827 | 0.000009 | MG321612.1.1372 | Bacteria;Bacteroidota;Bacteroidia;Bacteroidales;Tannerellaceae;Parabacteroides;Parabacteroides sp. | 88.5 | 1221 | -1 |
| metaRNA.PFemirge\_12905 | 819 | 0.000056 | HQ778039.1.1464 | Bacteria;Bacteroidota;Bacteroidia;Bacteroidales;Barnesiellaceae;Coprobacter;uncultured organism | 90.1 | 1305 | -1 |
| metaRNA.PFemirge\_11223 | 818 | 0.000214 | JN032891.1.1470 | Bacteria;Desulfobacterota;Desulfovibrionia;Desulfovibrionales;Desulfovibrionaceae;Desulfovibrio;uncultured bacterium | 91.4 | 1320 | -1 |
| metaRNA.PFemirge\_7296 | 814 | 0.000138 | EF403711.1.1488 | Bacteria;Bacteroidota;Bacteroidia;Bacteroidales;Bacteroidaceae;Bacteroides;uncultured bacterium | 90.5 | 1480 | -1 |
| metaRNA.PFemirge\_19176 | 813 | 0.000253 | JX198651.1.1513 | Bacteria;Firmicutes;Clostridia;Oscillospirales;Butyricicoccaceae;UCG-009;uncultured bacterium | 92.2 | 1383 | -1 |
| metaRNA.PFemirge\_23715 | 812 | 0.000282 | JX105679.1.1392 | Bacteria;Bacteroidota;Bacteroidia;Bacteroidales;Bacteroidaceae;Bacteroides;uncultured bacterium | 90.2 | 1310 | -1 |
| metaRNA.PFemirge\_14552 | 807 | 0.000054 | DQ676995.1.1498 | Bacteria;Actinobacteriota;Actinobacteria;Propionibacteriales;Propionibacteriaceae;Propionicicella;iron-reducing enrichment clone Cl-A3 | 98.3 | 1437 | -1 |
| metaRNA.PFemirge\_10528 | 806 | 0.000144 | GQ360012.1.1488 | Bacteria;Bacteroidota;Bacteroidia;Bacteroidales;Tannerellaceae;Macellibacteroides;uncultured bacterium | 87.4 | 1325 | -1 |
| metaRNA.PFemirge\_9685 | 800 | 0.000101 | KT952740.1.1467 | Bacteria;Verrucomicrobiota;Kiritimatiellae;Kiritimatiellales;Kiritimatiellaceae;uncultured;uncultured bacterium | 95.6 | 1367 | -1 |
| metaRNA.PFemirge\_4206 | 795 | 0.000185 | GQ503874.1.1501 | Bacteria;Desulfobacterota;Desulfovibrionia;Desulfovibrionales;Desulfovibrionaceae;Desulfovibrio;bacterium enrichment culture clone Ecwsrb037 | 91.4 | 1399 | -1 |
| metaRNA.PFemirge\_23221 | 794 | 0.000003 | HQ783986.1.1437 | Bacteria;Bacteroidota;Bacteroidia;Bacteroidales;Tannerellaceae;Parabacteroides;uncultured organism | 92.4 | 1362 | -1 |
| metaRNA.PFemirge\_14141 | 794 | 0.000050 | DQ905881.1.1578 | Bacteria;Bacteroidota;Bacteroidia;Bacteroidales;Bacteroidaceae;Bacteroides;uncultured bacterium | 92.3 | 1362 | -1 |
| metaRNA.PFemirge\_511 | 786 | 0.000319 | AQHW01000007.13.1527 | Bacteria;Bacteroidota;Bacteroidia;Bacteroidales;Tannerellaceae;Parabacteroides;Parabacteroides gordonii MS-1 | 78.0 | 1530 | -1 |
| metaRNA.PFemirge\_22812 | 781 | 0.000073 | DQ800679.1.1366 | Bacteria;Firmicutes;Clostridia;Lachnospirales;Lachnospiraceae;[Ruminococcus] gauvreauii group;uncultured bacterium | 93.0 | 1360 | -1 |
| metaRNA.PFemirge\_21386 | 775 | 0.000316 | DQ800244.1.1377 | Bacteria;Firmicutes;Clostridia;Oscillospirales;Ruminococcaceae;Faecalibacterium;uncultured bacterium | 92.3 | 1370 | -1 |
| metaRNA.PFemirge\_5633 | 773 | 0.000088 | CXSE01001446.1371.2879 | Bacteria;Bacteroidota;Bacteroidia;Bacteroidales;Bacteroidaceae;Bacteroides;human gut metagenome | 93.7 | 1401 | -1 |
| metaRNA.PFemirge\_14145 | 772 | 0.000012 | HQ813540.1.1430 | Bacteria;Bacteroidota;Bacteroidia;Bacteroidales;Tannerellaceae;Parabacteroides;uncultured organism | 93.8 | 1425 | -1 |
| metaRNA.PFemirge\_7265 | 765 | 0.000032 | AB238927.1.1486 | Bacteria;Bacteroidota;Bacteroidia;Bacteroidales;Tannerellaceae;Parabacteroides;Parabacteroides distasonis | 85.7 | 1473 | -1 |
| metaRNA.PFemirge\_4924 | 761 | 0.000568 | GQ360030.1.1497 | Bacteria;Proteobacteria;Gammaproteobacteria;Enterobacterales;Aeromonadaceae;Aeromonas;uncultured bacterium | 97.0 | 1495 | -1 |
| metaRNA.PFemirge\_27719 | 757 | 0.000142 | DQ456435.1.1452 | Bacteria;Bacteroidota;Bacteroidia;Bacteroidales;Bacteroidaceae;Bacteroides;uncultured bacterium | 93.2 | 1391 | -1 |
| metaRNA.PFemirge\_23832 | 754 | 0.000475 | EU766155.1.1353 | Bacteria;Bacteroidota;Bacteroidia;Bacteroidales;Bacteroidaceae;Bacteroides;uncultured bacterium | 95.7 | 1353 | -1 |
| metaRNA.PFemirge\_27887 | 747 | 0.000076 | HQ816006.1.1448 | Bacteria;Bacteroidota;Bacteroidia;Bacteroidales;Bacteroidaceae;Bacteroides;uncultured organism | 90.4 | 1390 | -1 |
| metaRNA.PFemirge\_9441 | 746 | 0.000053 | CP022655.4386843.4388399 | Bacteria;Firmicutes;Bacilli;Paenibacillales;Paenibacillaceae;Paenibacillus;Paenibacillus sp. RUD330 | 93.0 | 1422 | -1 |
| metaRNA.PFemirge\_16640 | 739 | 0.000098 | DQ793316.1.1393 | Bacteria;Bacteroidota;Bacteroidia;Bacteroidales;Tannerellaceae;Parabacteroides;uncultured bacterium | 93.4 | 1375 | -1 |
| metaRNA.PFemirge\_23310 | 733 | 0.000097 | DQ809036.1.1400 | Bacteria;Bacteroidota;Bacteroidia;Bacteroidales;Bacteroidaceae;Bacteroides;uncultured bacterium | 91.5 | 1208 | -1 |
| metaRNA.PFemirge\_22441 | 727 | 0.000050 | MH699149.1.1471 | Bacteria;Actinobacteriota;Actinobacteria;Micrococcales;Microbacteriaceae;Microbacterium;Mycobacterium sp. | 95.4 | 1287 | -1 |
| metaRNA.PFemirge\_22890 | 726 | 0.000077 | HQ819238.1.1451 | Bacteria;Bacteroidota;Bacteroidia;Bacteroidales;Bacteroidaceae;Bacteroides;uncultured organism | 93.7 | 1149 | -1 |
| metaRNA.PFemirge\_6134 | 725 | 0.000063 | JX457179.1.1492 | Bacteria;Firmicutes;Bacilli;Erysipelotrichales;Erysipelatoclostridiaceae;Erysipelatoclostridium;uncultured bacterium | 91.9 | 1413 | -1 |
| metaRNA.PFemirge\_11224 | 721 | 0.000318 | LT725663.1.1450 | Bacteria;Bacteroidota;Bacteroidia;Bacteroidales;Tannerellaceae;Parabacteroides;Parabacteroides sp. Marseille-P3668 | 91.0 | 1395 | -1 |
| metaRNA.PFemirge\_12625 | 703 | 0.000154 | CXSE01001446.1371.2879 | Bacteria;Bacteroidota;Bacteroidia;Bacteroidales;Bacteroidaceae;Bacteroides;human gut metagenome | 94.4 | 1330 | -1 |
| metaRNA.PFemirge\_16690 | 699 | 0.000006 | AY916206.1.1387 | Bacteria;Desulfobacterota;Desulfovibrionia;Desulfovibrionales;Desulfovibrionaceae;uncultured;uncultured bacterium | 91.2 | 1255 | -1 |
| metaRNA.PFemirge\_28188 | 698 | 0.000023 | FR749935.1.1522 | Bacteria;Firmicutes;Clostridia;Lachnospirales;Lachnospiraceae;[Eubacterium] fissicatena group;Faecalicatena fissicatena | 91.0 | 1367 | -1 |
| metaRNA.PFemirge\_3393 | 695 | 0.000032 | AGES01000014.73256.74770 | Bacteria;Bacteroidota;Bacteroidia;Bacteroidales;Tannerellaceae;Parabacteroides;Parabacteroides sp. HGS0025 | 92.2 | 1492 | -1 |
| metaRNA.PFemirge\_11903 | 694 | 0.000099 | CU926683.1.1355 | Bacteria;Bacteroidota;Bacteroidia;Bacteroidales;Tannerellaceae;Macellibacteroides;uncultured bacterium | 90.3 | 1291 | -1 |
| metaRNA.PFemirge\_30234 | 688 | 0.000078 | CP012706.2989623.2991162 | Bacteria;Bacteroidota;Bacteroidia;Bacteroidales;Bacteroidaceae;Bacteroides;Bacteroides fragilis | 90.0 | 1443 | -1 |
| metaRNA.PFemirge\_22396 | 688 | 0.000054 | DQ797995.1.1380 | Bacteria;Firmicutes;Clostridia;Oscillospirales;Oscillospiraceae;uncultured;uncultured bacterium | 90.3 | 1367 | -1 |
| metaRNA.PFemirge\_22505 | 678 | 0.000009 | AB510710.1.1461 | Bacteria;Bacteroidota;Bacteroidia;Bacteroidales;Bacteroidaceae;Bacteroides;Bacteroides thetaiotaomicron | 93.2 | 1211 | -1 |
| metaRNA.PFemirge\_11662 | 676 | 0.000003 | GQ360012.1.1488 | Bacteria;Bacteroidota;Bacteroidia;Bacteroidales;Tannerellaceae;Macellibacteroides;uncultured bacterium | 88.7 | 1390 | -1 |
| metaRNA.PFemirge\_22496 | 672 | 0.000324 | EU772885.1.1375 | Bacteria;Firmicutes;Clostridia;Lachnospirales;Lachnospiraceae;Agathobacter;uncultured bacterium | 89.9 | 1382 | -1 |
| metaRNA.PFemirge\_4201 | 667 | 0.000063 | GQ389185.1.1501 | Bacteria;Proteobacteria;Gammaproteobacteria;Pseudomonadales;Pseudomonadaceae;Pseudomonas;uncultured bacterium | 97.9 | 1400 | -1 |
| metaRNA.PFemirge\_16812 | 659 | 0.000006 | DQ825005.1.1403 | Bacteria;Bacteroidota;Bacteroidia;Bacteroidales;Bacteroidaceae;Bacteroides;uncultured bacterium | 91.7 | 1391 | -1 |
| metaRNA.PFemirge\_20664 | 657 | 0.000055 | LC094617.1.1418 | Bacteria;Actinobacteriota;Actinobacteria;Micrococcales;Microbacteriaceae;Leifsonia;Microbacteriaceae bacterium MAEY21 | 89.8 | 1293 | -1 |
| metaRNA.PFemirge\_17293 | 656 | 0.000099 | EU773321.1.1396 | Bacteria;Firmicutes;Clostridia;Oscillospirales;Oscillospiraceae;uncultured;uncultured bacterium | 93.3 | 1380 | -1 |
| metaRNA.PFemirge\_18963 | 654 | 0.000130 | CP011307.631177.632643 | Bacteria;Firmicutes;Clostridia;Oscillospirales;Oscillospiraceae;Intestinimonas;Intestinimonas butyriciproducens | 91.0 | 1176 | -1 |
| metaRNA.PFemirge\_1573 | 651 | 0.000053 | FPLP01011680.8.1529 | Bacteria;Proteobacteria;Gammaproteobacteria;Enterobacterales;Aeromonadaceae;Aeromonas;metagenome | 98.8 | 1496 | -1 |
| metaRNA.PFemirge\_12198 | 648 | 0.000070 | HQ761612.1.1444 | Bacteria;Bacteroidota;Bacteroidia;Bacteroidales;Tannerellaceae;Parabacteroides;uncultured organism | 90.3 | 1438 | -1 |
| metaRNA.PFemirge\_27670 | 648 | 0.000071 | ABVO01000045.3575.5089 | Bacteria;Bacteroidota;Bacteroidia;Bacteroidales;Bacteroidaceae;Bacteroides;Bacteroides eggerthii DSM 20697 | 92.7 | 1402 | -1 |
| metaRNA.PFemirge\_9241 | 644 | 0.000060 | AB612398.1.1470 | Bacteria;Verrucomicrobiota;Lentisphaeria;Victivallales;Victivallaceae;uncultured rumen bacterium | 94.6 | 1298 | -1 |
| metaRNA.PFemirge\_6691 | 643 | 0.000004 | FQSE01000007.1631802.1633308 | Bacteria;Bacteroidota;Bacteroidia;Bacteroidales;Tannerellaceae;Parabacteroides;Parabacteroides sp. Marseille-P3136 | 89.7 | 1481 | -1 |
| metaRNA.PFemirge\_11838 | 643 | 0.000075 | EU776216.1.1354 | Bacteria;Firmicutes;Clostridia;Lachnospirales;Lachnospiraceae;uncultured;uncultured bacterium | 92.6 | 1295 | -1 |
| metaRNA.PFemirge\_28106 | 638 | 0.000113 | GQ898343.1.1489 | Bacteria;Bacteroidota;Bacteroidia;Bacteroidales;Bacteroidaceae;Bacteroides;uncultured bacterium | 93.9 | 1484 | -1 |
| metaRNA.PFemirge\_12202 | 637 | 0.000068 | AB176219.1.1449 | Bacteria;Proteobacteria;Alphaproteobacteria;Sphingomonadales;Sphingomonadaceae;Novosphingobium;uncultured bacterium | 94.0 | 1291 | -1 |
| metaRNA.PFemirge\_26525 | 635 | 0.000032 | KP231747.1.1515 | Bacteria;Desulfobacterota;Desulfovibrionia;Desulfovibrionales;Desulfovibrionaceae;Desulfovibrio;uncultured Desulfovibrio sp. | 92.9 | 1207 | -1 |
| metaRNA.PFemirge\_28266 | 628 | 0.000003 | JQ083789.1.1511 | Bacteria;Desulfobacterota;Desulfovibrionia;Desulfovibrionales;Desulfovibrionaceae;Bilophila;uncultured bacterium | 91.1 | 1463 | -1 |
| metaRNA.PFemirge\_30232 | 627 | 0.000056 | AB574482.1.1489 | Bacteria;Bacteroidota;Bacteroidia;Bacteroidales;Tannerellaceae;Parabacteroides;Parabacteroides chinchillae | 86.7 | 1491 | -1 |
| metaRNA.PFemirge\_30259 | 611 | 0.000380 | MG367100.1.1491 | Bacteria;Firmicutes;Clostridia;Oscillospirales;Oscillospiraceae;Colidextribacter;uncultured Clostridiales Family IV bacterium | 90.9 | 1335 | -1 |
| metaRNA.PFemirge\_5654 | 609 | 0.000047 | AGZO01000045.56609.58123 | Bacteria;Bacteroidota;Bacteroidia;Bacteroidales;Tannerellaceae;Parabacteroides;Parabacteroides goldsteinii CL02T12C30 | 88.5 | 1450 | -1 |
| metaRNA.PFemirge\_6703 | 606 | 0.000042 | FP929047.78761.80257 | Bacteria;Actinobacteriota;Coriobacteriia;Coriobacteriales;Eggerthellaceae;Gordonibacter;Gordonibacter pamelaeae 7-10-1-b | 96.0 | 1384 | -1 |
| metaRNA.PFemirge\_17589 | 602 | 0.000056 | JN792358.1.1395 | Bacteria;Firmicutes;Clostridia;Peptostreptococcales-Tissierellales;Peptostreptococcaceae;Paraclostridium;uncultured Clostridium sp. | 94.8 | 1325 | -1 |
| metaRNA.PFemirge\_14692 | 601 | 0.000045 | HQ761439.1.1422 | Bacteria;Firmicutes;Clostridia;Clostridia vadinBB60 group;uncultured organism | 97.3 | 1283 | -1 |
| metaRNA.PFemirge\_10478 | 589 | 0.000063 | AGES01000014.73256.74770 | Bacteria;Bacteroidota;Bacteroidia;Bacteroidales;Tannerellaceae;Parabacteroides;Parabacteroides sp. HGS0025 | 92.9 | 1459 | -1 |
| metaRNA.PFemirge\_3472 | 584 | 0.000036 | EU939391.1.1539 | Bacteria;Firmicutes;Clostridia;Eubacteriales;Eubacteriaceae;Eubacterium;uncultured bacterium | 94.3 | 1413 | -1 |
| metaRNA.PFemirge\_24325 | 583 | 0.000037 | HAFU01069247.55.1411 | Bacteria;Proteobacteria;Gammaproteobacteria;Pseudomonadales;Pseudomonadaceae;Pseudomonas;Spumella-like flagellate JBL14 | 94.6 | 1343 | -1 |
| metaRNA.PFemirge\_16370 | 581 | 0.000246 | CP014030.1244709.1246258 | Bacteria;Proteobacteria;Gammaproteobacteria;Enterobacterales;Enterobacteriaceae;Citrobacter;Citrobacter sp. FDAARGOS\_156 | 94.5 | 1398 | -1 |
| metaRNA.PFemirge\_1728 | 581 | 0.000036 | KP231747.1.1515 | Bacteria;Desulfobacterota;Desulfovibrionia;Desulfovibrionales;Desulfovibrionaceae;Desulfovibrio;uncultured Desulfovibrio sp. | 92.8 | 1514 | -1 |
| metaRNA.PFemirge\_5628 | 579 | 0.000045 | JN391820.1.1494 | Bacteria;Proteobacteria;Gammaproteobacteria;Burkholderiales;Comamonadaceae;Sphaerotilus;uncultured bacterium | 97.1 | 1467 | -1 |
| metaRNA.PFemirge\_6685 | 577 | 0.000147 | EU663601.1.1489 | Bacteria;Bacteroidota;Bacteroidia;Bacteroidales;Marinifilaceae;Butyricimonas;uncultured bacterium | 93.4 | 1267 | -1 |
| metaRNA.PFemirge\_8147 | 575 | 0.000037 | KM650312.1.1481 | Bacteria;Firmicutes;Negativicutes;Acidaminococcales;Acidaminococcaceae;uncultured;uncultured Firmicutes bacterium | 92.4 | 1344 | -1 |
| metaRNA.PFemirge\_25820 | 563 | 0.000001 | EU454993.1.1404 | Bacteria;Firmicutes;Clostridia;Oscillospirales;Oscillospiraceae;uncultured;uncultured bacterium | 92.0 | 1262 | -1 |
| metaRNA.PFemirge\_11245 | 556 | 0.000110 | HQ783986.1.1437 | Bacteria;Bacteroidota;Bacteroidia;Bacteroidales;Tannerellaceae;Parabacteroides;uncultured organism | 92.4 | 1394 | -1 |
| metaRNA.PFemirge\_12955 | 551 | 0.000010 | HM779434.1.1483 | Bacteria;Proteobacteria;Gammaproteobacteria;Enterobacterales;Aeromonadaceae;Aeromonas;uncultured bacterium | 93.0 | 1333 | -1 |
| metaRNA.PFemirge\_21019 | 550 | 0.000048 | JN607058.1.1380 | Bacteria;Verrucomicrobiota;Chlamydiae;Chlamydiales;Parachlamydiaceae;Neochlamydia;uncultured bacterium | 93.2 | 1276 | -1 |
| metaRNA.PFemirge\_13494 | 550 | 0.000061 | KM205539.1.1438 | Bacteria;Firmicutes;Clostridia;Clostridiales;Clostridiaceae;Clostridium sensu stricto 3;uncultured soil bacterium | 97.8 | 1380 | -1 |
| metaRNA.PFemirge\_28268 | 549 | 0.000173 | CBVB010000006.254456.255954 | Bacteria;Bacteroidota;Bacteroidia;Bacteroidales;Bacteroidaceae;Bacteroides;Bacteroidaceae bacterium MS4 | 93.9 | 1504 | -1 |
| metaRNA.PFemirge\_18207 | 548 | 0.000010 | KX858536.1.1393 | Bacteria;Proteobacteria;Gammaproteobacteria;Pseudomonadales;Pseudomonadaceae;Pseudomonas;Pseudomonas sp. | 96.5 | 1273 | -1 |
| metaRNA.PFemirge\_23347 | 547 | 0.000124 | EU458567.1.1358 | Bacteria;Proteobacteria;Alphaproteobacteria;Rhodospirillales;uncultured;uncultured bacterium | 94.3 | 1273 | -1 |
| metaRNA.PFemirge\_26565 | 547 | 0.000107 | EU776269.1.1279 | Bacteria;Firmicutes;Clostridia;Oscillospirales;Hydrogenoanaerobacterium;uncultured bacterium | 91.9 | 1277 | -1 |
| metaRNA.PFemirge\_23415 | 547 | 0.000071 | AY643083.1.1446 | Bacteria;Bacteroidota;Bacteroidia;Bacteroidales;Rikenellaceae;Alistipes;Alistipes finegoldii | 91.3 | 1291 | -1 |
| metaRNA.PFemirge\_27888 | 542 | 0.000148 | MG367100.1.1491 | Bacteria;Firmicutes;Clostridia;Oscillospirales;Oscillospiraceae;Colidextribacter;uncultured Clostridiales Family IV bacterium | 94.3 | 1466 | -1 |
| metaRNA.PFemirge\_16302 | 535 | 0.000164 | EU775962.1.1393 | Bacteria;Bacteroidota;Bacteroidia;Bacteroidales;Bacteroidaceae;Bacteroides;uncultured bacterium | 91.3 | 1399 | -1 |
| metaRNA.PFemirge\_16968 | 533 | 0.000152 | HQ816198.1.1440 | Bacteria;Bacteroidota;Bacteroidia;Bacteroidales;Bacteroidaceae;Bacteroides;uncultured organism | 90.6 | 1272 | -1 |
| metaRNA.PFemirge\_28125 | 527 | 0.000035 | HQ761452.1.1440 | Bacteria;Bacteroidota;Bacteroidia;Bacteroidales;Tannerellaceae;Parabacteroides;uncultured organism | 92.6 | 1391 | -1 |
| metaRNA.PFemirge\_5019 | 521 | 0.000087 | HQ756942.1.1450 | Bacteria;Firmicutes;Clostridia;Lachnospirales;Lachnospiraceae;[Ruminococcus] gnavus group;uncultured organism | 93.2 | 1376 | -1 |
| metaRNA.PFemirge\_5554 | 516 | 0.000160 | FQXY01000005.366.1875 | Bacteria;Bacteroidota;Bacteroidia;Bacteroidales;Bacteroidaceae;Bacteroides;Bacteroides sp. Marseille-P3132 | 89.0 | 1497 | -1 |
| metaRNA.PFemirge\_13624 | 514 | 0.000038 | JN093128.1.1497 | Bacteria;Firmicutes;Clostridia;Clostridiales;Clostridiaceae;Clostridium sensu stricto 13;Clostridium punense | 97.9 | 1247 | -1 |
| metaRNA.PFemirge\_12903 | 508 | 0.000007 | AB510705.1.1459 | Bacteria;Bacteroidota;Bacteroidia;Bacteroidales;Bacteroidaceae;Bacteroides;Bacteroides ovatus | 93.0 | 1376 | -1 |
| metaRNA.PFemirge\_25536 | 501 | 0.000062 | AY916388.1.1341 | Bacteria;Proteobacteria;Alphaproteobacteria;Rhodospirillales;uncultured;uncultured bacterium | 91.5 | 1253 | -1 |
| metaRNA.PFemirge\_6164 | 499 | 0.000004 | LT223609.1.1490 | Bacteria;Bacteroidota;Bacteroidia;Bacteroidales;Tannerellaceae;Parabacteroides;Parabacteroides faecis | 92.2 | 1457 | -1 |
| metaRNA.PFemirge\_14375 | 499 | 0.000013 | HQ767134.1.1448 | Bacteria;Bacteroidota;Bacteroidia;Bacteroidales;Tannerellaceae;Parabacteroides;uncultured organism | 90.5 | 1362 | -1 |
| metaRNA.PFemirge\_24307 | 494 | 0.000051 | HQ800643.1.1431 | Bacteria;Bacteroidota;Bacteroidia;Bacteroidales;Bacteroidaceae;Bacteroides;uncultured organism | 88.0 | 1310 | -1 |
| metaRNA.PFemirge\_17247 | 493 | 0.000036 | KJ874069.1.1323 | Bacteria;Bacteroidota;Bacteroidia;Bacteroidales;Rikenellaceae;Rikenellaceae RC9 gut group;uncultured bacterium | 86.6 | 1239 | -1 |
| metaRNA.PFemirge\_28017 | 491 | 0.000059 | JX105679.1.1392 | Bacteria;Bacteroidota;Bacteroidia;Bacteroidales;Bacteroidaceae;Bacteroides;uncultured bacterium | 92.3 | 1239 | -1 |
| metaRNA.PFemirge\_15773 | 489 | 0.000014 | HQ697425.1.1459 | Bacteria;Proteobacteria;Alphaproteobacteria;Reyranellales;Reyranellaceae;Reyranella;uncultured bacterium | 92.7 | 1375 | -1 |
| metaRNA.PFemirge\_8537 | 487 | 0.000004 | CDZC01018442.2079.3583 | Bacteria;Bacteroidota;Bacteroidia;Bacteroidales;Bacteroidaceae;Bacteroides;gut metagenome | 90.7 | 1214 | -1 |
| metaRNA.PFemirge\_17357 | 487 | 0.000059 | AB554230.1.1487 | Bacteria;Bacteroidota;Bacteroidia;Bacteroidales;Rikenellaceae;Alistipes;Alistipes finegoldii | 91.6 | 1397 | -1 |
| metaRNA.PFemirge\_5717 | 486 | 0.000041 | LT629973.912339.913840 | Bacteria;Verrucomicrobiota;Verrucomicrobiae;Verrucomicrobiales;Akkermansiaceae;Akkermansia;Akkermansia glycaniphila | 98.8 | 1364 | -1 |
| metaRNA.PFemirge\_30228 | 480 | 0.000122 | JN547325.1.1576 | Eukaryota;Amorphea;Amoebozoa;Discosea;Flabellinia;Dactylopodida;Angulamoeba;uncultured eukaryote | 92.1 | 1298 | -1 |
| metaRNA.PFemirge\_26444 | 480 | 0.000026 | EU776450.1.1387 | Bacteria;Firmicutes;Clostridia;Oscillospirales;Ruminococcaceae;Ruminococcus;uncultured bacterium | 93.8 | 1282 | -1 |
| metaRNA.PFemirge\_23238 | 478 | 0.000034 | AQHW01000007.13.1527 | Bacteria;Bacteroidota;Bacteroidia;Bacteroidales;Tannerellaceae;Parabacteroides;Parabacteroides gordonii MS-1 | 91.7 | 1259 | -1 |
| metaRNA.PFemirge\_15034 | 475 | 0.000305 | JN619858.1.1417 | Bacteria;Bacteroidota;Bacteroidia;Bacteroidales;Rikenellaceae;Alistipes;uncultured bacterium | 93.2 | 1347 | -1 |
| metaRNA.PFemirge\_16632 | 475 | 0.000037 | DQ808503.1.1399 | Bacteria;Bacteroidota;Bacteroidia;Bacteroidales;Tannerellaceae;Parabacteroides;uncultured bacterium | 91.3 | 1404 | -1 |
| metaRNA.PFemirge\_12627 | 474 | 0.000040 | KC669122.1.1486 | Bacteria;Firmicutes;Clostridia;Lachnospirales;Lachnospiraceae;Epulopiscium;uncultured bacterium | 91.4 | 1319 | -1 |
| metaRNA.PFemirge\_4857 | 474 | 0.000005 | JN896612.1.1485 | Bacteria;Actinobacteriota;Actinobacteria;Corynebacteriales;Nocardiaceae;Nocardia;Nocardia sp. FSN271 | 96.7 | 1393 | -1 |
| metaRNA.PFemirge\_28411 | 473 | 0.000010 | CP012938.2352114.2353639 | Bacteria;Bacteroidota;Bacteroidia;Bacteroidales;Bacteroidaceae;Bacteroides;Bacteroides ovatus | 91.3 | 1409 | -1 |
| metaRNA.PFemirge\_1117 | 471 | 0.000229 | GQ871730.1.1506 | Bacteria;Proteobacteria;Gammaproteobacteria;Pseudomonadales;Pseudomonadaceae;Pseudomonas;uncultured bacterium | 87.6 | 1569 | -1 |
| metaRNA.PFemirge\_16518 | 470 | 0.000033 | KP231747.1.1515 | Bacteria;Desulfobacterota;Desulfovibrionia;Desulfovibrionales;Desulfovibrionaceae;Desulfovibrio;uncultured Desulfovibrio sp. | 90.0 | 1404 | -1 |
| metaRNA.PFemirge\_598 | 468 | 0.000032 | KX465194.1.1643 | Eukaryota;SAR;Alveolata;Ciliophora;Intramacronucleata;Litostomatea;Haptoria;Arcuospathidium;uncultured ciliate | 98.9 | 1419 | -1 |
| metaRNA.PFemirge\_17308 | 465 | 0.000014 | HM445960.1.1486 | Bacteria;Bacteroidota;Bacteroidia;Bacteroidales;Rikenellaceae;uncultured;uncultured bacterium | 90.3 | 1395 | -1 |
| metaRNA.PFemirge\_12452 | 465 | 0.000004 | HQ759402.1.1446 | Bacteria;Firmicutes;Clostridia;Clostridia UCG-014;uncultured organism | 91.0 | 1384 | -1 |
| metaRNA.PFemirge\_22798 | 463 | 0.000024 | DQ809702.1.1364 | Bacteria;Firmicutes;Clostridia;Peptostreptococcales-Tissierellales;Peptostreptococcaceae;uncultured;uncultured bacterium | 97.0 | 1356 | -1 |
| metaRNA.PFemirge\_28799 | 462 | 0.000088 | DQ355176.1.1464 | Bacteria;Desulfobacterota;Desulfovibrionia;Desulfovibrionales;Desulfovibrionaceae;Desulfovibrio;uncultured Desulfovibrio sp. | 91.7 | 1452 | -1 |
| metaRNA.PFemirge\_21888 | 462 | 0.000086 | GQ448098.1.1374 | Bacteria;Firmicutes;Clostridia;Christensenellales;Christensenellaceae;Christensenellaceae R-7 group;uncultured bacterium | 92.1 | 1298 | -1 |
| metaRNA.PFemirge\_11747 | 461 | 0.000053 | AJ629069.1.1499 | Bacteria;Firmicutes;Bacilli;Erysipelotrichales;Erysipelotrichaceae;Breznakia;Breznakia pachnodae | 88.9 | 1338 | -1 |
| metaRNA.PFemirge\_5036 | 460 | 0.000110 | CP003153.1901384.1902805 | Bacteria;Proteobacteria;Gammaproteobacteria;Burkholderiales;Rhodocyclaceae;Azospira;Azospira oryzae PS | 94.4 | 1339 | -1 |
| metaRNA.PFemirge\_15717 | 458 | 0.000002 | EU772041.1.1407 | Bacteria;Firmicutes;Negativicutes;Acidaminococcales;Acidaminococcaceae;Phascolarctobacterium;uncultured bacterium | 89.8 | 1351 | -1 |
| metaRNA.PFemirge\_15194 | 457 | 0.000074 | AJ576387.1.1452 | Bacteria;Firmicutes;Bacilli;Erysipelotrichales;Erysipelatoclostridiaceae;Erysipelatoclostridium;uncultured bacterium | 93.2 | 1219 | -1 |
| metaRNA.PFemirge\_24351 | 456 | 0.000045 | GQ417416.1.1463 | Bacteria;Proteobacteria;Gammaproteobacteria;Enterobacterales;Morganellaceae;Providencia;uncultured Providencia sp. | 87.6 | 1310 | -1 |
| metaRNA.PFemirge\_28024 | 455 | 0.000027 | EU762199.1.1370 | Bacteria;Bacteroidota;Bacteroidia;Bacteroidales;Bacteroidaceae;Bacteroides;uncultured bacterium | 92.9 | 1290 | -1 |
| metaRNA.PFemirge\_11899 | 450 | 0.000015 | BAAZ01000703.1371.2875 | Bacteria;Bacteroidota;Bacteroidia;Bacteroidales;Tannerellaceae;Parabacteroides;human gut metagenome | 89.4 | 1452 | -1 |
| metaRNA.PFemirge\_28178 | 448 | 0.000003 | JN559712.1.1498 | Bacteria;Bacteroidota;Bacteroidia;Bacteroidales;Bacteroidaceae;Bacteroides;uncultured bacterium | 87.2 | 1485 | -1 |
| metaRNA.PFemirge\_8407 | 444 | 0.000134 | EU234274.1.1501 | Bacteria;Proteobacteria;Gammaproteobacteria;Burkholderiales;Rhodocyclaceae;C39;uncultured bacterium | 94.4 | 1420 | -1 |
| metaRNA.PFemirge\_23414 | 443 | 0.000014 | CDTW01027470.3361.4864 | Bacteria;Bacteroidota;Bacteroidia;Bacteroidales;Rikenellaceae;Alistipes;gut metagenome | 93.2 | 1292 | -1 |
| metaRNA.PFemirge\_5990 | 443 | 0.000119 | CBVB010000006.254456.255954 | Bacteria;Bacteroidota;Bacteroidia;Bacteroidales;Bacteroidaceae;Bacteroides;Bacteroidaceae bacterium MS4 | 93.4 | 1428 | -1 |
| metaRNA.PFemirge\_22451 | 442 | 0.000005 | KF068124.1.1369 | Bacteria;Bacteroidota;Bacteroidia;Bacteroidales;Tannerellaceae;Parabacteroides;uncultured bacterium | 91.0 | 1370 | -1 |
| metaRNA.PFemirge\_16639 | 441 | 0.000058 | AB649277.1.1490 | Bacteria;Firmicutes;Bacilli;Erysipelotrichales;Erysipelotrichaceae;Faecalitalea;Absiella dolichum | 91.7 | 1390 | -1 |
| metaRNA.PFemirge\_2602 | 439 | 0.000036 | DQ234243.1.1534 | Bacteria;Proteobacteria;Gammaproteobacteria;Burkholderiales;Comamonadaceae;Sphaerotilus;uncultured Burkholderiales bacterium | 97.6 | 1400 | -1 |
| metaRNA.PFemirge\_11629 | 437 | 0.000050 | DQ456478.1.1451 | Bacteria;Firmicutes;Clostridia;Oscillospirales;Oscillospiraceae;Colidextribacter;uncultured bacterium | 93.9 | 1318 | -1 |
| metaRNA.PFemirge\_6783 | 428 | 0.000035 | AB447718.1.1450 | Bacteria;Firmicutes;Clostridia;Oscillospirales;Oscillospiraceae;Colidextribacter;uncultured bacterium | 91.6 | 1326 | -1 |
| metaRNA.PFemirge\_14075 | 423 | 0.000005 | CBVB010000006.254456.255954 | Bacteria;Bacteroidota;Bacteroidia;Bacteroidales;Bacteroidaceae;Bacteroides;Bacteroidaceae bacterium MS4 | 91.2 | 1315 | -1 |
| metaRNA.PFemirge\_12730 | 421 | 0.000104 | HQ761612.1.1444 | Bacteria;Bacteroidota;Bacteroidia;Bacteroidales;Tannerellaceae;Parabacteroides;uncultured organism | 93.9 | 1441 | -1 |
| metaRNA.PFemirge\_22986 | 418 | 0.000045 | CP024819.720120.721669 | Bacteria;Proteobacteria;Gammaproteobacteria;Enterobacterales;Enterobacteriaceae;Citrobacter;Citrobacter freundii | 88.6 | 1407 | -1 |
| metaRNA.PFemirge\_8123 | 418 | 0.000005 | FXDJ01000009.1407067.1408576 | Bacteria;Bacteroidota;Bacteroidia;Bacteroidales;Tannerellaceae;uncultured;Porphyromonadaceae bacterium MV12 | 89.3 | 1491 | -1 |
| metaRNA.PFemirge\_15735 | 410 | 0.000012 | HQ819238.1.1451 | Bacteria;Bacteroidota;Bacteroidia;Bacteroidales;Bacteroidaceae;Bacteroides;uncultured organism | 89.3 | 1295 | -1 |
| metaRNA.PFemirge\_25721 | 409 | 0.000441 | EU775240.1.1308 | Bacteria;Bacteroidota;Bacteroidia;Bacteroidales;Prevotellaceae;Prevotellaceae UCG-003;uncultured bacterium | 86.7 | 1305 | -1 |
| metaRNA.PFemirge\_16686 | 407 | 0.000010 | EU505366.1.1398 | Bacteria;Bacteroidota;Bacteroidia;Bacteroidales;Tannerellaceae;Parabacteroides;uncultured bacterium | 90.8 | 1397 | -1 |
| metaRNA.PFemirge\_1548 | 404 | 0.000008 | KP231747.1.1515 | Bacteria;Desulfobacterota;Desulfovibrionia;Desulfovibrionales;Desulfovibrionaceae;Desulfovibrio;uncultured Desulfovibrio sp. | 91.8 | 1515 | -1 |
| metaRNA.PFemirge\_3372 | 401 | 0.000322 | KJ600954.1.1517 | Bacteria;Proteobacteria;Gammaproteobacteria;Xanthomonadales;Xanthomonadaceae;Stenotrophomonas;uncultured bacterium | 93.9 | 1427 | -1 |
| metaRNA.PFemirge\_8831 | 401 | 0.000040 | AB574480.1.1486 | Bacteria;Bacteroidota;Bacteroidia;Bacteroidales;Bacteroidaceae;Bacteroides;Bacteroides faecichinchillae | 92.7 | 1446 | -1 |
| metaRNA.PFemirge\_14766 | 400 | 0.000059 | AGES01000014.73256.74770 | Bacteria;Bacteroidota;Bacteroidia;Bacteroidales;Tannerellaceae;Parabacteroides;Parabacteroides sp. HGS0025 | 94.7 | 1389 | -1 |
| metaRNA.PFemirge\_11159 | 398 | 0.000109 | HQ793252.1.1444 | Bacteria;Bacteroidota;Bacteroidia;Bacteroidales;Bacteroidaceae;Bacteroides;uncultured organism | 90.8 | 1320 | -1 |
| metaRNA.PFemirge\_30235 | 395 | 0.000216 | DQ799369.1.1397 | Bacteria;Bacteroidota;Bacteroidia;Bacteroidales;Bacteroidaceae;Bacteroides;uncultured bacterium | 84.8 | 1385 | -1 |
| metaRNA.PFemirge\_3735 | 389 | 0.000836 | MF175255.1.1500 | Bacteria;Firmicutes;Clostridia;Lachnospirales;Lachnospiraceae;Lachnoclostridium;Clostridium sp. | 93.7 | 1489 | -1 |
| metaRNA.PFemirge\_27748 | 388 | 0.000033 | GQ898302.1.1489 | Bacteria;Bacteroidota;Bacteroidia;Bacteroidales;Bacteroidaceae;Bacteroides;uncultured bacterium | 93.4 | 1398 | -1 |
| metaRNA.PFemirge\_27920 | 387 | 0.000064 | HQ806032.1.1447 | Bacteria;Bacteroidota;Bacteroidia;Bacteroidales;Tannerellaceae;Parabacteroides;uncultured organism | 91.0 | 1448 | -1 |
| metaRNA.PFemirge\_12037 | 387 | 0.000068 | HQ761612.1.1444 | Bacteria;Bacteroidota;Bacteroidia;Bacteroidales;Tannerellaceae;Parabacteroides;uncultured organism | 88.1 | 1444 | -1 |
| metaRNA.PFemirge\_12795 | 387 | 0.000251 | MG367100.1.1491 | Bacteria;Firmicutes;Clostridia;Oscillospirales;Oscillospiraceae;Colidextribacter;uncultured Clostridiales Family IV bacterium | 93.1 | 1322 | -1 |
| metaRNA.PFemirge\_11694 | 383 | 0.000035 | HQ681866.1.1451 | Bacteria;Bacteroidota;Bacteroidia;Bacteroidales;Bacteroidaceae;Bacteroides;uncultured bacterium | 91.8 | 1427 | -1 |
| metaRNA.PFemirge\_4975 | 380 | 0.000004 | JN559628.1.1499 | Bacteria;Bacteroidota;Bacteroidia;Bacteroidales;Bacteroidaceae;Bacteroides;uncultured bacterium | 90.8 | 1396 | -1 |
| metaRNA.PFemirge\_3072 | 379 | 0.000018 | EF511528.1.1486 | Bacteria;Proteobacteria;Gammaproteobacteria;Xanthomonadales;Xanthomonadaceae;Stenotrophomonas;uncultured bacterium | 94.0 | 1405 | -1 |
| metaRNA.PFemirge\_6704 | 378 | 0.000046 | MG367100.1.1491 | Bacteria;Firmicutes;Clostridia;Oscillospirales;Oscillospiraceae;Colidextribacter;uncultured Clostridiales Family IV bacterium | 94.3 | 1291 | -1 |
| metaRNA.PFemirge\_20894 | 377 | 0.000018 | DQ794702.1.1380 | Bacteria;Firmicutes;Clostridia;Lachnospirales;Lachnospiraceae;Lachnospiraceae UCG-004;uncultured bacterium | 92.1 | 1377 | -1 |
| metaRNA.PFemirge\_17917 | 374 | 0.000030 | AB721051.1.1697 | Eukaryota;Amorphea;Amoebozoa;SAR;Stramenopiles;Labyrinthulomycetes;Sorodiplophrys;uncultured freshwater eukaryote | 98.5 | 1238 | -1 |
| metaRNA.PFemirge\_27747 | 372 | 0.000017 | BAAZ01000703.1371.2875 | Bacteria;Bacteroidota;Bacteroidia;Bacteroidales;Tannerellaceae;Parabacteroides;human gut metagenome | 92.0 | 1301 | -1 |
| metaRNA.PFemirge\_7454 | 371 | 0.000029 | GQ480135.1.1486 | Bacteria;Bacteroidota;Bacteroidia;Chitinophagales;Saprospiraceae;Haliscomenobacter;uncultured bacterium | 95.3 | 1186 | -1 |
| metaRNA.PFemirge\_13192 | 366 | 0.000130 | EF454852.1.1475 | Bacteria;Desulfobacterota;Desulfovibrionia;Desulfovibrionales;Desulfovibrionaceae;uncultured;uncultured Desulfovibrionaceae bacterium | 88.5 | 1261 | -1 |
| metaRNA.PFemirge\_27061 | 361 | 0.000022 | AF480581.1.1470 | Bacteria;Actinobacteriota;Actinobacteria;Corynebacteriales;Mycobacteriaceae;Mycobacterium;Mycobacterium fortuitum | 94.6 | 1214 | -1 |
| metaRNA.PFemirge\_16933 | 359 | 0.000070 | EU778828.1.1398 | Bacteria;Bacteroidota;Bacteroidia;Bacteroidales;Tannerellaceae;Parabacteroides;uncultured bacterium | 93.0 | 1396 | -1 |
| metaRNA.PFemirge\_27879 | 359 | 0.000016 | CBVB010000006.254456.255954 | Bacteria;Bacteroidota;Bacteroidia;Bacteroidales;Bacteroidaceae;Bacteroides;Bacteroidaceae bacterium MS4 | 89.6 | 1379 | -1 |
| metaRNA.PFemirge\_16638 | 358 | 0.000657 | DQ799380.1.1399 | Bacteria;Bacteroidota;Bacteroidia;Bacteroidales;Rikenellaceae;Alistipes;uncultured bacterium | 93.1 | 1400 | -1 |
| metaRNA.PFemirge\_16808 | 357 | 0.000043 | AB554230.1.1487 | Bacteria;Bacteroidota;Bacteroidia;Bacteroidales;Rikenellaceae;Alistipes;Alistipes finegoldii | 89.3 | 1395 | -1 |
| metaRNA.PFemirge\_3901 | 357 | 0.000008 | HM630236.1.1451 | Bacteria;Bacteroidota;Bacteroidia;Bacteroidales;Rikenellaceae;dgA-11 gut group;uncultured bacterium | 87.9 | 1269 | -1 |
| metaRNA.PFemirge\_19940 | 355 | 0.000013 | DQ794177.1.1393 | Bacteria;Firmicutes;Clostridia;Oscillospirales;[Eubacterium] coprostanoligenes group;uncultured bacterium | 89.6 | 972 | -1 |
| metaRNA.PFemirge\_16117 | 354 | 0.000028 | HQ661204.1.1414 | Bacteria;Proteobacteria;Alphaproteobacteria;Holosporales;Holosporaceae;uncultured;uncultured bacterium | 92.9 | 1159 | -1 |
| metaRNA.PFemirge\_8463 | 354 | 0.000025 | GQ898302.1.1489 | Bacteria;Bacteroidota;Bacteroidia;Bacteroidales;Bacteroidaceae;Bacteroides;uncultured bacterium | 94.3 | 1319 | -1 |
| metaRNA.PFemirge\_19090 | 352 | 0.000036 | HQ777507.1.1443 | Bacteria;Firmicutes;Clostridia;Lachnospirales;Lachnospiraceae;Lachnospiraceae UCG-006;uncultured organism | 92.1 | 1170 | -1 |
| metaRNA.PFemirge\_25382 | 349 | 0.000016 | DQ456069.1.1449 | Bacteria;Bacteroidota;Bacteroidia;Bacteroidales;Bacteroidaceae;Bacteroides;uncultured bacterium | 88.1 | 1316 | -1 |
| metaRNA.PFemirge\_14112 | 344 | 0.000025 | JF830134.1.1428 | Bacteria;Proteobacteria;Alphaproteobacteria;Rhodobacterales;Rhodobacteraceae;Pseudorhodobacter;uncultured bacterium | 98.3 | 1235 | -1 |
| metaRNA.PFemirge\_14922 | 344 | 0.000022 | HQ119254.1.1464 | Bacteria;Proteobacteria;Alphaproteobacteria;Dongiales;Dongiaceae;Dongia;uncultured bacterium | 90.0 | 1219 | -1 |
| metaRNA.PFemirge\_16703 | 344 | 0.000022 | EU510318.1.1396 | Bacteria;Firmicutes;Clostridia;Oscillospirales;Oscillospiraceae;Colidextribacter;uncultured bacterium | 90.6 | 1296 | -1 |
| metaRNA.PFemirge\_30238 | 342 | 0.000025 | EF454852.1.1475 | Bacteria;Desulfobacterota;Desulfovibrionia;Desulfovibrionales;Desulfovibrionaceae;uncultured;uncultured Desulfovibrionaceae bacterium | 86.5 | 1262 | -1 |
| metaRNA.PFemirge\_15152 | 342 | 0.000041 | GQ449088.1.1384 | Bacteria;Bacteroidota;Bacteroidia;Bacteroidales;Bacteroidaceae;Bacteroides;uncultured bacterium | 94.0 | 1243 | -1 |
| metaRNA.PFemirge\_23419 | 341 | 0.000002 | AGES01000014.73256.74770 | Bacteria;Bacteroidota;Bacteroidia;Bacteroidales;Tannerellaceae;Parabacteroides;Parabacteroides sp. HGS0025 | 95.0 | 1359 | -1 |
| metaRNA.PFemirge\_5733 | 341 | 0.000056 | HQ785173.1.1448 | Bacteria;Bacteroidota;Bacteroidia;Bacteroidales;Bacteroidaceae;Bacteroides;uncultured organism | 92.2 | 1432 | -1 |
| metaRNA.PFemirge\_27990 | 341 | 0.000007 | GQ448371.1.1394 | Bacteria;Bacteroidota;Bacteroidia;Bacteroidales;Bacteroidaceae;Bacteroides;uncultured bacterium | 88.9 | 1281 | -1 |
| metaRNA.PFemirge\_17111 | 340 | 0.000035 | CP022049.2235979.2237530 | Bacteria;Proteobacteria;Gammaproteobacteria;Enterobacterales;Enterobacteriaceae;Citrobacter;Citrobacter braakii | 93.1 | 1396 | -1 |
| metaRNA.PFemirge\_28067 | 338 | 0.000010 | LT725663.1.1450 | Bacteria;Bacteroidota;Bacteroidia;Bacteroidales;Tannerellaceae;Parabacteroides;Parabacteroides sp. Marseille-P3668 | 93.1 | 1237 | -1 |
| metaRNA.PFemirge\_16925 | 337 | 0.000100 | DQ824259.1.1398 | Bacteria;Bacteroidota;Bacteroidia;Bacteroidales;Bacteroidaceae;Bacteroides;uncultured bacterium | 92.2 | 1304 | -1 |
| metaRNA.PFemirge\_7416 | 337 | 0.000030 | AB649276.1.1495 | Bacteria;Firmicutes;Clostridia;Lachnospirales;Lachnospiraceae;Anaerostignum;Anaerotignum propionicum | 94.5 | 1382 | -1 |
| metaRNA.PFemirge\_2689 | 335 | 0.000027 | AQHW01000007.13.1527 | Bacteria;Bacteroidota;Bacteroidia;Bacteroidales;Tannerellaceae;Parabacteroides;Parabacteroides gordonii MS-1 | 91.3 | 1501 | -1 |
| metaRNA.PFemirge\_30251 | 333 | 0.000064 | JQ248079.1.1513 | Bacteria;Firmicutes;Clostridia;Oscillospirales;Oscillospiraceae;Flavonifractor;uncultured bacterium | 88.2 | 1242 | -1 |
| metaRNA.PFemirge\_23621 | 329 | 0.000030 | HQ819316.1.1450 | Bacteria;Bacteroidota;Bacteroidia;Bacteroidales;Bacteroidaceae;Bacteroides;uncultured organism | 90.4 | 1359 | -1 |
| metaRNA.PFemirge\_16188 | 325 | 0.000021 | GQ417619.1.1463 | Bacteria;Proteobacteria;Gammaproteobacteria;Enterobacterales;Enterobacteriaceae;Citrobacter;uncultured Citrobacter sp. | 94.4 | 1326 | -1 |
| metaRNA.PFemirge\_20908 | 322 | 0.000012 | DQ805525.1.1380 | Bacteria;Firmicutes;Clostridia;Lachnospirales;Lachnospiraceae;Lachnospiraceae UCG-008;uncultured bacterium | 90.6 | 1380 | -1 |
| metaRNA.PFemirge\_8296 | 321 | 0.000066 | KP975338.1.1480 | Bacteria;Proteobacteria;Gammaproteobacteria;Enterobacterales;Aeromonadaceae;Aeromonas;uncultured bacterium | 94.4 | 1479 | -1 |
| metaRNA.PFemirge\_237 | 315 | 0.000025 | KJ563218.1.1771 | Eukaryota;Amorphea;Obazoa;Opisthokonta;Nucletmycea;Fungi;Blastocladiomycota;Incertae Sedis;Blastocladiomycetes;Blastocladiales;Physodermataceae;Paraphysoderma;Paraphysoderma sedebokerense | 97.0 | 1619 | -1 |
| metaRNA.PFemirge\_16574 | 314 | 0.000014 | GQ448730.1.1400 | Bacteria;Bacteroidota;Bacteroidia;Bacteroidales;Bacteroidaceae;Bacteroides;uncultured bacterium | 91.0 | 1173 | -1 |
| metaRNA.PFemirge\_24201 | 313 | 0.000316 | CP003153.1901384.1902805 | Bacteria;Proteobacteria;Gammaproteobacteria;Burkholderiales;Rhodocyclaceae;Azospira;Azospira oryzae PS | 89.6 | 1378 | -1 |
| metaRNA.PFemirge\_13650 | 312 | 0.000076 | JQ940836.1.1449 | Bacteria;Bacteroidota;Bacteroidia;Bacteroidales;Rikenellaceae;Alistipes;uncultured bacterium | 94.5 | 1436 | -1 |
| metaRNA.PFemirge\_3525 | 311 | 0.000015 | HQ697726.1.1505 | Bacteria;Proteobacteria;Gammaproteobacteria;Acidithiobacillales;Acidithiobacillaceae;KCM-B-112;uncultured bacterium | 92.9 | 1370 | -1 |
| metaRNA.PFemirge\_11834 | 310 | 0.000029 | KC887929.1.1472 | Bacteria;Proteobacteria;Gammaproteobacteria;Pseudomonadales;Pseudomonadaceae;Pseudomonas;proteobacterium K10 | 96.0 | 1288 | -1 |
| metaRNA.PFemirge\_24111 | 309 | 0.000032 | GQ492245.1.1349 | Bacteria;Firmicutes;Bacilli;Erysipelotrichales;Erysipelatoclostridiaceae;Candidatus Stoquefichus;uncultured bacterium | 94.1 | 1156 | -1 |
| metaRNA.PFemirge\_5076 | 304 | 0.000024 | JF808852.1.1503 | Bacteria;Proteobacteria;Gammaproteobacteria;Burkholderiales;Rhodocyclaceae;Azospira;uncultured bacterium | 93.6 | 1406 | -1 |
| metaRNA.PFemirge\_11872 | 304 | 0.000003 | JN680575.1.1450 | Bacteria;Bacteroidota;Bacteroidia;Bacteroidales;Dysgonomonadaceae;Dysgonomonas;uncultured Porphyromonadaceae bacterium | 88.7 | 1430 | -1 |
| metaRNA.PFemirge\_11161 | 303 | 0.000016 | EU828415.1.1454 | Bacteria;Proteobacteria;Gammaproteobacteria;Burkholderiales;Oxalobacteraceae;Janthinobacterium;uncultured bacterium | 93.2 | 1234 | -1 |
| metaRNA.PFemirge\_3510 | 300 | 0.000010 | GQ867427.1.1505 | Bacteria;Proteobacteria;Gammaproteobacteria;Enterobacterales;Enterobacteriaceae;Escherichia-Shigella;uncultured bacterium | 93.3 | 1495 | -1 |
| metaRNA.PFemirge\_17635 | 299 | 0.000012 | DQ794860.1.1394 | Bacteria;Firmicutes;Clostridia;Lachnospirales;Lachnospiraceae;[Eubacterium] oxidoreducens group;uncultured bacterium | 91.8 | 1391 | -1 |
| metaRNA.PFemirge\_5756 | 297 | 0.000092 | AY957942.1.1505 | Bacteria;Actinobacteriota;Actinobacteria;Corynebacteriales;Mycobacteriaceae;Mycobacterium;uncultured bacterium | 98.1 | 1391 | -1 |
| metaRNA.PFemirge\_30262 | 296 | 0.000015 | AB238925.1.1486 | Bacteria;Bacteroidota;Bacteroidia;Bacteroidales;Tannerellaceae;Parabacteroides;Parabacteroides distasonis | 90.5 | 1451 | -1 |
| metaRNA.PFemirge\_28197 | 295 | 0.000021 | LARG01000025.1.1402 | Bacteria;Bacteroidota;Bacteroidia;Bacteroidales;Rikenellaceae;Alistipes;Alistipes sp. N15.MGS-157 | 90.9 | 1313 | -1 |
| metaRNA.PFemirge\_23152 | 294 | 0.000010 | HQ778029.1.1454 | Bacteria;Firmicutes;Clostridia;Lachnospirales;Lachnospiraceae;Lachnoclostridium;uncultured organism | 90.7 | 1296 | -1 |
| metaRNA.PFemirge\_2576 | 293 | 0.000068 | DQ681227.1.1525 | Bacteria;Firmicutes;Clostridia;Lachnospirales;Lachnospiraceae;Robinsoniella;Robinsoniella peoriensis | 96.6 | 1468 | -1 |
| metaRNA.PFemirge\_2932 | 292 | 0.000015 | JF830177.1.1510 | Bacteria;Proteobacteria;Gammaproteobacteria;Pseudomonadales;Moraxellaceae;Acinetobacter;uncultured bacterium | 95.0 | 1501 | -1 |
| metaRNA.PFemirge\_26216 | 292 | 0.000003 | DQ795734.1.1397 | Bacteria;Firmicutes;Clostridia;Lachnospirales;Lachnospiraceae;[Ruminococcus] gauvreauii group;uncultured bacterium | 92.0 | 1292 | -1 |
| metaRNA.PFemirge\_8912 | 292 | 0.000005 | KC189649.1.1474 | Bacteria;Bacteroidota;Bacteroidia;Chitinophagales;Saprospiraceae;uncultured;uncultured bacterium | 97.4 | 1136 | -1 |
| metaRNA.PFemirge\_22806 | 291 | 0.000050 | LN650648.388047.389538 | Bacteria;Firmicutes;Clostridia;Peptostreptococcales-Tissierellales;Peptostreptococcaceae;Romboutsia;Romboutsia hominis | 94.3 | 1359 | -1 |
| metaRNA.PFemirge\_7425 | 291 | 0.000007 | JX457238.1.1481 | Bacteria;Bacteroidota;Bacteroidia;Bacteroidales;Rikenellaceae;Alistipes;uncultured bacterium | 88.7 | 1323 | -1 |
| metaRNA.PFemirge\_12067 | 291 | 0.000096 | AB574482.1.1489 | Bacteria;Bacteroidota;Bacteroidia;Bacteroidales;Tannerellaceae;Parabacteroides;Parabacteroides chinchillae | 89.1 | 1393 | -1 |
| metaRNA.PFemirge\_17925 | 290 | 0.003077 | DQ793765.1.1405 | Bacteria;Firmicutes;Clostridia;Oscillospirales;[Eubacterium] coprostanoligenes group;uncultured bacterium | 91.6 | 1287 | -1 |
| metaRNA.PFemirge\_2798 | 288 | 0.000019 | FPLS01017333.1.1416 | Bacteria;Proteobacteria;Gammaproteobacteria;Gammaproteobacteria Incertae Sedis;Unknown Family;metagenome | 97.1 | 1293 | -1 |
| metaRNA.PFemirge\_26412 | 287 | 0.000030 | DQ807523.1.1284 | Bacteria;Firmicutes;Clostridia;Lachnospirales;Lachnospiraceae;[Ruminococcus] torques group;uncultured bacterium | 91.2 | 1199 | -1 |
| metaRNA.PFemirge\_5047 | 285 | 0.000017 | MF085166.1.1529 | Bacteria;Proteobacteria;Gammaproteobacteria;Acidithiobacillales;Acidithiobacillaceae;KCM-B-112;uncultured bacterium | 98.0 | 1407 | -1 |
| metaRNA.PFemirge\_22623 | 284 | 0.000102 | DQ807369.1.1362 | Bacteria;Firmicutes;Clostridia;Peptostreptococcales-Tissierellales;Peptostreptococcaceae;uncultured;uncultured bacterium | 96.0 | 1363 | -1 |
| metaRNA.PFemirge\_6835 | 284 | 0.000022 | AY957892.1.1500 | Bacteria;Cyanobacteria;Vampirivibrionia;Obscuribacterales;Obscuribacteraceae;uncultured bacterium | 99.9 | 1244 | -1 |
| metaRNA.PFemirge\_6921 | 284 | 0.000052 | FQSE01000007.1631802.1633308 | Bacteria;Bacteroidota;Bacteroidia;Bacteroidales;Tannerellaceae;Parabacteroides;Parabacteroides sp. Marseille-P3136 | 88.1 | 1455 | -1 |
| metaRNA.PFemirge\_22900 | 284 | 0.000058 | FJ366291.1.1363 | Bacteria;Firmicutes;Clostridia;Oscillospirales;Oscillospiraceae;UCG-002;uncultured bacterium | 91.7 | 1279 | -1 |
| metaRNA.PFemirge\_27758 | 283 | 0.000017 | LT223609.1.1490 | Bacteria;Bacteroidota;Bacteroidia;Bacteroidales;Tannerellaceae;Parabacteroides;Parabacteroides faecis | 91.7 | 1392 | -1 |
| metaRNA.PFemirge\_12072 | 283 | 0.000009 | CP009057.2296841.2298384 | Bacteria;Bacteroidota;Bacteroidia;Bacteroidales;Bacteroidaceae;Bacteroides;Bacteroides dorei | 88.4 | 1322 | -1 |
| metaRNA.PFemirge\_12364 | 282 | 0.000025 | AB470343.1.1485 | Bacteria;Bacteroidota;Bacteroidia;Bacteroidales;Tannerellaceae;Parabacteroides;Parabacteroides gordonii | 90.6 | 1340 | -1 |
| metaRNA.PFemirge\_3476 | 282 | 0.000057 | FJ184339.1.1505 | Bacteria;Proteobacteria;Gammaproteobacteria;Xanthomonadales;Xanthomonadaceae;Stenotrophomonas;uncultured soil bacterium | 97.9 | 1406 | -1 |
| metaRNA.PFemirge\_11239 | 281 | 0.000029 | KM650266.1.1454 | Bacteria;Firmicutes;Clostridia;Oscillospirales;Oscillospiraceae;UCG-002;uncultured proteobacterium | 89.7 | 1413 | -1 |
| metaRNA.PFemirge\_15396 | 280 | 0.000009 | GFIG01012561.7.1417 | Bacteria;Proteobacteria;Gammaproteobacteria;Enterobacterales;Enterobacteriaceae;Escherichia-Shigella;Coregonus lavaretus (common whitefish) | 94.9 | 1325 | -1 |
| metaRNA.PFemirge\_24439 | 280 | 0.000029 | EU777165.1.1344 | Bacteria;Firmicutes;Clostridia;Oscillospirales;Oscillospiraceae;Oscillibacter;uncultured bacterium | 92.4 | 1252 | -1 |
| metaRNA.PFemirge\_27320 | 280 | 0.000082 | JF490035.1.1450 | Bacteria;Proteobacteria;Alphaproteobacteria;Rhodospirillales;Magnetospirillaceae;uncultured;alpha proteobacterium UT-2 | 90.8 | 1123 | -1 |
| metaRNA.PFemirge\_15280 | 280 | 0.000028 | JX221903.1.1461 | Bacteria;Proteobacteria;Alphaproteobacteria;Holosporales;Holosporaceae;uncultured;uncultured bacterium | 90.9 | 1227 | -1 |
| metaRNA.PFemirge\_13617 | 279 | 0.000003 | AY328682.1.1446 | Bacteria;Proteobacteria;Alphaproteobacteria;Caulobacterales;Hyphomonadaceae;uncultured;uncultured bacterium | 94.5 | 1296 | -1 |
| metaRNA.PFemirge\_20835 | 279 | 0.000027 | AE015928.3034714.3036195 | Bacteria;Bacteroidota;Bacteroidia;Bacteroidales;Bacteroidaceae;Bacteroides;Bacteroides thetaiotaomicron VPI-5482 | 91.9 | 1388 | -1 |
| metaRNA.PFemirge\_13713 | 279 | 0.000009 | AB218288.1.1449 | Bacteria;Bacteroidota;Bacteroidia;Bacteroidales;Tannerellaceae;Parabacteroides;uncultured Bacteroidaceae bacterium | 87.9 | 1434 | -1 |
| metaRNA.PFemirge\_20425 | 276 | 0.000055 | GQ448684.1.1384 | Bacteria;Bacteroidota;Bacteroidia;Bacteroidales;Rikenellaceae;Alistipes;uncultured bacterium | 93.2 | 1373 | -1 |
| metaRNA.PFemirge\_30229 | 274 | 0.000284 | HM017143.1.1714 | Eukaryota;Amorphea;Amoebozoa;Discosea;Flabellinia;Dactylopodida;Angulamoeba;Angulamoeba microcystivorans | 94.5 | 1434 | -1 |
| metaRNA.PFemirge\_2582 | 273 | 0.000049 | JN038827.1.1489 | Bacteria;Proteobacteria;Gammaproteobacteria;Burkholderiales;Sutterellaceae;uncultured;uncultured beta proteobacterium | 92.8 | 1401 | -1 |
| metaRNA.PFemirge\_1169 | 273 | 0.000023 | FR749938.1.1523 | Bacteria;Firmicutes;Clostridia;Lachnospirales;Lachnospiraceae;[Eubacterium] fissicatena group;Faecalicatena fissicatena | 91.4 | 1310 | -1 |
| metaRNA.PFemirge\_2507 | 272 | 0.000048 | JQ083789.1.1511 | Bacteria;Desulfobacterota;Desulfovibrionia;Desulfovibrionales;Desulfovibrionaceae;Bilophila;uncultured bacterium | 92.3 | 1501 | -1 |
| metaRNA.PFemirge\_27473 | 271 | 0.000022 | HG934468.2032323.2033833 | Bacteria;Bacteroidota;Bacteroidia;Bacteroidales;Rikenellaceae;Mucinivorans;Mucinivorans hirudinis | 90.1 | 1017 | -1 |
| metaRNA.PFemirge\_1634 | 271 | 0.000024 | GQ480160.1.1510 | Bacteria;Firmicutes;Negativicutes;Acidaminococcales;Acidaminococcaceae;Succinispira;uncultured bacterium | 89.9 | 1375 | -1 |
| metaRNA.PFemirge\_14522 | 270 | 0.000011 | LARG01000025.1.1402 | Bacteria;Bacteroidota;Bacteroidia;Bacteroidales;Rikenellaceae;Alistipes;Alistipes sp. N15.MGS-157 | 92.8 | 1203 | -1 |
| metaRNA.PFemirge\_6044 | 268 | 0.000013 | FJ542833.1.1492 | Bacteria;Verrucomicrobiota;Verrucomicrobiae;Chthoniobacterales;Terrimicrobiaceae;Terrimicrobium;uncultured Verrucomicrobiales bacterium | 95.6 | 1181 | -1 |
| metaRNA.PFemirge\_19378 | 268 | 0.000042 | GU322806.1.1435 | Bacteria;Firmicutes;Clostridia;Lachnospirales;Lachnospiraceae;Robinsoniella;Robinsoniella peoriensis | 91.0 | 1394 | -1 |
| metaRNA.PFemirge\_16483 | 267 | 0.000024 | FJ534971.1.1485 | Bacteria;Bacteroidota;Bacteroidia;Bacteroidales;Rikenellaceae;uncultured;uncultured bacterium | 92.8 | 1397 | -1 |
| metaRNA.PFemirge\_20446 | 267 | 0.000003 | HQ803101.1.1444 | Bacteria;Bacteroidota;Bacteroidia;Bacteroidales;Bacteroidaceae;Bacteroides;uncultured organism | 90.9 | 1029 | -1 |
| metaRNA.PFemirge\_4037 | 266 | 0.000034 | GU056304.1.1502 | Bacteria;Proteobacteria;Gammaproteobacteria;Burkholderiales;Rhodocyclaceae;Azonexus;Azonexus sp. enrichment culture clone ECC4-1 | 98.4 | 1416 | -1 |
| metaRNA.PFemirge\_26002 | 265 | 0.000049 | EU777328.1.1296 | Bacteria;Firmicutes;Bacilli;RF39;uncultured bacterium | 92.3 | 1293 | -1 |
| metaRNA.PFemirge\_7133 | 265 | 0.000004 | CP002345.1438824.1440339 | Bacteria;Bacteroidota;Bacteroidia;Bacteroidales;Paludibacteraceae;Paludibacter;Paludibacter propionicigenes WB4 | 87.9 | 1439 | -1 |
| metaRNA.PFemirge\_23236 | 264 | 0.000271 | HM630236.1.1451 | Bacteria;Bacteroidota;Bacteroidia;Bacteroidales;Rikenellaceae;dgA-11 gut group;uncultured bacterium | 90.2 | 1361 | -1 |
| metaRNA.PFemirge\_12566 | 263 | 0.000019 | KC189686.1.1429 | Bacteria;Proteobacteria;Alphaproteobacteria;Rhodobacterales;Rhodobacteraceae;Rhodobacter;uncultured bacterium | 94.9 | 1227 | -1 |
| metaRNA.PFemirge\_20578 | 262 | 0.000005 | DQ801443.1.1401 | Bacteria;Bacteroidota;Bacteroidia;Bacteroidales;Bacteroidaceae;Bacteroides;uncultured bacterium | 85.9 | 1379 | -1 |
| metaRNA.PFemirge\_16647 | 262 | 0.000019 | DQ808503.1.1399 | Bacteria;Bacteroidota;Bacteroidia;Bacteroidales;Tannerellaceae;Parabacteroides;uncultured bacterium | 93.1 | 1402 | -1 |
| metaRNA.PFemirge\_7082 | 261 | 0.000018 | AQHW01000007.13.1527 | Bacteria;Bacteroidota;Bacteroidia;Bacteroidales;Tannerellaceae;Parabacteroides;Parabacteroides gordonii MS-1 | 94.2 | 1488 | -1 |
| metaRNA.PFemirge\_28014 | 260 | 0.000010 | ADLN01000129.202.1639 | Bacteria;Firmicutes;Clostridia;Lachnospirales;Lachnospiraceae;Incertae Sedis;Hungatella hathewayi WAL-18680 | 93.0 | 1388 | -1 |
| metaRNA.PFemirge\_14769 | 260 | 0.000018 | KP231747.1.1515 | Bacteria;Desulfobacterota;Desulfovibrionia;Desulfovibrionales;Desulfovibrionaceae;Desulfovibrio;uncultured Desulfovibrio sp. | 93.1 | 1071 | -1 |
| metaRNA.PFemirge\_27653 | 259 | 0.000009 | AY721628.1.1491 | Bacteria;Desulfobacterota;Desulfovibrionia;Desulfovibrionales;Desulfovibrionaceae;Desulfovibrio;bacterium 'New Zealand D' | 91.1 | 1487 | -1 |
| metaRNA.PFemirge\_3711 | 258 | 0.000017 | HQ588840.1.1529 | Bacteria;Proteobacteria;Gammaproteobacteria;Pseudomonadales;Pseudomonadaceae;Pseudomonas;Pseudomonas sp. BZ77 | 95.4 | 1488 | -1 |
| metaRNA.PFemirge\_28036 | 256 | 0.000003 | GQ359970.1.1475 | Bacteria;Firmicutes;Clostridia;Clostridiales;Clostridiaceae;Clostridium sensu stricto 1;uncultured bacterium | 91.4 | 1375 | -1 |
| metaRNA.PFemirge\_17674 | 256 | 0.000028 | DQ808767.1.1394 | Bacteria;Firmicutes;Clostridia;Lachnospirales;Lachnospiraceae;Lachnospiraceae UCG-008;uncultured bacterium | 93.0 | 1379 | -1 |
| metaRNA.PFemirge\_20308 | 253 | 0.000025 | AF502202.1.1376 | Bacteria;Proteobacteria;Alphaproteobacteria;Rickettsiales;Mitochondria;uncultured bacterium | 99.7 | 1080 | -1 |
| metaRNA.PFemirge\_7034 | 252 | 0.000006 | JX457157.1.1486 | Bacteria;Bacteroidota;Bacteroidia;Bacteroidales;Tannerellaceae;Parabacteroides;uncultured bacterium | 91.2 | 1409 | -1 |
| metaRNA.PFemirge\_20627 | 252 | 0.000004 | AB649276.1.1495 | Bacteria;Firmicutes;Clostridia;Lachnospirales;Lachnospiraceae;Anaerostignum;Anaerotignum propionicum | 88.7 | 1298 | -1 |
| metaRNA.PFemirge\_8044 | 251 | 0.000160 | CP012937.757465.758990 | Bacteria;Bacteroidota;Bacteroidia;Bacteroidales;Bacteroidaceae;Bacteroides;Bacteroides thetaiotaomicron | 88.5 | 1498 | -1 |
| metaRNA.PFemirge\_24647 | 250 | 0.000018 | KP231747.1.1515 | Bacteria;Desulfobacterota;Desulfovibrionia;Desulfovibrionales;Desulfovibrionaceae;Desulfovibrio;uncultured Desulfovibrio sp. | 84.2 | 1312 | -1 |
| metaRNA.PFemirge\_5910 | 249 | 0.000239 | JF830174.1.1513 | Bacteria;Proteobacteria;Gammaproteobacteria;Enterobacterales;Enterobacteriaceae;Plesiomonas;uncultured bacterium | 88.9 | 1312 | -1 |
| metaRNA.PFemirge\_5519 | 248 | 0.000005 | BAAY01003437.486.1979 | Bacteria;Firmicutes;Clostridia;Lachnospirales;Lachnospiraceae;uncultured;human gut metagenome | 93.4 | 1314 | -1 |
| metaRNA.PFemirge\_20784 | 248 | 0.000026 | EU763324.1.1351 | Bacteria;Firmicutes;Clostridia;Lachnospirales;Lachnospiraceae;Lachnospiraceae NK4A136 group;uncultured bacterium | 90.2 | 1304 | -1 |
| metaRNA.PFemirge\_24301 | 247 | 0.000003 | FR749936.1.1524 | Bacteria;Firmicutes;Clostridia;Lachnospirales;Lachnospiraceae;[Eubacterium] fissicatena group;Faecalicatena fissicatena | 95.0 | 1240 | -1 |
| metaRNA.PFemirge\_22315 | 245 | 0.000005 | EU773345.1.1375 | Bacteria;Firmicutes;Clostridia;Clostridiales;Clostridiaceae;Clostridium sensu stricto 1;uncultured bacterium | 92.8 | 1367 | -1 |
| metaRNA.PFemirge\_1829 | 245 | 0.000025 | LARG01000024.765.2171 | Bacteria;Bacteroidota;Bacteroidia;Bacteroidales;Rikenellaceae;Alistipes;Alistipes sp. N15.MGS-157 | 92.4 | 1323 | -1 |
| metaRNA.PFemirge\_24040 | 244 | 0.000018 | AJ289162.1.1486 | Bacteria;Proteobacteria;Gammaproteobacteria;Pseudomonadales;Cellvibrionaceae;Cellvibrio;Cellvibrio gandavensis | 98.9 | 1322 | -1 |
| metaRNA.PFemirge\_6413 | 243 | 0.000035 | JQ940836.1.1449 | Bacteria;Bacteroidota;Bacteroidia;Bacteroidales;Rikenellaceae;Alistipes;uncultured bacterium | 91.7 | 1335 | -1 |
| metaRNA.PFemirge\_20554 | 243 | 0.000011 | MG367100.1.1491 | Bacteria;Firmicutes;Clostridia;Oscillospirales;Oscillospiraceae;Colidextribacter;uncultured Clostridiales Family IV bacterium | 87.5 | 1261 | -1 |
| metaRNA.PFemirge\_19321 | 243 | 0.000007 | CP022754.4526807.4528346 | Bacteria;Bacteroidota;Bacteroidia;Bacteroidales;Tannerellaceae;Parabacteroides;Parabacteroides sp. CT06 | 94.3 | 1388 | -1 |
| metaRNA.PFemirge\_10492 | 242 | 0.000020 | MF145698.1.1460 | Bacteria;Firmicutes;Clostridia;Peptostreptococcales-Tissierellales;Peptostreptococcaceae;Terrisporobacter;Terrisporobacter glycolicus | 95.7 | 1389 | -1 |
| metaRNA.PFemirge\_14690 | 242 | 0.000022 | HQ751117.1.1433 | Bacteria;Firmicutes;Clostridia;Lachnospirales;Lachnospiraceae;Lachnospiraceae UCG-004;uncultured organism | 91.2 | 1413 | -1 |
| metaRNA.PFemirge\_8975 | 240 | 0.000015 | KF441638.1.1495 | Bacteria;Bacteroidota;Bacteroidia;Flavobacteriales;Flavobacteriaceae;Flavobacterium;Flavobacterium sp. 7A-207 | 97.2 | 1278 | -1 |
| metaRNA.PFemirge\_12309 | 238 | 0.000016 | HQ740432.1.1447 | Bacteria;Bacteroidota;Bacteroidia;Bacteroidales;Marinifilaceae;Odoribacter;uncultured organism | 93.5 | 1268 | -1 |
| metaRNA.PFemirge\_25491 | 238 | 0.000005 | HQ781428.1.1455 | Bacteria;Firmicutes;Clostridia;Lachnospirales;Lachnospiraceae;uncultured;uncultured organism | 92.4 | 1149 | -1 |
| metaRNA.PFemirge\_28304 | 238 | 0.000021 | GQ448867.1.1381 | Bacteria;Bacteroidota;Bacteroidia;Bacteroidales;Tannerellaceae;Parabacteroides;uncultured bacterium | 92.8 | 1331 | -1 |
| metaRNA.PFemirge\_18472 | 231 | 0.000010 | EU772357.1.1394 | Bacteria;Bacteroidota;Bacteroidia;Bacteroidales;p-251-o5;uncultured bacterium | 89.4 | 1194 | -1 |
| metaRNA.PFemirge\_19894 | 230 | 0.000009 | JX198613.1.1484 | Bacteria;Firmicutes;Clostridia;Lachnospirales;Lachnospiraceae;[Ruminococcus] gauvreauii group;uncultured bacterium | 91.0 | 1305 | -1 |
| metaRNA.PFemirge\_22671 | 229 | 0.000052 | CP022115.2965538.2967082 | Bacteria;Proteobacteria;Gammaproteobacteria;Burkholderiales;Aquaspirillaceae;Laribacter;Laribacter hongkongensis | 93.5 | 1297 | -1 |
| metaRNA.PFemirge\_15658 | 228 | 0.000008 | JN653056.1.1473 | Bacteria;Desulfobacterota;Desulfovibrionia;Desulfovibrionales;Desulfovibrionaceae;Desulfovibrio;uncultured Desulfovibrionaceae bacterium | 89.7 | 1328 | -1 |
| metaRNA.PFemirge\_11419 | 228 | 0.000039 | HQ806032.1.1447 | Bacteria;Bacteroidota;Bacteroidia;Bacteroidales;Tannerellaceae;Parabacteroides;uncultured organism | 91.3 | 1452 | -1 |
| metaRNA.PFemirge\_14656 | 227 | 0.000017 | LK056647.1.1423 | Bacteria;Proteobacteria;Alphaproteobacteria;Sphingomonadales;Sphingomonadaceae;Novosphingobium;Novosphingobium piscinae | 92.9 | 1290 | -1 |
| metaRNA.PFemirge\_27669 | 226 | 0.000009 | AB574482.1.1489 | Bacteria;Bacteroidota;Bacteroidia;Bacteroidales;Tannerellaceae;Parabacteroides;Parabacteroides chinchillae | 92.1 | 1491 | -1 |
| metaRNA.PFemirge\_30258 | 225 | 0.000571 | DQ456246.1.1451 | Bacteria;Bacteroidota;Bacteroidia;Bacteroidales;Bacteroidaceae;Bacteroides;uncultured bacterium | 86.9 | 1295 | -1 |
| metaRNA.PFemirge\_18586 | 223 | 0.000031 | DQ856503.1.1485 | Bacteria;Bacteroidota;Bacteroidia;Bacteroidales;Bacteroidaceae;Bacteroides;uncultured bacterium | 93.9 | 1256 | -1 |
| metaRNA.PFemirge\_20099 | 222 | 0.000012 | EU771929.1.1386 | Bacteria;Firmicutes;Clostridia;Lachnospirales;Lachnospiraceae;Lachnospiraceae UCG-008;uncultured bacterium | 90.9 | 1382 | -1 |
| metaRNA.PFemirge\_26979 | 221 | 0.000065 | DQ815225.1.1263 | Bacteria;Desulfobacterota;Desulfovibrionia;Desulfovibrionales;Desulfovibrionaceae;uncultured;uncultured bacterium | 92.2 | 1242 | -1 |
| metaRNA.PFemirge\_2460 | 220 | 0.000052 | U82813.1.1467 | Bacteria;Desulfobacterota;Desulfovibrionia;Desulfovibrionales;Desulfovibrionaceae;Bilophila;Bilophila wadsworthia | 91.0 | 1454 | -1 |
| metaRNA.PFemirge\_16170 | 219 | 0.000009 | LT725663.1.1450 | Bacteria;Bacteroidota;Bacteroidia;Bacteroidales;Tannerellaceae;Parabacteroides;Parabacteroides sp. Marseille-P3668 | 89.8 | 1231 | -1 |
| metaRNA.PFemirge\_12489 | 219 | 0.000063 | CBVB010000006.254456.255954 | Bacteria;Bacteroidota;Bacteroidia;Bacteroidales;Bacteroidaceae;Bacteroides;Bacteroidaceae bacterium MS4 | 88.5 | 1447 | -1 |
| metaRNA.PFemirge\_8965 | 219 | 0.000036 | KP231747.1.1515 | Bacteria;Desulfobacterota;Desulfovibrionia;Desulfovibrionales;Desulfovibrionaceae;Desulfovibrio;uncultured Desulfovibrio sp. | 93.2 | 1464 | -1 |
| metaRNA.PFemirge\_28444 | 218 | 0.000111 | KC503998.1.1475 | Bacteria;Firmicutes;Bacilli;Bacillales;Planococcaceae;Lysinibacillus;Lysinibacillus sphaericus | 84.9 | 1356 | -1 |
| metaRNA.PFemirge\_17214 | 217 | 0.000011 | HQ681866.1.1451 | Bacteria;Bacteroidota;Bacteroidia;Bacteroidales;Bacteroidaceae;Bacteroides;uncultured bacterium | 88.6 | 1356 | -1 |
| metaRNA.PFemirge\_26413 | 217 | 0.000001 | GQ448053.1.1387 | Bacteria;Firmicutes;Clostridia;Peptostreptococcales-Tissierellales;Anaerovoracaceae;[Eubacterium] nodatum group;uncultured bacterium | 90.9 | 1281 | -1 |
| metaRNA.PFemirge\_22403 | 217 | 0.000019 | KR822473.1.1454 | Bacteria;Firmicutes;Clostridia;Peptostreptococcales-Tissierellales;Anaerovoracaceae;[Eubacterium] nodatum group;bacterium PM5-7 | 91.1 | 1174 | -1 |
| metaRNA.PFemirge\_28668 | 217 | 0.000017 | AJ867049.1.1500 | Bacteria;Desulfobacterota;Desulfovibrionia;Desulfovibrionales;Desulfovibrionaceae;Bilophila;Bilophila wadsworthia | 91.4 | 1505 | -1 |
| metaRNA.PFemirge\_26324 | 216 | 0.000002 | DQ800679.1.1366 | Bacteria;Firmicutes;Clostridia;Lachnospirales;Lachnospiraceae;[Ruminococcus] gauvreauii group;uncultured bacterium | 92.2 | 1288 | -1 |
| metaRNA.PFemirge\_16650 | 215 | 0.000005 | EF096599.1.1399 | Bacteria;Firmicutes;Clostridia;Lachnospirales;Lachnospiraceae;Lachnospiraceae NK4A136 group;uncultured bacterium | 89.1 | 1301 | -1 |
| metaRNA.PFemirge\_19845 | 215 | 0.000019 | LN564853.1.1322 | Bacteria;Proteobacteria;Alphaproteobacteria;Rhizobiales;Rhizobiaceae;Shinella;uncultured bacterium | 93.3 | 1261 | -1 |
| metaRNA.PFemirge\_4015 | 214 | 0.000023 | FJ269095.1.1502 | Bacteria;Bacteroidota;Bacteroidia;Bacteroidales;Bacteroidaceae;Bacteroides;iron-reducing bacterium enrichment culture clone HN-HFO47 | 91.8 | 1435 | -1 |
| metaRNA.PFemirge\_18749 | 214 | 0.000101 | EU454443.1.1393 | Bacteria;Firmicutes;Clostridia;Lachnospirales;Lachnospiraceae;Lachnospiraceae UCG-008;uncultured bacterium | 91.0 | 1296 | -1 |
| metaRNA.PFemirge\_13667 | 213 | 0.000145 | CP003274.2181216.2182629 | Bacteria;Bacteroidota;Bacteroidia;Bacteroidales;Rikenellaceae;Alistipes;Alistipes finegoldii DSM 17242 | 91.6 | 1333 | -1 |
| metaRNA.PFemirge\_6283 | 213 | 0.000007 | HQ772765.1.1442 | Bacteria;Bacteroidota;Bacteroidia;Bacteroidales;Bacteroidaceae;Bacteroides;uncultured organism | 90.1 | 1406 | -1 |
| metaRNA.PFemirge\_12921 | 213 | 0.000010 | KF773008.1.1446 | Bacteria;Firmicutes;Clostridia;Lachnospirales;Lachnospiraceae;Hungatella;uncultured bacterium | 89.4 | 1308 | -1 |
| metaRNA.PFemirge\_5940 | 213 | 0.000090 | FP929032.881290.882801 | Bacteria;Bacteroidota;Bacteroidia;Bacteroidales;Rikenellaceae;Alistipes;Alistipes shahii WAL 8301 | 92.6 | 1484 | -1 |
| metaRNA.PFemirge\_10253 | 213 | 0.000037 | KM650565.1.1462 | Bacteria;Desulfobacterota;Desulfovibrionia;Desulfovibrionales;Desulfovibrionaceae;Desulfovibrio;uncultured proteobacterium | 90.5 | 1343 | -1 |
| metaRNA.PFemirge\_2566 | 213 | 0.000026 | ABYH01000014.137.1655 | Bacteria;Bacteroidota;Bacteroidia;Bacteroidales;Tannerellaceae;Parabacteroides;Parabacteroides johnsonii DSM 18315 | 93.8 | 1491 | -1 |
| metaRNA.PFemirge\_25886 | 212 | 0.000008 | KM360180.1.1492 | Bacteria;Firmicutes;Clostridia;Lachnospirales;Lachnospiraceae;Lachnoclostridium;Clostridium sp. BPY5 | 92.0 | 1279 | -1 |
| metaRNA.PFemirge\_12601 | 212 | 0.000023 | GALZ01007484.194.1638 | Bacteria;Cyanobacteria;Cyanobacteriia;Chloroplast;Actinidia chinensis | 96.8 | 1354 | -1 |
| metaRNA.PFemirge\_27313 | 212 | 0.000197 | EU466098.1.1227 | Bacteria;Firmicutes;Bacilli;Erysipelotrichales;Erysipelotrichaceae;Turicibacter;uncultured bacterium | 88.3 | 1227 | -1 |
| metaRNA.PFemirge\_6590 | 211 | 0.000028 | CCFF01000015.520926.522414 | Bacteria;Firmicutes;Clostridia;Clostridiales;Clostridiaceae;Clostridium sensu stricto 13;Clostridium sp. CL-6 | 99.0 | 1174 | -1 |
| metaRNA.PFemirge\_22117 | 210 | 0.000063 | EF400740.1.1495 | Bacteria;Firmicutes;Clostridia;Oscillospirales;Butyricicoccaceae;UCG-009;uncultured bacterium | 91.6 | 1268 | -1 |
| metaRNA.PFemirge\_1478 | 209 | 0.000009 | EF179838.1.1507 | Bacteria;Proteobacteria;Gammaproteobacteria;Enterobacterales;Aeromonadaceae;Aeromonas;uncultured bacterium | 94.7 | 1493 | -1 |
| metaRNA.PFemirge\_10811 | 209 | 0.000021 | JN012636.1.1457 | Bacteria;Firmicutes;Clostridia;Lachnospirales;Lachnospiraceae;uncultured;uncultured bacterium | 89.3 | 1286 | -1 |
| metaRNA.PFemirge\_24043 | 207 | 0.000018 | EU777150.1.1350 | Bacteria;Firmicutes;Clostridia;Lachnospirales;Lachnospiraceae;Lachnospiraceae UCG-004;uncultured bacterium | 89.2 | 1203 | -1 |
| metaRNA.PFemirge\_7939 | 207 | 0.000028 | AB034016.1.1515 | Bacteria;Cyanobacteria;Vampirivibrionia;Gastranaerophilales;uncultured rumen bacterium 4C0d-2 | 93.1 | 1468 | -1 |
| metaRNA.PFemirge\_18548 | 205 | 0.000043 | CBVB010000006.254456.255954 | Bacteria;Bacteroidota;Bacteroidia;Bacteroidales;Bacteroidaceae;Bacteroides;Bacteroidaceae bacterium MS4 | 92.5 | 1392 | -1 |
| metaRNA.PFemirge\_15795 | 204 | 0.000015 | EU453128.1.1406 | Bacteria;Firmicutes;Clostridia;Lachnospirales;Lachnospiraceae;Lachnospiraceae UCG-004;uncultured bacterium | 89.8 | 1416 | -1 |
| metaRNA.PFemirge\_11241 | 204 | 0.000058 | AGES01000014.73256.74770 | Bacteria;Bacteroidota;Bacteroidia;Bacteroidales;Tannerellaceae;Parabacteroides;Parabacteroides sp. HGS0025 | 91.1 | 1454 | -1 |
| metaRNA.PFemirge\_17428 | 204 | 0.000005 | DQ809267.1.1395 | Bacteria;Bacteroidota;Bacteroidia;Bacteroidales;Marinifilaceae;Butyricimonas;uncultured bacterium | 93.2 | 1393 | -1 |
| metaRNA.PFemirge\_25689 | 202 | 0.000012 | KP182152.1.1434 | Bacteria;Proteobacteria;Alphaproteobacteria;Rhizobiales;Rhizobiales Incertae Sedis;Phreatobacter;Phreatobacter sp. 61DPR27 | 93.4 | 1016 | -1 |
| metaRNA.PFemirge\_15940 | 202 | 0.000017 | KF704746.1.1383 | Bacteria;Proteobacteria;Alphaproteobacteria;Rhizobiales;Rhizobiaceae;Shinella;Shinella zoogloeoides | 96.6 | 1302 | -1 |
| metaRNA.PFemirge\_3431 | 202 | 0.000028 | JX576025.1.1509 | Bacteria;Proteobacteria;Gammaproteobacteria;Burkholderiales;Rhodocyclaceae;Azospira;uncultured Azospira sp. | 93.6 | 1486 | -1 |
| metaRNA.PFemirge\_15497 | 202 | 0.000007 | FJ658504.1.1402 | Bacteria;Proteobacteria;Gammaproteobacteria;Enterobacterales;Morganellaceae;Morganella;uncultured bacterium | 95.9 | 1397 | -1 |
| metaRNA.PFemirge\_20331 | 202 | 0.000022 | DQ799377.1.1384 | Bacteria;Firmicutes;Clostridia;Oscillospirales;Ruminococcaceae;Ruminococcus;uncultured bacterium | 92.4 | 1293 | -1 |
| metaRNA.PFemirge\_4671 | 201 | 0.000009 | HQ819316.1.1450 | Bacteria;Bacteroidota;Bacteroidia;Bacteroidales;Bacteroidaceae;Bacteroides;uncultured organism | 88.8 | 1243 | -1 |
| metaRNA.PFemirge\_21668 | 201 | 0.000001 | AM183061.1.1440 | Bacteria;Firmicutes;Clostridia;Clostridiales;Clostridiaceae;Clostridium sensu stricto 1;uncultured bacterium | 95.3 | 1292 | -1 |
| metaRNA.PFemirge\_15643 | 201 | 0.000014 | HM630228.1.1408 | Bacteria;Firmicutes;Clostridia;Clostridia vadinBB60 group;uncultured bacterium | 96.6 | 1222 | -1 |
| metaRNA.PFemirge\_24079 | 200 | 0.000050 | EF098038.1.1349 | Bacteria;Firmicutes;Clostridia;Lachnospirales;Lachnospiraceae;Lachnospiraceae NC2004 group;uncultured bacterium | 90.5 | 1121 | -1 |
| metaRNA.PFemirge\_12765 | 199 | 0.000006 | JX105679.1.1392 | Bacteria;Bacteroidota;Bacteroidia;Bacteroidales;Bacteroidaceae;Bacteroides;uncultured bacterium | 89.2 | 1240 | -1 |
| metaRNA.PFemirge\_27169 | 199 | 0.000020 | KJ615121.1.1238 | Bacteria;Proteobacteria;Alphaproteobacteria;Rhizobiales;Beijerinckiaceae;Bosea;uncultured bacterium | 94.2 | 1235 | -1 |
| metaRNA.PFemirge\_27760 | 198 | 0.000012 | DQ799425.1.1396 | Bacteria;Bacteroidota;Bacteroidia;Bacteroidales;Tannerellaceae;Parabacteroides;uncultured bacterium | 94.9 | 1397 | -1 |
| metaRNA.PFemirge\_22501 | 197 | 0.000008 | HQ761492.1.1450 | Bacteria;Firmicutes;Clostridia;Lachnospirales;Lachnospiraceae;[Ruminococcus] torques group;uncultured organism | 94.1 | 1308 | -1 |
| metaRNA.PFemirge\_16811 | 196 | 0.000008 | DQ797856.1.1399 | Bacteria;Bacteroidota;Bacteroidia;Bacteroidales;Rikenellaceae;Alistipes;uncultured bacterium | 91.0 | 1396 | -1 |
| metaRNA.PFemirge\_16462 | 195 | 0.000050 | EU765827.1.1360 | Bacteria;Bacteroidota;Bacteroidia;Bacteroidales;Bacteroidaceae;Bacteroides;uncultured bacterium | 91.8 | 1283 | -1 |
| metaRNA.PFemirge\_27718 | 195 | 0.000020 | AB214328.1.1484 | Bacteria;Bacteroidota;Bacteroidia;Bacteroidales;Bacteroidaceae;Bacteroides;Bacteroides intestinalis | 93.8 | 1442 | -1 |
| metaRNA.PFemirge\_15548 | 194 | 0.000132 | FJ269051.1.1487 | Bacteria;Bacteroidota;Bacteroidia;Bacteroidales;Bacteroidaceae;Bacteroides;iron-reducing bacterium enrichment culture clone HN11 | 89.1 | 1409 | -1 |
| metaRNA.PFemirge\_6786 | 193 | 0.000015 | HM630236.1.1451 | Bacteria;Bacteroidota;Bacteroidia;Bacteroidales;Rikenellaceae;dgA-11 gut group;uncultured bacterium | 89.7 | 1315 | -1 |
| metaRNA.PFemirge\_1217 | 192 | 0.000014 | AM490748.1.1504 | Bacteria;Myxococcota;Polyangia;mle1-27;uncultured bacterium | 96.0 | 1362 | -1 |
| metaRNA.PFemirge\_99 | 191 | 0.000013 | M97959.3.1817 | Eukaryota;SAR;Stramenopiles;Ochrophyta;Phaeophyceae;Fucales;Fucus;Fucus distichus | 99.2 | 1672 | -1 |
| metaRNA.PFemirge\_12199 | 189 | 0.000012 | GQ898302.1.1489 | Bacteria;Bacteroidota;Bacteroidia;Bacteroidales;Bacteroidaceae;Bacteroides;uncultured bacterium | 91.1 | 1419 | -1 |
| metaRNA.PFemirge\_13618 | 188 | 0.000005 | GQ480160.1.1510 | Bacteria;Firmicutes;Negativicutes;Acidaminococcales;Acidaminococcaceae;Succinispira;uncultured bacterium | 89.9 | 1371 | -1 |
| metaRNA.PFemirge\_21700 | 188 | 0.000030 | EU466086.1.1375 | Bacteria;Firmicutes;Clostridia;Lachnospirales;Lachnospiraceae;Dorea;uncultured bacterium | 92.4 | 1286 | -1 |
| metaRNA.PFemirge\_12901 | 186 | 0.000019 | CP012937.2383895.2385420 | Bacteria;Bacteroidota;Bacteroidia;Bacteroidales;Bacteroidaceae;Bacteroides;Bacteroides thetaiotaomicron | 92.0 | 1379 | -1 |
| metaRNA.PFemirge\_9589 | 184 | 0.000015 | KP687196.1.1423 | Bacteria;Proteobacteria;Alphaproteobacteria;Holosporales;Holosporaceae;uncultured;uncultured bacterium | 89.2 | 1253 | -1 |
| metaRNA.PFemirge\_26443 | 182 | 0.000017 | HQ385555.1.1318 | Bacteria;Proteobacteria;Alphaproteobacteria;Rhodobacterales;Rhodobacteraceae;Rhodobacter;uncultured bacterium | 96.1 | 1207 | -1 |
| metaRNA.PFemirge\_23786 | 182 | 0.000024 | KC331222.1.1485 | Bacteria;Actinobacteriota;Actinobacteria;Micrococcales;Microbacteriaceae;Leucobacter;uncultured bacterium | 93.6 | 1325 | -1 |
| metaRNA.PFemirge\_26911 | 182 | 0.000000 | EU775615.1.1257 | Bacteria;Firmicutes;Clostridia;Peptostreptococcales-Tissierellales;Peptostreptococcaceae;Romboutsia;uncultured bacterium | 96.7 | 1153 | -1 |
| metaRNA.PFemirge\_16289 | 179 | 0.000095 | JX105679.1.1392 | Bacteria;Bacteroidota;Bacteroidia;Bacteroidales;Bacteroidaceae;Bacteroides;uncultured bacterium | 93.7 | 1401 | -1 |
| metaRNA.PFemirge\_11095 | 179 | 0.000044 | HQ756569.1.1448 | Bacteria;Bacteroidota;Bacteroidia;Bacteroidales;Bacteroidaceae;Bacteroides;uncultured organism | 88.7 | 1447 | -1 |
| metaRNA.PFemirge\_22786 | 178 | 0.000038 | LT725663.1.1450 | Bacteria;Bacteroidota;Bacteroidia;Bacteroidales;Tannerellaceae;Parabacteroides;Parabacteroides sp. Marseille-P3668 | 93.2 | 1348 | -1 |
| metaRNA.PFemirge\_6032 | 178 | 0.000096 | EU663602.1.1492 | Bacteria;Bacteroidota;Bacteroidia;Bacteroidales;Marinifilaceae;uncultured;uncultured bacterium | 91.2 | 1492 | -1 |
| metaRNA.PFemirge\_18220 | 178 | 0.000008 | AB746664.1.1536 | Bacteria;Firmicutes;Clostridia;Lachnospirales;Lachnospiraceae;Lachnospiraceae NK4A136 group;uncultured bacterium | 88.5 | 1310 | -1 |
| metaRNA.PFemirge\_20191 | 178 | 0.000004 | DQ805627.1.1385 | Bacteria;Firmicutes;Clostridia;Lachnospirales;Lachnospiraceae;Lachnospiraceae UCG-004;uncultured bacterium | 91.0 | 1378 | -1 |
| metaRNA.PFemirge\_11867 | 176 | 0.000139 | U81750.1.1482 | Bacteria;Firmicutes;Negativicutes;Acidaminococcales;Acidaminococcaceae;uncultured;uncultured bacterium | 90.5 | 1362 | -1 |
| metaRNA.PFemirge\_24922 | 175 | 0.000023 | LC094617.1.1418 | Bacteria;Actinobacteriota;Actinobacteria;Micrococcales;Microbacteriaceae;Leifsonia;Microbacteriaceae bacterium MAEY21 | 95.2 | 1217 | -1 |
| metaRNA.PFemirge\_9053 | 174 | 0.000013 | FPLP01013193.1.1403 | Bacteria;Cyanobacteria;Vampirivibrionia;Obscuribacterales;Obscuribacteraceae;Candidatus Obscuribacter;metagenome | 97.7 | 1279 | -1 |
| metaRNA.PFemirge\_21157 | 174 | 0.000045 | EU542517.1.1444 | Bacteria;Firmicutes;Clostridia;Oscillospirales;Oscillospiraceae;Intestinimonas;uncultured bacterium | 92.6 | 1156 | -1 |
| metaRNA.PFemirge\_26159 | 174 | 0.000042 | AB721051.1.1697 | Eukaryota;Amorphea;Amoebozoa;SAR;Stramenopiles;Labyrinthulomycetes;Sorodiplophrys;uncultured freshwater eukaryote | 98.2 | 446 | -1 |
| metaRNA.PFemirge\_19219 | 174 | 0.000015 | FR749936.1.1524 | Bacteria;Firmicutes;Clostridia;Lachnospirales;Lachnospiraceae;[Eubacterium] fissicatena group;Faecalicatena fissicatena | 94.2 | 1386 | -1 |
| metaRNA.PFemirge\_13109 | 173 | 0.000020 | KY122019.1.1428 | Bacteria;Proteobacteria;Alphaproteobacteria;Rhizobiales;Rhizobiaceae;Mesorhizobium;Mesorhizobium sp. | 89.7 | 1430 | -1 |
| metaRNA.PFemirge\_24271 | 172 | 0.000007 | LC094617.1.1418 | Bacteria;Actinobacteriota;Actinobacteria;Micrococcales;Microbacteriaceae;Leifsonia;Microbacteriaceae bacterium MAEY21 | 90.6 | 1299 | -1 |
| metaRNA.PFemirge\_28041 | 171 | 0.000006 | DQ824296.1.1398 | Bacteria;Bacteroidota;Bacteroidia;Bacteroidales;Bacteroidaceae;Bacteroides;uncultured bacterium | 92.4 | 1183 | -1 |
| metaRNA.PFemirge\_23661 | 166 | 0.000023 | AB486052.1.1370 | Bacteria;Firmicutes;Clostridia;Oscillospirales;Oscillospiraceae;Colidextribacter;uncultured bacterium | 92.3 | 1277 | -1 |
| metaRNA.PFemirge\_13981 | 166 | 0.000054 | GQ448213.1.1432 | Bacteria;Firmicutes;Clostridia;Lachnospirales;Lachnospiraceae;Lachnospiraceae NK4A136 group;uncultured bacterium | 92.6 | 1313 | -1 |
| metaRNA.PFemirge\_19805 | 166 | 0.000004 | EF400740.1.1495 | Bacteria;Firmicutes;Clostridia;Oscillospirales;Butyricicoccaceae;UCG-009;uncultured bacterium | 93.4 | 1215 | -1 |
| metaRNA.PFemirge\_16300 | 164 | 0.000027 | DQ809036.1.1400 | Bacteria;Bacteroidota;Bacteroidia;Bacteroidales;Bacteroidaceae;Bacteroides;uncultured bacterium | 91.6 | 1332 | -1 |
| metaRNA.PFemirge\_11100 | 164 | 0.000006 | HQ783986.1.1437 | Bacteria;Bacteroidota;Bacteroidia;Bacteroidales;Tannerellaceae;Parabacteroides;uncultured organism | 89.1 | 1400 | -1 |
| metaRNA.PFemirge\_13330 | 163 | 0.000023 | LC028724.1.1522 | Bacteria;Firmicutes;Clostridia;Lachnospirales;Lachnospiraceae;Lachnospiraceae FCS020 group;uncultured bacterium | 92.6 | 1295 | -1 |
| metaRNA.PFemirge\_1864 | 163 | 0.000004 | GQ480160.1.1510 | Bacteria;Firmicutes;Negativicutes;Acidaminococcales;Acidaminococcaceae;Succinispira;uncultured bacterium | 90.0 | 1432 | -1 |
| metaRNA.PFemirge\_12490 | 162 | 0.000030 | JQ940836.1.1449 | Bacteria;Bacteroidota;Bacteroidia;Bacteroidales;Rikenellaceae;Alistipes;uncultured bacterium | 87.7 | 1304 | -1 |
| metaRNA.PFemirge\_7126 | 161 | 0.000043 | AB916502.1.1486 | Bacteria;Bacteroidota;Bacteroidia;Bacteroidales;Marinifilaceae;Butyricimonas;Butyricimonas paravirosa | 90.5 | 1392 | -1 |
| metaRNA.PFemirge\_2918 | 161 | 0.000022 | GQ871719.1.1509 | Bacteria;Firmicutes;Bacilli;Erysipelotrichales;Erysipelotrichaceae;Faecalitalea;uncultured bacterium | 92.5 | 1265 | -1 |
| metaRNA.PFemirge\_23861 | 161 | 0.000009 | HQ143300.1.1507 | Bacteria;Proteobacteria;Gammaproteobacteria;Enterobacterales;Enterobacteriaceae;Escherichia-Shigella;uncultured bacterium | 93.3 | 1356 | -1 |
| metaRNA.PFemirge\_9530 | 161 | 0.000005 | CP022151.430175.431728 | Bacteria;Proteobacteria;Gammaproteobacteria;Enterobacterales;Enterobacteriaceae;Citrobacter;Citrobacter freundii | 94.8 | 1354 | -1 |
| metaRNA.PFemirge\_6842 | 160 | 0.000023 | DQ238611.1.1497 | Bacteria;Firmicutes;Clostridia;Clostridiales;Clostridiaceae;Clostridium sensu stricto 1;uncultured bacterium | 94.5 | 1465 | -1 |
| metaRNA.PFemirge\_18543 | 159 | 0.000023 | EU937930.1.1450 | Bacteria;Proteobacteria;Alphaproteobacteria;Sphingomonadales;Sphingomonadaceae;Novosphingobium;uncultured bacterium | 93.1 | 1341 | -1 |
| metaRNA.PFemirge\_19358 | 159 | 0.000346 | JQ248079.1.1513 | Bacteria;Firmicutes;Clostridia;Oscillospirales;Oscillospiraceae;Flavonifractor;uncultured bacterium | 93.2 | 1237 | -1 |
| metaRNA.PFemirge\_23640 | 159 | 0.000013 | GQ898343.1.1489 | Bacteria;Bacteroidota;Bacteroidia;Bacteroidales;Bacteroidaceae;Bacteroides;uncultured bacterium | 93.0 | 1365 | -1 |
| metaRNA.PFemirge\_16443 | 156 | 0.000064 | CP012801.6655317.6656834 | Bacteria;Bacteroidota;Bacteroidia;Bacteroidales;Bacteroidaceae;Bacteroides;Bacteroides cellulosilyticus | 90.2 | 1242 | -1 |
| metaRNA.PFemirge\_22745 | 156 | 0.000012 | JX105679.1.1392 | Bacteria;Bacteroidota;Bacteroidia;Bacteroidales;Bacteroidaceae;Bacteroides;uncultured bacterium | 91.0 | 1330 | -1 |
| metaRNA.PFemirge\_15856 | 156 | 0.000016 | EF024902.1.1792 | Eukaryota;Archaeplastida;Chloroplastida;Charophyta;Phragmoplastophyta;Streptophyta;Embryophyta;Tracheophyta;Spermatophyta;Magnoliophyta;Phaseoleae environmental sample | 95.0 | 776 | -1 |
| metaRNA.PFemirge\_4007 | 155 | 0.000023 | EU560783.1.1502 | Bacteria;Proteobacteria;Gammaproteobacteria;Enterobacterales;Enterobacteriaceae;Escherichia-Shigella;uncultured bacterium | 95.0 | 1396 | -1 |
| metaRNA.PFemirge\_28079 | 155 | 0.000005 | EU775962.1.1393 | Bacteria;Bacteroidota;Bacteroidia;Bacteroidales;Bacteroidaceae;Bacteroides;uncultured bacterium | 87.1 | 1300 | -1 |
| metaRNA.PFemirge\_5226 | 155 | 0.000012 | KC633480.1.1496 | Bacteria;Proteobacteria;Gammaproteobacteria;Burkholderiales;Comamonadaceae;Acidovorax;uncultured beta proteobacterium | 97.0 | 1397 | -1 |
| metaRNA.PFemirge\_21077 | 155 | 0.000002 | EF097622.1.1400 | Bacteria;Firmicutes;Clostridia;Oscillospirales;Oscillospiraceae;Colidextribacter;uncultured bacterium | 94.0 | 1131 | -1 |
| metaRNA.PFemirge\_16056 | 154 | 0.000004 | DQ809036.1.1400 | Bacteria;Bacteroidota;Bacteroidia;Bacteroidales;Bacteroidaceae;Bacteroides;uncultured bacterium | 90.5 | 1402 | -1 |
| metaRNA.PFemirge\_30252 | 154 | 0.000015 | DQ807465.1.1395 | Bacteria;Firmicutes;Clostridia;Oscillospirales;Oscillospiraceae;uncultured;uncultured bacterium | 86.4 | 1370 | -1 |
| metaRNA.PFemirge\_15909 | 152 | 0.000018 | HQ385613.1.1405 | Bacteria;Proteobacteria;Gammaproteobacteria;Gammaproteobacteria Incertae Sedis;Unknown Family;uncultured bacterium | 94.7 | 1402 | -1 |
| metaRNA.PFemirge\_6131 | 152 | 0.000014 | JN688021.1.1446 | Bacteria;Bacteroidota;Bacteroidia;Bacteroidales;Rikenellaceae;uncultured;bacterium enrichment culture clone M02 | 88.4 | 1451 | -1 |
| metaRNA.PFemirge\_21898 | 152 | 0.000007 | HM556834.1.1374 | Bacteria;Cyanobacteria;Cyanobacteriia;Chloroplast;uncultured bacterium | 98.1 | 1374 | -1 |
| metaRNA.PFemirge\_14600 | 151 | 0.000012 | JF824799.1.1423 | Bacteria;Bacteroidota;Bacteroidia;Bacteroidales;Rikenellaceae;Alistipes;Alistipes timonensis JC136 | 92.1 | 1386 | -1 |
| metaRNA.PFemirge\_5013 | 151 | 0.000028 | KC836026.1.1497 | Bacteria;Proteobacteria;Gammaproteobacteria;Burkholderiales;Rhodocyclaceae;Dechlorobacter;uncultured bacterium | 97.6 | 1414 | -1 |
| metaRNA.PFemirge\_24245 | 151 | 0.000002 | EU777145.1.1347 | Bacteria;Firmicutes;Clostridia;Christensenellales;Christensenellaceae;Christensenellaceae R-7 group;uncultured bacterium | 92.9 | 1279 | -1 |
| metaRNA.PFemirge\_17158 | 149 | 0.000007 | FQSE01000007.1631802.1633308 | Bacteria;Bacteroidota;Bacteroidia;Bacteroidales;Tannerellaceae;Parabacteroides;Parabacteroides sp. Marseille-P3136 | 92.7 | 1396 | -1 |
| metaRNA.PFemirge\_16875 | 149 | 0.000021 | HQ785173.1.1448 | Bacteria;Bacteroidota;Bacteroidia;Bacteroidales;Bacteroidaceae;Bacteroides;uncultured organism | 88.5 | 1221 | -1 |
| metaRNA.PFemirge\_24983 | 149 | 0.000020 | CDZJ01005172.1679.3161 | Bacteria;Proteobacteria;Alphaproteobacteria;Rhodospirillales;uncultured;gut metagenome | 83.9 | 1294 | -1 |
| metaRNA.PFemirge\_16949 | 148 | 0.000007 | HQ761452.1.1440 | Bacteria;Bacteroidota;Bacteroidia;Bacteroidales;Tannerellaceae;Parabacteroides;uncultured organism | 89.4 | 1299 | -1 |
| metaRNA.PFemirge\_23134 | 148 | 0.000040 | EU773851.1.1360 | Bacteria;Firmicutes;Clostridia;Peptostreptococcales-Tissierellales;Peptostreptococcaceae;uncultured;uncultured bacterium | 92.9 | 1358 | -1 |
| metaRNA.PFemirge\_6554 | 146 | 0.000060 | HM630239.1.1414 | Bacteria;Firmicutes;Bacilli;Erysipelotrichales;Erysipelatoclostridiaceae;Erysipelatoclostridium;uncultured bacterium | 91.1 | 1241 | -1 |
| metaRNA.PFemirge\_17352 | 146 | 0.000022 | HQ681866.1.1451 | Bacteria;Bacteroidota;Bacteroidia;Bacteroidales;Bacteroidaceae;Bacteroides;uncultured bacterium | 92.4 | 1344 | -1 |
| metaRNA.PFemirge\_18986 | 145 | 0.000116 | DQ798058.1.1390 | Bacteria;Firmicutes;Clostridia;Lachnospirales;Lachnospiraceae;[Ruminococcus] torques group;uncultured bacterium | 90.6 | 1387 | -1 |
| metaRNA.PFemirge\_26799 | 145 | 0.000024 | DQ808071.1.1266 | Bacteria;Firmicutes;Negativicutes;Acidaminococcales;Acidaminococcaceae;Phascolarctobacterium;uncultured bacterium | 91.5 | 1220 | -1 |
| metaRNA.PFemirge\_16774 | 144 | 0.000075 | GQ448331.1.1393 | Bacteria;Bacteroidota;Bacteroidia;Bacteroidales;Bacteroidaceae;Bacteroides;uncultured bacterium | 89.5 | 1234 | -1 |
| metaRNA.PFemirge\_16901 | 144 | 0.000019 | EU511065.1.1410 | Bacteria;Firmicutes;Clostridia;Oscillospirales;Oscillospiraceae;uncultured;uncultured bacterium | 93.5 | 1394 | -1 |
| metaRNA.PFemirge\_30246 | 143 | 0.000023 | EF402450.1.1486 | Bacteria;Bacteroidota;Bacteroidia;Bacteroidales;Bacteroidaceae;Bacteroides;uncultured bacterium | 86.4 | 1284 | -1 |
| metaRNA.PFemirge\_7308 | 143 | 0.000010 | FR749935.1.1522 | Bacteria;Firmicutes;Clostridia;Lachnospirales;Lachnospiraceae;[Eubacterium] fissicatena group;Faecalicatena fissicatena | 92.7 | 1368 | -1 |
| metaRNA.PFemirge\_26340 | 142 | 0.000017 | HM630205.1.1449 | Bacteria;Bacteroidota;Bacteroidia;Bacteroidales;Rikenellaceae;Rikenella;uncultured bacterium | 88.1 | 1285 | -1 |
| metaRNA.PFemirge\_27279 | 142 | 0.000031 | LC094617.1.1418 | Bacteria;Actinobacteriota;Actinobacteria;Micrococcales;Microbacteriaceae;Leifsonia;Microbacteriaceae bacterium MAEY21 | 93.9 | 1157 | -1 |
| metaRNA.PFemirge\_27638 | 142 | 0.000027 | GU935303.1.1474 | Bacteria;Firmicutes;Bacilli;Bacillales;Bacillaceae;Bacillus;Bacillus sp. EK-I67 | 92.0 | 915 | -1 |
| metaRNA.PFemirge\_25952 | 142 | 0.000006 | CP022754.4692906.4694445 | Bacteria;Bacteroidota;Bacteroidia;Bacteroidales;Tannerellaceae;Parabacteroides;Parabacteroides sp. CT06 | 92.4 | 828 | -1 |
| metaRNA.PFemirge\_25902 | 141 | 0.000006 | EU767529.1.1356 | Bacteria;Bacteroidota;Bacteroidia;Bacteroidales;Bacteroidaceae;Bacteroides;uncultured bacterium | 88.3 | 1182 | -1 |
| metaRNA.PFemirge\_634 | 141 | 0.000009 | FM177879.1.1437 | Bacteria;Proteobacteria;Alphaproteobacteria;Rhizobiales;Rhizobiaceae;Shinella;Shinella fusca | 82.5 | 1552 | -1 |
| metaRNA.PFemirge\_23074 | 140 | 0.000021 | HQ761492.1.1450 | Bacteria;Firmicutes;Clostridia;Lachnospirales;Lachnospiraceae;[Ruminococcus] torques group;uncultured organism | 92.3 | 1239 | -1 |
| metaRNA.PFemirge\_14455 | 140 | 0.000015 | HQ818958.1.1459 | Bacteria;Bacteroidota;Bacteroidia;Bacteroidales;Bacteroidaceae;Bacteroides;uncultured organism | 93.7 | 1324 | -1 |
| metaRNA.PFemirge\_26028 | 140 | 0.000012 | HQ775105.1.1456 | Bacteria;Firmicutes;Clostridia;Lachnospirales;Lachnospiraceae;Hungatella;uncultured organism | 88.3 | 1270 | -1 |
| metaRNA.PFemirge\_16157 | 140 | 0.000021 | DQ794175.1.1406 | Bacteria;Bacteroidota;Bacteroidia;Bacteroidales;Bacteroidaceae;Bacteroides;uncultured bacterium | 93.4 | 1400 | -1 |
| metaRNA.PFemirge\_10947 | 139 | 0.000006 | DQ355176.1.1464 | Bacteria;Desulfobacterota;Desulfovibrionia;Desulfovibrionales;Desulfovibrionaceae;Desulfovibrio;uncultured Desulfovibrio sp. | 90.9 | 1384 | -1 |
| metaRNA.PFemirge\_15966 | 139 | 0.000007 | FJ534957.1.1501 | Bacteria;Firmicutes;Clostridia;Lachnospirales;Lachnospiraceae;Lachnospiraceae FCS020 group;uncultured bacterium | 92.9 | 1402 | -1 |
| metaRNA.PFemirge\_23284 | 139 | 0.000031 | KP944118.1.1420 | Bacteria;Bacteroidota;Bacteroidia;Bacteroidales;Bacteroidaceae;Bacteroides;Bacteroides thetaiotaomicron | 93.6 | 1304 | -1 |
| metaRNA.PFemirge\_16459 | 138 | 0.000006 | DQ807222.1.1401 | Bacteria;Bacteroidota;Bacteroidia;Bacteroidales;Bacteroidaceae;Bacteroides;uncultured bacterium | 90.6 | 1404 | -1 |
| metaRNA.PFemirge\_18878 | 137 | 0.000005 | EU775230.1.1391 | Bacteria;Firmicutes;Clostridia;Lachnospirales;Lachnospiraceae;Lachnospiraceae AC2044 group;uncultured bacterium | 92.3 | 1293 | -1 |
| metaRNA.PFemirge\_30231 | 136 | 0.000042 | CBVB010000006.254456.255954 | Bacteria;Bacteroidota;Bacteroidia;Bacteroidales;Bacteroidaceae;Bacteroides;Bacteroidaceae bacterium MS4 | 91.6 | 1322 | -1 |
| metaRNA.PFemirge\_23226 | 135 | 0.000034 | DQ809036.1.1400 | Bacteria;Bacteroidota;Bacteroidia;Bacteroidales;Bacteroidaceae;Bacteroides;uncultured bacterium | 91.9 | 1303 | -1 |
| metaRNA.PFemirge\_2056 | 135 | 0.000009 | DQ817541.1.1479 | Bacteria;Proteobacteria;Gammaproteobacteria;Burkholderiales;Chromobacteriaceae;Crenobacter;uncultured bacterium | 95.3 | 1417 | -1 |
| metaRNA.PFemirge\_12036 | 133 | 0.000018 | AGES01000014.73256.74770 | Bacteria;Bacteroidota;Bacteroidia;Bacteroidales;Tannerellaceae;Parabacteroides;Parabacteroides sp. HGS0025 | 92.4 | 1451 | -1 |
| metaRNA.PFemirge\_25582 | 133 | 0.000010 | CP020388.637686.639237 | Bacteria;Proteobacteria;Gammaproteobacteria;Enterobacterales;Enterobacteriaceae;Pluralibacter;Pluralibacter gergoviae | 81.4 | 1492 | -1 |
| metaRNA.PFemirge\_4824 | 133 | 0.000005 | KX589545.1.1498 | Bacteria;Firmicutes;Bacilli;Alicyclobacillales;Alicyclobacillaceae;Tumebacillus;uncultured bacterium | 90.0 | 1427 | -1 |
| metaRNA.PFemirge\_23650 | 133 | 0.000000 | EU776022.1.1355 | Bacteria;Firmicutes;Clostridia;Peptostreptococcales-Tissierellales;Peptostreptococcaceae;uncultured;uncultured bacterium | 93.1 | 1343 | -1 |
| metaRNA.PFemirge\_16323 | 133 | 0.000005 | EU461102.1.1401 | Bacteria;Firmicutes;Clostridia;Lachnospirales;Lachnospiraceae;Dorea;uncultured bacterium | 88.1 | 1398 | -1 |
| metaRNA.PFemirge\_24401 | 132 | 0.000007 | HQ761612.1.1444 | Bacteria;Bacteroidota;Bacteroidia;Bacteroidales;Tannerellaceae;Parabacteroides;uncultured organism | 91.8 | 1135 | -1 |
| metaRNA.PFemirge\_13374 | 132 | 0.000003 | HQ762410.1.1446 | Bacteria;Bacteroidota;Bacteroidia;Bacteroidales;Tannerellaceae;Parabacteroides;uncultured organism | 90.8 | 1442 | -1 |
| metaRNA.PFemirge\_14889 | 132 | 0.000011 | EU834774.1.1419 | Bacteria;Proteobacteria;Alphaproteobacteria;Rhizobiales;Rhizobiales Incertae Sedis;Phreatobacter;uncultured bacterium | 96.0 | 1390 | -1 |
| metaRNA.PFemirge\_10712 | 131 | 0.000005 | JX198651.1.1513 | Bacteria;Firmicutes;Clostridia;Oscillospirales;Butyricicoccaceae;UCG-009;uncultured bacterium | 90.7 | 1329 | -1 |
| metaRNA.PFemirge\_26364 | 131 | 0.000031 | EU775261.1.1286 | Bacteria;Firmicutes;Clostridia;Lachnospirales;Lachnospiraceae;Agathobacter;uncultured bacterium | 91.8 | 1139 | -1 |
| metaRNA.PFemirge\_5136 | 131 | 0.000008 | FM213077.1.1512 | Bacteria;Proteobacteria;Gammaproteobacteria;Xanthomonadales;Rhodanobacteraceae;Tahibacter;uncultured bacterium | 97.0 | 1398 | -1 |
| metaRNA.PFemirge\_18380 | 131 | 0.000005 | EU461103.1.1392 | Bacteria;Firmicutes;Clostridia;Lachnospirales;Lachnospiraceae;Lachnospiraceae UCG-008;uncultured bacterium | 92.5 | 1394 | -1 |
| metaRNA.PFemirge\_11693 | 130 | 0.000022 | DQ800054.1.1388 | Bacteria;Firmicutes;Clostridia;Oscillospirales;Oscillospiraceae;uncultured;uncultured bacterium | 91.3 | 1299 | -1 |
| metaRNA.PFemirge\_23725 | 128 | 0.000005 | AATF01000761.367.1874 | Bacteria;Firmicutes;Clostridia;Oscillospirales;Ruminococcaceae;Ruminococcus;mouse gut metagenome | 89.9 | 1349 | -1 |
| metaRNA.PFemirge\_25608 | 128 | 0.000012 | EU776025.1.1313 | Bacteria;Firmicutes;Bacilli;Erysipelotrichales;Erysipelotrichaceae;Holdemanella;uncultured bacterium | 91.0 | 1307 | -1 |
| metaRNA.PFemirge\_7333 | 128 | 0.000021 | EF400726.1.1488 | Bacteria;Bacteroidota;Bacteroidia;Bacteroidales;Marinifilaceae;Odoribacter;uncultured bacterium | 90.6 | 1467 | -1 |
| metaRNA.PFemirge\_8548 | 128 | 0.000006 | EF404130.1.1478 | Bacteria;Firmicutes;Clostridia;Oscillospirales;Ruminococcaceae;Ruminococcus;uncultured bacterium | 91.7 | 1348 | -1 |
| metaRNA.PFemirge\_8249 | 126 | 0.000024 | DQ816804.1.1464 | Bacteria;Firmicutes;Clostridia;Peptostreptococcales-Tissierellales;Anaerovoracaceae;Anaerovorax;uncultured bacterium | 95.8 | 1305 | -1 |
| metaRNA.PFemirge\_10119 | 125 | 0.000008 | AY721628.1.1491 | Bacteria;Desulfobacterota;Desulfovibrionia;Desulfovibrionales;Desulfovibrionaceae;Desulfovibrio;bacterium 'New Zealand D' | 93.1 | 1443 | -1 |
| metaRNA.PFemirge\_7109 | 124 | 0.000012 | GQ448528.1.1397 | Bacteria;Firmicutes;Clostridia;Clostridia UCG-014;uncultured bacterium | 95.0 | 1150 | -1 |
| metaRNA.PFemirge\_11222 | 124 | 0.000001 | DQ673521.1.1485 | Bacteria;Firmicutes;Clostridia;Lachnospirales;Lachnospiraceae;Lachnospiraceae NK4A136 group;uncultured rumen bacterium | 88.8 | 1372 | -1 |
| metaRNA.PFemirge\_28136 | 124 | 0.000016 | EU772907.1.1401 | Bacteria;Bacteroidota;Bacteroidia;Bacteroidales;Bacteroidaceae;Bacteroides;uncultured bacterium | 90.1 | 1389 | -1 |
| metaRNA.PFemirge\_21594 | 124 | 0.000009 | EU778426.1.1385 | Bacteria;Firmicutes;Clostridia;Clostridiales;Clostridiaceae;Clostridium sensu stricto 1;uncultured bacterium | 93.5 | 1375 | -1 |
| metaRNA.PFemirge\_16959 | 123 | 0.000007 | JF709929.1.1398 | Bacteria;Firmicutes;Bacilli;Exiguobacterales;Exiguobacteraceae;Exiguobacterium;uncultured bacterium | 97.0 | 1337 | -1 |
| metaRNA.PFemirge\_11902 | 123 | 0.000016 | HQ785173.1.1448 | Bacteria;Bacteroidota;Bacteroidia;Bacteroidales;Bacteroidaceae;Bacteroides;uncultured organism | 90.2 | 1345 | -1 |
| metaRNA.PFemirge\_24781 | 123 | 0.000004 | DQ805650.1.1385 | Bacteria;Firmicutes;Clostridia;Lachnospirales;Lachnospiraceae;Lachnospiraceae UCG-008;uncultured bacterium | 92.4 | 1258 | -1 |
| metaRNA.PFemirge\_3590 | 123 | 0.000010 | HQ748426.1.1449 | Bacteria;Firmicutes;Clostridia;Lachnospirales;Lachnospiraceae;Lachnoclostridium;uncultured organism | 90.7 | 1330 | -1 |
| metaRNA.PFemirge\_4086 | 122 | 0.000010 | U14932.1.1433 | Bacteria;Spirochaetota;Brachyspirae;Brachyspirales;Brachyspiraceae;Brachyspira;Brachyspira hyodysenteriae | 95.9 | 1281 | -1 |
| metaRNA.PFemirge\_17488 | 121 | 0.000014 | DQ856503.1.1485 | Bacteria;Bacteroidota;Bacteroidia;Bacteroidales;Bacteroidaceae;Bacteroides;uncultured bacterium | 92.0 | 1246 | -1 |
| metaRNA.PFemirge\_7079 | 121 | 0.000003 | AB574479.1.1484 | Bacteria;Bacteroidota;Bacteroidia;Bacteroidales;Bacteroidaceae;Bacteroides;Bacteroides stercorirosoris | 89.6 | 1271 | -1 |
| metaRNA.PFemirge\_11104 | 121 | 0.000018 | HQ774886.1.1445 | Bacteria;Bacteroidota;Bacteroidia;Bacteroidales;Barnesiellaceae;Coprobacter;uncultured organism | 88.0 | 1295 | -1 |
| metaRNA.PFemirge\_19772 | 120 | 0.000019 | EU776035.1.1400 | Bacteria;Proteobacteria;Gammaproteobacteria;Enterobacterales;Enterobacteriaceae;Escherichia-Shigella;uncultured bacterium | 93.4 | 1385 | -1 |
| metaRNA.PFemirge\_7175 | 120 | 0.000004 | AB470343.1.1485 | Bacteria;Bacteroidota;Bacteroidia;Bacteroidales;Tannerellaceae;Parabacteroides;Parabacteroides gordonii | 89.1 | 1411 | -1 |
| metaRNA.PFemirge\_4771 | 120 | 0.000016 | JN391840.1.1499 | Bacteria;Proteobacteria;Gammaproteobacteria;Burkholderiales;Comamonadaceae;uncultured;uncultured bacterium | 95.9 | 1476 | -1 |
| metaRNA.PFemirge\_20173 | 120 | 0.000015 | DQ795028.1.1385 | Bacteria;Firmicutes;Clostridia;Oscillospirales;Oscillospiraceae;UCG-002;uncultured bacterium | 93.2 | 1379 | -1 |
| metaRNA.PFemirge\_13173 | 119 | 0.000011 | HQ761613.1.1445 | Bacteria;Bacteroidota;Bacteroidia;Bacteroidales;Bacteroidaceae;Bacteroides;uncultured organism | 91.8 | 1261 | -1 |
| metaRNA.PFemirge\_18287 | 119 | 0.000026 | DQ807591.1.1392 | Bacteria;Bacteroidota;Bacteroidia;Bacteroidales;Bacteroidaceae;Bacteroides;uncultured bacterium | 90.6 | 1390 | -1 |
| metaRNA.PFemirge\_6300 | 118 | 0.000007 | FN428761.1.1496 | Bacteria;Bacteroidota;Bacteroidia;Chitinophagales;Chitinophagaceae;Edaphobaculum;uncultured bacterium | 96.1 | 1345 | -1 |
| metaRNA.PFemirge\_23421 | 118 | 0.000006 | JQ940836.1.1449 | Bacteria;Bacteroidota;Bacteroidia;Bacteroidales;Rikenellaceae;Alistipes;uncultured bacterium | 92.8 | 1339 | -1 |
| metaRNA.PFemirge\_16724 | 117 | 0.000006 | EU775673.1.1399 | Bacteria;Proteobacteria;Gammaproteobacteria;Enterobacterales;Enterobacteriaceae;Escherichia-Shigella;uncultured bacterium | 94.0 | 1394 | -1 |
| metaRNA.PFemirge\_9028 | 117 | 0.000019 | AB552872.1.1469 | Bacteria;Verrucomicrobiota;Verrucomicrobiae;Verrucomicrobiales;Verrucomicrobiaceae;Brevifollis;Brevifollis gellanilyticus | 96.5 | 1354 | -1 |
| metaRNA.PFemirge\_12790 | 117 | 0.000229 | JX198651.1.1513 | Bacteria;Firmicutes;Clostridia;Oscillospirales;Butyricicoccaceae;UCG-009;uncultured bacterium | 91.9 | 1325 | -1 |
| metaRNA.PFemirge\_23392 | 117 | 0.000003 | HQ756942.1.1450 | Bacteria;Firmicutes;Clostridia;Lachnospirales;Lachnospiraceae;[Ruminococcus] gnavus group;uncultured organism | 92.3 | 1292 | -1 |
| metaRNA.PFemirge\_13214 | 117 | 0.000008 | EU283367.1.1440 | Bacteria;Bacteroidota;Bacteroidia;Chitinophagales;Saprospiraceae;Haliscomenobacter;uncultured Bacteroidetes bacterium | 92.8 | 1369 | -1 |
| metaRNA.PFemirge\_500 | 116 | 0.000003 | GQ480160.1.1510 | Bacteria;Firmicutes;Negativicutes;Acidaminococcales;Acidaminococcaceae;Succinispira;uncultured bacterium | 89.8 | 1439 | -1 |
| metaRNA.PFemirge\_20523 | 116 | 0.000009 | HQ764606.1.1446 | Bacteria;Bacteroidota;Bacteroidia;Bacteroidales;Bacteroidaceae;Bacteroides;uncultured organism | 90.6 | 1304 | -1 |
| metaRNA.PFemirge\_2761 | 116 | 0.000011 | KC748467.1.1433 | Bacteria;Firmicutes;Bacilli;Lactobacillales;Streptococcaceae;Streptococcus;Streptococcus iniae | 96.2 | 1220 | -1 |
| metaRNA.PFemirge\_26334 | 116 | 0.000019 | EU778872.1.1287 | Bacteria;Firmicutes;Clostridia;Lachnospirales;Lachnospiraceae;Lachnospiraceae UCG-004;uncultured bacterium | 93.0 | 1285 | -1 |
| metaRNA.PFemirge\_21127 | 115 | 0.000013 | CP024676.4151197.4152748 | Bacteria;Proteobacteria;Gammaproteobacteria;Enterobacterales;Enterobacteriaceae;Citrobacter;Citrobacter pasteurii | 95.1 | 1094 | -1 |
| metaRNA.PFemirge\_23766 | 115 | 0.000010 | EF153297.1.1392 | Bacteria;Proteobacteria;Gammaproteobacteria;Enterobacterales;Yersiniaceae;Serratia;uncultured bacterium | 94.9 | 1301 | -1 |
| metaRNA.PFemirge\_23065 | 115 | 0.000024 | CP012801.6655317.6656834 | Bacteria;Bacteroidota;Bacteroidia;Bacteroidales;Bacteroidaceae;Bacteroides;Bacteroides cellulosilyticus | 95.5 | 1162 | -1 |
| metaRNA.PFemirge\_24461 | 114 | 0.000011 | HQ761492.1.1450 | Bacteria;Firmicutes;Clostridia;Lachnospirales;Lachnospiraceae;[Ruminococcus] torques group;uncultured organism | 89.8 | 1293 | -1 |
| metaRNA.PFemirge\_13120 | 114 | 0.000006 | GQ175524.1.1451 | Bacteria;Bacteroidota;Bacteroidia;Bacteroidales;Rikenellaceae;Rikenella;uncultured bacterium | 89.6 | 1256 | -1 |
| metaRNA.PFemirge\_27885 | 113 | 0.000004 | AQHW01000007.13.1527 | Bacteria;Bacteroidota;Bacteroidia;Bacteroidales;Tannerellaceae;Parabacteroides;Parabacteroides gordonii MS-1 | 91.7 | 1490 | -1 |
| metaRNA.PFemirge\_21199 | 113 | 0.000006 | AJ421915.1.1402 | Bacteria;Proteobacteria;Gammaproteobacteria;Burkholderiales;Comamonadaceae;Sphaerotilus;uncultured beta proteobacterium | 97.6 | 1318 | -1 |
| metaRNA.PFemirge\_25403 | 113 | 0.000012 | EU775698.1.1374 | Bacteria;Firmicutes;Clostridia;Clostridiales;Clostridiaceae;Clostridium sensu stricto 1;uncultured bacterium | 92.7 | 1268 | -1 |
| metaRNA.PFemirge\_27844 | 113 | 0.000018 | CEBY01043209.9650.11156 | Bacteria;Bacteroidota;Bacteroidia;Bacteroidales;Rikenellaceae;Alistipes;gut metagenome | 91.4 | 1483 | -1 |
| metaRNA.PFemirge\_15841 | 112 | 0.000011 | AB307664.1.1358 | Bacteria;Proteobacteria;Alphaproteobacteria;Rhodospirillales;Magnetospirillaceae;uncultured;uncultured bacterium | 89.2 | 1282 | -1 |
| metaRNA.PFemirge\_23699 | 112 | 0.000008 | AB088925.1.1354 | Bacteria;Bacteroidota;Bacteroidia;Bacteroidales;Williamwhitmaniaceae;Rs-D38 termite group;uncultured Bacteroidaceae bacterium | 88.0 | 1289 | -1 |
| metaRNA.PFemirge\_12098 | 112 | 0.000009 | KJ725364.1.1442 | Bacteria;Proteobacteria;Gammaproteobacteria;Burkholderiales;Chromobacteriaceae;Vogesella;Vibrio cholerae | 94.1 | 1365 | -1 |
| metaRNA.PFemirge\_26078 | 112 | 0.000014 | EU459250.1.1293 | Bacteria;Firmicutes;Clostridia;Peptostreptococcales-Tissierellales;Peptostreptococcaceae;Paeniclostridium;uncultured bacterium | 96.1 | 1282 | -1 |
| metaRNA.PFemirge\_11184 | 112 | 0.000013 | CP003153.1901384.1902805 | Bacteria;Proteobacteria;Gammaproteobacteria;Burkholderiales;Rhodocyclaceae;Azospira;Azospira oryzae PS | 92.3 | 1319 | -1 |
| metaRNA.PFemirge\_24632 | 111 | 0.000006 | JF490034.1.1450 | Bacteria;Proteobacteria;Alphaproteobacteria;Rhodospirillales;Magnetospirillaceae;uncultured;alpha proteobacterium UT-4 | 90.5 | 1196 | -1 |
| metaRNA.PFemirge\_16556 | 111 | 0.000006 | EF096958.1.1403 | Bacteria;Bacteroidota;Bacteroidia;Bacteroidales;Bacteroidaceae;Bacteroides;uncultured bacterium | 86.2 | 1401 | -1 |
| metaRNA.PFemirge\_9673 | 111 | 0.000013 | FPLS01027059.10.1473 | Bacteria;Proteobacteria;Alphaproteobacteria;Rhizobiales;Devosiaceae;Devosia;metagenome | 91.7 | 1450 | -1 |
| metaRNA.PFemirge\_27764 | 111 | 0.000008 | LT725663.1.1450 | Bacteria;Bacteroidota;Bacteroidia;Bacteroidales;Tannerellaceae;Parabacteroides;Parabacteroides sp. Marseille-P3668 | 93.2 | 1379 | -1 |
| metaRNA.PFemirge\_30242 | 109 | 0.000027 | JQ940836.1.1449 | Bacteria;Bacteroidota;Bacteroidia;Bacteroidales;Rikenellaceae;Alistipes;uncultured bacterium | 83.2 | 1243 | -1 |
| metaRNA.PFemirge\_23451 | 109 | 0.000011 | EU475681.1.1364 | Bacteria;Firmicutes;Clostridia;Peptostreptococcales-Tissierellales;Peptostreptococcaceae;uncultured;uncultured bacterium | 89.8 | 1378 | -1 |
| metaRNA.PFemirge\_26243 | 109 | 0.000042 | EU778905.1.1289 | Bacteria;Firmicutes;Clostridia;Oscillospirales;Ruminococcaceae;Faecalibacterium;uncultured bacterium | 89.0 | 1227 | -1 |
| metaRNA.PFemirge\_6989 | 108 | 0.000076 | HG934468.2032323.2033833 | Bacteria;Bacteroidota;Bacteroidia;Bacteroidales;Rikenellaceae;Mucinivorans;Mucinivorans hirudinis | 90.0 | 1481 | -1 |
| metaRNA.PFemirge\_10180 | 108 | 0.000008 | CP000875.5447186.5448660 | Bacteria;Chloroflexi;Chloroflexia;Chloroflexales;Herpetosiphonaceae;Herpetosiphon;Herpetosiphon aurantiacus DSM 785 | 99.4 | 1137 | -1 |
| metaRNA.PFemirge\_17433 | 108 | 0.000009 | HQ772787.1.1440 | Bacteria;Firmicutes;Clostridia;Lachnospirales;Lachnospiraceae;Agathobacter;uncultured organism | 91.7 | 1380 | -1 |
| metaRNA.PFemirge\_15906 | 108 | 0.000034 | HQ767134.1.1448 | Bacteria;Bacteroidota;Bacteroidia;Bacteroidales;Tannerellaceae;Parabacteroides;uncultured organism | 93.1 | 1374 | -1 |
| metaRNA.PFemirge\_7266 | 107 | 0.000005 | EU774419.1.1295 | Bacteria;Firmicutes;Clostridia;Christensenellales;Christensenellaceae;Christensenellaceae R-7 group;uncultured bacterium | 90.8 | 1293 | -1 |
| metaRNA.PFemirge\_23113 | 107 | 0.000004 | HQ806032.1.1447 | Bacteria;Bacteroidota;Bacteroidia;Bacteroidales;Tannerellaceae;Parabacteroides;uncultured organism | 92.0 | 1230 | -1 |
| metaRNA.PFemirge\_3258 | 107 | 0.000008 | CP012938.2956562.2958087 | Bacteria;Bacteroidota;Bacteroidia;Bacteroidales;Bacteroidaceae;Bacteroides;Bacteroides ovatus | 87.9 | 1320 | -1 |
| metaRNA.PFemirge\_10948 | 107 | 0.000015 | AGZO01000045.56609.58123 | Bacteria;Bacteroidota;Bacteroidia;Bacteroidales;Tannerellaceae;Parabacteroides;Parabacteroides goldsteinii CL02T12C30 | 93.7 | 1383 | -1 |
| metaRNA.PFemirge\_6605 | 106 | 0.000009 | JF160900.1.1352 | Bacteria;Firmicutes;Clostridia;Oscillospirales;Oscillospiraceae;uncultured;uncultured bacterium | 93.3 | 1269 | -1 |
| metaRNA.PFemirge\_21460 | 106 | 0.000044 | JX000037.1.1482 | Bacteria;Firmicutes;Clostridia;Oscillospirales;Oscillospiraceae;uncultured;uncultured bacterium | 91.8 | 1150 | -1 |
| metaRNA.PFemirge\_2385 | 106 | 0.000143 | JN038827.1.1489 | Bacteria;Proteobacteria;Gammaproteobacteria;Burkholderiales;Sutterellaceae;uncultured;uncultured beta proteobacterium | 93.0 | 1414 | -1 |
| metaRNA.PFemirge\_17717 | 106 | 0.000011 | DQ824247.1.1290 | Bacteria;Firmicutes;Clostridia;Lachnospirales;Lachnospiraceae;Lachnoclostridium;uncultured bacterium | 92.3 | 1292 | -1 |
| metaRNA.PFemirge\_11298 | 105 | 0.000012 | FQXY01000005.366.1875 | Bacteria;Bacteroidota;Bacteroidia;Bacteroidales;Bacteroidaceae;Bacteroides;Bacteroides sp. Marseille-P3132 | 89.1 | 1391 | -1 |
| metaRNA.PFemirge\_13651 | 105 | 0.000003 | LT725663.1.1450 | Bacteria;Bacteroidota;Bacteroidia;Bacteroidales;Tannerellaceae;Parabacteroides;Parabacteroides sp. Marseille-P3668 | 90.1 | 1339 | -1 |
| metaRNA.PFemirge\_4967 | 105 | 0.000011 | JN217178.1.1510 | Bacteria;Proteobacteria;Gammaproteobacteria;Acidithiobacillales;Acidithiobacillaceae;KCM-B-112;uncultured bacterium | 95.3 | 1398 | -1 |
| metaRNA.PFemirge\_11327 | 104 | 0.000006 | KM650913.1.1442 | Bacteria;Bacteroidota;Bacteroidia;Bacteroidales;Bacteroidaceae;Bacteroides;uncultured Bacteroidetes bacterium | 89.8 | 1386 | -1 |
| metaRNA.PFemirge\_23625 | 104 | 0.000016 | CU926592.1.1355 | Bacteria;Bacteroidota;Bacteroidia;Bacteroidales;Porphyromonadaceae;Porphyromonas;uncultured bacterium | 92.4 | 1356 | -1 |
| metaRNA.PFemirge\_13244 | 104 | 0.000017 | HQ761452.1.1440 | Bacteria;Bacteroidota;Bacteroidia;Bacteroidales;Tannerellaceae;Parabacteroides;uncultured organism | 93.4 | 1391 | -1 |
| metaRNA.PFemirge\_18308 | 104 | 0.000006 | BAAY01003437.486.1979 | Bacteria;Firmicutes;Clostridia;Lachnospirales;Lachnospiraceae;uncultured;human gut metagenome | 93.6 | 1300 | -1 |
| metaRNA.PFemirge\_22898 | 104 | 0.000015 | HQ785589.1.1441 | Bacteria;Bacteroidota;Bacteroidia;Bacteroidales;Bacteroidaceae;Bacteroides;uncultured organism | 91.3 | 1292 | -1 |
| metaRNA.PFemirge\_30245 | 102 | 0.000213 | JX105679.1.1392 | Bacteria;Bacteroidota;Bacteroidia;Bacteroidales;Bacteroidaceae;Bacteroides;uncultured bacterium | 86.4 | 1371 | -1 |
| metaRNA.PFemirge\_23138 | 102 | 0.000015 | EU465589.1.1360 | Bacteria;Firmicutes;Clostridia;Lachnospirales;Lachnospiraceae;Lachnospiraceae NK4A136 group;uncultured bacterium | 92.6 | 1305 | -1 |
| metaRNA.PFemirge\_5138 | 102 | 0.000654 | FN794258.1.1496 | Bacteria;Proteobacteria;Gammaproteobacteria;Burkholderiales;Rhodocyclaceae;Azospira;uncultured bacterium | 96.6 | 1403 | -1 |
| metaRNA.PFemirge\_26187 | 102 | 0.000008 | HQ748426.1.1449 | Bacteria;Firmicutes;Clostridia;Lachnospirales;Lachnospiraceae;Lachnoclostridium;uncultured organism | 92.5 | 1290 | -1 |
| metaRNA.PFemirge\_18465 | 102 | 0.000004 | FR749946.1.1524 | Bacteria;Firmicutes;Clostridia;Lachnospirales;Lachnospiraceae;[Eubacterium] fissicatena group;Faecalicatena contorta | 92.0 | 1320 | -1 |
| metaRNA.PFemirge\_19936 | 102 | 0.000026 | EU771244.1.1399 | Bacteria;Firmicutes;Clostridia;Lachnospirales;Lachnospiraceae;Lachnospiraceae NC2004 group;uncultured bacterium | 91.4 | 1238 | -1 |
| metaRNA.PFemirge\_27610 | 101 | 0.000006 | EU475758.1.1213 | Bacteria;Firmicutes;Clostridia;Lachnospirales;Lachnospiraceae;Lachnospiraceae UCG-008;uncultured bacterium | 95.6 | 1098 | -1 |
| metaRNA.PFemirge\_8971 | 101 | 0.000011 | AB198564.1.1344 | Bacteria;Elusimicrobiota;Elusimicrobia;Elusimicrobiales;Elusimicrobiaceae;Elusimicrobium;uncultured Termite group 1 bacterium | 96.7 | 1159 | -1 |
| metaRNA.PFemirge\_27531 | 101 | 0.000013 | AGES01000014.73256.74770 | Bacteria;Bacteroidota;Bacteroidia;Bacteroidales;Tannerellaceae;Parabacteroides;Parabacteroides sp. HGS0025 | 94.8 | 1193 | -1 |
| metaRNA.PFemirge\_18125 | 100 | 0.000005 | DQ793316.1.1393 | Bacteria;Bacteroidota;Bacteroidia;Bacteroidales;Tannerellaceae;Parabacteroides;uncultured bacterium | 92.2 | 1219 | -1 |
| metaRNA.PFemirge\_11165 | 100 | 0.000012 | EU616637.1.1399 | Bacteria;Bacteroidota;Bacteroidia;Bacteroidales;Rikenellaceae;Rikenella;uncultured Rikenellaceae bacterium | 89.5 | 1223 | -1 |
| metaRNA.PFemirge\_350 | 100 | 0.000007 | AF083009.1.1734 | Eukaryota;Amorphea;Obazoa;Opisthokonta;Holozoa;Choanozoa;Metazoa;Animalia;BCP clade;Bilateria;Protostomia;Ecdysozoa;Nematozoa;Nematoda;Chromadorea;Rhabditida;Diploscapter sp. PS1897 | 100.0 | 1399 | -1 |
| metaRNA.PFemirge\_4408 | 100 | 0.000004 | ATXW01000003.334809.336317 | Bacteria;Bacteroidota;Bacteroidia;Bacteroidales;Rikenellaceae;Rikenella;Rikenella microfusus DSM 15922 | 87.9 | 1492 | -1 |
| metaRNA.PFemirge\_23481 | 100 | 0.000008 | KJ584639.1.1330 | Bacteria;Proteobacteria;Alphaproteobacteria;Rhizobiales;Beijerinckiaceae;Bosea;Bosea sp. A17 | 93.7 | 1260 | -1 |
| metaRNA.PFemirge\_25956 | 100 | 0.000062 | EU471853.1.1298 | Bacteria;Firmicutes;Clostridia;Christensenellales;Christensenellaceae;Christensenellaceae R-7 group;uncultured bacterium | 93.0 | 1286 | -1 |
| metaRNA.PFemirge\_26295 | 99 | 0.000002 | FJ534957.1.1501 | Bacteria;Firmicutes;Clostridia;Lachnospirales;Lachnospiraceae;Lachnospiraceae FCS020 group;uncultured bacterium | 87.1 | 1263 | -1 |
| metaRNA.PFemirge\_12327 | 99 | 0.000005 | HQ772765.1.1442 | Bacteria;Bacteroidota;Bacteroidia;Bacteroidales;Bacteroidaceae;Bacteroides;uncultured organism | 90.3 | 1389 | -1 |
| metaRNA.PFemirge\_6401 | 98 | 0.000005 | HQ819238.1.1451 | Bacteria;Bacteroidota;Bacteroidia;Bacteroidales;Bacteroidaceae;Bacteroides;uncultured organism | 91.4 | 1298 | -1 |
| metaRNA.PFemirge\_16525 | 98 | 0.000007 | AY452004.1.1502 | Bacteria;Firmicutes;Clostridia;Lachnospirales;Lachnospiraceae;Lachnospiraceae UCG-004;uncultured bacterium | 90.2 | 1241 | -1 |
| metaRNA.PFemirge\_18350 | 98 | 0.000004 | DQ676995.1.1498 | Bacteria;Actinobacteriota;Actinobacteria;Propionibacteriales;Propionibacteriaceae;Propionicicella;iron-reducing enrichment clone Cl-A3 | 92.2 | 1276 | -1 |
| metaRNA.PFemirge\_8230 | 98 | 0.000014 | GU304321.1.1480 | Bacteria;Firmicutes;Clostridia;Lachnospirales;Lachnospiraceae;Lachnospiraceae UCG-008;uncultured rumen bacterium | 90.7 | 1469 | -1 |
| metaRNA.PFemirge\_20745 | 97 | 0.000017 | DQ797886.1.1381 | Bacteria;Firmicutes;Clostridia;Lachnospirales;Lachnospiraceae;Lachnospiraceae UCG-004;uncultured bacterium | 91.8 | 1379 | -1 |
| metaRNA.PFemirge\_16730 | 97 | 0.000011 | EU776960.1.1399 | Bacteria;Proteobacteria;Gammaproteobacteria;Enterobacterales;Enterobacteriaceae;Escherichia-Shigella;uncultured bacterium | 96.3 | 1397 | -1 |
| metaRNA.PFemirge\_12807 | 97 | 0.000003 | HQ806032.1.1447 | Bacteria;Bacteroidota;Bacteroidia;Bacteroidales;Tannerellaceae;Parabacteroides;uncultured organism | 90.0 | 1387 | -1 |
| metaRNA.PFemirge\_9592 | 97 | 0.000007 | AB509219.1.1467 | Bacteria;Firmicutes;Negativicutes;Acidaminococcales;Acidaminococcaceae;Phascolarctobacterium;uncultured bacterium | 93.2 | 1359 | -1 |
| metaRNA.PFemirge\_13867 | 97 | 0.000006 | HQ761492.1.1450 | Bacteria;Firmicutes;Clostridia;Lachnospirales;Lachnospiraceae;[Ruminococcus] torques group;uncultured organism | 89.8 | 1377 | -1 |
| metaRNA.PFemirge\_11728 | 97 | 0.000004 | HQ810916.1.1454 | Bacteria;Firmicutes;Clostridia;Oscillospirales;Butyricicoccaceae;UCG-009;uncultured organism | 90.7 | 1370 | -1 |
| metaRNA.PFemirge\_13802 | 97 | 0.000006 | DQ659895.1.1400 | Bacteria;Actinobacteriota;Actinobacteria;Corynebacteriales;Nocardiaceae;Nocardia;Nocardia abscessus | 96.5 | 1343 | -1 |
| metaRNA.PFemirge\_23597 | 96 | 0.000021 | LT725663.1.1450 | Bacteria;Bacteroidota;Bacteroidia;Bacteroidales;Tannerellaceae;Parabacteroides;Parabacteroides sp. Marseille-P3668 | 90.3 | 1357 | -1 |
| metaRNA.PFemirge\_7092 | 96 | 0.000017 | AGES01000014.73256.74770 | Bacteria;Bacteroidota;Bacteroidia;Bacteroidales;Tannerellaceae;Parabacteroides;Parabacteroides sp. HGS0025 | 87.5 | 1334 | -1 |
| metaRNA.PFemirge\_12927 | 96 | 0.000008 | AB574479.1.1484 | Bacteria;Bacteroidota;Bacteroidia;Bacteroidales;Bacteroidaceae;Bacteroides;Bacteroides stercorirosoris | 88.6 | 1391 | -1 |
| metaRNA.PFemirge\_25313 | 96 | 0.000007 | KP876564.1.1454 | Bacteria;Proteobacteria;Alphaproteobacteria;Reyranellales;Reyranellaceae;Reyranella;uncultured bacterium | 95.8 | 980 | -1 |
| metaRNA.PFemirge\_17976 | 96 | 0.000032 | FJ534957.1.1501 | Bacteria;Firmicutes;Clostridia;Lachnospirales;Lachnospiraceae;Lachnospiraceae FCS020 group;uncultured bacterium | 91.6 | 1304 | -1 |
| metaRNA.PFemirge\_27575 | 95 | 0.000002 | JF160900.1.1352 | Bacteria;Firmicutes;Clostridia;Oscillospirales;Oscillospiraceae;uncultured;uncultured bacterium | 95.6 | 776 | -1 |
| metaRNA.PFemirge\_5190 | 95 | 0.000008 | JQ923952.1.1496 | Bacteria;Proteobacteria;Gammaproteobacteria;Burkholderiales;Rhodocyclaceae;Candidatus Accumulibacter;uncultured bacterium | 95.2 | 1403 | -1 |
| metaRNA.PFemirge\_11750 | 95 | 0.000002 | AGZO01000045.56609.58123 | Bacteria;Bacteroidota;Bacteroidia;Bacteroidales;Tannerellaceae;Parabacteroides;Parabacteroides goldsteinii CL02T12C30 | 88.8 | 1452 | -1 |
| metaRNA.PFemirge\_9100 | 95 | 0.000003 | KC432217.1.1330 | Bacteria;Firmicutes;Clostridia;Clostridiales;Clostridiaceae;Clostridium sensu stricto 3;uncultured bacterium | 95.0 | 1332 | -1 |
| metaRNA.PFemirge\_6145 | 95 | 0.000003 | AB547646.1.1488 | Bacteria;Bacteroidota;Bacteroidia;Bacteroidales;Bacteroidaceae;Bacteroides;Bacteroides rodentium | 93.5 | 1394 | -1 |
| metaRNA.PFemirge\_4069 | 94 | 0.000010 | JN217164.1.1510 | Bacteria;Proteobacteria;Gammaproteobacteria;Acidithiobacillales;Acidithiobacillaceae;KCM-B-112;uncultured bacterium | 92.1 | 1411 | -1 |
| metaRNA.PFemirge\_18858 | 94 | 0.000018 | AB649276.1.1495 | Bacteria;Firmicutes;Clostridia;Lachnospirales;Lachnospiraceae;Anaerostignum;Anaerotignum propionicum | 91.8 | 1390 | -1 |
| metaRNA.PFemirge\_19453 | 94 | 0.000002 | JX872321.1.1469 | Bacteria;Proteobacteria;Gammaproteobacteria;Enterobacterales;Aeromonadaceae;Aeromonas;uncultured bacterium | 91.4 | 1273 | -1 |
| metaRNA.PFemirge\_27849 | 94 | 0.000032 | HQ806032.1.1447 | Bacteria;Bacteroidota;Bacteroidia;Bacteroidales;Tannerellaceae;Parabacteroides;uncultured organism | 92.6 | 1454 | -1 |
| metaRNA.PFemirge\_10927 | 93 | 0.000021 | KC669122.1.1486 | Bacteria;Firmicutes;Clostridia;Lachnospirales;Lachnospiraceae;Epulopiscium;uncultured bacterium | 92.8 | 1246 | -1 |
| metaRNA.PFemirge\_1733 | 93 | 0.000003 | GQ480160.1.1510 | Bacteria;Firmicutes;Negativicutes;Acidaminococcales;Acidaminococcaceae;Succinispira;uncultured bacterium | 90.3 | 1492 | -1 |
| metaRNA.PFemirge\_23125 | 93 | 0.000014 | AB649276.1.1495 | Bacteria;Firmicutes;Clostridia;Lachnospirales;Lachnospiraceae;Anaerostignum;Anaerotignum propionicum | 87.7 | 1397 | -1 |
| metaRNA.PFemirge\_19102 | 93 | 0.000009 | FJ534957.1.1501 | Bacteria;Firmicutes;Clostridia;Lachnospirales;Lachnospiraceae;Lachnospiraceae FCS020 group;uncultured bacterium | 91.6 | 1267 | -1 |
| metaRNA.PFemirge\_28012 | 93 | 0.000012 | JQ317253.1.1380 | Bacteria;Bacteroidota;Bacteroidia;Bacteroidales;Bacteroidaceae;Bacteroides;Bacteroides sp. CannelCatfish9 | 93.4 | 1159 | -1 |
| metaRNA.PFemirge\_26246 | 93 | 0.000009 | FJ542833.1.1492 | Bacteria;Verrucomicrobiota;Verrucomicrobiae;Chthoniobacterales;Terrimicrobiaceae;Terrimicrobium;uncultured Verrucomicrobiales bacterium | 96.9 | 1095 | -1 |
| metaRNA.PFemirge\_1297 | 92 | 0.000008 | FPLP01012861.16.1542 | Bacteria;Verrucomicrobiota;Verrucomicrobiae;Verrucomicrobiales;Verrucomicrobiaceae;uncultured;metagenome | 94.1 | 1313 | -1 |
| metaRNA.PFemirge\_27871 | 92 | 0.000006 | AB264620.1.1463 | Bacteria;Firmicutes;Clostridia;Lachnospirales;Lachnospiraceae;Lachnospiraceae NC2004 group;Clostridiaceae bacterium bSSV31 | 93.9 | 1302 | -1 |
| metaRNA.PFemirge\_27128 | 92 | 0.000013 | EU774505.1.1241 | Bacteria;Firmicutes;Clostridia;Clostridia UCG-014;uncultured bacterium | 91.5 | 1230 | -1 |
| metaRNA.PFemirge\_23838 | 92 | 0.000015 | EU778697.1.1353 | Bacteria;Bacteroidota;Bacteroidia;Bacteroidales;Rikenellaceae;SP3-e08;uncultured bacterium | 88.2 | 1300 | -1 |
| metaRNA.PFemirge\_16271 | 92 | 0.000008 | HM630236.1.1451 | Bacteria;Bacteroidota;Bacteroidia;Bacteroidales;Rikenellaceae;dgA-11 gut group;uncultured bacterium | 89.4 | 1405 | -1 |
| metaRNA.PFemirge\_20522 | 92 | 0.000007 | BABG01001469.752.2264 | Bacteria;Firmicutes;Clostridia;Lachnospirales;Lachnospiraceae;Lachnoclostridium;human gut metagenome | 91.0 | 1136 | -1 |
| metaRNA.PFemirge\_23080 | 91 | 0.000007 | DQ816804.1.1464 | Bacteria;Firmicutes;Clostridia;Peptostreptococcales-Tissierellales;Anaerovoracaceae;Anaerovorax;uncultured bacterium | 93.0 | 1312 | -1 |
| metaRNA.PFemirge\_8730 | 91 | 0.000006 | AM930363.1.1475 | Bacteria;Firmicutes;Clostridia;Clostridiales;Clostridiaceae;Clostridium sensu stricto 1;uncultured bacterium | 95.7 | 1411 | -1 |
| metaRNA.PFemirge\_5928 | 91 | 0.000011 | KT952743.1.1492 | Bacteria;Bacteroidota;Bacteroidia;Bacteroidales;Rikenellaceae;Rikenella;uncultured bacterium | 89.3 | 1469 | -1 |
| metaRNA.PFemirge\_3004 | 91 | 0.000002 | AF371540.1.1455 | Bacteria;Firmicutes;Clostridia;Lachnospirales;Lachnospiraceae;Blautia;uncultured bacterium | 88.9 | 1230 | -1 |
| metaRNA.PFemirge\_21766 | 90 | 0.000003 | JX105679.1.1392 | Bacteria;Bacteroidota;Bacteroidia;Bacteroidales;Bacteroidaceae;Bacteroides;uncultured bacterium | 90.3 | 1347 | -1 |
| metaRNA.PFemirge\_25115 | 90 | 0.000008 | LN563559.1.1331 | Bacteria;Bacteroidota;Bacteroidia;Cytophagales;Amoebophilaceae;Candidatus Amoebophilus;uncultured bacterium | 97.1 | 1281 | -1 |
| metaRNA.PFemirge\_15469 | 90 | 0.000008 | EU777648.1.1410 | Bacteria;Firmicutes;Bacilli;Lactobacillales;Streptococcaceae;Lactococcus;uncultured bacterium | 95.4 | 1173 | -1 |
| metaRNA.PFemirge\_7984 | 90 | 0.000019 | KC994683.1.1482 | Bacteria;Bacteroidota;Bacteroidia;Cytophagales;Spirosomaceae;Runella;uncultured Runella sp. | 93.0 | 1380 | -1 |
| metaRNA.PFemirge\_16960 | 88 | 0.000007 | MF175255.1.1500 | Bacteria;Firmicutes;Clostridia;Lachnospirales;Lachnospiraceae;Lachnoclostridium;Clostridium sp. | 95.9 | 1388 | -1 |
| metaRNA.PFemirge\_4079 | 87 | 0.000011 | AY957925.1.1536 | Bacteria;Proteobacteria;Gammaproteobacteria;Legionellales;Legionellaceae;Legionella;uncultured bacterium | 95.4 | 1400 | -1 |
| metaRNA.PFemirge\_27396 | 87 | 0.000005 | KF464766.1.1221 | Bacteria;Proteobacteria;Gammaproteobacteria;Enterobacterales;Yersiniaceae;Yersinia;uncultured bacterium | 95.9 | 1206 | -1 |
| metaRNA.PFemirge\_10256 | 87 | 0.000017 | LT725663.1.1450 | Bacteria;Bacteroidota;Bacteroidia;Bacteroidales;Tannerellaceae;Parabacteroides;Parabacteroides sp. Marseille-P3668 | 92.4 | 1383 | -1 |
| metaRNA.PFemirge\_27211 | 87 | 0.000011 | FPLS01020683.16.1482 | Bacteria;Proteobacteria;Alphaproteobacteria;Sphingomonadales;Sphingomonadaceae;Novosphingobium;metagenome | 89.7 | 1230 | -1 |
| metaRNA.PFemirge\_28083 | 87 | 0.000016 | GQ898302.1.1489 | Bacteria;Bacteroidota;Bacteroidia;Bacteroidales;Bacteroidaceae;Bacteroides;uncultured bacterium | 94.6 | 965 | -1 |
| metaRNA.PFemirge\_26515 | 86 | 0.000007 | EU776450.1.1387 | Bacteria;Firmicutes;Clostridia;Oscillospirales;Ruminococcaceae;Ruminococcus;uncultured bacterium | 91.1 | 1282 | -1 |
| metaRNA.PFemirge\_21595 | 86 | 0.000006 | M59091.1.1455 | Bacteria;Firmicutes;Clostridia;Clostridiales;Clostridiaceae;Clostridium sensu stricto 1;Clostridium carnis | 93.8 | 1367 | -1 |
| metaRNA.PFemirge\_5586 | 86 | 0.000016 | CBVB010000006.254456.255954 | Bacteria;Bacteroidota;Bacteroidia;Bacteroidales;Bacteroidaceae;Bacteroides;Bacteroidaceae bacterium MS4 | 92.7 | 1309 | -1 |
| metaRNA.PFemirge\_25050 | 86 | 0.000004 | CP003274.119147.120561 | Bacteria;Bacteroidota;Bacteroidia;Bacteroidales;Rikenellaceae;Alistipes;Alistipes finegoldii DSM 17242 | 84.4 | 1294 | -1 |
| metaRNA.PFemirge\_19870 | 85 | 0.000021 | DQ810054.1.1385 | Bacteria;Firmicutes;Clostridia;Lachnospirales;Lachnospiraceae;Lachnoclostridium;uncultured bacterium | 93.6 | 1383 | -1 |
| metaRNA.PFemirge\_14514 | 85 | 0.000032 | KT952840.1.1494 | Bacteria;Firmicutes;Clostridia;Oscillospirales;Oscillospiraceae;Flavonifractor;uncultured bacterium | 91.8 | 1347 | -1 |
| metaRNA.PFemirge\_9231 | 84 | 0.000012 | GQ376177.1.1441 | Bacteria;Actinobacteriota;Actinobacteria;Corynebacteriales;Nocardiaceae;Nocardia;Nocardia lijiangensis | 95.2 | 1387 | -1 |
| metaRNA.PFemirge\_24547 | 84 | 0.000006 | EU468659.1.1353 | Bacteria;Firmicutes;Clostridia;Lachnospirales;Lachnospiraceae;uncultured;uncultured bacterium | 92.5 | 1259 | -1 |
| metaRNA.PFemirge\_25014 | 84 | 0.000142 | JX198651.1.1513 | Bacteria;Firmicutes;Clostridia;Oscillospirales;Butyricicoccaceae;UCG-009;uncultured bacterium | 91.6 | 1268 | -1 |
| metaRNA.PFemirge\_12011 | 84 | 0.000016 | FQXY01000005.366.1875 | Bacteria;Bacteroidota;Bacteroidia;Bacteroidales;Bacteroidaceae;Bacteroides;Bacteroides sp. Marseille-P3132 | 91.4 | 1390 | -1 |
| metaRNA.PFemirge\_12596 | 83 | 0.000015 | FN667250.1.1445 | Bacteria;Firmicutes;Clostridia;Oscillospirales;Ruminococcaceae;Anaerotruncus;uncultured compost bacterium | 94.1 | 1321 | -1 |
| metaRNA.PFemirge\_30255 | 83 | 0.000046 | DQ800679.1.1366 | Bacteria;Firmicutes;Clostridia;Lachnospirales;Lachnospiraceae;[Ruminococcus] gauvreauii group;uncultured bacterium | 91.1 | 1363 | -1 |
| metaRNA.PFemirge\_25565 | 82 | 0.000008 | JF266314.1.1315 | Bacteria;Cyanobacteria;Vampirivibrionia;Obscuribacterales;Obscuribacteraceae;uncultured bacterium | 95.2 | 1249 | -1 |
| metaRNA.PFemirge\_5805 | 82 | 0.000005 | DQ793316.1.1393 | Bacteria;Bacteroidota;Bacteroidia;Bacteroidales;Tannerellaceae;Parabacteroides;uncultured bacterium | 93.5 | 1216 | -1 |
| metaRNA.PFemirge\_18679 | 82 | 0.000010 | DQ810138.1.1391 | Bacteria;Firmicutes;Clostridia;Lachnospirales;Lachnospiraceae;Agathobacter;uncultured bacterium | 92.2 | 1385 | -1 |
| metaRNA.PFemirge\_22056 | 82 | 0.000025 | EF400740.1.1495 | Bacteria;Firmicutes;Clostridia;Oscillospirales;Butyricicoccaceae;UCG-009;uncultured bacterium | 93.5 | 1328 | -1 |
| metaRNA.PFemirge\_6260 | 81 | 0.000009 | FJ534957.1.1501 | Bacteria;Firmicutes;Clostridia;Lachnospirales;Lachnospiraceae;Lachnospiraceae FCS020 group;uncultured bacterium | 93.0 | 1379 | -1 |
| metaRNA.PFemirge\_4901 | 81 | 0.000008 | FJ823944.1.1497 | Bacteria;Proteobacteria;Gammaproteobacteria;Burkholderiales;Comamonadaceae;uncultured;uncultured Comamonas sp. | 93.6 | 1393 | -1 |
| metaRNA.PFemirge\_27917 | 81 | 0.000004 | BAAZ01000703.1371.2875 | Bacteria;Bacteroidota;Bacteroidia;Bacteroidales;Tannerellaceae;Parabacteroides;human gut metagenome | 91.3 | 1304 | -1 |
| metaRNA.PFemirge\_1572 | 81 | 0.000005 | FPLN01009549.9.1530 | Bacteria;Proteobacteria;Gammaproteobacteria;Enterobacterales;Enterobacteriaceae;Escherichia-Shigella;metagenome | 90.4 | 1385 | -1 |
| metaRNA.PFemirge\_11344 | 81 | 0.000006 | GCKC01172265.99.1551 | Bacteria;Proteobacteria;Alphaproteobacteria;Rickettsiales;Mitochondria;Gleditsia sinensis | 95.0 | 1382 | -1 |
| metaRNA.PFemirge\_24475 | 80 | 0.000031 | LC094617.1.1418 | Bacteria;Actinobacteriota;Actinobacteria;Micrococcales;Microbacteriaceae;Leifsonia;Microbacteriaceae bacterium MAEY21 | 89.9 | 1313 | -1 |
| metaRNA.PFemirge\_15445 | 80 | 0.000011 | DQ799866.1.1410 | Bacteria;Bacteroidota;Bacteroidia;Bacteroidales;Rikenellaceae;Alistipes;uncultured bacterium | 92.8 | 1407 | -1 |
| metaRNA.PFemirge\_3395 | 80 | 0.000008 | KU667224.1.1485 | Bacteria;Bacteroidota;Bacteroidia;Bacteroidales;Rikenellaceae;uncultured;uncultured bacterium | 89.4 | 1487 | -1 |
| metaRNA.PFemirge\_26583 | 80 | 0.000005 | EU771874.1.1278 | Bacteria;Firmicutes;Bacilli;Erysipelotrichales;Erysipelatoclostridiaceae;uncultured;uncultured bacterium | 92.6 | 1277 | -1 |
| metaRNA.PFemirge\_4892 | 80 | 0.000019 | HM630236.1.1451 | Bacteria;Bacteroidota;Bacteroidia;Bacteroidales;Rikenellaceae;dgA-11 gut group;uncultured bacterium | 90.3 | 1418 | -1 |
| metaRNA.PFemirge\_25338 | 80 | 0.000012 | AB934800.1.1322 | Bacteria;Proteobacteria;Alphaproteobacteria;Rhodospirillales;uncultured;uncultured bacterium | 91.6 | 1177 | -1 |
| metaRNA.PFemirge\_4780 | 80 | 0.000004 | LS483441.307060.308592 | Bacteria;Proteobacteria;Gammaproteobacteria;Enterobacterales;Aeromonadaceae;Aeromonas;Aeromonas caviae | 95.8 | 1328 | -1 |
| metaRNA.PFemirge\_20527 | 80 | 0.000002 | KT952726.1.1486 | Bacteria;Firmicutes;Clostridia;Oscillospirales;Ruminococcaceae;uncultured;uncultured bacterium | 90.4 | 1278 | -1 |
| metaRNA.PFemirge\_27532 | 80 | 0.000021 | AB534319.1.1609 | Eukaryota;Amorphea;Obazoa;Opisthokonta;Nucletmycea;Fungi;Dikarya;Ascomycota;Pezizomycotina;Dothideomycetes;Pleosporales;uncultured fungus | 97.5 | 955 | -1 |
| metaRNA.PFemirge\_27966 | 80 | 0.000003 | JX105679.1.1392 | Bacteria;Bacteroidota;Bacteroidia;Bacteroidales;Bacteroidaceae;Bacteroides;uncultured bacterium | 91.6 | 1317 | -1 |
| metaRNA.PFemirge\_22977 | 79 | 0.000003 | CZBH01000001.999036.1000544 | Bacteria;Bacteroidota;Bacteroidia;Bacteroidales;Rikenellaceae;Alistipes;Alistipes sp. cv1 | 92.2 | 1213 | -1 |
| metaRNA.PFemirge\_16332 | 79 | 0.000026 | EU469747.1.1392 | Bacteria;Firmicutes;Clostridia;Lachnospirales;Lachnospiraceae;28-4;uncultured bacterium | 90.9 | 1308 | -1 |
| metaRNA.PFemirge\_7058 | 79 | 0.000007 | EF632748.1.1502 | Bacteria;Firmicutes;Clostridia;Clostridiales;Clostridiaceae;Clostridium sensu stricto 8;uncultured bacterium | 99.0 | 1207 | -1 |
| metaRNA.PFemirge\_19539 | 79 | 0.000004 | EU773965.1.1389 | Bacteria;Firmicutes;Clostridia;Lachnospirales;Lachnospiraceae;[Ruminococcus] torques group;uncultured bacterium | 92.8 | 1296 | -1 |
| metaRNA.PFemirge\_22892 | 78 | 0.000014 | KT002331.1.1465 | Bacteria;Firmicutes;Clostridia;Peptostreptococcales-Tissierellales;Peptostreptococcaceae;Paeniclostridium;uncultured Clostridium sp. | 94.0 | 1361 | -1 |
| metaRNA.PFemirge\_27774 | 78 | 0.000013 | HQ681866.1.1451 | Bacteria;Bacteroidota;Bacteroidia;Bacteroidales;Bacteroidaceae;Bacteroides;uncultured bacterium | 94.3 | 1253 | -1 |
| metaRNA.PFemirge\_8057 | 78 | 0.000005 | AB470343.1.1485 | Bacteria;Bacteroidota;Bacteroidia;Bacteroidales;Tannerellaceae;Parabacteroides;Parabacteroides gordonii | 87.2 | 1329 | -1 |
| metaRNA.PFemirge\_28152 | 78 | 0.000012 | BAAZ01000703.1371.2875 | Bacteria;Bacteroidota;Bacteroidia;Bacteroidales;Tannerellaceae;Parabacteroides;human gut metagenome | 89.8 | 1346 | -1 |
| metaRNA.PFemirge\_25998 | 78 | 0.000014 | GQ358437.1.1400 | Bacteria;Firmicutes;Clostridia;Clostridia UCG-014;uncultured Ruminococcaceae bacterium | 93.6 | 1179 | -1 |
| metaRNA.PFemirge\_2228 | 77 | 0.000010 | MF040523.1.1509 | Bacteria;Verrucomicrobiota;Verrucomicrobiae;Chthoniobacterales;Terrimicrobiaceae;FukuN18 freshwater group;uncultured bacterium | 97.7 | 1164 | -1 |
| metaRNA.PFemirge\_14760 | 77 | 0.000012 | JQ885561.1.1458 | Bacteria;Proteobacteria;Gammaproteobacteria;Enterobacterales;Enterobacteriaceae;Enterobacter;uncultured Enterobacter sp. | 93.4 | 1229 | -1 |
| metaRNA.PFemirge\_16169 | 77 | 0.000005 | ABYH01000014.137.1655 | Bacteria;Bacteroidota;Bacteroidia;Bacteroidales;Tannerellaceae;Parabacteroides;Parabacteroides johnsonii DSM 18315 | 92.4 | 1220 | -1 |
| metaRNA.PFemirge\_26214 | 77 | 0.000007 | DQ797586.1.1394 | Bacteria;Firmicutes;Clostridia;Lachnospirales;Lachnospiraceae;[Eubacterium] eligens group;uncultured bacterium | 89.9 | 1214 | -1 |
| metaRNA.PFemirge\_26055 | 77 | 0.000012 | EU774249.1.1294 | Bacteria;Firmicutes;Clostridia;Oscillospirales;Butyricicoccaceae;uncultured;uncultured bacterium | 92.7 | 1287 | -1 |
| metaRNA.PFemirge\_24172 | 77 | 0.000017 | EU777338.1.1348 | Bacteria;Bacteroidota;Bacteroidia;Bacteroidales;Rikenellaceae;dgA-11 gut group;uncultured bacterium | 88.9 | 1309 | -1 |
| metaRNA.PFemirge\_6959 | 76 | 0.000005 | JN688021.1.1446 | Bacteria;Bacteroidota;Bacteroidia;Bacteroidales;Rikenellaceae;uncultured;bacterium enrichment culture clone M02 | 90.2 | 1448 | -1 |
| metaRNA.PFemirge\_18613 | 76 | 0.000046 | EU778864.1.1377 | Bacteria;Firmicutes;Clostridia;Lachnospirales;Lachnospiraceae;Lachnospiraceae UCG-008;uncultured bacterium | 91.2 | 1245 | -1 |
| metaRNA.PFemirge\_14304 | 76 | 0.000005 | HQ741327.1.1434 | Bacteria;Firmicutes;Clostridia;Clostridiales;Clostridiaceae;Clostridium sensu stricto 1;uncultured bacterium | 94.7 | 1321 | -1 |
| metaRNA.PFemirge\_23719 | 76 | 0.000008 | DQ799040.1.1399 | Bacteria;Bacteroidota;Bacteroidia;Bacteroidales;Bacteroidaceae;Bacteroides;uncultured bacterium | 93.4 | 1209 | -1 |
| metaRNA.PFemirge\_28135 | 76 | 0.000007 | GQ448867.1.1381 | Bacteria;Bacteroidota;Bacteroidia;Bacteroidales;Tannerellaceae;Parabacteroides;uncultured bacterium | 92.1 | 1387 | -1 |
| metaRNA.PFemirge\_25860 | 76 | 0.000002 | FJ365071.1.1328 | Bacteria;Proteobacteria;Alphaproteobacteria;Rhodospirillales;uncultured;uncultured bacterium | 92.7 | 1236 | -1 |
| metaRNA.PFemirge\_19085 | 75 | 0.000007 | AATF01000761.367.1874 | Bacteria;Firmicutes;Clostridia;Oscillospirales;Ruminococcaceae;Ruminococcus;mouse gut metagenome | 92.5 | 1304 | -1 |
| metaRNA.PFemirge\_23557 | 75 | 0.000007 | KF318250.1.1396 | Bacteria;Verrucomicrobiota;Verrucomicrobiae;Verrucomicrobiales;Akkermansiaceae;Akkermansia;uncultured Verrucomicrobia bacterium | 90.0 | 1282 | -1 |
| metaRNA.PFemirge\_22660 | 75 | 0.000007 | HM630236.1.1451 | Bacteria;Bacteroidota;Bacteroidia;Bacteroidales;Rikenellaceae;dgA-11 gut group;uncultured bacterium | 88.5 | 1190 | -1 |
| metaRNA.PFemirge\_24949 | 74 | 0.000006 | MG367100.1.1491 | Bacteria;Firmicutes;Clostridia;Oscillospirales;Oscillospiraceae;Colidextribacter;uncultured Clostridiales Family IV bacterium | 90.6 | 1180 | -1 |
| metaRNA.PFemirge\_19773 | 74 | 0.000009 | EU776080.1.1388 | Bacteria;Firmicutes;Clostridia;Lachnospirales;Lachnospiraceae;Lachnospiraceae NC2004 group;uncultured bacterium | 91.5 | 1299 | -1 |
| metaRNA.PFemirge\_7236 | 74 | 0.000005 | HQ761612.1.1444 | Bacteria;Bacteroidota;Bacteroidia;Bacteroidales;Tannerellaceae;Parabacteroides;uncultured organism | 92.0 | 1447 | -1 |
| metaRNA.PFemirge\_18757 | 74 | 0.000009 | EU850512.1.1455 | Bacteria;Firmicutes;Clostridia;Oscillospirales;Ruminococcaceae;Ruminococcus;uncultured rumen bacterium | 90.7 | 1166 | -1 |
| metaRNA.PFemirge\_22723 | 74 | 0.000013 | DQ810172.1.1365 | Bacteria;Firmicutes;Bacilli;Erysipelotrichales;Erysipelotrichaceae;Dielma;uncultured bacterium | 92.2 | 1198 | -1 |
| metaRNA.PFemirge\_17446 | 74 | 0.000009 | EU465626.1.1378 | Bacteria;Firmicutes;Clostridia;Lachnospirales;Lachnospiraceae;Lachnospiraceae NK4A136 group;uncultured bacterium | 90.8 | 1389 | -1 |
| metaRNA.PFemirge\_22310 | 74 | 0.000028 | AB298756.1.1471 | Bacteria;Firmicutes;Clostridia;Lachnospirales;Lachnospiraceae;Anaerostignum;Anaerotignum aminivorans | 94.3 | 1280 | -1 |
| metaRNA.PFemirge\_21184 | 74 | 0.000012 | HQ748426.1.1449 | Bacteria;Firmicutes;Clostridia;Lachnospirales;Lachnospiraceae;Lachnoclostridium;uncultured organism | 93.7 | 1344 | -1 |
| metaRNA.PFemirge\_21475 | 74 | 0.000087 | EU778290.1.1377 | Bacteria;Firmicutes;Clostridia;Lachnospirales;Lachnospiraceae;Lachnospiraceae NC2004 group;uncultured bacterium | 93.6 | 1288 | -1 |
| metaRNA.PFemirge\_2119 | 74 | 0.000005 | FPLS01033705.10.1524 | Bacteria;Planctomycetota;Planctomycetes;Pirellulales;Pirellulaceae;Pir4 lineage;metagenome | 97.6 | 1359 | -1 |
| metaRNA.PFemirge\_10848 | 73 | 0.000006 | GQ360012.1.1488 | Bacteria;Bacteroidota;Bacteroidia;Bacteroidales;Tannerellaceae;Macellibacteroides;uncultured bacterium | 88.2 | 1374 | -1 |
| metaRNA.PFemirge\_2903 | 73 | 0.000018 | EF400726.1.1488 | Bacteria;Bacteroidota;Bacteroidia;Bacteroidales;Marinifilaceae;Odoribacter;uncultured bacterium | 91.7 | 1489 | -1 |
| metaRNA.PFemirge\_2567 | 73 | 0.000039 | CP012938.2352114.2353639 | Bacteria;Bacteroidota;Bacteroidia;Bacteroidales;Bacteroidaceae;Bacteroides;Bacteroides ovatus | 93.4 | 1420 | -1 |
| metaRNA.PFemirge\_23708 | 73 | 0.000015 | EU616637.1.1399 | Bacteria;Bacteroidota;Bacteroidia;Bacteroidales;Rikenellaceae;Rikenella;uncultured Rikenellaceae bacterium | 89.6 | 1284 | -1 |
| metaRNA.PFemirge\_26253 | 73 | 0.000018 | LOQJ01002372.643.1942 | Eukaryota;Amorphea;Obazoa;Opisthokonta;Nucletmycea;Fungi;Dikarya;Ascomycota;Saccharomycotina;Saccharomycetes;Saccharomycetales;Saccharomycetaceae;Saccharomyces;Saccharomyces pastorianus | 96.9 | 1149 | -1 |
| metaRNA.PFemirge\_19535 | 72 | 0.000007 | EU772882.1.1401 | Bacteria;Firmicutes;Clostridia;Oscillospirales;Butyricicoccaceae;uncultured;uncultured bacterium | 93.4 | 1282 | -1 |
| metaRNA.PFemirge\_5128 | 72 | 0.000006 | CP012938.2352114.2353639 | Bacteria;Bacteroidota;Bacteroidia;Bacteroidales;Bacteroidaceae;Bacteroides;Bacteroides ovatus | 89.9 | 1390 | -1 |
| metaRNA.PFemirge\_16667 | 72 | 0.000017 | EU509830.1.1376 | Bacteria;Firmicutes;Clostridia;Oscillospirales;Ruminococcaceae;Ruminococcus;uncultured bacterium | 91.8 | 1397 | -1 |
| metaRNA.PFemirge\_19427 | 72 | 0.000020 | DQ824136.1.1393 | Bacteria;Bacteroidota;Bacteroidia;Bacteroidales;Bacteroidaceae;Bacteroides;uncultured bacterium | 93.4 | 1383 | -1 |
| metaRNA.PFemirge\_18550 | 71 | 0.000031 | FJ881357.1.1492 | Bacteria;Firmicutes;Clostridia;Oscillospirales;Butyricicoccaceae;UCG-009;uncultured bacterium | 91.4 | 1219 | -1 |
| metaRNA.PFemirge\_20318 | 71 | 0.000011 | DQ129271.1.1384 | Bacteria;Actinobacteriota;Actinobacteria;Propionibacteriales;Nocardioidaceae;Nocardioides;uncultured bacterium | 93.2 | 1214 | -1 |
| metaRNA.PFemirge\_1589 | 71 | 0.000007 | KC431919.1.1365 | Bacteria;Proteobacteria;Gammaproteobacteria;Legionellales;Legionellaceae;Legionella;uncultured bacterium | 91.0 | 1216 | -1 |
| metaRNA.PFemirge\_27744 | 70 | 0.000011 | EU772907.1.1401 | Bacteria;Bacteroidota;Bacteroidia;Bacteroidales;Bacteroidaceae;Bacteroides;uncultured bacterium | 91.6 | 1389 | -1 |
| metaRNA.PFemirge\_7563 | 70 | 0.000014 | AB470343.1.1485 | Bacteria;Bacteroidota;Bacteroidia;Bacteroidales;Tannerellaceae;Parabacteroides;Parabacteroides gordonii | 88.4 | 1322 | -1 |
| metaRNA.PFemirge\_7353 | 70 | 0.000002 | LT223609.1.1490 | Bacteria;Bacteroidota;Bacteroidia;Bacteroidales;Tannerellaceae;Parabacteroides;Parabacteroides faecis | 89.6 | 1352 | -1 |
| metaRNA.PFemirge\_22167 | 70 | 0.000009 | GQ448867.1.1381 | Bacteria;Bacteroidota;Bacteroidia;Bacteroidales;Tannerellaceae;Parabacteroides;uncultured bacterium | 93.9 | 919 | -1 |
| metaRNA.PFemirge\_27568 | 70 | 0.000007 | JN391753.1.1499 | Bacteria;Proteobacteria;Gammaproteobacteria;Burkholderiales;Comamonadaceae;uncultured;uncultured bacterium | 93.2 | 1200 | -1 |
| metaRNA.PFemirge\_21052 | 70 | 0.000003 | DQ796132.1.1379 | Bacteria;Firmicutes;Clostridia;Oscillospirales;Ruminococcaceae;Faecalibacterium;uncultured bacterium | 92.5 | 1284 | -1 |
| metaRNA.PFemirge\_25888 | 70 | 0.000015 | DQ807393.1.1301 | Bacteria;Firmicutes;Clostridia;Lachnospirales;Lachnospiraceae;Anaerostipes;uncultured bacterium | 92.6 | 1299 | -1 |
| metaRNA.PFemirge\_20993 | 70 | 0.000007 | GU302791.1.1495 | Bacteria;Firmicutes;Clostridia;Christensenellales;Christensenellaceae;Christensenellaceae R-7 group;uncultured rumen bacterium | 91.8 | 1292 | -1 |
| metaRNA.PFemirge\_18144 | 70 | 0.000007 | EU469473.1.1385 | Bacteria;Firmicutes;Clostridia;Lachnospirales;Lachnospiraceae;[Ruminococcus] gauvreauii group;uncultured bacterium | 89.3 | 1391 | -1 |
| metaRNA.PFemirge\_25367 | 70 | 0.000009 | CU920243.2.1323 | Bacteria;Planctomycetota;Planctomycetes;Planctomycetales;Rubinisphaeraceae;SH-PL14;uncultured bacterium | 95.7 | 1318 | -1 |
| metaRNA.PFemirge\_9931 | 69 | 0.000007 | EU772041.1.1407 | Bacteria;Firmicutes;Negativicutes;Acidaminococcales;Acidaminococcaceae;Phascolarctobacterium;uncultured bacterium | 88.4 | 1389 | -1 |
| metaRNA.PFemirge\_12069 | 69 | 0.000009 | GQ360012.1.1488 | Bacteria;Bacteroidota;Bacteroidia;Bacteroidales;Tannerellaceae;Macellibacteroides;uncultured bacterium | 90.6 | 1290 | -1 |
| metaRNA.PFemirge\_14416 | 69 | 0.000005 | KM651117.1.1438 | Bacteria;Firmicutes;Clostridia;Oscillospirales;[Eubacterium] coprostanoligenes group;uncultured Spirochaetes bacterium | 87.9 | 1260 | -1 |
| metaRNA.PFemirge\_23106 | 69 | 0.000008 | AQHW01000007.13.1527 | Bacteria;Bacteroidota;Bacteroidia;Bacteroidales;Tannerellaceae;Parabacteroides;Parabacteroides gordonii MS-1 | 94.0 | 1297 | -1 |
| metaRNA.PFemirge\_7723 | 69 | 0.000003 | AGES01000014.73256.74770 | Bacteria;Bacteroidota;Bacteroidia;Bacteroidales;Tannerellaceae;Parabacteroides;Parabacteroides sp. HGS0025 | 93.5 | 1228 | -1 |
| metaRNA.PFemirge\_6437 | 69 | 0.000006 | AB243814.1.1495 | Bacteria;Bacteroidota;Bacteroidia;Bacteroidales;Rikenellaceae;uncultured;uncultured bacterium | 89.7 | 1487 | -1 |
| metaRNA.PFemirge\_3348 | 69 | 0.000007 | KY445639.1.1525 | Bacteria;Proteobacteria;Gammaproteobacteria;Burkholderiales;Comamonadaceae;uncultured;Ideonella sp. | 95.7 | 1260 | -1 |
| metaRNA.PFemirge\_22972 | 68 | 0.000009 | LN561456.1.1394 | Bacteria;Firmicutes;Bacilli;Erysipelotrichales;Erysipelotrichaceae;Breznakia;uncultured bacterium | 93.6 | 1135 | -1 |
| metaRNA.PFemirge\_16817 | 68 | 0.000003 | DQ800771.1.1398 | Bacteria;Proteobacteria;Gammaproteobacteria;Enterobacterales;Enterobacteriaceae;Escherichia-Shigella;uncultured bacterium | 94.4 | 1394 | -1 |
| metaRNA.PFemirge\_20855 | 68 | 0.000009 | FJ901820.1.1378 | Bacteria;MBNT15;uncultured delta proteobacterium | 86.6 | 1237 | -1 |
| metaRNA.PFemirge\_12017 | 68 | 0.000004 | HQ779103.1.1440 | Bacteria;Firmicutes;Clostridia;Lachnospirales;Lachnospiraceae;Lachnospiraceae NC2004 group;uncultured organism | 88.4 | 1341 | -1 |
| metaRNA.PFemirge\_8290 | 68 | 0.000001 | GQ480160.1.1510 | Bacteria;Firmicutes;Negativicutes;Acidaminococcales;Acidaminococcaceae;Succinispira;uncultured bacterium | 90.5 | 1340 | -1 |
| metaRNA.PFemirge\_27729 | 67 | 0.000013 | HQ681866.1.1451 | Bacteria;Bacteroidota;Bacteroidia;Bacteroidales;Bacteroidaceae;Bacteroides;uncultured bacterium | 91.7 | 1380 | -1 |
| metaRNA.PFemirge\_12656 | 67 | 0.000021 | HQ767134.1.1448 | Bacteria;Bacteroidota;Bacteroidia;Bacteroidales;Tannerellaceae;Parabacteroides;uncultured organism | 95.0 | 1321 | -1 |
| metaRNA.PFemirge\_10288 | 67 | 0.000011 | DQ068853.1.1504 | Bacteria;Proteobacteria;Gammaproteobacteria;Enterobacterales;Enterobacteriaceae;Enterobacter;uncultured bacterium | 96.6 | 1390 | -1 |
| metaRNA.PFemirge\_15952 | 67 | 0.000004 | JX105679.1.1392 | Bacteria;Bacteroidota;Bacteroidia;Bacteroidales;Bacteroidaceae;Bacteroides;uncultured bacterium | 87.1 | 1313 | -1 |
| metaRNA.PFemirge\_14307 | 67 | 0.000006 | FR749935.1.1522 | Bacteria;Firmicutes;Clostridia;Lachnospirales;Lachnospiraceae;[Eubacterium] fissicatena group;Faecalicatena fissicatena | 92.4 | 1376 | -1 |
| metaRNA.PFemirge\_13718 | 67 | 0.000008 | DQ339713.1.1435 | Bacteria;Firmicutes;Clostridia;Clostridiales;Clostridiaceae;Clostridium sensu stricto 13;uncultured bacterium | 91.9 | 1345 | -1 |
| metaRNA.PFemirge\_21727 | 66 | 0.000007 | HQ815341.1.1435 | Bacteria;Firmicutes;Clostridia;Peptostreptococcales-Tissierellales;Peptostreptococcaceae;Intestinibacter;uncultured organism | 94.0 | 1283 | -1 |
| metaRNA.PFemirge\_4565 | 66 | 0.000003 | FJ868757.1.1502 | Bacteria;Bacteroidota;Bacteroidia;Bacteroidales;Porphyromonadaceae;Porphyromonas;uncultured bacterium | 93.7 | 1391 | -1 |
| metaRNA.PFemirge\_12531 | 66 | 0.000003 | AB547644.1.1485 | Bacteria;Bacteroidota;Bacteroidia;Bacteroidales;Bacteroidaceae;Bacteroides;Bacteroides oleiciplenus | 88.1 | 1376 | -1 |
| metaRNA.PFemirge\_21996 | 65 | 0.000030 | EU762749.1.1373 | Bacteria;Bacteroidota;Bacteroidia;Bacteroidales;Bacteroidaceae;Bacteroides;uncultured bacterium | 89.4 | 1192 | -1 |
| metaRNA.PFemirge\_22176 | 64 | 0.000008 | KP231747.1.1515 | Bacteria;Desulfobacterota;Desulfovibrionia;Desulfovibrionales;Desulfovibrionaceae;Desulfovibrio;uncultured Desulfovibrio sp. | 88.9 | 1373 | -1 |
| metaRNA.PFemirge\_2702 | 64 | 0.000002 | AQHW01000007.13.1527 | Bacteria;Bacteroidota;Bacteroidia;Bacteroidales;Tannerellaceae;Parabacteroides;Parabacteroides gordonii MS-1 | 91.1 | 1489 | -1 |
| metaRNA.PFemirge\_14238 | 64 | 0.000020 | HQ813653.1.1429 | Bacteria;Firmicutes;Clostridia;Oscillospirales;Ruminococcaceae;Subdoligranulum;uncultured organism | 92.9 | 1316 | -1 |
| metaRNA.PFemirge\_16199 | 64 | 0.000007 | HQ819238.1.1451 | Bacteria;Bacteroidota;Bacteroidia;Bacteroidales;Bacteroidaceae;Bacteroides;uncultured organism | 91.8 | 1224 | -1 |
| metaRNA.PFemirge\_12197 | 64 | 0.000012 | CP002345.1438824.1440339 | Bacteria;Bacteroidota;Bacteroidia;Bacteroidales;Paludibacteraceae;Paludibacter;Paludibacter propionicigenes WB4 | 88.4 | 1454 | -1 |
| metaRNA.PFemirge\_26323 | 64 | 0.000009 | EU508752.1.1287 | Bacteria;Firmicutes;Clostridia;Oscillospirales;Ruminococcaceae;uncultured;uncultured bacterium | 94.3 | 1277 | -1 |
| metaRNA.PFemirge\_28124 | 64 | 0.000008 | JX105679.1.1392 | Bacteria;Bacteroidota;Bacteroidia;Bacteroidales;Bacteroidaceae;Bacteroides;uncultured bacterium | 91.6 | 1373 | -1 |
| metaRNA.PFemirge\_27580 | 64 | 0.000012 | HM630236.1.1451 | Bacteria;Bacteroidota;Bacteroidia;Bacteroidales;Rikenellaceae;dgA-11 gut group;uncultured bacterium | 90.9 | 1202 | -1 |
| metaRNA.PFemirge\_22518 | 63 | 0.000005 | JX223620.1.1368 | Bacteria;Firmicutes;Clostridia;Clostridiales;Clostridiaceae;Clostridium sensu stricto 12;uncultured bacterium | 96.0 | 1362 | -1 |
| metaRNA.PFemirge\_4467 | 63 | 0.000005 | FR749935.1.1522 | Bacteria;Firmicutes;Clostridia;Lachnospirales;Lachnospiraceae;[Eubacterium] fissicatena group;Faecalicatena fissicatena | 95.3 | 1472 | -1 |
| metaRNA.PFemirge\_17423 | 63 | 0.000007 | DQ807770.1.1395 | Bacteria;Firmicutes;Clostridia;Lachnospirales;Lachnospiraceae;Roseburia;uncultured bacterium | 91.7 | 1393 | -1 |
| metaRNA.PFemirge\_23145 | 63 | 0.000025 | DQ793916.1.1399 | Bacteria;Bacteroidota;Bacteroidia;Bacteroidales;Bacteroidaceae;Bacteroides;uncultured bacterium | 90.6 | 1220 | -1 |
| metaRNA.PFemirge\_12393 | 63 | 0.000003 | CP009057.2296841.2298384 | Bacteria;Bacteroidota;Bacteroidia;Bacteroidales;Bacteroidaceae;Bacteroides;Bacteroides dorei | 89.3 | 1261 | -1 |
| metaRNA.PFemirge\_3834 | 62 | 0.000009 | JN038827.1.1489 | Bacteria;Proteobacteria;Gammaproteobacteria;Burkholderiales;Sutterellaceae;uncultured;uncultured beta proteobacterium | 92.1 | 1422 | -1 |
| metaRNA.PFemirge\_11772 | 62 | 0.000013 | HQ793063.1.1437 | Bacteria;Bacteroidota;Bacteroidia;Bacteroidales;Rikenellaceae;Alistipes;uncultured organism | 90.3 | 1226 | -1 |
| metaRNA.PFemirge\_26231 | 62 | 0.000007 | EU775041.1.1395 | Bacteria;Bacteroidota;Bacteroidia;Bacteroidales;Bacteroidaceae;Bacteroides;uncultured bacterium | 93.3 | 899 | -1 |
| metaRNA.PFemirge\_8356 | 62 | 0.000005 | HQ827970.1.1471 | Bacteria;Bacteroidota;Bacteroidia;Chitinophagales;Saprospiraceae;uncultured;uncultured bacterium | 99.4 | 1176 | -1 |
| metaRNA.PFemirge\_21028 | 62 | 0.000008 | JQ940836.1.1449 | Bacteria;Bacteroidota;Bacteroidia;Bacteroidales;Rikenellaceae;Alistipes;uncultured bacterium | 91.0 | 1339 | -1 |
| metaRNA.PFemirge\_23063 | 62 | 0.000006 | FJ366372.1.1363 | Bacteria;Bacteroidota;Bacteroidia;Bacteroidales;Rikenellaceae;Alistipes;uncultured bacterium | 91.4 | 1307 | -1 |
| metaRNA.PFemirge\_20799 | 61 | 0.000008 | EU772882.1.1401 | Bacteria;Firmicutes;Clostridia;Oscillospirales;Butyricicoccaceae;uncultured;uncultured bacterium | 91.5 | 1287 | -1 |
| metaRNA.PFemirge\_23902 | 61 | 0.000002 | HQ796347.1.1441 | Bacteria;Bacteroidota;Bacteroidia;Bacteroidales;Bacteroidaceae;Bacteroides;uncultured organism | 89.6 | 1285 | -1 |
| metaRNA.PFemirge\_27385 | 61 | 0.000009 | HQ793892.1.1442 | Bacteria;Bacteroidota;Bacteroidia;Bacteroidales;Bacteroidaceae;Bacteroides;uncultured organism | 91.1 | 1222 | -1 |
| metaRNA.PFemirge\_18876 | 61 | 0.000005 | FR749935.1.1522 | Bacteria;Firmicutes;Clostridia;Lachnospirales;Lachnospiraceae;[Eubacterium] fissicatena group;Faecalicatena fissicatena | 94.0 | 1297 | -1 |
| metaRNA.PFemirge\_18627 | 60 | 0.000006 | DQ798901.1.1391 | Bacteria;Firmicutes;Clostridia;Lachnospirales;Lachnospiraceae;Dorea;uncultured bacterium | 93.2 | 1303 | -1 |
| metaRNA.PFemirge\_26884 | 60 | 0.000016 | KC358008.1.1260 | Bacteria;Bacteroidota;Bacteroidia;Bacteroidales;Barnesiellaceae;uncultured;uncultured bacterium | 87.3 | 1261 | -1 |
| metaRNA.PFemirge\_11285 | 59 | 0.000004 | LT725663.1.1450 | Bacteria;Bacteroidota;Bacteroidia;Bacteroidales;Tannerellaceae;Parabacteroides;Parabacteroides sp. Marseille-P3668 | 92.3 | 1453 | -1 |
| metaRNA.PFemirge\_12065 | 59 | 0.000003 | AGES01000014.73256.74770 | Bacteria;Bacteroidota;Bacteroidia;Bacteroidales;Tannerellaceae;Parabacteroides;Parabacteroides sp. HGS0025 | 89.8 | 1447 | -1 |
| metaRNA.PFemirge\_15056 | 58 | 0.000153 | CP016762.4991311.4992838 | Bacteria;Proteobacteria;Gammaproteobacteria;Enterobacterales;Enterobacteriaceae;Citrobacter;Citrobacter freundii | 91.3 | 1387 | -1 |
| metaRNA.PFemirge\_3064 | 58 | 0.000093 | DQ793316.1.1393 | Bacteria;Bacteroidota;Bacteroidia;Bacteroidales;Tannerellaceae;Parabacteroides;uncultured bacterium | 92.6 | 1399 | -1 |
| metaRNA.PFemirge\_8465 | 58 | 0.000009 | EU469747.1.1392 | Bacteria;Firmicutes;Clostridia;Lachnospirales;Lachnospiraceae;28-4;uncultured bacterium | 91.3 | 1306 | -1 |
| metaRNA.PFemirge\_12306 | 58 | 0.000010 | JQ924024.1.1501 | Bacteria;Proteobacteria;Gammaproteobacteria;Burkholderiales;Rhodocyclaceae;Dechlorosoma;uncultured bacterium | 93.9 | 1389 | -1 |
| metaRNA.PFemirge\_20655 | 58 | 0.000007 | EU508987.1.1386 | Bacteria;Firmicutes;Clostridia;Oscillospirales;Ruminococcaceae;Ruminococcus;uncultured bacterium | 92.5 | 1377 | -1 |
| metaRNA.PFemirge\_23233 | 58 | 0.000006 | HQ761746.1.1448 | Bacteria;Bacteroidota;Bacteroidia;Bacteroidales;Tannerellaceae;Parabacteroides;uncultured organism | 88.0 | 1357 | -1 |
| metaRNA.PFemirge\_3615 | 58 | 0.000018 | FJ393116.1.1506 | Bacteria;Proteobacteria;Gammaproteobacteria;Burkholderiales;Rhodocyclaceae;Azospira;uncultured Azospira sp. | 93.5 | 1423 | -1 |
| metaRNA.PFemirge\_24886 | 57 | 0.000009 | JN680616.1.1459 | Bacteria;Firmicutes;Clostridia;Oscillospirales;Butyricicoccaceae;UCG-009;uncultured Ruminococcaceae bacterium | 93.0 | 1162 | -1 |
| metaRNA.PFemirge\_7707 | 57 | 0.000007 | EU104138.1.1438 | Bacteria;Bacteroidota;Bacteroidia;Chitinophagales;Saprospiraceae;Haliscomenobacter;uncultured bacterium | 99.7 | 1187 | -1 |
| metaRNA.PFemirge\_26455 | 57 | 0.000004 | EU765828.1.1335 | Bacteria;Firmicutes;Clostridia;Oscillospirales;Ruminococcaceae;Faecalibacterium;uncultured bacterium | 90.1 | 1281 | -1 |
| metaRNA.PFemirge\_25787 | 57 | 0.000008 | FR799325.1.1443 | Bacteria;Proteobacteria;Alphaproteobacteria;Rhizobiales;Xanthobacteraceae;Azorhizobium;Azorhizobium oxalatiphilum | 89.0 | 1186 | -1 |
| metaRNA.PFemirge\_5371 | 57 | 0.000002 | DQ817541.1.1479 | Bacteria;Proteobacteria;Gammaproteobacteria;Burkholderiales;Chromobacteriaceae;Crenobacter;uncultured bacterium | 92.1 | 1455 | -1 |
| metaRNA.PFemirge\_20200 | 57 | 0.000004 | AATF01000761.367.1874 | Bacteria;Firmicutes;Clostridia;Oscillospirales;Ruminococcaceae;Ruminococcus;mouse gut metagenome | 93.0 | 1294 | -1 |
| metaRNA.PFemirge\_8156 | 56 | 0.000003 | CP014223.544406.545927 | Bacteria;Firmicutes;Clostridia;Lachnospirales;Lachnospiraceae;Anaerostignum;Anaerotignum propionicum DSM 1682 | 91.3 | 1396 | -1 |
| metaRNA.PFemirge\_2928 | 56 | 0.000009 | HM779545.1.1509 | Bacteria;Proteobacteria;Gammaproteobacteria;Pseudomonadales;Pseudomonadaceae;Pseudomonas;uncultured bacterium | 99.8 | 1268 | -1 |
| metaRNA.PFemirge\_17978 | 56 | 0.000014 | FR749935.1.1522 | Bacteria;Firmicutes;Clostridia;Lachnospirales;Lachnospiraceae;[Eubacterium] fissicatena group;Faecalicatena fissicatena | 93.3 | 1389 | -1 |
| metaRNA.PFemirge\_6196 | 56 | 0.000004 | CXSE01001446.1371.2879 | Bacteria;Bacteroidota;Bacteroidia;Bacteroidales;Bacteroidaceae;Bacteroides;human gut metagenome | 90.7 | 1305 | -1 |
| metaRNA.PFemirge\_23142 | 56 | 0.000006 | FR669217.1.1452 | Bacteria;Firmicutes;Clostridia;Oscillospirales;Oscillospiraceae;Colidextribacter;uncultured bacterium | 92.6 | 1277 | -1 |
| metaRNA.PFemirge\_18330 | 56 | 0.000006 | LT598589.1.1424 | Bacteria;Firmicutes;Clostridia;Oscillospirales;Butyricicoccaceae;UCG-008;Intestinibacillus massiliensis | 92.8 | 1290 | -1 |
| metaRNA.PFemirge\_2123 | 56 | 0.000004 | HG917472.1.1448 | Bacteria;Proteobacteria;Gammaproteobacteria;Burkholderiales;Chitinimonadaceae;Chitinivorax;uncultured bacterium | 96.6 | 1284 | -1 |
| metaRNA.PFemirge\_23882 | 56 | 0.000006 | HM630236.1.1451 | Bacteria;Bacteroidota;Bacteroidia;Bacteroidales;Rikenellaceae;dgA-11 gut group;uncultured bacterium | 88.9 | 1357 | -1 |
| metaRNA.PFemirge\_23333 | 55 | 0.000004 | DQ793316.1.1393 | Bacteria;Bacteroidota;Bacteroidia;Bacteroidales;Tannerellaceae;Parabacteroides;uncultured bacterium | 88.3 | 1360 | -1 |
| metaRNA.PFemirge\_13924 | 55 | 0.000022 | AGXU01000010.481004.482520 | Bacteria;Bacteroidota;Bacteroidia;Bacteroidales;Bacteroidaceae;Bacteroides;Bacteroides ovatus CL03T12C18 | 93.1 | 1374 | -1 |
| metaRNA.PFemirge\_2115 | 54 | 0.000008 | FR749935.1.1522 | Bacteria;Firmicutes;Clostridia;Lachnospirales;Lachnospiraceae;[Eubacterium] fissicatena group;Faecalicatena fissicatena | 94.6 | 1470 | -1 |
| metaRNA.PFemirge\_6529 | 54 | 0.000007 | GQ871713.1.1490 | Bacteria;Bacteroidota;Bacteroidia;Bacteroidales;Rikenellaceae;Alistipes;uncultured bacterium | 93.2 | 1291 | -1 |
| metaRNA.PFemirge\_1876 | 54 | 0.000003 | AB649277.1.1490 | Bacteria;Firmicutes;Bacilli;Erysipelotrichales;Erysipelotrichaceae;Faecalitalea;Absiella dolichum | 94.3 | 1170 | -1 |
| metaRNA.PFemirge\_2017 | 54 | 0.000013 | EF403674.1.1517 | Bacteria;Firmicutes;Negativicutes;Acidaminococcales;Acidaminococcaceae;Phascolarctobacterium;uncultured bacterium | 90.9 | 1512 | -1 |
| metaRNA.PFemirge\_20039 | 54 | 0.000011 | EU765284.1.1347 | Bacteria;Firmicutes;Clostridia;Lachnospirales;Lachnospiraceae;Lachnospiraceae UCG-008;uncultured bacterium | 92.3 | 1265 | -1 |
| metaRNA.PFemirge\_25432 | 54 | 0.000006 | GQ133618.1.1320 | Bacteria;Firmicutes;Bacilli;Erysipelotrichales;Erysipelotrichaceae;Erysipelothrix;uncultured bacterium | 91.6 | 1277 | -1 |
| metaRNA.PFemirge\_27746 | 54 | 0.000012 | FQSE01000007.1631802.1633308 | Bacteria;Bacteroidota;Bacteroidia;Bacteroidales;Tannerellaceae;Parabacteroides;Parabacteroides sp. Marseille-P3136 | 91.9 | 1225 | -1 |
| metaRNA.PFemirge\_27927 | 54 | 0.000007 | JN680650.1.1446 | Bacteria;Firmicutes;Clostridia;Oscillospirales;Ruminococcaceae;Ruminococcus;uncultured Lachnospiraceae bacterium | 92.3 | 1406 | -1 |
| metaRNA.PFemirge\_20181 | 53 | 0.000003 | AATF01000761.367.1874 | Bacteria;Firmicutes;Clostridia;Oscillospirales;Ruminococcaceae;Ruminococcus;mouse gut metagenome | 90.7 | 1304 | -1 |
| metaRNA.PFemirge\_26250 | 53 | 0.000011 | AB599904.1.1384 | Bacteria;Proteobacteria;Alphaproteobacteria;Sphingomonadales;Sphingomonadaceae;Novosphingobium;alpha proteobacterium INB3 | 96.8 | 1206 | -1 |
| metaRNA.PFemirge\_8438 | 53 | 0.000003 | AF245350.1.1489 | Bacteria;Proteobacteria;Gammaproteobacteria;Burkholderiales;Rhodocyclaceae;uncultured;uncultured bacterium SA35 | 95.1 | 1459 | -1 |
| metaRNA.PFemirge\_10840 | 53 | 0.000008 | JX198651.1.1513 | Bacteria;Firmicutes;Clostridia;Oscillospirales;Butyricicoccaceae;UCG-009;uncultured bacterium | 91.2 | 1241 | -1 |
| metaRNA.PFemirge\_4063 | 53 | 0.000001 | JF808852.1.1503 | Bacteria;Proteobacteria;Gammaproteobacteria;Burkholderiales;Rhodocyclaceae;Azospira;uncultured bacterium | 94.7 | 1270 | -1 |
| metaRNA.PFemirge\_30263 | 53 | 0.000013 | KP231747.1.1515 | Bacteria;Desulfobacterota;Desulfovibrionia;Desulfovibrionales;Desulfovibrionaceae;Desulfovibrio;uncultured Desulfovibrio sp. | 92.2 | 1504 | -1 |
| metaRNA.PFemirge\_11626 | 53 | 0.000007 | HQ793063.1.1437 | Bacteria;Bacteroidota;Bacteroidia;Bacteroidales;Rikenellaceae;Alistipes;uncultured organism | 89.7 | 1345 | -1 |
| metaRNA.PFemirge\_8104 | 53 | 0.000009 | MG367106.1.1496 | Bacteria;Firmicutes;Clostridia;Lachnospirales;Lachnospiraceae;Lachnoclostridium;uncultured Clostridiales Family XIV bacterium | 92.1 | 1400 | -1 |
| metaRNA.PFemirge\_16372 | 53 | 0.000003 | EU775986.1.1401 | Bacteria;Bacteroidota;Bacteroidia;Bacteroidales;Bacteroidaceae;Bacteroides;uncultured bacterium | 93.2 | 1396 | -1 |
| metaRNA.PFemirge\_6115 | 52 | 0.000010 | HAEN01058188.41.1551 | Bacteria;Proteobacteria;Gammaproteobacteria;Burkholderiales;Comamonadaceae;Rhizobacter;Bragasellus peltatus | 95.2 | 1409 | -1 |
| metaRNA.PFemirge\_21042 | 52 | 0.000014 | CBVB010000006.254456.255954 | Bacteria;Bacteroidota;Bacteroidia;Bacteroidales;Bacteroidaceae;Bacteroides;Bacteroidaceae bacterium MS4 | 90.8 | 721 | -1 |
| metaRNA.PFemirge\_3497 | 52 | 0.000015 | FPLS01048670.18.1522 | Bacteria;Bacteroidota;Bacteroidia;Bacteroidales;Paludibacteraceae;uncultured;metagenome | 87.8 | 1348 | -1 |
| metaRNA.PFemirge\_23846 | 52 | 0.000006 | HQ761714.1.1441 | Bacteria;Bacteroidota;Bacteroidia;Bacteroidales;Tannerellaceae;Parabacteroides;uncultured organism | 87.2 | 1360 | -1 |
| metaRNA.PFemirge\_12029 | 51 | 0.000002 | FLKF01000010.1355.2888 | Bacteria;Firmicutes;Negativicutes;Veillonellales-Selenomonadales;Sporomusaceae;Anaerosinus;Clostridiales bacterium Marseille-P2411 | 89.6 | 1357 | -1 |
| metaRNA.PFemirge\_22739 | 51 | 0.000005 | EU775026.1.1400 | Bacteria;Bacteroidota;Bacteroidia;Bacteroidales;Bacteroidaceae;Bacteroides;uncultured bacterium | 88.7 | 1314 | -1 |
| metaRNA.PFemirge\_12661 | 51 | 0.000008 | HQ800643.1.1431 | Bacteria;Bacteroidota;Bacteroidia;Bacteroidales;Bacteroidaceae;Bacteroides;uncultured organism | 93.9 | 1205 | -1 |
| metaRNA.PFemirge\_1497 | 51 | 0.000007 | LN995859.1.1478 | Bacteria;Verrucomicrobiota;Chlamydiae;Chlamydiales;Parachlamydiaceae;uncultured;Neochlamydia sp. Trut23-12-2015\_Venoge-Embouchure | 92.1 | 1172 | -1 |
| metaRNA.PFemirge\_14697 | 50 | 0.000010 | FJ880485.1.1495 | Bacteria;Firmicutes;Clostridia;Lachnospirales;Lachnospiraceae;uncultured;uncultured bacterium | 91.3 | 1284 | -1 |
| metaRNA.PFemirge\_530 | 50 | 0.000008 | KT072190.1.1634 | Eukaryota;SAR;Stramenopiles;Labyrinthulomycetes;Amphitraemidae;uncultured eukaryote | 99.3 | 685 | -1 |
| metaRNA.PFemirge\_5247 | 50 | 0.000004 | LC036226.1.1496 | Bacteria;Bacteroidota;Bacteroidia;Bacteroidales;Paludibacteraceae;H1;uncultured Paludibacter sp. | 86.4 | 1303 | -1 |
| metaRNA.PFemirge\_1882 | 50 | 0.000004 | AY957923.1.1520 | Bacteria;Proteobacteria;Gammaproteobacteria;Burkholderiales;Comamonadaceae;Curvibacter;uncultured bacterium | 94.6 | 1416 | -1 |
| metaRNA.PFemirge\_18240 | 50 | 0.000015 | X71855.1.1501 | Bacteria;Firmicutes;Clostridia;Lachnospirales;Lachnospiraceae;Lachnoclostridium;[Clostridium] xylanolyticum | 92.3 | 1292 | -1 |
| metaRNA.PFemirge\_7726 | 50 | 0.000003 | AB517712.1.1448 | Bacteria;Bacteroidota;Bacteroidia;Bacteroidales;Marinilabiliaceae;Saccharicrinis;Saccharicrinis fermentans | 88.2 | 1387 | -1 |
| metaRNA.PFemirge\_24603 | 50 | 0.000004 | HQ748426.1.1449 | Bacteria;Firmicutes;Clostridia;Lachnospirales;Lachnospiraceae;Lachnoclostridium;uncultured organism | 92.9 | 1284 | -1 |
| metaRNA.PFemirge\_5015 | 50 | 0.000007 | HQ766029.1.1454 | Bacteria;Firmicutes;Clostridia;Oscillospirales;Ruminococcaceae;Subdoligranulum;uncultured organism | 88.9 | 1448 | -1 |
| metaRNA.PFemirge\_21756 | 49 | 0.000003 | GQ449092.1.1375 | Bacteria;Firmicutes;Clostridia;Lachnospirales;Lachnospiraceae;Tyzzerella;uncultured bacterium | 93.5 | 1286 | -1 |
| metaRNA.PFemirge\_22906 | 49 | 0.000002 | FQ658622.1.1363 | Bacteria;Proteobacteria;Gammaproteobacteria;Burkholderiales;Hydrogenophilaceae;Thiobacillus;uncultured soil bacterium | 87.0 | 1258 | -1 |
| metaRNA.PFemirge\_21161 | 49 | 0.000003 | JQ317253.1.1380 | Bacteria;Bacteroidota;Bacteroidia;Bacteroidales;Bacteroidaceae;Bacteroides;Bacteroides sp. CannelCatfish9 | 90.6 | 1380 | -1 |
| metaRNA.PFemirge\_15432 | 49 | 0.000003 | KY190699.1.1411 | Bacteria;Proteobacteria;Alphaproteobacteria;Sphingomonadales;Sphingomonadaceae;Sphingomonas;uncultured bacterium | 91.8 | 1312 | -1 |
| metaRNA.PFemirge\_26303 | 49 | 0.000003 | CR933273.1.1287 | Bacteria;Firmicutes;Clostridia;Lachnospirales;Lachnospiraceae;Lachnospiraceae UCG-004;uncultured bacterium | 93.3 | 1282 | -1 |
| metaRNA.PFemirge\_16410 | 49 | 0.000002 | AB574482.1.1489 | Bacteria;Bacteroidota;Bacteroidia;Bacteroidales;Tannerellaceae;Parabacteroides;Parabacteroides chinchillae | 91.0 | 1302 | -1 |
| metaRNA.PFemirge\_6250 | 49 | 0.000010 | MG367100.1.1491 | Bacteria;Firmicutes;Clostridia;Oscillospirales;Oscillospiraceae;Colidextribacter;uncultured Clostridiales Family IV bacterium | 94.1 | 1291 | -1 |
| metaRNA.PFemirge\_27705 | 49 | 0.000033 | HM630236.1.1451 | Bacteria;Bacteroidota;Bacteroidia;Bacteroidales;Rikenellaceae;dgA-11 gut group;uncultured bacterium | 91.6 | 1405 | -1 |
| metaRNA.PFemirge\_15683 | 49 | 0.000008 | DQ809572.1.1407 | Bacteria;Firmicutes;Bacilli;Erysipelotrichales;Erysipelotrichaceae;Holdemanella;uncultured bacterium | 91.4 | 1320 | -1 |
| metaRNA.PFemirge\_25233 | 48 | 0.000006 | JN680650.1.1446 | Bacteria;Firmicutes;Clostridia;Oscillospirales;Ruminococcaceae;Ruminococcus;uncultured Lachnospiraceae bacterium | 93.3 | 1269 | -1 |
| metaRNA.PFemirge\_3096 | 48 | 0.000003 | JTDA01000039.175.1712 | Bacteria;Bacteroidota;Bacteroidia;Bacteroidales;Barnesiellaceae;Coprobacter;Coprobacter secundus | 89.0 | 1493 | -1 |
| metaRNA.PFemirge\_26046 | 48 | 0.000009 | FR669217.1.1452 | Bacteria;Firmicutes;Clostridia;Oscillospirales;Oscillospiraceae;Colidextribacter;uncultured bacterium | 91.6 | 1293 | -1 |
| metaRNA.PFemirge\_21241 | 48 | 0.000018 | GQ451215.1.1487 | Bacteria;Firmicutes;Clostridia;Oscillospirales;Ruminococcaceae;Ruminococcus;uncultured bacterium | 94.2 | 1306 | -1 |
| metaRNA.PFemirge\_15025 | 48 | 0.000015 | FR749935.1.1522 | Bacteria;Firmicutes;Clostridia;Lachnospirales;Lachnospiraceae;[Eubacterium] fissicatena group;Faecalicatena fissicatena | 93.0 | 1440 | -1 |
| metaRNA.PFemirge\_9234 | 48 | 0.000003 | CP012068.1576987.1578505 | Bacteria;Firmicutes;Clostridia;Peptostreptococcales-Tissierellales;Anaerovoracaceae;[Eubacterium] nodatum group;Eubacterium sulci ATCC 35585 | 92.2 | 1205 | -1 |
| metaRNA.PFemirge\_19080 | 48 | 0.000003 | HQ778371.1.1457 | Bacteria;Firmicutes;Clostridia;Lachnospirales;Lachnospiraceae;Lachnoclostridium;uncultured organism | 88.9 | 1188 | -1 |
| metaRNA.PFemirge\_21667 | 48 | 0.000015 | HQ792908.1.1451 | Bacteria;Firmicutes;Clostridia;Oscillospirales;Butyricicoccaceae;UCG-008;uncultured organism | 89.7 | 1274 | -1 |
| metaRNA.PFemirge\_23053 | 48 | 0.000013 | HQ748426.1.1449 | Bacteria;Firmicutes;Clostridia;Lachnospirales;Lachnospiraceae;Lachnoclostridium;uncultured organism | 91.5 | 1284 | -1 |
| metaRNA.PFemirge\_18993 | 48 | 0.000006 | DQ798990.1.1390 | Bacteria;Firmicutes;Clostridia;Lachnospirales;Lachnospiraceae;[Ruminococcus] torques group;uncultured bacterium | 92.4 | 1285 | -1 |
| metaRNA.PFemirge\_15303 | 48 | 0.000004 | EF098082.1.1412 | Bacteria;Bacteroidota;Bacteroidia;Bacteroidales;Muribaculaceae;uncultured bacterium | 87.8 | 1406 | -1 |
| metaRNA.PFemirge\_7796 | 48 | 0.000017 | FJ393129.1.1501 | Bacteria;Proteobacteria;Gammaproteobacteria;Burkholderiales;Rhodocyclaceae;Azospira;uncultured Azospira sp. | 91.5 | 1377 | -1 |
| metaRNA.PFemirge\_23222 | 47 | 0.000005 | CBVB010000006.254456.255954 | Bacteria;Bacteroidota;Bacteroidia;Bacteroidales;Bacteroidaceae;Bacteroides;Bacteroidaceae bacterium MS4 | 92.8 | 1143 | -1 |
| metaRNA.PFemirge\_10860 | 47 | 0.000017 | GQ480160.1.1510 | Bacteria;Firmicutes;Negativicutes;Acidaminococcales;Acidaminococcaceae;Succinispira;uncultured bacterium | 90.6 | 1364 | -1 |
| metaRNA.PFemirge\_14643 | 47 | 0.000003 | JN680597.1.1451 | Bacteria;Bacteroidota;Bacteroidia;Bacteroidales;Rikenellaceae;Alistipes;uncultured Rikenellaceae bacterium | 91.5 | 1406 | -1 |
| metaRNA.PFemirge\_4742 | 47 | 0.000004 | GU293185.1.1498 | Bacteria;Proteobacteria;Gammaproteobacteria;Enterobacterales;Alteromonadaceae;Rheinheimera;uncultured bacterium | 94.1 | 1491 | -1 |
| metaRNA.PFemirge\_13047 | 47 | 0.000042 | HQ760530.1.1440 | Bacteria;Bacteroidota;Bacteroidia;Bacteroidales;Tannerellaceae;Parabacteroides;uncultured organism | 89.9 | 1409 | -1 |
| metaRNA.PFemirge\_22967 | 46 | 0.000002 | DQ809859.1.1362 | Bacteria;Firmicutes;Clostridia;Oscillospirales;Oscillospiraceae;uncultured;uncultured bacterium | 92.0 | 1355 | -1 |
| metaRNA.PFemirge\_25146 | 46 | 0.000005 | AY983363.1.1360 | Bacteria;Bacteroidota;Bacteroidia;Bacteroidales;Bacteroidaceae;Bacteroides;uncultured bacterium | 92.6 | 1224 | -1 |
| metaRNA.PFemirge\_21949 | 46 | 0.000007 | DQ799101.1.1373 | Bacteria;Firmicutes;Clostridia;Oscillospirales;Ruminococcaceae;Faecalibacterium;uncultured bacterium | 92.5 | 1287 | -1 |
| metaRNA.PFemirge\_17569 | 46 | 0.000004 | FJ715990.1.1502 | Bacteria;Proteobacteria;Gammaproteobacteria;Enterobacterales;Aeromonadaceae;Aeromonas;uncultured bacterium | 92.6 | 1401 | -1 |
| metaRNA.PFemirge\_30249 | 46 | 0.000019 | AY452004.1.1502 | Bacteria;Firmicutes;Clostridia;Lachnospirales;Lachnospiraceae;Lachnospiraceae UCG-004;uncultured bacterium | 86.2 | 1336 | -1 |
| metaRNA.PFemirge\_19061 | 46 | 0.000014 | FR749936.1.1524 | Bacteria;Firmicutes;Clostridia;Lachnospirales;Lachnospiraceae;[Eubacterium] fissicatena group;Faecalicatena fissicatena | 93.7 | 1386 | -1 |
| metaRNA.PFemirge\_12401 | 46 | 0.000009 | HM159970.1.1478 | Bacteria;Proteobacteria;Gammaproteobacteria;Enterobacterales;Aeromonadaceae;Aeromonas;uncultured Aeromonas sp. | 92.3 | 1254 | -1 |
| metaRNA.PFemirge\_21922 | 45 | 0.000006 | DQ794241.1.1398 | Bacteria;Bacteroidota;Bacteroidia;Bacteroidales;Rikenellaceae;Alistipes;uncultured bacterium | 87.5 | 1227 | -1 |
| metaRNA.PFemirge\_19200 | 45 | 0.000005 | EU772114.1.1390 | Bacteria;Firmicutes;Clostridia;Peptostreptococcales-Tissierellales;Anaerovoracaceae;[Eubacterium] nodatum group;uncultured bacterium | 92.7 | 1389 | -1 |
| metaRNA.PFemirge\_24794 | 45 | 0.000006 | HM630236.1.1451 | Bacteria;Bacteroidota;Bacteroidia;Bacteroidales;Rikenellaceae;dgA-11 gut group;uncultured bacterium | 87.7 | 1213 | -1 |
| metaRNA.PFemirge\_26690 | 45 | 0.000003 | LRVM01000024.42.1563 | Bacteria;Firmicutes;Clostridia;Lachnospirales;Lachnospiraceae;Anaerostignum;[Clostridium] neopropionicum | 93.7 | 1212 | -1 |
| metaRNA.PFemirge\_13003 | 45 | 0.000004 | CP003153.1901384.1902805 | Bacteria;Proteobacteria;Gammaproteobacteria;Burkholderiales;Rhodocyclaceae;Azospira;Azospira oryzae PS | 92.7 | 1270 | -1 |
| metaRNA.PFemirge\_12307 | 44 | 0.000001 | JN688021.1.1446 | Bacteria;Bacteroidota;Bacteroidia;Bacteroidales;Rikenellaceae;uncultured;bacterium enrichment culture clone M02 | 87.5 | 1324 | -1 |
| metaRNA.PFemirge\_27292 | 44 | 0.000002 | EU772602.1.1240 | Bacteria;Proteobacteria;Gammaproteobacteria;Enterobacterales;Enterobacteriaceae;Escherichia-Shigella;uncultured bacterium | 95.4 | 1120 | -1 |
| metaRNA.PFemirge\_30256 | 44 | 0.000020 | HQ815291.1.1415 | Bacteria;Firmicutes;Clostridia;Lachnospirales;Lachnospiraceae;[Ruminococcus] gauvreauii group;uncultured organism | 91.2 | 1291 | -1 |
| metaRNA.PFemirge\_20156 | 44 | 0.000004 | KU161317.1.1411 | Bacteria;Proteobacteria;Gammaproteobacteria;Enterobacterales;Enterobacteriaceae;Citrobacter;Yokenella regensburgei | 95.8 | 981 | -1 |
| metaRNA.PFemirge\_8284 | 44 | 0.000003 | LN810638.1.1401 | Bacteria;Bacteroidota;Bacteroidia;Cytophagales;Spirosomaceae;Flectobacillus;Flectobacillus rhizosphaerae | 94.2 | 1145 | -1 |
| metaRNA.PFemirge\_5576 | 44 | 0.000002 | GQ871737.1.1494 | Bacteria;Firmicutes;Clostridia;Oscillospirales;Ruminococcaceae;Anaerofilum;uncultured bacterium | 91.4 | 1341 | -1 |
| metaRNA.PFemirge\_3087 | 44 | 0.000005 | JF513192.1.1510 | Bacteria;Proteobacteria;Gammaproteobacteria;Pseudomonadales;Moraxellaceae;Acinetobacter;Acinetobacter baumannii | 96.1 | 1498 | -1 |
| metaRNA.PFemirge\_26363 | 44 | 0.000004 | EU774348.1.1286 | Bacteria;Firmicutes;Clostridia;Clostridiales;Clostridiaceae;Clostridium sensu stricto 1;uncultured bacterium | 94.8 | 1245 | -1 |
| metaRNA.PFemirge\_25653 | 44 | 0.000004 | JN038827.1.1489 | Bacteria;Proteobacteria;Gammaproteobacteria;Burkholderiales;Sutterellaceae;uncultured;uncultured beta proteobacterium | 91.0 | 1207 | -1 |
| metaRNA.PFemirge\_5707 | 44 | 0.000006 | KX508033.1.1494 | Bacteria;Proteobacteria;Gammaproteobacteria;Burkholderiales;Comamonadaceae;Aquabacterium;uncultured bacterium | 95.5 | 1411 | -1 |
| metaRNA.PFemirge\_10801 | 44 | 0.000001 | HQ774646.1.1457 | Bacteria;Proteobacteria;Gammaproteobacteria;Burkholderiales;Sutterellaceae;Parasutterella;uncultured organism | 91.9 | 1366 | -1 |
| metaRNA.PFemirge\_16721 | 44 | 0.000004 | EU775302.1.1379 | Bacteria;Firmicutes;Clostridia;Lachnospirales;Lachnospiraceae;Lachnospiraceae AC2044 group;uncultured bacterium | 92.5 | 1304 | -1 |
| metaRNA.PFemirge\_20165 | 43 | 0.000003 | DQ793508.1.1385 | Bacteria;Firmicutes;Clostridia;Oscillospirales;UCG-011;uncultured bacterium | 92.4 | 1288 | -1 |
| metaRNA.PFemirge\_26665 | 43 | 0.000003 | CR933278.1.1273 | Bacteria;Firmicutes;Clostridia;Lachnospirales;Lachnospiraceae;Lachnospiraceae UCG-004;uncultured bacterium | 92.5 | 1174 | -1 |
| metaRNA.PFemirge\_16878 | 43 | 0.000013 | AM230650.1.1368 | Bacteria;Bacteroidota;Bacteroidia;Bacteroidales;Bacteroidaceae;Bacteroides;Bacteroides xylanisolvens | 88.5 | 1303 | -1 |
| metaRNA.PFemirge\_23135 | 43 | 0.000009 | EU475681.1.1364 | Bacteria;Firmicutes;Clostridia;Peptostreptococcales-Tissierellales;Peptostreptococcaceae;uncultured;uncultured bacterium | 94.0 | 1361 | -1 |
| metaRNA.PFemirge\_27006 | 42 | 0.000032 | X81125.1.1482 | Bacteria;Firmicutes;Clostridia;Oscillospirales;Oscillospiraceae;Colidextribacter;[Clostridium] viride | 92.8 | 1233 | -1 |
| metaRNA.PFemirge\_13282 | 42 | 0.000002 | GQ360012.1.1488 | Bacteria;Bacteroidota;Bacteroidia;Bacteroidales;Tannerellaceae;Macellibacteroides;uncultured bacterium | 86.9 | 1428 | -1 |
| metaRNA.PFemirge\_21473 | 42 | 0.000005 | EU465560.1.1391 | Bacteria;Firmicutes;Clostridia;Lachnospirales;Lachnospiraceae;Lachnospiraceae AC2044 group;uncultured bacterium | 91.7 | 1319 | -1 |
| metaRNA.PFemirge\_8488 | 42 | 0.000006 | HM445966.1.1478 | Bacteria;Firmicutes;Bacilli;Erysipelotrichales;Erysipelotrichaceae;Erysipelothrix;uncultured bacterium | 91.9 | 1408 | -1 |
| metaRNA.PFemirge\_2664 | 42 | 0.000003 | JN868790.1.1523 | Bacteria;Proteobacteria;Gammaproteobacteria;Burkholderiales;Comamonadaceae;uncultured;uncultured bacterium | 95.7 | 1387 | -1 |
| metaRNA.PFemirge\_24494 | 41 | 0.000005 | EU777237.1.1343 | Bacteria;Firmicutes;Clostridia;Lachnospirales;Lachnospiraceae;Lachnospiraceae UCG-008;uncultured bacterium | 92.6 | 1344 | -1 |
| metaRNA.PFemirge\_11292 | 41 | 0.000008 | AB574482.1.1489 | Bacteria;Bacteroidota;Bacteroidia;Bacteroidales;Tannerellaceae;Parabacteroides;Parabacteroides chinchillae | 87.5 | 1389 | -1 |
| metaRNA.PFemirge\_17982 | 41 | 0.000003 | GQ898302.1.1489 | Bacteria;Bacteroidota;Bacteroidia;Bacteroidales;Bacteroidaceae;Bacteroides;uncultured bacterium | 89.9 | 1393 | -1 |
| metaRNA.PFemirge\_3979 | 41 | 0.000007 | AY452004.1.1502 | Bacteria;Firmicutes;Clostridia;Lachnospirales;Lachnospiraceae;Lachnospiraceae UCG-004;uncultured bacterium | 89.2 | 1295 | -1 |
| metaRNA.PFemirge\_16743 | 41 | 0.000008 | CP003153.1901384.1902805 | Bacteria;Proteobacteria;Gammaproteobacteria;Burkholderiales;Rhodocyclaceae;Azospira;Azospira oryzae PS | 96.2 | 1388 | -1 |
| metaRNA.PFemirge\_16362 | 41 | 0.000010 | EU772907.1.1401 | Bacteria;Bacteroidota;Bacteroidia;Bacteroidales;Bacteroidaceae;Bacteroides;uncultured bacterium | 92.2 | 1391 | -1 |
| metaRNA.PFemirge\_13476 | 41 | 0.000005 | FR749935.1.1522 | Bacteria;Firmicutes;Clostridia;Lachnospirales;Lachnospiraceae;[Eubacterium] fissicatena group;Faecalicatena fissicatena | 94.1 | 1295 | -1 |
| metaRNA.PFemirge\_21142 | 40 | 0.000004 | AB610559.1.1481 | Bacteria;Firmicutes;Clostridia;Lachnospirales;Lachnospiraceae;Lachnoclostridium;Clostridium sp. K7 | 94.0 | 1285 | -1 |
| metaRNA.PFemirge\_9376 | 40 | 0.000013 | EF688147.1.1469 | Bacteria;Firmicutes;Clostridia;Oscillospirales;UCG-010;uncultured bacterium | 93.1 | 1080 | -1 |
| metaRNA.PFemirge\_35 | 40 | 0.000002 | AF011457.1.2048 | Eukaryota;Discoba;Discicristata;Heterolobosea;Tetramitia;Naegleria;Naegleria galeacystis | 99.6 | 1743 | -1 |
| metaRNA.PFemirge\_22960 | 40 | 0.000002 | AM936169.1.1362 | Bacteria;Proteobacteria;Gammaproteobacteria;Immundisolibacterales;Immundisolibacteraceae;Immundisolibacter;uncultured Chromatiales bacterium | 92.9 | 1289 | -1 |
| metaRNA.PFemirge\_22551 | 40 | 0.000002 | CP014223.544406.545927 | Bacteria;Firmicutes;Clostridia;Lachnospirales;Lachnospiraceae;Anaerostignum;Anaerotignum propionicum DSM 1682 | 91.7 | 1288 | -1 |
| metaRNA.PFemirge\_18865 | 40 | 0.000003 | EU772683.1.1391 | Bacteria;Firmicutes;Clostridia;Lachnospirales;Lachnospiraceae;Blautia;uncultured bacterium | 92.7 | 1239 | -1 |
| metaRNA.PFemirge\_20021 | 40 | 0.000069 | FJ393126.1.1488 | Bacteria;Bacteroidota;Bacteroidia;Bacteroidales;Bacteroidaceae;Bacteroides;uncultured Bacteroides sp. | 90.8 | 1234 | -1 |
| metaRNA.PFemirge\_22735 | 40 | 0.000002 | DQ799369.1.1397 | Bacteria;Bacteroidota;Bacteroidia;Bacteroidales;Bacteroidaceae;Bacteroides;uncultured bacterium | 91.7 | 1193 | -1 |
| metaRNA.PFemirge\_3320 | 39 | 0.000004 | KP411849.1.1479 | Bacteria;Myxococcota;Myxococcia;Myxococcales;Myxococcaceae;P3OB-42;uncultured bacterium | 99.3 | 1227 | -1 |
| metaRNA.PFemirge\_11130 | 39 | 0.000005 | HQ793691.1.1450 | Bacteria;Bacteroidota;Bacteroidia;Bacteroidales;Bacteroidaceae;Bacteroides;uncultured organism | 88.5 | 1231 | -1 |
| metaRNA.PFemirge\_23880 | 39 | 0.000002 | JX198651.1.1513 | Bacteria;Firmicutes;Clostridia;Oscillospirales;Butyricicoccaceae;UCG-009;uncultured bacterium | 91.4 | 1164 | -1 |
| metaRNA.PFemirge\_1784 | 39 | 0.000005 | CP002345.1438824.1440339 | Bacteria;Bacteroidota;Bacteroidia;Bacteroidales;Paludibacteraceae;Paludibacter;Paludibacter propionicigenes WB4 | 87.9 | 1335 | -1 |
| metaRNA.PFemirge\_22064 | 39 | 0.000005 | KF104611.1.1373 | Bacteria;Proteobacteria;Gammaproteobacteria;Xanthomonadales;Xanthomonadaceae;Stenotrophomonas;uncultured bacterium | 95.0 | 1372 | -1 |
| metaRNA.PFemirge\_19862 | 39 | 0.000005 | BAAY01003437.486.1979 | Bacteria;Firmicutes;Clostridia;Lachnospirales;Lachnospiraceae;uncultured;human gut metagenome | 91.9 | 1291 | -1 |
| metaRNA.PFemirge\_1632 | 39 | 0.000008 | FPLL01003728.5.1485 | Bacteria;Proteobacteria;Gammaproteobacteria;Xanthomonadales;Rhodanobacteraceae;Tahibacter;metagenome | 95.1 | 1290 | -1 |
| metaRNA.PFemirge\_28072 | 39 | 0.000003 | AY212697.1.1526 | Bacteria;Proteobacteria;Gammaproteobacteria;Burkholderiales;Rhodocyclaceae;Azospira;uncultured bacterium | 94.6 | 1296 | -1 |
| metaRNA.PFemirge\_19494 | 39 | 0.000005 | BAAU01002038.417.1926 | Bacteria;Firmicutes;Clostridia;Lachnospirales;Lachnospiraceae;uncultured;human gut metagenome | 91.7 | 1303 | -1 |
| metaRNA.PFemirge\_23516 | 39 | 0.000004 | CP003274.119147.120561 | Bacteria;Bacteroidota;Bacteroidia;Bacteroidales;Rikenellaceae;Alistipes;Alistipes finegoldii DSM 17242 | 92.1 | 1162 | -1 |
| metaRNA.PFemirge\_24669 | 38 | 0.000001 | FR749937.1.1523 | Bacteria;Firmicutes;Clostridia;Lachnospirales;Lachnospiraceae;[Eubacterium] fissicatena group;Faecalicatena fissicatena | 94.8 | 1335 | -1 |
| metaRNA.PFemirge\_775 | 38 | 0.000004 | CABV01003337.48680.50227 | Bacteria;Verrucomicrobiota;Verrucomicrobiae;Verrucomicrobiales;Rubritaleaceae;Luteolibacter;Oikopleura dioica | 99.3 | 1215 | -1 |
| metaRNA.PFemirge\_23694 | 38 | 0.000013 | AB088925.1.1354 | Bacteria;Bacteroidota;Bacteroidia;Bacteroidales;Williamwhitmaniaceae;Rs-D38 termite group;uncultured Bacteroidaceae bacterium | 86.6 | 1357 | -1 |
| metaRNA.PFemirge\_27894 | 38 | 0.000013 | ADLN01000129.202.1639 | Bacteria;Firmicutes;Clostridia;Lachnospirales;Lachnospiraceae;Incertae Sedis;Hungatella hathewayi WAL-18680 | 90.8 | 1321 | -1 |
| metaRNA.PFemirge\_20325 | 38 | 0.000004 | DQ795713.1.1384 | Bacteria;Firmicutes;Clostridia;Lachnospirales;Lachnospiraceae;Lachnospiraceae UCG-008;uncultured bacterium | 92.5 | 1276 | -1 |
| metaRNA.PFemirge\_3312 | 38 | 0.000004 | GU390143.1.1290 | Bacteria;Bacteroidota;Bacteroidia;Bacteroidales;Prolixibacteraceae;BSV13;uncultured bacterium | 85.3 | 1201 | -1 |
| metaRNA.PFemirge\_8952 | 38 | 0.000006 | CP011914.3962957.3964425 | Bacteria;Firmicutes;Clostridia;Eubacteriales;Eubacteriaceae;Eubacterium;Eubacterium limosum | 94.2 | 1175 | -1 |
| metaRNA.PFemirge\_25640 | 38 | 0.000004 | HQ767447.1.1440 | Bacteria;Bacteroidota;Bacteroidia;Bacteroidales;Bacteroidaceae;Bacteroides;uncultured organism | 87.9 | 1194 | -1 |
| metaRNA.PFemirge\_18121 | 38 | 0.000002 | EU773113.1.1391 | Bacteria;Bacteroidota;Bacteroidia;Bacteroidales;Rikenellaceae;Rikenellaceae RC9 gut group;uncultured bacterium | 89.7 | 1190 | -1 |
| metaRNA.PFemirge\_11749 | 38 | 0.000005 | HQ767134.1.1448 | Bacteria;Bacteroidota;Bacteroidia;Bacteroidales;Tannerellaceae;Parabacteroides;uncultured organism | 89.5 | 1330 | -1 |
| metaRNA.PFemirge\_17727 | 38 | 0.000002 | EU472520.1.1399 | Bacteria;Firmicutes;Clostridia;Lachnospirales;Lachnospiraceae;Lachnospiraceae AC2044 group;uncultured bacterium | 91.8 | 1309 | -1 |
| metaRNA.PFemirge\_19789 | 38 | 0.000002 | JN688021.1.1446 | Bacteria;Bacteroidota;Bacteroidia;Bacteroidales;Rikenellaceae;uncultured;bacterium enrichment culture clone M02 | 90.2 | 1211 | -1 |
| metaRNA.PFemirge\_23154 | 38 | 0.000006 | CP028523.1101513.1103019 | Bacteria;Firmicutes;Clostridia;Peptostreptococcales-Tissierellales;Peptostreptococcaceae;Clostridioides;Clostridioides difficile | 92.9 | 1359 | -1 |
| metaRNA.PFemirge\_13894 | 38 | 0.000003 | AB294338.1.1433 | Bacteria;Bacteroidota;Bacteroidia;Cytophagales;Cytophagaceae;Cytophaga;uncultured bacterium | 92.0 | 1258 | -1 |
| metaRNA.PFemirge\_27913 | 38 | 0.000002 | JX105679.1.1392 | Bacteria;Bacteroidota;Bacteroidia;Bacteroidales;Bacteroidaceae;Bacteroides;uncultured bacterium | 90.5 | 1355 | -1 |
| metaRNA.PFemirge\_10739 | 37 | 0.000004 | FJ393118.1.1494 | Bacteria;Firmicutes;Clostridia;Oscillospirales;Oscillospiraceae;uncultured;uncultured Clostridiales bacterium | 92.7 | 1157 | -1 |
| metaRNA.PFemirge\_21967 | 37 | 0.000007 | DQ824480.1.1373 | Bacteria;Actinobacteriota;Coriobacteriia;Coriobacteriales;Eggerthellaceae;Eggerthella;uncultured bacterium | 94.0 | 1159 | -1 |
| metaRNA.PFemirge\_23903 | 37 | 0.000006 | ADLN01000129.202.1639 | Bacteria;Firmicutes;Clostridia;Lachnospirales;Lachnospiraceae;Incertae Sedis;Hungatella hathewayi WAL-18680 | 94.2 | 1146 | -1 |
| metaRNA.PFemirge\_3326 | 37 | 0.000003 | GQ388996.1.1510 | Bacteria;Proteobacteria;Gammaproteobacteria;Pseudomonadales;Pseudomonadaceae;Pseudomonas;uncultured bacterium | 93.4 | 1401 | -1 |
| metaRNA.PFemirge\_23318 | 37 | 0.000006 | HQ783986.1.1437 | Bacteria;Bacteroidota;Bacteroidia;Bacteroidales;Tannerellaceae;Parabacteroides;uncultured organism | 89.9 | 1241 | -1 |
| metaRNA.PFemirge\_13158 | 37 | 0.000003 | GQ867583.1.1492 | Bacteria;Bacteroidota;Bacteroidia;Bacteroidales;Bacteroidaceae;Bacteroides;uncultured bacterium | 90.1 | 1167 | -1 |
| metaRNA.PFemirge\_24874 | 37 | 0.000003 | HM630210.1.1461 | Bacteria;Firmicutes;Bacilli;RF39;uncultured bacterium | 95.7 | 1281 | -1 |
| metaRNA.PFemirge\_20055 | 37 | 0.000008 | DQ807211.1.1386 | Bacteria;Firmicutes;Clostridia;Oscillospirales;Ruminococcaceae;Incertae Sedis;uncultured bacterium | 90.7 | 1286 | -1 |
| metaRNA.PFemirge\_16610 | 36 | 0.000002 | GQ480160.1.1510 | Bacteria;Firmicutes;Negativicutes;Acidaminococcales;Acidaminococcaceae;Succinispira;uncultured bacterium | 87.9 | 1196 | -1 |
| metaRNA.PFemirge\_27209 | 36 | 0.000001 | AB610575.1.1461 | Bacteria;Firmicutes;Clostridia;Peptostreptococcales-Tissierellales;Peptostreptococcaceae;Romboutsia;Clostridium sp. K39 | 98.9 | 788 | -1 |
| metaRNA.PFemirge\_10325 | 36 | 0.000006 | HQ804310.1.1461 | Bacteria;Firmicutes;Clostridia;Lachnospirales;Lachnospiraceae;Lachnospiraceae UCG-004;uncultured organism | 89.2 | 1363 | -1 |
| metaRNA.PFemirge\_1026 | 36 | 0.000005 | FPLP01007766.16.1549 | Bacteria;Myxococcota;Polyangia;Haliangiales;Haliangiaceae;Haliangium;metagenome | 94.8 | 1182 | -1 |
| metaRNA.PFemirge\_27870 | 36 | 0.000005 | FQXY01000005.366.1875 | Bacteria;Bacteroidota;Bacteroidia;Bacteroidales;Bacteroidaceae;Bacteroides;Bacteroides sp. Marseille-P3132 | 90.7 | 1485 | -1 |
| metaRNA.PFemirge\_30248 | 36 | 0.000013 | AB554233.1.1486 | Bacteria;Bacteroidota;Bacteroidia;Bacteroidales;Rikenellaceae;Alistipes;Alistipes shahii | 88.8 | 1400 | -1 |
| metaRNA.PFemirge\_2466 | 36 | 0.000003 | FR749935.1.1522 | Bacteria;Firmicutes;Clostridia;Lachnospirales;Lachnospiraceae;[Eubacterium] fissicatena group;Faecalicatena fissicatena | 92.0 | 1382 | -1 |
| metaRNA.PFemirge\_837 | 36 | 0.000003 | AM934838.1.1381 | Bacteria;Dadabacteria;Dadabacteriia;Dadabacteriales;uncultured candidate division SBR1093 bacterium | 99.9 | 1381 | -1 |
| metaRNA.PFemirge\_25223 | 36 | 0.000006 | FJ825522.1.1343 | Bacteria;Firmicutes;Clostridia;Peptostreptococcales-Tissierellales;Peptostreptococcaceae;Romboutsia;uncultured bacterium | 92.9 | 1250 | -1 |
| metaRNA.PFemirge\_24895 | 36 | 0.000008 | EF644509.1.1469 | Bacteria;Firmicutes;Clostridia;Oscillospirales;Oscillospiraceae;Colidextribacter;uncultured bacterium | 90.1 | 1269 | -1 |
| metaRNA.PFemirge\_12523 | 36 | 0.000091 | HQ819316.1.1450 | Bacteria;Bacteroidota;Bacteroidia;Bacteroidales;Bacteroidaceae;Bacteroides;uncultured organism | 90.4 | 1192 | -1 |
| metaRNA.PFemirge\_7189 | 35 | 0.000004 | EU136696.1.1944 | Bacteria;Bacteroidota;Bacteroidia;Bacteroidales;Bacteroidaceae;Bacteroides;Bacteroides massiliensis | 87.1 | 1405 | -1 |
| metaRNA.PFemirge\_16860 | 35 | 0.000009 | LARG01000025.1.1402 | Bacteria;Bacteroidota;Bacteroidia;Bacteroidales;Rikenellaceae;Alistipes;Alistipes sp. N15.MGS-157 | 92.8 | 1396 | -1 |
| metaRNA.PFemirge\_12361 | 34 | 0.000007 | FXDJ01000009.1407067.1408576 | Bacteria;Bacteroidota;Bacteroidia;Bacteroidales;Tannerellaceae;uncultured;Porphyromonadaceae bacterium MV12 | 87.7 | 1458 | -1 |
| metaRNA.PFemirge\_10705 | 34 | 0.000004 | KJ624065.29957.31422 | Bacteria;Cyanobacteria;Cyanobacteriia;Chloroplast;Trachydiscus minutus | 99.6 | 1131 | -1 |
| metaRNA.PFemirge\_21054 | 34 | 0.000002 | DQ796609.1.1391 | Bacteria;Firmicutes;Clostridia;Oscillospirales;Ruminococcaceae;Faecalibacterium;uncultured bacterium | 93.6 | 1289 | -1 |
| metaRNA.PFemirge\_22134 | 34 | 0.000006 | GQ359970.1.1475 | Bacteria;Firmicutes;Clostridia;Clostridiales;Clostridiaceae;Clostridium sensu stricto 1;uncultured bacterium | 91.4 | 1368 | -1 |
| metaRNA.PFemirge\_20262 | 34 | 0.000004 | EU778633.1.1385 | Bacteria;Firmicutes;Clostridia;Lachnospirales;Lachnospiraceae;Moryella;uncultured bacterium | 92.5 | 1242 | -1 |
| metaRNA.PFemirge\_24650 | 34 | 0.000006 | GU174132.1.1481 | Bacteria;Firmicutes;Clostridia;Lachnospirales;Lachnospiraceae;Lachnospiraceae UCG-009;uncultured bacterium | 92.1 | 1263 | -1 |
| metaRNA.PFemirge\_12328 | 34 | 0.000003 | HQ795096.1.1444 | Bacteria;Bacteroidota;Bacteroidia;Bacteroidales;Bacteroidaceae;Bacteroides;uncultured organism | 86.8 | 1413 | -1 |
| metaRNA.PFemirge\_20218 | 34 | 0.000002 | EU459185.1.1385 | Bacteria;Firmicutes;Clostridia;Lachnospirales;Lachnospiraceae;Epulopiscium;uncultured bacterium | 93.3 | 1186 | -1 |
| metaRNA.PFemirge\_11718 | 34 | 0.000005 | HQ741466.1.1426 | Bacteria;Firmicutes;Clostridia;Clostridiales;Clostridiaceae;Clostridium sensu stricto 1;uncultured bacterium | 92.5 | 1316 | -1 |
| metaRNA.PFemirge\_20441 | 34 | 0.000029 | HM630236.1.1451 | Bacteria;Bacteroidota;Bacteroidia;Bacteroidales;Rikenellaceae;dgA-11 gut group;uncultured bacterium | 89.0 | 1190 | -1 |
| metaRNA.PFemirge\_23794 | 33 | 0.000008 | JN703378.1.1490 | Bacteria;Bacteroidota;Bacteroidia;Bacteroidales;Barnesiellaceae;Coprobacter;Coprobacter fastidiosus | 86.6 | 1354 | -1 |
| metaRNA.PFemirge\_5611 | 33 | 0.000002 | CP022151.4897185.4898740 | Bacteria;Proteobacteria;Gammaproteobacteria;Enterobacterales;Enterobacteriaceae;Citrobacter;Citrobacter freundii | 94.9 | 1399 | -1 |
| metaRNA.PFemirge\_24309 | 33 | 0.000005 | EU772665.1.1358 | Bacteria;Firmicutes;Clostridia;Lachnospirales;Lachnospiraceae;Lachnospiraceae UCG-008;uncultured bacterium | 93.7 | 1225 | -1 |
| metaRNA.PFemirge\_18128 | 33 | 0.000009 | EU772234.1.1393 | Bacteria;Firmicutes;Clostridia;Christensenellales;Christensenellaceae;Christensenellaceae R-7 group;uncultured bacterium | 93.2 | 1293 | -1 |
| metaRNA.PFemirge\_5397 | 33 | 0.000004 | FPLP01013193.1.1403 | Bacteria;Cyanobacteria;Vampirivibrionia;Obscuribacterales;Obscuribacteraceae;Candidatus Obscuribacter;metagenome | 95.5 | 1192 | -1 |
| metaRNA.PFemirge\_5097 | 33 | 0.000004 | EF406661.1.1497 | Bacteria;Firmicutes;Clostridia;Lachnospirales;Lachnospiraceae;[Ruminococcus] torques group;uncultured bacterium | 94.7 | 1395 | -1 |
| metaRNA.PFemirge\_27410 | 33 | 0.000001 | KF798941.1.1503 | Bacteria;Proteobacteria;Gammaproteobacteria;Enterobacterales;Enterobacteriaceae;Citrobacter;uncultured bacterium | 93.8 | 884 | -1 |
| metaRNA.PFemirge\_15917 | 33 | 0.000003 | FJ534957.1.1501 | Bacteria;Firmicutes;Clostridia;Lachnospirales;Lachnospiraceae;Lachnospiraceae FCS020 group;uncultured bacterium | 92.7 | 1342 | -1 |
| metaRNA.PFemirge\_17303 | 33 | 0.000008 | JX105679.1.1392 | Bacteria;Bacteroidota;Bacteroidia;Bacteroidales;Bacteroidaceae;Bacteroides;uncultured bacterium | 90.9 | 1215 | -1 |
| metaRNA.PFemirge\_5798 | 32 | 0.000004 | GQ448246.1.1389 | Bacteria;Bacteroidota;Bacteroidia;Bacteroidales;Bacteroidaceae;Bacteroides;uncultured bacterium | 90.8 | 1396 | -1 |
| metaRNA.PFemirge\_27319 | 32 | 0.000003 | EF626688.1.1481 | Bacteria;Proteobacteria;Gammaproteobacteria;Xanthomonadales;Xanthomonadaceae;Luteimonas;Luteimonas aquatica | 93.1 | 1168 | -1 |
| metaRNA.PFemirge\_23431 | 32 | 0.000015 | HM630236.1.1451 | Bacteria;Bacteroidota;Bacteroidia;Bacteroidales;Rikenellaceae;dgA-11 gut group;uncultured bacterium | 87.7 | 1361 | -1 |
| metaRNA.PFemirge\_3014 | 32 | 0.000005 | JX222815.1.1491 | Bacteria;Proteobacteria;Gammaproteobacteria;Burkholderiales;Comamonadaceae;Aquabacterium;uncultured bacterium | 93.9 | 1414 | -1 |
| metaRNA.PFemirge\_18253 | 32 | 0.000499 | FJ534957.1.1501 | Bacteria;Firmicutes;Clostridia;Lachnospirales;Lachnospiraceae;Lachnospiraceae FCS020 group;uncultured bacterium | 93.3 | 1398 | -1 |
| metaRNA.PFemirge\_1445 | 32 | 0.000005 | GU303678.1.1524 | Bacteria;Firmicutes;Bacilli;RF39;uncultured rumen bacterium | 91.6 | 1177 | -1 |
| metaRNA.PFemirge\_9837 | 32 | 0.000002 | HQ789722.1.1442 | Bacteria;Bacteroidota;Bacteroidia;Bacteroidales;Bacteroidaceae;Bacteroides;uncultured organism | 94.8 | 1347 | -1 |
| metaRNA.PFemirge\_17065 | 32 | 0.000009 | EU464167.1.1397 | Bacteria;Bacteroidota;Bacteroidia;Bacteroidales;Barnesiellaceae;uncultured;uncultured bacterium | 89.1 | 1285 | -1 |
| metaRNA.PFemirge\_25314 | 32 | 0.000004 | EU511665.1.1382 | Bacteria;Firmicutes;Clostridia;Lachnospirales;Lachnospiraceae;Lachnospiraceae UCG-008;uncultured bacterium | 91.2 | 1161 | -1 |
| metaRNA.PFemirge\_22129 | 32 | 0.000002 | ADLN01000129.202.1639 | Bacteria;Firmicutes;Clostridia;Lachnospirales;Lachnospiraceae;Incertae Sedis;Hungatella hathewayi WAL-18680 | 91.0 | 1373 | -1 |
| metaRNA.PFemirge\_14306 | 32 | 0.000004 | FR749937.1.1523 | Bacteria;Firmicutes;Clostridia;Lachnospirales;Lachnospiraceae;[Eubacterium] fissicatena group;Faecalicatena fissicatena | 95.5 | 1366 | -1 |
| metaRNA.PFemirge\_16460 | 32 | 0.000006 | DQ824384.1.1400 | Bacteria;Bacteroidota;Bacteroidia;Bacteroidales;Bacteroidaceae;Bacteroides;uncultured bacterium | 88.7 | 1361 | -1 |
| metaRNA.PFemirge\_17776 | 32 | 0.000002 | CU926683.1.1355 | Bacteria;Bacteroidota;Bacteroidia;Bacteroidales;Tannerellaceae;Macellibacteroides;uncultured bacterium | 88.0 | 1209 | -1 |
| metaRNA.PFemirge\_20887 | 32 | 0.000008 | LN573922.1.1381 | Bacteria;Planctomycetota;Planctomycetes;Gemmatales;Gemmataceae;uncultured;uncultured bacterium | 97.9 | 838 | -1 |
| metaRNA.PFemirge\_28066 | 32 | 0.000004 | EU765828.1.1335 | Bacteria;Firmicutes;Clostridia;Oscillospirales;Ruminococcaceae;Faecalibacterium;uncultured bacterium | 93.7 | 1252 | -1 |
| metaRNA.PFemirge\_22148 | 32 | 0.000002 | EU655834.1.1314 | Bacteria;Firmicutes;Clostridia;Oscillospirales;Ruminococcaceae;UBA1819;uncultured bacterium | 91.5 | 1266 | -1 |
| metaRNA.PFemirge\_6700 | 31 | 0.000006 | FJ951888.1.1489 | Bacteria;Bacteroidota;Bacteroidia;Bacteroidales;Rikenellaceae;Alistipes;uncultured bacterium | 89.0 | 1488 | -1 |
| metaRNA.PFemirge\_15328 | 31 | 0.000002 | AB554230.1.1487 | Bacteria;Bacteroidota;Bacteroidia;Bacteroidales;Rikenellaceae;Alistipes;Alistipes finegoldii | 91.0 | 1245 | -1 |
| metaRNA.PFemirge\_19580 | 31 | 0.000003 | HQ761492.1.1450 | Bacteria;Firmicutes;Clostridia;Lachnospirales;Lachnospiraceae;[Ruminococcus] torques group;uncultured organism | 91.6 | 1289 | -1 |
| metaRNA.PFemirge\_5958 | 31 | 0.000005 | AGES01000014.73256.74770 | Bacteria;Bacteroidota;Bacteroidia;Bacteroidales;Tannerellaceae;Parabacteroides;Parabacteroides sp. HGS0025 | 88.7 | 1412 | -1 |
| metaRNA.PFemirge\_21900 | 31 | 0.000011 | HQ802642.1.1374 | Bacteria;Firmicutes;Clostridia;Lachnospirales;Lachnospiraceae;Epulopiscium;uncultured organism | 89.9 | 1322 | -1 |
| metaRNA.PFemirge\_5680 | 31 | 0.000003 | AGTS01000010.187.1721 | Bacteria;Proteobacteria;Gammaproteobacteria;Burkholderiales;Comamonadaceae;Acidovorax;Acidovorax sp. NO-1 | 94.6 | 1411 | -1 |
| metaRNA.PFemirge\_25985 | 31 | 0.000003 | DQ677005.1.1395 | Bacteria;Firmicutes;Clostridia;Lachnospirales;Lachnospiraceae;Lachnoclostridium;Clostridium sp. Iso-A1 | 95.8 | 1226 | -1 |
| metaRNA.PFemirge\_19798 | 30 | 0.000002 | GQ448680.1.1388 | Bacteria;Firmicutes;Clostridia;Lachnospirales;Lachnospiraceae;uncultured bacterium | 89.9 | 1303 | -1 |
| metaRNA.PFemirge\_20726 | 30 | 0.000004 | FJ269051.1.1487 | Bacteria;Bacteroidota;Bacteroidia;Bacteroidales;Bacteroidaceae;Bacteroides;iron-reducing bacterium enrichment culture clone HN11 | 90.2 | 1324 | -1 |
| metaRNA.PFemirge\_3265 | 30 | 0.000001 | AB261128.1.1492 | Bacteria;Bacteroidota;Bacteroidia;Bacteroidales;Tannerellaceae;Parabacteroides;Parabacteroides johnsonii | 86.7 | 1325 | -1 |
| metaRNA.PFemirge\_25175 | 30 | 0.000001 | KF500761.1.1329 | Bacteria;Proteobacteria;Gammaproteobacteria;Burkholderiales;uncultured;uncultured bacterium | 94.6 | 1203 | -1 |
| metaRNA.PFemirge\_22204 | 30 | 0.000004 | HQ761492.1.1450 | Bacteria;Firmicutes;Clostridia;Lachnospirales;Lachnospiraceae;[Ruminococcus] torques group;uncultured organism | 91.9 | 1322 | -1 |
| metaRNA.PFemirge\_20056 | 30 | 0.000005 | MS821754.1.1508 | Bacteria;Firmicutes;Clostridia;Lachnospirales;Lachnospiraceae;[Eubacterium] fissicatena group;Faecalicatena contorta | 90.9 | 1391 | -1 |
| metaRNA.PFemirge\_7880 | 30 | 0.000005 | CBVB010000006.254456.255954 | Bacteria;Bacteroidota;Bacteroidia;Bacteroidales;Bacteroidaceae;Bacteroides;Bacteroidaceae bacterium MS4 | 89.3 | 1332 | -1 |
| metaRNA.PFemirge\_17705 | 30 | 0.000003 | AB746664.1.1536 | Bacteria;Firmicutes;Clostridia;Lachnospirales;Lachnospiraceae;Lachnospiraceae NK4A136 group;uncultured bacterium | 90.6 | 1292 | -1 |
| metaRNA.PFemirge\_24550 | 30 | 0.000001 | ADLN01000129.202.1639 | Bacteria;Firmicutes;Clostridia;Lachnospirales;Lachnospiraceae;Incertae Sedis;Hungatella hathewayi WAL-18680 | 90.5 | 1290 | -1 |
| metaRNA.PFemirge\_6511 | 30 | 0.000007 | HQ757204.1.1439 | Bacteria;Bacteroidota;Bacteroidia;Bacteroidales;Bacteroidaceae;Bacteroides;uncultured organism | 87.5 | 1315 | -1 |
| metaRNA.PFemirge\_18796 | 30 | 0.000002 | EF400740.1.1495 | Bacteria;Firmicutes;Clostridia;Oscillospirales;Butyricicoccaceae;UCG-009;uncultured bacterium | 93.5 | 1286 | -1 |
| metaRNA.PFemirge\_21002 | 30 | 0.000004 | HQ748426.1.1449 | Bacteria;Firmicutes;Clostridia;Lachnospirales;Lachnospiraceae;Lachnoclostridium;uncultured organism | 93.0 | 1326 | -1 |
| metaRNA.PFemirge\_26488 | 30 | 0.000004 | EU772735.1.1293 | Bacteria;Firmicutes;Clostridia;Oscillospirales;Butyricicoccaceae;uncultured;uncultured bacterium | 95.4 | 1210 | -1 |
| metaRNA.PFemirge\_30254 | 30 | 0.000016 | KP944118.1.1420 | Bacteria;Bacteroidota;Bacteroidia;Bacteroidales;Bacteroidaceae;Bacteroides;Bacteroides thetaiotaomicron | 90.5 | 1150 | -1 |
| metaRNA.PFemirge\_25227 | 29 | 0.000002 | AB626942.1.1521 | Bacteria;Firmicutes;Clostridia;Lachnospirales;Lachnospiraceae;Lachnospiraceae UCG-008;uncultured bacterium | 89.9 | 1301 | -1 |
| metaRNA.PFemirge\_26536 | 29 | 0.000003 | LC028726.1.1525 | Bacteria;Firmicutes;Clostridia;Lachnospirales;Lachnospiraceae;uncultured;uncultured bacterium | 91.4 | 1175 | -1 |
| metaRNA.PFemirge\_6254 | 29 | 0.000005 | AY321958.1.1429 | Bacteria;Actinobacteriota;Coriobacteriia;Coriobacteriales;Eggerthellaceae;Eggerthella;Eggerthella sinensis | 93.8 | 1176 | -1 |
| metaRNA.PFemirge\_26764 | 28 | 0.000004 | HQ756942.1.1450 | Bacteria;Firmicutes;Clostridia;Lachnospirales;Lachnospiraceae;[Ruminococcus] gnavus group;uncultured organism | 89.3 | 1275 | -1 |
| metaRNA.PFemirge\_25825 | 28 | 0.000004 | DQ808514.1.1304 | Bacteria;Bacteroidota;Bacteroidia;Bacteroidales;Bacteroidaceae;Bacteroides;uncultured bacterium | 90.9 | 1104 | -1 |
| metaRNA.PFemirge\_16270 | 28 | 0.000003 | FJ957875.1.1447 | Bacteria;Firmicutes;Clostridia;Lachnospirales;Lachnospiraceae;Lachnoclostridium;[Clostridium] saccharolyticum | 93.4 | 1367 | -1 |
| metaRNA.PFemirge\_23234 | 28 | 0.000001 | HQ778533.1.1446 | Bacteria;Bacteroidota;Bacteroidia;Bacteroidales;Tannerellaceae;Parabacteroides;uncultured organism | 93.1 | 1311 | -1 |
| metaRNA.PFemirge\_12504 | 28 | 0.000006 | KM650913.1.1442 | Bacteria;Bacteroidota;Bacteroidia;Bacteroidales;Bacteroidaceae;Bacteroides;uncultured Bacteroidetes bacterium | 91.5 | 1421 | -1 |
| metaRNA.PFemirge\_17240 | 28 | 0.000004 | KF318232.1.1402 | Bacteria;Bacteroidota;Bacteroidia;Bacteroidales;Bacteroidaceae;Bacteroides;uncultured Bacteroidetes bacterium | 88.8 | 1246 | -1 |
| metaRNA.PFemirge\_20401 | 28 | 0.000002 | EU775304.1.1384 | Bacteria;Firmicutes;Clostridia;Lachnospirales;Lachnospiraceae;Lachnospiraceae UCG-008;uncultured bacterium | 91.1 | 1301 | -1 |
| metaRNA.PFemirge\_21457 | 28 | 0.000003 | EU475534.1.1401 | Bacteria;Firmicutes;Clostridia;Lachnospirales;Lachnospiraceae;Lachnospiraceae AC2044 group;uncultured bacterium | 90.1 | 1289 | -1 |
| metaRNA.PFemirge\_24146 | 28 | 0.000003 | DQ801177.1.1348 | Bacteria;Bacteroidota;Bacteroidia;Bacteroidales;Bacteroidaceae;Bacteroides;uncultured bacterium | 90.8 | 1310 | -1 |
| metaRNA.PFemirge\_28453 | 28 | 0.000003 | FJ366372.1.1363 | Bacteria;Bacteroidota;Bacteroidia;Bacteroidales;Rikenellaceae;Alistipes;uncultured bacterium | 89.2 | 1365 | -1 |
| metaRNA.PFemirge\_26869 | 28 | 0.000003 | LT598589.1.1424 | Bacteria;Firmicutes;Clostridia;Oscillospirales;Butyricicoccaceae;UCG-008;Intestinibacillus massiliensis | 92.4 | 1250 | -1 |
| metaRNA.PFemirge\_9528 | 28 | 0.000003 | HQ317401.1.1485 | Bacteria;Cyanobacteria;Cyanobacteriia;Chloroplast;Haematococcus lacustris | 99.7 | 1018 | -1 |
| metaRNA.PFemirge\_10921 | 27 | 0.000003 | KC961269.1.1495 | Bacteria;Proteobacteria;Gammaproteobacteria;Burkholderiales;Comamonadaceae;Hydrogenophaga;uncultured bacterium | 94.5 | 1367 | -1 |
| metaRNA.PFemirge\_8590 | 27 | 0.000001 | AB470343.1.1485 | Bacteria;Bacteroidota;Bacteroidia;Bacteroidales;Tannerellaceae;Parabacteroides;Parabacteroides gordonii | 92.5 | 1415 | -1 |
| metaRNA.PFemirge\_21495 | 27 | 0.000001 | FJ534957.1.1501 | Bacteria;Firmicutes;Clostridia;Lachnospirales;Lachnospiraceae;Lachnospiraceae FCS020 group;uncultured bacterium | 91.7 | 1301 | -1 |
| metaRNA.PFemirge\_6540 | 27 | 0.000006 | FXDJ01000009.1407067.1408576 | Bacteria;Bacteroidota;Bacteroidia;Bacteroidales;Tannerellaceae;uncultured;Porphyromonadaceae bacterium MV12 | 89.3 | 1443 | -1 |
| metaRNA.PFemirge\_22979 | 27 | 0.000007 | CP016102.16052.17542 | Bacteria;Firmicutes;Clostridia;Peptostreptococcales-Tissierellales;Peptostreptococcaceae;Clostridioides;Clostridioides difficile | 93.7 | 1357 | -1 |
| metaRNA.PFemirge\_7721 | 27 | 0.000014 | CYSK01000016.265066.266548 | Bacteria;Firmicutes;Clostridia;Clostridiales;Clostridiaceae;Clostridium sensu stricto 8;Clostridiaceae bacterium mt10 | 95.8 | 1223 | -1 |
| metaRNA.PFemirge\_5789 | 26 | 0.000006 | CP009628.2620010.2621547 | Bacteria;Firmicutes;Bacilli;Bacillales;Bacillaceae;Bacillus;Bacillus cereus ATCC 4342 | 89.5 | 1320 | -1 |
| metaRNA.PFemirge\_12736 | 26 | 0.000007 | HQ779103.1.1440 | Bacteria;Firmicutes;Clostridia;Lachnospirales;Lachnospiraceae;Lachnospiraceae NC2004 group;uncultured organism | 93.1 | 1369 | -1 |
| metaRNA.PFemirge\_12525 | 26 | 0.000002 | CP003274.2181216.2182629 | Bacteria;Bacteroidota;Bacteroidia;Bacteroidales;Rikenellaceae;Alistipes;Alistipes finegoldii DSM 17242 | 94.0 | 1226 | -1 |
| metaRNA.PFemirge\_27621 | 26 | 0.000005 | FJ477666.1.1463 | Bacteria;Proteobacteria;Gammaproteobacteria;Enterobacterales;Enterobacteriaceae;Escherichia-Shigella;uncultured Pantoea sp. | 96.0 | 849 | -1 |
| metaRNA.PFemirge\_19442 | 26 | 0.000006 | HQ681763.1.1459 | Bacteria;Firmicutes;Clostridia;Oscillospirales;Ruminococcaceae;Ruminococcus;uncultured bacterium | 92.7 | 1294 | -1 |
| metaRNA.PFemirge\_21816 | 26 | 0.000002 | EU775041.1.1395 | Bacteria;Bacteroidota;Bacteroidia;Bacteroidales;Bacteroidaceae;Bacteroides;uncultured bacterium | 89.2 | 1374 | -1 |
| metaRNA.PFemirge\_16359 | 25 | 0.000008 | DQ807222.1.1401 | Bacteria;Bacteroidota;Bacteroidia;Bacteroidales;Bacteroidaceae;Bacteroides;uncultured bacterium | 90.9 | 1257 | -1 |
| metaRNA.PFemirge\_6116 | 25 | 0.000006 | GQ871713.1.1490 | Bacteria;Bacteroidota;Bacteroidia;Bacteroidales;Rikenellaceae;Alistipes;uncultured bacterium | 87.0 | 1433 | -1 |
| metaRNA.PFemirge\_19201 | 25 | 0.000002 | DQ800695.1.1386 | Bacteria;Firmicutes;Clostridia;Lachnospirales;Lachnospiraceae;Lachnoclostridium;uncultured bacterium | 90.2 | 1392 | -1 |
| metaRNA.PFemirge\_26124 | 25 | 0.000002 | EU775698.1.1374 | Bacteria;Firmicutes;Clostridia;Clostridiales;Clostridiaceae;Clostridium sensu stricto 1;uncultured bacterium | 95.3 | 1287 | -1 |
| metaRNA.PFemirge\_10106 | 25 | 0.000002 | FR749935.1.1522 | Bacteria;Firmicutes;Clostridia;Lachnospirales;Lachnospiraceae;[Eubacterium] fissicatena group;Faecalicatena fissicatena | 93.1 | 1378 | -1 |
| metaRNA.PFemirge\_26291 | 25 | 0.000003 | EU775302.1.1379 | Bacteria;Firmicutes;Clostridia;Lachnospirales;Lachnospiraceae;Lachnospiraceae AC2044 group;uncultured bacterium | 92.8 | 1215 | -1 |
| metaRNA.PFemirge\_24538 | 25 | 0.000003 | DQ803378.1.1392 | Bacteria;Firmicutes;Clostridia;Lachnospirales;Lachnospiraceae;uncultured;uncultured bacterium | 90.5 | 1347 | -1 |
| metaRNA.PFemirge\_3621 | 25 | 0.000002 | CDZS01002192.2025.3531 | Bacteria;Bacteroidota;Bacteroidia;Bacteroidales;Bacteroidaceae;Bacteroides;gut metagenome | 88.9 | 1318 | -1 |
| metaRNA.PFemirge\_15283 | 25 | 0.000001 | LT725663.1.1450 | Bacteria;Bacteroidota;Bacteroidia;Bacteroidales;Tannerellaceae;Parabacteroides;Parabacteroides sp. Marseille-P3668 | 84.9 | 1440 | -1 |
| metaRNA.PFemirge\_27926 | 25 | 0.000006 | AGES01000014.73256.74770 | Bacteria;Bacteroidota;Bacteroidia;Bacteroidales;Tannerellaceae;Parabacteroides;Parabacteroides sp. HGS0025 | 88.4 | 1340 | -1 |
| metaRNA.PFemirge\_21456 | 25 | 0.000004 | EU771988.1.1377 | Bacteria;Firmicutes;Clostridia;Oscillospirales;Ruminococcaceae;Fournierella;uncultured bacterium | 92.2 | 1365 | -1 |
| metaRNA.PFemirge\_11197 | 25 | 0.000002 | HQ741370.1.1454 | Bacteria;Firmicutes;Clostridia;Lachnospirales;Lachnospiraceae;Epulopiscium;uncultured bacterium | 91.5 | 1310 | -1 |
| metaRNA.PFemirge\_17871 | 24 | 0.000036 | GQ867454.1.1492 | Bacteria;Firmicutes;Clostridia;Oscillospirales;Ruminococcaceae;Fournierella;uncultured bacterium | 94.6 | 1305 | -1 |
| metaRNA.PFemirge\_26180 | 24 | 0.000003 | EU655834.1.1314 | Bacteria;Firmicutes;Clostridia;Oscillospirales;Ruminococcaceae;UBA1819;uncultured bacterium | 92.9 | 1228 | -1 |
| metaRNA.PFemirge\_16589 | 24 | 0.000005 | CU926683.1.1355 | Bacteria;Bacteroidota;Bacteroidia;Bacteroidales;Tannerellaceae;Macellibacteroides;uncultured bacterium | 90.5 | 1168 | -1 |
| metaRNA.PFemirge\_27994 | 24 | 0.000003 | DQ824289.1.1398 | Bacteria;Bacteroidota;Bacteroidia;Bacteroidales;Bacteroidaceae;Bacteroides;uncultured bacterium | 91.1 | 1394 | -1 |
| metaRNA.PFemirge\_9797 | 24 | 0.000003 | CP003153.1901384.1902805 | Bacteria;Proteobacteria;Gammaproteobacteria;Burkholderiales;Rhodocyclaceae;Azospira;Azospira oryzae PS | 93.9 | 1221 | -1 |
| metaRNA.PFemirge\_17783 | 24 | 0.000002 | ADLN01000129.202.1639 | Bacteria;Firmicutes;Clostridia;Lachnospirales;Lachnospiraceae;Incertae Sedis;Hungatella hathewayi WAL-18680 | 92.9 | 1211 | -1 |
| metaRNA.PFemirge\_25672 | 24 | 0.000001 | GEEG01002153.1.1395 | Bacteria;Bacteroidota;Bacteroidia;Bacteroidales;Tannerellaceae;Parabacteroides;Monocercomonoides sp. PA203 | 91.7 | 1241 | -1 |
| metaRNA.PFemirge\_27752 | 24 | 0.000024 | FQSE01000007.1631802.1633308 | Bacteria;Bacteroidota;Bacteroidia;Bacteroidales;Tannerellaceae;Parabacteroides;Parabacteroides sp. Marseille-P3136 | 91.6 | 1403 | -1 |
| metaRNA.PFemirge\_26399 | 24 | 0.000003 | GQ448690.1.1389 | Bacteria;Firmicutes;Clostridia;Lachnospirales;Lachnospiraceae;[Eubacterium] oxidoreducens group;uncultured bacterium | 91.6 | 1176 | -1 |
| metaRNA.PFemirge\_8137 | 24 | 0.000002 | KC682971.1.1497 | Bacteria;Planctomycetota;Planctomycetes;Planctomycetales;Rubinisphaeraceae;SH-PL14;uncultured bacterium | 99.8 | 1345 | -1 |
| metaRNA.PFemirge\_22189 | 24 | 0.000024 | DQ806157.1.1383 | Bacteria;Firmicutes;Clostridia;Oscillospirales;Ruminococcaceae;Faecalibacterium;uncultured bacterium | 91.3 | 1282 | -1 |
| metaRNA.PFemirge\_11040 | 23 | 0.000003 | FN563303.1.1455 | Bacteria;Firmicutes;Clostridia;Lachnospirales;Lachnospiraceae;Eisenbergiella;uncultured bacterium | 92.3 | 1254 | -1 |
| metaRNA.PFemirge\_27051 | 23 | 0.000002 | EU777807.1.1400 | Bacteria;Proteobacteria;Gammaproteobacteria;Enterobacterales;Enterobacteriaceae;Escherichia-Shigella;uncultured bacterium | 96.4 | 1125 | -1 |
| metaRNA.PFemirge\_25380 | 23 | 0.000002 | EU778073.1.1322 | Bacteria;Bacteroidota;Bacteroidia;Bacteroidales;Rikenellaceae;Alistipes;uncultured bacterium | 93.1 | 1249 | -1 |
| metaRNA.PFemirge\_19051 | 23 | 0.000004 | FR749935.1.1522 | Bacteria;Firmicutes;Clostridia;Lachnospirales;Lachnospiraceae;[Eubacterium] fissicatena group;Faecalicatena fissicatena | 94.6 | 1388 | -1 |
| metaRNA.PFemirge\_25880 | 23 | 0.000002 | EU772065.1.1302 | Bacteria;Bacteroidota;Bacteroidia;Bacteroidales;Prevotellaceae;Prevotellaceae UCG-004;uncultured bacterium | 87.2 | 1297 | -1 |
| metaRNA.PFemirge\_19034 | 23 | 0.000004 | HQ802049.1.1451 | Bacteria;Firmicutes;Clostridia;Lachnospirales;Lachnospiraceae;uncultured;uncultured organism | 91.0 | 1157 | -1 |
| metaRNA.PFemirge\_12520 | 22 | 0.000004 | ASTA01000042.17380.18894 | Bacteria;Firmicutes;Clostridia;Oscillospirales;Ruminococcaceae;Anaerotruncus;Anaerotruncus sp. G3(2012) | 91.2 | 1321 | -1 |
| metaRNA.PFemirge\_26585 | 22 | 0.000003 | AATF01000761.367.1874 | Bacteria;Firmicutes;Clostridia;Oscillospirales;Ruminococcaceae;Ruminococcus;mouse gut metagenome | 90.9 | 1274 | -1 |
| metaRNA.PFemirge\_3185 | 22 | 0.000003 | FJ184323.1.1507 | Bacteria;Proteobacteria;Gammaproteobacteria;Xanthomonadales;Xanthomonadaceae;Stenotrophomonas;uncultured soil bacterium | 96.3 | 1485 | -1 |
| metaRNA.PFemirge\_7284 | 22 | 0.000002 | KF773008.1.1446 | Bacteria;Firmicutes;Clostridia;Lachnospirales;Lachnospiraceae;Hungatella;uncultured bacterium | 93.2 | 1392 | -1 |
| metaRNA.PFemirge\_26408 | 22 | 0.000001 | DQ793376.1.1284 | Bacteria;Firmicutes;Clostridia;Oscillospirales;Butyricicoccaceae;Butyricicoccus;uncultured bacterium | 92.4 | 1285 | -1 |
| metaRNA.PFemirge\_26157 | 22 | 0.000005 | HQ815341.1.1435 | Bacteria;Firmicutes;Clostridia;Peptostreptococcales-Tissierellales;Peptostreptococcaceae;Intestinibacter;uncultured organism | 92.9 | 1285 | -1 |
| metaRNA.PFemirge\_11478 | 21 | 0.000002 | LT725663.1.1450 | Bacteria;Bacteroidota;Bacteroidia;Bacteroidales;Tannerellaceae;Parabacteroides;Parabacteroides sp. Marseille-P3668 | 89.3 | 1246 | -1 |
| metaRNA.PFemirge\_27324 | 21 | 0.000004 | KF697581.1.1227 | Bacteria;Nitrospirota;Nitrospiria;Nitrospirales;Nitrospiraceae;Nitrospira;uncultured bacterium | 93.1 | 1020 | -1 |
| metaRNA.PFemirge\_22720 | 21 | 0.000001 | DQ798128.1.1365 | Bacteria;Firmicutes;Clostridia;Oscillospirales;Ruminococcaceae;Ruminococcus;uncultured bacterium | 90.9 | 1172 | -1 |
| metaRNA.PFemirge\_13515 | 21 | 0.000014 | AB649276.1.1495 | Bacteria;Firmicutes;Clostridia;Lachnospirales;Lachnospiraceae;Anaerostignum;Anaerotignum propionicum | 89.2 | 1207 | -1 |
| metaRNA.PFemirge\_19506 | 21 | 0.000005 | EU475815.1.1389 | Bacteria;Firmicutes;Clostridia;Lachnospirales;Lachnospiraceae;Lachnospiraceae UCG-008;uncultured bacterium | 94.0 | 1381 | -1 |
| metaRNA.PFemirge\_16322 | 21 | 0.000001 | DQ799369.1.1397 | Bacteria;Bacteroidota;Bacteroidia;Bacteroidales;Bacteroidaceae;Bacteroides;uncultured bacterium | 87.0 | 1405 | -1 |
| metaRNA.PFemirge\_16163 | 21 | 0.000017 | DQ799557.1.1402 | Bacteria;Bacteroidota;Bacteroidia;Bacteroidales;Bacteroidaceae;Bacteroides;uncultured bacterium | 92.6 | 1398 | -1 |
| metaRNA.PFemirge\_19857 | 21 | 0.000002 | EU778864.1.1377 | Bacteria;Firmicutes;Clostridia;Lachnospirales;Lachnospiraceae;Lachnospiraceae UCG-008;uncultured bacterium | 91.5 | 1382 | -1 |
| metaRNA.PFemirge\_19766 | 20 | 0.000003 | DQ802739.1.1400 | Bacteria;Firmicutes;Clostridia;Oscillospirales;Ruminococcaceae;Incertae Sedis;uncultured bacterium | 92.9 | 1385 | -1 |
| metaRNA.PFemirge\_26540 | 20 | 0.000005 | FR749935.1.1522 | Bacteria;Firmicutes;Clostridia;Lachnospirales;Lachnospiraceae;[Eubacterium] fissicatena group;Faecalicatena fissicatena | 92.8 | 1284 | -1 |
| metaRNA.PFemirge\_23960 | 20 | 0.000001 | HQ781017.1.1452 | Bacteria;Bacteroidota;Bacteroidia;Bacteroidales;Bacteroidaceae;Bacteroides;uncultured organism | 93.3 | 1140 | -1 |
| metaRNA.PFemirge\_23470 | 19 | 0.000003 | GQ135998.1.1368 | Bacteria;Firmicutes;Clostridia;Peptostreptococcales-Tissierellales;Anaerovoracaceae;uncultured;uncultured bacterium | 92.4 | 982 | -1 |
| metaRNA.PFemirge\_26597 | 19 | 0.000001 | DQ795751.1.1277 | Bacteria;Firmicutes;Clostridia;Oscillospirales;Butyricicoccaceae;Butyricicoccus;uncultured bacterium | 94.7 | 1274 | -1 |
| metaRNA.PFemirge\_10289 | 19 | 0.000002 | CP026235.4663857.4665408 | Bacteria;Proteobacteria;Gammaproteobacteria;Enterobacterales;Enterobacteriaceae;Citrobacter;Citrobacter freundii complex sp. CFNIH3 | 94.9 | 1369 | -1 |
| metaRNA.PFemirge\_27707 | 19 | 0.000010 | HQ764819.1.1444 | Bacteria;Bacteroidota;Bacteroidia;Bacteroidales;Bacteroidaceae;Bacteroides;uncultured organism | 92.3 | 1349 | -1 |
| metaRNA.PFemirge\_20677 | 18 | 0.000001 | EU794122.1.1443 | Bacteria;Firmicutes;Clostridia;Oscillospirales;Ruminococcaceae;uncultured;uncultured Ruminococcaceae bacterium | 92.8 | 1289 | -1 |
| metaRNA.PFemirge\_7277 | 18 | 0.000003 | DQ677005.1.1395 | Bacteria;Firmicutes;Clostridia;Lachnospirales;Lachnospiraceae;Lachnoclostridium;Clostridium sp. Iso-A1 | 92.6 | 1307 | -1 |
| metaRNA.PFemirge\_2446 | 18 | 0.000014 | EU158190.1.1510 | Bacteria;Firmicutes;Clostridia;Oscillospirales;Hydrogenoanaerobacterium;Hydrogenoanaerobacterium saccharovorans | 90.6 | 1376 | -1 |
| metaRNA.PFemirge\_23962 | 18 | 0.000004 | AB447718.1.1450 | Bacteria;Firmicutes;Clostridia;Oscillospirales;Oscillospiraceae;Colidextribacter;uncultured bacterium | 93.4 | 1156 | -1 |
| metaRNA.PFemirge\_26618 | 18 | 0.000000 | EU777679.1.1373 | Bacteria;Firmicutes;Clostridia;Clostridiales;Clostridiaceae;Clostridium sensu stricto 1;uncultured bacterium | 93.5 | 1257 | -1 |
| metaRNA.PFemirge\_19175 | 18 | 0.000002 | EU454993.1.1404 | Bacteria;Firmicutes;Clostridia;Oscillospirales;Oscillospiraceae;uncultured;uncultured bacterium | 91.2 | 1389 | -1 |
| metaRNA.PFemirge\_4290 | 18 | 0.000046 | CP029554.2592470.2594015 | Bacteria;Proteobacteria;Gammaproteobacteria;Burkholderiales;Chromobacteriaceae;Chromobacterium;Chromobacterium sp. IIBBL 274-1 | 93.0 | 1394 | -1 |
| metaRNA.PFemirge\_12919 | 18 | 0.000002 | JN012610.1.1443 | Bacteria;Firmicutes;Clostridia;Oscillospirales;Ruminococcaceae;Anaerotruncus;uncultured bacterium | 90.6 | 1241 | -1 |
| metaRNA.PFemirge\_25548 | 18 | 0.000004 | DQ809036.1.1400 | Bacteria;Bacteroidota;Bacteroidia;Bacteroidales;Bacteroidaceae;Bacteroides;uncultured bacterium | 88.0 | 1204 | -1 |
| metaRNA.PFemirge\_7187 | 18 | 0.000001 | AGES01000014.73256.74770 | Bacteria;Bacteroidota;Bacteroidia;Bacteroidales;Tannerellaceae;Parabacteroides;Parabacteroides sp. HGS0025 | 93.8 | 1491 | -1 |
| metaRNA.PFemirge\_6423 | 17 | 0.000004 | JX198651.1.1513 | Bacteria;Firmicutes;Clostridia;Oscillospirales;Butyricicoccaceae;UCG-009;uncultured bacterium | 90.4 | 1349 | -1 |
| metaRNA.PFemirge\_17678 | 17 | 0.000001 | FR749936.1.1524 | Bacteria;Firmicutes;Clostridia;Lachnospirales;Lachnospiraceae;[Eubacterium] fissicatena group;Faecalicatena fissicatena | 94.8 | 1375 | -1 |
| metaRNA.PFemirge\_12177 | 17 | 0.000004 | AF371590.1.1437 | Bacteria;Firmicutes;Clostridia;Lachnospirales;Lachnospiraceae;uncultured;uncultured bacterium | 89.5 | 1370 | -1 |
| metaRNA.PFemirge\_26073 | 17 | 0.000001 | EU771988.1.1377 | Bacteria;Firmicutes;Clostridia;Oscillospirales;Ruminococcaceae;Fournierella;uncultured bacterium | 91.0 | 1285 | -1 |
| metaRNA.PFemirge\_2588 | 17 | 0.000002 | AY126453.1.1511 | Bacteria;Proteobacteria;Gammaproteobacteria;Burkholderiales;Rhodocyclaceae;Azospira;Azospira sp. PCC | 92.6 | 1339 | -1 |
| metaRNA.PFemirge\_27950 | 17 | 0.000002 | HF558386.1.1480 | Bacteria;Firmicutes;Clostridia;Lachnospirales;Lachnospiraceae;Lachnospiraceae NC2004 group;Bacteroides xylanolyticus | 90.3 | 1391 | -1 |
| metaRNA.PFemirge\_5673 | 17 | 0.000025 | LGUD01000026.3673.5196 | Bacteria;Proteobacteria;Gammaproteobacteria;Burkholderiales;Rhodocyclaceae;Dechlorosoma;Rhodocyclaceae bacterium Paddy-1 | 93.0 | 1422 | -1 |
| metaRNA.PFemirge\_24892 | 17 | 0.000001 | AB547650.1.1476 | Bacteria;Bacteroidota;Bacteroidia;Bacteroidales;Tannerellaceae;Parabacteroides;Parabacteroides goldsteinii | 93.5 | 963 | -1 |
| metaRNA.PFemirge\_19967 | 16 | 0.000019 | HQ739338.1.1431 | Bacteria;Bacteroidota;Bacteroidia;Bacteroidales;Bacteroidaceae;Bacteroides;uncultured bacterium | 92.4 | 1242 | -1 |
| metaRNA.PFemirge\_22783 | 16 | 0.000004 | JN653038.1.1443 | Bacteria;Bacteroidota;Bacteroidia;Bacteroidales;Dysgonomonadaceae;uncultured;uncultured Porphyromonadaceae bacterium | 87.5 | 1299 | -1 |
| metaRNA.PFemirge\_6359 | 16 | 0.000004 | JX222815.1.1491 | Bacteria;Proteobacteria;Gammaproteobacteria;Burkholderiales;Comamonadaceae;Aquabacterium;uncultured bacterium | 97.7 | 1284 | -1 |
| metaRNA.PFemirge\_26451 | 16 | 0.000003 | JQ608060.1.1417 | Bacteria;Firmicutes;Clostridia;Oscillospirales;Butyricicoccaceae;UCG-009;bacterium NLAE-zl-C349 | 92.6 | 1160 | -1 |
| metaRNA.PFemirge\_11376 | 16 | 0.000004 | FR749937.1.1523 | Bacteria;Firmicutes;Clostridia;Lachnospirales;Lachnospiraceae;[Eubacterium] fissicatena group;Faecalicatena fissicatena | 92.8 | 1452 | -1 |
| metaRNA.PFemirge\_20801 | 16 | 0.000002 | LC028724.1.1522 | Bacteria;Firmicutes;Clostridia;Lachnospirales;Lachnospiraceae;Lachnospiraceae FCS020 group;uncultured bacterium | 90.1 | 1306 | -1 |
| metaRNA.PFemirge\_10872 | 16 | 0.000001 | EU772882.1.1401 | Bacteria;Firmicutes;Clostridia;Oscillospirales;Butyricicoccaceae;uncultured;uncultured bacterium | 92.6 | 1287 | -1 |
| metaRNA.PFemirge\_16440 | 16 | 0.000003 | CP026332.126495.128022 | Bacteria;Proteobacteria;Gammaproteobacteria;Pseudomonadales;Pseudomonadaceae;Pseudomonas;Pseudomonas sp. XWY-1 | 96.1 | 1397 | -1 |
| metaRNA.PFemirge\_2521 | 16 | 0.000002 | EF679189.1.1505 | Bacteria;Proteobacteria;Gammaproteobacteria;Enterobacterales;Aeromonadaceae;Aeromonas;uncultured Aeromonas sp. | 96.1 | 1310 | -1 |
| metaRNA.PFemirge\_3806 | 16 | 0.000001 | GEEG01002153.1.1395 | Bacteria;Bacteroidota;Bacteroidia;Bacteroidales;Tannerellaceae;Parabacteroides;Monocercomonoides sp. PA203 | 89.3 | 1259 | -1 |
| metaRNA.PFemirge\_18904 | 16 | 0.000005 | EU775302.1.1379 | Bacteria;Firmicutes;Clostridia;Lachnospirales;Lachnospiraceae;Lachnospiraceae AC2044 group;uncultured bacterium | 92.4 | 1243 | -1 |
| metaRNA.PFemirge\_28095 | 15 | 0.000008 | LT223609.1.1490 | Bacteria;Bacteroidota;Bacteroidia;Bacteroidales;Tannerellaceae;Parabacteroides;Parabacteroides faecis | 91.9 | 1405 | -1 |
| metaRNA.PFemirge\_20738 | 15 | 0.000001 | AB574480.1.1486 | Bacteria;Bacteroidota;Bacteroidia;Bacteroidales;Bacteroidaceae;Bacteroides;Bacteroides faecichinchillae | 90.1 | 1311 | -1 |
| metaRNA.PFemirge\_4839 | 15 | 0.000003 | CP003153.1901384.1902805 | Bacteria;Proteobacteria;Gammaproteobacteria;Burkholderiales;Rhodocyclaceae;Azospira;Azospira oryzae PS | 95.4 | 1242 | -1 |
| metaRNA.PFemirge\_5887 | 15 | 0.000002 | KM360180.1.1492 | Bacteria;Firmicutes;Clostridia;Lachnospirales;Lachnospiraceae;Lachnoclostridium;Clostridium sp. BPY5 | 90.0 | 1405 | -1 |
| metaRNA.PFemirge\_24437 | 15 | 0.000003 | HQ777443.1.1457 | Bacteria;Bacteroidota;Bacteroidia;Bacteroidales;Bacteroidaceae;Bacteroides;uncultured organism | 90.4 | 1294 | -1 |
| metaRNA.PFemirge\_18649 | 15 | 0.000001 | DQ804757.1.1391 | Bacteria;Firmicutes;Clostridia;Lachnospirales;Lachnospiraceae;[Ruminococcus] torques group;uncultured bacterium | 89.9 | 1388 | -1 |
| metaRNA.PFemirge\_20037 | 14 | 0.000002 | EU778864.1.1377 | Bacteria;Firmicutes;Clostridia;Lachnospirales;Lachnospiraceae;Lachnospiraceae UCG-008;uncultured bacterium | 92.7 | 1382 | -1 |
| metaRNA.PFemirge\_10448 | 14 | 0.000001 | FJ374184.1.1494 | Bacteria;Bacteroidota;Bacteroidia;Bacteroidales;Dysgonomonadaceae;uncultured;uncultured bacterium | 89.5 | 1264 | -1 |
| metaRNA.PFemirge\_24291 | 14 | 0.000002 | AB261128.1.1492 | Bacteria;Bacteroidota;Bacteroidia;Bacteroidales;Tannerellaceae;Parabacteroides;Parabacteroides johnsonii | 90.0 | 944 | -1 |
| metaRNA.PFemirge\_12400 | 14 | 0.000001 | AM930333.1.1446 | Bacteria;Halanaerobiaeota;Halanaerobiia;Halanaerobiales;Halanaerobiaceae;Halocella;uncultured bacterium | 96.8 | 1359 | -1 |
| metaRNA.PFemirge\_5919 | 14 | 0.000002 | KJ807937.1.1493 | Bacteria;Proteobacteria;Gammaproteobacteria;Burkholderiales;Comamonadaceae;uncultured;uncultured bacterium | 92.3 | 1385 | -1 |
| metaRNA.PFemirge\_10002 | 14 | 0.000002 | HQ761612.1.1444 | Bacteria;Bacteroidota;Bacteroidia;Bacteroidales;Tannerellaceae;Parabacteroides;uncultured organism | 88.6 | 1379 | -1 |
| metaRNA.PFemirge\_18633 | 14 | 0.000002 | DQ799802.1.1391 | Bacteria;Firmicutes;Clostridia;Lachnospirales;Lachnospiraceae;Lachnospiraceae UCG-008;uncultured bacterium | 92.3 | 1286 | -1 |
| metaRNA.PFemirge\_7319 | 14 | 0.000002 | JX222921.1.1476 | Bacteria;Firmicutes;Clostridia;Peptostreptococcales-Tissierellales;Anaerovoracaceae;Anaerovorax;uncultured bacterium | 94.1 | 1281 | -1 |
| metaRNA.PFemirge\_17932 | 13 | 0.000003 | HQ761492.1.1450 | Bacteria;Firmicutes;Clostridia;Lachnospirales;Lachnospiraceae;[Ruminococcus] torques group;uncultured organism | 94.8 | 1296 | -1 |
| metaRNA.PFemirge\_11870 | 13 | 0.000005 | EU775041.1.1395 | Bacteria;Bacteroidota;Bacteroidia;Bacteroidales;Bacteroidaceae;Bacteroides;uncultured bacterium | 91.4 | 1337 | -1 |
| metaRNA.PFemirge\_15465 | 13 | 0.000001 | AF371590.1.1437 | Bacteria;Firmicutes;Clostridia;Lachnospirales;Lachnospiraceae;uncultured;uncultured bacterium | 91.3 | 1220 | -1 |
| metaRNA.PFemirge\_4081 | 13 | 0.000002 | KF135448.1.1499 | Bacteria;Proteobacteria;Gammaproteobacteria;Pseudomonadales;Pseudomonadaceae;Pseudomonas;Pseudomonas sp. J1-5-3 | 93.8 | 1496 | -1 |
| metaRNA.PFemirge\_12416 | 13 | 0.000004 | JRGF01000044.3.1537 | Bacteria;Bacteroidota;Bacteroidia;Bacteroidales;Rikenellaceae;Alistipes;Alistipes inops | 91.7 | 1388 | -1 |
| metaRNA.PFemirge\_18132 | 13 | 0.000001 | EU454507.1.1407 | Bacteria;Firmicutes;Clostridia;Oscillospirales;Butyricicoccaceae;UCG-009;uncultured bacterium | 90.0 | 1215 | -1 |
| metaRNA.PFemirge\_20640 | 13 | 0.000002 | EU466072.1.1394 | Bacteria;Firmicutes;Clostridia;Christensenellales;Christensenellaceae;Christensenellaceae R-7 group;uncultured bacterium | 91.9 | 1300 | -1 |
| metaRNA.PFemirge\_24081 | 12 | 0.000001 | KT952737.1.1475 | Bacteria;Firmicutes;Clostridia;Oscillospirales;Ruminococcaceae;Pygmaiobacter;uncultured bacterium | 89.5 | 1278 | -1 |
| metaRNA.PFemirge\_26285 | 12 | 0.000002 | EU772528.1.1373 | Bacteria;Firmicutes;Clostridia;Lachnospirales;Lachnospiraceae;Lachnospiraceae NC2004 group;uncultured bacterium | 93.2 | 1175 | -1 |
| metaRNA.PFemirge\_26517 | 11 | 0.000001 | EU775320.1.1281 | Bacteria;Firmicutes;Clostridia;Oscillospirales;Ruminococcaceae;Ruminococcus;uncultured bacterium | 94.3 | 1273 | -1 |
| metaRNA.PFemirge\_15075 | 11 | 0.000001 | FM204998.1.1455 | Bacteria;Firmicutes;Clostridia;Lachnospirales;Lachnospiraceae;Lachnospiraceae NC2004 group;uncultured Clostridium sp. | 92.3 | 1200 | -1 |
| metaRNA.PFemirge\_16100 | 11 | 0.000002 | HQ763114.1.1444 | Bacteria;Bacteroidota;Bacteroidia;Bacteroidales;Bacteroidaceae;Bacteroides;uncultured organism | 90.8 | 1313 | -1 |
| metaRNA.PFemirge\_4544 | 11 | 0.000000 | BAAZ01000703.1371.2875 | Bacteria;Bacteroidota;Bacteroidia;Bacteroidales;Tannerellaceae;Parabacteroides;human gut metagenome | 91.3 | 1236 | -1 |
| metaRNA.PFemirge\_25220 | 10 | 0.000001 | AY494079.33242.34761 | Bacteria;Proteobacteria;Alphaproteobacteria;Rickettsiales;Mitochondria;Fucus vesiculosus | 99.7 | 1181 | -1 |
| metaRNA.PFemirge\_5105 | 10 | 0.000003 | JQ924024.1.1501 | Bacteria;Proteobacteria;Gammaproteobacteria;Burkholderiales;Rhodocyclaceae;Dechlorosoma;uncultured bacterium | 92.4 | 1412 | -1 |
| metaRNA.PFemirge\_13123 | 10 | 0.000001 | HQ741361.1.1434 | Bacteria;Firmicutes;Clostridia;Lachnospirales;Lachnospiraceae;Epulopiscium;uncultured bacterium | 94.0 | 1310 | -1 |
| metaRNA.PFemirge\_10925 | 10 | 0.000001 | HQ741595.1.1456 | Bacteria;Firmicutes;Clostridia;Lachnospirales;Lachnospiraceae;Epulopiscium;uncultured bacterium | 93.0 | 1406 | -1 |
| metaRNA.PFemirge\_13831 | 10 | 0.000002 | HQ741592.1.1434 | Bacteria;Firmicutes;Clostridia;Lachnospirales;Lachnospiraceae;Epulopiscium;uncultured bacterium | 94.2 | 1303 | -1 |
| metaRNA.PFemirge\_12794 | 10 | 0.000013 | EF071142.1.1509 | Bacteria;Bacteroidota;Bacteroidia;Bacteroidales;Bacteroidaceae;Bacteroides;uncultured Bacteroidetes bacterium | 89.6 | 1222 | -1 |
| metaRNA.PFemirge\_16522 | 10 | 0.000074 | EU772076.1.1400 | Bacteria;Firmicutes;Clostridia;Lachnospirales;Lachnospiraceae;Lachnospiraceae UCG-008;uncultured bacterium | 93.6 | 1303 | -1 |
| metaRNA.PFemirge\_5891 | 10 | 0.000001 | HQ767134.1.1448 | Bacteria;Bacteroidota;Bacteroidia;Bacteroidales;Tannerellaceae;Parabacteroides;uncultured organism | 90.4 | 1270 | -1 |
| metaRNA.PFemirge\_3270 | 9 | 0.000002 | CP002345.1438824.1440339 | Bacteria;Bacteroidota;Bacteroidia;Bacteroidales;Paludibacteraceae;Paludibacter;Paludibacter propionicigenes WB4 | 88.7 | 1495 | -1 |
| metaRNA.PFemirge\_21528 | 9 | 0.000002 | HQ795338.1.1448 | Bacteria;Firmicutes;Clostridia;Oscillospirales;Ruminococcaceae;Subdoligranulum;uncultured organism | 92.5 | 1288 | -1 |
| metaRNA.PFemirge\_11932 | 9 | 0.000002 | MF383477.1.1442 | Bacteria;Proteobacteria;Gammaproteobacteria;Pseudomonadales;Pseudomonadaceae;Pseudomonas;Pseudomonas sp. | 99.3 | 1203 | -1 |
| metaRNA.PFemirge\_21215 | 8 | 0.000007 | DQ799921.1.1390 | Bacteria;Firmicutes;Clostridia;Oscillospirales;Ruminococcaceae;Faecalibacterium;uncultured bacterium | 90.7 | 1284 | -1 |
| metaRNA.PFemirge\_26218 | 8 | 0.000001 | FJ957875.1.1447 | Bacteria;Firmicutes;Clostridia;Lachnospirales;Lachnospiraceae;Lachnoclostridium;[Clostridium] saccharolyticum | 93.7 | 1284 | -1 |
| metaRNA.PFemirge\_20408 | 8 | 0.000001 | EU779425.1.1388 | Bacteria;Firmicutes;Clostridia;Oscillospirales;[Eubacterium] coprostanoligenes group;uncultured bacterium | 93.7 | 1280 | -1 |
| metaRNA.PFemirge\_20621 | 8 | 0.000002 | DQ810054.1.1385 | Bacteria;Firmicutes;Clostridia;Lachnospirales;Lachnospiraceae;Lachnoclostridium;uncultured bacterium | 92.6 | 1292 | -1 |
| metaRNA.PFemirge\_23926 | 8 | 0.000005 | GQ135950.1.1352 | Bacteria;Firmicutes;Clostridia;Clostridiales;Clostridiaceae;Clostridium sensu stricto 1;uncultured bacterium | 93.8 | 1254 | -1 |
| metaRNA.PFemirge\_3956 | 8 | 0.000000 | JQ084791.1.1476 | Bacteria;Firmicutes;Clostridia;Oscillospirales;Ruminococcaceae;Anaerotruncus;uncultured bacterium | 93.0 | 1321 | -1 |
| metaRNA.PFemirge\_20030 | 8 | 0.000000 | DQ905437.1.1471 | Bacteria;Firmicutes;Clostridia;Oscillospirales;Ruminococcaceae;uncultured;uncultured bacterium | 92.3 | 1277 | -1 |
| metaRNA.PFemirge\_6206 | 8 | 0.000006 | HQ739338.1.1431 | Bacteria;Bacteroidota;Bacteroidia;Bacteroidales;Bacteroidaceae;Bacteroides;uncultured bacterium | 90.1 | 1318 | -1 |
| metaRNA.PFemirge\_19434 | 8 | 0.000001 | EF096560.1.1389 | Bacteria;Firmicutes;Clostridia;Lachnospirales;Lachnospiraceae;Lachnospiraceae UCG-008;uncultured bacterium | 92.5 | 1374 | -1 |
| metaRNA.PFemirge\_26828 | 8 | 0.000001 | KF945050.1.1264 | Bacteria;Myxococcota;Polyangia;Haliangiales;Haliangiaceae;Haliangium;uncultured bacterium | 99.7 | 652 | -1 |
| metaRNA.PFemirge\_14287 | 8 | 0.000002 | EU104080.1.1428 | Bacteria;Bacteroidota;Bacteroidia;Chitinophagales;Saprospiraceae;uncultured;uncultured bacterium | 97.9 | 1138 | -1 |
| metaRNA.PFemirge\_30265 | 6 | 0.000015 | AB574482.1.1489 | Bacteria;Bacteroidota;Bacteroidia;Bacteroidales;Tannerellaceae;Parabacteroides;Parabacteroides chinchillae | 86.7 | 1402 | -1 |
| metaRNA.PFemirge\_26292 | 6 | 0.000001 | FR749937.1.1523 | Bacteria;Firmicutes;Clostridia;Lachnospirales;Lachnospiraceae;[Eubacterium] fissicatena group;Faecalicatena fissicatena | 95.2 | 1185 | -1 |
| metaRNA.PFemirge\_4551 | 6 | 0.000003 | EU560770.1.1499 | Bacteria;Proteobacteria;Gammaproteobacteria;Pseudomonadales;Pseudomonadaceae;Pseudomonas;uncultured bacterium | 95.9 | 1479 | -1 |
| metaRNA.PFemirge\_16700 | 5 | 0.000000 | HM630205.1.1449 | Bacteria;Bacteroidota;Bacteroidia;Bacteroidales;Rikenellaceae;Rikenella;uncultured bacterium | 87.1 | 1181 | -1 |
| metaRNA.PFemirge\_20985 | 5 | 0.000000 | AB746664.1.1536 | Bacteria;Firmicutes;Clostridia;Lachnospirales;Lachnospiraceae;Lachnospiraceae NK4A136 group;uncultured bacterium | 90.2 | 1308 | -1 |
| metaRNA.PFemirge\_19950 | 5 | 0.000001 | EU774669.1.1387 | Bacteria;Firmicutes;Clostridia;Peptostreptococcales-Tissierellales;Anaerovoracaceae;[Eubacterium] nodatum group;uncultured bacterium | 95.1 | 1286 | -1 |
| metaRNA.PFemirge\_10915 | 4 | 0.000003 | GU222249.1.1456 | Bacteria;Proteobacteria;Gammaproteobacteria;Burkholderiales;Comamonadaceae;Simplicispira;uncultured bacterium | 94.4 | 1384 | -1 |
| metaRNA.PFemirge\_30233 | 4 | 0.000082 | HM630179.1.1443 | Bacteria;Cyanobacteria;Vampirivibrionia;Gastranaerophilales;uncultured bacterium | 93.4 | 1445 | -1 |
| metaRNA.PFemirge\_25519 | 3 | 0.000007 | KM251124.1.1542 | Bacteria;Firmicutes;Clostridia;Oscillospirales;Hydrogenoanaerobacterium;uncultured bacterium | 92.4 | 1320 | -1 |
| metaRNA.PFemirge\_30244 | 3 | 0.000034 | EU775962.1.1393 | Bacteria;Bacteroidota;Bacteroidia;Bacteroidales;Bacteroidaceae;Bacteroides;uncultured bacterium | 90.1 | 1397 | -1 |
| metaRNA.PFemirge\_8549 | 1 | 0.000000 | JN680618.1.1455 | Bacteria;Firmicutes;Clostridia;Oscillospirales;Butyricicoccaceae;UCG-008;uncultured Ruminococcaceae bacterium | 93.9 | 1318 | -1 |
| metaRNA.PFemirge\_20171 |  | 0.000000 | FJ368696.1.1354 | Bacteria;Firmicutes;Clostridia;Oscillospirales;Oscillospiraceae;uncultured;uncultured bacterium | 90.2 | 1169 | -1 |


##### SSU rRNA from trusted contigs

Full-length SSU rRNA seqeunces extracted from trusted contigs file, matched to SILVA database by Vsearch.

| OTU | Mapped | DB hit | Taxonomy | % ID | Alnlen | Evalue |
| --- | --- | --- | --- | --- | --- | --- |
| metaRNA.PFtrusted\_13 | 91903 | EU773260.1.1381 | Bacteria;Firmicutes;Clostridia;Lachnospirales;Lachnospiraceae;Tyzzerella;uncultured bacterium | 87.5 | 1090 | -1 |
| metaRNA.PFtrusted\_8 | 76492 | EF401177.1.1491 | Bacteria;Firmicutes;Clostridia;Lachnospirales;Lachnospiraceae;uncultured;uncultured bacterium | 82.7 | 1161 | -1 |
| metaRNA.PFtrusted\_9 | 60409 | EF401177.1.1491 | Bacteria;Firmicutes;Clostridia;Lachnospirales;Lachnospiraceae;uncultured;uncultured bacterium | 87.7 | 1150 | -1 |
| metaRNA.PFtrusted\_10 | 41455 | FM957479.1.1497 | Bacteria;Proteobacteria;Gammaproteobacteria;Enterobacterales;Vibrionaceae;Vibrio;Vibrio sp. MY-2008-U67 | 84.9 | 1156 | -1 |
| metaRNA.PFtrusted\_12 | 2128 | HQ813979.1.1426 | Bacteria;Firmicutes;Clostridia;Lachnospirales;Lachnospiraceae;uncultured;uncultured organism | 84.3 | 1113 | -1 |
| metaRNA.PFtrusted\_11 | 831 | BAAU01002038.417.1926 | Bacteria;Firmicutes;Clostridia;Lachnospirales;Lachnospiraceae;uncultured;human gut metagenome | 89.5 | 1055 | -1 |
| metaRNA.PFtrusted\_14 | 38 | BAAU01002038.417.1926 | Bacteria;Firmicutes;Clostridia;Lachnospirales;Lachnospiraceae;uncultured;human gut metagenome | 88.2 | 1147 | -1 |


##### Taxonomic affiliation of unassembled SSU rRNA reads

Approximate overview of taxonomic composition for reads that did NOT assemble into full-length sequences, based on mapping hits to SILVA SSU rRNA database with BBmap.

Taxonomy summarized at level 4. Only displaying taxa with > 3 reads mapped.

| Taxon | Reads |
| --- | --- |
| Bacteria;Bacteroidota;Bacteroidia;Bacteroidales | 107875 |
| Eukaryota;SAR;Rhizaria;Retaria | 51670 |
| Bacteria;Proteobacteria;Gammaproteobacteria;Enterobacterales | 44241 |
| Bacteria;Desulfobacterota;Desulfovibrionia;Desulfovibrionales | 33947 |
| Eukaryota;SAR;Stramenopiles;(Stramenopiles) | 27273 |
| Bacteria;(Bacteria);(Bacteria);(Bacteria) | 26397 |
| Eukaryota;SAR;(SAR);(SAR) | 24419 |
| Bacteria;Firmicutes;Clostridia;Oscillospirales | 21853 |
| Eukaryota;SAR;Stramenopiles;Opalinata | 21174 |
| Eukaryota;(Eukaryota);(Eukaryota);(Eukaryota) | 17102 |
| Bacteria;Firmicutes;Clostridia;Lachnospirales | 13341 |
| Eukaryota;Amorphea;Amoebozoa;Archamoebae | 12349 |
| Bacteria;Actinobacteriota;Actinobacteria;Micrococcales | 8667 |
| Eukaryota;Amorphea;Obazoa;Opisthokonta | 8359 |
| Bacteria;Firmicutes;Negativicutes;Acidaminococcales | 6784 |
| Bacteria;Bacteroidota;Bacteroidia;(Bacteroidia) | 6722 |
| Eukaryota;Amorphea;Amoebozoa;Discosea | 6635 |
| Eukaryota;SAR;Alveolata;Ciliophora | 5169 |
| Bacteria;Proteobacteria;Gammaproteobacteria;Burkholderiales | 4301 |
| Bacteria;Proteobacteria;Alphaproteobacteria;(Alphaproteobacteria) | 4170 |
| Bacteria;Cyanobacteria;Vampirivibrionia;Gastranaerophilales | 3907 |
| Bacteria;Firmicutes;(Firmicutes);(Firmicutes) | 3722 |
| Eukaryota;SAR;Stramenopiles;Incertae Sedis | 3566 |
| Bacteria;Bacteroidota;Bacteroidia;Flavobacteriales | 3548 |
| Bacteria;Proteobacteria;Alphaproteobacteria;Rhodospirillales | 3310 |
| Bacteria;Firmicutes;Clostridia;Peptostreptococcales-Tissierellales | 3214 |
| Bacteria;Firmicutes;Clostridia;(Clostridia) | 2604 |
| Bacteria;Proteobacteria;(Proteobacteria);(Proteobacteria) | 2241 |
| Bacteria;Firmicutes;Bacilli;Erysipelotrichales | 2025 |
| Bacteria;Fusobacteriota;Fusobacteriia;Fusobacteriales | 1798 |
| Bacteria;Proteobacteria;Alphaproteobacteria;Rhizobiales | 1415 |
| Eukaryota;Archaeplastida;Chloroplastida;Charophyta | 1312 |
| Bacteria;Desulfobacterota;(Desulfobacterota);(Desulfobacterota) | 1261 |
| Bacteria;Firmicutes;Bacilli;RF39 | 1243 |
| Bacteria;Bacteroidota;Bacteroidia;Chitinophagales | 1200 |
| Bacteria;Firmicutes;Clostridia;Clostridia UCG-014 | 1170 |
| Bacteria;Firmicutes;Bacilli;(Bacilli) | 1093 |
| Bacteria;Firmicutes;Bacilli;Bacillales | 1035 |
| Bacteria;Firmicutes;Negativicutes;Veillonellales-Selenomonadales | 1014 |
| Bacteria;Verrucomicrobiota;Verrucomicrobiae;Verrucomicrobiales | 991 |
| Bacteria;Actinobacteriota;Actinobacteria;(Actinobacteria) | 883 |
| Bacteria;Proteobacteria;Alphaproteobacteria;Holosporales | 870 |
| Bacteria;Firmicutes;Negativicutes;(Negativicutes) | 745 |
| Eukaryota;Amorphea;Amoebozoa;uncultured eukaryote | 675 |
| Eukaryota;SAR;Stramenopiles;MAST-3 | 629 |
| Bacteria;Verrucomicrobiota;Chlamydiae;Chlamydiales | 606 |
| Bacteria;Proteobacteria;Gammaproteobacteria;(Gammaproteobacteria) | 594 |
| Bacteria;Verrucomicrobiota;Verrucomicrobiae;(Verrucomicrobiae) | 548 |
| Eukaryota;Discoba;Discicristata;Euglenozoa | 523 |
| Bacteria;Proteobacteria;Gammaproteobacteria;Pseudomonadales | 516 |
| Bacteria;Firmicutes;Bacilli;Lactobacillales | 500 |
| Eukaryota;SAR;Stramenopiles;Labyrinthulomycetes | 468 |
| Eukaryota;Amorphea;(Amorphea);(Amorphea) | 453 |
| Bacteria;Firmicutes;Clostridia;Clostridiales | 404 |
| Eukaryota;Amorphea;Amoebozoa;(Amoebozoa) | 391 |
| Bacteria;Firmicutes;Clostridia;Christensenellales | 385 |
| Bacteria;Synergistota;Synergistia;Synergistales | 377 |
| Eukaryota;SAR;Rhizaria;Cercozoa | 352 |
| Bacteria;Cyanobacteria;Cyanobacteriia;Chloroplast | 328 |
| Bacteria;Proteobacteria;Gammaproteobacteria;Legionellales | 275 |
| Bacteria;Actinobacteriota;Actinobacteria;Corynebacteriales | 274 |
| Eukaryota;Amorphea;Amoebozoa;SAR | 264 |
| Bacteria;Proteobacteria;Alphaproteobacteria;Rickettsiales | 252 |
| Eukaryota;SAR;Stramenopiles;Ochrophyta | 252 |
| Bacteria;Actinobacteriota;Actinobacteria;Actinomycetales | 245 |
| Bacteria;Cyanobacteria;Cyanobacteriia;Cyanobacteriales | 242 |
| Bacteria;Acidobacteriota;Vicinamibacteria;Vicinamibacterales | 241 |
| Bacteria;Proteobacteria;Gammaproteobacteria;Xanthomonadales | 221 |
| Bacteria;Bacteroidota;Bacteroidia;Sphingobacteriales | 218 |
| Bacteria;Verrucomicrobiota;Verrucomicrobiae;Chthoniobacterales | 217 |
| Bacteria;Deferribacterota;Deferribacteres;Deferribacterales | 212 |
| Bacteria;Bacteroidota;Bacteroidia;Cytophagales | 212 |
| Eukaryota;SAR;Stramenopiles;Peronosporomycetes | 205 |
| Bacteria;Firmicutes;Bacilli;Paenibacillales | 201 |
| Bacteria;Verrucomicrobiota;Lentisphaeria;Victivallales | 192 |
| Eukaryota;Amorphea;Amoebozoa;Incertae Sedis | 176 |
| Bacteria;Desulfobacterota;Desulfobulbia;Desulfobulbales | 175 |
| Bacteria;Proteobacteria;Gammaproteobacteria;Diplorickettsiales | 174 |
| Bacteria;Proteobacteria;Alphaproteobacteria;Paracaedibacterales | 153 |
| Bacteria;Proteobacteria;Alphaproteobacteria;Dongiales | 151 |
| Bacteria;Verrucomicrobiota;Verrucomicrobiae;Opitutales | 123 |
| Bacteria;Proteobacteria;Alphaproteobacteria;Sphingomonadales | 116 |
| Bacteria;Acidobacteriota;Holophagae;Subgroup 7 | 115 |
| Bacteria;Desulfobacterota;Desulfuromonadia;Geobacterales | 114 |
| Eukaryota;Archaeplastida;Chloroplastida;Chlorophyta | 108 |
| Bacteria;Proteobacteria;Alphaproteobacteria;Kiloniellales | 106 |
| Bacteria;Proteobacteria;Alphaproteobacteria;Rhodobacterales | 106 |
| Bacteria;Myxococcota;Polyangia;Polyangiales | 104 |
| Bacteria;Proteobacteria;Alphaproteobacteria;Azospirillales | 95 |
| Bacteria;Firmicutes;Bacilli;Staphylococcales | 89 |
| Bacteria;Firmicutes;Clostridia;Monoglobales | 80 |
| Bacteria;Planctomycetota;Phycisphaerae;S-70 | 78 |
| Bacteria;Planctomycetota;Planctomycetes;Pirellulales | 74 |
| Bacteria;Actinobacteriota;Coriobacteriia;Coriobacteriales | 72 |
| Eukaryota;SAR;Alveolata;Protalveolata | 68 |
| Bacteria;Actinobacteriota;Actinobacteria;Propionibacteriales | 68 |
| Bacteria;Firmicutes;Clostridia;Eubacteriales | 65 |
| Bacteria;Firmicutes;Bacilli;RsaHf231 | 65 |
| Bacteria;Proteobacteria;Alphaproteobacteria;Reyranellales | 65 |
| Bacteria;Proteobacteria;Gammaproteobacteria;Gammaproteobacteria Incertae Sedis | 63 |
| Bacteria;Proteobacteria;Alphaproteobacteria;Caedibacterales | 62 |
| Bacteria;Firmicutes;Bacilli;Haloplasmatales | 58 |
| Bacteria;Actinobacteriota;Actinobacteria;PeM15 | 56 |
| Bacteria;Proteobacteria;Gammaproteobacteria;Coxiellales | 55 |
| Bacteria;Proteobacteria;Alphaproteobacteria;uncultured | 54 |
| Bacteria;Cyanobacteria;Vampirivibrionia;Obscuribacterales | 52 |
| Bacteria;Firmicutes;Clostridia;Clostridia vadinBB60 group | 51 |
| Bacteria;Firmicutes;Clostridia;Peptococcales | 51 |
| Bacteria;Verrucomicrobiota;Verrucomicrobiae;Pedosphaerales | 50 |
| Bacteria;Proteobacteria;Alphaproteobacteria;Elsterales | 50 |
| Bacteria;Verrucomicrobiota;Lentisphaeria;(Lentisphaeria) | 44 |
| Bacteria;Bdellovibrionota;Bdellovibrionia;Bdellovibrionales | 44 |
| Bacteria;Bacteroidota;(Bacteroidota);(Bacteroidota) | 43 |
| Bacteria;Proteobacteria;Gammaproteobacteria;Ectothiorhodospirales | 42 |
| Eukaryota;SAR;Alveolata;(Alveolata) | 42 |
| Bacteria;Firmicutes;Desulfitobacteriia;Desulfitobacteriales | 41 |
| Bacteria;Planctomycetota;Planctomycetes;Gemmatales | 40 |
| Bacteria;Proteobacteria;Alphaproteobacteria;Caulobacterales | 37 |
| Bacteria;Spirochaetota;Brachyspirae;Brachyspirales | 37 |
| Bacteria;Actinobacteriota;Acidimicrobiia;Microtrichales | 36 |
| Bacteria;Acidobacteriota;(Acidobacteriota);(Acidobacteriota) | 36 |
| Bacteria;Firmicutes;Desulfotomaculia;Desulfotomaculales | 34 |
| Bacteria;Patescibacteria;Microgenomatia;uncultured sediment bacterium | 34 |
| Eukaryota;Amorphea;Amoebozoa;metagenome | 34 |
| Bacteria;Planctomycetota;vadinHA49;uncultured bacterium | 34 |
| Bacteria;Planctomycetota;Planctomycetes;Planctomycetales | 33 |
| Bacteria;Actinobacteriota;(Actinobacteriota);(Actinobacteriota) | 31 |
| Bacteria;Dependentiae;Babeliae;Babeliales | 31 |
| Bacteria;Proteobacteria;Alphaproteobacteria;Zavarziniales | 30 |
| Bacteria;Actinobacteriota;Actinobacteria;Pseudonocardiales | 30 |
| Bacteria;Verrucomicrobiota;Kiritimatiellae;Kiritimatiellales | 28 |
| Bacteria;Planctomycetota;Planctomycetes;Isosphaerales | 26 |
| Bacteria;Bacteroidota;Kapabacteria;Kapabacteriales | 26 |
| Bacteria;Proteobacteria;Alphaproteobacteria;Acetobacterales | 25 |
| Eukaryota;Amorphea;Amoebozoa;Tubulinea | 25 |
| Bacteria;Firmicutes;Bacilli;Thermoactinomycetales | 24 |
| Bacteria;Nitrospirota;Thermodesulfovibrionia;uncultured | 23 |
| Bacteria;Cyanobacteria;Cyanobacteriia;(Cyanobacteriia) | 22 |
| Bacteria;Firmicutes;Bacilli;Exiguobacterales | 22 |
| Bacteria;Deinococcota;Deinococci;Deinococcales | 22 |
| Bacteria;Planctomycetota;Phycisphaerae;Phycisphaerales | 20 |
| Bacteria;Desulfobacterota;Desulfuromonadia;Desulfuromonadales | 20 |
| Eukaryota;uncultured Entamoeba;(uncultured Entamoeba);(uncultured Entamoeba) | 19 |
| Bacteria;Firmicutes;Clostridia;uncultured | 19 |
| Eukaryota;Archaeplastida;Chloroplastida;(Chloroplastida) | 18 |
| Bacteria;RCP2-54;metagenome;(metagenome) | 18 |
| Bacteria;Firmicutes;Bacilli;Brevibacillales | 18 |
| Bacteria;Acidobacteriota;Holophagae;(Holophagae) | 18 |
| Bacteria;Firmicutes;Thermoanaerobacteria;Thermoanaerobacterales | 18 |
| Eukaryota;SAR;Alveolata;uncultured | 18 |
| Eukaryota;Archaeplastida;Rhodophyceae;Bangiales | 18 |
| Bacteria;Patescibacteria;Gracilibacteria;metagenome | 18 |
| Bacteria;Myxococcota;Myxococcia;Myxococcales | 17 |
| Bacteria;Actinobacteriota;Actinobacteria;Streptomycetales | 17 |
| Eukaryota;SAR;Alveolata;Apicomplexa | 16 |
| Bacteria;Bdellovibrionota;Oligoflexia;0319-6G20 | 16 |
| Bacteria;Verrucomicrobiota;Verrucomicrobiae;Arctic97B-4 marine group | 16 |
| Eukaryota;Amorphea;Amoebozoa;Schizoplasmodiida | 16 |
| Bacteria;Desulfobacterota;Desulfuromonadia;(Desulfuromonadia) | 16 |
| Bacteria;Myxococcota;Polyangia;Haliangiales | 16 |
| Bacteria;Patescibacteria;Gracilibacteria;uncultured bacterium | 15 |
| Bacteria;Proteobacteria;Gammaproteobacteria;Steroidobacterales | 15 |
| Bacteria;Myxococcota;Polyangia;mle1-27 | 14 |
| Bacteria;Verrucomicrobiota;Verrucomicrobiae;Methylacidiphilales | 14 |
| Bacteria;Proteobacteria;Alphaproteobacteria;AT-s3-44 | 14 |
| Bacteria;Desulfobacterota;Syntrophia;Syntrophales | 14 |
| Bacteria;Proteobacteria;Gammaproteobacteria;Granulosicoccales | 13 |
| Bacteria;Proteobacteria;Gammaproteobacteria;Methylococcales | 13 |
| Bacteria;Patescibacteria;Gracilibacteria;(Gracilibacteria) | 13 |
| Bacteria;Dadabacteria;Dadabacteriia;Dadabacteriales | 13 |
| Eukaryota;Excavata;Metamonada;Preaxostyla | 12 |
| Bacteria;Calditrichota;Calditrichia;Calditrichales | 12 |
| Eukaryota;Cryptophyceae;Cryptomonadales;Cryptomonas | 12 |
| Bacteria;Patescibacteria;Microgenomatia;Candidatus Roizmanbacteria | 12 |
| Bacteria;Hydrogenedentes;Hydrogenedentia;Hydrogenedentiales | 12 |
| Bacteria;Campylobacterota;Campylobacteria;Campylobacterales | 11 |
| Bacteria;Acidobacteriota;Subgroup 22;uncultured bacterium | 11 |
| Bacteria;Firmicutes;Clostridia;Gracilibacteraceae | 11 |
| Bacteria;Proteobacteria;Gammaproteobacteria;uncultured | 11 |
| Bacteria;Actinobacteriota;Thermoleophilia;Gaiellales | 11 |
| Bacteria;Spirochaetota;Spirochaetia;Spirochaetales | 11 |
| Bacteria;Chloroflexi;Anaerolineae;SBR1031 | 10 |
| Bacteria;Firmicutes;Bacilli;Aneurinibacillales | 10 |
| Eukaryota;SAR;Rhizaria;(Rhizaria) | 10 |
| Bacteria;Proteobacteria;Gammaproteobacteria;Acidithiobacillales | 10 |
| Bacteria;Actinobacteriota;Actinobacteria;Streptosporangiales | 10 |
| Bacteria;Chloroflexi;Chloroflexia;Chloroflexales | 10 |
| Bacteria;Verrucomicrobiota;(Verrucomicrobiota);(Verrucomicrobiota) | 9 |
| Eukaryota;Amorphea;Amoebozoa;Gracilipodida | 8 |
| Bacteria;Firmicutes;Moorellia;Desulfitibacterales | 8 |
| Bacteria;Proteobacteria;Alphaproteobacteria;Kordiimonadales | 8 |
| Bacteria;Desulfobacterota;uncultured;uncultured delta proteobacterium | 8 |
| Bacteria;RCP2-54;uncultured bacterium;(uncultured bacterium) | 8 |
| Eukaryota;SAR;Stramenopiles;MAST-12 | 8 |
| Bacteria;Myxococcota;Polyangia;(Polyangia) | 8 |
| Bacteria;Methylomirabilota;Methylomirabilia;Methylomirabilales | 7 |
| Bacteria;Proteobacteria;Alphaproteobacteria;Sneathiellales | 7 |
| Bacteria;Gemmatimonadota;Gemmatimonadetes;Gemmatimonadales | 7 |
| Bacteria;Proteobacteria;Alphaproteobacteria;Defluviicoccales | 7 |
| Bacteria;Abditibacteriota;Abditibacteria;Abditibacteriales | 7 |
| Bacteria;Proteobacteria;Gammaproteobacteria;Cardiobacteriales | 7 |
| Bacteria;Myxococcota;Polyangia;Blfdi19 | 7 |
| Bacteria;Campylobacterota;Desulfurellia;Desulfurellales | 6 |
| Bacteria;Firmicutes;Symbiobacteriia;Symbiobacteriales | 6 |
| Bacteria;Myxococcota;bacteriap25;metagenome | 6 |
| Bacteria;Proteobacteria;Gammaproteobacteria;EV818SWSAP88 | 6 |
| Bacteria;Acidobacteriota;Blastocatellia;Blastocatellales | 6 |
| Bacteria;Firmicutes;Dethiobacteria;Dethiobacterales | 6 |
| Bacteria;Myxococcota;bacteriap25;uncultured bacterium | 6 |
| Bacteria;Proteobacteria;Gammaproteobacteria;Thiotrichales | 6 |
| Bacteria;Actinobacteriota;Actinobacteria;Bifidobacteriales | 6 |
| Bacteria;Firmicutes;Bacilli;Acholeplasmatales | 5 |
| Bacteria;Firmicutes;Syntrophomonadia;Syntrophomonadales | 5 |
| Bacteria;Firmicutes;Negativicutes;uncultured | 5 |
| Bacteria;Bdellovibrionota;Oligoflexia;Oligoflexales | 5 |
| Bacteria;Sva0485;uncultured bacterium;(uncultured bacterium) | 5 |
| Bacteria;Desulfobacterota;Desulfobacteria;Desulfatiglandales | 5 |
| Bacteria;Firmicutes;Incertae Sedis;Gelria | 5 |
| Bacteria;Firmicutes;Clostridia;Caldicoprobacterales | 5 |
| Bacteria;Chloroflexi;Chloroflexia;Thermomicrobiales | 5 |
| Bacteria;Firmicutes;D8A-2;uncultured bacterium | 5 |
| Bacteria;Patescibacteria;Parcubacteria;CG1-02-40-25 | 4 |
| Bacteria;Myxococcota;(Myxococcota);(Myxococcota) | 4 |
| Eukaryota;Amorphea;Amoebozoa;Mycamoeba | 4 |
| Bacteria;Patescibacteria;CPR2;uncultured eubacterium WCHB1-60 | 4 |
| Bacteria;Chloroflexi;TK10;metagenome | 4 |
| Bacteria;Chloroflexi;Ktedonobacteria;Ktedonobacterales | 4 |
| Bacteria;Proteobacteria;Alphaproteobacteria;Micavibrionales | 4 |
| Bacteria;Actinobacteriota;Thermoleophilia;Solirubrobacterales | 4 |
| Bacteria;Firmicutes;Clostridia;Thermincolales | 4 |
| Bacteria;Desulfobacterota;uncultured;uncultured Desulfuromonadales bacterium | 4 |
| Bacteria;Cyanobacteria;(Cyanobacteria);(Cyanobacteria) | 4 |
| Bacteria;Nitrospirota;Nitrospiria;Nitrospirales | 4 |
| Bacteria;Proteobacteria;Gammaproteobacteria;Piscirickettsiales | 4 |
| Bacteria;Planctomycetota;Phycisphaerae;MSBL9 | 4 |
| Eukaryota;Haptophyta;T58;uncultured marine haptophyte | 4 |
| Bacteria;Marinimicrobia (SAR406 clade);uncultured bacterium;(uncultured bacterium) | 4 |
| Bacteria;Desulfobacterota;Desulfuromonadia;PB19 | 4 |
| Bacteria;Planctomycetota;Phycisphaerae;Tepidisphaerales | 4 |
| Bacteria;Actinobacteriota;Actinobacteria;Frankiales | 4 |
| Eukaryota;Discoba;Discicristata;Heterolobosea | 4 |
| Bacteria;Acidobacteriota;Subgroup 25;uncultured Acidobacteria bacterium | 4 |
| Bacteria;Planctomycetota;BD7-11;uncultured bacterium | 4 |

##### Please cite...

Harald R Gruber-Vodicka, Brandon KB Seah, Elmar Pruesse. 2019 (preprint). phyloFlash - Rapid SSU rRNA profiling and targeted assembly from metagenomes. bioRxiv 521922; doi: https://doi.org/10.1101/521922

###### Cite dependencies when used

- Quast C. et al. 2013. The SILVA ribosomal RNA gene database project: improved data processing and web-based tools. *Nucl. Acids Res.* 41: D590-D596. doi:10.1093/nar/gks1219. Homepage
- Bushnell B. BBMap. Online: https://sourceforge.net/projects/bbmap/
- Bankevich A., et al. 2012. SPAdes: A New Genome Assembly Algorithm and Its Applications to Single-Cell Sequencing. *J. Comput. Biol.* 19 (5): 455-477. doi:10.1089/cmb.2012.0021. Homepage.
- Katoh K., Standley D.M.. 2013. MAFFT Multiple Sequence Alignment Software Version 7: Improvements in Performance and Usability. *Mol. Biol. Evol.* 30 (4): 772-780. doi:10.1093/molbev/mst010. Homepage
- Rognes T. et al. 2016. VSEARCH: a versatile open source tool for metagenomics. *PeerJ* 4: e2584. doi:10.7717/peerj.2584. Homepage
- Miller C.S. et al. 2011. EMIRGE: reconstruction of full-length ribosomal genes from microbial community short read sequencing data. *Genome Biol.* 12: R44. doi:10.1186/gb-2011-12-5-r44. Homepage
- Kopylova E., Noé L., Touzet H. 2012. SortMeRNA: fast and accurate filtering of ribosomal RNAs in metatranscriptomic data. *Bioinformatics* 28 (24): 3211-3217. doi: 10.1093/bioinformatics/bts611. Homepage
- Bedtools. Online: http://bedtools.readthedocs.io/en/latest/
- Seemann T. Barrnap. Online: https://github.com/tseemann/barrnap
- Wheeler T.J., Eddy S.R. 2013. nhmmer: DNA homology search with profile HMMs. *Bioinformatics* 29 (19): 2487-2489. doi:10.1093/bioinformatics/btt403 Homepage
