## Supplementary material for "Gut microbial ecology of Xenopus tadpoles across life stages": Figure_S13: Figure_S13_metaDNA.kegg.minpath.html

Javascript must be enabled to view this page.

magnitude
magnitudeUnassigned

metaDNA.krona.kegg.minpath

5733.84252960327

233.113125041353

39.1172610333333

39.1172610333333

62.9410917746113

44.4062910588218

18.5348007157895

131.054772233408

30.1349003013699

21.0894987373016

75.9352914947369

3.8950817

112.853512191111

81.6094829966667

81.6094829966667

31.2440291944444

31.2440291944444

53.0138210010204

46.0956354581633

46.0956354581633

6.91818554285714

6.91818554285714

4905.0160593892

93.1997656382644

14.8803693659574

24.6702642744227

21.4351422266667

10.5433189403846

21.670670830833

92.0052206846732

53.2186230088005

38.7865976758727

315.083464045368

83.587878385529

35.4618414634956

64.2706435555556

81.0491840891751

50.713916551613

376.333730258646

75.0905185925

34.1234757081476

22.5042575623519

27.236021240555

50.8009226124958

27.7017160447778

23.5768054218536

43.2209390586468

53.7909624458368

18.2881115714809

384.14467696047

32.5778674877065

84.5341156701594

22.3324680661692

111.153880099067

44.2608735626117

35.7107804453007

17.412853575

20.9314296544559

15.2304084

190.944466407043

11.47223575

36.5538405

27.5353142539788

115.383075903064

790.426086015655

31.6595742118475

55.7859833593431

82.6601650630421

28.9943094148984

90.7490184393756

37.6702829019066

41.5329643192273

46.0350099356873

46.8300739402249

82.6729651926714

46.328823403464

78.3367852246556

28.156363090345

93.0137675189655

844.742394435465

79.253395122807

28.2772869098246

226.032085706358

84.7616400114035

39.3397152

158.287669603333

89.8572411999999

52.457038623405

86.4763220583333

877.212139660856

46.8037365997308

39.599522104767

124.188598760847

57.366616375465

69.7013893028732

54.5360708925884

64.529882214277

59.5994530485424

87.7051245090288

92.3568689239488

118.255628116805

62.5692488119827

325.281564322063

22.2027873074638

23.1090750498632

18.0003779744444

25.7978695336111

34.5364568066807

201.63499765

127.245211679635

65.9873662880289

61.2578453916065

488.397339281061

142.558184365337

20.2849876155195

34.7024309575141

82.199775683858

102.120399506395

34.7087787249063

34.1261856480074

28.7777760861899

8.91882069333333

101.470213539238

86.9643372678095

54.663723204

16.9242686571429

15.3763454066667

14.5058762714286

14.5058762714286

0

0

0

328.375798441349

42.8483623233333

42.8483623233333

285.527436118015

46.3906273575758

37.1360825

202.00072626044
