## Supplementary material for "Gut microbial ecology of Xenopus tadpoles across life stages": Figure_S14: Figure_S14_metaDNA.metacyc.minpath.html

Javascript must be enabled to view this page.

magnitude
magnitudeUnassigned

metaDNA.krona.metacyc.minpath

33493.2533488257

44.405881035131

44.405881035131

39.2592349896384

22.7968714827052

16.4623635069332

241.931757295942

241.931757295942

0

0

0

0

0

0

0

0

16169.0181674559

186.855191397256

44.5712059174273

55.0523328733035

55.0523328733035

39.3397152

47.8919374065249

1216.74808397311

1166.13208002311

562.540797666667

22.8671306355333

113.386402392348

26.1041977761767

441.233551552381

50.61600395

687.857379415498

29.4290470502137

658.428332365285

27.8087112160405

9.38269293226951

18.426018283771

9.26012836666667

241.292953161529

220.865568934559

20.4273842269697

19.8964787670833

19.8964787670833

16.3457484180392

16.3457484180392

22.1948959621947

22.1948959621947

38.0942548509087

20.8678748509087

17.22638

89.5296332770578

30.292464

28.6161647

28.6161647

20.165006462963

20.165006462963

18.1392269916667

18.1392269916667

3.5567379

19.8547322018978

73.3811960892381

23.3857033055785

25.2202945765625

24.7751982070971

371.294688625287

21.2958518416667

13.6608690916667

7.63498275

163.485692568647

52.9115973268024

28.6821646433608

24.2294326834416

106.751552591845

34.8937874043269

45.7728829675177

26.08488222

3.82254265

3.82254265

186.513144214974

43.0550098771281

43.3371033336582

36.3970154111111

18.5593881

17.8376273111111

50.6345486430762

13.08946695

979.011908005131

979.011908005131

52.086061025

78.4569941922882

28.8025829120261

16.9700846119054

5.0752346

27.6090920683566

29.9131460407797

35.5584828548505

61.6587453439381

44.0419147055681

17.61683063837

190.919154432494

15.375983535

26.3428648919737

15.4562160921429

17.36657232

51.7297909638984

37.0750022342105

14.6547887296879

13.5747381481481

61.0802664109918

22.0550334907197

16.0797335621088

22.9454993581633

40.2377076407777

23.8552399332653

14.019254075

12.848828625

113.969251336637

3.0918165

59.4093183910227

51.468116445614

124.562610142947

50.0067604708802

40.5815521500478

33.9742975220186

2004.06121019362

35.0811026889286

35.0811026889286

12.8178397090909

12.8178397090909

27.6241143306391

27.6241143306391

76.9986241306929

57.4493225363636

19.5493015943293

164.25906309

164.25906309

20.8400102361111

114.118028361538

114.118028361538

26.5045809313297

17.1769816813297

9.32759925

49.1124004941033

49.1124004941033

101.776539556991

66.8073494055556

22.7071535846591

12.2620365667763

404.06649828664

68.5642914148148

225.99420415

12.7688350857143

96.7391676361111

173.639677863601

38.7711097518488

81.5400319288461

53.328536182906

314.831235020928

47.1435362969968

245.388449629313

22.2992490946185

44.6904193294872

44.6904193294872

216.906839757754

17.529214256756

75.1144483266667

58.1908286531612

66.0723485211696

77.0523117767442

77.0523117767442

31.3873975375

18.9182726125

12.469124925

75.6230684196078

14.618447912963

17.288897425641

17.288897425641

4.82411333333333

4.82411333333333

1665.44603048595

866.441878495312

165.740029995312

700.7018485

73.1992433716312

24.2312287716312

48.9680146

13.4298452375

81.3545627682217

17.1489957484849

49.5567198

14.6488472197368

29.1840031611111

15.08815145

14.0958517111111

100.095495711667

100.095495711667

28.7356003894737

28.7356003894737

298.854781055921

229.4971061375

63.2635074684211

6.09416745

29.3547118826087

24.2779027826087

5.0768091

137.5551683125

7.2407401

7.2407401

796.990635218588

157.580143254167

248.916785661111

248.916785661111

194.260074425

54.6567112361111

4.3521015

13.1290779571429

54.0333455388889

14.1403529

11.5169660388889

28.3760266

8.0689081

10.9569616

46.2623254118108

33.7542846472744

12.5080407645363

232.358247238889

17.2575752565789

12.7141646065789

4.54341065

4.0751637

8.7092195

8.7092195

77.2020233133334

77.2020233133334

345.117999245971

14.8541539

4.0080005

10.8461534

118.020210715363

49.9233568718009

27.8783535380833

40.2185003054784

25.8489241166667

25.8489241166667

16.5515737487719

169.843136765169

169.843136765169

80.116742050464

46.2992106380948

33.8175314123692

39.6524875922912

17.3210507127083

12.0309874718897

20.7218689378161

51.4053617795264

10.144426462963

10.144426462963

41.2609353165635

626.546028333334

626.546028333334

626.546028333334

618.135773033334

8.4102553

344.356673216981

188.38392485128

188.38392485128

9.31318704621711

146.659561319484

74.7846173347222

14.0924451651786

18.996807805

38.7856910145833

474.713605082039

10.3824846

7.44315344566667

280.525004306086

82.214534932276

198.31046937381

105.018514728056

17.005434

17.005434

24.7692849490547

21.8471310031758

7.72259805

27.8603930981736

12.4196578135582

15.4407352846154

15.4407352846154

1976.34057738447

37.600039533687

37.600039533687

1018.06795753982

27.16354475

20.3633671568438

64.2260160391304

64.2260160391304

64.2260160391304

118.423887506822

20.1038260927152

14.6816978

14.6816978

20.341792207451

20.341792207451

16.2186834675936

16.2186834675936

27.5623170140625

11.726711125

15.8356058890625

19.515570925

18.566265

18.566265

738.01012980256

135.746184837256

7.3051757

58.2506778741296

18.3036503733333

48.6878591036264

3.930108

465.786473914215

31.3147472844618

25.0328208693009

13.7518976978723

13.7518976978723

13.7518976978723

11.2809231714286

11.2809231714286

11.2809231714286

895.639759441666

17.8597224416667

877.780036999999

877.780036999999

1055.21916359371

602.366065393764

66.5335356340418

22.457788775

27.8566054064103

16.2191414526316

11.91223795

11.91223795

39.419204575

39.419204575

236.2194517125

11.5213496

236.760285922222

59.1887179646086

44.2109151646086

23.3566458846591

20.8542692799495

14.9778028

26.4261033791667

367.238276856169

30.5201814236203

11.07985985

11.07985985

13.7252762

13.7252762

16.20319426

18.9600018

18.9600018

11.6226461416667

25.3757158741987

84.1504169916498

27.6233899743102

30.6050802554349

25.9219467619048

119.931088206111

16.5556026395833

43.1155194152778

31.7238660729167

28.5361000783333

35.6698961089224

18.36636155375

17.3035345551724

3273.28199559392

117.273477140617

34.6392778166667

522.6879127925

8.9439551

513.7439576925

3.5730996

3.5730996

3.5730996

11.74571494

11.74571494

11.74571494

44.3505454

4.6781609

4.6781609

4.6875571

57.331945

41.5514437

41.5514437

15.7805013

45.4362526

45.4362526

248.072269980296

55.6994865

30.2879142862963

106.385382694

82.9298967162222

23.4554859777778

55.6994865

151.761223393333

426.842513094444

266.955070345066

239.560855825

27.394214520066

6.1015195

13.042446

6.0083335

6.0083335

29.6561571576663

9.89239109583333

19.763766061833

21.0209294833333

8.80141075

1244.01051525

997.4025031

983.923337

13.4791661

246.60801215

246.60801215

246.60801215

4.60566385

4.60566385

178.81970603521

14.0201041

14.0201041

67.3380170507895

32.6247075909764

32.6247075909764

16.2643834131987

16.3603241777778

31.132536049

13.9044451166667

17.2280909323333

33.7043412444444

44.6467861302521

44.6467861302521

30.40047221

30.40047221

30.40047221

4.5105816

25.88989061

277.937745925644

29.836308320625

29.836308320625

27.0078707419674

56.6971953135131

164.396371549539

20.5307540951225

25.0903423806615

9.2291555

14.5555979700348

94.9905216037202

997.834701410808

56.6723033502246

29.6462527599079

27.0260505903167

21.60941594

21.60941594

35.6391232120445

35.6391232120445

9.85475081699346

9.85475081699346

160.193780828889

160.193780828889

282.021918274456

139.566572346142

54.7861443448976

41.1654848679456

46.5037167154708

57.8427339982496

12.0281313260274

14.9143692722222

30.9002334

37.7144146611577

11.2367423

25.8385517469556

6.69648703352941

195.597604939366

46.108539831206

29.1911925110468

83.6245618592004

36.6733107379128

23.1997102703704

23.1997102703704

23.1997102703704

73.7171640385714

73.7171640385714

242.5675436597

26.031380276875

12.01603585

14.015344426875

216.536163382825

61.4162443529101

155.119919029915

35.3824557163265

35.3824557163265

252.071055133421

67.049236993421

185.02181814

14938.9778418278

2013.66922036969

57.2681711

101.056875175374

65.3921632856515

49.0797697523182

49.0797697523182

6.543131

6.543131

9.76926253333333

9.76926253333333

35.6647118897222

35.6647118897222

30.6113166780027

16.6093329136667

27.8176480476389

128.295700167715

33.4836692645151

26.7204173700805

68.0916135331195

934.7461549

184.709314853928

39.4344771063158

39.4344771063158

47.2409885793103

42.6576551793103

42.6576551793103

4.5833334

4.5833334

30.2252739417355

22.9580772417355

7.2671967

20.9334646142857

20.9334646142857

20.9334646142857

46.8751106122807

46.8751106122807

174.796224637241

37.4790458728007

37.4790458728007

37.4790458728007

9.7780699

9.7780699

34.4907487051948

34.4907487051948

58.16898862144

58.16898862144

58.16898862144

34.8793715378055

24.8582264378055

10.0211451

318.012300784639

10.11332095

34.6637808253134

14.2027752089713

13.4848330282895

13.4848330282895

28.9894231871265

7.7734257

75.7735519145238

36.13444782365

35.1935743046588

26.3750467333333

35.3081211087719

39.746181111484

39.746181111484

1172.26526252878

116.918796636842

1033.87561734749

961.816913780317

36.1038220803181

925.713091699999

32.4732740944444

32.4732740944444

39.5854294727273

39.5854294727273

21.4708485444444

644.503679027968

413.380758002085

21.106581697371

41.9785028682437

94.9403923511243

56.0879903884594

38.8524019626649

132.047162332418

37.6333182226191

11.071857847619

14.1984686

12.362991775

85.6748005303087

44.4584501521619

41.2163503781468

67.4248698838804

50.0804226631579

50.0804226631579

17.3444472207225

12.2134735666667

151.484577575336

21.2167907131579

90.9125722073045

67.4209882489711

23.4915839583333

20.8607517154407

18.4944629394328

1699.51959221761

40.3796889108696

40.3796889108696

40.3796889108696

25.05357103

25.05357103

17.50199058125

4.1412425

13.36074808125

1466.4061634691

72.6959872510904

72.6959872510904

11.5940571333333

11.5940571333333

11.5940571333333

1382.11611908467

96.9911295745833

21.31503015

75.6760994245833

22.575898217929

22.575898217929

314.652398715023

29.9927735436936

284.659625171329

47.4957512333333

47.4957512333333

22.4034677246032

22.4034677246032

214.760406213393

171.064132778581

13.9234275222083

45.119840078125

37.6769314440625

27.7049729390625

46.6389607951231

539.340516631429

40.998528776261

34.7572461209877

115.361249656593

14.8995954833333

31.8647908207408

48.6587318247856

53.9780708420926

137.436905028241

32.6124861209877

28.7729119574074

237.492043167125

75.1768989095238

18.9168098078125

18.7603438097561

31.9141665357143

66.0334785546673

17.7062115329849

8.98413401666667

150.178178226393

123.255230900507

40.3156040556137

45.2265095261438

37.71311731875

26.9229473258852

40.2842688693451

40.2842688693451

2339.74427044385

2339.74427044385

121.982708019631

59.2652974332479

62.7174105863828

91.1289427916667

91.1289427916667

67.8197818738824

67.8197818738824

279.38007308913

40.3120708267007

52.6588739617943

105.177845944204

37.0315002557325

44.1997821006982

152.726973162787

27.524079609375

30.65437744275

30.0632226642857

50.1503039219907

14.334989524386

58.7875362631172

58.7875362631172

63.486647064

63.486647064

87.5487232613435

36.5008229617031

51.0479002996404

42.9468265210233

42.9468265210233

105.390913133217

62.2903258409091

43.1005872923077

40.8486794777778

16.0275938

24.8210856777778

53.7007921559379

28.7574777089474

24.9433144469906

19.4571117384615

19.4571117384615

475.177146237349

447.8363053375

27.3408408998497

42.7787118097826

16.2605583347826

19.846359425

6.67179405

237.7478843125

237.7478843125

65.8319758880221

65.8319758880221

53.556190226402

53.556190226402

279.446653417815

20.439525480519

36.4802389332055

222.526889004091

3794.03063941603

12.2169276

27.0694858

43.9241436289474

325.059540705416

236.668374566667

41.8723795106383

19.1007938380823

6.4955758

4.8208618

16.1015551900285

376.501650226818

202.905973194843

15.0026153

32.7086675333333

19.5787693173913

78.69753413125

27.60809075

28.4212066961905

28.4212066961905

16.7913661

185.350364550699

2.6637427

67.303370222541

31.913218875

35.390151347541

75.0905185925

75.0905185925

732.439898259685

74.3458374043858

74.3458374043858

68.8433265016667

26.3470503533333

42.4962761483333

491.150061127919

166.998326219048

118.465259813161

186.378126624281

19.3083484714286

33.5162678857143

33.5162678857143

64.58440534

64.58440534

317.253463290309

93.0255727776396

26.6646988513745

60.7750159262652

5.585858

178.769318200904

45.4585723117647

36.1345844729318

22.3690619642857

13.7655225086461

1245.30215751337

200.121782838199

176.941634600665

7.87282706666667

15.3073211708678

11.3559313934211

11.3559313934211

112.445624101542

36.0715327708333

21.9748403875

14.0966923833333

15.5954281212683

31.3678439744734

15.5954281212683

15.7724158532051

29.4108192349673

156.921329144153

69.9106634465033

87.0106656976496

182.483192998731

32.3357848807571

19.4087373

14.0615543427635

23.6769060213542

79.4858410757936

35.9806256404762

14.977408900641

28.5278065346764

13.5143693780627

210.864591272991

30.0913892021578

17.7162374541667

163.056964616667

262.622798875028

18.4590626357143

79.6709805410847

52.2476766159515

52.2476766159515

23.2401419058246

89.0049371764524

81.1012100521648

18.0332675868182

44.5281285658011

24.7177556908784

19.8103728749228

18.5398138995455

27.3856968371429

27.3856968371429

30.8700080384615

30.8700080384615

22.0137557365714

22.0137557365714

34.6693265376278

22.115152610776

12.5541739268518

56.2886652545454

49.35388006

21.3362689773504

14.8953000153846

3.5426361

69.5383783366749

13.3573418420259

13.3573418420259

43.1157661872961

13.7997896170259

13.7997896170259

29.3159765702703

13.0652703073529

213.577516499796

42.5297626062808

42.5297626062808

88.9108163247312

45.9518152666667

42.9590010580645

41.6851871394737

40.4517504293103

331.696608133993

89.7232116293851

41.7080831803819

14.7450310090909

33.2700974399123

53.0836507883655

53.0836507883655

23.486588875

82.7633474545096

4.55607292

27.3051139098684

50.9021606246412

26.0376443928819

36.251416508851

20.350748485

5.63124581

14.719502675

624.69162593709

33.8619653459412

17.3916704352193

22.2359775301242

22.2359775301242

23.10818875

23.10818875

57.3189738738889

149.276511408049

27.9777671054348

27.9777671054348

27.6238894885939

38.1005447531915

31.8339788285507

23.7403312322778

23.7403312322778

88.21142455

233.286914043868

71.9540748291375

21.3760034241379

21.3760034241379

5.7203403

5.7203403

52.9730321465625

52.9730321465625

81.2634633440299

44.4170903740299

36.84637297

25.6721332208468

25.6721332208468

1389.50270368148

384.594344145577

271.112984449951

48.7051080994375

50.4228212133333

30.6903506579772

87.4112523956364

53.8834520835664

113.481359695626

113.481359695626

50.040406419462

63.4409532761643

174.485662353462

39.9019718806452

53.5622205293904

45.6708835612903

15.5720570513636

19.7785293307721

428.847923643561

296.513080122228

104.398812776509

88.6739618127692

103.440305532949

132.334843521333

132.334843521333

132.334843521333

401.57477353888

144.696854731498

144.696854731498

45.1997072839805

55.7401397740934

55.7401397740934

43.7570076734238

161.309777150896

24.8500232245625

50.11443631875

50.11443631875

86.3453176075833

27.4030152645833

25.159725223

33.78257712

95.568141656487

45.0183438294788

50.5497978270082

649.820321481279

463.8409838

36.5885924061086

16.7013883666667

43.9040106702703

43.9040106702703

51.388237305733

12.1546160325

12.1546160325

25.2424929
