## Supplementary material for "Gut microbial ecology of Xenopus tadpoles across life stages": Figure_S15

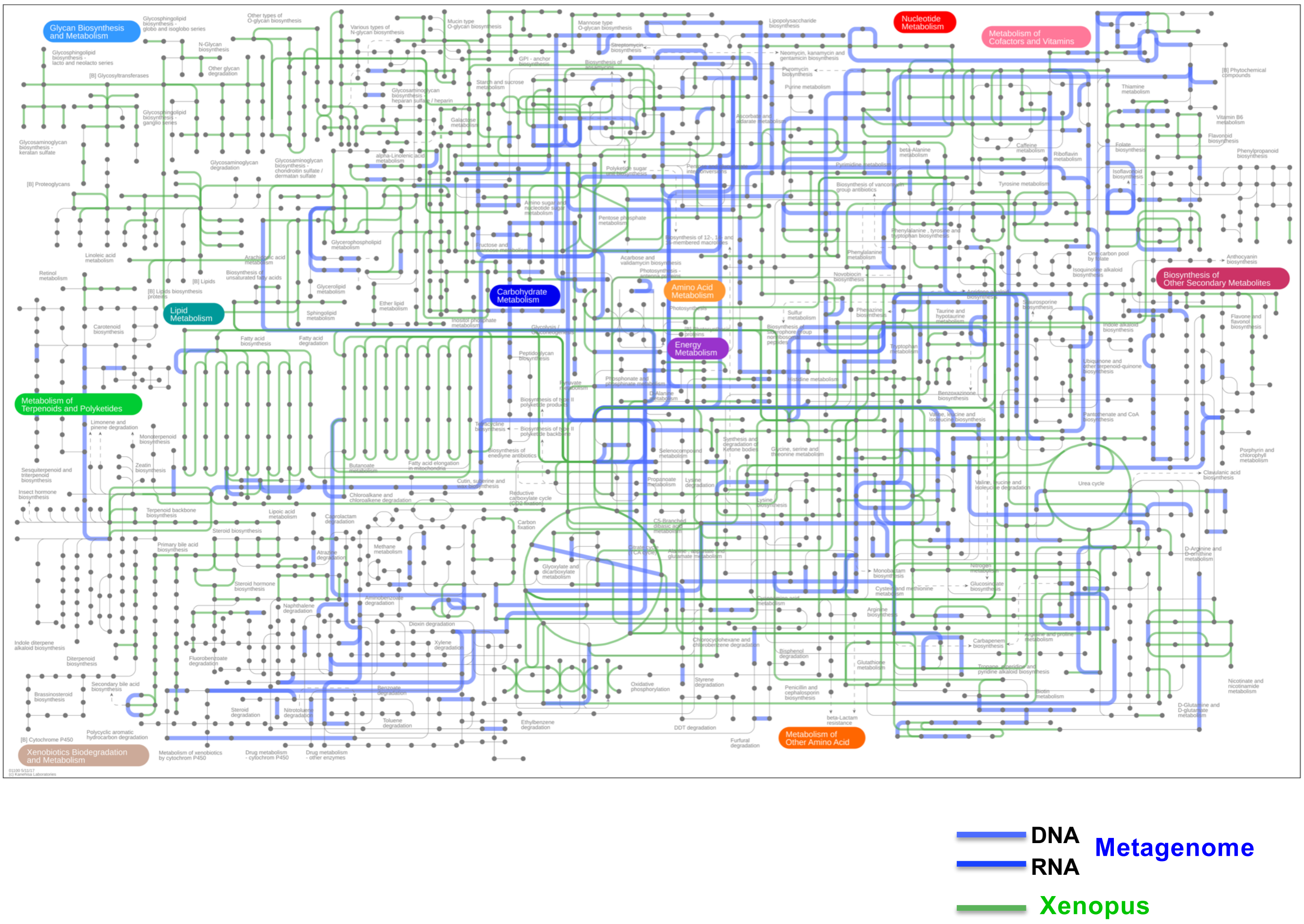


**Figure S15: Metabolic map of *X. tropicalis* genome and its gut metagenome.**
This metabolic pathway highlights in green the metabolic pathways predicted from the *Xenopus* genome and in blue those predicted from the tadpole gut metagenome. This map can be interactively accessed at https://pathways.embl.de/selection/pWbci4bo871W8Qm8XKF
