## Supplementary material for "Gut microbial ecology of Xenopus tadpoles across life stages": Figure_S16

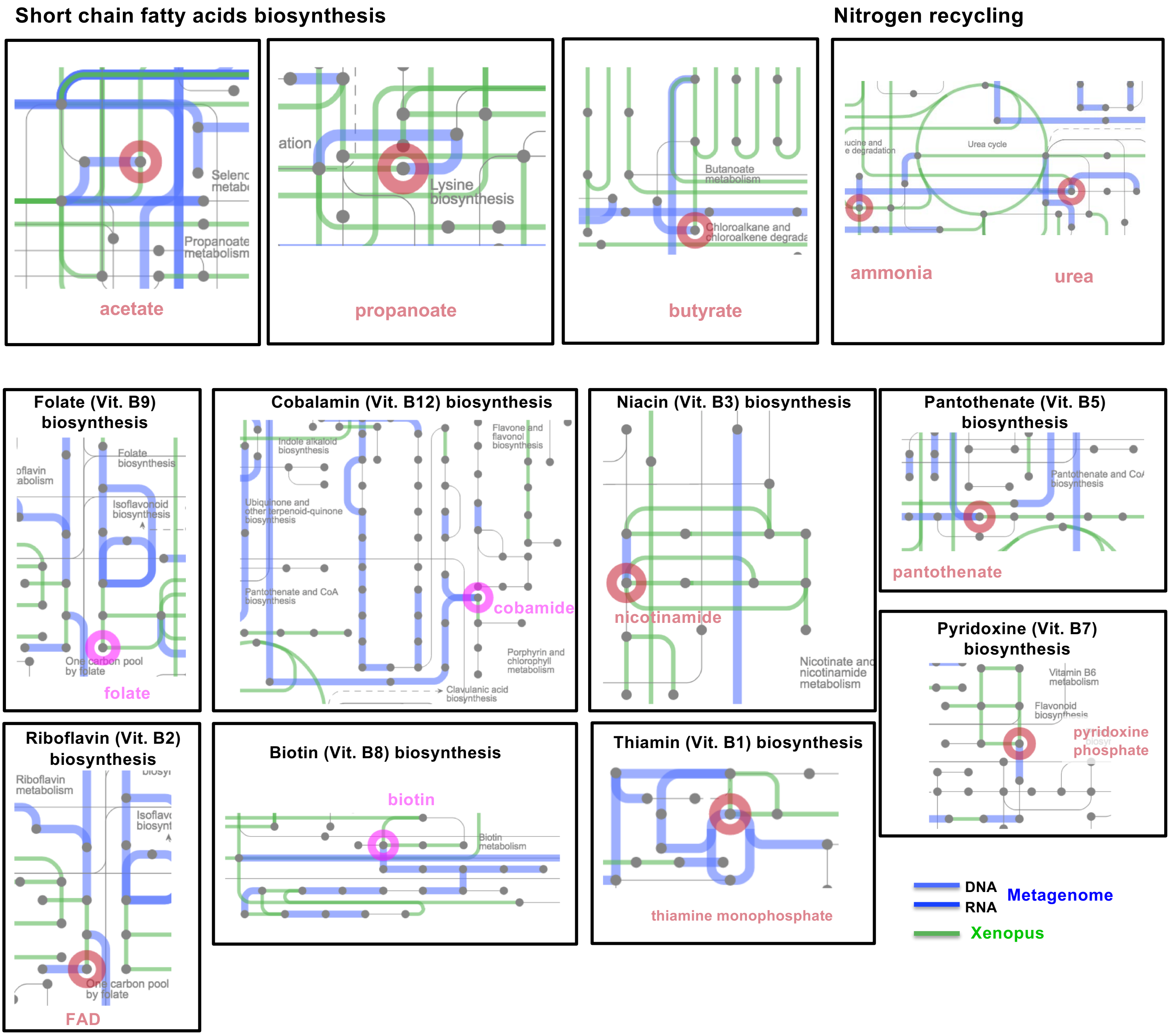


**Figure S16: Selected cases of metabolic potential of *X. tropicalis* genome and its gut metagenome.**
Fragments of biosynthetic pathways for common short-chain fatty acids, nitrogen recycling and B-vitamins is depicted. These metabolic pathways highlight in green the pathways predicted from the *Xenopus* genome and in blue those predicted from the tadpole gut metagenome. This map can be interactively accessed at https://pathways.embl.de/selection/pWbci4bo871W8Qm8XKF
